# *The Binary Cellular Biology of Human Growth*: I. Quantifying the Creation and Growth of the Body and its Parts

**DOI:** 10.1101/2025.01.22.634248

**Authors:** Philip Chodrow, Jessica Su, Daniel Lee, Neil He, Ruben De Man, Ankur Tiwari, William Mannherz, Luca Citi, Rashi Gupta, Zifan Gu, David E Cantonwine, Thomas F McElrath, Henning Tiemeier, James Selib Michaelson

**Affiliations:** Department of Computer Science, Middlebury College, Middlebury, VT 05753; Ultrasound Systems, Philips; Stanford University, 450 Jane Stanford Way, Stanford, CA, USA; Department of Mathematics, Yale University, New Haven, CT; Yale University School of Medicine, New Haven, CT, USA; Department of Surgery, University of Texas Health Science Center San Antonio; Harvard/MIT MD-PhD Program, Boston MA; School of Computer Science and Electronic Engineering (CSEE), University of Essex, Wivenhoe Park, Colchester CO4 3SQ; Laboratory for Quantitative Medicine, Massachusetts General Hospital, Boston MA; University of Texas Southwestern Medical Center, Dallas, Texas; Division of Maternal-Fetal Medicine, Department of Obstetrics Gynecology, Brigham and Women’s Hospital, Boston MA; Department of Social and Behavioral Sciences, Harvard TH Chan School of Public Health, Boston, MA; Laboratory for Quantitative Medicine, and Departments of Pathology and Surgery, Massachusetts General Hospital, Department of Pathology, Harvard Medical School, Boston MA USA; Department of Biostatistics, Boston University; Massachusetts Institute of Technology, 77 Massachusetts Avenue, Cambridge, MA, USA

## Abstract

Assessing and understanding the ***Growth*** and ***Development*** of animals generally, and of human fetuses specifically, provides essential information for the management of the health of the newborn and its mother. Here we show that a consideration of the formation of the body, in units of numbers of cells, ***N***, and the change in these numbers by the binary either/or decisions of cells to divide or not, an approach we call ***Binary Cellular Analysis***, provides new insights and new equations for quantifying the growth and development of the body and its parts. These equations include: 1) The ***Binary Cellular Universal Growth Equation***, which captures growth from conception to adulthood, and provides a method for biologically-based, data-driven, ***Growth Curves***; 2) The ***Binary Cellular Universal Mitotic Fraction Equation***, which lies within the ***Binary Cellular Universal Growth Equation***, and captures the mechanism of growth to adult size, control of cell division; 3) The ***Binary Cellular Allometric Growth Equation***, which captures the creation and growth of the ***Tissues, Organs***, and ***Anatomical Structures*** of the body, and how the embryo creates each ***Body Part*** from a ***Single Founder Cell***; and, 4) ***Binary Cellular Estimated Fetal Weight Equations***, derived from the ***Binary Cellular Allometric Growth Equation***, which captures the relationship between ***Ultrasound Measurement*** and the ***Size*** of the body as a whole. These equations capture the developmental process of ***Cellular Selection***, resulting from ***Differential Cellular Proliferation***, which molds the formation of the body from a single fertilized ovum into a multicellular animal. The parameters of these equations both capture the differences in ***Size*** and ***Growth*** between animals across the taxonomic spectrum, and between human fetuses, as well as identify the individual mechanisms of mitosis that drive ***Growth*** and ***Development***. From the ***Binary Cellular Universal Growth Equation***, the ***Growth*** of human fetuses can be measured, thus providing a much-needed tool for understanding the biological forces that cause a fetus to grow to ***Small, Average***, or ***Large Size*** at birth, for providing the basis of biologically-based, data-driven, ***Growth Curves***, and for assisting the management of the obstetric care of the fetus and its mother.

## INTRODUCTION

### The Biology and Medicine of *Growth*

***Growth*** is the essential feature of animal life. Driven by cell division, animal life begins with all cells dividing, ***Exponential Growth***, then almost all cells dividing, then fewer cells dividing, guiding the ***Growth*** of the animal body to its recognizable ***Size***. It is this fundamental embryological process of the macroscopic increase in ***Size***, driven by the microscopic increase in ***Cell Number*** (***N***), that lies behind the successful and unsuccessful growth of humans before and after birth.

### The Biology and Medicine of *Cellular Selection*

***Cellular Selection***, resulting from ***Differential Cellular Proliferation***, or ***Differential Cell Death***, molds the formation of the body from a single cell, the fertilized ovum, into a multicellular animal.^1-3^, If the magnitude of ***Cellular Selection*** is ideal, animals will grow to ideal size. If there is ***Too Much***, or ***Too Little Cellular Selection***, animals, and especially human fetuses, will become ***Too Big***, or ***Too Small***. Thus, as we shall see here, ***Cellular Selection*** gives us a measure which links the macroscopic quantity of ***Fetal Size*** to the microscopic events of ***Mitosis*** ^9-11^ ***and Apoptosis***,^4^ and with which we can understand, and manage, the obstetric care of the fetus and its mother.

### The Evolutionary Origin of *Growth* and *Cellular Selection*

More than half a billion year ago, our single-cell ancestor gave rise to the first multi-cellular animal by learning how to relinquish uncontrolled ***Exponential Growth*** in exchange for suppression of cell division.^5,6^ It was this transformation of the ***Natural Selection*** of the ***Competing Cells*** of ***Single-Cell Species***, into the ***Cellular Selection*** of the ***Cooperating Cells*** of ***Multicellular Species***, that makes ***Growth*** to specific ***Sizes*** possible.^1-3^

As we shall see here, and in the next two papers in this series,^7,8^ gaining insight into this biologically deep, and ancient, process of ***Cellular Selection***, across the animal spectrum, and comprehending this information in a mathematical form that reliably captures the actual ***Growth*** of real animals, and real humans, can provide the obstetrician with the practical information needed to bring the child and its mother into the world with the best possible health.

### The Nature of *Growth*, and the Current Math for Capturing *Growth*

As we grow older, we grow bigger, and we grow slower, by progressively more and more of our cells becoming ***Quiescent***^,9-11^ ***Biologists have long searched for an equation that fits, and explains, such animal Growth***, and its role in human health.^12-31^ Unfortunately, none of these equations have been found to do a good job at capturing the actual growth of real animals, including humans.^32-34^

The absence of an equation that can accurately and precisely gauge the growth of humans, in fetal life and beyond, has posed a formidable challenge to the construction of the tools needed for guiding obstetric care.^35-38^ For example, growth charts^39-51^ have frequently had to rely on polynomial equations as curve fitting devices, unlinked to the biology of cell division. This has made the creation of a ***Universal Reference*** for ***Fetal Growth Charts***, which could capture the growth of the variety of fetuses, and the variety of populations in which they are born, only a hoped-for possibility.^52^

### The Nature of *Body Part Creation* and *Growth*, and the Current Math for *Body Part Creation* and *Growth*

Early in life, embryos create ***Body Parts***. Biologists have long searched for a mathematical framework that fits, and explains, the ***Creation*** and ***Growth*** of these ***Tissues, Organs***, and ***Anatomical Structures***^.104-106^ ***Precisely how our Bodies*** make our ***Body Parts*** from ***Founder Cells***, and how they guide the resulting ***Body Parts*** to their mature ***Sizes***, has long been obscure. Capturing the information of the relationship between the ***Size*** of the ***Whole Body***, and the ***Size*** of the ***Parts*** of the ***Body***, is of particular importance in obstetrical practice, where the size of the fetus is estimated from ***Ultrasound Measurement*** of these ***Body Parts***.

### *Binary Cellular Analysis* Provides a New Way to Capture *Size, Growth*, and *Creation*, of the *Body* and its *Parts*

To solve the problems outlined in the previous sections, we have examined data on the ***Size*** and ***Growth*** of developing animals and humans in units of numbers of cells, ***N***.^3,53^,54 We call this method ***Binary Cellular Analysis***, as it examines the ***Binary*** choices made by our cells to divide, or not, and how these choices determine the number of cells in our ***Bodies*** and our ***Body Parts***, that is, our ***Tissues, Organs***, and ***Anatomical Structures***, and from this, their ***Size*** and ***Growth***.

As we shall see in this communication, and the two papers that accompany this communication^,7,8^ such a ***Binary Cellular*** approach allows us to derive biologically-based, accurate, and precise methods and equations for capturing the ***Size, Growth***, and ***Creation***, of our ***Bodies*** and our ***Parts***. The equations include:

1. The ***Binary Cellular Universal Growth Equation***, which captures growth from conception to adulthood. The ***Binary Cellular Universal Growth Equation*** provides a method for creating biologically-based, datadriven, ***Growth Curves***.^38^***-***52
2. ***The Binary Cellular Universal Mitotic Fraction Equation***, which lies within the ***Binary Cellular Universal Growth Equation***. The ***Binary Cellular Universal Mitotic Fraction Equation*** captures the mechanism of growth to adult size, control of cell division, that is, ***Cellular Selection***^.1-3^
3. ***The Binary Cellular Allometric Growth Equation***, which captures the creation and growth of the ***Tissues, Organs***, and ***Anatomical Structures*** of the body. The ***Binary Cellular Allometric Growth Equation***, captures how embryos create these ***Body Parts*** by ***Cellular Selection*** of ***Single Founder Cells*** and regulates the size of these ***Body Parts*** by ***Cellular Selection*** of the ***Founder Cells***’ progeny^.1-3^
4. ***Binary Cellular Estimated Fetal Weight Equations***, derived from the ***Binary Cellular Allometric Growth Equation***, which captures the relationship between ***Ultrasound Measurements*** and the ***Size*** of the body as a whole.

As we shall see in this communication, and the two papers that accompany this communication,^7,8^ these ***Binary Cellular Equations*** make it possible to make more accurate, precise, and actionable estimates of ***Fetal Size*** and ***Growth*** from ***Ultrasound Measurements***. As a result, these ***Binary Cellular Equations*** can help the obstetrician guide each pregnancy to a ***Birthweight*** and a ***Pregnancy Length***^55^ with the highest chances of survival and health.

In the second, following paper in this series,^8^ we shall see that ***Binary Cellular Estimated Fetal Weight Equations*** for estimating human ***Fetal Weight*** provide improved assessments of ***Fetal Weight***, in comparison to other currently used ***Estimated Fetal Weight Equations***. We shall also see how ***Binary Cellular Mathematics*** can be put to work to improve the assessment of the ***Size*** and ***Growth*** of the human fetus from ***Ultrasound Measurement***.

In the third, final paper in this series,^8^ we shall see that ***Binary Cellular Equations*** for estimating ***Fetal Weight*** from ***Ultrasound Measurements***, and for capturing ***Fetal Growth*** from conception onward, allows for the practical measurement and prediction of human fetuses that become ***Large, Average***, and ***Small*** at birth. This ***Binary Cellular Analysis*** approach provides a way to identify the biological mechanisms that drive human fetuses to being born ***Too Big*** or ***Too Small***, as well as for identifying new possibilities for the detection and pharmacological treatment of fetuses that are growing ***Too Slowly*** and ***Too Rapidly***^.56,57^,***58 Binary Cellular Analysis*** also provides new ways to help obstetricians guide each fetus to a ***Birthweight*** and ***Fetal Age*** that has the highest possible chance of a favorable outcome. Finally, we shall see how the ***Binary Cellular Universal Growth Equation***, which captures growth from conception to adulthood, provides a method for creating biologically-based, data-driven, ***Growth Curves***.^52^

## METHODS

To examine data on the growth of animal bodies, we assembled data on the number of cells in the whole animal, ***N***_***w***_, by age, ***t***, in days from fertilization until maturity, for 13 species of animals: zebrafish (***Danio rerio***),^59^ European sea bass (***Dicentrarchus labrax***),^60,61^ mice (***Mus musculus***),^62,63^ rats (***Rattus norvegicus***),^64^ cows (***Bos taurus***),^65,66^ bobwhite quail (***Colinus virgianus***),^67,68^ domestic chickens (***Gallus gallus domesticus***),^69,70^ turkeys (***Meleagris gallopavo***),^71^ geese (***Anser anser***),^67^ nematode worms (***C. elegans***,^89^ ***M incognita***^90^), frogs (***Rana pipiens***),^72^ clams (***Merceneria mercenaria***).^73,74^,75 and humans (***Homo sapiens***),^76,77^,78 Data on the numbers of cells in early embryos, from fertilization onward, were available for mice,^79,80^ rats,^81^ cows,^82^ chickens,^83^ turkeys,^84^ nematode worms,^89,90^ clams,^74^ fish.^85,86^ and humans.^87^ Data on the numbers of cells in later embryos were assembled from published values in units of weight or volume by taking advantage of the finding that there are about 10^8^ cells/gram-cc.^88^ Excel files (SUPPLEMENT-TABLE A2) characterizing these basic growth data (“**Basic Data Files**”), and the calculations made from these data (“**Calculations Files**”), are available on request. Also available to interested readers are excel files with the data on the growth of the parts of the body that we examined (see below).

To examine data on the growth of the parts of animal bodies, we assembled values of ***N***_***p***_ vs. ***N***_***w***_ (the number of cells in a body part, ***N***_***p***_, and in the body as a whole, ***N***_***w***_, at various points in time) for each of the ***Body Parts*** that have been characterized in the ***Cell Lineage Charts*** of ***Caenorhabditis elegans***^89^and ***Meloidogyne incognita***^90^, nematode worms, as well as for ***Oikopleura dioica***, chordate tunicates.^91^ We also assembled values of ***N***_***p***_ vs. ***N***_***w***_ for the ***Body Parts*** evident in embryos and juveniles of larger animals, again relying on the observation that there are about 10^8^ cells in every gram of tissue.^92,88^ We assembled these values for the ***Body Parts*** of chicks (gizzards, livers, hearts, kidneys),^93^ rats (livers, brains, kidneys, forelegs, ears, stomachs, spinal cords),^94^ zebrafish (eye lens),^95-98^ mice (livers, brains, kidneys, forelegs),^94,99^ clams, goldfish, and humans (brains, livers, kidneys, lungs, pancreases, adrenals, thymuses, spleens, lower extremities, upper extremities, stomach, heart, intestines).^100^

***Gestational Age***, the time since the last menstrual period, has long been the most common way for thinking about the age of the human fetus. Another commonly used term is ***Conceptional Age***, but the date of conception is not always known. As we shall be examining fetal, childhood and adult growth data, as well as growth data of animals such as mollusks, tunicates, and nematodes, which don’t have a fetal period, we shall be using the simplest term, ***Age***, with modifiers to indicate the period examined, which, for our work here, will be ***Fetal Age***.

To calculate human ***Fetal Age***, we have adopted and adapted the INTERGROWTH Equation for estimating ***Fetal Age*** from ***Crown to Rump Length*** (***CRL***)^101^:

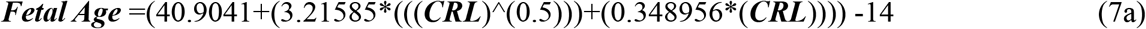

We call this expression, the ***CRL Fetal Age Equation*** (#7a). The 14 days subtracted from the INTERGROWTH value converts the values from the ***Gestational Age*** to ***Fetal Age***.

Data were analyzed in Microsoft Excel.

Equations used here and in the related manuscripts are numbered (1), (2), etc. When equations are listed, but in different form from their parent equations, or specify the details of a parameter in a parent equation, they are given the same number as the parent equation, followed by a letter, that is (2b), (2c), (2d), etc. Equations only used in the SUPPLEMENT, and not in this manuscript, are listed in numbers starting from (100).

## RESULTS

### *Binary Cellular Analysis*: Capturing *Size, Growth*, and *Creation*, of the *Body* and its *Parts* in *Cell Numbers, N*

Here, we report our studies of examining data on the ***Size, Growth***, and ***Creation***, of the ***Body*** and its ***Parts*** (***Tissues, Organs***, and ***Anatomical Structures***) in units of numbers of cells, ***N***. We call this method ***Binary Cellular Analysis***, as it examines the ***Binary*** choices made by our cells to divide, or not, and how these choices determine the number of cells in our ***Bodies*** and our ***Body Parts***, and from this, their ***Size*** and ***Growth***.

### *Binary Cellular Analysis* of the *Growth* of the Body

As we grow older, we grow bigger, and we grow slower. Two centuries ago, Verhulst provided the first expression for such ***Density Dependent Growth***, the ***Logistic Equation***, which envisages growth as occurring exponentially, with a linear decline in the rate of growth, in continuous units of weight, volume, and length, ***w***.^13^ Many ***Density Dependent Growth Equations*** have been considered since Verhulst - the ***Gompertz***,^14^ the ***Von Bertanalfy***,^15^ the ***West***,^17^ to name just a few - none of which has been found to accurately capture the growth of real animals, including humans (Equations #100-104 in SUPPLEMENT)^.32-34^

### Data for *Binary Cellular Analysis* of the *Growth* of the Body

To solve this problem, we turned to actual growth data, to let real information on the growth of real animals tell us what the best ***Density Dependent Growth Equation*** might be, indeed, whether there even is such a single equation that matches the growth of all animals.^54^ To carry out the ***Binary Cellular Analysis*** of growth, we assembled data on the number of cells in the whole animal, ***N***_***w***_, by age, ***t***, in days from fertilization (***t*=0**) until maturity, for 13 species of animals: zebrafish (***Danio rerio***), European sea bass (***Dicentrarchus labrax***), mice (***Mus musculus***), rats (***Rattus norvegicus***), cows (***Bos taurus***), bobwhite quail (***Colinus virgianus***), domestic chickens (***Gallus gallus domesticus***), turkeys (***Meleagris gallopavo***), geese (***Anser anser***), nematode worms (***C. elegans, M incognita***), frogs (***Rana pipiens***), clams (***Merceneria mercenaria***), and humans (***Homo sapiens***) (see METHODS section). Graphs of these various datapoints for animal size’s (***N***_***w***_) versus time (***t***), are shown in **Figure 1** and **Supplement-Figures-I.1** and **4**, and the methods for assembling these data can be found in the SUPPLEMENT.

**Figure 1.**
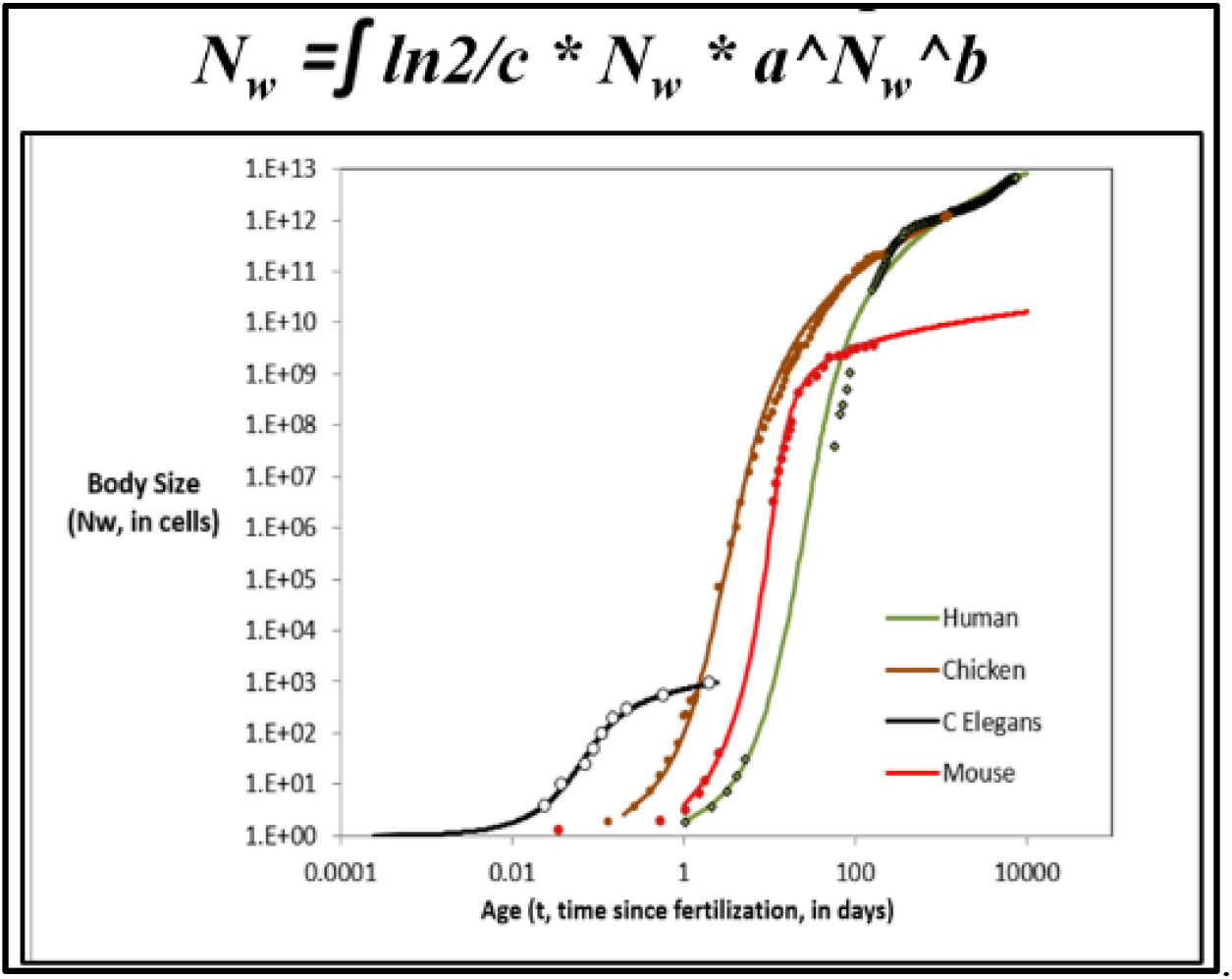
The ***Binary Cellular Universal Growth Equation***. Actual Size/Age datapoints for humans, chickens, nematodes, and mice, in units of numbers of cells, ***N***_***w***_, shown in dots; the ***Binary Cellular Universal Growth Equation***, with the appropriate ***a, b***, and ***c*** parameters for each of these species (SUPPLEMENT-TABLE I), shown in curves.

### The S-shaped Growth Curve

Graphing the number of cells in the body as a whole, ***N***_***w***_, vs. age, ***t***, from fertilization until maturity, reveals, for each of these 13 species, that animal growth occurs by ***S-shaped*** growth curves (**Figure 1** and **Supplement-Figures-I.1** and **4**). This is most easily viewed on log-log plots, because the greatest part of growth occurs rapidly at the beginning of life. For example, in terms of numbers of cells, ***N***_***w***_, humans grow about a hundred-billion-fold in the womb and about a hundred-fold after birth. As we shall see below, the reason why all of these creatures have similar ***S-shaped*** growth curves is that they all grow by an equation of the same form, the ***Binary Cellular Universal Growth Equation***.

### The Mitotic Fraction

Having assembled this growth data in units of numbers of cells, ***N***, we are now in a position to calculate how cell division created these numbers of cells. We have done so by calculating what fraction of cells must have been dividing to account for the growth in each of these animals, a number we call the ***Mitotic Fraction, m***.

Details for how the ***Mitotic Fraction, m***, is calculated can be found in the SUPPLEMENT, but one can easily visualize the process by imagining the case of a child’s growth from fertilization into maturity. Many IVF studies have found that human embryos starting from a single fertilized ovum on day1, and typically become ∼2cells on day2, ∼4cells on day3, ∼8cells on day4, and so on. These numbers suggest that the ***Cell Cycle Time, c***, of a human is about **1** day, and that right after conception, the ***Mitotic Fraction, m*≈1**. Thus, at the beginning of life, almost all cells are dividing, doubling in number, ***N***, once a day. Nine months later, we have an 8-pound baby girl, and one thing we know for sure is that the next day, she doesn’t weigh 16 pounds! If our 8-pound newborn baby girl became 8.8 pounds on the day after birth, we could calculate the ***Mitotic Fraction*** as being 0.1 (***m***≈0.1), since 10% of the child’s cells would have to divide for her to grow from 8.0 pounds to 8.8 pounds. If our newborn baby girl weighed 8.08 pounds the day after birth, ***m*≈**0.01. If our newborn baby girl weighed 8.008 pounds the day after birth, ***m*≈**0.001, and so on.

### As Animals Grow in Size, *N*_*w*_, the *Mitotic Fraction, m*, Declines Rapidly

We carried out calculations by the ***Mitotic Fraction Method*** of the ***Mitotic Fraction***, from fertilization until maturity, for a great range of animals, including nematodes, frogs, chickens, cows, geese, quail, turkeys, mice, rats, fish, mollusks, and humans (**Figure 2** and **Supplement-Figures-I.2** and **3)**. These graphs reveal, for all of these animals, that growth is close to exponential during the first few cell divisions (***m*≈1**), but soon begins its “slippery slope” descent, resulting in a rapid and accelerating decline in the value of ***m***, reaching values of **∼10**^**-2**^ to **∼10**^**-6**^ by the time adult size is reached.

**Figure 2.**
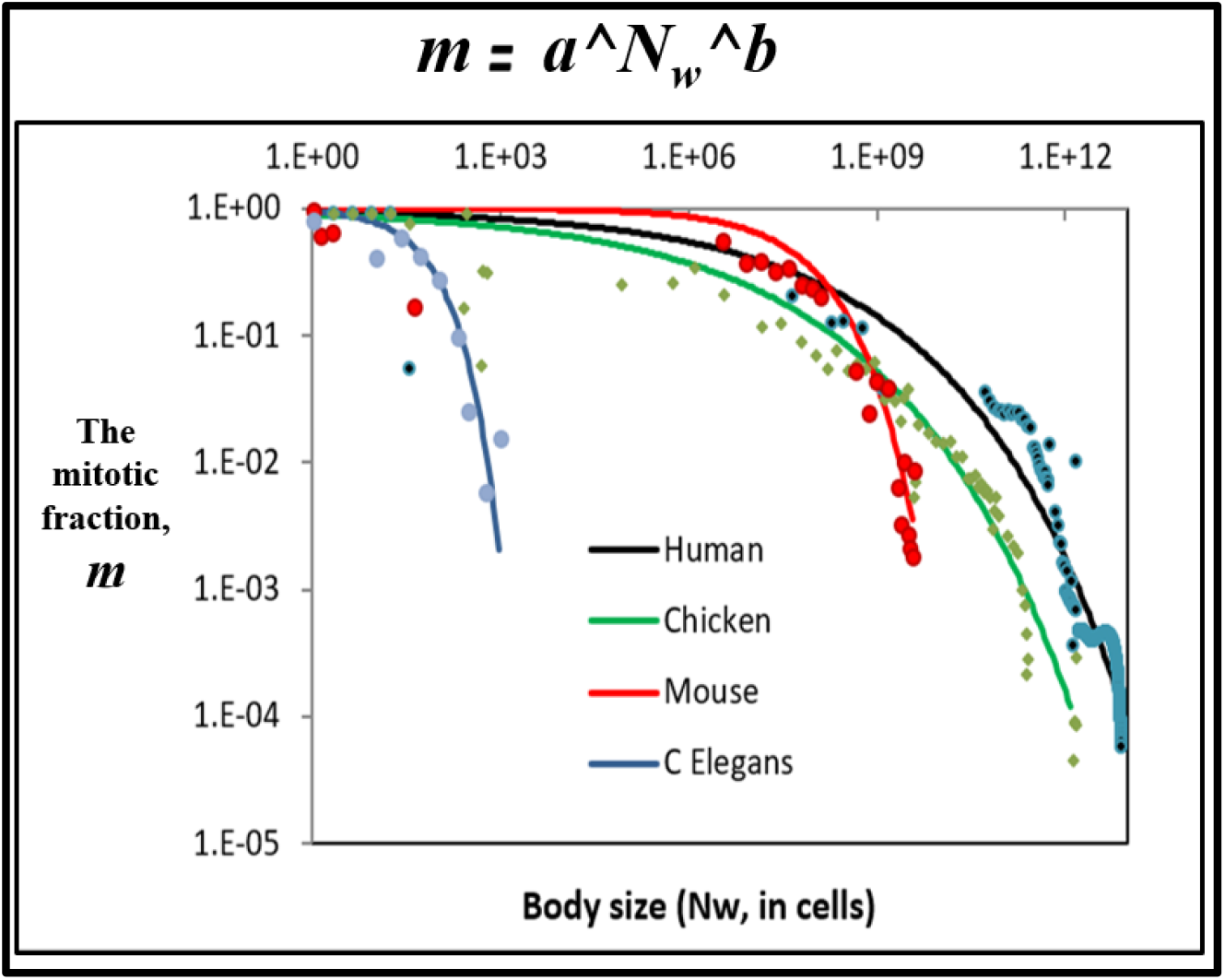
The ***Binary Cellular Universal Mitotic Fraction Equation***. Values of the ***Mitotic Fraction, m***, calculated by the ***Mitotic Fraction Method***, as a function of animal size, ***N***_***w***_. Data points for animal size, in integer units of numbers of cells, ***N***_***w***_, vs. the ***Mitotic Fraction, m***, from fertilization, until maturity, for humans, chickens, ***C elegans*** nematodes, and mice.

### The *Binary Cellular Universal Mitotic Fraction Equation* Captures the Decline in the *Mitotic Fraction*

The basis for the relationship between the ***Mitotic Fraction, m***, and ***N***_***w***_ becomes evident by graphing the **log(*N***_***w***_) vs **log(−log(*m*))**, revealing roughly straight rows of dots (**Supplement-Figure-I.2**). This means (see the SUPPLEMENT for the math) that the relationship between the ***Mitotic Fraction, m***, and ***N***_***w***_ can be captured with the expression:

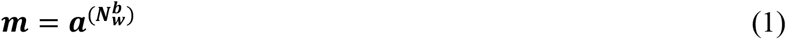

We call this expression the ***Binary Cellular Universal Mitotic Fraction Equation*** (#1), as we have found that it closely captures the value of the ***Mitotic Fraction, m***, the fraction of cells that must be dividing to account for the growth of the body, from fertilization until maturity, for each of the 13 species noted above, including humans (**SUPPLEMENT-TABLE A1**). The parameters ***a*** and ***b*** reflect the specific growth characteristics of each animal.

### The Binary Cellular Universal Mitotic Fraction Equation Links Growth to Endocrinology

What might be the mechanism behind the decline in the ***Mitotic Fraction, m***, by the ***Binary Cellular Universal Mitotic Fraction Equation*** (#1)? A helpful insight emerged from the mathematical examination of the discrete allocation of hormones and growth factors among the cells of an embryo growing in a constant volume, such as a bird’s egg or mammal’s uterus.^3^

Consider the case of the embryo’s cells producing an inhibitory growth factor and carrying a receptor for those inhibitory molecules. Early in development, when our idealized embryo is just a few cells, those few cells would produce only a small number of inhibitory molecules, the concentration of these inhibitory molecules would be low, and very few cells would have bound the number of inhibitory molecules needed to prevent cell division. Thus, at the beginning of development, most cells would divide, with the value of the ***Mitotic Fraction, m***, being close to **1**. However, as the embryo grows, more and more cells are present to produce inhibitory molecules, the concentration of these inhibitory molecules would increase, more and more cells would have bound the number of inhibitory molecules needed to prevent cell division, fewer and fewer cells would be able to divide, and the value of the ***Mitotic Fraction, m***, would decline.

Remarkably, when we rephrase this idealized case in mathematical terms (see SUPPLEMENT), such a decline in the ***Mitotic Fraction, m***, can be seen to occur in exactly the form as the ***Binary Cellular Universal Mitotic Fraction Equation*** (#1). Furthermore, all of the biochemical features of endocrine action appear in the “***a***” ***Parameter*** of this equation:

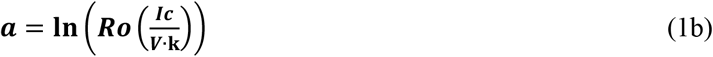

**Ic** is the number of inhibitory growth factor molecules made by each cell, ***V*** is the volume in which the embryo develops, such as an egg or a uterus, and **Ro** is the number of receptors per cell.

### The Binary Cellular Universal Growth Equation

Taking into account one more parameter, ***c***, the ***Cell Cycle Time***, the average length of time that it takes a cell to divide, allowed us to derive this ***Density Dependent*** expression for the how fast animals grow:

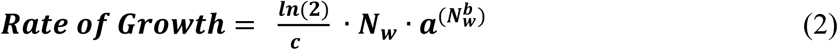

Integration (see the SUPPLEMENT) yields the relationship between ***age, t***, and ***size, N***:

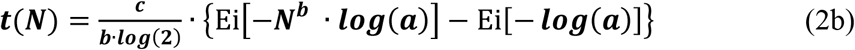

and the relationship between ***size, N***, and ***age, t***:

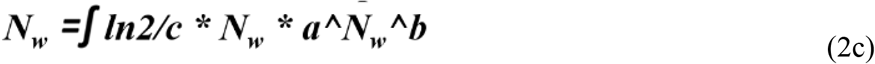

We have called these variations different forms of the ***Binary Cellular Universal Growth Equation*** (#2), as we found it to closely capture growth, from fertilization until maturity, for each 13 species listed above (SUPPLEMENT-TABLE A1).The values of the ***a, b*** and ***c*** parameters of the ***Binary Cellular Universal Growth Equation*** (#2) capture the specific growth characteristics of each animal (Details can be found in the **SUPPLEMENT** and Reference54). For humans, ***a*≈0.943, *b*≈0.169** and ***c*≈1.016** (in units of days) (**Figures 1**,**3**,**4, Supplement-Figure-III.2**). The resulting growth curve for humans, chickens, nematodes, and mice, are shown in **Figure 1, Supplement-Figure-I.4, SUPPLEMENT-TABLE A1**, and for a human fetus in **Figure 3** and **Supplement-Figure-III.2**.

**Figure 3.**
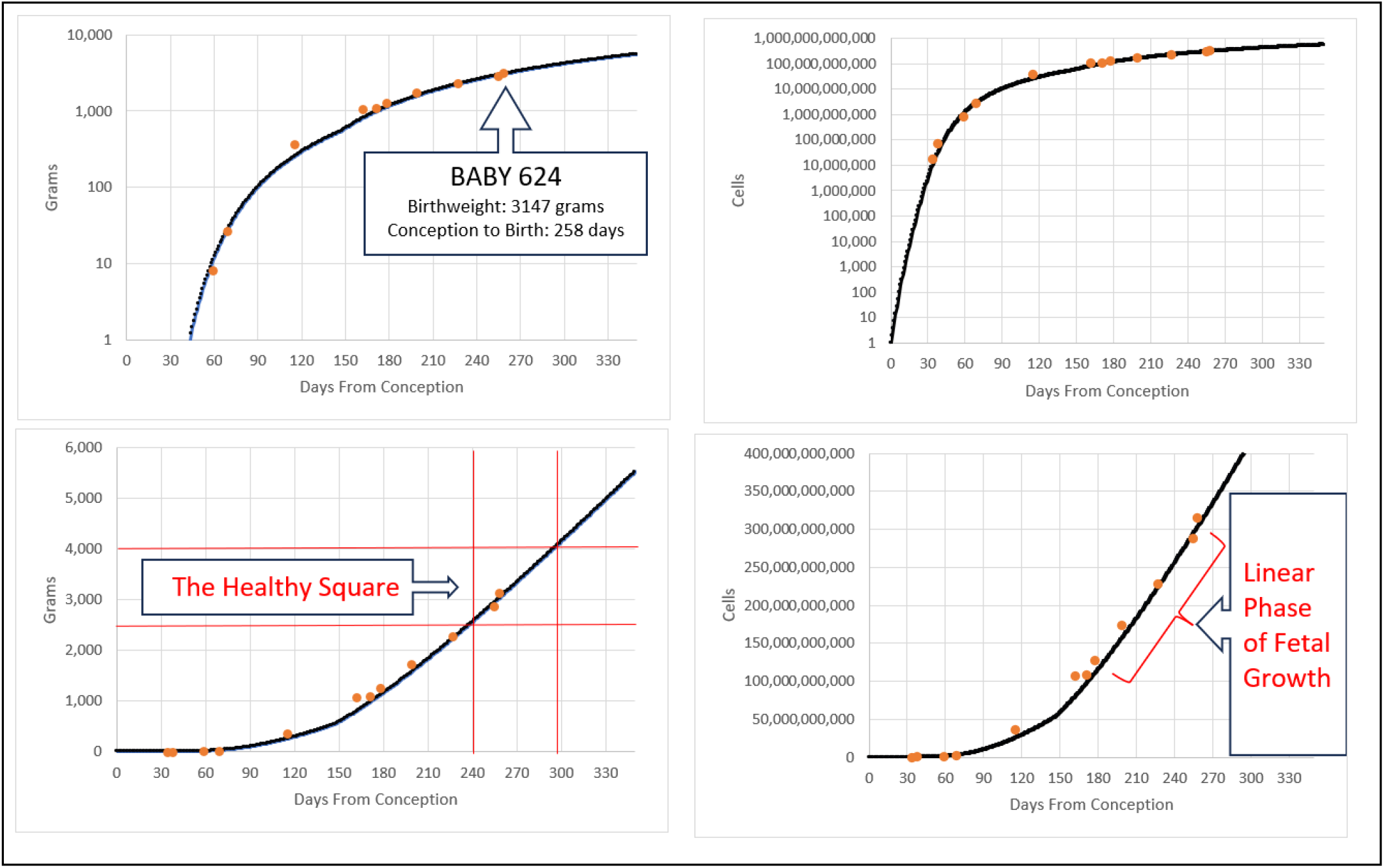
***Binary Cellular Fetal Growth Charts*** for an average size baby, arriving at a ***Healthy Square*** bounded by a conventionally healthy ***Birthweight***, below 4,000 grams and above 2,500 grams, and at a healthy ***Fetal Age***, more than 230 days after conception, the time by the ***Binary Cellular Universal Growth Equation*** when an ***Average Fetus*** would reach 2,500 grams. The last datapoint identifies birthweight. Other values are from ultrasound, as described in the next paper in this series.^7^ The dark line is the ***Binary Cellular Universal Growth Equation***

**Figure 4.**
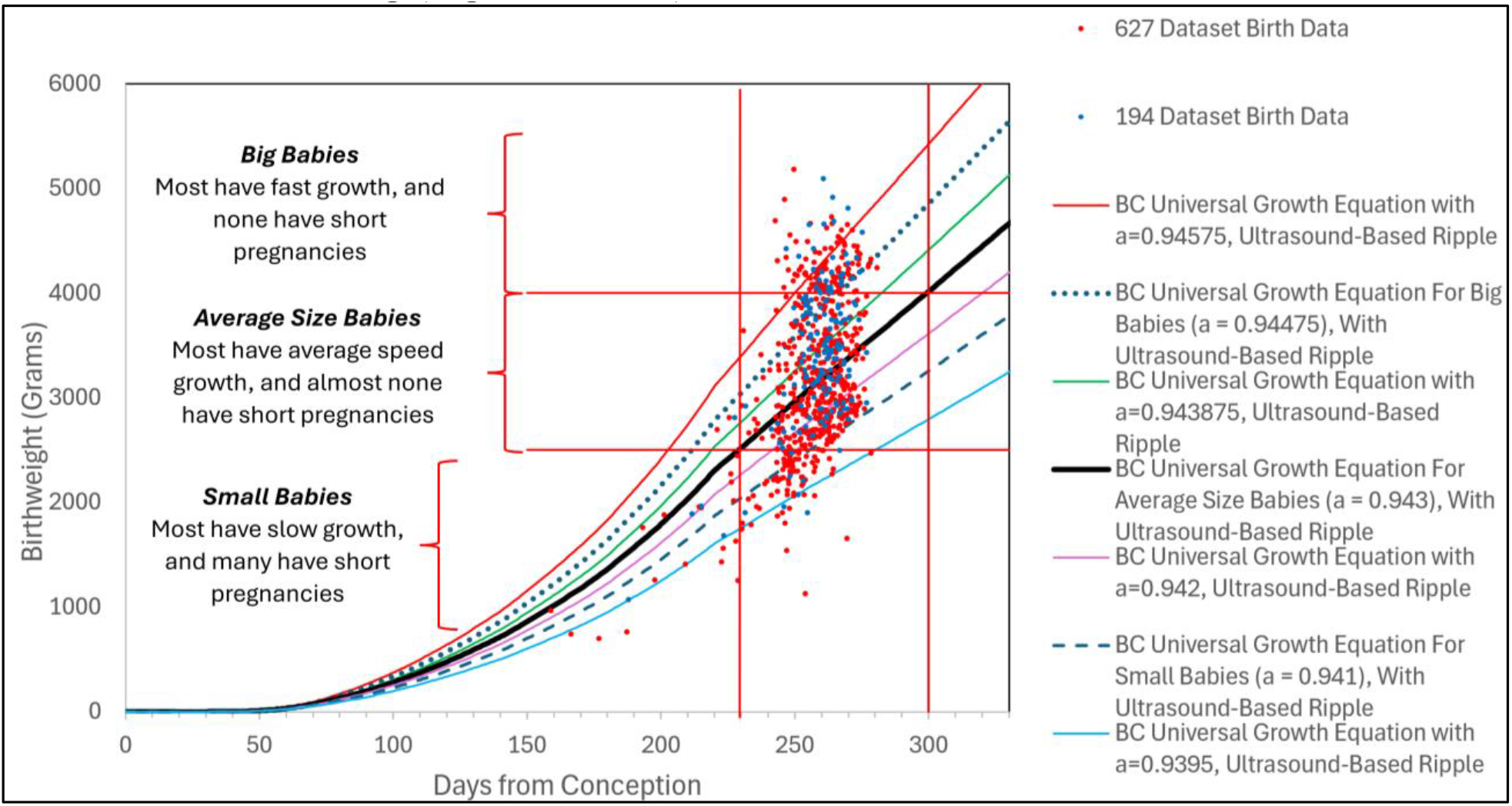
A ***Binary Cellular Fetal Growth Chart*** with ***Fetal Growth Curves*** differing in the ***“a” Parameter*** of the ***Binary Cellular Universal Growth Equation***, together with the ***Birthweights*** of a set of babies studied in the third paper in this series.^8^

### *Integration* and *Empirical Ripple Adjustment* of the *Binary Cellular Universal Growth Equation*

The curves in **Figure 1** for capturing the ***Size*** of animals, ***N***_***w***,_ over ***time, t***, by the ***Binary Cellular Universal Growth Equation*** (#2), were created by the Runge–Kutta method. A simpler way to execute such integrations, which we shall use here, breaks down growth into chunks of ***time***, Δ***t***_***C***_, in units of ***Cell Cycle Times, c***. This can be done in Excel, which also permits the fine scale fit of the ***Binary Cellular Universal Growth Equation*** (#2) to data, a method we call ***Empirical Ripple Adjustments*** (***R***) (**Figures 3** and **4;** for details, see the SUPPLEMENT).

### The Curve of the *Binary Cellular Universal Growth Equation* is Quite Similar to Other Fetal Growth Curves

Comparisons can be seen in **Supplement-Figure III.37** of the ***Binary Cellular Universal Growth Equation*** (with the ***a, b, c*** and ***R*** parameters/factors for average human fetal growth), the 50^th^ percentile of the NICHD ***Fetal Growth Curve***, and the 50^th^ percentile of the INTERGROWTH-21 ***Fetal Growth Curve***, revealing them to be remarkably similar. Comparison of ***Weight-Gain Rate*** based on the ***Binary Cellular Universal Growth Equation*** (#2), with the 50^th^ percentile of the NICHD^102^ and INTERGROWTH-21^103^ ***Fetal Growth Velocity Curves***, again revealed their similarity (**Supplement-Figure III.38**).

### The ∼Linear Phase of Human Fetal Growth

Over the last three months of pregnancy, when most fetuses have exceeded 1500 grams (∼150 billion cells), the ***Binary Cellular Universal Growth Equation*** (#2) flattens out (**Figures 3, 4, Supplement-Figures-I.16**,**17**,**18**,**21**, and **III.2**). By linear regression, this occurs with a slope of **26.038** and a correlation coefficient of ***r***^***2***^=**0.9996** (**Supplement-Figure-I.19**). In simple terms, from ∼7 to ∼10 months after ***conception***, the average fetus increases by about 26 grams, and about 3 billion cells, each day. We call this the ***∼Linear Phase of Human Fetal Growth***. The ***∼Linear Phase*** doesn’t extend back in the first 6 months of life, and it appears to end after birth (**Supplement-Figure-I.20**).

### The *Binary Cellular Universal Growth Equation* Gives Biologically-Based Data-Driven *Growth Curves*

The ***Binary Cellular Universal Growth Equation*** (#2) affords the opportunity to create biologically-based, data-driven, ***Fetal Growth Curves***. Shown in **Figure 4** and **Supplement-Figure-I.21** are various ***Binary Cellular Universal Growth Equations*** (#2) with different values for “***a****”* ***Parameter***, which, as described in the third paper in this series, captures fetus-to-fetus variation in growth.^8^

### The *Binary Cellular Universal Growth Equation* Can Be Used to Asess *Fetal Growth*

***Binary Cellular Universal Growth Equation Curves***, with various values for “***a****”* ***Parameter***, can be compared with ***Birthweights*** and ***Fetal Weights*** at their ***Times Since Conception*** (**Figures 4**), as determined by the ***CRL Fetal Age Equation*** (#7a). (The ***Binary Cellular Equations*** for making the calculations are derived, described, and tested, below, and in the next two papers in this series.^7,8^) We call the images that capture this information ***Binary Cellular Fetal Growth Charts***, and, as we shall see in the third paper in this series, they provide a useful way to predict and understand fetuses that are growing ***Too Slow*** or ***Too Fast***, leading to babies being born ***Too Small*** or ***Too Big*** (**Figures 3** and **4**).^8^

### *Binary Cellular Analysis* of the *Creation* and *Growth* of *Body Parts*

Embryos create ***Body Parts***. Obstetricians rely upon the ultrasound measurement of these ***Body Parts*** to gauge the ***Size*** and ***Growth*** of fetuses. Thus, the practical task of managing the health of the fetus and its mother reaches down to the ancient and fundamental biology of the creation of the ***Tissues, Organs***, and ***Anatomical Structures*** of the body, to arrive at actionable estimates of the ***Size*** of the fetal body.

### The Allometric Growth Equation

For more than a century,^104-106^ biologists have observed that the ***Size, p***, of many parts of the body, when compared with the ***Size, w***, of the body as a whole, frequently appear on log-log graphs as straight rows of ***p***-***w***-pairs, that is:

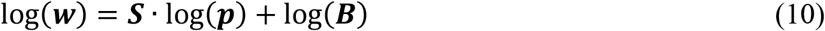

### This observation is called *Allometric Growth*, and the *Allometric Growth Equation. Binary Cellular Analysis* of *Allometric Growth*

Why parts of the body grow in this ***Allometric*** fashion, and what the ***S*** and ***B*** parameters the ***Allometric Growth Equation*** mean, have long been mysteries.^105,106^ To solve these problems, we again return to the methodology of ***Binary Cellular Analysis***, to examine, in units of numbers of cells, ***N***, the creation and growth of the various parts of the body, ***N***_***p***_, relative to the body as a whole, ***N***_***w***_, examined on log-log graphs.

### Data for Binary Cellular Analysis of Allometric Growth

We assembled values of ***N***_***p***_ vs. ***N***_***w***_ for ***Body Parts***, (***Tissues, Organs***, and ***Anatomical Structures***) for animals for which every cell has been characterized from the single fertilized ovum onward, providing ***Cell Lineage Charts*** (**Figure 5** and **Supplement-Figures-I.26&32, Supplement-Figures-I.24-27**). These species include the nematode worms ***Caenorhabditis elegans*** and ***Meloidogyne incognita*** (**Figure 5** and **Supplement-Figure-I.26 Supplement-Figures-I.24-26**), and the chordate tunicate ***Oikopleura dioica*** (**Supplement-Figure-I.27**) (see METHODS section for references).

**Figure 5:**
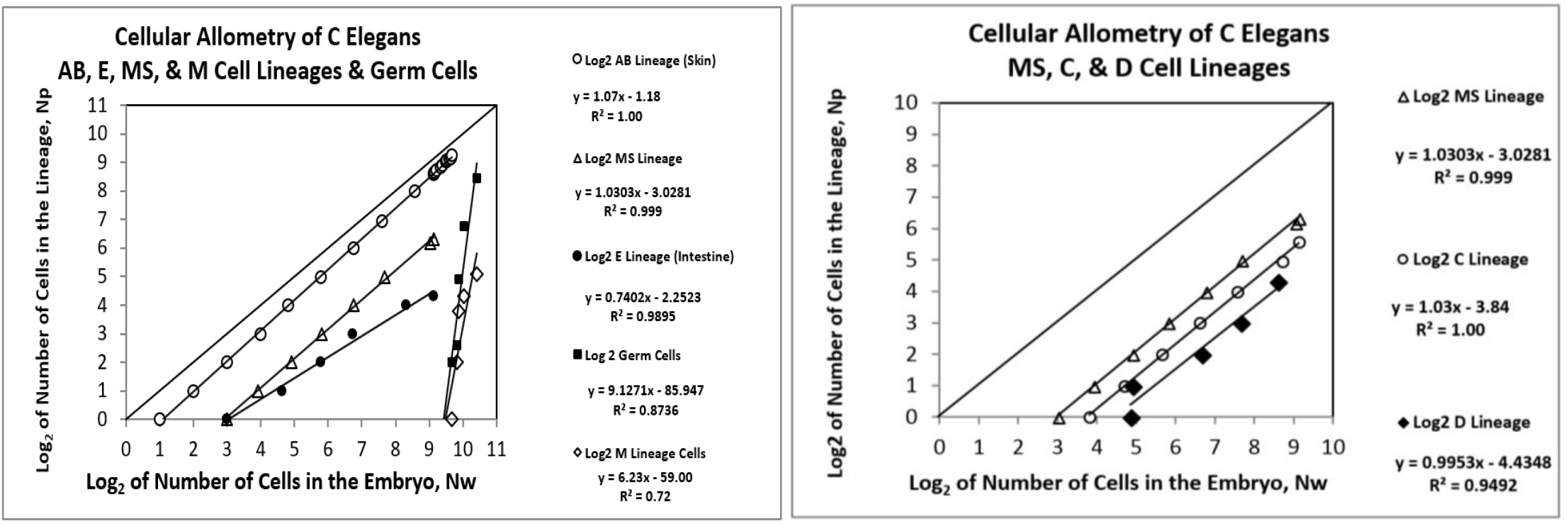
***Binary Cellular Analysis* of *Allometric Growth* of *Tissues, Organs***, and ***Anatomical Structures*** of **developing *C elegans* nematode**. The **AB** lineage forms the skin, the **E** lineage the intestine, the **MS** lineage the mesoderm, the **C** lineage the body wall muscle.

We also assembled values of ***N***_***p***_ vs. ***N***_***w***_ for ***Body Parts***, (***Tissues, Organs***, and ***Anatomical Structures***) for animals for which we lack information on every mitotic relationship early in development, but for which there is abundant information on ***Body Part Size*** later in development. These species (and the body parts examined) include: chicks (gizzards, livers, hearts, kidneys), rats (livers, brains, kidneys, forelegs, ears, stomachs, spinal cords), zebrafish (eye lens), mice (livers, brains, kidneys, forelegs), goldfish, and humans (brains, livers, kidneys, lungs, pancreases, adrenals, thymuses, spleens, lower extremities, upper extremities, stomach, heart, intestines) (see METHODS section for references) (**Figure 6** and **Supplement-Figure-I.27, Supplement-Figures-I.28-33**).

**Figure 6:**
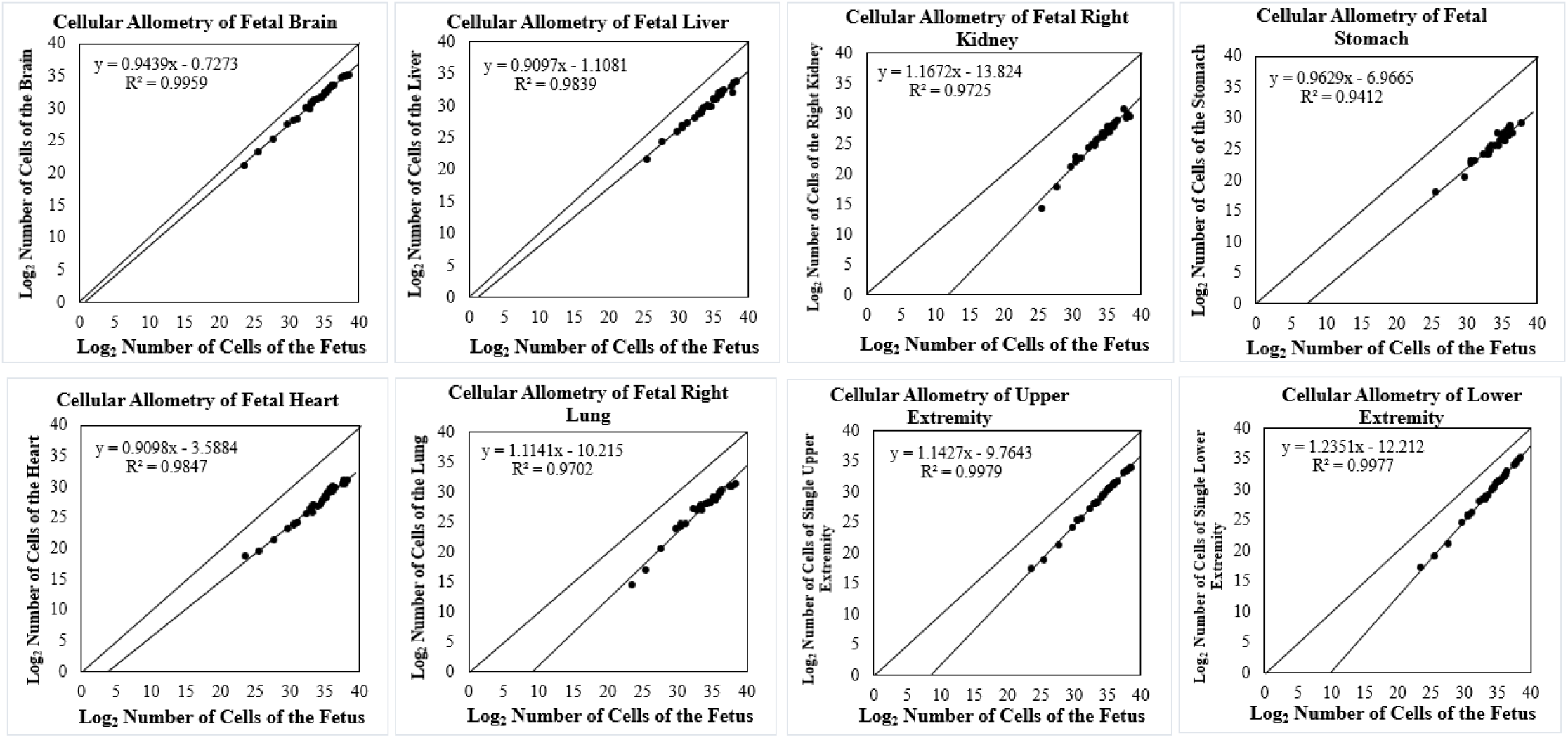
***Binary Cellular Analysis* of *Allometric Growth* of *Tissues, Organs***, and ***Anatomical Structures*** of the developing human fetus, from autopsy data. For additional human relative growth data, see **Supplement-Figures-I.31** and **32**

### The Binary Cellular Allometric Growth Equation

When this growth data, in units of numbers of cells, ***N***, was displayed on log-log graphs (**Figures 6, 7** and **Supplement-Figures-I.24-33**, including data on human body-part size measured at autopsy^100^ (**Figure 6, Supplement-Figures-I.28-33**), these ***N***_***p***_-***N***_***w***_-pairs (***Body Part Size***, vs ***Whole Body Size***) almost always appear as straight rows of dots, or:

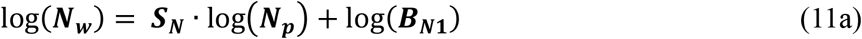

which is equivalent to:

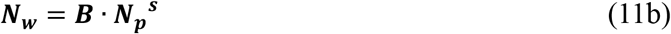

**Figure 7:**
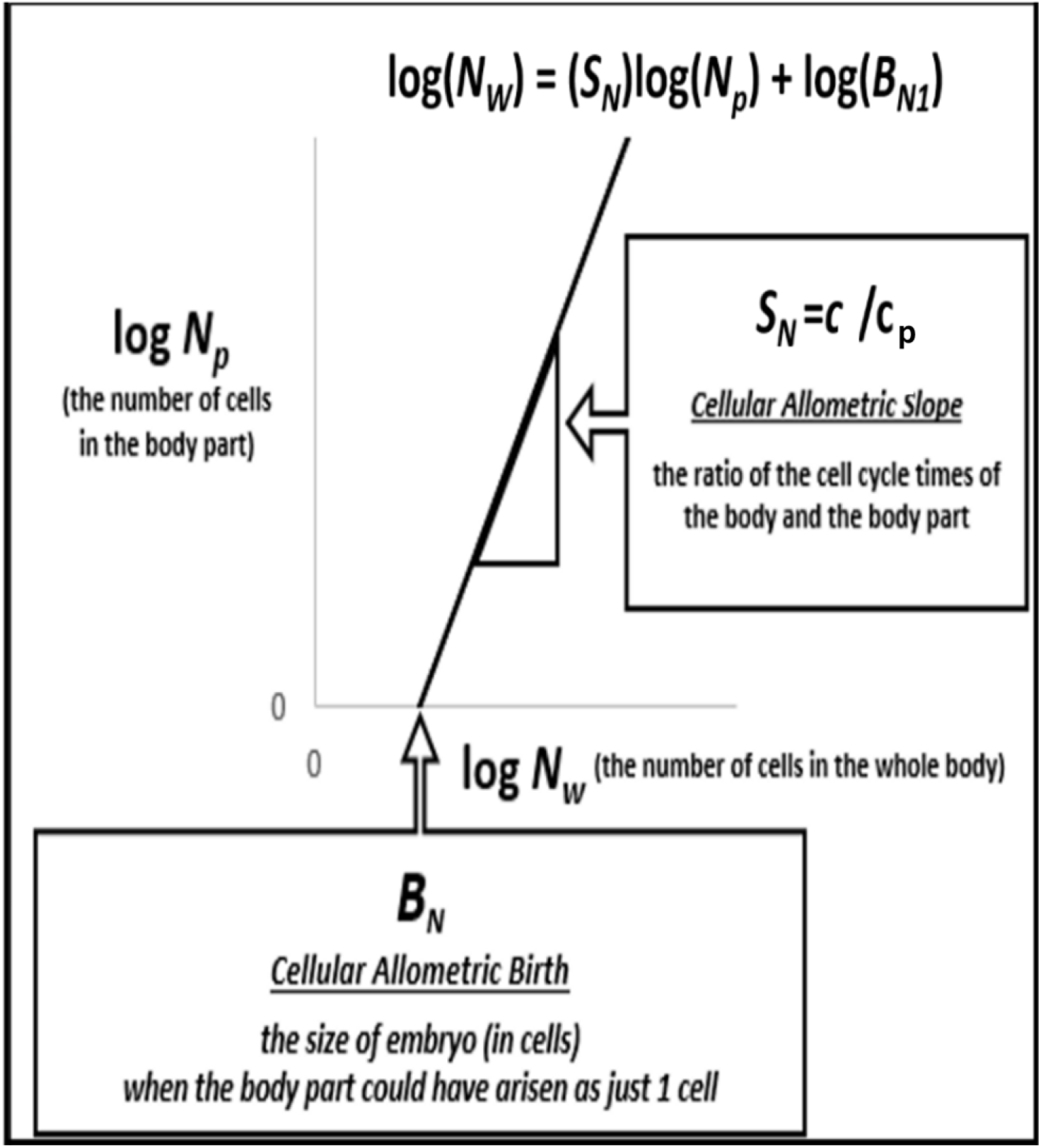
**The *Binary Cellular Allometric Growth Equation*** (#11)

We call this the ***Binary Cellular Allometric Growth Equation*** (**Figure 7**). We call ***S***_***N***_ the ***Binary Cellular Allometric Slope***, and ***B***_***N1***_ the ***Binary Cellular Allometric Birth***.

Note that this this equation was not constructed to ***fit*** the data we have on the growth of the ***Tissues, Organs***, and ***Anatomical Structures*** of the body, but, rather, was derived ***from*** this data. Thus, it doesn’t ***imitate*** the pattern by which animals create their ***Body Parts***, and determines their ***Sizes***, but ***reveals*** the way animals accomplish these tasks.

As we shall see, the value of this equation’s ***B***_***N1***_ parameter will show us that animals give rise to each ***Body Part*** from a single ***Founder Cell***, that is, ***N***_***p***_***=*1** when the embryo is ***B***_***N1***_ cells in size. We shall also see that the ***S***_***N***_ parameter captures whether a ***Body Part*** is growing faster (***S***_***N***_>**1**), or slower (***S***_***N***_<**1**), or at the same speed (***S***_***N***_=**1**), as the body as a whole, as determined by a ***Cell-Heritable*** change in the ***Cell Cycle Time, c***_***p***_, of the ***Founder Cell*** at its ***Binary Cellular Allometric Birth, B***_***N1***_.

### *Binary Cellular Analysis* Reveals that *Body Parts* Appear to Arise from *Single Founder Cells*

#### *Body Parts* Formation from *Single Founder Cells*, as Seen From the Value of *B*_*N1*_

Perhaps the most striking result that we have learned from the ***Binary Cellular Analysis*** of the many ***Body Parts*** of the many ***Species*** we have studied is that each ***Body Part*** appears to arise from a ***Single Founder Cell***. This occurs at the ***Body Part’s Binary Cellular Allometric Birth, B***_***N1***_ (**Figures 5, 6, Supplement-Figures-I.24-33**).

#### *Body Parts* and *Founder Cells* in Animals With Full *Cell Lineage Charts*

When we examined ***Body Parts*** of animals for which we have full ***Cell Lineage Charts***, showing all mitotic relations from conception onward, the data have always shown that the ***Binary Cellular Allometric Growth Equation*** points to where each ***Body Part*** has arisen from a ***Single Founder Cell*** at its ***Binary Cellular Allometric Birth, B***_***N1***_ (**Figure 5** and **Supplement-Figure-I.26, Supplement-Figures-I.24-27**). Note, for example, how the nematode ***C elegans* AB *Founder Cell***, whose progeny go on to form the **AB *lineage*** (the skin), arises when the embryo is just **2** cells in size, that is, at its ***Binary Cellular Allometric Birth*** of ***B***_***N1***_**≈2** (i.e., ***N***_***w***_**≈2** when ***N***_***p***_**≈1, Figure 5, Supplement-Figure-I.26, Supplement-Figures-I.24-26**). Similarly, note how the ***C elegans* E *Founder Cell***, whose progeny go on to form the **E *lineage*** (the intestine), arises when the embryo is **8** cells in size, that is, at its ***Binary Cellular Allometric Birth*** of ***B***_***N1***_**≈8** (i.e., ***N***_***w***_**≈8** when ***N***_***p***_**≈1**). Many other such examples of the creation of ***Body Parts*** from ***Single Founder Cells*** can be seen in **Figure 5** and **Supplement-Figures-I.24-27**.

#### *Body Parts* and *Founder Cells* in Animals Without Full *Cell Lineage Charts*

When we examined ***Body Parts*** of ***Vertebrate*** animals for which we don’t have full cell lineage data showing all mitotic relations from conception onward, but for which have abundant information on the size of ***Body Parts*** later in development, the data have always shown that the ***Binary Cellular Allometric Growth Equation*** points to where we would expect to find a ***Single Founder Cell*** at its ***Binary Cellular Allometric Birth, B***_***N1***_, early in development, and never when embryos are smaller than **2** cells. (**Figure 6** and **Supplement-Figures-I.27-33**). Note, for example, in **Figure 6** and **Supplement-Figure-I.32**, for the human heart ***B***_***N1***_**≈8**, suggesting that the heart could have arisen from a ***Single Founder Cell*** when the embryo was **∼8** cells in size. For the liver, ***B***_***N1***_**≈4**, for each kidney, ***B***_***N1***_**≈*2***,***000***, for the stomach, ***B***_***N1***_**≈100**, for the brain, ***B***_***N1***_**≈2**, for each lung, ***B***_***N1***_**≈500**, for the arms, ***B***_***N1***_**≈*500***, for the legs, ***B***_***N1***_**≈*1***,***000*** (**Figure 6** and **Supplement-Figure-I.32**). Many other such cases, in a great range of vertebrates, can be seen in **Supplement-Figures-I.28-32**.

#### *Body Part* Growth, *S*_*N*_, is Set by *Cell-Heritable* Change in *Cell Cycle Time, c*_*p*_, of the *Founder Cell*

The ***Binary Cellular Allometric Slope, S***_***N***_ of the ***Binary Cellular Allometric Growth Equation*** captures whether a ***Body Part*** is growing faster (***S***_***N***_>**1**), or slower (***S***_***N***_<**1**), or at the same speed (***S***_***N***_=**1**), as the body as a whole. The mathematical dissection of the ***Binary Cellular Allometric Growth Equation***, whose details can be found in the SUPPLEMENT, tells us that this arises from a ***Cell-Heritable*** change in the ***Cell Cycle Time, c***_***p***_, of the ***Founder Cell*** at its ***Binary Cellular Allometric Births, B***_***N1***_:

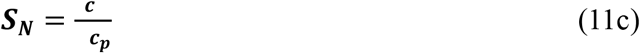

That the ***Cell Cycle Time, c***_***p***_, of the cells in a ***Body Part*** is due to a ***Cell-Heritable*** change from the ***Cell Cycle Time, c***, of the cells in the ***Body*** from which is arises can be seen in the straightness of the rows of ***N***_***p***_ vs ***N***_***w***_ dots appearing on the log-log graphs (**Supplement-Figures-I.26&32**, and **Supplement-Figures-I.24-33)**. This reveals that the length of time that it takes a ***Founder Cell*** to divide, and to give rise to the ***Body Part’s*** first two cells, is the same length of time that it takes the ***Progeny*** of that ***Founder Cell*** to divide, that is, all of the subsequent cells that make up that ***Body Part***.

#### *Binary Cellular Estimated Fetal Weight Equations* from *Binary Cellular Allometric Growth Equation*

We can now employ ***Binary Cellular Analysis*** to derive a new, biologically based, method for calculating ***Fetal Weight*** from ***Ultrasound Measurements***. Let us begin with the ***Binary Cellular Allometric Growth Equation*** (#11b) to examine the ***Allometric*** relationship between individual ***Fetal Ultrasound Measurements*** (***U***) and ***Birthweights*** (***B***_***w***_):

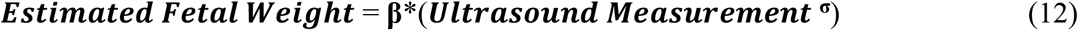

We call this expression the ***1***^***st***^***-Step-Binary Cellular Estimated Fetal Weight Equation***. In the next paper in this series,^7^ we shall put this equation to work, and test it, against patient data, revealing that the most promising of these expressions, the ***Abdominal Circumference Binary Cellular Estimated Fetal Weight Equation***, performs better than other currently used ***Fetal Weight Equations***, while also yielding up new methods for improving ultrasound size assessment.

#### *Binary Cellular Allometry*: A Way to Find the Best *Ultrasound* Measures For Estimating *Fetal Weight*

For decades, it has been realized that some ***Ultrasound Measurements***, such as ***Abdominal Circumference***, are better for estimating ***Fetal Weight*** than others. The ***Binary Cellular Allometric Slope, S***_***N***_ determined by the ***Cell-Heritable*** Change in ***Body Part Cell Cycle Time, c***_***p***_, provides an explanation for why some body measures are better that others, as ***Body Parts*** that grow more rapidly, relative to the body as a whole, will give a stronger signal than slow growing ***Body Parts***. This means that a closer examination of the allometric relationship between various body measures, such as those shown in **Supplement-Figures-I.31** and **32**, can help find those ideal ***Ultrasound Measurements***, a point to be expanded upon in the DISCUSSION of the next paper in this series.^7^

## DISCUSSION

### *Binary Cellular Analysis* Provides New Equations That Describe *Growth* and *Development*

Here we have seen that a consideration of animal and human bodies in units of numbers of cells, ***N***, an approach we call ***Binary Cellular Analysis***, provides:

- A new framework for examining embryonic development, ***Binary Cellular Biology***.
- New methods for measuring the occurrence of ***Cellular Selection***, resulting from ***Differential Cellular Proliferation***, the force which molds the formation of the body from a single fertilized ovum into a multicellular animal.^1,2^
- New ways to help obstetricians guide each fetus to a ***Birthweight*** and ***Pregnancy Length*** that has the highest possible chance of a favorable outcome.
- New equations for comprehending ***Growth*** and ***Development*** from conception to adulthood.

These ***Binary Cellular Equations*** include:

1. The ***Binary Cellular Universal Growth Equation***, which captures growth from conception to adulthood, and provides a method for creating biologically-based, data-driven, ***Growth Curves***.
2. The ***Binary Cellular Universal Mitotic Fraction Equation***, which lies within the ***Binary Cellular Universal Growth Equation***, and captures how growth to adult size is determined by the decline in the fraction of cells dividing, the ***Mitotic Fraction***.
3. The ***Binary Cellular Allometric Growth Equation***, which captures the ***Creation*** and ***Growth*** of the ***Tissues, Organs***, and ***Anatomical Structures*** of the body.
4. The ***Binary Cellular Estimated Fetal Weight Equation***, derived from the ***Binary Cellular Allometric Growth Equation***, which captures the relationship between ***Ultrasound Measurements*** and the ***Size*** of the body as a whole.

These ***Binary Cellular Equations*** were not made up to ***fit*** growth data, but were derived ***from*** growth data. Thus, they don’t ***imitate*** the pattern by which animals ***Grow*** and ***Develop***, but ***reveal*** the way, and the processes, by which, we ***Grow*** and ***Develop***.

### *Binary Cellular Analysis* Reveals How We Grow

How does the body grow from a single fertilized ovum into a creature of mature ***Size***? Capturing this information on the ***Growth*** of a ***Fetus*** is of particular importance in obstetrical practice, where being able to understand, and predict babies being born ***Too Small***, or ***Too Big*** provides essential information for the management of the health of the newborn and its mother.

### The ***Binary Cellular Universal Growth Equation*** Reveals How We Grow

As we have seen here, by considering growth data in units of numbers of cells, ***N***, that is, ***Binary Cellular Analysis***, we have been able to discover, for a great range of animals, including nematodes, frogs, chickens, cows, geese, quail, turkeys, mice, rats, fish, mollusks, and humans, from conception to adulthood, that growth occurs by the same expression: the ***Binary Cellular Universal Growth Equation***.

Mathematically, each of the parameters (***a, b, c***) of the ***Binary Cellular Universal Growth Equation*** (#2) has a specific impact on the shape of the body’s “***S-shaped***” growth curve. The ***“c” Parameter***, the ***Cell Cycle Time***, determines the speed of growth, expanding or contracting the “***S***” like an accordion. The ***a*** and ***b*** parameters determine the curviness of the “***S***”.

Biologically, each of the parameters of the ***Binary Cellular Universal Growth Equation*** (#2) is linked to a specific aspect of cell division. The ***“c” Parameter***, the ***Cell Cycle Time***, describes how long it takes cells to complete mitosis, once they have begun cell division. The ***Mitotic Fraction, m***, reflects how many cells are engaged in mitosis; ***m*** decreases as we become larger, as determined by the ***a*** and ***b*** parameters of ***Binary Cellular Universal Mitotic Fraction Equation*** (#1), which lies within the ***Binary Cellular Universal Growth Equation*** (#2).

### The ***Binary Cellular Universal Mitotic Fraction Equation*** Links Fetal Growth to Endocrinology

Why, across the animal spectrum, is growth driven by the decline in the ***Mitotic Fraction***, the fraction of the body’s cell that are dividing, by the ***Binary Cellular Universal Mitotic Fraction Equation***? One intriguing hint arose from the mathematical examination of the idealized case of the growth of cells that produce inhibitory growth factor molecules, acting in a constant volume, such as an egg or a uterus. What emerged from this exercise is that the fraction of cells not having bound one or more such molecule declined as the embryo increased in size, in exactly the same form at the ***Binary Cellular Universal Mitotic Fraction Equation***. Thus, little more than the conventional thermodynamics of all chemical reactions is sufficient to account for growth by the ***Binary Cellular Universal Mitotic Fraction Equation***, a finding that calls out for experimental examination. This also raises the intriguing possibility that the endocrine pharmacology revealed by such an analysis might well be employed to reverse abnormal growth.

Key to the result of this mathematical exercise is growth within a constant volume. Virtually all animals start out life in the constant volume of an egg or a uterus. Perhaps the invention of the egg was the critical requirement needed for the emergence of the first multicellular animal from its single cell ancestor.^107^

For human fetuses, this finding of an idealized case of ligand action in a constant volume motivates a consideration of hormones, such as insulin, leading to downstream action of inhibitory growth factor molecules, such as TGF. The examination of the genetics of such candidate molecules^108,109^,110 or their laboratory values,^111,112^ may shed light on such a possibility.

As we shall see in the third paper in this series,^8^ ***Binary Cellular Analysis*** of ultrasound data reveals that ***Small, Average***, and ***Large*** newborns were ***Slow-, Average-*** and ***Fast***-***Growing*** fetuses, differing by the ***“a” Parameter*** of the ***Binary Cellular Universal Growth Equation***, the parameter that modeling suggests could reflect the endocrinological control of fetal growth. These findings reveal possibilities for the detection and pharmacological treatment of fetuses that grow ***Too Fast*** or ***Too Slowly***, as well as for new ways to help obstetricians guide each fetus to a ***Birthweight*** and ***Fetal Age*** that has the lowest possible chance of complication.

### *Binary Cellular Analysis* Reveals How Control of Growth, by *Cell Cycle Time, c*. is Set by *Genome Size*

What might be the mechanism behind the ***Cell Cycle Time***, the ***“c” Parameter*** of the ***Binary Cellular Universal Growth Equation*** (#2), which not only determines how long it takes a cell to divide, but also determines how rapidly the body as a whole grows? Biochemically, the principal determinant of the ***Cell Cycle Time, c***, is the amount of DNA that the cell contains. The more DNA a cell has to copy, the longer it takes to divide. This was first found by Van’t Hof and Sparrow, who discovered a linear relationship between the total amount of amount of DNA and ***Cell Cycle Time*** for plant cells.^113^ Their observation has subsequently been confirmed in many studies of many types of organisms.^114,115^

### *Binary Cellular Analysis* Reveals How *Junk DNA* Can Control the *Speed of Growth*

For most animals, the largest part of the genome is non-coding DNA, sometimes called ***Junk DNA***,^116^ no doubt imprecisely named.^117,118^ The amount of this non-coding DNA has been found to be correlated with growth in salamanders,^119^ anurans,^120^ amphibians,^121^ insects,^122^ and copepods.^123^

The speed of growth has a profound impact on survival.^124,125^ Human fetuses that grow to full-size in 10 months, or in 8 months, have a much lower chance of survival than fetuses that reach optimal size in 9 months.^126^ Non-coding DNA, which determines genome size, is very susceptible to duplication or deletion, and thus provides populations of animals with abundant genetic variation in genome size.^127-129^ Perhaps this genetic variation in genome size leads to genetic variation in the ***Cell Cycle Time, c***, which then leads to genetic variation in the speed of body growth. Such genetic variation would appear to give species a powerful resource to draw upon, so that they can evolve, by the bitter reality of Darwinian selection, to growth rates that give them the greatest chance of survival. Indeed, this makes us wonder whether ***Junk DNA*** may owe its very existence to its role in determining the speed of growth.

### *Binary Cellular Analysis* Reveals the Biological Logic of the *∼Linear Phase of Human Fetal Growth*

One of the striking findings of our mathematical examination of human fetal growth is that the ***Binary Cellular Universal Growth Equation*** (#2) flattens into a roughly constant rate of growth in the last months of pregnancy, a quality we call the ***∼Linear Phase of Human Fetal Growth***. In simple terms, in the last months of pregnancy, an average size fetus increases by about 26 grams each day. This observation of linearity of growth in the third trimester was observed by Mongelli and colleagues in 2015,^130^ a quality they called “quasi-linear”, a feature that has continued to attract use and interest.^131^

The ***∼Linear Phase of Human Fetal Growth*** does not appear to be a quirky accident, as it is determined by the various parameters (***a, b***, and ***c***) of the ***Binary Cellular Universal Growth Equation*** (#2), each of which is specified genetically: ***a*** and ***b*** most likely by the hormones and growth factor molecules that regulate cell division, and their genes, and ***c***, the ***Cell Cycle Time***, by genome size.^132^ Clearly, the human growth curve has been molded by evolution to occur at a remarkably constant, linear rate, over the last months of pregnancy.

A plausible hypothesis for the linearity of growth at the end of pregnancy is that when a fetus reaches the rate of growth that occurs at the maximally possible rate of provision of nutrients, then Darwinism would drive our species to a ***Binary Cellular Universal Growth Equation*** (#2), with its genetically determined parameters, ***a, b***, and ***c***, yielding the ideally maximal constant rate of growth. Departure from this ideally maximal constant rate of growth would be expected in the past to reduce the fitness of the future child and its mother, and to be eradicated by ***Natural Selection***. As we shall see in the in the third paper in this series, ***Binary Cellular Analysis*** reveals precisely this force behind human babies that are born ***Too Small*** and ***Too Big***.^8^ ***Binary Cellular Analysis*** provides us with a range of options for reducing these unfortunate possibilities, as discussed below, and outlined in greater detail in the third paper in this series.^8^

### *Binary Cellular Analysis* Reveals How We Make Our *Body Parts*

How does the body make its ***Tissues, Organs***, and ***Anatomical Structures***? ***Capturing*** this information of the relationship between the ***Size*** of the ***Body***, and the ***Size*** of its ***Body Parts***, is of particular importance in obstetrical practice, where ***Fetal Weight*** is estimated from ***Ultrasound Measurement*** of these ***Body Parts***.

### The *Binary Cellular Allometric Growth Equation* Reveals How We Make Our *Body Parts*

By again considering data on growth, in units of numbers of cells, ***N***, the method we call ***Binary Cellular Analysis***, we have been able to discover that the size of the various parts of the body, ***N***_***p***_, relative to the body as a whole, ***N***_***w***_, almost always fits an equation of the same form: the ***Binary Cellular Allometric Growth Equation***. The ***Parameters*** of the ***Binary Cellular Allometric Growth Equation***, the ***Binary Cellular Allometric Birth*** (***B***_***N1***_), and the ***Binary Cellular Allometric Slope*** (***S***_***N***_), capture the basic biology of ***Body Part*** formation (**Figure 2** and **Supplement-Figure-I.3**). The ***Binary Cellular Allometric Birth*** (***B***_***N1***_), lies where the ***Binary Cellular Allometric Growth Equation*** crosses the ***x***-axis, thus where log(***N***_***p***_)**=0**, thus where ***N***_***p***_**=1**, that is, when the ***Body Part*** could have been a ***Single Founder Cell***. The the ***Binary Cellular Allometric Slope*** (***S***_***N***_), appears on log-log graphs as a measure of whether a ***Body Part*** is growing faster (***S***_***N***_>**1**), slower (***S***_***N***_<**1**), or at the same speed (***S***_***N***_=**1**), as the body as a whole. ***S***_***N***_ is determined by ***Cell-Heritable*** change in ***Cell Cycle Time, c***_***p***_, of each ***Founder Cell***.

### The Binary Cellular Allometric Growth Equation Reveals Body Parts Arise from Single Founder Cells

From our ***Binary Cellular Analysis*** of the growth ***Body Parts*** of those animals for which we have full cell lineage data (**Supplement-Figures-I.24-27**), nematode worms and tunicates, the data have always shown that the origin of each ***Body Part*** arises from a ***Single Founder Cell*** at its ***Binary Cellular Allometric Birth, B***_***N1***_. Tunicates are of special importance here, as they are members of the chordate phylum to which we belong.

From our ***Binary Cellular Analysis*** of the growth ***Body Parts*** of those animals for which we don’t have full cell lineage data, but for whom we have abundant information on the size of ***Body Parts*** later in development, principally vertebrates, we have always found that the ***Binary Cellular Allometric Growth Equation*** points down to a ***Binary Cellular Allometric Birth, B***_***N1***_, where we would expect to find a ***Single Founder Cell***, that is, early in development and never when embryos are smaller than **2** cells (**Supplement-Figures-I.27-33**). This provides the information for where 4D cell imaging can be carried out to tell us whether the ***Body Part*** does, in fact, arise from a ***Single Founder Cell***.^133^ The single ***Founder Cell*** option is also most likely in terms of the evolutionary principal of ***Parsimony***^,134,135^ ***whose mathematical demonstration can be found in the SUPPLEMENT***.

### *Data from the Literature Suggesting Body Parts* Arising from a *Few* or *Single Founder Cells*

While few of the large structures from late in development have been measured to the ***B***_***N1***_ intersection point, the zebrafish eye lens comes close, as Greiling and Clark documented its growth from just 8 cells, when the embryo is 16 hours post fertilization.^96^ As can be seen in **Supplement-Figure-I.33, *Binary Cellular Analysis*** of their data reveal that the zebrafish eye lens also grows by the ***Binary Cellular Allometric Growth Equation***, pointing down to a ***Binary Cellular Allometric Birth*** when the embryo was about 2000 cell in size, and thus ***B***_***N1***_**≈2000** when ***N***_***p*≈**_**1**.

Other studies that have also found that many large anatomical structures arise from small numbers of cells.^136,137^ These studies include recent CRISPR/Cas9 ***cell lineage*** labeling studies.^138,139^

Many parts of the ***Drosophila*** body have also been observed from small numbers of cells (**Supplement-Figures-I.38-41**).^140-142^ The spiracular anlage^143^, the pole cells (which form the germ cells)^144-146^, and genital ***Imaginal Discs***^147,148^ ***have been seen from 2 cells onward. Imaging before these points in time was not carried out, so these structures might well have arisen from single cells, and the mathematical calculation of the Binary Cellular Allometric Birth, B***_***N1***_ for each of these body parts tells us when we should go look for the creation of these ***Body Parts***. Verma and Cohen^144^ have made videos that show the growth of ***Drosophila*** histoblast nests, which form the mass of the body, back to as few as 4 cells at the beginning of their videos; one wonders what would have been seen if one turned on the camera before then. Such a simple data collection exercise cries out for experimental examination. Furthermore, these videos make clear that histoblast nest cells increase in number by cell division rather than by recruiting unrelated cells.^144^ For a deeper review of cellular origins of Drosophila ***Body Parts***, see the SUPPLEMENT.

### The Binary Cellular Allometric Growth Equation Reveals Body Part Size Is Set by Cell Cycle Time, c_p_ *Body Part* Growth, *S*_*N*_, is Set By *Cell-Heritable* Change in *Cell Cycle Time, c*_*p*_, in *Founder Cells*

By examining ***Growth*** of our ***Tissues, Organs***, and ***Anatomical Structures*** in units of numbers of cells, ***N***, we have been able to see that the ***Speed of Growth*** of each ***Body Part***, captured by its ***Binary Cellular Allometric Slope, S***_***N***_, is determined by a ***Cell-Heritable*** change in the ***Cell Cycle Time, c***_***p***_, of its ***Founder Cell*** at its ***Binary Cellular Allometric Birth, B***_***N1***_. That the change in the ***Cell Cycle Time, c***_***p***_, of each ***Founder Cell*** is ***Cell-Heritable*** is made visible by the log-linearity of ***Body Part*** growth captured by ***Binary Cellular Allometric Growth Equation*** on log-log graphs, revealing the new value of ***c***_***p***_ arising in each ***Founder Cell*** continues in the ***Founder Cell’s*** progeny.

### *Body Part* Change in the *Cell Cycle Time, c*_*p*_, and *DNA Methylation*

There are a number of ***Cell-Heritable*** biological processes that influence the time it takes for a cell to divide, of which DNA methylation has been the most widely studied. ^149,150^ Methylated DNA takes longer to copy than unmethylated DNA, and thus methylation causes cells to take more time to divide.

### *Body Part* Change in the *Cell Cycle Time, c*_*p*_, is But One Example of *Cell-Heritable* Change

Of course, body parts display many ***Cell-Heritable*** changes in their phenotypes. Perhaps the most familiar examples are the ***Cell-Heritable*** changes in the expression of genes that mark the cell types that characterize the specific features of each of the body’s ***Tissues, Organs***, and ***Anatomical Structures***.^3^

### *Binary Cellular Allometric Growth Equation* Gives New and Better *Estimated Fetal Weight Equations*

As we have seen here, and will explore in greater detail in the next paper in this series,^7^ the ***Binary Cellular Allometric Growth Equation***, can give rise to ***Binary Cellular Estimated Fetal Weight Equations***, which can perform better than other currently used ***Fetal Weight Equations***, while also yielding up new methods for improving ultrasound size assessment.

### *Binary Cellular Analysis* Provides Practical Methods for *Managing Human Fetal Growth*

As we have seen here, and in the next two papers in this series, these new ***Binary Cellular Equations*** have at least six practical applications in health:

1. As we shall see in the next paper in this series,^7^ the ***Binary Cellular Estimated Fetal Weight Equations*** described above provides better, more accurate and more precise equations and methods for measuring of the ***Size*** of human fetuses from fetal ultrasound measurements.
2. As we shall see in the third paper in this series,^8^ the ***Binary Cellular Equations*** described above provides better, more accurate, and more precise, equations and methods for detecting the abnormal ***Growth*** of human fetuses that leads to birthweights that are ***Too Small*** or ***Too Big***.
3. As we shall see also in the third paper in this series,^8^ ***Binary Cellular Mathematics***, and specifically the ***Binary Cellular Universal Growth Equation***, provides the basis for deriving biologically-based, data-driven, ***Fetal Growth Curves***.
4. As we shall also see in the third paper in this series,^8^ the ***Binary Cellular Equations*** described above reveals previously hidden forces by which human fetuses grow to birthweights that are ***Too Small*** or ***Too Big***, specifically, growth that is ***Too Slow*** or ***Too Fast***, both of which map to the ***“a” Parameter*** of the ***Binary Cellular Universal Mitotic Fraction Equation***. Mathematical examination also reported in the third paper in this series indicates that the ***“a” Parameter*** tracks to the endocrinological forces that control growth.
5. As we shall also see in the third paper in this series,^8^ the ***Binary Cellular Equations*** described above reveals previously hidden opportunities for managing human fetuses that grow ***Too Slow*** or ***Too Fast***, so as to lower the chance of each fetus being born ***Too Small*** or ***Too Big***.
6. ***Binary Cellular Analysis*** also has the potential to examine childhood growth.

These practical applications will be reviewed in greater detail in the DISCUSSION sections of the second and third paper in this series.^7,8^

### Web-based Calculators Provide *Binary Cellular* Calculations That Aid Obstetric Care

A first-generation Web-based calculator for calculating ***Fetal Weight*** from the ***Abdominal Circumference Binary Cellular Estimated Fetal Weight Equation***, is now available at “https://kidzgrowth.com“. Encoding the additional ***Binary Cellular*** calculations described here is straightforward,^151^ and underway, including calculation of the speed of fetal growth from multiple, sequential, ***Ultrasound Measurements***, in units of ***Weight-Gain Rate*** (grams/day), and in units of the value of the ***“a” Parameter*** of the ***Binary Cellular Universal Growth Equation***, as well as projected values ***Fetal Weight*** in the days and weeks ahead, displayed on a ***Binary Cellular Fetal Growth Chart***, with the statistical limits of reliability also shown. Also to be provided by these calculators are tools for performance feedback, comparing each ***Ultrasound Measurement*** to the ultimate ***Birthweight***, so that sonographers and their supervisors can review performance over the long term, as well as the performance of each individual case. These web-based calculators make the results of these ***Binary Cellular Equations*** widely available for aiding in obstetric care of individual pregnancies.

## Supporting information

SUPPLEMENT Data and Details

## ACKNOWLEDGEMENTS

Many thanks for the superb collaborative contributions to this work of Daniel DiCorpo,^15^ Tahmid Ahmed,^16^ and Luke Huang,^17^

