## SUPPLEMENT Data and Details for "*The Binary Cellular Biology of Human Growth*: I. Quantifying the Creation and Growth of the Body and its Parts"

#### ***The Binary Cellular Biology of Human Growth*: II. Calculating the Size of the Human Fetus from the Size of its Parts**

#### ***The Binary Cellular Biology of Human Growth*: III. Identifying, Comprehending, and Ameliorating Normal and Abnormal Fetal Size and Growth**

James Selib Michaelson[1]

.

[1] Laboratory for Quantitative Medicine, and Departments of Pathology and Surgery, Massachusetts General Hospital and Department of Pathology, Harvard Medical School, Boston MA USA Mailing Address: 89r Bass Point Rd, Nahant, MA 01908,

### ***The Binary Cellular Biology of Human Growth*: I. Quantifying the Creation and Growth of the Body and its Parts**


**ABSTRACT**

Assessing and understanding the ***Growth*** and ***Development*** of animals generally, and of human fetuses specifically, provides essential information for the management of the health of the newborn and its mother. Here we show that a consideration of the formation of the body, in units of numbers of cells, ***N***, and the change in these numbers by the binary either/or decisions of cells to divide or not, an approach we call ***Binary Cellular Analysis***, provides new equations for quantifying the growth and development of the body and its parts. These equations include: 1) The ***Binary Cellular Universal Growth Equation***, which captures growth from conception to adulthood, and provides a method for biologically-based, data-driven, ***Growth Curves***; 2) The ***Binary Cellular Universal Mitotic Fraction Equation***, which lies within the ***Binary Cellular Universal Growth Equation***, and captures the mechanism of growth to adult size, control of cell division; 3) The ***Binary Cellular Allometric Growth Equation***, which captures the creation and growth of the ***tissues***, ***organs***, and ***anatomical structures*** of the body, and how the embryo creates each ***Body Part*** from a ***Single Founder Cell***; and, 4) ***Binary Cellular Estimated Fetal Weight Equations***, derived from the ***Binary Cellular Allometric Growth Equation***, which captures the relationship between ***Ultrasound Measurement*** and the ***Size*** of the body as a whole. These equations capture ***Cellular Selection***, resulting from ***Differential Cellular Proliferation***, which molds the formation of the body from a single fertilized ovum into a multicellular animal. The parameters of these equations capture differences in ***Size*** and ***Growth*** between animals across the taxonomic spectrum, and between human fetuses. From the ***Binary Cellular Universal Growth Equation***, the ***Growth*** of human fetuses can be measured, thus providing a much-needed tool for understanding the biological forces that cause a fetus to grow to ***Small***, ***Average***, or ***Large Size*** at birth, for providing the basis of biologically-based, data-driven, ***Growth Curves***, as well as for assisting the management of the obstetric care of the fetus and its mother.

#### INTRODUCTION

The Biology and Medicine of ***Growth*** and ***Cellular Selection***

***Growth*** is the essential feature of animal life. Driven by cell division, animal life begins with all cells dividing, ***Exponential Growth***, then almost all cells dividing, then progressively more cells becoming ***Quiescent***,[97-99] guiding the ***Growth*** of the animal body to its recognizable ***Size***. It is this fundamental embryological process of ***Cellular Selection***,[1,2,3] resulting from ***Differential Cellular Proliferation***, that is, some cells being ***Mitotic*** while others remain ***Quiescent***, which molds the ***Growth*** and ***Development*** of all animals, and lies behind the successful and unsuccessful growth of humans before and after birth.

The Evolutionary Origin of ***Growth*** and ***Cellular Selection***

More than half a billion year ago, our single-cell ancestor gave rise to the first multi-cellular animal by learning how to give up uncontrolled ***Exponential Growth*** in exchange for suppression of cell division.[4,5] It was this transformation of the ***Natural Selection*** of the ***Competing Cells*** of single-cell species, into the ***Cellular Selection*** of the ***Cooperating Cells*** of multicellular bodies, that makes ***Growth*** to specific ***Sizes*** possible. As we shall see here, and in the next two papers in this series,[6,7] gaining insight into this biologically deep, and ancient, process of ***Cellular Selection***, across the animal spectrum, and comprehending this information in a mathematical form that reliably captures the actual ***Growth*** of real animals, and real humans, can provide the obstetrician with the practical information needed to bring the child and its mother into the world with the best possible health.

The Nature of ***Growth***, and the Current Math for Capturing ***Growth***.

As we grow older, we grow bigger, and we grow slower. Biologists have long searched for an equation that fits, and explains, such animal ***Growth***, and its role in human health.[8-27] Unfortunately, none of these equations have been found to do a good job at capturing the actual growth of real animals, including humans.[28-30]

The absence of an equation that can accurately and precisely gauge the growth of humans, in fetal life and beyond, has posed a formidable challenge to the construction of the tools needed for guiding obstetric care.[31-34] For example, growth charts[35-47] have frequently had to rely on polynomial equations as curve fitting devices, unlinked to the biology of cell division. This has made the creation of a ***Universal Reference*** for ***Fetal Growth Charts***, which could capture the growth of the variety of fetuses, and the variety of populations in which they are born, only a hoped-for possibility.[48]

The Nature of ***Body Part Creation*** and ***Growth***, and the Current Math for ***Body Part Creation*** and ***Growth***

Early in life, embryos create ***Body Parts***. Biologists have long searched for a mathematical framework that fits, and explains, the ***Creation*** and ***Growth*** of these ***tissues***, ***organs***, and ***anatomical structures***.[121-123] Precisely how our ***Bodies*** make our ***Body Parts*** from ***Founder Cells***, and how they guide the resulting ***Body Parts*** to their mature ***Sizes***, has long been obscure. Capturing the information of the relationship between the ***Size*** of the ***Whole Body***, and the ***Size*** of the ***Parts*** of the ***Body***, is of particular importance in obstetrical practice, where the size of the fetus is estimated from ***Ultrasound Measurement*** of these ***Body Parts***.

***Binary Cellular Analysis*** Provides a New Way to Capture ***Size***, ***Growth***, and ***Creation***, of the ***Body*** and its ***Parts***.

To solve the problems outlined in the previous sections, we have examined data on the ***Size*** and ***Growth*** of developing animals and humans in units of numbers of cells, ***N***.[3,100,113] We call this method ***Binary Cellular Analysis***, as it examines the ***Binary*** choices made by our cells to divide, or not, and how these choices determine the number of cells in our ***Bodies*** and our ***Body Parts***.

As we shall see in this communication, and the two papers that accompany this communication,[6,7] such a ***Binary Cellular*** approach allows us to derive biologically-based, accurate, and precise ***Binary Cellular Equations*** for capturing the ***Size***, ***Growth***, and ***Creation***, of our ***Bodies*** and our ***Parts***, and for the assessment of the ***Size***, ***Growth***, and ***Age*** of developing animals, especially for humans. The equations include:

1. The ***Binary Cellular Universal Growth Equation***, which captures growth from conception to adulthood, and provides a method for creating biologically-based, data-driven, ***Growth Curves***.[34-48]
2. The ***Binary Cellular Universal Mitotic Fraction Equation***, which lies within the ***Binary Cellular Universal Growth Equation***, and captures the mechanism of growth to adult size, control of cell division, that is, **Cellular Selection**.
3. The ***Binary Cellular Allometric Growth Equation***, which captures the creation and growth of the ***tissues***, ***organs***, and ***anatomical structures*** of the body relative to the body as a whole, and how the embryo creates each ***Body Part*** from a ***Single Founder Cell***.
4. ***Binary Cellular Estimated Fetal Weight Equations***, derived from the ***Binary Cellular Allometric Growth Equation***, which captures the relationship between ***Ultrasound Measurements*** and the ***Size*** of the body as a whole.

As we shall see in this communication, and the two papers that accompany this communication,[6,7] these ***Binary Cellular Equations*** make it possible to make more accurate, precise, and actionable estimates of fetal ***Size*** and ***Growth*** from ***Ultrasound Measurement***, so as to help obstetricians guide each pregnancy to a ***Birthweight*** and ***Fetal Age*** at birth, a ***Pregnancy Length***,[49] with the highest chances of survival and health

In the second, following paper in this series,[7]we shall see that ***Binary Cellular Estimated Fetal Weight Equations*** for estimating human ***Fetal Weight*** provides improved assessment of ***Fetal Weight***, in comparison to other currently used ***Estimated Fetal Weight Equations***. We shall also see how ***Binary Cellular Mathematics*** can be put to work to improve the assessment of the ***Size*** and ***Growth*** of the human fetus from ***Ultrasound Measurement***.

In the third, final paper in this series,[7] we shall see that ***Binary Cellular Equations*** for estimating ***Fetal Weight*** from ***Ultrasound Measurements***, and for capturing ***Fetal Growth*** from conception onward, allows for the practical measurement and prediction of human fetuses that become ***Large***, ***Average***, and ***Small*** at birth. This ***Binary Cellular Fetal Growth Analysis*** provides a way to identify the biological mechanisms that drive human fetuses to being born ***Too Large*** or ***Too Small***, as well as for identifying new possibilities for the detection and pharmacological treatment of fetuses that are growing too slowly.[50,51,52] ***Binary Cellular Fetal Growth Analysis*** also provides new ways to help obstetricians guide each fetus to a ***Birthweight*** and ***Fetal Age*** that has the highest possible chance of a favorable outcome. Finally, we shall see how the ***Binary Cellular Universal Growth Equation***, which captures growth from conception to adulthood, provides a method for creating biologically-based, data-driven, ***Growth Curves***.[48]

#### METHODS

To examine data on the growth of animal bodies, we assembled data on the number of cells in the whole animal, $N_w$, by age, ***t***, in days from fertilization until maturity, for 13 species of animals: zebrafish (***Danio rerio***),[53] European sea bass (***Dicentrarchus labrax***),[54,55] mice (***Mus musculus***),[56,57] rats (***Rattus norvegicus***),[58] cows (***Bos taurus***),[59,60] bobwhite quail (***Colinus virgianus***),[61,62] domestic chickens (***Gallus gallus domesticus***),[63,64] turkeys (***Meleagris gallopavo***),[65] geese (***Anser anser***),[61] nematode worms (***C. elegans***,[83] ***M incognita***[84]), frogs (***Rana pipiens***),[66] clams (***Merceneria mercenaria***).[67,68,69] and humans (***Homo sapiens***),[70,71,72] Data on the numbers of cells in early embryos, from fertilization onward, were available for mice,[73,74] rats,[75] cows,[76] chickens,[77] turkeys,[78] nematode worms,[83,84] clams,[68] fish.[79,80] and humans.[81] Data on the numbers of cells in later embryos were assembled from published values in units of weight or volume by taking advantage of the finding that there are about $10^8$ cells/gram-cc.[82] Excel files (TABLE A2) characterizing these basic growth data ("**Basic Data Files**"), and the calculations made from these data ("**Calculations Files**"), are available on request. Also available to interested readers are excel files with the data on the growth of the parts of the body that we examined (see below).

To examine data on the growth of the parts of animal bodies, we assembled values of $N_p$ vs. $N_w$ (the number of cells in a body part, $N_p$, and in the body as a whole, $N_w$, at various points in time) for each of the ***Body Parts*** that have been characterized in the ***Cell Lineage Charts*** of ***Caenorhabditis elegans***[83]and ***Meloidogyne incognita***[84], nematode worms, as well as for ***Oikopleura dioica***, chordate tunicates (**Supplement-Figure-Z I.**S A9-A11).[85] We also assembled values of $N_p$ vs. $N_w$ for the ***Body Parts*** evident in embryos and juveniles of larger animals, again relying on the observation that there are about $10^8$ cells in every gram of tissue.[86,82] We assembled these values for the ***Body Parts*** of chicks (gizzards, livers, hearts, kidneys),[87] rats (livers, brains, kidneys, forelegs, ears, stomachs, spinal cords),[88] zebrafish (eye lens),[89-92] mice (livers, brains, kidneys, forelegs),[88,93] clams, goldfish, and humans (brains, livers, kidneys, lungs, pancreases, adrenals, thymuses, spleens, lower extremities, upper extremities, stomach, heart, intestines).[94]

Data were analyzed in Microsoft Excel.

Equations used here and in the related manuscripts are numbered (1), (2), etc. When the same equation is listed, but in different form, the various forms are given the same number, followed by a letter, that is (2b), (2c), (2d), etc. Equations only used in this supplement, and not in the related manuscript, are listed in numbers starting from (100).

#### RESULTS

***Binary Cellular Analysis***: Capturing ***Size***, ***Growth***, and ***Creation***, of the ***Body*** and its ***Parts*** in ***Cell Numbers***, ***N***.

Here, we report our studies of examining data on the ***Size*** and ***Growth*** of developing animals and humans in units of numbers of cells, ***N***, so as to decipher the occurrence, and mechanisms, by which the body determines its ***Size***, ***Growth***, and the ***Creation*** of its ***Parts*** (***tissues***, ***organs***, and ***anatomical structures***). We call this method ***Binary Cellular Analysis***, as it examines the ***Binary*** choices made by our cells to divide, or not, and how these choices determine the number of cells in our ***Bodies*** and our ***Body Parts***.

The Current Math for the ***Growth*** and ***Size***: The Body in Continuous Units of Width or Height or Weight, ***w***

As we grow older, we grow bigger, and we grow slower. Two centuries ago, Verhulst provided the first expression for such ***Density Dependent Growth***, the ***Logistic Equation***, which envisages growth as occurring exponentially, with a linear decline in the rate of growth, in continuous units of weight, volume, and length, ***w***:

$$\boldsymbol{Rate\ of\ Growth = Growth\ Rate\ Parameter \cdot Whole\ Body\ Size \cdot Growth\ Modifier} \quad (100)$$

or, more simply:

$$\frac{dw}{dt} = r \cdot w \cdot M \quad (101)$$

where ***w*** is the ***Size*** of the ***Whole Body***, which increases over time, and ***M*** is the ***Growth Modifier***, that decreases as we get larger. By leaving ***M*** undefined, this is sort of the Swiss-Army Knife of ***Density Dependent Growth Equations***, which Verhulst filled with:

$$M = \left(1 - \frac{k}{w}\right) \quad (102)$$

where ***k*** is the final size of the body in adulthood, a term frequently called the ***Carrying Capacity***. Combining Equations #101 and #102 gives us the ***Logistic Equation***, in condensed form:

$$\frac{dw}{dt} = r \cdot w \cdot \left(1 - \frac{k}{w}\right) \quad (103)$$

Note that when we are very small, the fraction ***k/w*** is negligible, ***M*** becomes close to **1**, and the size of the body, ***w***, just doubles, and doubles, and doubles, i.e. ***Exponential Growth***. But when we become full size, that is when ***k*** equals ***w***, ***k/w*** becomes **1**, ***M*** becomes **0**, and the rate of growth is **0**. So, the ***Logistic Equation*** captures the "growing slower as we grow bigger" quality of animal growth, with growth stopping when we reach adult size, ***k***. Regrettably, it doesn't do such a good job of capturing the growth of real creatures, including humans,[28-30] a problem we shall address next.

***Binary Cellular Analysis*** of ***Growth*** and ***Size***: The Body in Discrete Units of Numbers of Cells, $N_w$

Many ***Density Dependent Growth Equations*** have been considered since Verhulst - the ***Gompertz***,[10] the ***Von Bertanalfy***,[11] the ***West***,[13] to name just a few - none of which has been found to accurately capture the growth of real animals, including humans.[28-30] To solve this problem, we turned to actual growth data, to let real information on the growth of real animals tell us what the best ***Density Dependent Growth Equation*** might be.[100] ***Binary Cellular Analysis*** gives us a way to accomplish this, by reformatting of Equation #101 into units of numbers of cells, ***N***:

$$\frac{dN_w}{dt} = \left\{\frac{ln(2)}{c}\right\} \cdot N_w \cdot m \quad (104)$$

Where $N_w$ is the number of cells in the body as a whole. Now, our ***Growth Rate Parameter*** is {the ***Natural Log*** of **2** divided by the ***Cell Cycle Time***, ***c***, the amount of time that is takes a cell to divide, if it is engaged in mitosis}, while ***m***, the ***Mitotic Fraction***, is the fraction of cells that must be dividing to account for the growth of the body.

Data for the ***Binary Cellular Analysis*** of the Growth of the Whole Body.

To carry out the ***Binary Cellular Analysis*** of growth, we assembled data on the number of cells in the whole animal, $N_w$, by age, ***t***, in days from fertilization (***t*=0**) until maturity, for 13 species of animals: zebrafish (***Danio rerio***), European sea bass (***Dicentrarchus labrax***), mice (***Mus musculus***), rats (***Rattus norvegicus***), cows (***Bos taurus***), bobwhite quail (***Colinus virgianus***), domestic chickens (***Gallus gallus domesticus***), turkeys (***Meleagris gallopavo***), geese (***Anser anser***), nematode worms (***C. elegans***, ***M incognita***), frogs (***Rana pipiens***), clams (***Merceneria mercenaria***), and humans (***Homo sapiens***) (see METHODS section). Graphs of these various datapoints for animal size's ($N_w$) versus time (***t***), are shown in **Supplement-Figure-I.1**.

Graphing the number of cells in the body as a whole, $N_w$, vs. age, ***t***, from fertilization until maturity, reveals, for each of these 13 species, that animal growth occurs by ***S-shaped*** growth curves (**Supplement-Figure-I.1**). This is most easily viewed on log-log plots, because the greatest part of growth occurs rapidly at the beginning of life. For example, in terms of numbers of cells, $N_w$, humans grow about 11 orders of magnitude in the womb and about 2 orders of magnitude after birth. As we shall see below, the reason why all of these creatures have similar ***S-shaped*** growth curves is that they all grow by an equation of the same form, the ***Binary Cellular Universal Growth Equation***.

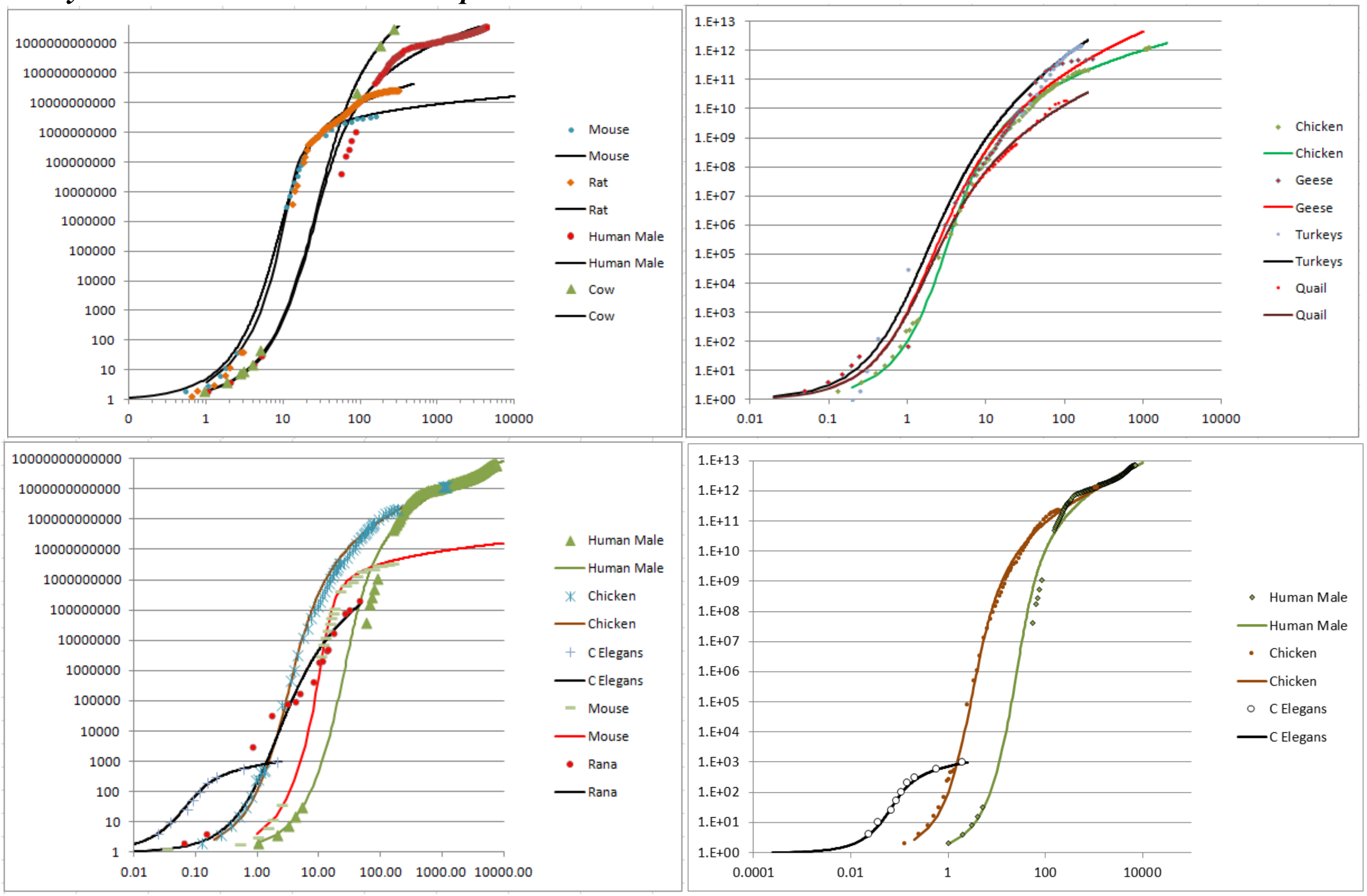


**Supplement-Figure-I.1**:**Growth, in units of numbers of cells, $N_w$, from fertilization until maturity, in units of days, *t*.** Datapoints for animal size's ($N_w$) versus time (***t***), for ***C elegans***, chickens, mice, turkeys, quail, geese, frogs, and humans, and their fit to the numerically integrated form of the ***Binary Cellular Universal Growth Equation*** (#2)**.**

The ***Binary Cellular Analysis*** of ***Growth*** and ***Size***: Measuring the ***Mitotic Fraction***

Returning to Equation #104 above, we expect that the value of the ***Mitotic Fraction*** will decline as the body gets bigger, an expectation that is fulfilled with actual growth data. Indeed, as we shall also see below, growth data will show that the decline in the ***Mitotic Fraction***, ***m***, that occurs as we increase in ***Size, N***, can be captured by the same equation, the ***Binary Cellular Universal Mitotic Fraction Equation***, in a great range of animals, including nematodes, frogs, chickens, cows, geese, quail, turkeys, mice, rats, fish, mollusks, and humans (**Supplement-Figures-I.2-3**).

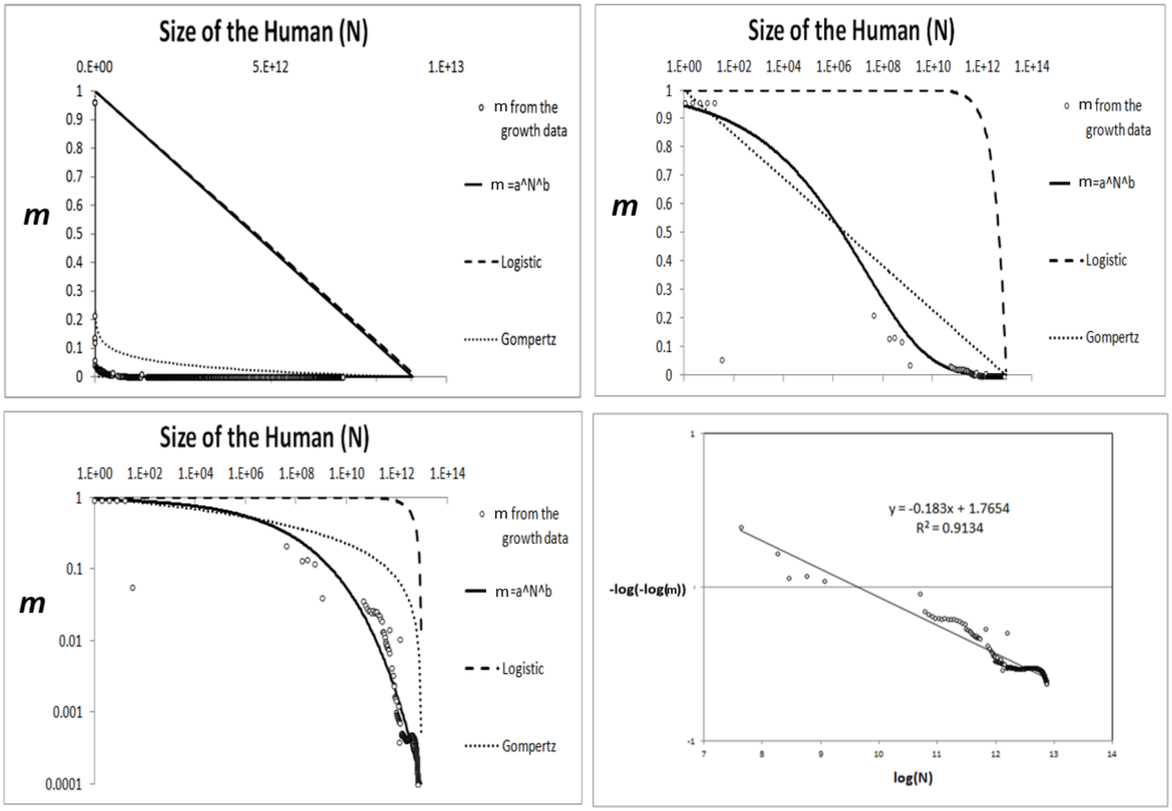


**Supplement-Figure-I.2: Decline in fraction of cells dividing, the *Mitotic Fraction*, *m*, calculated by the *Mitotic Fraction Method*, that occurs as animals increase in size, as seen by the *Binary Cellular Universal Mitotic Fraction Equation*.** Data points for size, $N_w$, in integer units of numbers of cells, vs. the ***Mitotic Fraction***, ***m***, from fertilization, until maturity, for humans.

Let us now take a look at how we make these calculations. Many IVF studies have found that human embryos starting from a single fertilized ovum on day-1, and typically become ~2 cells on day-2, ~4 cells on day-3, ~8 cells on day-4, and so on. These numbers suggest that the ***Cell Cycle Time***, ***c***, of a human is about **1** day, and that right after conception, the ***Mitotic Fraction***, ***m*≈1**. Thus, at the beginning of life, almost all cells are dividing, doubling in number, ***N***, once a day. Nine months later, we have a lovely 8-pound baby girl, and one thing we know for sure is that the next day, she doesn't weigh 16 pounds! If our 8-pound newborn baby girl became 8.8 pounds on the day after birth, we could calculate the ***Mitotic Fraction*** as being 0.1 (***m***≈0.1), since 10% of the child's cells would have to divide for her to grow from 8.0 pounds to 8.8 pounds. If our newborn baby girl weighed 8.08 pounds the day after birth, ***m***=0.01. If our newborn baby girl weighed 8.008 pounds the day after birth, ***m***=0.001, and so on.

Basic Features Of Growth And Mitosis:***Exponential Growth*, *Biotic Potential*, *Mitotic Fraction***, and ***Quiescence***

Because cells increase in number by cell division, they are capable of ***Exponential Growth***, in which the size of an organism, $N_w$ (in terms of cell number), with age, ***t***, can be thought of in terms of its ***Rate of Growth***:

$$\frac{dN_w}{dt} = \frac{ln(2)}{c} \cdot N_w \tag{110}$$

where ***c*** is the average ***Cell Cycle Time*** among those cells that are dividing, which, for ***Exponential Growth***, is all cells.

Integration of Equation #100, translates the integer nature of cells into the continuous appearance of growth:

$$N_w = e^{\left(\frac{ln(2)}{c}\right) \cdot t} \tag{111}$$

Thus, data on the relationship between cell number, $N_w$, and time, ***t***, at the very beginning of development, when all cells are dividing, can give one a practical measure of the animal's ***Cell Cycle Time, c***.

Of course, ***Exponential Growth*** (Equation #100b) is ultimately unsustainable, a fact captured by the concept of the ***biotic potential***.[95] The departure from the ***biotic potential*** of our cells is a fundamental feature of animal growth, which can be captured by introducing the term ***m***, the ***Mitotic Fraction***, giving us this expression for the ***Rate of Growth***:

$$\frac{dN_w}{dt} = \frac{ln(2)}{c} \cdot N_w \cdot m(N_w) \tag{112}$$

Equation #112 embraces all forms of growth, but without specificity, as the manner by which the ***Mitotic Fraction***, $m(N_w)$, changes with the animal size, $m(N_w)$, is completely unspecified. As we shall see below, this lack of specificity gives the Equation #112 the power to accomplish such things as leading to a way to calculate how the ***Mitotic Fraction***, ***m***, changes as we grow (see the ***Mitotic Fraction Method*** below). Additionally, insertion of the ***Mitotic Fraction***, $m(N_w)$, into the ***Exponential Growth*** equation (#111), converts it into Equation #112, and gives us the general form of a ***density dependent growth*** equation, which balances the unlimited quality of ***Exponential Growth*** ($\frac{ln(2)}{c} \cdot N_w$) with the inevitable limit of resources ($m(N_w)$).

Although our treatment of the ***Mitotic Fraction***, ***m***, relies on an obsessively rigorous adherence to this mathematical definition in Equation #112, from time to time we will revert, for the purposes of clarity, to speaking of ***m*** as a rough measure of the ***Growth Fraction***, ***G***, the fraction of cells dividing, which is usually measured by the cytogenetic analysis of the uptake of DNA precursors.[96] The two abstractions of ***Mitotic Fraction***, ***m***, and ***Growth Fraction***, ***G***, are obviously closely, but not precisely, related, since other factors (cell death, cell size, cell density) play roles in growth that are likely to be secondary to cell division. Definitionally, the ***Mitotic Fraction*** is a measure of the amount of actual growth that occurs as a fraction of the growth that would have occurred had each cell divided, ignoring other effects, such as cell death and cell size, a plausible approximation, as mitosis is growth's principal driving force. Thus, the ***Mitotic Fraction m*** can be thought of as the main component of the ***growth fraction***, ***G***, the precise fraction of cells that are dividing. Operationally, this imprecision has little practical impact, as it will be cancelled out when we reintroduce ***m*** into new equations of growth that we shall derive below.

When we think imprecisely of the ***Mitotic Fraction***, ***m***, as the fraction of cells dividing:

$$m \approx N_{Wd}/(N_{Wd} + N_{Wq}) \tag{113}$$

where $N_{Wd}$ is the number of cells dividing and $N_{Wq}$ is the number of cells that have ceased from dividing, a process known as ***Mitotic Quiescence***. ***Quiescence*** is the complex biochemical mechanism that occurs within cells to prevent them from progressing to mitosis, which is set in motion by inhibitory and stimulators growth factor molecules, hormones, and other signals that cells use to communicate.[97-99]

The ***Mitotic Fraction Method***: Measuring the ***Mitotic Fraction***, ***m***, from Fertilization Until Maturity.

The value the ***Mitotic Fraction***, ***m***, for animals of various sizes, $N_w$, can be calculated by reversing ***Equation*** #100c:

$$m = \frac{dN_w}{dt} \Big/ \left[\frac{ln(2)}{c} \cdot N_w\right] \tag{3b}$$

We call the practical implementation of this equation the "***Mitotic Fraction Method***". To execute these ***Mitotic Fraction Method*** calculations, we use our assembled growth data to calculate the value of the ***Mitotic Fraction***, ***m***, (column M of the excel files listed in "**Calculations Files**" column of Table 1) by:

$$m_i = \frac{2N'(t_i)}{r(N_i+N_{i+1})} \tag{120}$$

The same iteration was also carried out based on an exponential calculation, rather than on the linear calculation show above, yielding essentially the same outcome (not shown).

Equation #101 occasionally resulted in a small number of data points with $m > 1$. Since this is both theoretically and numerically impossible (the biological meaning would be one cell dividing to give rise to more than two cells, which is nonsensical), these values of ***m*** were manually overwritten with a value of $m = .999$ noted by orange highlights in the excel spreadsheets.

As Animals Grow in Size, $N_w$, the ***Mitotic Fraction***, ***m***, Declines Rapidly.

Our ***Mitotic Fraction Method*** calculations, based on the animal growth data noted above (Table 1), characterized how the value of the ***Mitotic Fraction***, ***m***, declines as growth occurs, that is, as $N_w$ increases, from fertilization until maturity, for nematodes, chickens, cows, geese, quail, turkeys, fish, clams, mice, rats, humans, and many other animals (**Supplement-Figures-I.2-3**). Graphs of the values of the ***Mitotic Fraction***, ***m***, calculated by the ***Mitotic Fraction Method***, for each of the diverse species of animals noted above, reveal that growth is close to exponential during the first few cell divisions ($m \approx 1$), but soon begins its "slippery slope" descent, resulting in a rapid and accelerating decline in the value of ***m***, reaching values of $10^{-2}$ to $10^{-6}$ by the time adult size is reached (**Supplement-Figures-I.2-3**).

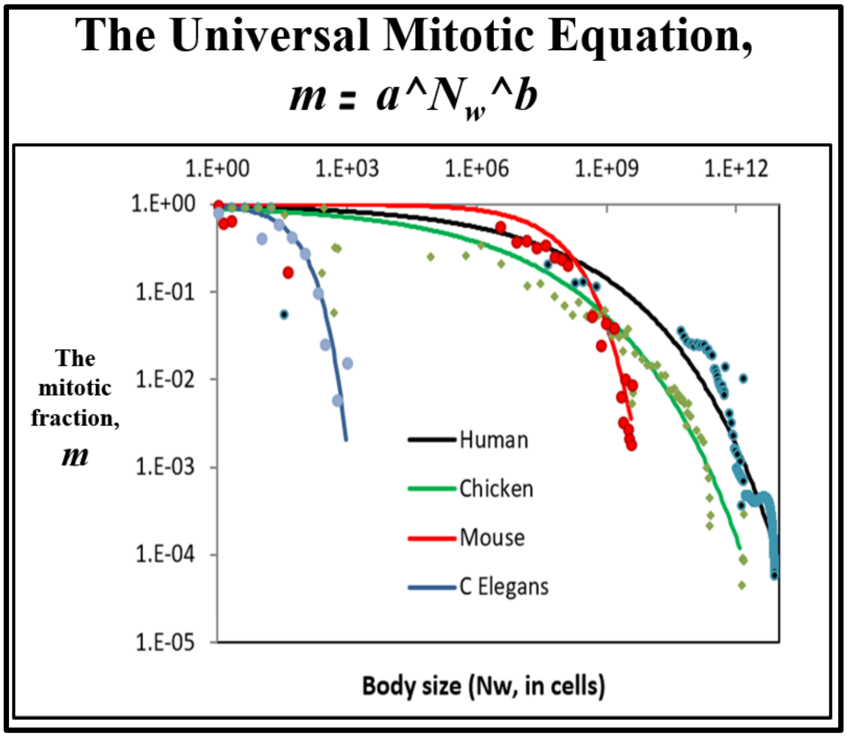


**Supplement-Figure-I.3: Values of the *Mitotic Fraction*, *m*, calculated by the *Mitotic Fraction Method*, as a function of animal size, $N_w$, as shown at various scales.** Data points for animal size, $N_w$, in integer units of numbers of cells, vs. the ***Mitotic Fraction***, ***m***, from fertilization, until maturity, for humans, chickens, ***C elegans*** nematodes, and mice.

The ***Binary Cellular Biology*** of ***Growth*** and ***Size***: The ***Binary Cellular Universal Mitotic Fraction Equation***

The basis for the relationship between the ***Mitotic Fraction***, $\boldsymbol{m}$, and $\boldsymbol{N_w}$ becomes evident by graphing the $\boldsymbol{\log(N_w)}$ vs $-\boldsymbol{\log(-\log(m))}$, revealing roughly straight rows of dots (**Supplement-Figure-I.2**). This means (see below for the math) that the relationship between the ***Mitotic Fraction***, $\boldsymbol{m}$, and $\boldsymbol{N_w}$ can be captured with the expression:

$$\boldsymbol{m = a^{(N_w^b)}} \tag{1}$$

We call this expression the ***Binary Cellular Universal Mitotic Fraction Equation*** (#1), as it accurately captures the value of the ***Mitotic Fraction, m***, the fraction of cells that must be dividing to account for the growth of the body, from fertilization until maturity, for each of the 13 species noted above, including humans (TABLE A1). The parameters $\boldsymbol{a}$ and $\boldsymbol{b}$ reflect the specific growth characteristics of each animal.

The best fit values for $\boldsymbol{a}$ and $\boldsymbol{b}$ for each species were calculated as regression inputs of $\boldsymbol{\log N_i}$ and $\boldsymbol{-\ln(-\ln g_i)}$ using Excel's LINEST function. Values for $\boldsymbol{a}$ and $\boldsymbol{b}$ were calculated from the LINEST output (in columns P-Q of the excel files listed in Table 1) together with the $r^2$ value. This relationship is well borne out, as indicated by the high $r^2$ values from non-linear regressions for each of the eleven animals listed in the TABLE A2

Because Equation #1 so closely captures the relationship between the ***Mitotic Fraction***, $\boldsymbol{m}$, and size, $\boldsymbol{N_w}$, from fertilization until maturity, for all of the animals we have examined, we call this expression the "***Binary Cellular Universal Mitotic Fraction Equation***". ("Universal"; from the Latin "universalis"; Etymology: "uni (Latin, "one"), Versum (Latin, "turned"); thus something "turned into one"). Note that the ***Binary Cellular Universal Mitotic Fraction Equation***, and the ***Binary Cellular Universal Growth Equation***, that is derived from it, and which we shall describe below, holds for all of the animals that we have examined, and for all of the sizes we have examined, from the first fertilized cell onward.

Of course, the practical execution of these ***Mitotic Fraction*** calculations can get a bit complicated. However, details, and values for the ***Mitotic Fraction m***, as a function of ***Body Size, N***, for all of the species listed above, from ***conception*** ($\boldsymbol{N_w=1}$ when $\boldsymbol{t=0}$), to adulthood ($\boldsymbol{N_w \approx 10^{12}}$ for humans), can be found in Reference 100.

The ***Binary Cellular Biology*** of ***Growth*** and ***Size***: The ***Mitotic Fraction*** is a Measure of ***Cellular Selection***

***Cellular Selection***, resulting from ***Differential Cellular Proliferation***,[97-99] molds the progeny of single fertilized ova into multicellular animals.[100,105,114] Calculating the ***Mitotic Fraction, m***, from growth data, allows us to make measurements of this fundamental force of embryonic development.

The ***Mitotic Fraction Log*(*Log*)/*Log Test***

As we noted above, when we graphed the ***Log***(-***Log***) of the ***Mitotic Fraction*** against the ***Body Size*** $N_w$, from ***conception*** ($N_w$=**1**) to adulthood ($N_w \approx 10^{12}$ for humans), we got straight-lines (**Supplement-Figure-I.4**), and that this indicated the existence of the ***Binary Cellular Universal Mitotic Fraction Equation*** (#1). To see why this is the case, consider this arithmetic:

$$m = a\text{^}(N\text{^}b) \tag{90}$$

$$-Log(m) = -Log\ (a\text{^}(N\text{^}b)) \tag{91}$$

$$-Log(m) = (N\text{^}b) \ * \ (-Log\ (a) \tag{92}$$

$$Log\ (-Log(m)) = Log\ (\ (N\text{^}b) \ * \ (-Log\ (a)\ ) \tag{93}$$

$$Log\ (-Log(m)) = Log(\ (N\text{^}b)\ ) + Log\ (-Log\ (a)\ ) \tag{94}$$

$$Log\ (-Log(m)) = b * Log(\ (N) + Log\ (-Log\ (a)\ ) \tag{95}$$

$$Log\ (-Log(m)) = b * Log(\ (N) + Z \tag{96}$$

where

$$Z = Log\ (-Log\ (a) \tag{97}$$

Since the calculations we carried out used ***Log*** to the base 10: (98)

$$-Log(a) = 10\text{^}Z \tag{99}$$

$$a = 10\text{^}(-10\text{^}Z) \tag{999}$$

Thus the ***Mitotic Fraction Log*(*Log*)/*Log Line*** has a slope that reflects the ***b*** parameter of the ***Binary Cellular Mitotic Fraction Equation*** (#4), and a height that reflects the ***"a" Parameter***. It follows that carrying out a linear regression of comparisons of ***Mitotic Fraction***, ***m***, data against ***fetal size***, ***N***, data, on such a ***Log*(*Log*)/*Log*** graph, provides a test for whether the relationship conforms with the ***Binary Cellular Mitotic Fraction Equation*** (#4). Such a test, revealing this to be the case for human growth data can be seen in **Supplement-Figure-I.2**. In the third paper in this series, we shall again that this ***Mitotic Fraction Log*(*Log*)/*Log Test*** is useful, not only in identifying the biologically deep, and ancient, ways that different animals grow to different sizes, at different rates, but also, in tracking the growth of human fetuses that become ***Small, Average***, and ***Big Babies*** to the single ***"a" Parameter***, of the ***Binary Cellular Mitotic Fraction Equation*** (**Supplement-Figure-III.23**).

The ***Binary Cellular Biology*** of ***Growth*** and ***Size***: The ***Binary Cellular Universal Growth Equation***,

Taking into account one more parameter, ***c***, the ***Cell Cycle Time***, the average length of time that it takes a cell to divide, allowed us to derive this ***Density Dependent*** expression for the how fast animals grow:

$$\boldsymbol{Rate\ of\ Growth} = \frac{\boldsymbol{ln(2)}}{\boldsymbol{c}} \cdot \boldsymbol{N_w} \cdot \boldsymbol{a}^{(\boldsymbol{N_w^b})} \tag{2}$$

Integration (se below) yields the relationship between ***age***, ***t***, and ***size***, ***N***:

$$\boldsymbol{t(N)} = \frac{\boldsymbol{c}}{\boldsymbol{b \cdot log(2)}} \cdot \{\mathrm{Ei}[-\boldsymbol{N^b} \cdot \boldsymbol{log(a)}] - \mathrm{Ei}[-\boldsymbol{log(a)}]\} \tag{2b}$$

and the relationship between ***size***, ***N***, and ***age***, ***t***:

$$N_w = \int ln2/c * N_w * a\hat{\ }N_w\hat{\ }b \tag{2c}$$

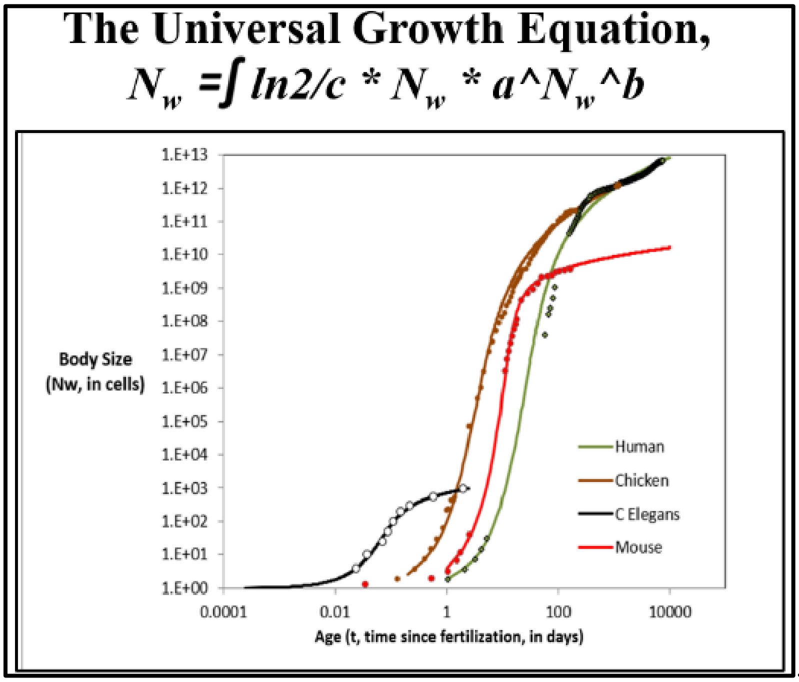


**Supplement-Figure-I.4** The ***Binary Cellular Universal Growth Equation***.
Actaul Size/Age datapoints for humans, chickens, nematodes, and mice shown in dots; the ***Binary Cellular Universal Growth Equation***, with the appropriate ***a, b***, and ***c*** parameters for each of these species (TABLE I), shown in curves.

We have called these variations different forms of the ***Binary Cellular Universal Growth Equation*** (#2), as we found it to accurately capture growth, from fertilization until maturity, for each 13 species listed above, again noting that "Universal" is from the Latin "universalis", "uni" being "one", with "Versum" being "turned", thus something "turned into one" (TABLE A1).[100]

The values of the ***a***, ***b*** and ***c*** parameters of the ***Binary Cellular Universal Growth Equation*** (#2) capture the specific growth characteristics of each animal (Details can be found in Reference 100). For humans, ***a*≈0.943**, ***b*≈0.169** and ***c*≈1.016** (in units of days). The resulting growth curve for humans, chickens, nematodes, and mice, can be seen in **Supplement-Figure-I.4**, **TABLE A1**. As we shall see below, the ***Binary Cellular Universal Growth Equation*** (#2) provides us with a basis from which we can build practical mathematical expressions for deciphering and managing ***Human Fetal Growth***.

While the ***Binary Cellular Universal Growth Equation*** (#2) for capturing animal and human growth over time (**Supplement-Figure-I.4**) is expressed in units of number of cells, $N_w$, conversion to grams, ***g***, is easily accomplished, as there are approximately $10^8$ cells in every gram of tissue (see below).[101] When cell number values, $N_w$, are derived from weight data, conversion back to grams cancels out any error in the estimation of cells per gram. Furthermore, even if we convert numbers of cells, ***N***, into grams, ***g***, to derive practical expressions for comprehending growth, these expressions still possess a well-hidden, but still available, biologically rooted mathematics for comprehending the biochemical mechanisms of growth.

**TABLE A1**

Parameters and goodness-of-fit metrics to the

***Universal Growth*** and ***Universal Mitotic Fraction Equations***. From Ref 100

| Organism | Genus species | *Fit of the data to the Rate of Growth* *m=N*a^(N^b)* (*c=Cell Cycle Time*, in days) | | | | | *Fit of the data to growth* *N*=∫[log(2)/*c*]**N*a^(N^b)* |
|---|---|---|---|---|---|---|---|
| | | *c* | *a* | *b* | $r^2$ (linear fit) | $r^2$ (log fit) | $r^2$ (log fit) |
| Humans (♂) | *Homo sapiens* | 1.016 | 0.943 | 0.169 | 89.9% | 95.0% | 99.5% |
| Frogs | *Rana pipiens* | 0.070 | 0.925 | 0.269 | 57.0% | 82.3% | 97.3% |
| Nematodes | *C. elegans* | 0.011 | 0.946 | 0.680 | 66.7% | 91.6% | 99.8% |
| Chickens | *Gallus gallus* | 0.127 | 0.906 | 0.166 | 80.8% | 97.8% | 99.8% |
| Cows | *Bos taurus* | 0.921 | 0.882 | 0.128 | 76.8% | 95.1% | 100.0% |
| Geese | *Anser anser* | 0.048 | 0.898 | 0.170 | 77.0% | 92.9% | 97.1% |
| Mice | *Mus musculus* | 0.508 | 0.990 | 0.281 | 65.0% | 96.2% | 99.7% |
| Quail | *Colinus virgianus* | 0.292 | 0.795 | 0.126 | 76.5% | 95.3% | 94.2% |
| Rats | *Rattus norvegicus* | 0.417 | 0.950 | 0.197 | 66.5% | 90.6% | 99.5% |
| Clams | *M. mercenaria* | 0.033 | 0.962 | 0.275 | 79.0% | 91.0% | 99.5% |
| Turkeys | *Meleagris gallopavo* | 0.028 | 0.757 | 0.122 | 65.3% | 92.2% | 99.7% |

Zebrafish (*Danio rerio*) and European sea bass (*Dicentrarchus labrax*) also show these features of growth (data not shown)

**TABLE A2**

**FILES WITH GROWTH DATA, FOR DISTRIBUTION TO INTERESTED READERS**

| Species | Genus Species | Basic Data Files | Calculations Files |
|---|---|---|---|
| Human Males | *Homo sapiens* | Boys Basic Growth Data jsm 2 12 2013.xlsx | 140330 Human males.xlsx |
| Frogs | *Rana pipiens* | Rana Basic Growth Data jsm 2 15b 2013.xlsx | 140330 Rana Pipiens.xlsx |
| Nematodes | *C. elegans* | BASIC FORM Celegans SingleSheet jsm 2 13 13 LogMethod.xlsx | 140426 C elegans.xlsx |
| Chickens | *Gallus gallus* | CHikcens Aggrey and Byerley and cleavage Data 2 14 2013.xlsx | 140426 Chickens.xlsx |
| Cows | *Bos taurus* | COW Basic Growth Data jsm 1 27 2013.xlsx | 140426 Cows.xlsx |
| Geese | *Anser anser* | BASIC FORM Geese SingleSheet jsm 2 14 2013 Log Method.xlsx | 140426 Geese.xlsx |
| Mice | *Mus musculus* | BASIC FORM MicePoiley M AKR jsm 1 28 2013 Log&LinearMethods.xlsx | 140426 Mice.xlsx |
| Quail | *Colinus virgianus* | BASIC FORM Quail SingleSheet jsm 2 16 2013.xlsx | 140426 Quail.xlsx |
| Rats | *Rattus norvegicus* | RAT Basic Growth Data jsm 2 14 2013.xlsx | 140426 Rats.xlsx |
| Turkey | *Meleagris gallopavo* | BASIC FORM TURKEYwithOvaposition SingleSheet jsm 2 8 2013.xlsx | 140426 Turkey.xlsx |
| Clams | *Merceneria mercenaria* | BASIC FORM LOG METHOD mercenaria 1 11 2013 v2 from Boys SingleSheet jsm 2 12 2013 | 140705 Mercenaria.xlsx |
| LOCATION: | | C:_0\My Papers\Growth_DRAFTS__final Excell Files_Basic Data 7 7 14 | |

The Relationship Between Size, $N_w$, and Age, $t$, is Well Fit by the ***Binary Cellular Universal Growth Equation***:

Combining Equation #3 and #4 leads to this expression for describing the rate of animal growth:

$$\frac{dN_w}{dt} = \frac{ln(2)}{c} \cdot N_w \cdot a^{(N_w^b)} \tag{5a}$$

The relationship between size, $N_w$, and age, $t$, is captured by numerical integration of Equation #5, where the age, $t$, at fertilization is ~0, and the age at death is $d$:

$$N_w = \int_{t=0}^{t=d} \frac{ln(2)}{c} \cdot N_w \cdot a^{(N_w^b)} \tag{5b}$$

Numerical integration of Equation #5, to derive Equation #5b, was carried out 4th order Runge-Kutta method, employed manually in Excel to carry out such growth curve reconstructions, so as to examine relationship between Size ($N_w$) and Age ($t$). The step size $h$ (cell b11 of the excel files listed in "**Calculations Files**" column of the excel files enumerated in Table 1) was set such that 10,000 steps was used for each organism. At each $t_i$ (column W), the model value of $N_i$ was calculated recursively in column X as

$$N_i = N_{i-1} + \frac{h(k_1+2k_2+2k_3+k_4)}{6} \tag{200}$$

where:

$$\text{if: } f(N) = N' = rNg \tag{201}$$

$$k_1 = f(N_i) \tag{202}$$

$$k_2 = f\left(N_{i-1} + \frac{k_1}{2}h\right) \tag{203}$$

$$k_3 = f\left(N_{i-1} + \frac{k_2}{2}h\right) \tag{204}$$

$$k_4 = f(N_{i-1} + k_3 h) \tag{205}$$

The values of each $k_i$ are calculated in columns Z-AC of the excel files listed in "**Calculations Files**" column of the table above. Excel's VLOOKUP and CORREL functions were then used to calculate correlations of observed and model **ln** $g$ and **ln** $N$ (columns AM-AU). These correlations, as well as the regression correlation coefficient, were used as quality-of-fit metrics for the model and recorded in the summary columns A-B in the excel files.

The fit of Equation #5b, the numerically integrated form of Equation #5, to the various datapoints for animal size's ($N_w$) versus time ($t$), are shown in **Supplement-Figure-I.3**. The strength of this relationship was confirmed by the high $r^2$ values for each of the eleven animals listed in the Table 2: humans, frogs, nematodes, chickens, cows, geese, mice, quail, rats, turkeys, fish, and clams.

Closed Form Solution to The Integration of the ***Binary Cellular Universal Growth Equation***

A closed form solution to the integration of the ***Binary Cellular Universal Growth Equation*** (#2), can also be seen by considering:

$$\frac{dN}{dt} = r\,N\,a^{(N^b)} \tag{301}$$

where $r = \frac{\log(2)}{c}$. Rearranging, one obtains:

$$N^{-1}\,a^{(-N^b)}\,dN = r\,dt \tag{302}$$

which can be integrated as:

$$\int_1^{N_w} N^{-1}\,a^{(-N^b)}\,dN = \int_0^t r\,d\tau \tag{303}$$

Leading to the growth equation:

$$t = \left[\mathrm{Ei}\left(-N_w^b \log(a)\right) - \mathrm{Ei}(-\log(a))\right]\frac{c}{b\,\log(2)} \tag{3}$$

where $t = 0$ when $N_w = 0$, and **Ei** is the exponential integral function[102].

Equation #3 can be considered as a continuous approximation of the expected time required by a Poisson model with individual cell growth $ra^{(N^b)}$.

We show this by considering the random variable $t_N$ representing the time required to reach the number of cells *N*, starting from a single cell. This variable has expected value:

$$E[t_N] = \sum_{n=2}^{N} E[t_n - t_{n-1}] \tag{305}$$

We assume that each one of the *n* cells that make up the organism at a given time behaves according to a Poisson process with rate $ra^{(n^b)}$ and that they are all independent. As a result, the event represented by the growth of a new cell out of any of the *n* current cells behaves according to a Poisson process with rate $n\,r\,a^{(n^b)}$ (this can be easily seen by considering the cumulative distribution function of the exponential distribution). Therefore:

$$\overline{t_N} = E[t_N] = \sum_{n=2}^{N} \frac{1}{nra^{(n^b)}} \tag{306}$$

$$\sigma_N = \sqrt{\mathrm{var}(t_N)} = \sum_{n=2}^{N} \frac{1}{\left(nra^{(n^b)}\right)^2} \tag{307}$$

where $\overline{t_N}$ and $\sigma_N$ are, respectively, the mean and standard deviation of the time required to reach the number of cells *N*. The continuous version in #6 and the discrete (point-process) version in #306 provide very similar results. For example, using $c = 0.070$, $a = 0.925$, $b = 0.269$, we get the behavior shown in **Supplement-Figure-I.**1 for the continuous version in red, and the discrete version in blue (mean ± std) (**Supplement-Figure-I.**7).

Density Dependent Growth Equations

A number of ***density-dependent*** growth equations have been developed to capture either the growth of the number of organisms in a population,[103,104] or the growth of the size of individual organisms,[9-13] none of which accurately fit the growth of real animals.[28-30] The simplest of these ***density-dependent*** growth equations, the logistic, occurs by $\boldsymbol{m}$ declining linearly as the size of the organism, $\boldsymbol{N_w}$, increases.[9] In the ***Gompertz equation***[10], $\boldsymbol{m}$ declines as the log of $\boldsymbol{N_w}$. Similarly, for many other equations, including the ***von Bertalanffy***[11], the ***Richards***[12], and ***West***[13] equations, the value of $\boldsymbol{m}$ declines monotonically as size, $\boldsymbol{N_w}$, increases.

The relationship of the ***Mitotic Fraction*** (***m***) and size ($\boldsymbol{N_w}$) to the ***logistic equation***.

***Density dependent*** growth is captured by the ***logistic equation***[9] in the form:

$$\boldsymbol{m = 1 - \frac{N_w}{K}} \tag{401}$$

The relationship of the ***Mitotic Fraction*** (***m***) and size ($\boldsymbol{N_w}$) to the ***Gompertz equation***.

***Density dependent*** growth is captured by the ***Gompertz equation***,[10] in the form:

$$\boldsymbol{m = \left(\frac{1}{\log K}\right) \log\left(\frac{N_w}{K}\right)} \tag{402}$$

The relationship of the ***Mitotic Fraction*** (***m***) and size ($\boldsymbol{N_w}$) to the ***West equation***.

***Density dependent*** growth is captured by the ***West equation***,[13] for total mass of the organism (***M***), in the form:

$$\boldsymbol{\frac{dM}{dt} = aM^{\frac{3}{4}}\left(1 - \left(\frac{M}{K}\right)^{\frac{3}{4}}\right)} \tag{403}$$

Units of mass are unspecified, so we can treat Equation #403 such that the units of mass (***M***) of our organism are in units of "the weight of a cell" ($\boldsymbol{N_w}$). West's equation then becomes:

$$\boldsymbol{\frac{dN_w}{dt} = aN^{\frac{3}{4}}\left(1 - \left(\frac{N_w}{k}\right)^{\frac{3}{4}}\right)} \tag{404}$$

By re-arranging:

$$\boldsymbol{\frac{dN_w}{dt} = \left[\left(\frac{\log 2}{c}\right) N\right]\left[\left(\frac{ac}{\log 2}\right) N^{-\frac{3}{4}}\right]\left[1 - \left(\frac{N_w}{k}\right)^{\frac{3}{4}}\right]} \tag{405}$$

Thus, for any organism growing by the ***West equation***, the fraction of cells dividing, ***m***:

$$m = \left[\left(\frac{ac}{\log 2}\right) N_W^{-\frac{3}{4}}\right]\left[1 - \left(\frac{N_W}{k}\right)^{\frac{3}{4}}\right] \tag{406}$$

Equation #406 can be thought of as the ***West equation***, in terms of the fraction of cells dividing, ***m***.

Fit of the Relationship of the ***Mitotic Fraction*** (***m***) and Size ($N_w$), to the ***Gompertz***, ***Logistic***, and ***West Equations***.

As can be seen in **Supplement-Figure-I.2**, over the full range from conception to adult size, neither the Gompertz (Equation #402), nor the logistic (Equation #401) provides a close characterization of the behavior of the ***Mitotic Fraction***, ***m***, with respect to $N_w$. As can be seen in **Supplement-Figure-I.5**, while the ***West equation*** can be used to quite accurately capture growth over its full expanse, to do so requires $m > 1$ early in development, which is mathematically valid, but biologically impossible, since individual cells would have to give rise to more than two daughter cells in a single ***Cell Cycle Time***. However, both our ***Binary Cellular Universal Growth Equation*** (Equation #2), and the ***West equation*** (#406), work quite well (**Supplement-Figure-I.**5) later in development. This means that our ***Binary Cellular Universal Growth Equation*** is capable of achieving growth within the limits of the blood supply's capacity to support growth, as captured by the ***West equation***.[13]

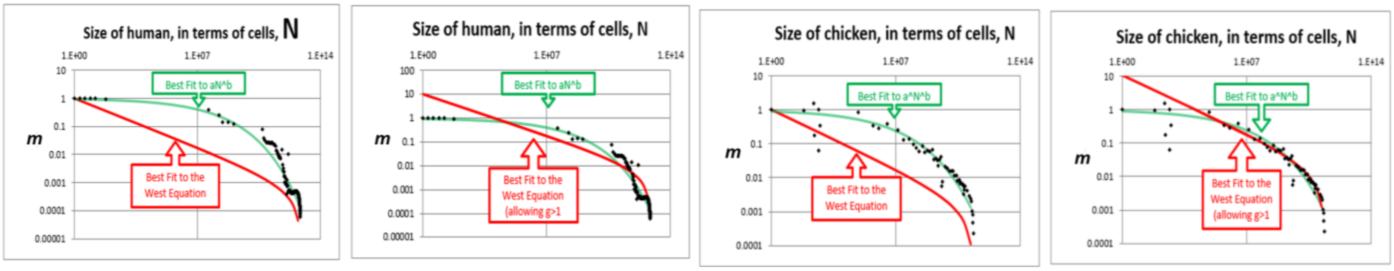


**Supplement-Figure-I.5**: **Comparison of the fit of *Mitotic Fraction*, *m*, as a function of size, $N_w$, with human growth data, to the *Binary Cellular Universal Mitotic Fraction Equation* (green) and *West equation* (red).**

Fit of the Relationship Between Size ($N_w$) and Age (***t***) to the ***Gompertz***, ***logistic***, and ***West equations***.

We tested our data on body size, $N_w$, and age, ***t***, against three of the most widely used density dependent growth equations (the ***Gompertz***, the ***logistic***, and the ***West*** [a variation of the ***von Bertalanffy***]) (**SupplementFigure-I.**6). None of these growth equations fit the actual growth data as well as the ***Binary Cellular Universal Growth Equation*** (**Supplement-Figure-I.**6).

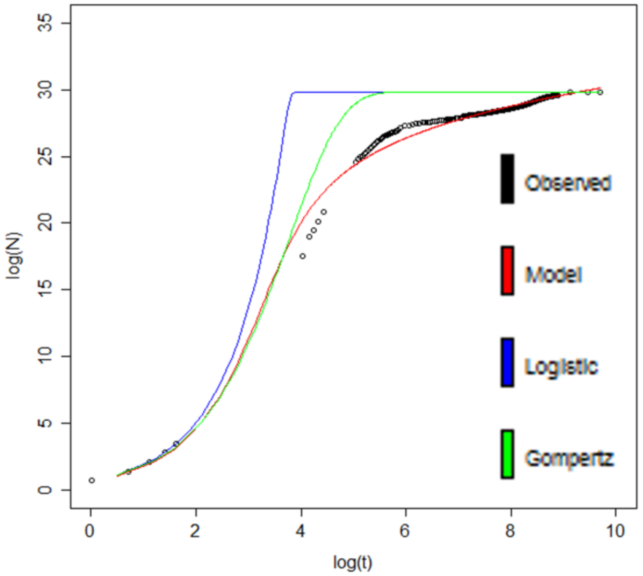


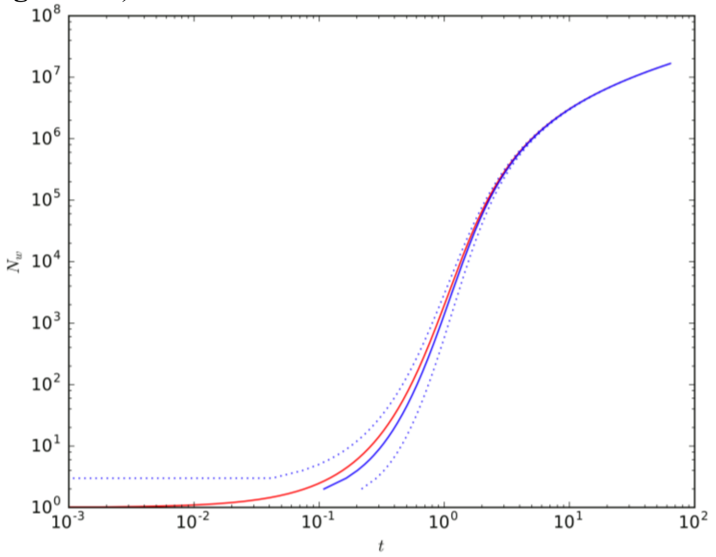


**Supplement-Figure-I.6**: **Fit of human growth to the *Binary Cellular Universal Growth Equation* (red), the *logistic equation* (blue), and the *Gompertz equation* (green).** Data values as black circles.

**Supplement-Figure-I.7:**
**Analysis of the closed form integration of the *Binary Cellular Universal Growth Equation* (#3).**

The ***Mitotic Fraction***, ***m***, is Linked to Signaling Molecules.

What might be the mechanism behind the ***Mitotic Fraction***, ***m***, the fraction of cells that divide, which declines as we increase in size? What could cause a fraction of the body's cells to enter into a state of ***mitotic quiescence***, the internal block to cell division, which is set in motion by external signaling molecules?[97-99] Curiously, as we shall see below, this decline in the ***Mitotic Fraction***, by the ***Binary Cellular Universal Mitotic Fraction Equation*** (#1), $\boldsymbol{m = a^{(N_w^b)}}$, might well trace its origin to the deceptively simple and unobvious discrete allocation of ligand molecule among cells[105]. To picture how this might occur, consider the case of an embryo growing in a constant volume, such as a bird's egg or mammal's uterus, whose cells produce an inhibitory growth factor and carry a receptor for that inhibitory molecule. Early in development, when our idealized embryo is just a few cells, those few cells would produce only a small number of inhibitory molecules, and thus the concentration of these inhibitory molecules would be low, and thus very few cells would have bound the number of inhibitory molecules needed to prevent cell division. Thus, at the beginning of development, most cells would divide, with the value of the ***Mitotic Fraction***, ***m***, being close to **1**. However, as the embryo grows, more and more cells are present to produce inhibitory molecules, the concentration of these inhibitory molecules would increase, more and more cells would have bound the number of inhibitory molecules needed to prevent cell division, fewer and fewer cells would be able to divide, and the value of the ***Mitotic Fraction***, ***m***, would decline. Remarkably, as we shall see next, when we rephrase this idealized case in mathematical terms, such a decline in the ***Mitotic Fraction***, ***m***, can be seen to occur in exactly the form of the ***Binary Cellular Universal Mitotic Fraction Equation*** (#1), $\boldsymbol{m = a^{(N_w^b)}}$ .

Let us consider growth factor molecules in terms of integers.[105] Thus, when growth factor molecules bind to cells, they shall display a discrete allocation among cells, that is, as a Poisson process, which can be comprehended in terms of a Poisson probability (Equation #611 below).

Let **Ic** be the number of inhibitory growth factor molecule made by each cell which makes the molecule. This number is a constant, being the result of the rates of synthesis and decay, most likely being first order reactions.

Let ***N*** be the number of cells in the organism. We shall consider growth from conception, that is, from when ***N*****=1**.

Let $\boldsymbol{N^b}$ be the number of cells producing the inhibitory growth factor molecules. If every cell produces the molecules, $\boldsymbol{b = 1}$. If only the cells on the surface produce the molecule, $\boldsymbol{b = \frac{2}{3}}$, since this captures the relationship between the surface of an isomorphic 3-dimensional shape, such as a sphere, and its volume.

Let ***V*** be the volume in which the embryo develops. For a bird ***V*** is the volume of the egg; for a mammal it is the volume of the uterus.

The total number of molecules of **I** in the body is $\boldsymbol{Ic \cdot N^b}$.

**[I]** is the concentration of inhibitory growth factor molecule in the body, assuming uniform diffusion.

Let us now put all this into mathematical form.

It follows that:

$$[\mathbf{I}] = \frac{Ic \cdot N^b}{V} \tag{601}$$

$$[\mathbf{I}] = \left(Ic \cdot N^b\right)\left(\frac{1}{V}\right) \tag{602}$$

Let $\mathbf{P_i}$ be the probability of a receptor molecule binding an inhibitory growth factor molecule, where **[R]** is the concentration of receptor molecules.

$$\mathbf{P_i} = \frac{[\mathbf{IR}]}{[\mathbf{R}]} \tag{603}$$

$$\frac{[\mathbf{I}][\mathbf{R}]}{[\mathbf{IR}]} = \mathbf{k} \tag{604}$$

according to the Law of Mass Action, thus

$$\frac{[\mathbf{IR}]}{[\mathbf{R}]} = \frac{[\mathbf{I}]}{\mathbf{k}} \tag{605}$$

$$\mathbf{P_i} = [\mathbf{I}] \cdot \frac{\mathbf{1}}{\mathbf{k}} \tag{606}$$

$$\mathbf{P_i} = \left(\mathbf{Ic} \cdot \mathbf{N^b}\right)\left(\frac{\mathbf{1}}{\mathbf{V}}\right)\left(\frac{\mathbf{1}}{\mathbf{k}}\right) \tag{607}$$

$$\mathbf{P_i} = \mathbf{N^b}(\mathbf{Ic})\left(\frac{\mathbf{1}}{\mathbf{V}}\right)\left(\frac{\mathbf{1}}{\mathbf{k}}\right) \tag{608}$$

$$\mathbf{P_i} = \mathbf{N^b}\left(\frac{\mathbf{Ic}}{\mathbf{V \cdot k}}\right) \tag{609}$$

$$\mathbf{P_i} = \left(\frac{\mathbf{Ic}}{\mathbf{V \cdot k}}\right)\mathbf{N^b} \tag{610}$$

Let **Ro** be the number of receptors per cell.

We shall use the Poisson probability[106] to calculate the value of ***m***:

$$\mathbf{P(x; \mu)} = \frac{(\mathbf{e^{-\mu}})(\mathbf{\mu^x})}{\mathbf{x!}} \tag{611}$$

where **x** is the actual number of successes that result from the experiment.

**e**: A constant equal to approximately 2.71828

**μ**: The mean number of successes that occur in a specified region.

**x**: The actual number of successes that occur in a specified region.

$\mathbf{P(x; \mu)}$: The Poisson probability that exactly **x** successes occur in a Poisson experiment, when the mean number of successes is **μ**.

It follows that

$$\boldsymbol{m} = \mathbf{P(x; u)} = \frac{(\mathbf{e^{-\mu}})(\mathbf{\mu^x})}{\mathbf{x!}} \tag{612}$$

It follows that $\boldsymbol{x} = \mathbf{0}$ (since we want to calculate the chance of a cell binding **0** molecules of inhibitory growth factor).

And where $\mathbf{\mu} = \mathbf{Ro} \cdot \mathbf{P_i}$ (since the mean number of successes for each cell is the number of receptors, time the chance that a molecule will bind one of those receptors):

$$\boldsymbol{m} = \mathbf{P(x; \mu)} = \frac{(\mathbf{e^{-\mu}})(\mathbf{\mu^0})}{\mathbf{0!}} \tag{613}$$

$$\boldsymbol{m} = \mathbf{P(x; \mu)} = \frac{(\mathbf{e^{-\mu}})(\mathbf{1})}{\mathbf{1}} \tag{614}$$

$$\boldsymbol{m} = \mathbf{P(x; \mu)} = \mathbf{e^{-\mu}} \tag{615}$$

$$\boldsymbol{m} = \mathbf{P(x; \mu)} = \mathbf{e^{-Ro \cdot P_i}} \tag{616}$$

or

$$\boldsymbol{m} = \mathbf{e^{-RoP_i}} \tag{617}$$

It follows that

$$\boldsymbol{m} = \mathbf{e}^{-\mathbf{Ro}\left(\frac{\mathbf{Ic}}{\mathbf{V \cdot k}}\right)(\mathbf{N^b})} \tag{618}$$

Let $\boldsymbol{Z} = \boldsymbol{Ro}\left(\frac{\boldsymbol{Ic}}{\boldsymbol{V}\cdot\mathbf{k}}\right)$

$$\boldsymbol{m} = \mathbf{e^{-Z(N^b)}} \tag{619}$$

Let $\boldsymbol{a} = \boldsymbol{e^{-Z}}$

$$\boldsymbol{m} = \boldsymbol{a}^{(\mathbf{N^b})} \tag{4}$$

Which is the ***Binary Cellular Universal Mitotic Fraction Equation***!

This leads to our ***Binary Cellular Universal Growth Equation***, when combined with Equation #5a:

$$\frac{\boldsymbol{dN}}{\boldsymbol{dt}} = \frac{\mathbf{ln(2)}}{\boldsymbol{c}} \cdot \boldsymbol{N} \cdot \boldsymbol{a}^{(\boldsymbol{N^b})} \tag{5a}$$

where

$$\boldsymbol{a} = \mathbf{ln}\left(\boldsymbol{Ro}\left(\frac{\boldsymbol{Ic}}{\boldsymbol{V}\cdot\mathbf{k}}\right)\right) \tag{620}$$

Note that all of the biochemistry (the rate of growth factor synthesis, abundance of receptors, dissociation constants, etc.) is in the ***"a" Parameter***, which we shall see in the third paper in this series, provides the key to predicting the size at which a human fetus is born, while all of the geometry (size of the tissues that making the growth factor in relationship to the size of the organism) is in the parameter ***b*** (see "Let $\boldsymbol{N^b}$ be …" above).

#### The ***Binary Cellular Biology*** of Human ***Growth***

Let us now take a closer look at human growth.

##### Terminology for ***Fetal Growth***: ***Age***, ***Gestational Age***, ***Conceptional Age***, ***Fetal Age***

***Gestational Age***, the time since the last menstrual period, has long been the most common way for thinking about the age of the human fetus. Another commonly used term is ***Conceptional Age***, but the date of conception is not always known. As we shall be examining fetal, childhood and adult growth data, as well as growth data of animals such as mollusks, tunicates, and nematodes, which don't have a fetal period, we shall be using the simplest term, ***Age***, with modifiers to indicate the period examined, which, for our work here, will be ***Fetal Age***.

##### Terminology for ***Fetal Growth***: ***Cells***, ***Grams***, ***Mitotic Fraction***, ***Weight-Gain Rate***, ***Growth Velocity***

As we have noted above, while ***Binary Cellular Analysis,*** and specifically, the ***Binary Cellular Universal Growth Equation*** (#2), captures human growth over time, in units of number of cells, ***N*** (**Supplement-Figure-I.10**). Conversion to grams, ***g***, is also easily accomplished, as there are approximately $10^8$ cells in every gram of tissue.[82] Such a conversion to grams, ***g***, still retains an easily accessible, if perhaps temporarily hidden, biologically rooted mathematics for comprehending the biochemical mechanisms of growth, as we shall see in all three papers in this series**.** Further, when cell number values, ***N***, are derived from weight data, conversion back to grams, ***g***, cancels out any error in the estimation of cells per gram.

***Human Fetal Size*** and ***Growth*** can thus be examined in units of ***Cells***, ***Mitotic Fraction***, ***Grams***, or ***Weight-Gain Rate*** in units of ***Cells***/***Day*** or ***Grams***/***Day***, allowing comparison to other commonly used standards for assessing fetal developments, while retaining the analytical power of ***Binary Cellular Analysis***. We shall see this especially in the third paper in this series.

The term ***Fetal Growth Velocity*** is sometime used, but we shall not be using it for several reasons. ***First***, although it has a long history in the study of growth, it is, in fact, a misnomer, as velocity technically is used to describe motion with a direction (a vector), such as a satellite, car, or bullet, while the growth of a fetus has no such directionality. ***Second***, in practice, there is no standardized definition of ***Fetal Growth Velocity***, and it has been used in various ways. ***Fetal Growth Velocity*** is sometime used to describe grams/time, based on ***Estimated Fetal Weight***,[107,108] and sometimes in millimeters/time, for ultrasound measurements.[109] Partarp Sovio and Smith capture ***Fetal Growth Velocity*** by measuring the change in the Z score of various ultrasound measurements between 28 and 37 weeks of gestational age.[110]

Of course, fetal growth is not linear, so ***Weight-Gain Rate***, in units of ***Cells***/***Day*** or ***Grams***/***Day***, even for the same fetus at various time points, will vary as to when the measurements are made. However, as Mongelli and colleagues[111] have noted, fetal growth is relatively constant at the end of pregnancy, a quality they call "quasi-linearity". As well shall see below, and examine further in the third paper in this series,[7] we also see this linearity, which we refer to as the ***~Linear Phase of Human Fetal Growth***. This linearity allows a first approximation of growth to be assessed in terms that we shall refer to as the ***Weight-Gain Rate***, as we shall see below, and in the third paper in this series.

Data for ***Human Fetal Growth***

A variety of sources provide us with information on human ***Fetal Growth***. For early in development, a number of studies have counted the number of cells from conception onward, for patients that have utilized ***In Vitro Fertilization*** (***IVF***).[81] For later in development, the quantitative information on the growth of fetuses can be seen from ***Fetal Ultrasound*** [7] and ***Fetal Autopsy*** data.[116,117] Values from two ***Fetal Autopsy*** studies[116,117] are shown in **Supplement-Figure-I.8** below. Values from ***Fetal Ultrasound*** data will be examined in the third paper in this series.[7]

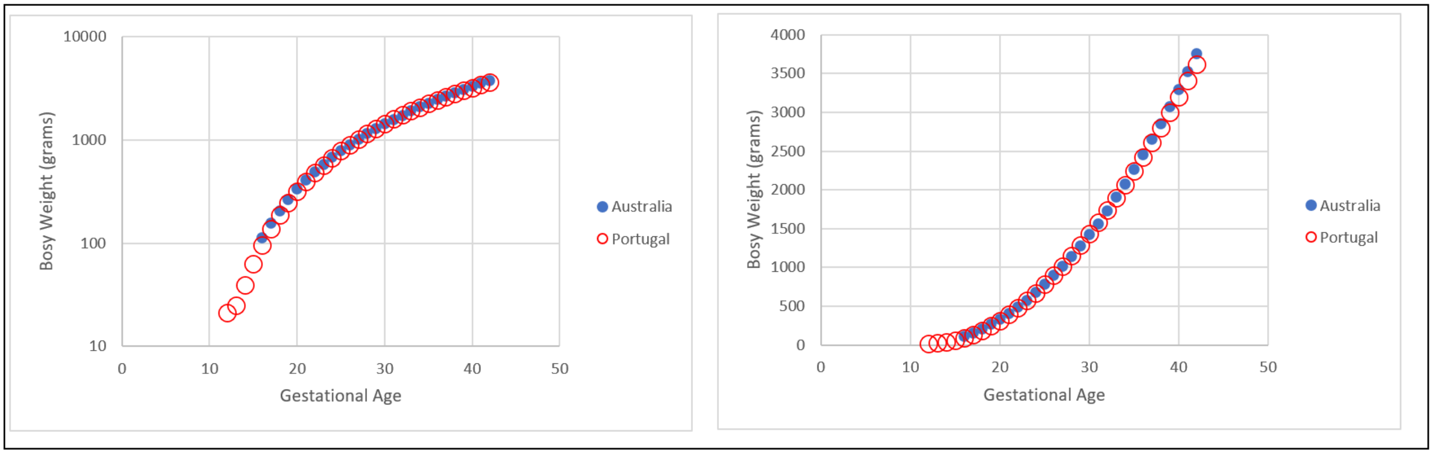


**Supplement-Figure-I.8**. Human fetal growth, in grams, from two fetal autopsy studies.[116,117]

In **Supplement-Figure-I.9** below, we can see growth, in units of number of cells, at the beginning of life, from the 1st cell onward, as has been measured by counting cells as embryos grow following ***IVF*** in the first week of life,[81] as well as from 3 months to 9 months, as has been measured by fetal autopsy.[116,117] Cell numbers for the autopsy data were estimated by $10^8$ cells/gram.[101] Values are shown at various scales.

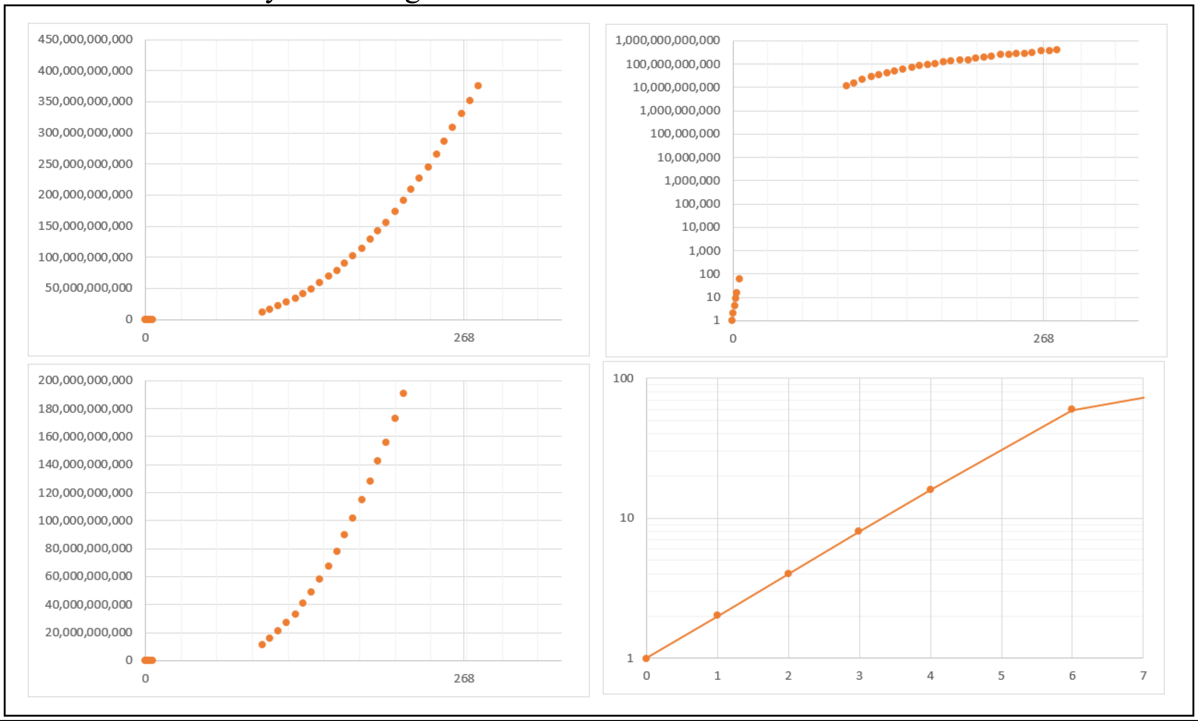


**Supplement-Figure-I.9** Human growth, in units of numbers of cells, ***N***, from fertilization until 280 days of age, where births occur, on average, at 268 days. Data in the first week of life from studies of embryos created by IVF,[81] while data later in pregnancy determined form fetal autopsy data,[116,117] in grams, converted to cell numbers,[116,117] based on the approximations of $10^8$ cells/gram[82]

***Excel Numerical Integration*** of the ***Binary Cellular Universal Growth Equation***

The curves in **Supplement-Figures-I.1** and **4** above for determining the ***Size*** of animals, $N_w$, over ***time***, ***t***, from this ***Binary Cellular Universal Growth Equation*** (#2), were created by the Runge–Kutta method. A simpler way to execute such integrations, which we shall use here, breaks down growth into chunks of ***time***, $\Delta t_{Cw}$, in units of ***Cell Cycle Times***, $c_w$. This can be done, in excel, in row, after row, and thus in ***Cell Cycle Time***, $c_w$, after ***Cell Cycle Time***, $c_w$, calculating, in each step, the increase in number of cells in each row, $\Delta t_{Cw}$, with the ***Binary Cellular Universal Mitotic Fraction Equation*** (#1). We call this the ***Excel Numerical Integration of the Binary Cellular Universal Growth Equation*** (#2). In **Supplement-Figure-I.10** below are shown, at various scales, such excel-created integrations of the ***Binary Cellular Universal Growth Equation***, for human growth, from conception ($t$=**0**) to the first months after birth ($t$=**330** days), with the parameters noted above, $a$=**0.943**, $b$=**0.169** and $c$=**0.9893**.

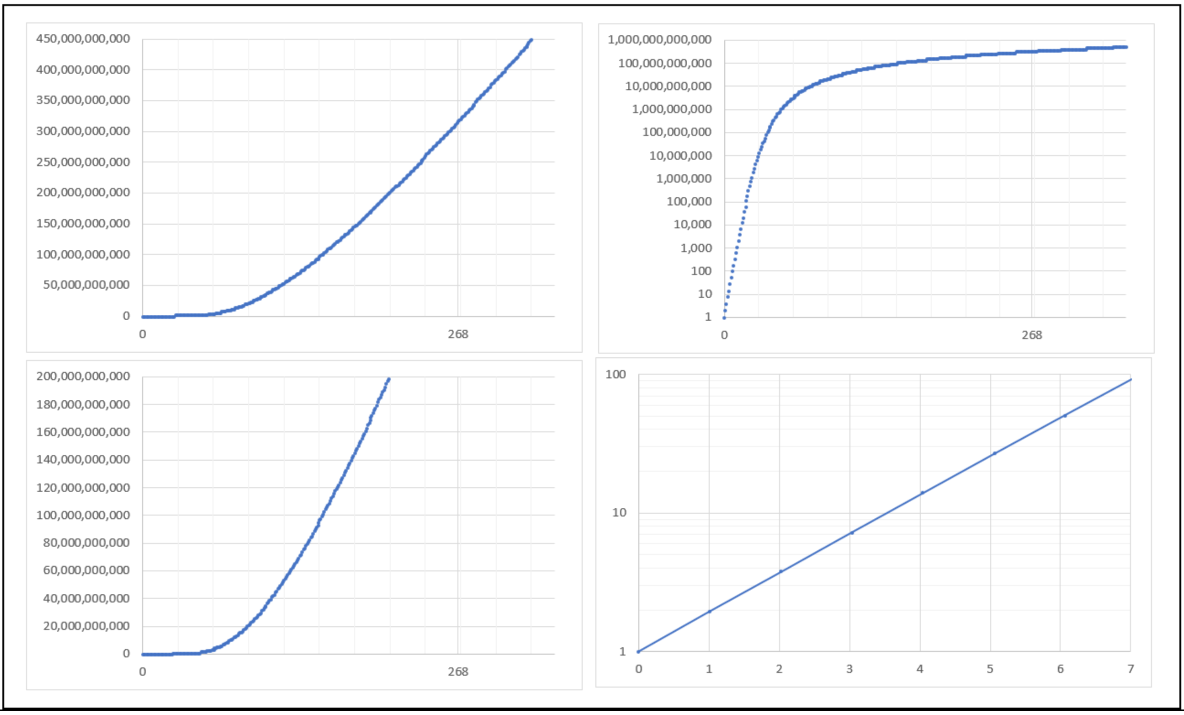


<u>**Supplement-Figure-I.10**</u>. ***Excel Numerical Integration*** of the ***Binary Cellular Universal Growth Equation*** for human growth, in units of numbers of cells, ***N***, from conception ($t$ =**0**) to shortly after birth ($t$=**330** days), where births occur, on average, at 268 days, with the parameters noted above, $a$=**0.943**, $b$=**0.169** and $c$= **0.9893**, calculated by excel.

**_Excel Numerical Integration_** and Data Comparison: the **_Binary Cellular Universal Growth Equation_**

We can now compare ***Fetal Size***, from ***IVF*** and ***Fetal Autopsy*** data (**Supplement-Figure-I.9**), with the ***Excel Numerical Integration*** of the ***Binary Cellular Universal Growth Equation*** (**Supplement-Figure-I.10**). This reveals that the ***Binary Cellular Universal Growth Equation*** (#2) does a remarkably good job at capturing growth from fertilization onward. Indeed, this equation covers a range, for humans, from 1 cell to a trillion+ cells (~0.00000001 grams to ~100,000+grams) with an accuracy of ~1-in-a-100-billion (**Supplement-Figure-I.11**).

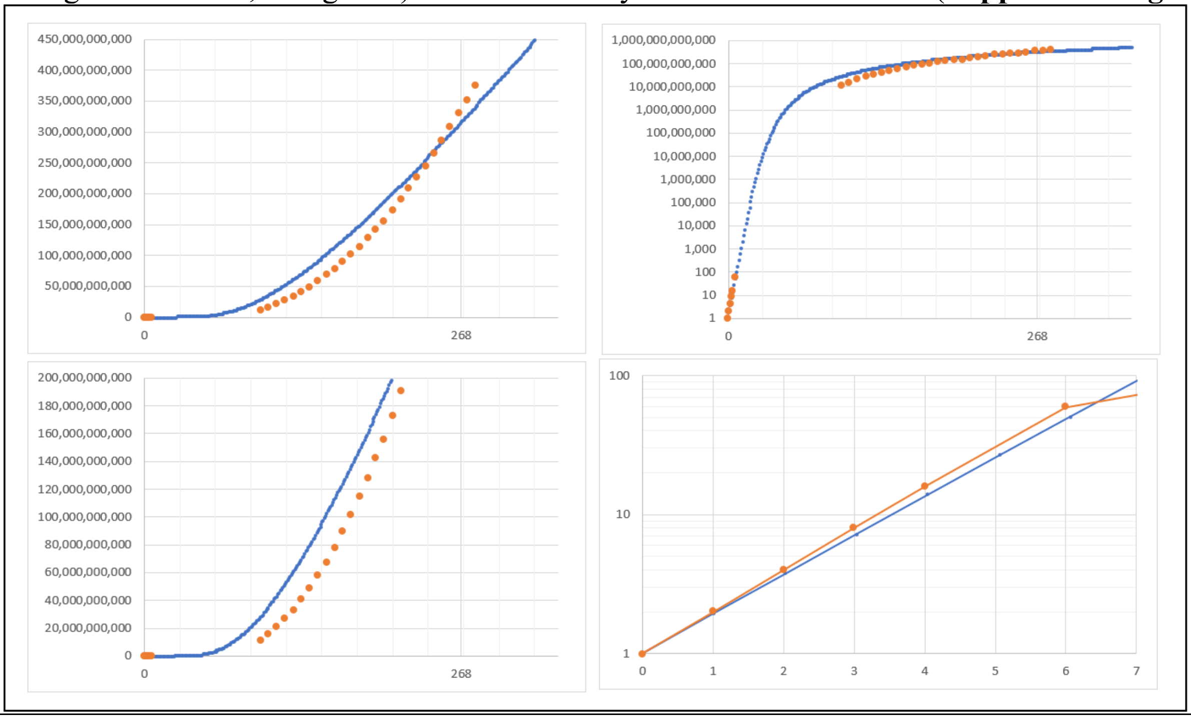


**<u>Supplement-Figure-I.11</u>**. Comparison of human growth, calculated by the ***Binary Cellular Universal Growth Equation***, with values determined for actual human ***Size*** measurements, both in units of numbers of cells, ***N***. Calculations of the ***Binary Cellular Universal Growth Equation*** were carried out by ***Excel Numerical Integration***, with the parameters ***a*≈0.943**, ***b*≈0.169** and ***c*≈1.016**. Data in the first week of life from studies of embryos created by IVF,[81] while data later in pregnancy determined form fetal autopsy data,[116,117] in grams, converted to cell numbers, based on the approximations of $10^8$ cells/gram.[82]

***Ripples in Growth***

Like all equations in biology or physics, however good they may be at capturing the world, the world always gives us small tweaks of reality, which deviate from the math that describes us. This has a long tradition in science, such as Newton's Law's of motion, which provides a perfect equation for the earth traveling around the sun (the so-called 2-body problem), while also revealing little bumps when the moon pushes us here and there (the 3-body problem). We call these minor deviations from the ***Binary Cellular Universal Growth Equation***, ***Ripples in Growth***, ***R***, the most pronounced of which occurs at puberty (**Supplement-Figure-I.12**).

A number of biological processes can lie behind these ***Ripples in Growth***. The hormonal effects of puberty on the acceleration of growth are well known and characterized. Another likely source of ***Ripples in Growth*** may well be parts of the body growing faster, or slower, than the body as a whole. As we shall see below, many parts of the body display such differential growth, ascribable to ***Founder Cells*** undergoing ***Cell-Heritable*** changes in their ***Cell Cycle Times***, $c_p$, changes, which persist in their progeny. Arms and legs provide one of the more dramatic cases of such ***Differential Cellular Proliferation***, the most common form of ***Cellular Selection***, which plays such a vital role in forming our bodies.[112,113,114] As we shall see below, the fact that these cells appear to have a shorter ***Cell Cycle Time***, $c_p$, than the cells in the rest of the body, $c_w$, accounts for why limbs grow out of the body. This gives the body as a whole a slightly larger rate of the growth than just the trunk of the body.

At the other end of the spectrum, ***Cell Death*** molds the shape of the body, in such tasks as removing cells between the fingers.[112,114,115] The occurrence of such ***Apoptosis*** should be expected to lead to negative ***Ripples in Growth***. Thus, ***Cell Death*** will lower the number of cells of which we are comprised, making us grow slower than what would be expected from the ***Binary Cellular Universal Growth Equation*** alone.

All of these ***Ripples in Growth***, ***R***, can be captured mathematically by such methods as Fourier transform computation, a task we shall leave for a later day. To extract this second layer of growth information, we derived an expression, Equation #500, to measure the magnitude and timing of these growth residuals, $\boldsymbol{\delta_j}$:

$$m = A^{N^B} + \sum \delta_j \qquad (500)$$

where each $\delta_j$ models a wave-like deviation away from the trend (the ***Binary Cellular Universal Growth Equation*** #2) that corresponds to an interpretable life-event in the growth of the organism. The appropriate functional forms for the $\delta_j$ may be determined through regression of the residuals of observed ***m*** against the ***Binary Cellular Universal Growth Equation*** (#2).

Indeed, a graph of these $\delta_j$ values, as a function of the size of the developing human male, $N_w$, which is shown in **Supplement-Figure-I.12** bears out our suspicion that these ***ripples*** identify biologically relevant processes, such as the right-most peak (at $\log N_w \approx 29.3$), which corresponds to puberty.

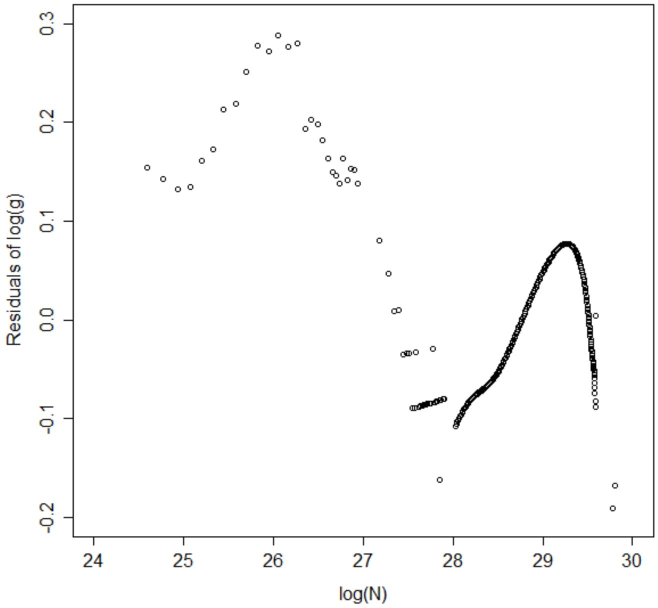


**<u>Supplement-Figure-I. 12</u>: Wave-like deviations ("*Ripples*") from the *Binary Cellular Universal Growth Equation curve*, for human growth.**

***Ripples in Human Fetal Growth***

For the purpose of refining human fetal growth, here we shall see that ***Empirical*** approximations of these ***Ripples*** can provide perfectly practical refinements of human fetal growth curves, optimizing ***Binary Cellular Growth Mathematics*** for the tasks at hand, as we shall see below (**Supplement-Figures I.13-15**). ***Ripples in Growth*** can be readily added to the ***Binary Cellular Universal Growth Equation*** when examined by ***Excel Numerical Integration***.

In **Supplement-Figure-I.13**, we can see in two minor ***Ripples in Growth***, the first slightly below the ***Binary Cellular Universal Growth Equation*** before 250 days after conception, and then slightly above the ***Binary Cellular Universal Growth Equation*** after 250 days after conception. These ***Ripples*** could be included with the ***Excel Numerical Integration*** of the ***Binary Cellular Universal Growth Equation*** by adding a small decrease in ***Mitotic Fraction*** (~20%) to the early part of pregnancy, and a small increase in ***Mitotic Fraction*** (~30%) to the later part of pregnancy.

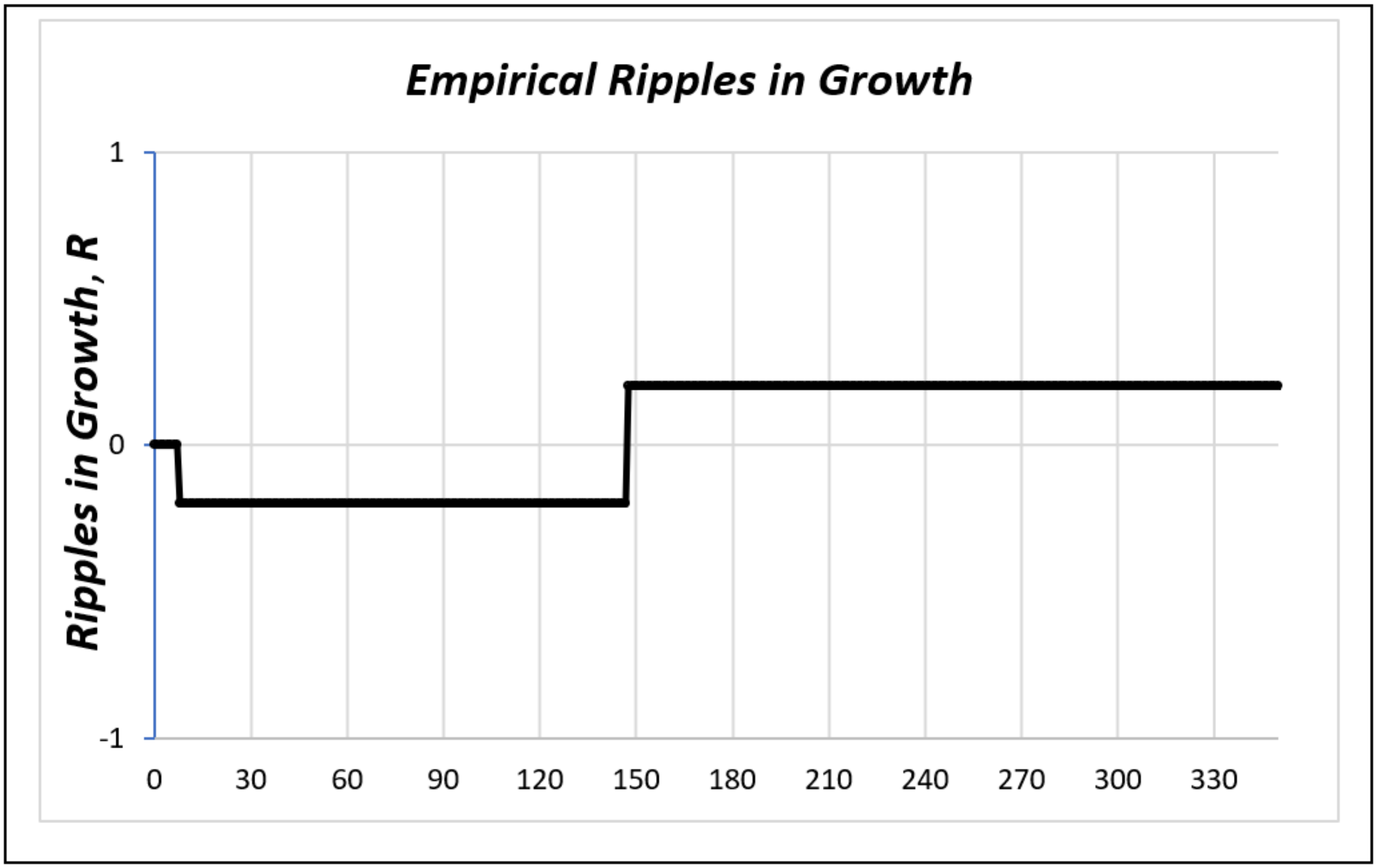


**Supplement-Figure-I.13**. The ***Empirical Ripples in Growth***, used to fit the ***Binary Cellular Universal Growth Equation*** (#2) to the fetal autopsy growth data.[116,117]

The resulting growth curve provides a close fit to the data from both autopsy studies (**Supplement-Figures I.14** and **I.15**; Portugal[116]; Australia[117]). This ***Empirical*** curve-fitting approach doesn't tell us very much about the biology behind these fetal ***Ripples in Growth***. However, it does provide a perfectly good job for accurately capturing fetal growth from conception until birth.

Here, we have utilized a simple additive step function for capturing the ***Empirical Ripples in Growth***, manually adjusted to achieve a close fit to the data. This makes possible a quite satisfactory fit of actual growth data to the {***Binary Cellular Universal Growth Equation*** (#2) plus the fetus's ***Ripples in Growth***}(**Supplement-Figure-I.8a** and **b**) In the third paper in this series, we shall see how this ***Empirical Ripple*** approach can be refined further, with ultrasound data, and a multiplicative rather than additive ***Ripple***. We can also easily envisage a smoother, more rigorous, wave-like set of ***Ripples in Growth***, rather than the clunky step functions shown in **Supplement-Figure-I.8a** and **b**, which could be estimated, and then put to work by such methods as Fourier transform computation, an opportunity we shall save for a later day.

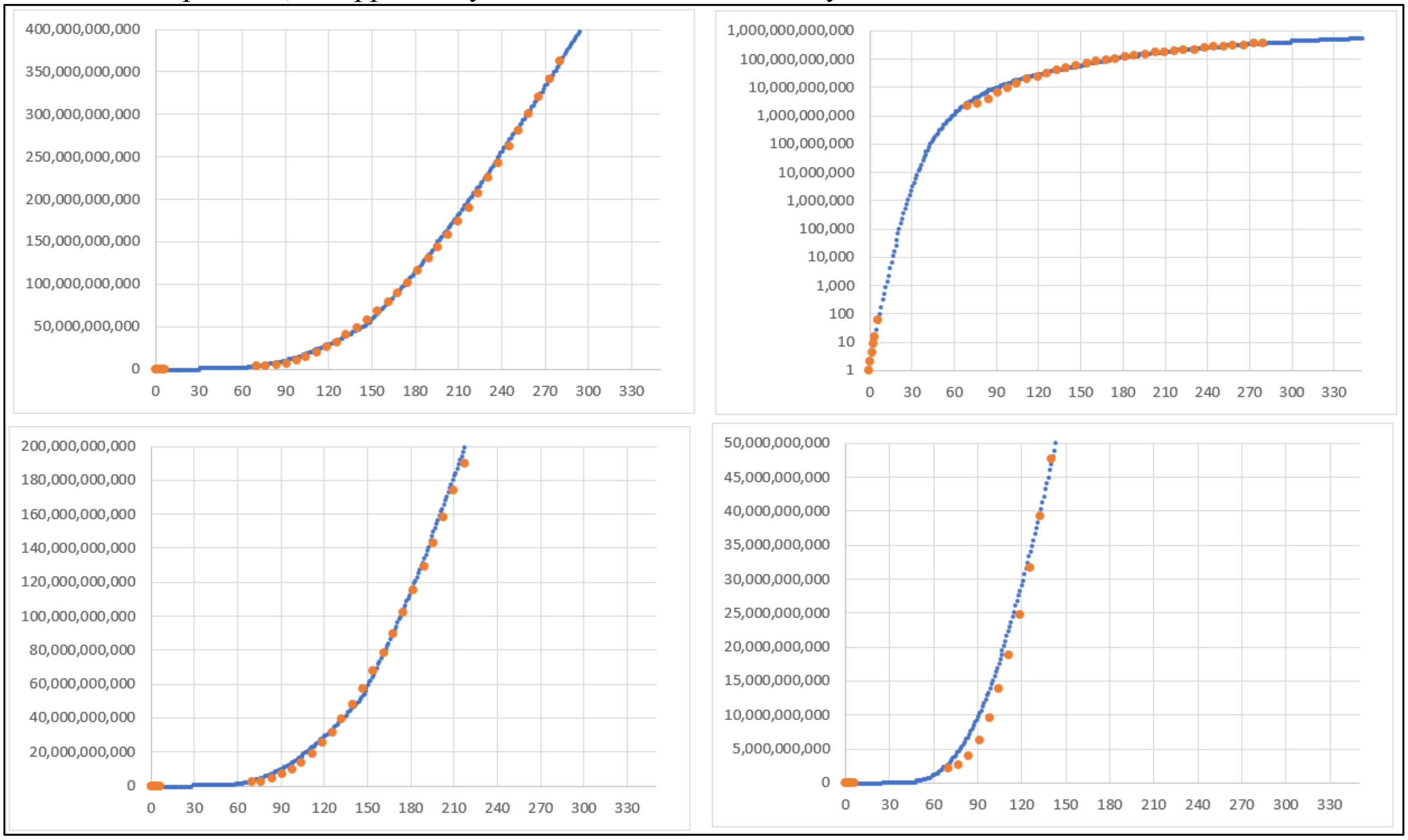


__**Supplement-Figure-I.14**__. Comparison of the human growth curve calculated by the ***Binary Cellular Universal Growth Equation*** with (blue line), with ***Empirical*** curve-fitting described in the text, with values determined for actual human ***Size*** measurements, in units of numbers of cells, ***N*** (orange dots), with data in the first week of life from studies of embryos created by IVF,[81] while data later in pregnancy determined form fetal autopsy data, in grams, converted to cell numbers, based on the approximations of $10^8$ cells/gram, from Portugal.[116]

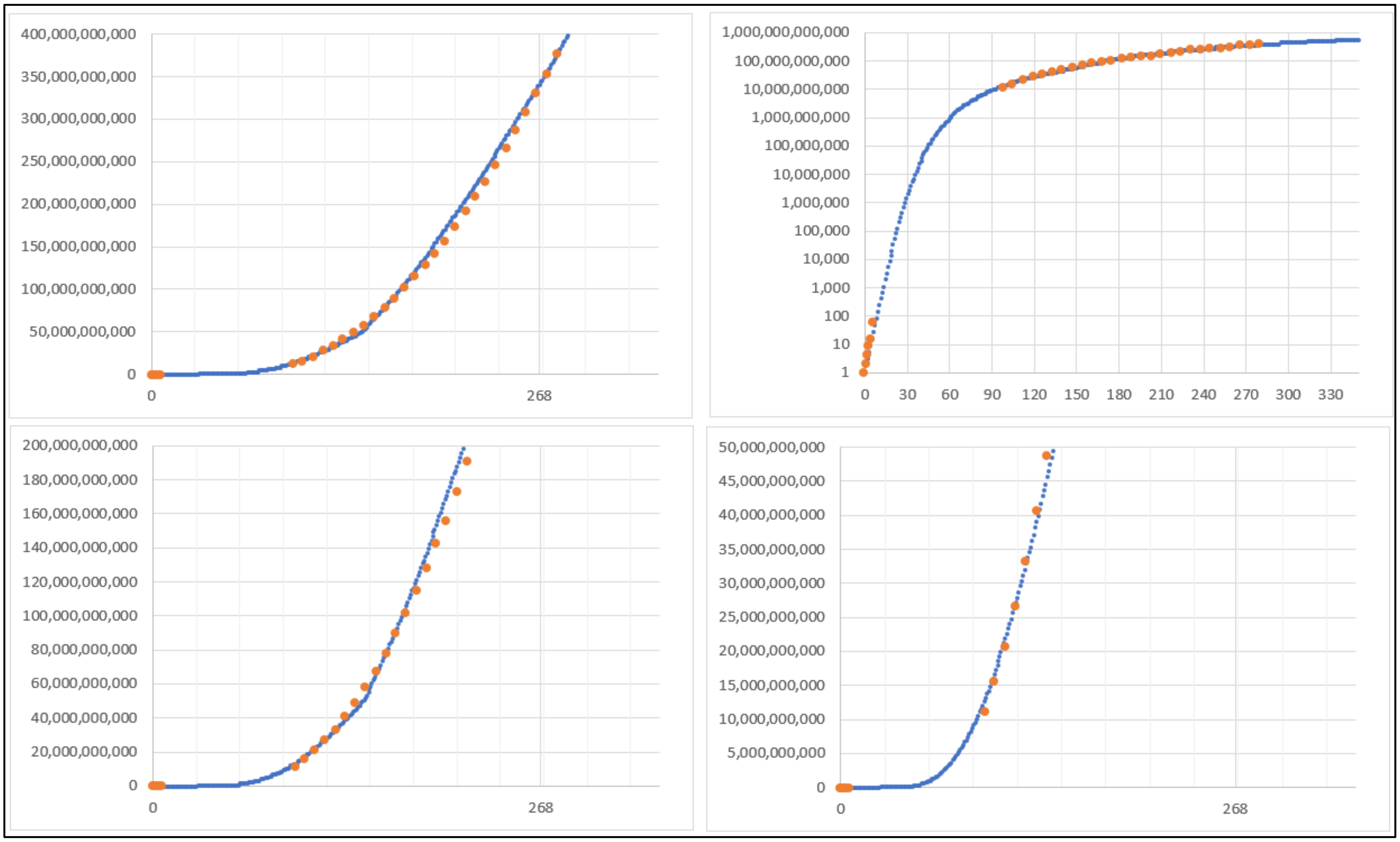


**Supplement-Figure-I.15**. Comparison of the human growth curve calculated by the ***Binary Cellular Universal Growth Equation*** with (blue line), with ***Empirical*** curve-fitting described in the text, with values determined for actual human ***Size*** measurements, in units of numbers of cells, ***N*** (orange dots), with data in the first week of life from studies of embryos created by IVF,[81] while data later in pregnancy determined form fetal autopsy data, in grams, converted to cell numbers, based on the approximations of $10^8$ cells/gram, from Australia.[117]

The ***~Linear Phase of Human Fetal Growth***

The approximation of fetal growth becomes even simpler, and indeed, strikingly informative, when we examine the ***Binary Cellular Universal Growth Equation*** (#2) over the last three months of pregnancy (**Supplement-Figures I.16-18**). From about 7 months until 10 months after ***conception***, when most fetuses have exceeded 1500 grams in weight (~150 billion cells), growth is highly, and conveniently, extraordinarily, if nor precisely, linear, which this why we call it the ***~Linear Phase of Human Fetal Growth***.

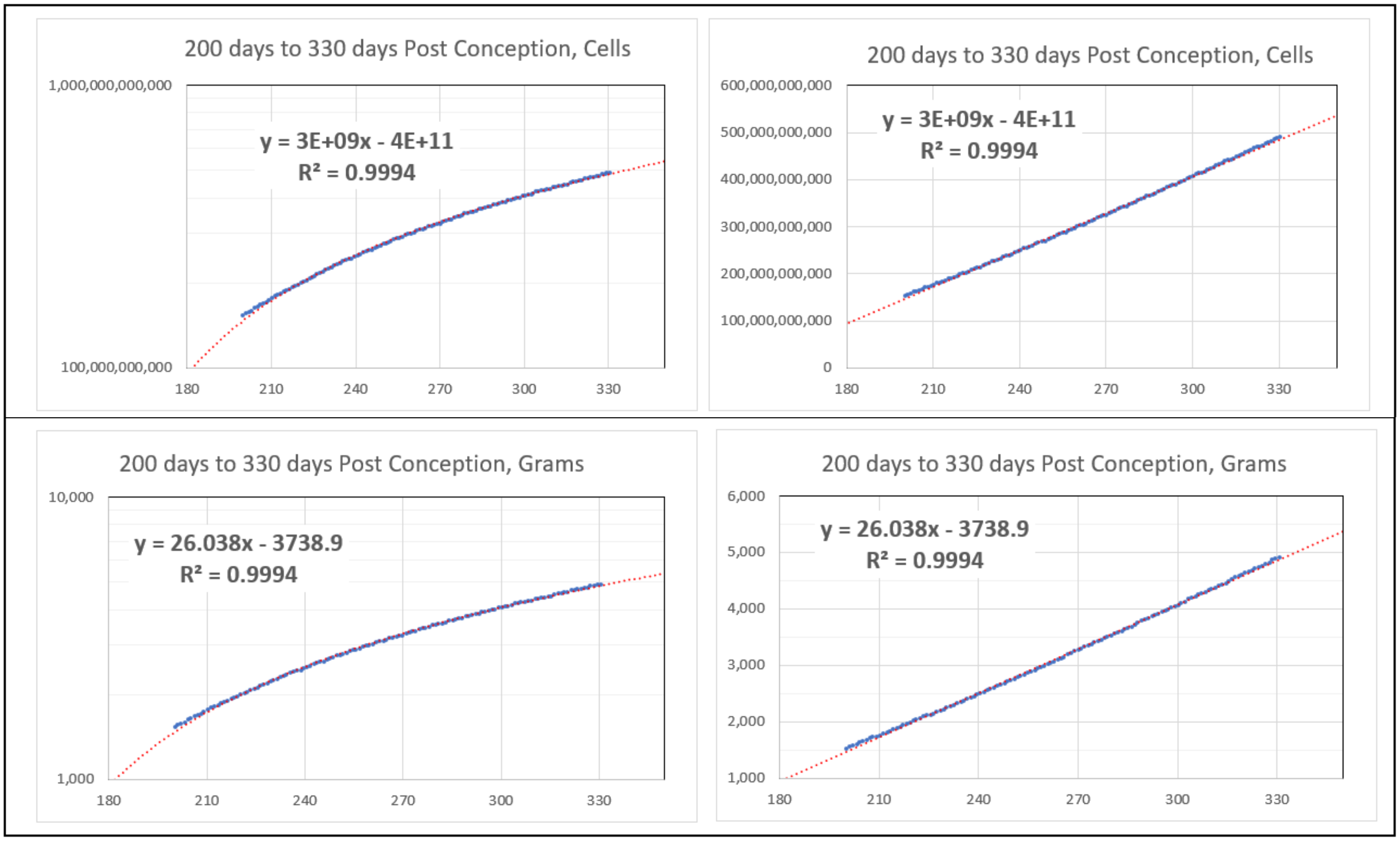


**Supplement-Figure-I.16**. The ***~Linear Phase of Human Fetal Growth*** The human growth curve calculated by the ***Binary Cellular Universal Growth Equation*** with (blue line), with ***Empirical Ripples*** adjustment shown in shown in **Supplement-Figure-I.13** (red line). Shown from 180 days to 360 days post conception, with linear regressions examined from 200 days to 330 days post conception. Births occur, on average, at 268 days.

It follows that, with taking ***Ripple*** refinement into account, the in the last 3 months of pregnancy:

$$\textit{\textbf{Fetal Size}} \text{ (in grams)} \approx 26.038*(\boldsymbol{t}) - 3738.9 \qquad (501)$$

and

$$\textit{\textbf{Fetal Size}} \text{ (in numbers of cells)} \approx (26.038*(\boldsymbol{t}) - 3738.9) * 1e8 \qquad (502)$$

where $\boldsymbol{t}$**=0** at ***conception***, in units of days.

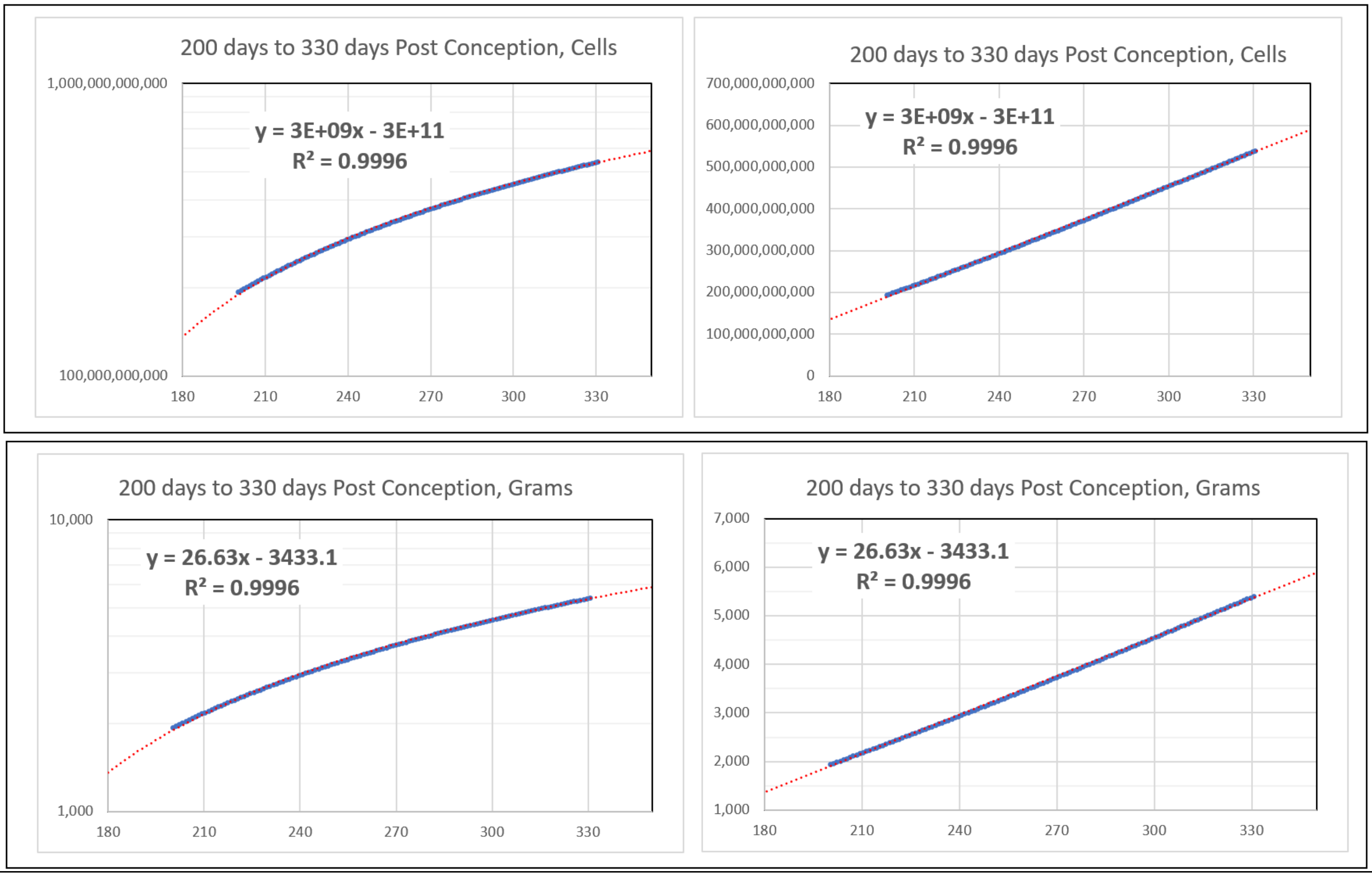


**Supplement-Figure-I.17**. The ***~Linear Phase of Human Fetal Growth***. The human growth curve calculated by the ***Binary Cellular Universal Growth Equation*** with (blue line), without ***Empirical Ripples*** adjustment. Shown from 180 days to 360 days post conception, with linear regressions examined from 200 days to 330 days post conception. Births occur, on average, at 268 days.

It follows that, without taking ***Ripple*** refinement into account, in the last 3 months of pregnancy:

***Fetal Size*** (in grams) ≈ 26.63*(***t***) – 3433.1 (503)

and

***Fetal Size*** (in numbers of cells) ≈ (26.63*(***t***) – 3433.1) * 1e8 (504)

where ***t*=0** at ***conception***, in units of days.

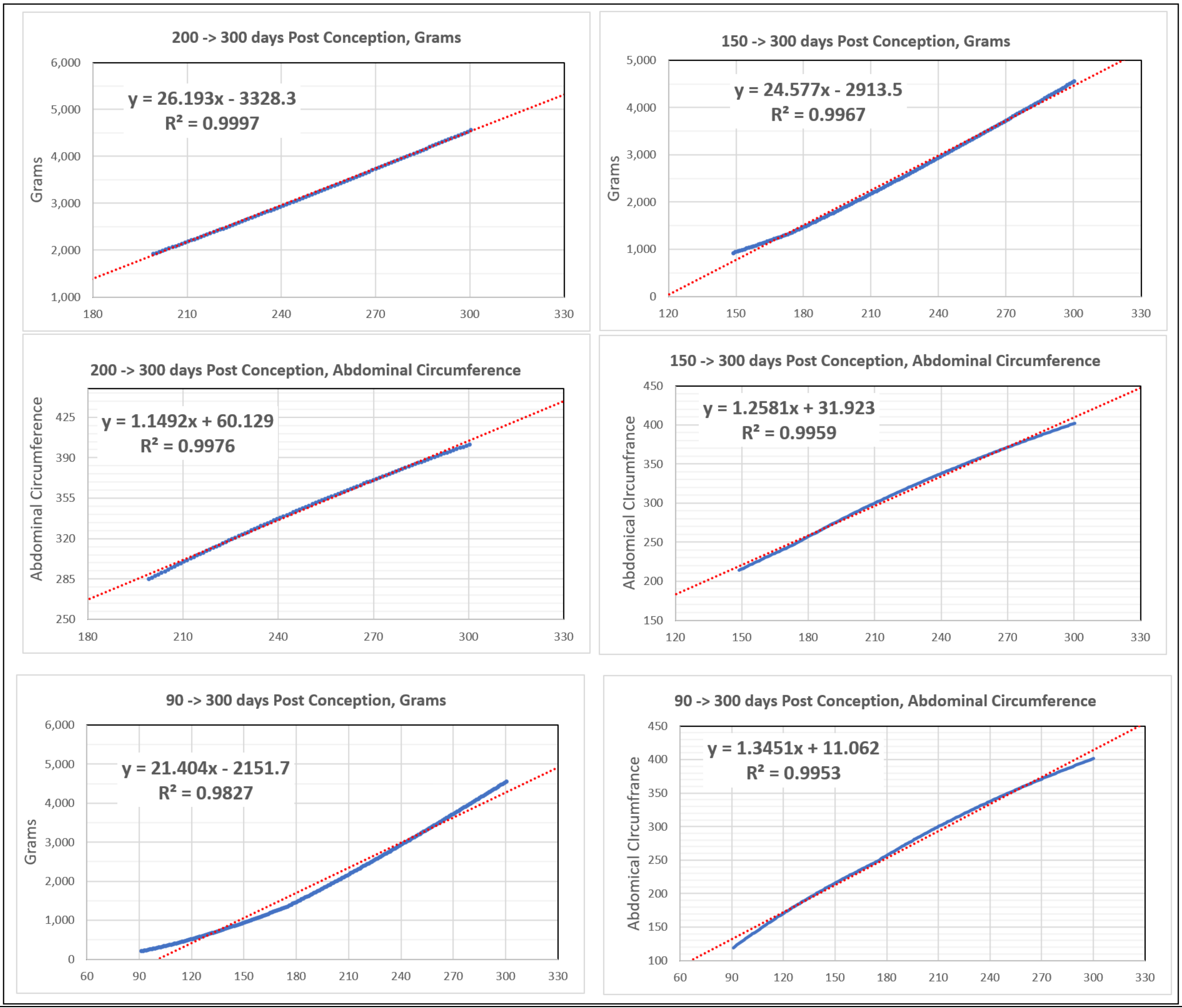


**Supplement-Figure-I.18**. The ***~Linear Phase of Human Fetal Growth***. The human growth curve calculated by the ***Binary Cellular Universal Growth Equation*** with (blue line), without ***Empirical Ripples*** adjustment. Linear regressions examined from 200 days to 300 days post conception and 150 days to 300 days post conception. Births occur, on average, at 268 days. Curvature appears when values are examined below 120 days.

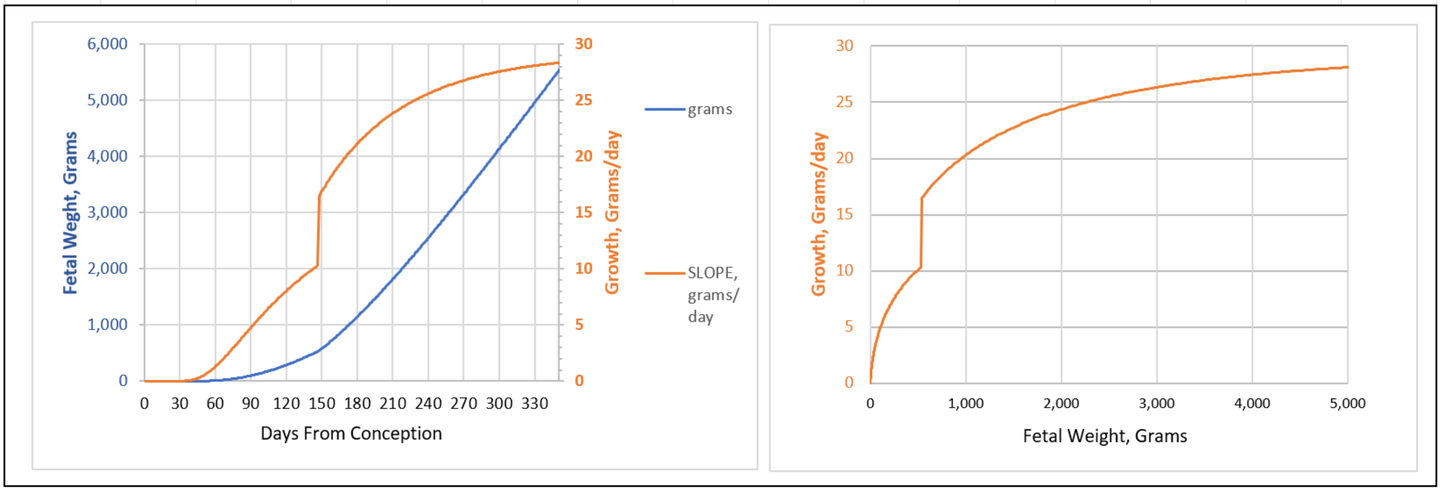


**Supplement-Figure-I.19**.
***Weight-Gain Rate***, increase ***Fetal Weight***, per day, in grams (Brown Lines), by ***Fetal Age*** (left) and ***Fetal Weight*** (right), from the ***Binary Cellular Universal Growth Equation*** with the ***Empirical Ripples*** adjustment (blue line). Note the during the ***~Linear Phase of Human Fetal Growth*** (red line), ***Weight-Gain Rate***, is between 20 and 27 grams/day.

In simple terms, in the last months of pregnancy, each day, the fetus increases by about 26 grams in weight, and about 3 billion cells (**Supplement-Figure-I.19**). This ***~Linear Phase of Human Fetal Growth***, which we see over the last 3 months of pregnancy, doesn't extend back in the first 6 months of life, and it appears to end after birth (**Supplement-Figure-I.20**).

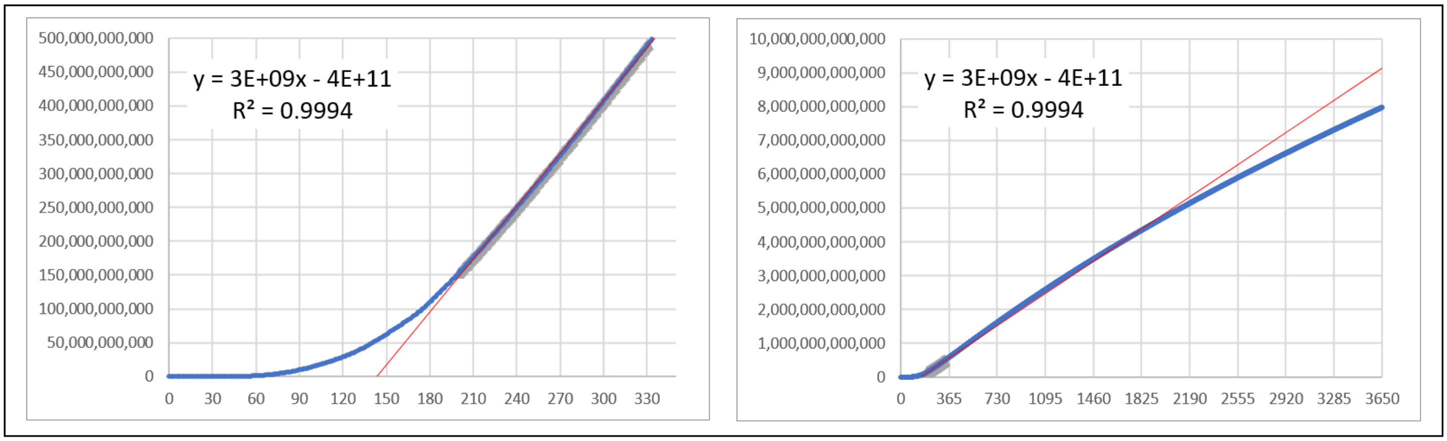


**Supplement-Figure-I.20**. The limits of the ***~Linear Phase of Human Fetal Growth***. The human growth curve calculated by the ***Binary Cellular Universal Growth Equation*** (#2) with (blue line), with ***Empirical*** curve-fitting described in the text from conception to 330 days post conception (left), from conception to 3650 days post conception (right), with linear regressions examined from 200 days to 330 days post conception. Births occur, on average, at 268 days.

***Binary Cellular Fetal Growth Analysis*** Gives Biologically, Data-Driven, ***Fetal Growth Curves***.

The ***Binary Cellular Universal Growth Equation*** (#2) affords the opportunity to create biologically, data-driven, ***Fetal Growth Curves*** (**Supplement-Figure-I.21a**). Shown in **Supplement-Figure-I.21a** are various ***Binary Cellular Universal Growth Equations*** (#2) with different values for ***"a"Parameter***, which, as described in the third paper in this series,[7] is likely to be the critical component for capturing such growth variety.

Growth curves can also now be constructed varying in the other parameters, ***b***, or ***c***, or in the modifier, ***R***, for the examination of fetal growth. Also show in **Supplement-Figure-I.21a** are ***Birthweights*** of babies in our dataset, also to be examined in greater detail in the third paper in this series,[7] and, of course, ***Estimated Fetal Weights*** calculated from ultrasound data can also be examined to determine which parameter values best capture the growth, and variety of growth, that occurs among fetuses.

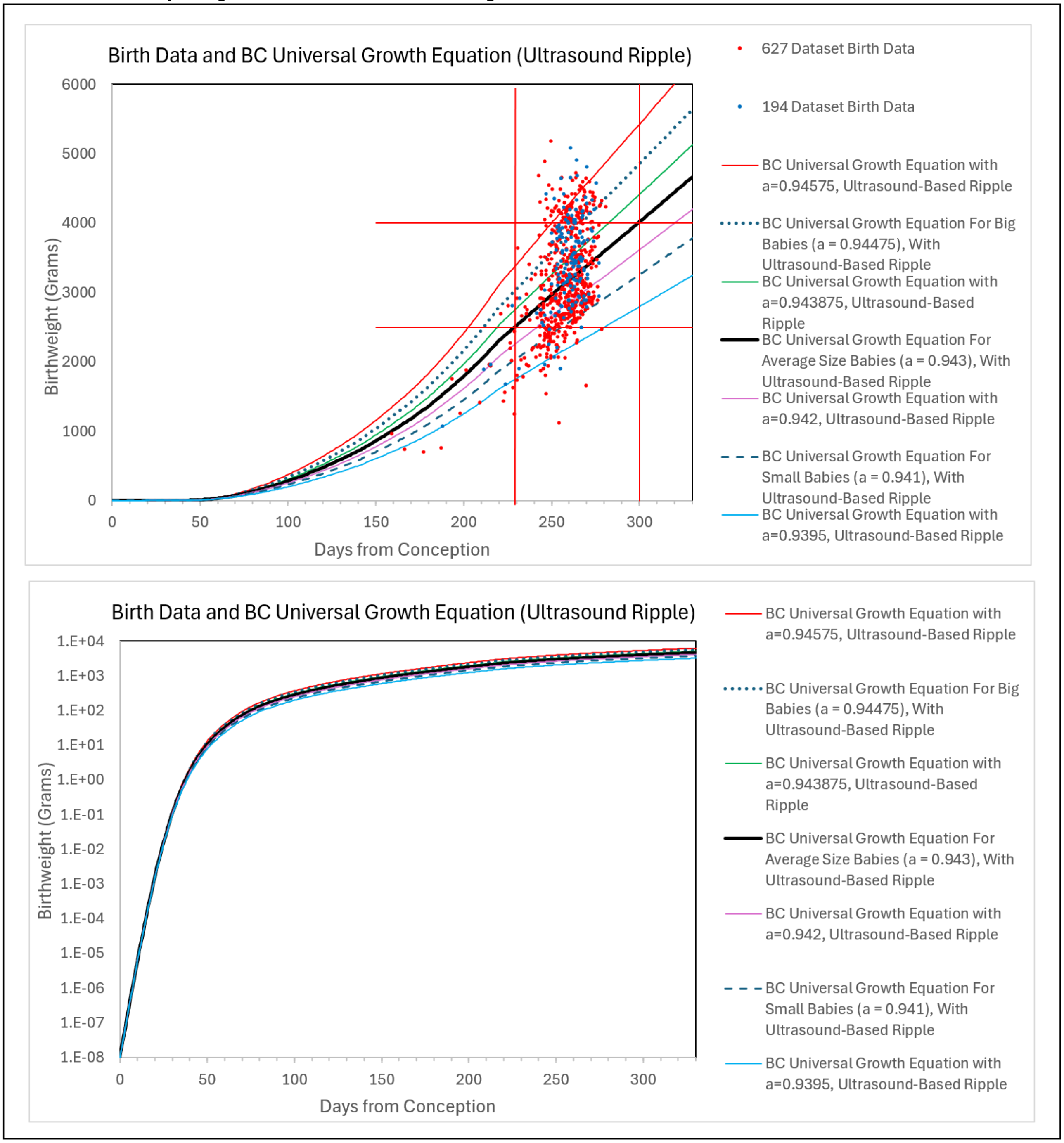


**Supplement-Figure-I.21a**. ***Fetal Growth Curves***, differing in the ***"a" Parameter*** of the ***Binary Cellular Universal Growth Equation***.

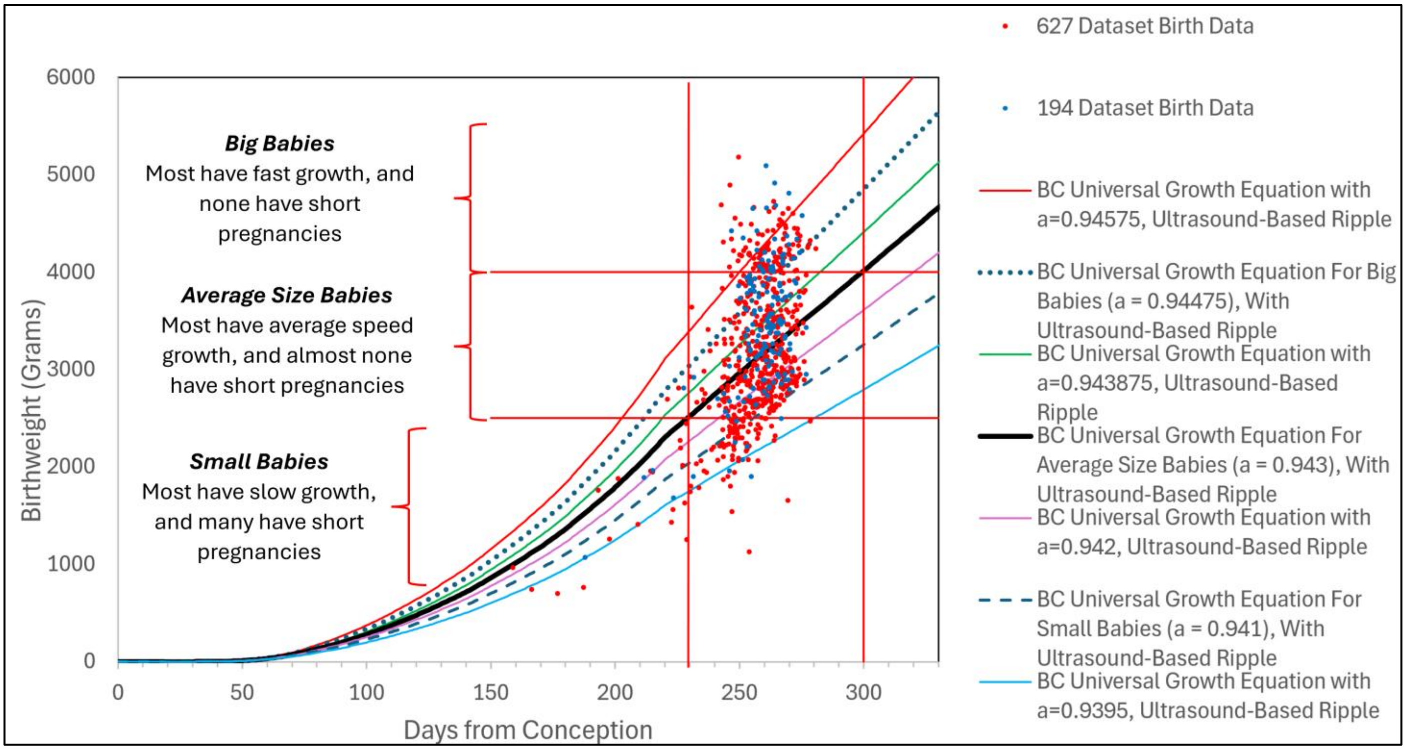


**Supplement-Figure-I.21b**. ***Fetal Growth Curves***, differing in the ***"a" Parameter*** of the ***Binary Cellular Universal Growth Equation***. The ***"a" Parameter*** values shown in this figure were taken from ***Average***, ***Small***, and ***Big Baby Groups***, plus values above, below, and in between those values.

***Weight-Gain Rate*** and ***Fetal Growth Velocity***

As can be seen in **Supplement-Figure-I.22** ***Weight-Gain Rate*** of the ***Binary Cellular Universal Growth Equation*** (#2) is about 24 grams a day at 210 days post conception, which is about 26 grams per day at 240 days post conception, and about 27 grams a day at 270 days post conception., a period of time we have called the ***~Linear Phase of Human Fetal Growth***. Mongelli and colleagues[118,111] have also noted this relative constancy of fetal growth at the end of pregnancy, a quality they call "quasi-linearity".

Growth charts examined in other studies also display, like the ***Binary Cellular Universal Growth Equation*** (#2), a rough, if not precise, similar amount of growth per day in the last months of pregnancy. For the NICHD growth chart, this is about 30 grams a day at 210 days post conception, and about 32 grams per day at 240 days post conception, and about 25 grams a day at 270 days post conception.[119,107] For the INTERGROWTH chart,[120,109] this is about 31 grams a day at 210 days post conception, and about 30 grams per day at 240 days post conception, and about 25 grams a day at 250 days post conception.

In the next paper in this series,[7] we shall show the values, when further refined with ultrasound-based size data (**Supplement-Figure-III.2**). Calculations of ***Weight-Gain Rates*** of the ***Binary Cellular Universal Growth Equation*** (#2), of the ***Binary Cellular Universal Growth Equations*** (#2) with different values for ***"a"Parameter***, (as well as the ***b***, or ***c***, parameters, or the ***R*** modifier) reflecting various centiles in specific populations, can now also be constructed, as the data become available.

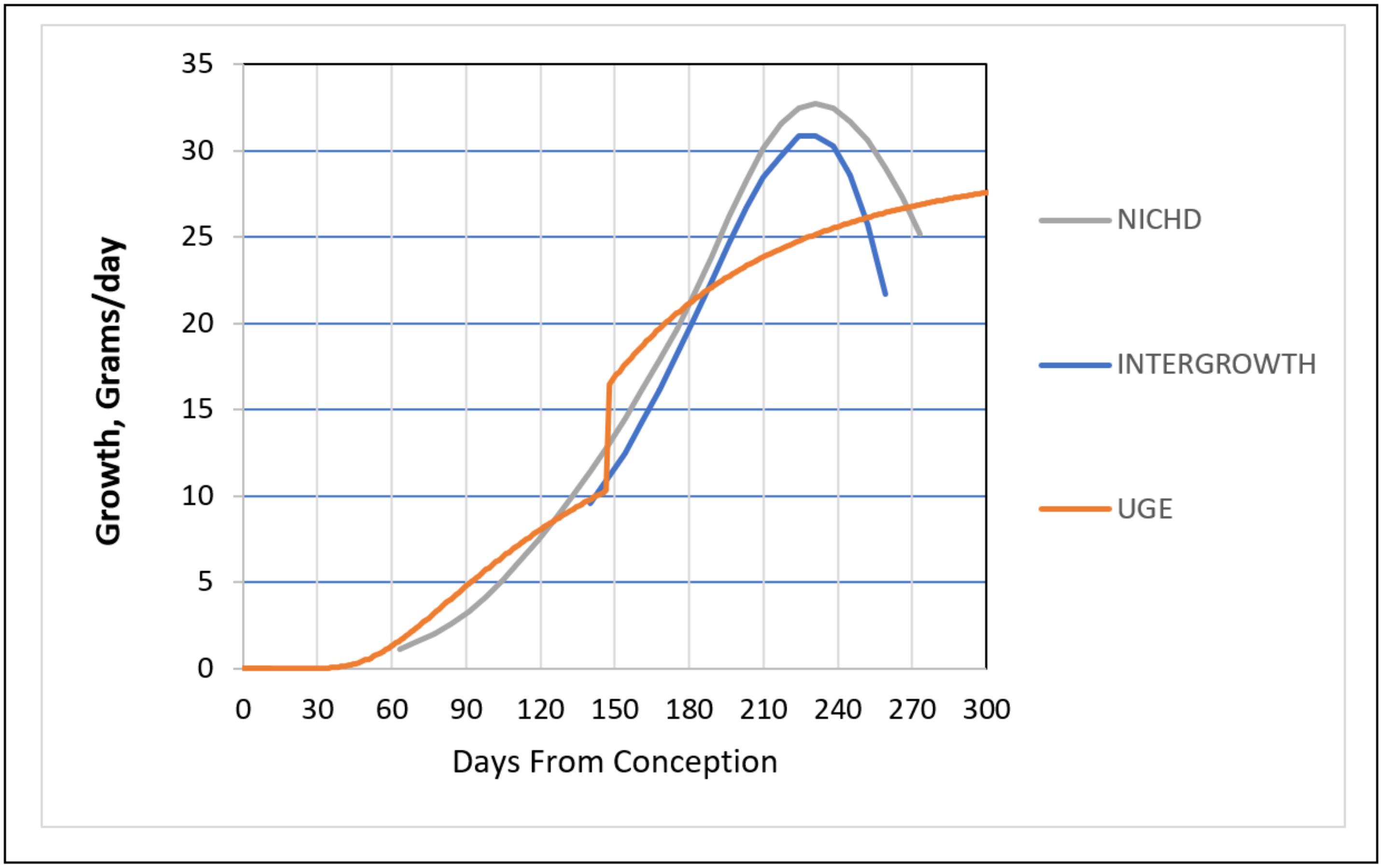


**Supplement-Figure-I.22**.

***Weight-Gain Rate***, the increase ***Fetal Weight***, per day, in grams (Brown Line), by ***Fetal Age*** (left) from the ***Binary Cellular Universal Growth Equation*** (#2) with the ***Empirical Ripples*** adjustment (blue line), characterizing the ***~Linear Phase of Human Fetal Growth*** (red line), together with data from INTERGROWTH[109,120] (Grey line) and NICHD[107,119] (Blue Line) growth charts. In the last paper in this series,[7] we shall examine the potential to use fetal ultrasound to refine, smooth, and define with greater precision, this first derivative of the ***Binary Cellular Universal Growth Equation*** (#2).

#### The ***Binary Cellular Biology*** of ***Body Part Creation*** and ***Growth***

Embryos create ***Body Parts***. Obstetricians rely upon the ultrasound measurement of these ***Body Parts*** to gauge the ***Size*** and ***Growth*** of fetuses. Thus, the practical task of managing the health of the fetus and its mother reaches down to the ancient and fundamental biology of the creation of the ***tissues***, ***organs***, and ***anatomical structures*** of the body, to arrive at actionable estimates of the size of the fetal body. Next, we shall outline how, with ***Binary Cellular Analysis***, the fundamental mechanism of ***Body Part*** formation can be deciphered, and how this information can be brought to the practical task of estimating human ***Fetal Weight*** from ultrasound measurement of these ***Body Parts***.

##### The ***Allometric Growth Equation***

For more than a century,[121-123] biologists have observed that the ***Size***, $\boldsymbol{p}$, of many parts of the body, when compared with the ***Size***, $\boldsymbol{w}$, of the body as a whole, frequently appear on log-log graphs as straight rows of ***p-w***-pairs, that is:

$$\log(\boldsymbol{w}) = \boldsymbol{S} \cdot \log(\boldsymbol{p}) + \log(\boldsymbol{B}) \qquad (10a)$$

which is equivalent to:

$$\boldsymbol{w} = \boldsymbol{B} \cdot \boldsymbol{p}^{\boldsymbol{s}} \qquad (10b)$$

This observation is called ***Allometric Growth***, and the ***Allometric Growth Equation***. Here we shall call $\boldsymbol{S}$ the ***Allometric Slope***, and $\boldsymbol{B}$, the ***Allometric Birth***, for reasons that will become evident as we examine the cellular basis of ***Allometric Growth*** next.

##### ***Binary Cellular Allometric Growth Analysis***

Why parts of the body grow in this ***Allometric*** fashion, and what $\boldsymbol{S}$ and $\boldsymbol{B}$ mean in terms of the biology of body-part formation and growth, have long been mysteries.[122,123] To solve these problems, we again return to the methodology of ***Binary Cellular Analysis***, to examine, in units of numbers of cells, $\boldsymbol{N}$, the creation and growth of the various parts of the body, $\boldsymbol{N_p}$, relative to the body as a whole, $\boldsymbol{N_w}$, examined on log-log graphs.[100] We call this method ***Binary Cellular Allometric Growth Analysis***.

We assembled values of $\boldsymbol{N_p}$ vs. $\boldsymbol{N_w}$ for each of the ***Body Parts*** from the three species of animals that have been characterized in the ***Cell Lineage Charts***, and thus telling us mitotic relationships for every cell going back to the single fertilized ovum (**Supplement-Figures-I.24-27**). These species include the nematode worms ***Caenorhabditis elegans*** and ***Meloidogyne incognita*** (**Supplement-Figures-I.24-26**), and the chordate tunicate ***Oikopleura dioica*** (**Supplement-Figure-I.27**) (see METHODS section for references).

We also assembled values of $\boldsymbol{N_p}$ vs. $\boldsymbol{N_w}$ for 25 ***Body Parts***, (***tissues***, ***organs***, and ***anatomical structures***) of vertebrate embryos and juveniles, again relying on the observation that there are about $10^8$ cells in every gram of tissue. These species (and the body parts examined) include: chicks (gizzards, livers, hearts, kidneys), rats (livers, brains, kidneys, forelegs, ears, stomachs, spinal cords), zebrafish (eye lens), mice (livers, brains, kidneys, forelegs), goldfish, and humans (brains, livers, kidneys, lungs, pancreases, adrenals, thymuses, spleens, lower extremities, upper extremities, stomach, heart, intestines) (see METHODS section for references) (**Supplement-Figures-I.28-33**).

##### The ***Binary Cellular Allometric Growth Equation***

When these $\boldsymbol{N_p}$-$\boldsymbol{N_w}$-pairs were displayed on log-log graphs (**Supplement-Figures-I.24-33**),[100] including data in the growth of various human body-parts measured in human fetal autopsy studies (**Supplement-Figures-I.28-33**),[94] they almost always appear as straight rows of dots, or:

$$\log(\boldsymbol{N_w}) = \boldsymbol{S_N} \cdot \log(\boldsymbol{N_p}) + \log(\boldsymbol{B_{N1}}) \quad \text{where} \quad \boldsymbol{S_N} = \frac{\boldsymbol{c_p}}{\boldsymbol{c_w}} \qquad (11a)$$

which is equivalent to:

$$\boldsymbol{N_w} = \boldsymbol{B} \cdot \boldsymbol{N_p}^{\boldsymbol{s}} \qquad (11b)$$

We call this the ***Binary Cellular Allometric Growth Equation*** (**Supplement-Figure-I.3**). We call $\boldsymbol{B_{N1}}$ the ***Binary Cellular Allometric Birth***, and $\boldsymbol{S_N}$ the ***Binary Cellular Allometric Slope***. As we shall see next, when data on the growth of many ***Body Parts***, from many animals, are examined by ***Binary Cellular Allometric Growth Analysis***, what emerges is that that animals make their ***tissues***, ***organs***, and ***anatomical structures*** from ***Single Founder Cells***, at their ***Binary Cellular Allometric Births***, $\boldsymbol{B_{N1}}$, each growing at the rate of their ***Binary Cellular Allometric Slopes***, $\boldsymbol{S_N}$, as determined by their ***Cell Cycle Times***, $\boldsymbol{c_p}$.

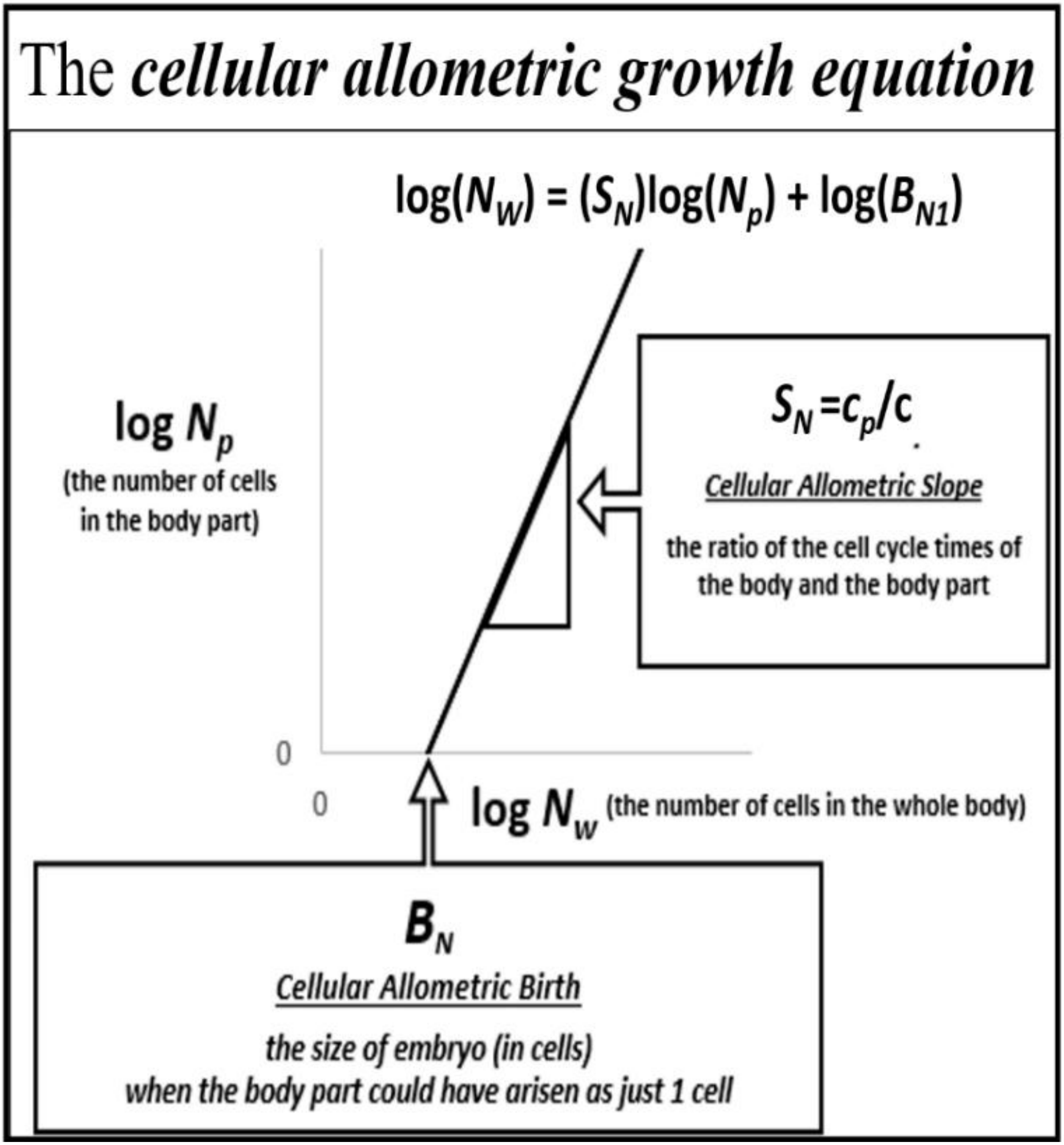


**Supplement-Figure-I.23: The *Binary Cellular Allometric Growth Equation*** (#11)

***Binary Cellular Analysis*** Reveals that ***Body Parts*** Arise from ***Single Founder Cells***.

Body Parts are Formed from ***Single Founder Cells***, as Seen From the Value of $B_{N1}$

Perhaps the most striking result that we have learned from the ***Binary Cellular Allometric Growth Analysis*** of the many ***Body Parts***, from many animals, we have studied (**Supplement-Figures-I.24-23**) is that each ***Body Part*** appears to arise from a ***Single Founder Cell*** at its ***Binary Cellular Allometric Birth***, $B_{N1}$. This is visualized by transforming ***Body Part*** growth data into $N_p$-$N_w$-pairs, and examining them on log-log graphs. On these graphs, we can see that the ***Binary Cellular Allometric Birth***, $B_{N1}$, of each ***Body Part,*** lies where the ***Binary Cellular Allometric Growth Equation*** crosses the $x$-axis, thus where $\log(N_p)=0$, thus where $N_p=1$, that is, when the ***Body Part*** could have been a ***Single Cell*** (**Supplement-Figure-I.3**). Many such examples pointing to the creation of ***Body Parts*** from single ***Founder Cells*** can be seen in the ***Binary Cellular Allometric Growth Analyses*** of these embryonic structures of developing nematodes, tunicates, chicks, mice, rats, zebrafish, and human embryos (**Supplement-Figures-I.27-33**).

Data from Animals with ***Cell Lineage Charts*** Show ***Body Parts*** Arise from ***Single Founder Cells***.

When we examined ***Body Parts*** animals for which we have full ***Cell Lineage Charts***, the data have always shown that the ***Binary Cellular Allometric Growth Equation*** points to where ***Body Part*** has arisen from a ***Single Founder Cell*** at its ***Binary Cellular Allometric Birth***, $B_{N1}$ (**Supplement-Figures-I.24-27**). These animals are nematode worms ***Caenorhabditis elegans*** and ***Meloidogyne incognita***, and tunicates ***Oikopleura dioica***, members of the chordate phylum to which we and our fellow vertebrates belong.

Note, for example, in **Supplement-Figures-I.24-26**, how the nematode ***C elegans*** **AB** ***Founder Cell*** arises when the embryo is just **2** cells in size, whose progeny go on to form the **AB** ***lineage***, which forms the skin from the ***Binary Cellular Allometric Birth*** of $B_{N1}$**≈2** (i.e., $N_w$**≈2** when $N_p$**≈1**). Similarly, note how the ***C elegans*** **E** ***Founder Cell*** arises when the embryo is **8** cells in size, whose progeny go on to form the **E** ***lineage***, which forms the intestine from the ***Binary Cellular Allometric Birth*** of $B_{N1}$**≈8** (i.e., $N_w$**≈8** when $N_p$**≈1**). Many other such examples of the creation of ***Body Parts*** from ***Single Founder Cells*** can be seen in **Supplement-Figures-I.24-27**.

Data from ***Vertebrate*** Animals Show ***Body Parts*** Appear to Arise from ***Single Founder Cells***.

When we examined ***Body Parts*** of ***Vertebrate*** animals for which we don't have full cell lineage data, the data have always shown that the ***Binary Cellular Allometric Growth Equation*** points to where we would expect to find a ***Single Founder Cell*** at its ***Binary Cellular Allometric Birth***, $B_{N1}$. These species (and the body parts examined) include: chicks (gizzards, livers, hearts, kidneys), rats (livers, brains, kidneys, forelegs, ears, stomachs, spinal cords), zebrafish (eye lens), mice (livers, brains, kidneys, forelegs), goldfish, and humans (brains, livers, kidneys, lungs, pancreases, adrenals, thymuses, spleens, lower extremities, upper extremities, stomach, heart, intestines) (see METHODS section for references) (**Supplement-Figures-I.28-32**).

Note, for example, in **Supplement-Figure-I.28**, $B_{N1}$**≈8** for the chicken heart, suggesting that the heart could have arisen from just 1 cell, when the embryo was **~8** cells in size. For the chicken liver, $B_{N1}$**≈2,000**, suggesting that the liver could have arisen from just **1** cell, when the embryo was **~2,000** cells in size (**Supplement-Figure-I.28**). Many other such cases can be seen in **Supplement-Figures-I.28-32**.

Note, also, in **Supplement-Figure-I.32**, $B_{N1}$**≈8** for the human heart, suggesting that the heart could have arisen from just 4 cells, when the embryo was ~**16** cells in size. For the liver, $B_{N1}$**≈4**, suggesting that the liver could have arisen from just **1** cell, when the embryo was ~**4** cells in size, for each kidney, $B_{N1}$***≈2,000***, for the stomach, $B_{N1}$**≈100**, for the brain, $B_{N1}$**≈2**, for each lung, $B_{N1}$**≈500**, for the arms, $B_{N1}$***≈500***, for the legs, $B_{N1}$***≈1,000***.

We Have Found No Cases Where the Creation of ***Body Parts*** Arises from ***Multiple Founder Cells***.

Of course, one cannot definitively distinguish between establishment of ***Body Parts*** from ***Single*** vs ***Multiple Founder Cells*** when we lack imaging data, which is usually the case for data on vertebrate ***Body Part*** growth late in development (**Supplement-Figures-I.27-33**). Nevertheless, in case after case, the projected values of $B_{N1}$ derived from actual growth data have favored the single ***Founder Cell*** option. The single ***Founder Cell*** option is also the most likely in terms of ***Parsimony***,[124,125] as we shall see below. Ultimately, the absence of such evidence simply calls out for settling the issue with actual observation, and ***Binary Cellular Allometric Growth Analysis*** provides us with a guide for carrying out the imaging studies that will make the counting of cells possible.

Many Others Have Observed ***Body Parts*** Arising from a ***Few*** or ***Single Founder Cells***.

While few of the large structures from late in development have been measured to the $B_{N1}$ intersection point, the zebrafish eye lens comes close, as Greiling and Clark documented its growth from just 8 cells, when the embryo is 16 hours post fertilization.[90] As can be seen in **Supplement-Figure-I.33**, ***Binary Cellular Analysis*** of their data reveal that this body parts also growths by the ***Binary Cellular Allometric Growth Equation***, pointing down to a ***Binary Cellular Allometric Birth*** when the embryo was about 2000 cell in size, and thus $B_{N1}$**≈2000** when $N_p$**≈1**. A variety of other such observations have also been noted in many research studies, and will be reviewed in the DISCUSSION.

***Body Parts*** Can be Made With 1 ***Founder Cell*** With Ease, or More Than 1 with Difficulty.

Why would body parts arise from just a ***Single Founder Cell***? Evolution favors the simplest way forward, a principal called ***Maximal Parsimony***,[124,125] and constructing a body part from a ***Single Founder Cell*** is that simplest way forward. This could easily be seen in a calculation of an idealized ***Body Part*** that is made from **3** ***Founder Cells***, arising early in development, when growth is close to exponential (**Supplement-Figure-I.34**).

As can be seen in the top two graphs of **Supplement-Figure-I.34**, if all three ***Founder Cells***, arise at the same time, and have the same ***Cell Cycle Time***, $c_p$, their growth together will be indistinguishable from a part made of **1** ***Founder Cell*** (once it gets up to 3+ cells!). Furthermore, as can be seen in the other graphs of **Supplement-Figure-I.34**, no matter when the three ***Founder Cells*** are born, or what their ***Cell Cycle Times***, $c_p$, are, the aggregate growth of the part will still appear as a quite straight row of datapoints on a log graph. Thus, the fact that the growth of a ***Body Part*** points back to a single ***Binary Cellular Allometric Birth***, doesn't mean that it arose from a single cell; we still have to do the microscopy work to see what actually happened.[90,126]

This excise also shows, however, that ***Body Parts*** made of more than a single ***Founder Cell*** do have some challenges. Note for example, that ***Founder Cells*** born after the first ***Founder Cell***, or having a lower ***Cell Cycle Time***, $c_p$, than the fastest growing ***Founder Cell***, will soon become irrelevant, and the ***Body Part*** will become functionally monoclonal. A ***Founder Cell*** born three ***Cell Cycle Times***, **3**x*c*, after the first ***Founder Cells***, which in a human or a mouse is about three days, will make up only about 10% of the part, while a ***Founder Cell*** born eight ***Cell Cycle Times***, **8**x*c*, after the first ***Founder Cell*** will make up only about 1% of the part. ***Founder Cells*** that have ***Cell Cycle Times***, $c_p$, slower than the fastest ***Founder Cell*** will also soon lose in the race to make up a significant percentage of the part. Thus, when evolution calls out for the creation of a body part, accomplishing the task with a ***Single Founder Cell*** would be the way forward most likely to achieve that goal.

#### ***Binary Cellular Analysis*** Reveals that ***Body Part*** Growth is Set by Change in the ***Cell Cycle Time, $c_p$***.

##### Body Parts Grow at Various Speeds from ***Single Founder Cells***, as Seen from the Value of $S_N$

Another striking result that we have learned from the ***Binary Cellular Allometric Growth Analysis*** of the many ***Body Parts***, from many animals, we have studied (**Supplement-Figures-I.24-33**) is that that the speed of the growth of each ***Body Part***, and thus its final ***Size***, is captured by the ***Binary Cellular Allometric Slope***, $S_N$, of the ***Binary Cellular Allometric Growth Equation***. The value of $S_N$, the ***Binary Cellular Allometric Slope*** appears on log-log graphs as a measure of whether a ***Body Part*** is growing faster ($S_N$>**1**), slower ($S_N$<**1**), or at the same speed ($S_N$=**1**), as the body as a whole.

##### ***Body Part Allometric Growth*** Results From the ***Binary Cellular Universal Growth Equation***.

Recall that the ***Binary Cellular Allometric Growth Equation*** captures the rate of growth of the whole embryo, while the rate of growth for each body-part is:

$$\frac{dN_p}{dt} = \frac{ln(2)}{c_p} \cdot N_p \cdot (m_p) \qquad (650)$$

where $c_p$ is the ***Cell Cycle Time***, and $m_p$ the ***Mitotic Fraction*** of the cells of the ***Body Part***. Thus, the ratio of the rate of growth of the body to the body-part is:

$$\frac{dN_w}{dN_p} = \left[\frac{ln(2)}{c} \cdot N_w \cdot (m)\right] \cdot \left[\frac{ln(2)}{c_p} \cdot N_p \cdot (m_p)\right]^{-1} \quad \text{, where} \quad m = a^{(N_w^b)} \quad \text{and} \quad m_p = a_p^{(N_w^{bp})} \qquad (651)$$

Consider the case where $m_p$=$m$ and $c_p$=$c$, that is, where a ***Body Part*** has the same ***Mitotic Fraction*** and ***Cell Cycle Time*** as the embryo, and starts sometime later than the first cell:

$$\frac{dN_w}{N_w} = \frac{dN_p}{N_p} \qquad (652)$$

Integration[127] reveals:

$$log(N_w) = \log(N_p) + \log(B_N) \qquad (652)$$

which is the allometric relationship, with $S_N$=**1.** Thus, growth of a part of the body by the ***Binary Cellular Allometric Growth Equation*** is the consequence of ***Founder Cells*** giving rise to cell lineages in animals growing by the ***Universal Growth Equation***.

##### ***Binary Cellular Allometric Growth Equation***: ***Body Part*** Growth is Set by ***Cell Cycle Time***.

Now, let us consider the case where $m_p$=$m$ but $c_p \neq c$, that is, where a ***Body Part*** has the same ***Mitotic Fraction*** as the embryo but a different ***Cell Cycle Time***:

$$\frac{dN_w}{N_w} = \frac{c_p}{c} \cdot \frac{dN_p}{N_p} \qquad (653)$$

Integration reveals:

$$log(N_w) = \frac{c_p}{c} \cdot \log(N_p) + \log(B_N) \qquad (653)$$

where:

$$S_N = \frac{c}{c_p} \qquad (655)$$

which is the allometric relationship, with $S_N$,≠**1**. Thus, $S_N$, the ***Binary Cellular Allometric Slope*** of the ***Binary Cellular Allometric Growth Equation***, can be traced to the ***Cell Cycle Time***, $c_p$ in each part of the body (**Supplement-Figure-I.3**).

On the other hand, similar analysis revealed the ***Mitotic Fraction, $m_p$***, that is, a change in the fraction of cell dividing, to be an unlikely source for allometric log-linearity. Indeed, changes in the value of either ***a*** or ***b*** of the ***Universal Mitotic Fraction Equation*** (#4), which sets the value of $m_p$, does not result in straight lines on log-log graphs but in curves (not shown).

##### ***Body Part*** Growth, $S_N$, is Set by ***Cell-Heritable*** Change in ***Cell Cycle Time***, $c_p$, of ***Founder Cell***

That the value of the ***Cell Cycle Time***, $c_p$, of the cells in a ***Body Part*** is ***Cell-Heritable*** can be seen in the straightness of the rows of $N_p$ vs $N_w$ dots appearing on the log-log graphs shown in **Supplement-Figures-I.24-33**. This reveals that the length of time that it takes a ***Founder Cell*** to divide, and to give rise to its first two cells, is the same length of time that it takes the ***Progeny*** of that ***Founder Cell*** to divide, that is, all of the subsequent cells that make up that ***Body Part***.

***Binary Cellular Analysis*** Provides Quantitative Information on How ***Cellular Selection*** Creates the Body.

***Cellular Selection***, resulting from ***Differential Cellular Proliferation***, or ***Differential Cell Death***, molds the formation of the body from a single fertilized ovum into a multicellular animal.[1,2] The action of ***Cellular Selection*** can be seen to be at work throughout the formation of the animal body, which can now be measured by ***Binary Cellular Analysis***.

Thus, the role of ***Cellular Selection*** in the overall ***Growth*** of the body can be measured by the ***Mitotic Fraction***, ***m***, of the ***Binary Cellular Universal Mitotic Fraction Equation*** (**Supplement-Figure-I.3**). As we have seen, and will examine in greater detail in the third paper in this series,[7] the role of ***Cellular Selection***, measured by the decline in the value of the ***Mitotic Fraction***, ***m***, that occurs from conception to maturity, and the ***a*** and ***b Parameters*** of the ***Binary Cellular Universal Mitotic Fraction Equation*** that determines this decline, captures the differences in ***Size*** and ***Growth*** between animals across the taxonomic spectrum, and between human fetuses.

Similarly, the role of ***Cellular Selection*** in the innumerable fine scale details of ***Body Part*** formation can be measured by the ***Binary Cellular Allometric Birth***, $B_{N1}$, and the ***Binary Cellular Allometric Slope***, $S_N$, of the ***Binary Cellular Allometric Growth Equation*** (**Supplement-Figure-I.23**). As we have seen, the ***Binary Cellular Allometric Birth***, $B_{N1}$, is marked by the ***Selection*** of a ***Single Founder Cell*** among many cells in the body, at its ***Binary Cellular Allometric Birth***, when the body is comprised of $B_{N1}$ cells (i.e. $N_w=B_{N1}$ when $N_p=1$). As we have also seen, the ***Binary Cellular Allometric Slope***, $S_N$, is marked by the ***Selection*** of the ***Progeny*** of the ***Founder Cell***, made possibe by a ***Cell Heritable Change*** in the ***Cell Cycle Time*** $c_p$, which occurs in the ***Founder Cell*** at its ***Binary Cellular Allometric Birth***, $B_{N1}$, and is thus inherited in the ***Progeny*** of that ***Founder Cell***, which will make up that ***Body Part***.

***Case Study*** in the ***Binary Cellular Analysis*** of ***Cellular Selection***: ***Limb Morphogenesis***

***Limb Morphogenesis***, Captured by Descriptive Embryology

To take but one example, ***Limb Morphogenesis***, our ***Arms*** and ***Legs*** arise because the cells where these structures will appear divide more abundantly than the cells in the other areas of our flanks. We first see the results of such ***Cellular Selection*** in the appearance of ***Limb Buds***, and then ultimately in the appearance of our ***Arms*** and ***Legs***. ***Differential Cell Death*** sculpts the details of these structures, by accomplishing such tasks as forming our fingers and toes by eliminating the cells in between them.

***Limb Morphogenesis***, Captured by ***Binary Cellular Biology***

***Binary Cellular Analysis***, the consideration of the formation of the body, in units of numbers of cells, ***N***, provides us with quantitative information on how such ***Cellular Selection*** creates the body. To return to our example of ***Limb Morphogenesis***, ***Binary Cellular Analysis*** of the formation of the limbs of rats, mice, and humans (**Supplement-Figures-I.29-31**) extends the descriptive embryology of our ***Arms*** and ***Legs***, into the quantitative realm, provides us with all sorts of new information about how our bodies made these structures.

One thing that the ***Binary Cellular Analysis*** shown in these figures has told us is that the ***Binary Cellular Allometric Slope***, $S_N$, of our ***Legs*** is greater than our ***Arms***, due to a shorter ***Cell-Heritable*** change in the ***Cell Cycle Time***, $c_p$, of our ***Legs*** in comparison to our ***Arms***. Thus, ***Binary Cellular Analysis*** shows us that ***Cellular Selection***, resulting from ***Differential Cellular Proliferation***, not only results in the very existence of our ***Arms*** and ***Legs***, it also shows us why are our ***Legs*** are bigger than our ***Arms***. Embryologists also made a 5th limb by putting various things where a limb wouldn't otherwise be, so we are capable of experimentally inducing ***Cellular Selection***, resulting from ***Differential Cellular Proliferation*** to make these monsters.[1]

Another thing that the ***Binary Cellular Analysis*** shown in these figures has told us is that the ***Binary Cellular Allometric Birth***, $B_{N1}$ of our ***Arms*** and ***Legs*** occurs when we are 100-300 cells in size. Whether our ***Arms*** and ***Legs*** arise from ***Single Founder Cells*** at that time can't be seen in the data we have, but at least ***Binary Cellular Analysis*** again tells us where we should go make microscopy images and count cells. As we have noted above, for every ***Body Part***, in every animal, for which we have full cell lineage data (nematodes and tunicates), the data have always shown that the origin of each ***Body Part*** arises from a ***Single Founder Cell*** at its ***Binary Cellular Allometric Birth***, $B_{N1}$., which is most definitely the ***Cellular Selection*** of a very few cells out of many cells.

***Body Part*** Size to ***Body Part*** Size: Their ***Allometric*** Relationships, Revealed by ***Binary Cellular Analysis***

Let us now consider the growth of two parts of the body, $p_1$ and $p_2$, both of which display allometric growth with reference to the growth of the body as a whole, $w$.

For part 1:

$$\log(N_w) = \left(\frac{S_{N1}}{c}\right)\log(N_{p1}) + \log(B_{N1}) \tag{719}$$

For part 2:

$$\log(N_w) = \left(\frac{S_{N2}}{c}\right)\log(N_{p2}) + \log(B_{N2}) \tag{720}$$

It follows that:

$$\log(N_w) = (S_{N1})\log(N_{p1}) + \log(B_{N1}) = (S_{N2})\log(N_{p2}) + \log(B_{N2}) \tag{721a}$$

which is equivalent to:

$$\log(N_{p1}) = \left(\frac{S_{N2}}{S_{N1}}\right)\log(N_{p2}) + \log\left(\frac{B_{N2}}{B_{N1}}\right) \tag{721b}$$

In other words, if two parts of the body show log-linear allometric growth with respect to the size of the body as a whole, they will also show log-linear allometric growth with respect to each other, with each part's ***Binary Cellular Allometric Birth***, $B_N$ and ***Binary Cellular Allometric Slope***, $S_N$ repackaged into the slope and intercept of this new variation on the ***Binary Cellular Allometric Growth Equation***.

It follows that if the ***Body Parts*** in question are in units of mass or volume or length, we can repeat the treatment outlined in the previous section. Thus:

$$\log(p_1) = \left(\frac{1}{S_{1,2}}\right)\log(p_2) + \log(B_{1,2}) \tag{722a}$$

which is equivalent to:

$$p_1 = B_{1,2} \cdot p_2^{\frac{1}{S_{1,2}}} \tag{722b}$$

The Geometry of the ***Binary Cellular Allometric Growth Equation***

A simple geometric examination can show that if the ***Size*** of body-part is related to the ***Size*** of the body as a whole in an ***Allometric*** fashion, then this ***Allometric*** relationship extends to other measurements of the body-part, such as diameter or circumference, if these geometrical values relate in the most common, ordinary, Euclidian fashion. For example, the circumference of a sphere is $2/3^{rds}$ the power of the volume of the sphere, which means that a circumference measure of a ***Body Part*** will retain the ***Allometric*** relationship of a part of the body, This quality makes it possible to relate not only the ***Size*** of a body-part, but other measurements of that body-part as well, such as circumference or length, thus allowing us to use this mathematics in the analysis of ***Fetal Ultrasound Measurements***, as we shall do below.

Forming ***Binary Cellular Estimated Fetal Weight Equations***.

We can now put ***Binary Cellular Analysis*** to work to derive a new, biologically based, method for calculating ***Fetal Weight*** from ***Ultrasound Measurements***. Let us begin with the ***Binary Cellular Allometric Growth*** and ***Allometric Growth Equations*** (#10 and #11) to examine the ***Allometric*** relationship between individual ***Fetal Ultrasound Measurements*** (***U***) and ***Birthweights*** ($B_w$):

$$\log(\textit{Estimated Fetal Weight}) = \sigma \cdot \log(\textit{Ultrasound Measurement}) + \log(\beta) \tag{12a}$$

which is equivalent to:

$$\textit{Estimated Fetal Weight} = \beta{*}(\textit{Ultrasound Measurement}^{\,\sigma}) \tag{12b}$$

We call this expression the ***1st-Steps-Binary Cellular Estimated Fetal Weight Equation*** (#12). In the next paper in this series, we shall put this equation to work, and test it, against patient data. To correct for the small size-correlated distortions that many have noted may appear in such calculations, one final parameter, $\gamma$ is added:

$$\textit{Fetal Weight} = \beta{*}(\textit{Ultrasound Measurement}^{\,\sigma}) - \gamma \tag{14a}$$

or

$$EFW = \beta{*}(U^{\sigma}) - \gamma \tag{14b}$$

We call Equation #14 the ***Binary Cellular Estimated Fetal Weight Equation***. As we shall see in the next paper in this series,[6] in many ways, ***Binary Cellular Estimated Fetal Weight Equations*** provide superior estimations of ***Fetal Weight*** than other currently used ***Estimated Fetal Weight Equations***.

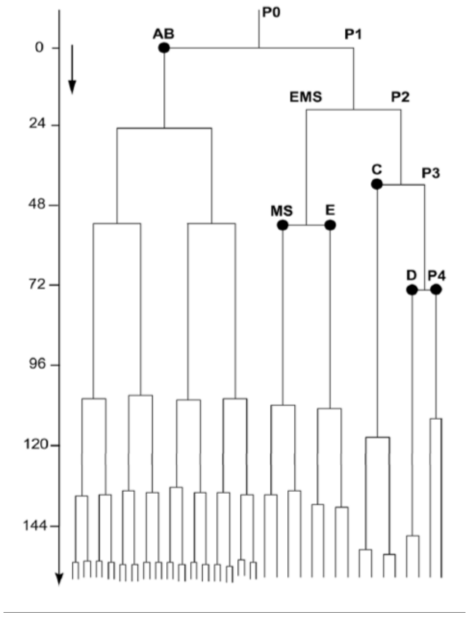


**Supplement-Figure-I.24**: ***Cell lineage chart*** of the ***cell lineages*** of the developing root knot nematode, ***Meloidogyne incognita***. ***Cell lineage chart*** and data from Calderón-Urrea et al[84,100]

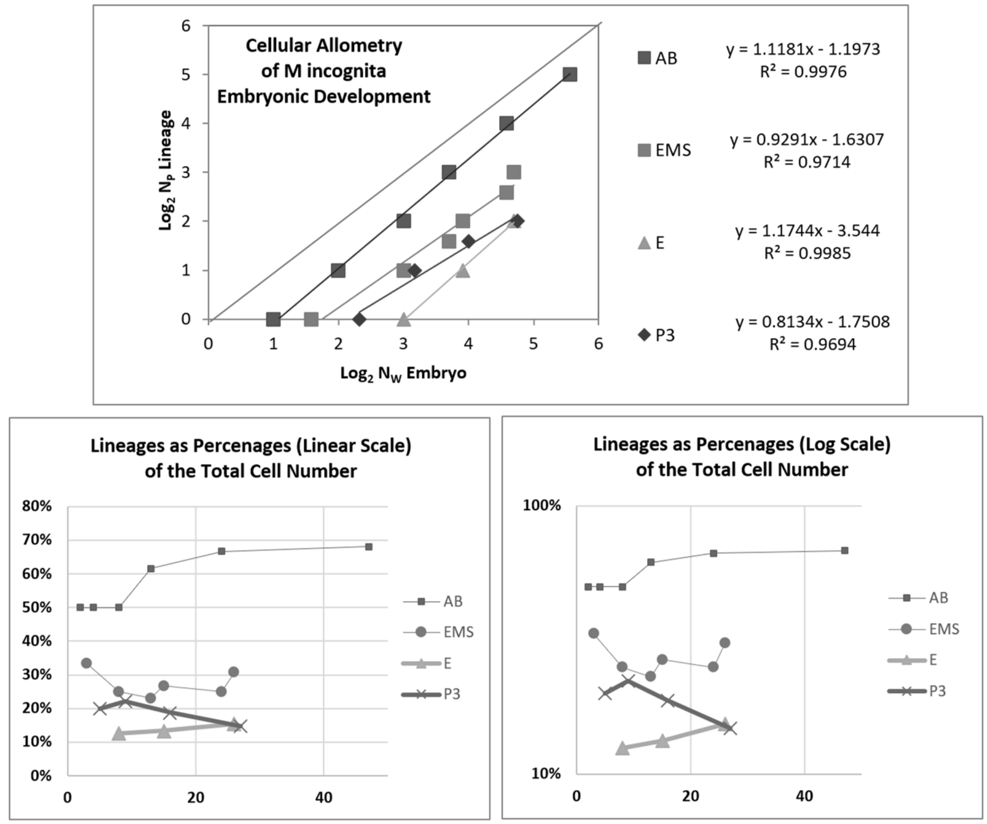


**Supplement-Figure-I.25**: ***Binary Cellular Allometric Lineage Growth Analysis*** of the ***cell lineages*** of the developing root knot nematode, ***Meloidogyne incognita***.[100,85]

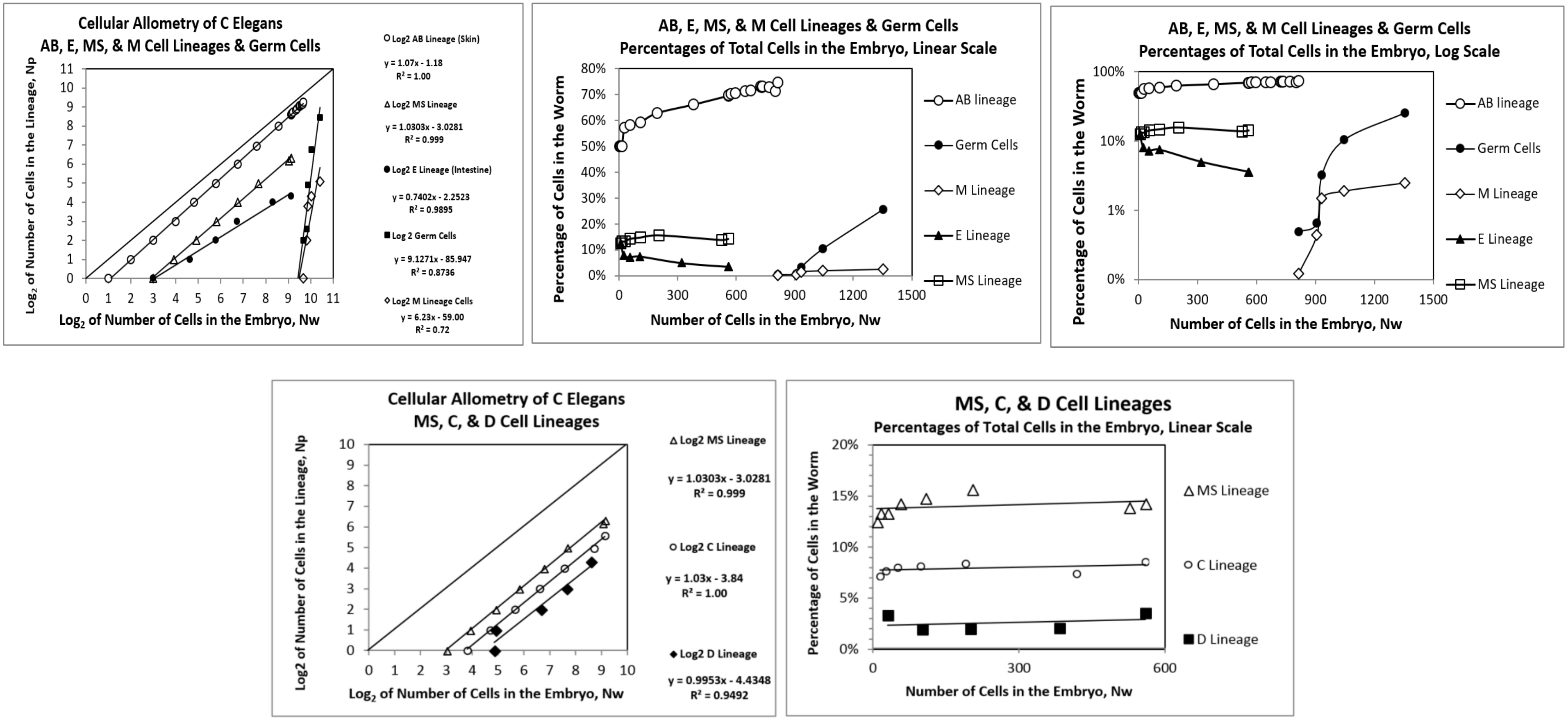


**<u>Supplement-Figure-I.26</u>**: ***Binary Cellular Allometric Lineage Growth Analysis*** **of the** ***cell lineages*** **of the developing** ***C elegans*** **nematode**,[100]

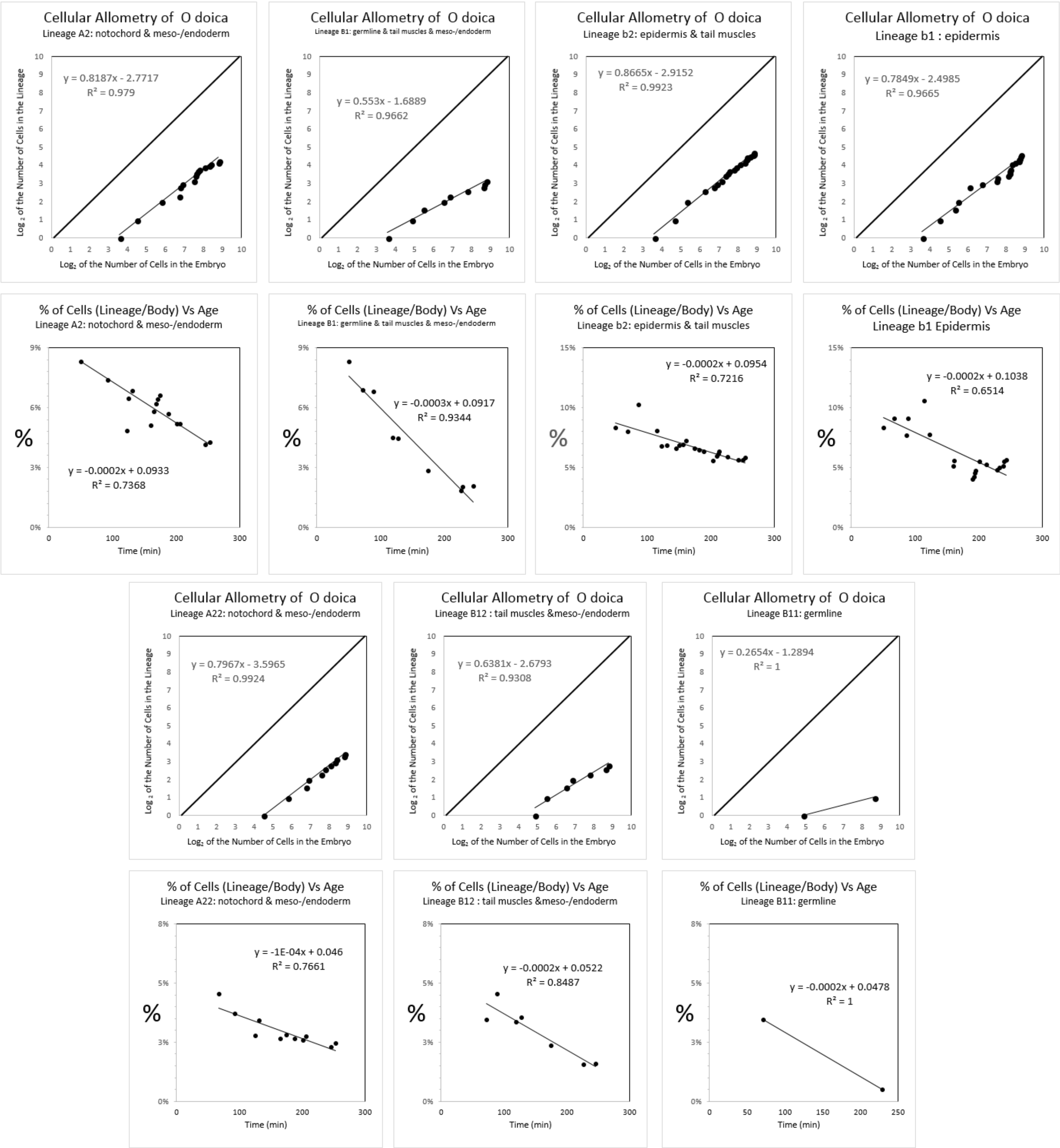


<u>**Supplement-Figure-I.27**</u>**:** ***Binary Cellular Allometric Lineage Growth Analysis*** **of the** ***cell lineages*** **of the chordate tunicate** ***Oikopleura dioica*****.** Unlike ***C elegans*** and ***M incognita***, all of the cases of ***Cellular Selection*** for this species reflect reductions in cell number, although this could be due to the limited size of the dataset rather than a biological difference. ***Cell lineage chart*** data from Stach et al[85,100]

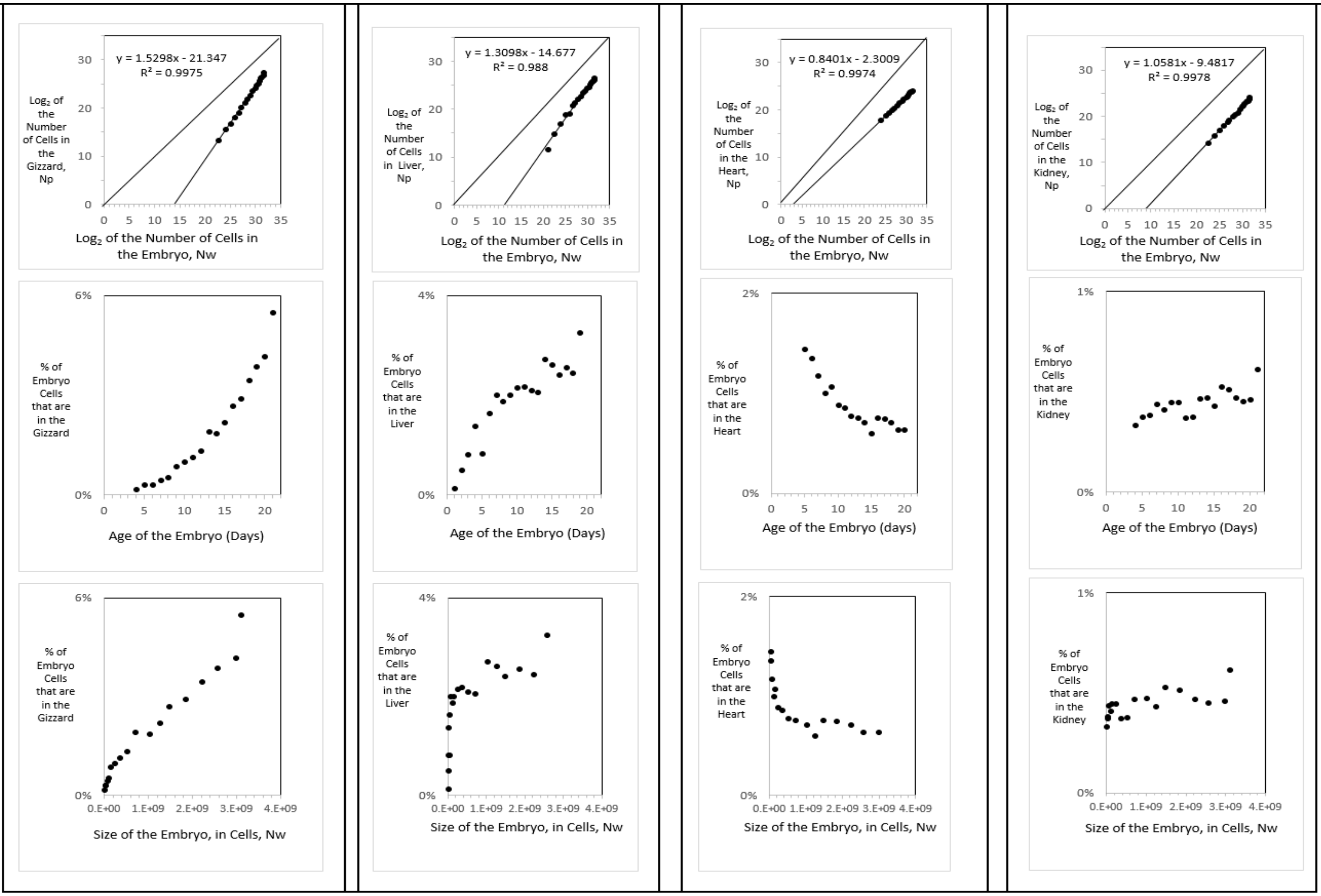


| | % | age | Size of Embryo in Cells | % | age | Size of Embryo in Cells, N | % | age | Size of Embryo in Cells, N | % | age | Size of Embryo in Cells, N |
|---|---|---|---|---|---|---|---|---|---|---|---|---|
| smallest value | 0.17% | 4 | 6.E+06 | 0.14% | 3 | 2.E+06 | 1.35% | 6 | 3.E+07 | 0.33% | 4 | 6.E+06 |
| largest value | 5.49% | 21 | 3.E+09 | 3.26% | 21 | 3.E+09 | 0.64% | 20 | 3.E+09 | 0.61% | 21 | 3.E+09 |
| % increase absolute | 5.32% | | 3.E+09 | 3.12% | | 3.E+09 | -0.72% | | 3.E+09 | 0.28% | | 3.E+09 |
| % increase relative | 33 | | 516 | 23 | | 1475 | 0.47 | | 87 | 2 | | 516 |
| absolute increase relative | 16,990 | | | 33,633 | | | 41 | | | 950 | | |

**<u>Supplement-Figure-I.28</u>: *Binary Cellular Allometric Growth Analysis* of *tissues*, *organs*, and *anatomical structures* of the chick embryo.** Growth of the parts of the chick embryonic gizzard, liver, heart, and kidney, in units of numbers of cells, in comparison to the number of cells in the embryo as a whole shown on log-log graphs, showing how the fraction of cells comprising in the gizzard and liver increases as the number of cells in the embryo increase with development and how the fraction of cells comprising in the heart decreases as the number of cells in the embryo increase with development, and how the fraction of cells comprising in the kidney remains roughly constant as the number of cells in the embryo increases with development.[100,]

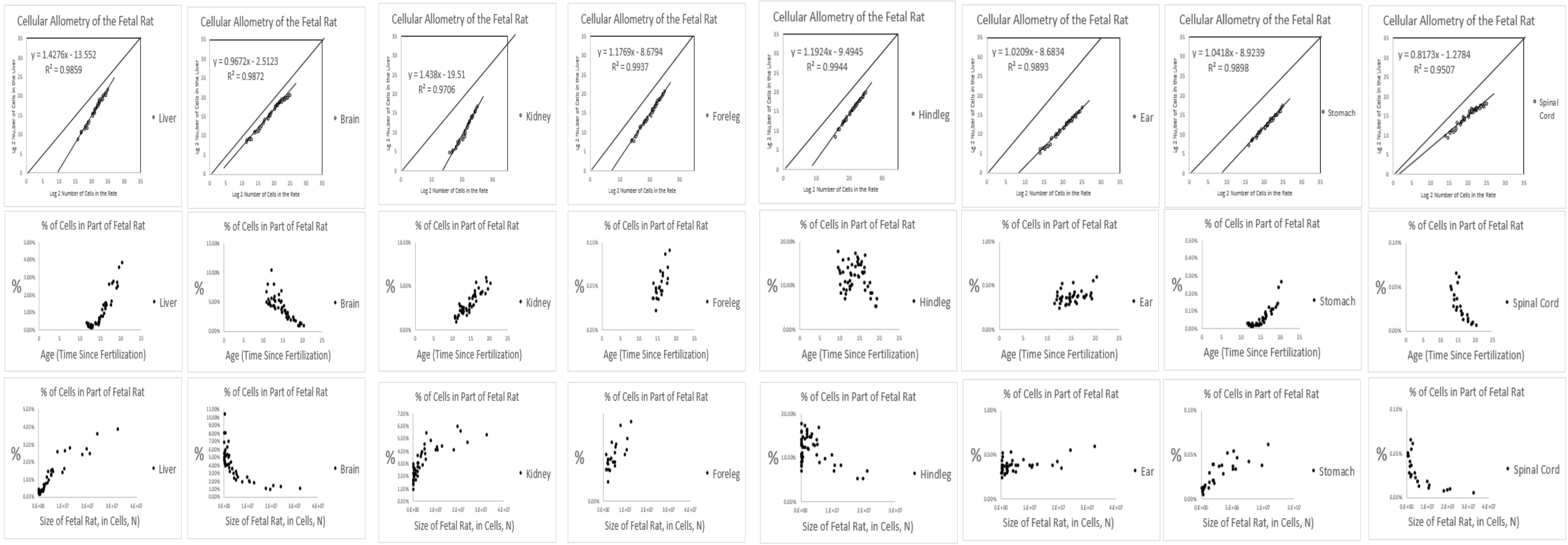


| | % | age | Size of Embryo in Cells, N |
|---|---|---|---|
| first value | 0.46% | 12 | 5.E+04 |
| last value | 3.89% | 20 | 3.E+07 |
| ; increase absolute | 3.43% | | 3.E+07 |
| % increase relative | 8.51 | | 589 |
| te increase relative | 5,012 | | |

| | % | age | Size of Embryo in Cells, N |
|---|---|---|---|
| first value | 6.92% | 11 | 7.E+04 |
| last value | 1.10% | 20 | 3.E+07 |
| % decrease absolute | -5.82% | | 3.E+07 |
| % decrease relative | 6.31 | | 5.E+02 |

| | % | age | Size of Embryo in Cells, N |
|---|---|---|---|
| first value | 1.70% | 11 | 7.E+04 |
| last value | 5.37% | 20 | 3.E+07 |
| % increase absolute | 3.67% | | 3.E+07 |
| % increase relative | 3.16 | | 5.E+02 |
| absolute increase relative | 1,549 | | |

| | % | age | Size of Embryo in Cells, N |
|---|---|---|---|
| first value | 0.04% | 14 | 1.E+06 |
| last value | 0.21% | 20 | 3.E+07 |
| % increase absolute | 0.17% | | 3.E+07 |
| % increase relative | 5.62 | | 29 |
| absolute increase relative | 162 | | |

| | % | age | Size of Embryo in Cells, N |
|---|---|---|---|
| first value | 17.78% | 12 | 5.E+04 |
| last value | 5.25% | 20 | 3.E+07 |
| % decrease absolute | -12.53% | | 3.E+07 |
| % decrease relative | 3.39 | | 589 |

| | % | age | Size of Embryo in Cells, N |
|---|---|---|---|
| first value | 0.30% | 12 | 5.E+04 |
| last value | 0.60% | 20 | 3.E+07 |
| % increase absolute | 0.30% | | 3.E+07 |
| % increase relative | 2.00 | | 589 |
| absolute increase relative | 1.175 | | |

| | % | age | Size of Embryo in Cells, N |
|---|---|---|---|
| first value | 0.03% | 12 | 5.E+04 |
| last value | 0.27% | 20 | 3.E+07 |
| % increase absolute | 0.24% | | 3.E+07 |
| % increase relative | 8.13 | | 589 |
| absolute increase relative | 4.786 | | |

| | % | age | Size of Embryo in Cells, N |
|---|---|---|---|
| first value | 0.05% | 12 | 5.E+04 |
| last value | 0.01% | 20 | 3.E+07 |
| % decrease absolute | -0.04% | | 3.E+07 |
| % decrease relative | 0.13 | | 589 |

**Supplement-Figure-I.29**: ***Binary Cellular Allometric Growth Analysis* of tissues, organs, and anatomical structures of the rat embryo**. Growth of ***Body Parts*** of the rat embryonic liver, brain, kidney and foreleg, in units of numbers of cells, in comparison to the number of cells in the embryo as a whole, are shown on log-log graphs. Note how the fraction of cells comprising in the liver kidney and foreleg increases as the number of cells in the embryo increase with development, while the fraction of cells comprising the brain decreases.[100]

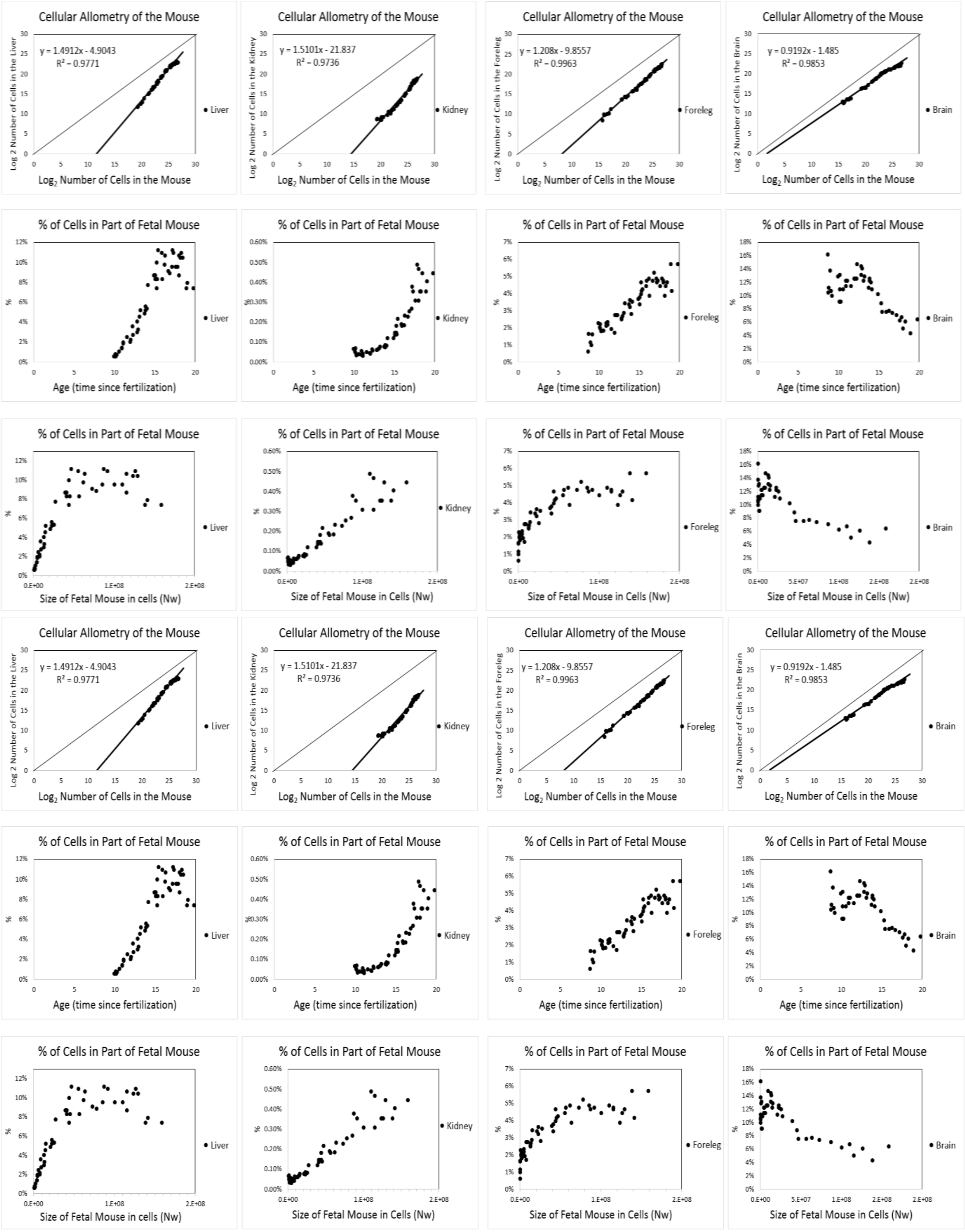


**Supplement-Figure-I.30:** ***Binary Cellular Allometric Growth Analysis*** **of** ***tissues***, ***organs***, and ***anatomical structures*** **of the mouse embryo.**[100]

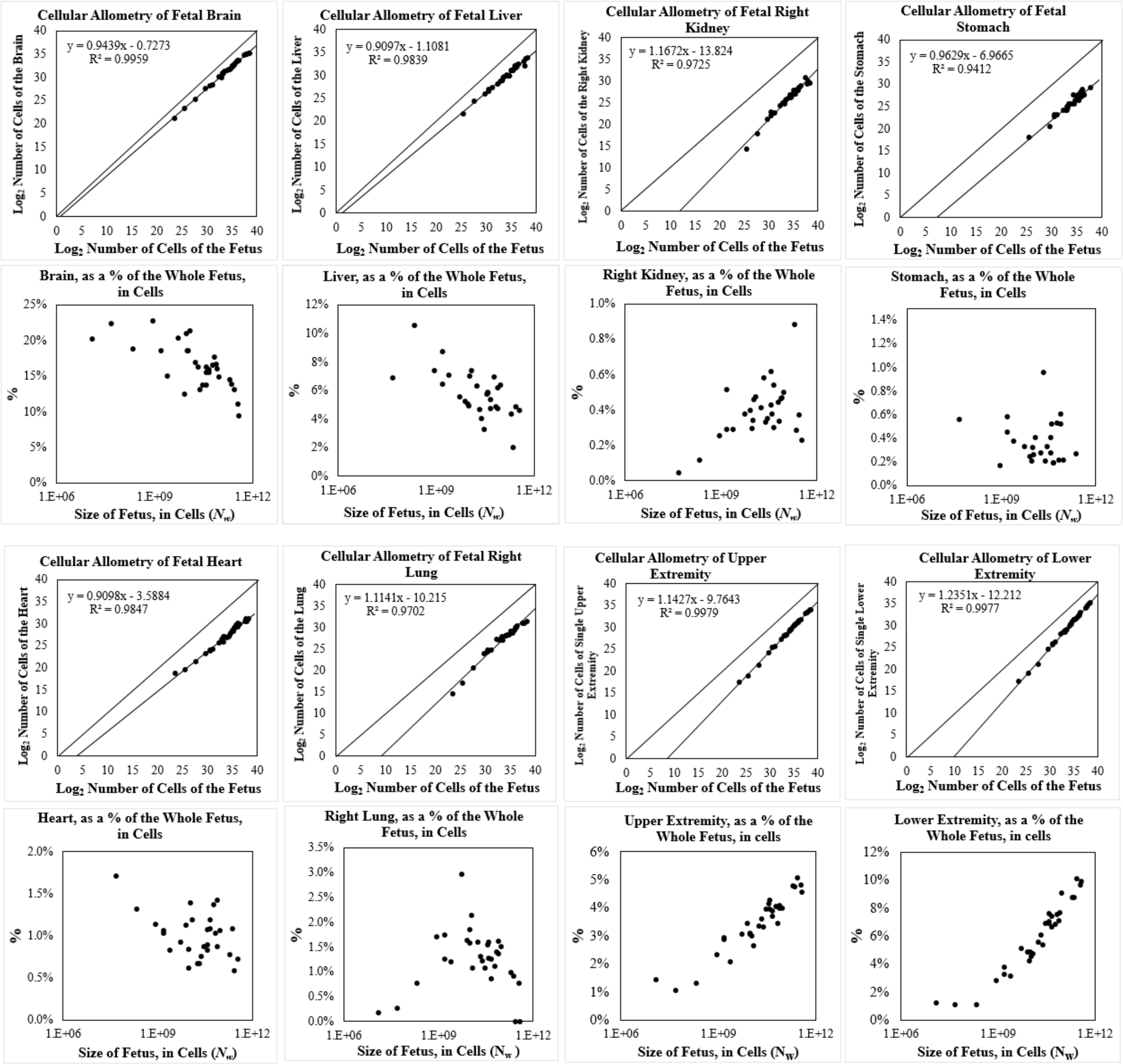


**Supplement-Figure-I.32**: ***Binary Cellular Allometric Growth Analysis* of *tissues*, *organs*, and *anatomical structures* of the developing human fetuses, from autopsy data, of individual fetuses**.[100] For additional human relative growth data, see **Supplement-Figure-I.31**

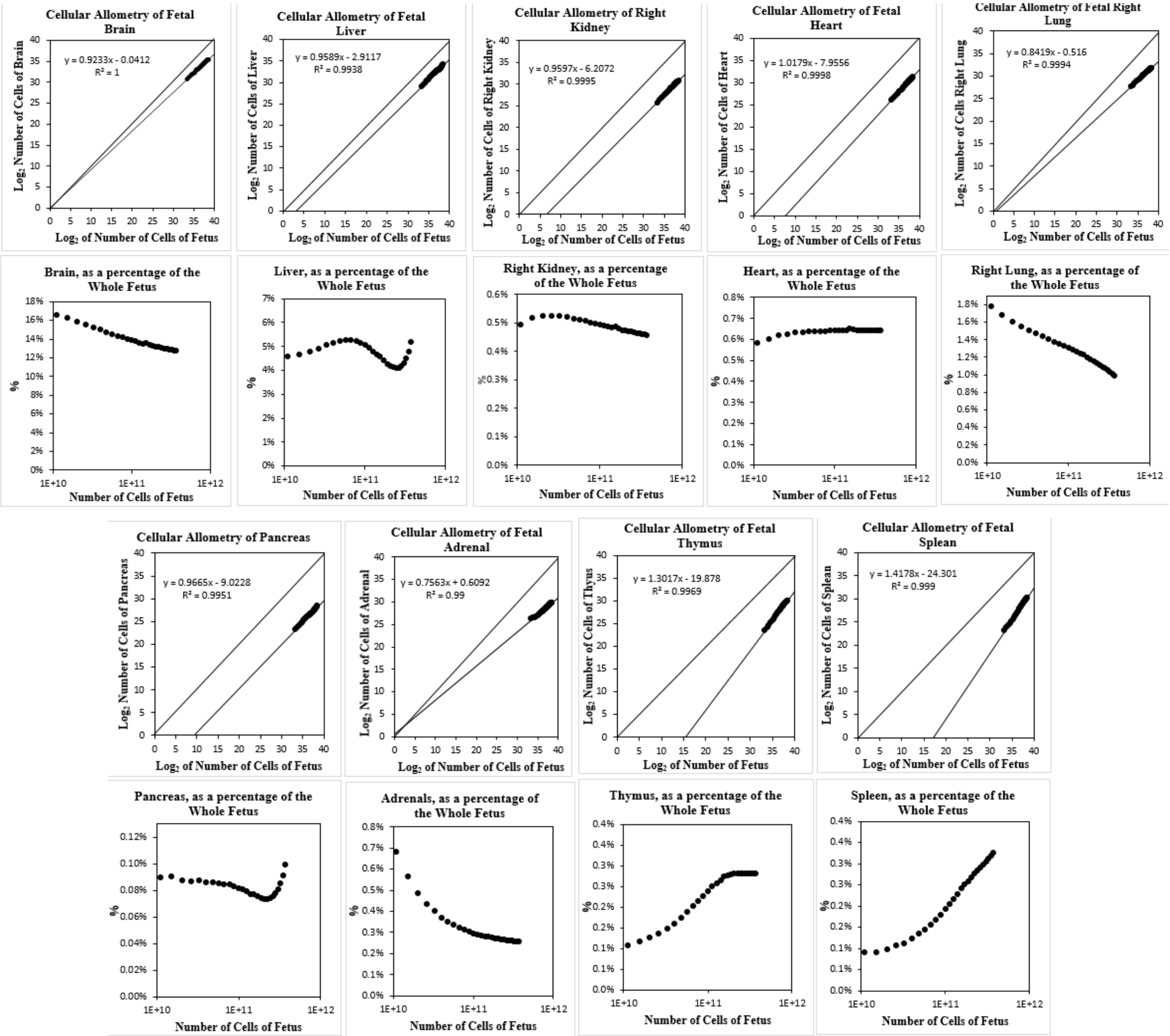


**Supplement-Figure-I.32**: ***Binary Cellular Allometric Growth Analysis* of *tissues*, *organs*, and *anatomical structures* of the developing human fetuses, from autopsy data, averaged values**.[94,128] For additional human relative growth data, see **Supplement-Figure-Z I.** A15[100,129] For additional human relative growth data, see **Supplement-Figure-I.30**

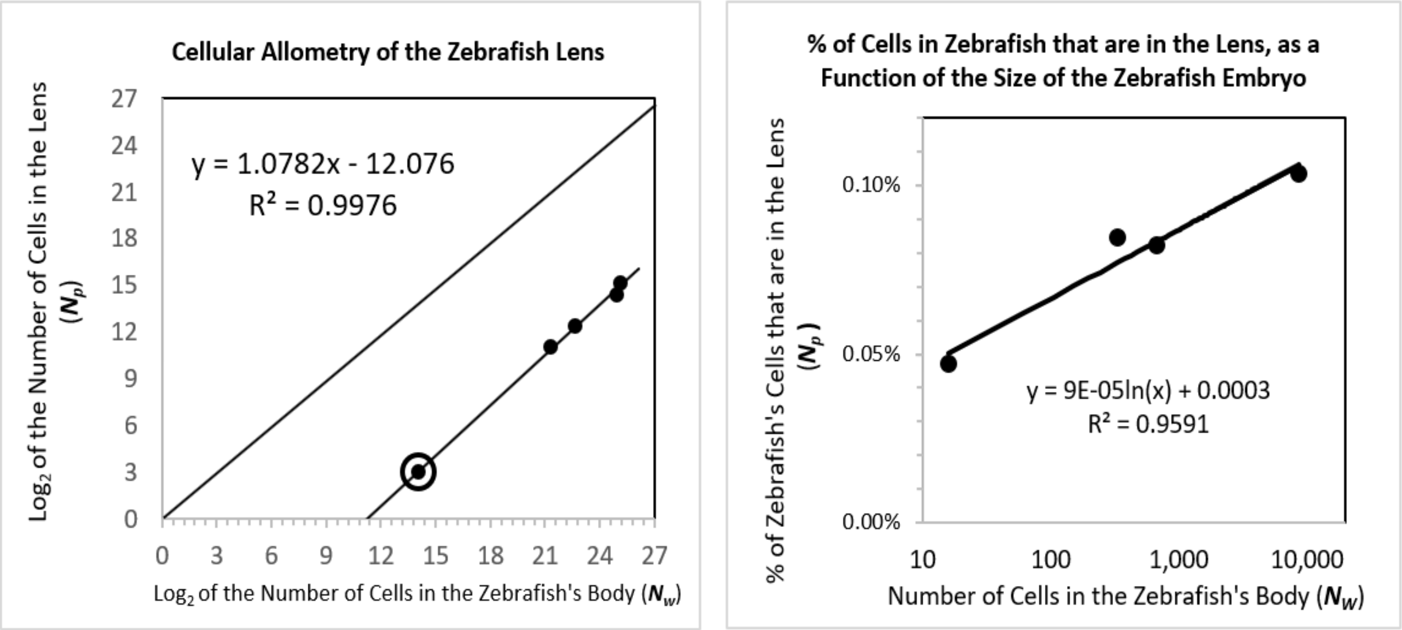


**<u>Supplement-Figure-I.33</u>:** ***Binary Cellular Allometric Growth Analysis*** **of the Zebrafish Lens.**[89,90,91,92]

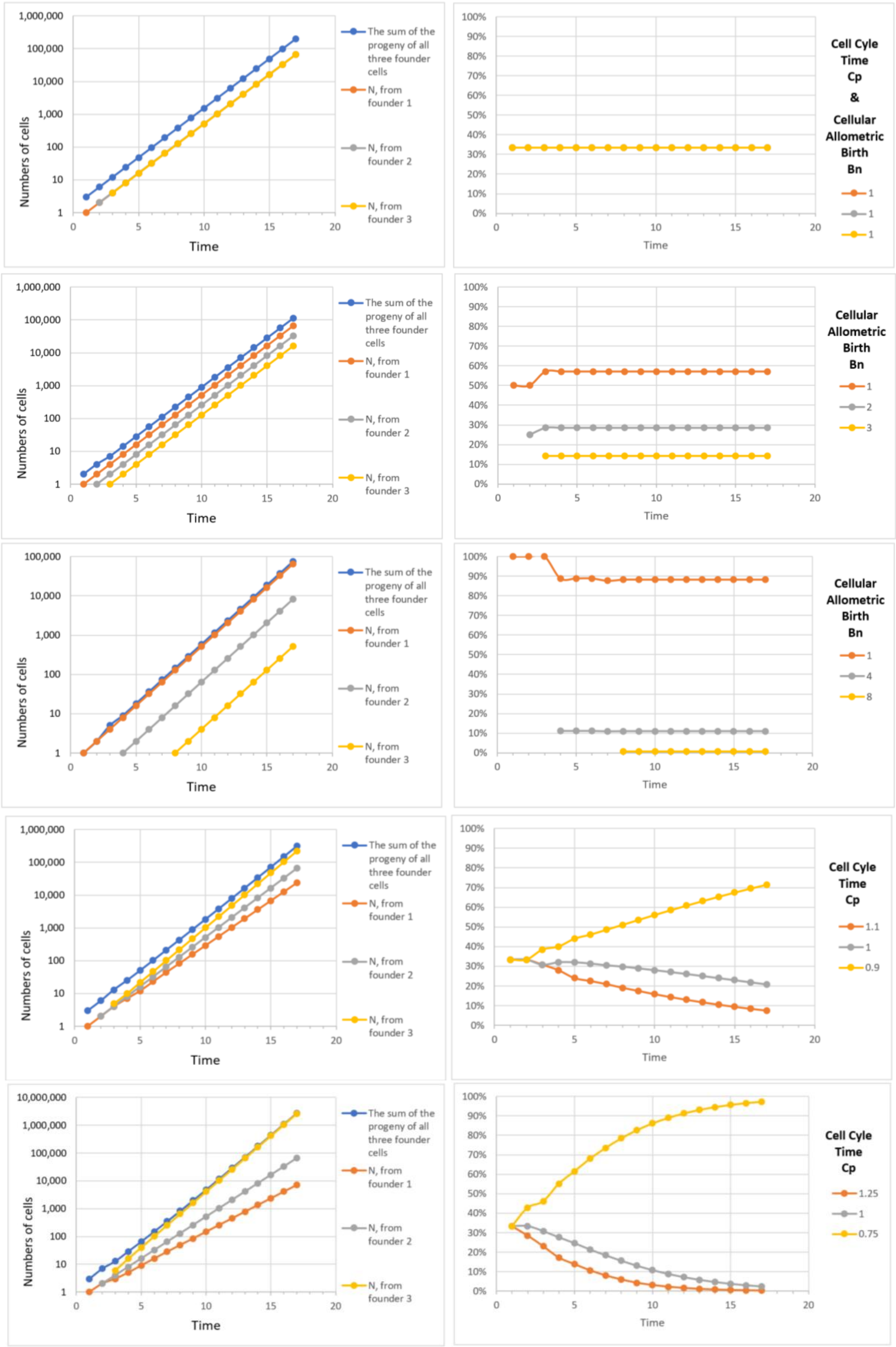


**Supplement-Figure-I.34**: **Calculations for an idealized part of the body that is made from 3 *Founder Cells*, arising early in development.**

***Binary Cellular*** Estimation of ***Fetal Weight*** from ***CRL*** Measurements Early in Pregnancy

The ***Crown to Rump Length Binary Cellular Fetal Weight*** (***CRL-BC EFW***) ***Equation***

To develop a method for estimating ***Fetal Weight***, early in pregnancy, we began by using ***Excel Numerical Integration*** of the ***Binary Cellular Universal Growth Equation*** (#2), incorporating the ***Empirical Ripples in Growth***, to calculate the number of cells, at each point in time (in units of ***Cell Cycle Times***, $c_w$), and from this, ***Fetal Weight***, at each point in time, from conception (***t*=0**) onward, while, for each iteration, excel is also used to calculate the ***Crown to Rump Length*** (***CRL***), from each of those ***Fetal Age*** values, from the ***CRL from Fetal Age Equation*** (#7b).

***Allometric Analysis*** was then used to examine the relationship of the ***Crown to Rump Length*** (***CRL***) and ***Fetal Weight*** values generated by this method, that is, by comparing the log of ***Crown to Rump Length*** (***CRL***) to the log of ***Fetal Weight*** (**Supplement-Figure-I.35**). Regression leads directly to this expression:

***Fetal Weight*=0.002751693*(CRL)^2.3103** (14d)

We call this expression the ***Crown to Rump Length Binary Cellular Fetal Weight*** (***CRL-BC EFW***) ***Equation*** (#14d) (Brown Line in **Supplement-Figure-I.36**). This expression, which conforms quite closely to the values of ***Fetal Weight***, and ***Crown to Rump Length*** (***CRL***) generated by the ***Excel Numerical Integration*** (Blue Line in **Supplement-Figure-I.36**), provides us with a way to estimate ***Fetal Weight*** early in pregnancy from and ***Crown to Rump Length*** (***CRL***) ***Ultrasound Measurement***.

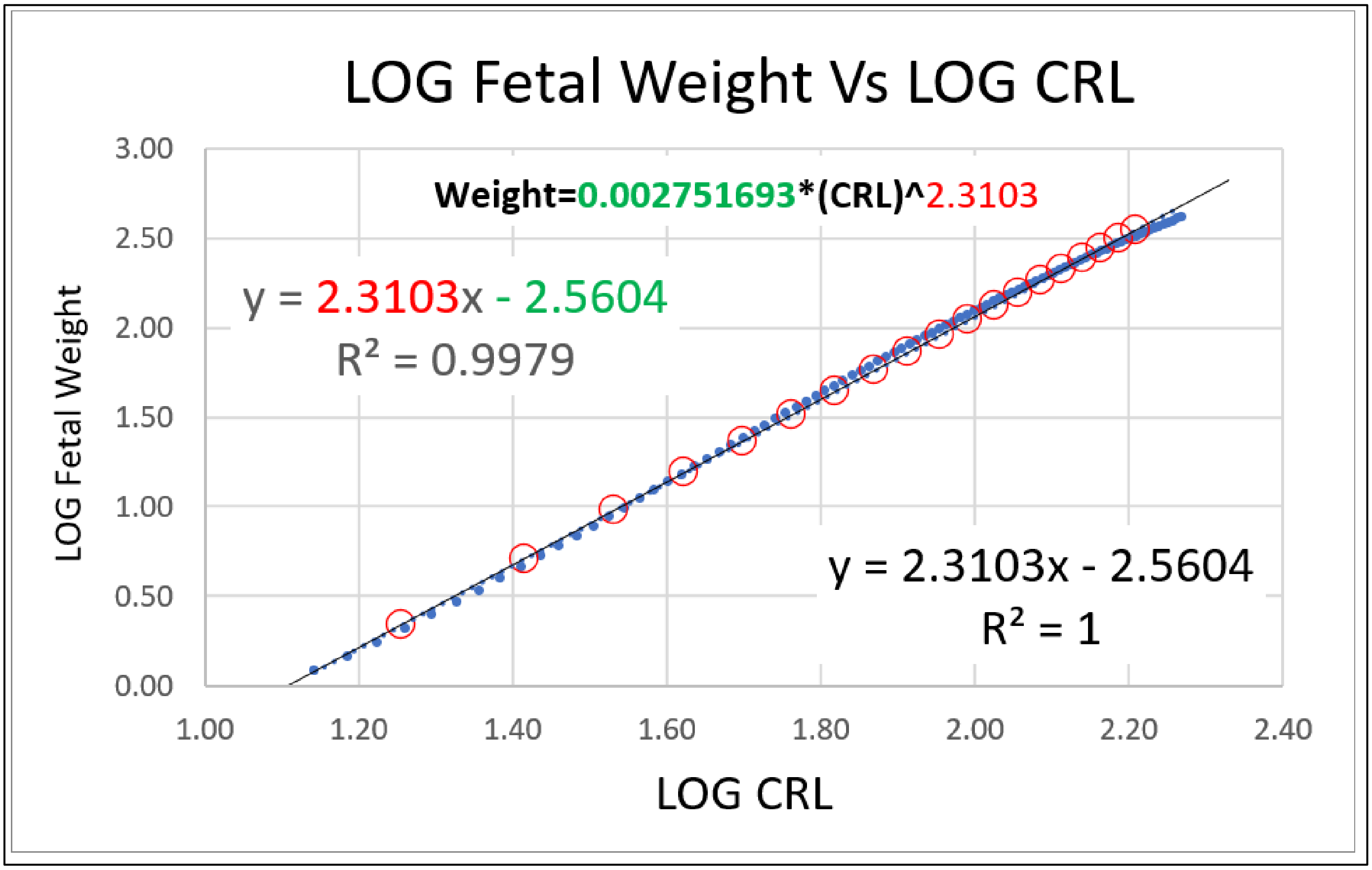


**Supplement-Figure-I.35** ***Crown to Rump Length Binary Cellular Fetal Weight*** (***CRL-BC EFW***) ***Equation*** (#14d)

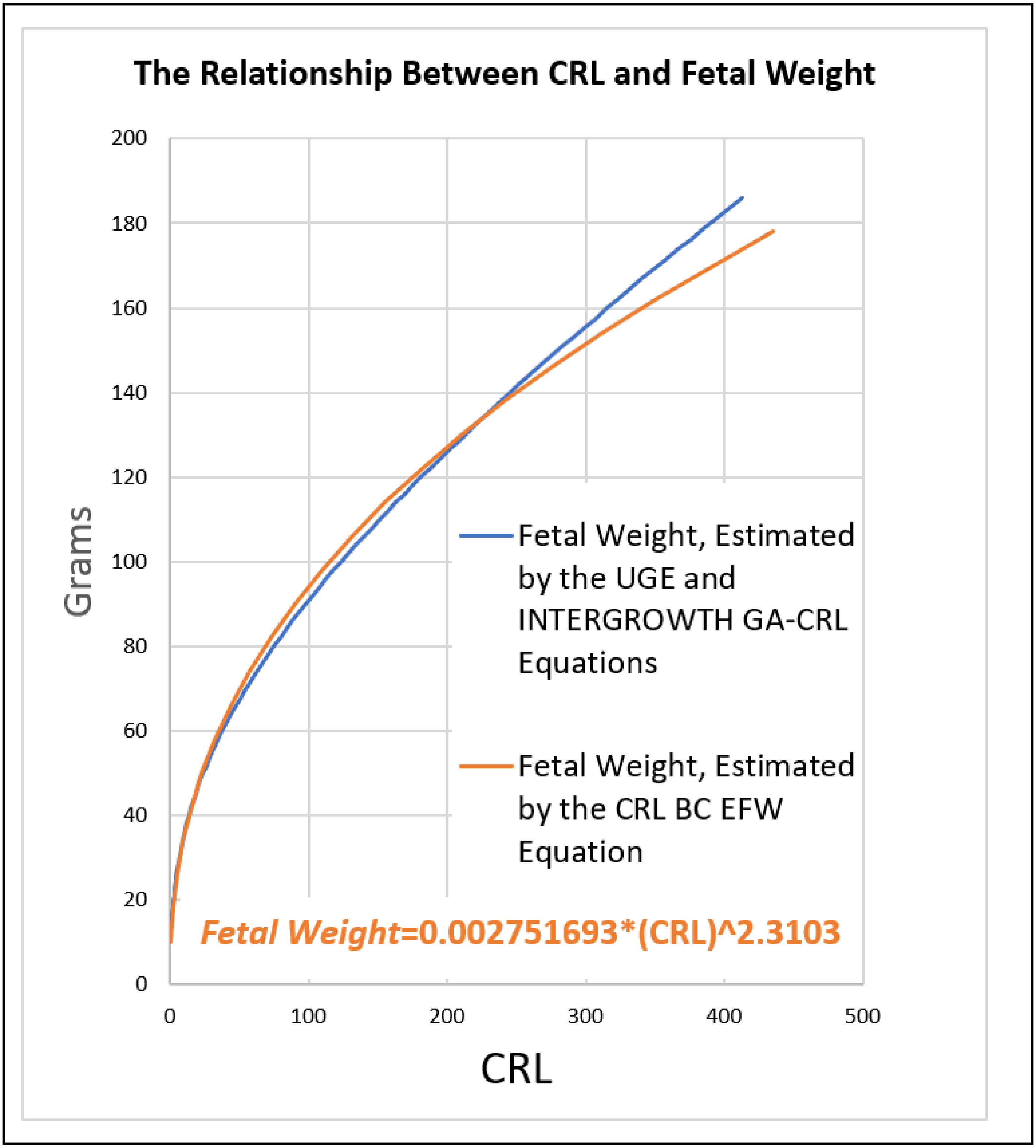


**<u>Supplement-Figure-I.36</u>** The Relationship Between Crown to Rump Length (CRL) and Fetal Weight, as captured by the ***Crown to Rump Length Binary Cellular Fetal Weight*** (***CRL-BC EFW***) ***Equation*** (#14d).

The ***Binary Cellular Mathematics*** of ***Fetal Age***

Terminology for ***Fetal Growth***: ***Age***, ***Gestational Age***, ***Conceptional Age, Fetal Age***.

***Gestational Age***, the time since the last menstrual period, has long been the most common way for thinking about the age of the human fetus. Another commonly used term is ***Conceptional Age***, but the date of conception is not always known. As we shall be examining fetal, childhood and adult growth data, as well as growth data of animals such as mollusks, tunicates, and nematodes, which don't have a fetal period, we shall be using the simplest term, ***Age***, with modifiers to indicate the period examined, which, for our work here, will be ***Fetal Age***. Here we shall calculate ***Fetal Age*** from ***Crown to Rump Length*** (***CRL***) ***Ultrasound*** measurements.

Estimation of ***Fetal Age*** from ***Crown to Rump Length*** (***CRL***): The ***CRL Fetal Age Equation***

To calculate ***Fetal Age***, we have adopted and adapted the INTERGROWTH Equation for estimating ***Fetal Age*** from ***Crown to Rump Length*** (***CRL***)[130]:

$$\textit{Fetal Age} = (40.9041+(3.21585*(((\textit{CRL})\hat{}(0.5)))+(0.348956*(\textit{CRL})))) - 14 \qquad (7a)$$

We call this expression, the ***CRL Fetal Age Equation*** (#7a). The 14 days subtracted from the INTERGROWTH value converts the values from the ***Gestational Age*** to ***Fetal Age***.

Estimation of ***Crown to Rump Length*** (***CRL***) from ***Fetal Age***: ***CRL from Fetal Age Equation***

The inverse for the mean of ***Crown to Rump Length*** (***CRL***), as a function of ***Fetal Age***, is also provided by the INTERGROWTH group:

$$\textit{CRL} = -50.6562 + (0.815118 * (\textit{Fetal Age} + 14)) + (0.00535302 * (\textit{Fetal Age} + 14)^2) \qquad (7b)$$

We call this expression, the ***CRL from Fetal Age Equation*** (#7b). Again, the 14 days added to the ***Fetal Age*** values converts the values from the ***Gestational Age*** metric of INTERGROWTH.

The ***Binary Cellular Fetal Growth Analysis*** of Individual Human Fetuses from Conception Onward

With the ***CRL Fetal Age Equation*** (#7a) for estimating the date of conception, and the ***Binary Cellular Universal Growth Equation*** (#2), we can visualize the growth that would be expected for an average fetus from its conception onward (**Supplement-Figure-I.37**). We call this approach to the examination of the growth of individual fetuses ***Binary Cellular Fetal Growth Analysis***, and the images that capture this information ***Binary Cellular Fetal Growth Charts*** (**Supplement-Figure-I.39**). As we shall see, such a ***Binary Cellular Fetal Growth Analysis*** can make it possible to characterize the growth pathways of individual fetuses into newborns, as well as to gain insight into how most fetuses become normal size at birth, while others emerge as small newborns, and others emerge as large newborns.[6] We shall be able to carry out the ***Binary Cellular Fetal Growth Analysis*** of individual fetuses once we have developed the methods for transforming ***Fetal Ultrasound Measurements*** into ***Fetal Size*** estimates, as we shall do next.[7]

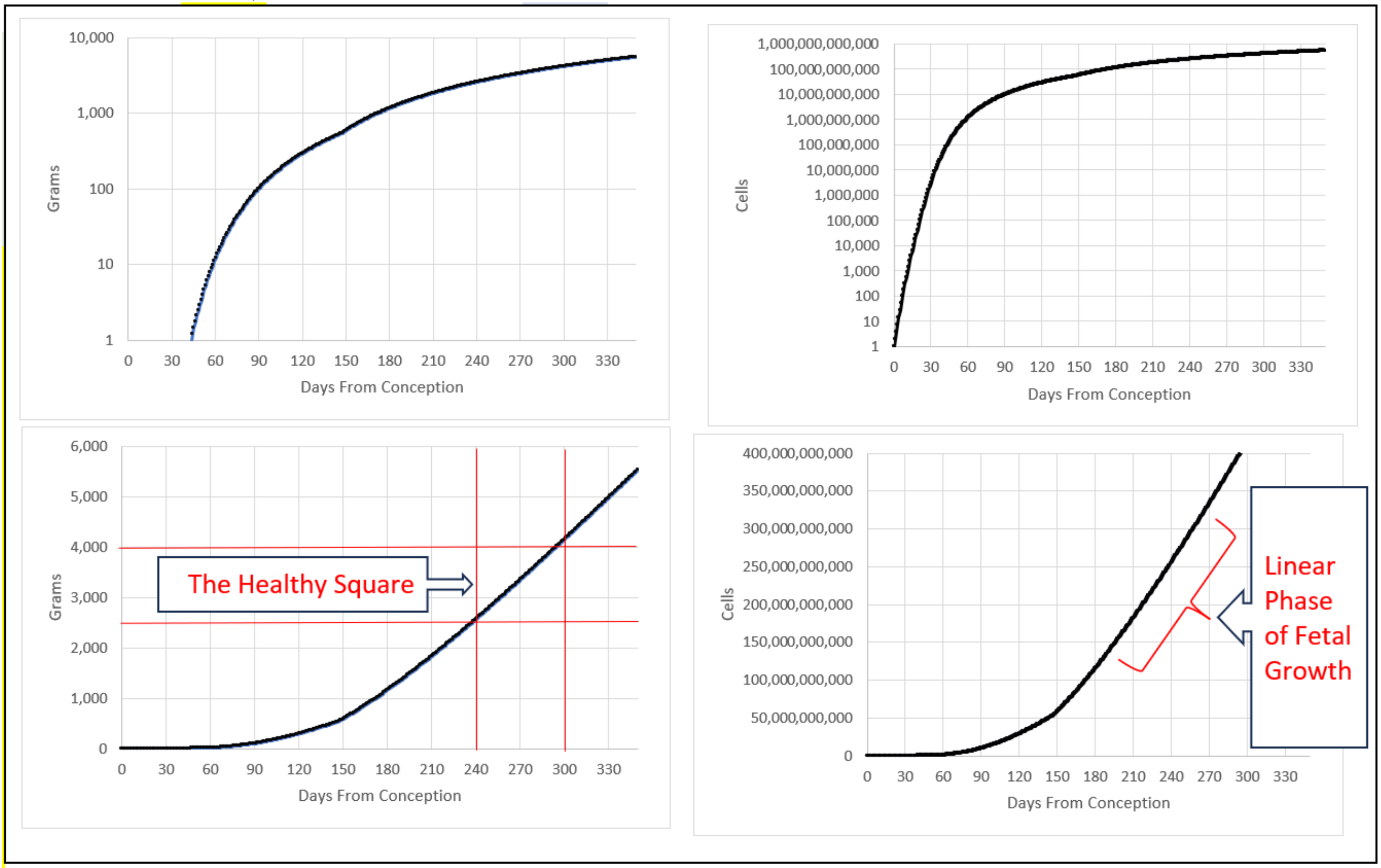


**Supplement-Figure-I.37**. A ***Binary Cellular Fetal Growth Chart***, showing the ***Binary Cellular Universal Growth Equation*** (#2), rooted on the date of conception, estimated by the ***CRL Fetal Age Equation*** (#7a).

***Binary Cellular Analysis* of *Drosophila Growth and Development***

To carry the growth and development of drosophila, we assembled values for the numbers of cells in the fly as a whole, $N_w$, from fertilization onward, and in the fly's various parts, $N_p$ (**Supplement-Figure-**I.38). Many of these parts grow from small structures, which contain few cells, including discs, histoblast nests, and pole cells (which will give rise to the fly's germ cells), as we have found in many studies scattered though the literature over the past 80 years.[131-145]

Numbers of cells in the fly as a whole, $N_w$.

Measuring the size of the body in units of numbers of cells, $N_w$, and gauging growth from these values, is complicated by the quirky nature of drosophila growth, which is marked by periods of massive cell suicide, followed by regrowth from small structures, as well as by the mass polyploidization that goes on as flies develop. Nonetheless, the relationship between cell number, $N_w$, and age, $t$, has been quite accurately measured for the first 6000 cells by Zoloka and Erk (1975).[131] As we shall see, for our first look, we don't require precise values for $N_w$ beyond this point, but rough estimates can be made from measured of body mass by Sawala and Gould[132] (who kindly provided raw data) and from DNA content.[133]

Numbers of cells in the body parts of the fly, $N_p$.

There has been considerable uncertainty as to whether parts of the fly's body arise from single ***Founder Cells*** (clones) or multiple ***Founder Cells*** (polyclones)[134] addressed in many studies of challenging effort that used ***Cell-Heritable*** making techniques, such as gynandromorphs, or mitotic recombination, to track these parts to their origins. We shall return below to this area of scholarship, and its relationship to the work described here, once we have presented the findings based on the raw numbers of cells in the fly as a whole, $N_w$, from fertilization onward, and in its various parts, $N_p$, and time, based on cell counting studies.

Many of the body parts of flies have been observed from small numbers of cells (**Supplement-Figure-**I.38). Both the spiracular anlage and the pole cells (which form the germ cells)[131196] have been seen from 2 cells onward. Genital imaginal discs have been observed from 2 cells[135,136], while other discs have been observed from as few as 12 cells. Superb data on the numbers of cell is imaginal discs and histoblast nests have been reported in studies by Madhaven and Madhaven (1980),[137] Madhavan and Schneiderman (1977),[138] Graves and Schubiger (1982),[139] Bate and Arias (1991),[140], and Bryant and Levinson (1985).[141] These researchers reported that the number of cells, $N_p$, in antenna discs (33 cells to 48 cells), eye discs (40 cells to 250 cells), wing discs (24 cells to 4500 cells), haltere discs (12 cells to 63 cells), prothoracic leg discs (36 cells to 54 cells), mesothoracic leg discs (42 cells to 85 cells), metathoracic leg discs (45 cells to 89 cells), over various points in time from as early as 13 hours until 120 hours after fertilization. For histoblast nests, cell numbers, $N_p$, have been counted from as small as 4 cells. The spiracular anlage has been tracked from 2 cells.[138] From 118 to 138 hours of life we have data on the three anterior dorsal histoblast nests from 15 to 466 cells, on the three posterior dorsal histoblast nests from 5 to 110 cells, and on the three ventral nests, with numbers ranging from 11 to 284 cells.[138] Verma and Cohen[142] have provided videos that show the growth of these histoblast nest back to as few as 4 cells at the beginning of their videos; one wonders what would have been seen if one turned on the camera before then, and such a simple data collection exercise cries out for experimental examination. Furthermore, these videos make clear that nest cells increase in number by cell division rather than by recruiting unrelated cells, and that simply starting the videos earlier would allow one to see the ***Founder Cells*** that give rise to these structures. The population of pole cells, which gives rise to the germ cells of the fly, have been counted back to 2 cells by Rabinowitz (1941)[143] while Zoloka and Erk (1975)[131] have counted the number pole cells from 12 cells, together with the number of cells in embryo as a whole from the first cell until ~6000 cells.

<u>Log-Linearity of the increase in the numbers of cells in various body parts of the fly, $N_p$.</u>

When examined on log graphs of cell number, $N_p$, versus time, $t$ (where $t$=**0** when $N_w$=**1**, **Supplement-Figure-**I.38), most of the parts of the fly's body display striking exponential growth, which appears as straight-rows of datapoints on log-graphs, below 10,000 cells, indicating that they aren't much modified by a ***Mitotic Fraction*** until this size point. The exponential lines of pole cells, and disc cells, point back to ***Cellular Allometric Births in Time***, $B_T$, close to the first moments of fly life. The exponential lines of growth for the histoblast nests and the spiracular anlage point back to ***Cellular Allometric Births in Time***, $B_T$, to between 100 and 125 hours of development. Zolokar's and Erk's data[131] on pole cells numbers show a remarkable log-log linearity by the ***Cellular Allometric Growth Equation*** by ***Cellular Allometric Batch Growth Analysis*** ($r^2$=**0.96**, not shown).

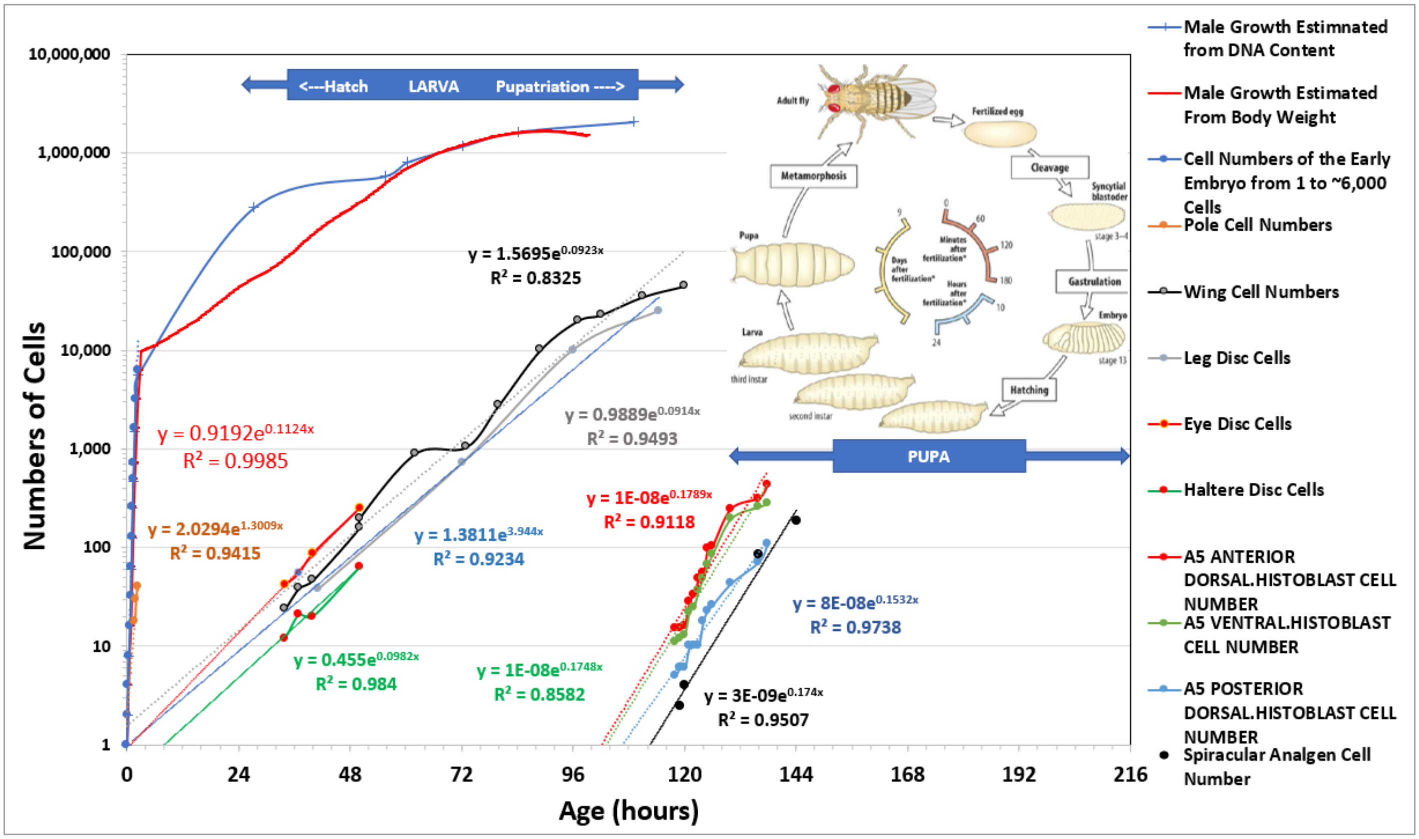


**<u>Supplement-Figure-I.38</u>**

There are dramatic differences in the slopes of these exponential lines, indicating that these body parts have a variety of ***cell cycle times***, $c_p$, which account for changes in the relative size of these parts that occur as the fly matures (**Supplement-Figure-**I.38). These differences in slope lead to dramatic changes in the relative sizes of each of these body parts.

The remarkable log-linearity of growth below 10,000 cells, followed by the beginning of the reduction in exponential log-linearity can be seen more carefully when we display just a single imaginal disc on both logarithmic (**Supplement-Figure-**I.39) and linear (**Supplement-Figure-**I.40) scales. The reduction in the proliferative potential of discs and histoblast nests above 10,000 cells in size would allow their growth at these larger sizes to be amenable to the ***Mitotic Fraction Method***, another possibility worthy of experimental and computational examination.

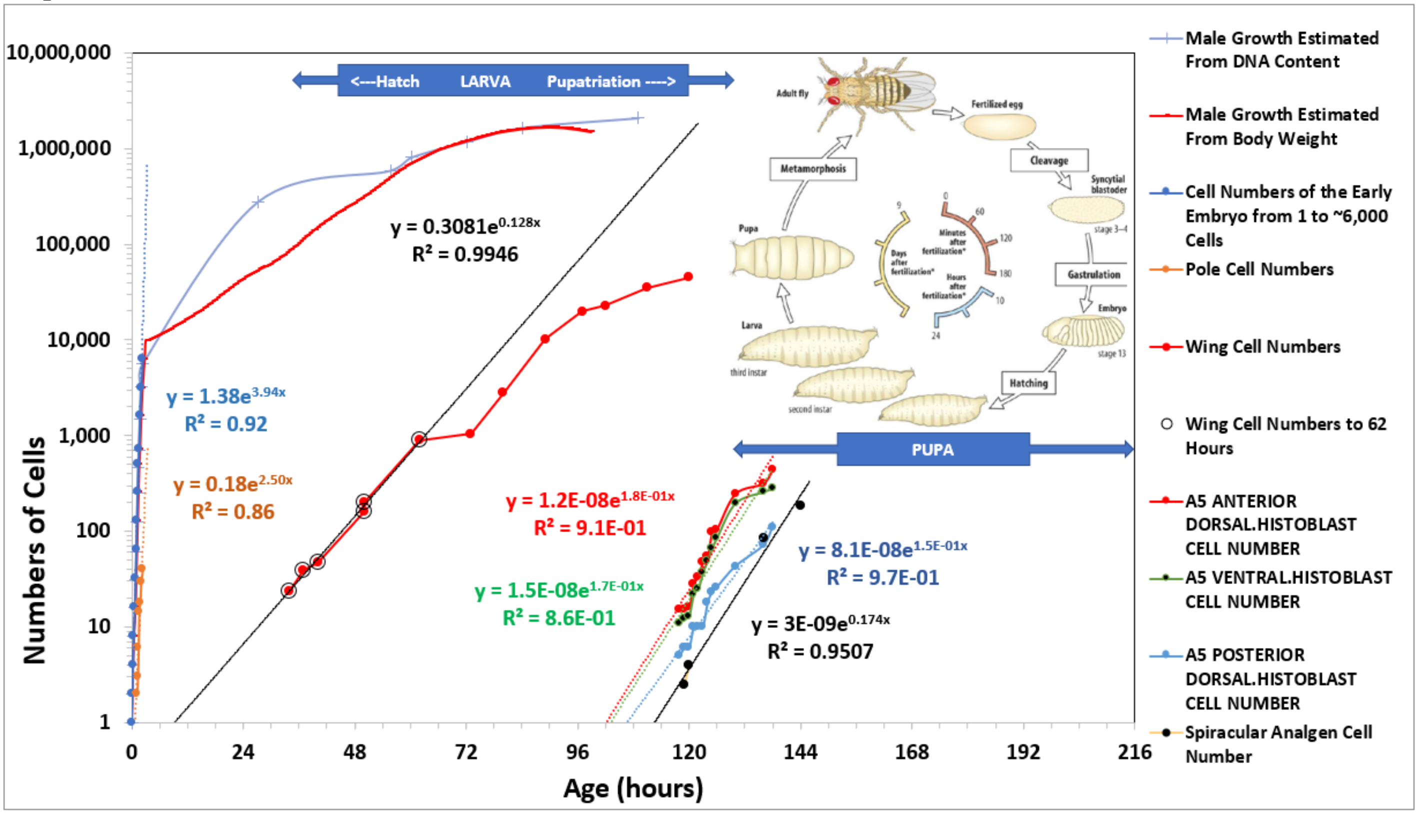

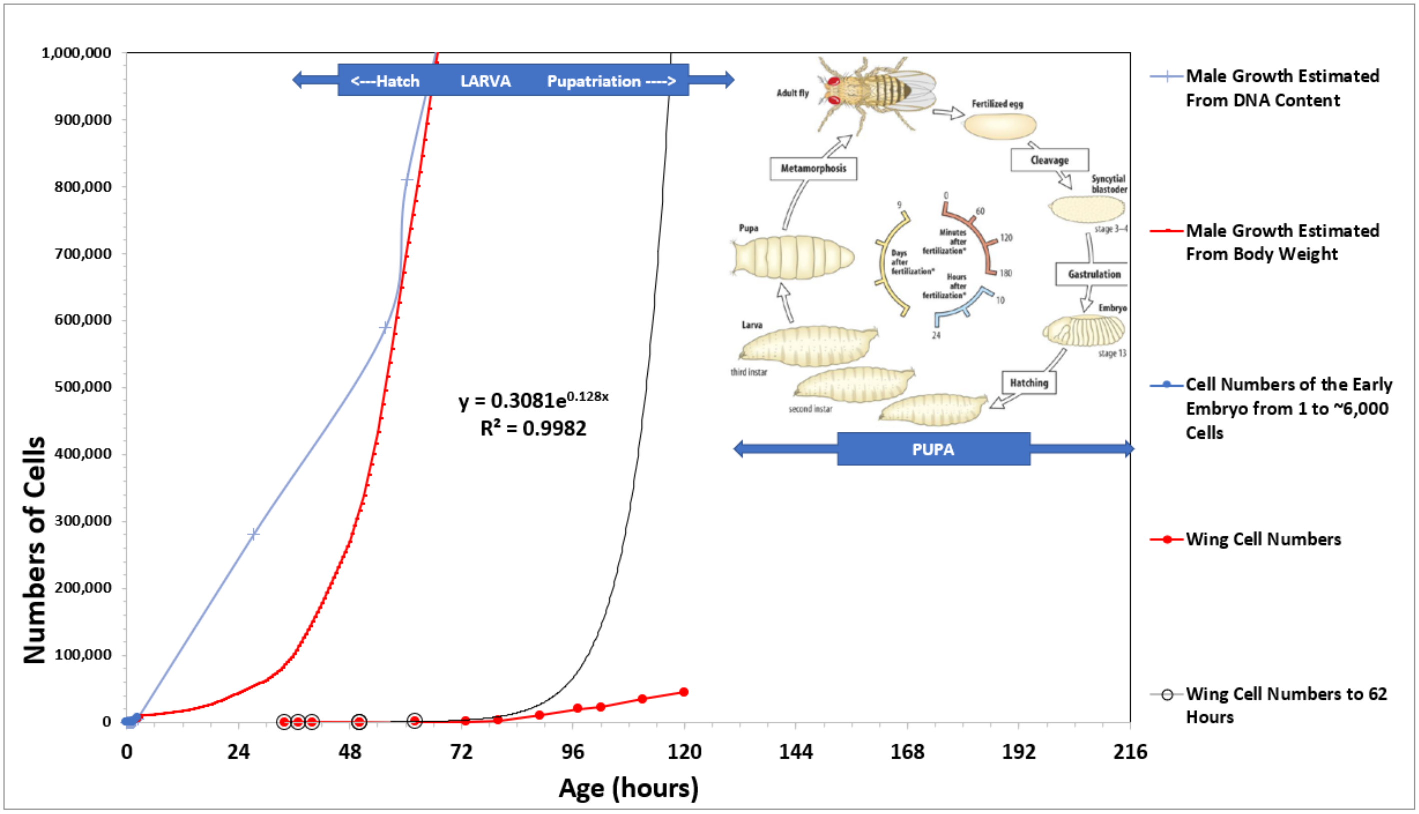


**Supplement-Figure-I.39**

<u>Clones Within Clones</u>

<u>Drosophila</u>

As can be seen in Equation #722 below, if each of two parts of the body are seen by ***Cellular Allometric Growth Analysis*** to conform to the ***Cellular Allometric Growth Equation*** (#7), when each is compared with the body as a whole, they will also conform to the ***Cellular Allometric Growth Equation*** (#7) when compared with each other. An actual example of this can be seen in **Supplement-Figure-**I.40-left, in which we display the number of cells in the anterior part of the drosophila wing, $N_{p\text{-}a}$, in comparison to the number of cells in the wing as a whole, $N_{p\text{-}w}$. These values were generated from measurements on the area of the whole wing disc, and of its anterior part, that have been collected by Parker and Shingleton,[144] translated into likely cell numbers, $N_{p\text{-}a}$, and $N_{p\text{-}w}$, with cell number per area data derived by Ulrike.[145]

These log-log comparisons of $N_{p\text{-}a}$ (number of cells in the anterior part of the drosophila wing), and $N_{p\text{-}w}$ (number of cells in the wing as a whole), shown in **Supplement-Figure-**I.39-left reveal that the ***Cellular Allometric Growth Equation*** captures the growth of the anterior part of the wing, pointing to a ***Cellular Allometric Birth***, $B_N$, growing from when the wing as a whole was 8 cells in size ($B_{NI}=2^3=8$), should the anterior part have arisen from a single ***founder cell***. Furthermore, the ***Cellular Allometric Slope***, $S_{Np}$, is greater than **1**, indicating an increase in speed of the ***cell cycle time***, $c_p$, of the cells in the anterior part of the wing, thus accounting for the progressively increasing relative size of the anterior part of the wing in relationship to the wing as a whole (**Supplement-Figure-**I.39-right). We call this process "***clones within clones***', a mechanism which provides animals a way to create functional diversity.

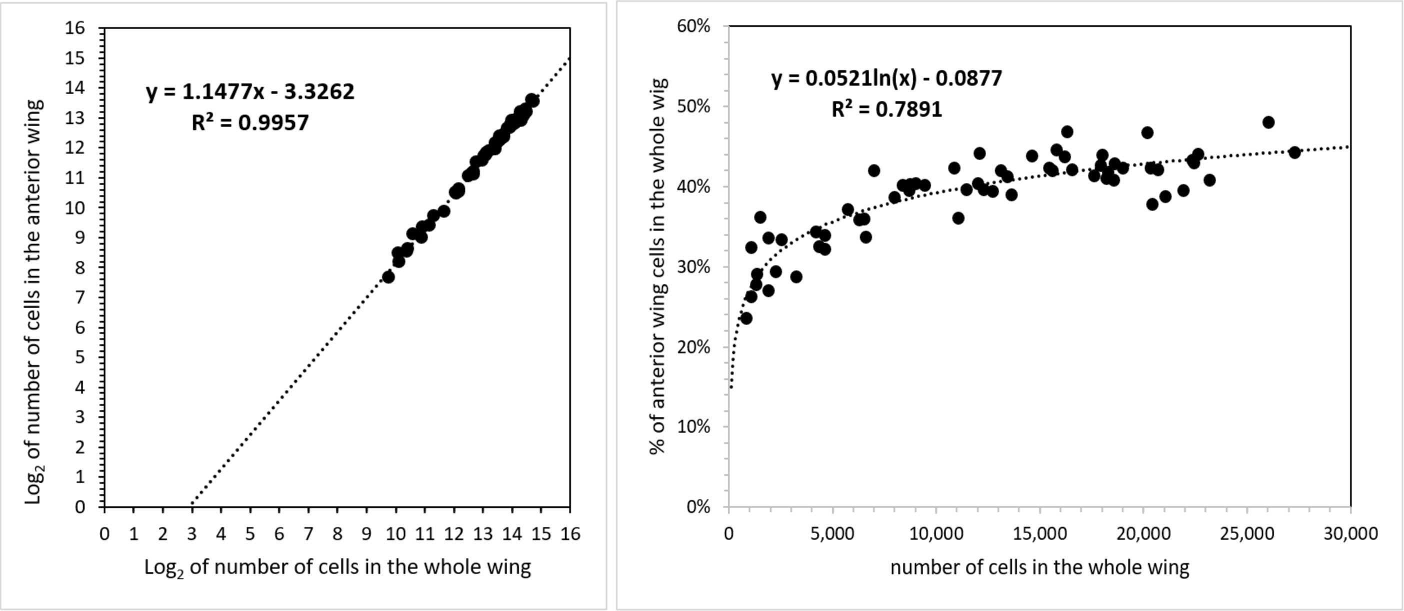


**Supplement-Figure-I.40**

Whether the anterior part of the wing grew from a single ***founder cell***, at the ***Cellular Allometric Birth***, $\boldsymbol{B_N}$, or from more than **1** ***founder cell***, after the ***Cellular Allometric Birth***, $\boldsymbol{B_N}$, cannot be known from the data we have, since the values for $\boldsymbol{N_{p\text{-}a}}$, and $\boldsymbol{N_{p\text{-}w}}$ only point down to that origin at when the wing as a whole was 8 cells in size ($\boldsymbol{B_{N1}}$=$\mathbf{2^3}$=**8**). Nonetheless, this poses an eminently answerable question, which could be settled by light sheet microscopy*, again providing motivation for experiment.

These observations are also strikingly complementary to the long appreciated finding the anterior and posterior parts of the fly's wing are clonally sperate entities, called ***compartments***.[192]

Mammals

The formation of ***clones within clones*** occurs in many instances of embryonic development of many animals. For example, the melanocytes of the skin of mice appear to be derived from 34 ***Founder-Cells*** (17 on each side of the body) created around the $12^{th}$ day of life.[146] We also see ***clones within clones*** at work in the immune system, where ***clonal selection*** selects individual cells and expand them up to large populations of cells through the action of antigens stimulating cell division by attaching to antibodies or T-cell receptors at the cell surface. **,[147] Similar processes may well occur in other systems, such as in the production of plasma proteins by the liver, in which each of the hundred-or-so blood proteins produced by the liver appears to be made by separate cells, in separate clones, that arise as patches continuously throughout life.* Anatomically, the creation of such functional diversity by the creation of ***clones within clones*** has been seen time after time, such as in the clones that form the crypts of the intestine[148], the sequential clones of spermatocytes that produce the spermatozoa[149], and the many anatomical structures that seem to be made of countless subunits, such as in the lungs, the liver, the kidney, and so on. ***Binary Cellular Analysis*** seems poised to make sense of how cells create such modular structures.

Mitotic Recombination

By mitotic recombination induced by irradiation at ~3 hours (we are unaware of any such studies carried out successfully earlier; embryos irradiated before that time point die at a very high rate), Weishchaus and Gehring (1971)[150] have identified that 6 ***Founder Cells*** were marked for the leg disks (3 hours), while Postlethwait and Schneiderman (1971)[151] have identified that 8 ***Founder Cells*** were marked for the leg disks (3 hours), and Bryant (1971) [152] has identified 10 ***Founder Cells*** were marked for the wing disks. (Bryant also saw 11 ***Founder Cells*** were marked for the wing disks when marked at 2 hours, a datapoint that requires confirmation, since the standard errors were quite large, and Weishchaus and Gehring (1971)[150] and others have found high levels of embryonic and cell death by irradiation at this time).

Taken together, these findings reflect discs nests arising from small number of ***Founder Cells***. Are these small number of ***Founder Cells*** the sizes at which these structures are created, or could these body parts arise from even smaller numbers of precursor cells? Several studies have carried out mitotic recombination induced by irradiation at various points in time, and these data for legs, wings, and antenna discs are shown in FIGURE A24 below. As Larsen-Rapport (1986)[153] points out "if the recombination event occurs at the time of X irradiation, the genetically marked cell itself will have to divide (some 15 hr after irradiation) before it can be used for certain purposes like testing for cell lineage restriction to one or another structure" (and our ***Binary Cellular Analysis*** shows imaginal disc doubling time in this range: **Supplement-Figure-**I.38), so we have added the 15 hours. Remarkably, when displayed on log graphs, these number of ***Founder Cells*** marked by X-radiation point back to 1 ***Founder Cell*** early in the development of the drosophila embryo, precisely where the log graphs from cell counts point (**Supplement-Figure-**I.38).

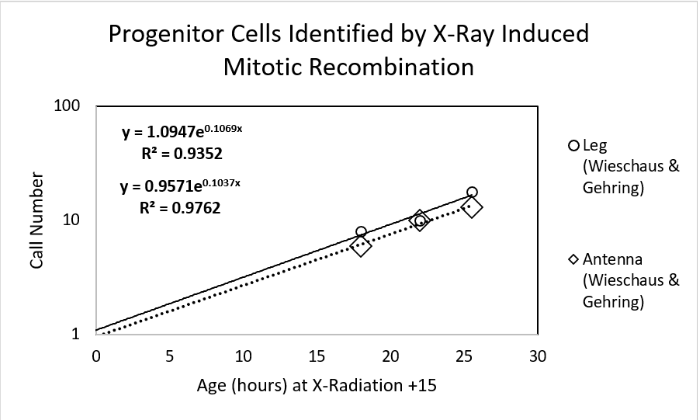


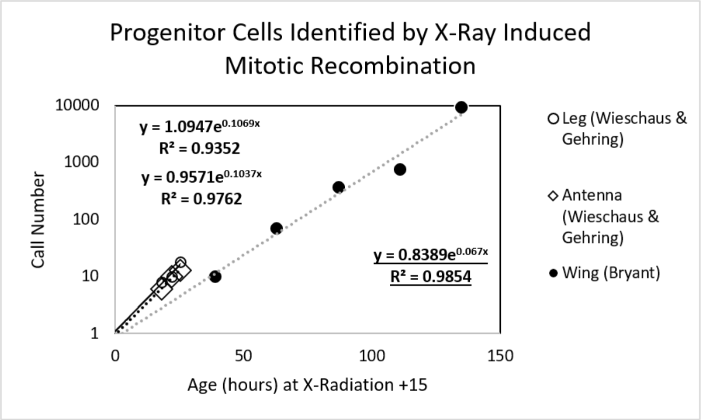


**Supplement-Figure-I.41**

Do these graphs tell us that discs arise from single ***Founder Cells***? We regard the best way to treat these graphs is to consider them as intriguing findings and useful hints for where we should look, by high resolution 4D microscopy, for when these parts might arisen.

Gynandromorph Marking

Using cell marking by gynandromorphs, Madhavan and Schneiderman (1977)[138] have identified that 8-9 ***Founder Cells*** were marked for the antenna disks, Bryant and Schneiderman (1969)[154] have identified that 20 ***Founder Cells*** were marked for the leg disks, Ripo1l (1972)[155] has identified that 47 ***Founder Cells*** were marked for wing disks, Morata and Garcia-Bellido (1976)[156] have identified that 6 ***Founder Cells*** were marked for haltere disks, Schupbach et al (1976) have identified that 9 ***Founder Cells*** that were marked for female genital discs disks and 7 ***Founder Cells*** for male genital discs.

These findings again reflect discs arising from small number of ***Founder Cells***. Unfortunately, these numbers are not precise, ***first*** because their calculation assumes that all parts of the body grow at the same rate, while the ***Binary Cellular Analysis*** of the growth and development of the drosophila body and its parts shown in FIGURES A20-A22 has revealed that they have dramatically different ***Cell Cycle Times*** and rates of growth; and, ***second***, because gynandromorph ***Founder Cell*** number calculations assume that the genetic change always takes place at the same time in the same cells. Indeed, details in the literature make clear that these changes occur over a range of time during early development. For example, Garcia-Bellido and Meriman (1969)[157] suggest that "Gynandromorphs are generally thought to result from nondisjunction or chromosome elimination at the first zygotic division." and Hotta and Benzer (1972)[158] note that "The method of generating Drosophila mosaics used in this paper depends on loss of an unstable, ring-X chromosome … during the first division of the zygote nucleus of a female embryo". However, Zalokar et al (1980)[159] reports that such losses are common in the first division, but also occur in subsequent divisions. As he notes "The proportion of male nuclei ('by loss of the ring-X chromosome') varied from 80.8 to 0.4%, indicating that there must have been more than one loss of the ring-X in most of the eggs and that losses occurred as late as the ninth division". Portin's analysis (1978)[160] of the phenotype of gynandromorphs seems to support this, as he states "two-thirds of the gynandromorphs arise as a consequence of first division loss, whereas one third arise as a consequence of X chromosome loss in a later division.". Of course, these concerns are relevant for an earlier era, and we now have high resolution 4D microscopy, * which can define the originating events with precision. The results of the ***Binary Cellular Analysis*** we have presented here again point us to where we should look.

#### DISCUSSION

***Binary Cellular Analysis*** Provides New Equations That Describe ***Growth*** and ***Development***.

Here we have seen that a consideration of animal and human bodies in units of numbers of cells, ***N***, an approach we call ***Binary Cellular Analysis***, provides a new framework for examining embryonic development, ***Binary Cellular Biology***, new methods for measuring the occurrence of ***Cellular Selection***, resulting from ***Differential Cellular Proliferation***, a force which molds the formation of the body from a single fertilized ovum into a multicellular animal, and new equations for comprehending ***Growth*** and ***Development*** from conception to adulthood. These ***Binary Cellular Equations*** include:

1. The ***Binary Cellular Universal Growth Equation*** captures growth from conception to adulthood, and provides a method for creating biologically-based, data-driven, ***Growth Curves***.
2. The ***Binary Cellular Universal Mitotic Fraction Equation***, which lies within the ***Binary Cellular Universal Growth Equation***, captures how growth to adult size is determined by the decline in the fraction of cells dividing, the ***Mitotic Fraction***. The ***Mitotic Fraction*** is a body-wide measure of ***Cellular Selection***, resulting from ***Differential Cellular Proliferation***.
3. The ***Binary Cellular Allometric Growth Equation*** captures the ***Creation*** and ***Growth*** of the ***Tissues***, ***Organs***, and ***Anatomical Structures*** of the body. The parameters of the ***Binary Cellular Allometric Growth Equation*** provide measures of the action of ***Cellular Selection***, the mechanism by which the ***Body*** creates its ***Parts*** and determines their ***Sizes***.
4. The ***Binary Cellular Estimated Fetal Weight Equation***, derived from the ***Binary Cellular Allometric Growth Equation***, captures the relationship between ***Ultrasound Measurements*** and the ***Size*** of the body as a whole.

These ***Binary Cellular Equations*** were not made up to ***fit*** growth data, but were derived ***from*** growth data. Thus, they don't ***imitate*** the pattern by which animals ***Grow*** and ***Develop***, but ***reveal*** the way, and the processes, by which we ***Grow*** and ***Develop***.

***Binary Cellular Analysis*** Describes How Animals Grow: The ***Binary Cellular Universal Growth Equation***

We start out as a single fertilized ovum, then 2 cells, then 4, then a thousand cells if we are a nematode worm, or a trillion cells if we are a human. As grow bigger by this process of cellular proliferation, we also grow slower. How does this occur? By considering growth data in units of numbers of cells, ***N***, the method we call ***Binary Cellular Analysis***, we have been able to discover that growth from conception to adulthood, in a great range of animals (nematodes, frogs, chickens, cows, geese, quail, turkeys, mice, rats, fish, mollusks, and humans) occurs by the same expression: the ***Binary Cellular Universal Growth Equation***.

The ***Binary Cellular Universal Growth Equation*** Describes How Growth Occurs

Mathematically, each of the parameters (***a***, ***b***, ***c***) of the ***Binary Cellular Universal Growth Equation*** (#2) has a specific impact on the shape of the body's "***S-shaped***" growth curve. The ***a*** and ***b*** parameters determine the curviness of the "***S***". The ***c*** parameter, the ***Cell Cycle Time***, determines the speed of growth, expanding or contracting the "***S***" like an accordion. As we shall see in the third paper in this series, the ***"a" Parameter*** of the ***Binary Cellular Universal Growth Equation*** provides the key to predicting the size at which each human fetus is born, which is why we call it the ***"a"*** "***Birthweight***" ***Parameter***.

The ***Binary Cellular Universal Growth Equation*** Describes How Mitotic Regulation Drives Growth.

Biologically, each of the parameters of the ***Binary Cellular Universal Growth Equation*** (#2) is linked to a specific aspect of cell division. The ***Mitotic Fraction***, ***m***, reflects how many cells are engaged in mitosis; ***m*** itself, which decreases as we become larger, is determined by the ***a*** and ***b*** parameters of ***Binary Cellular Universal Mitotic Fraction Equation*** (#1), which lies within the ***Binary Cellular Universal Growth Equation*** (#2). The ***c*** parameter, the ***Cell Cycle Time***, of the ***Binary Cellular Universal Growth Equation*** (#2) describes how long it takes cells to divide, once they have begun cell division.

***Binary Cellular Universal Mitotic Fraction Equation*** Describes Fetal Growth's Link to Endocrinology

Why, across the animal spectrum, is growth driven by the decline in the fraction of the body's cell dividing, the ***Mitotic Fraction***, by the ***Binary Cellular Universal Mitotic Fraction Equation***? One intriguing hint arose from the mathematical examination of the idealized case of the growth of cells that produce inhibitory growth factor molecules, acting in a constant volume, such as an egg or a uterus. What emerged from this exercise is that as this imaginary embryo grew, the fraction of cells not having bound one or more such molecules took on exactly the same form at the ***Binary Cellular Universal Mitotic Fraction Equation***. Thus, little more than the conventional thermodynamics of all chemical reactions is sufficient to account for growth by the ***Binary Cellular Universal Mitotic Fraction Equation***, a finding that calls out for experimental examination. This also raises the intriguing possibility little more than endocrine pharmacology may be brought to bear on instances of abnormal growth.

Key to the result of this mathematical exercise is growth within a constant volume. Virtually all animals start out life in the constant volume of an egg or a uterus. Perhaps the invention of the egg was the critical requirement needed for the emergence of the first multicellular animal from its single cell ancestor.[161]

For human fetuses, this finding of an idealized case of ligand action in a constant volume motivates a consideration of hormones, such as insulin, leading to downstream action of inhibitory growth factor molecules, such as TGF, acting in the uterus. Perhaps an examination of the genetics of such candidates[162,163,164] or their laboratory values,[165,166] may shed light on such a possibility.

As we shall see in the third paper in this series,[7] ***Binary Cellular Analysis*** of ultrasound data from of ***Small***, ***Average***, and ***Large*** newborns reveals that they were ***Slow***, ***Average*** and ***Fast*** growing fetuses differing by the ***"a" Parameter*** of the ***Binary Cellular Universal Growth Equation***, the parameter that modeling suggests could reflect the endocrinological control of fetal growth. These findings reveal possibilities for the detection and pharmacological treatment of fetuses that grow too fast or too slowly, as well as for new ways to help obstetricians guide each fetus to a ***Birthweight*** and ***Fetal Age*** that has the lowest possible chance of complication.

***Binary Cellular Analysis*** Describes Control of Growth by the ***Cell Cycle Time, c***. as Set by ***Genome Size***

What might be the mechanism behind the ***Cell Cycle Time***, the ***c*** parameter of the ***Binary Cellular Universal Growth Equation*** (#2), which not only determines how long it takes a cell to divide, but also determines how rapidly the body as a whole grows? Biochemically, the principal determinant of the ***Cell Cycle Time***, ***c***, is the amount of DNA that the cell contains. The more DNA a cell has to copy, the longer it takes to divide. This was first found by Van't Hof and Sparrow, who discovered a linear relationship between the total amount of amount of DNA and ***Cell Cycle Time*** for plant cells.[167] Their observation has subsequently been confirmed in many studies of many types of organisms.[168,169]

***Binary Cellular Analysis*** Describes How ***Junk DNA*** Controls the ***Speed of Growth***

For most animals, the largest part of the genome is non-coding DNA, sometimes called ***Junk DNA***,[170] no doubt imprecisely named.[171,172] The amount of this non-coding DNA has been found to be correlated with growth in salamanders,[173] anurans,[174] amphibians,[175] insects,[176] and copepods.[177]

The speed of growth has a profound impact on survival.[178,179] Human fetuses that grow to full-size in 10 months, or in 8 months, have a much lower chance of survival than fetuses that reach optimal size in 9 months.[180] Non-coding DNA, which determines genome size, is very susceptible to duplication or deletion, and thus provides populations of animals abundant genetic variation in genome size.[181-183]

Perhaps this genetic variation in genome size leads to genetic variation in the ***Cell Cycle Time***, ***c***, which leads to genetic variation in the speed of body growth. Such genetic variation would appear to give species a powerful resource to draw upon, so that they can evolve, by the bitter reality of Darwinian selection, to growth rates that give them the greatest chance of survival. Indeed, this makes us wonder whether ***Junk DNA*** may owe its very existence to its role in determining the speed of growth.

***Binary Cellular Analysis*** Describes the Biological Logic of the ***~Linear Phase of Human Fetal Growth***

One of the striking findings of our mathematical examination of human fetal growth is that the ***Binary Cellular Universal Growth Equation*** (#2) flattens into a roughly constant rate of growth in the last months of pregnancy, a quality we call the ***~Linear Phase of Human Fetal Growth***. In simple terms, in the last months of pregnancy, an average size fetus increases by about 26 grams each day. This observation of linearity of growth in the third trimester was observed by Mongelli and colleagues in 2015,[111] a quality they called "quasi-linear", a feature that has continued to attract use and interest.[184]

The ***~Linear Phase of Human Fetal Growth*** does not appear to be a quirky accident, as it is determined by the various parameters (***a***, ***b***, and ***c***) of the ***Binary Cellular Universal Growth Equation*** (#2), each of which is specified genetically: ***a*** and ***b*** most likely by the hormones and growth factor molecules that regulate cell division, and their genes;[100] and ***c***, the ***Cell Cycle Time***, by genome size.[185] Clearly, the human growth curve has been molded by evolution to occur at a remarkably constant, linear rate, over the last months of pregnancy. A plausible hypothesis for this linearity is that when a fetus reaches the rate of growth that occurs at the maximally possible rate of provision of nutrients, then Darwinism would drive our species to a ***Binary Cellular Universal Growth Equation*** (#2), with its genetically determined parameters, yielding the ideally maximal constant rate of growth. Departure from this ideally maximal constant rate of growth would be expected in the past to reduce the fitness of the future child and its mother, and to be eradicated by ***Natural Selection***, as well as to be expected in the present to reduce the health of the future child and its mother. As we have shall see in the in the third paper in this series, ***Binary Cellular Analysis*** reveals precisely this force behind human babies that are born ***Too Small*** and ***Too Big***.[7] As we have shall also discuss below, and examine in detail in the third paper in this series, ***Binary Cellular Analysis*** provides us with a range of options for reducing these unfortunate possibilities.

***Binary Cellular Analysis*** Describes How the Body Makes its ***Parts***

***Binary Cellular Allometric Growth Analysis*** Provides a Method for Deciphering ***Body Part Formation***

How does the body make its ***tissues***, ***organs***, and ***anatomical structures***? Capturing this information of the relationship between the ***Size*** of the ***Whole Body***, and the ***Size*** of its ***Body Parts***, is of particular importance in obstetrical practice, where the size of the fetus is estimated from ***Ultrasound Measurement*** of these ***Body Parts***. To solve this problem, we again return to ***Binary Cellular Analysis***, to examine, in units of numbers of cells, ***N***, the size of the various parts of the body, $N_p$, relative to the body as a whole, $N_w$, examined on log-log graphs.[100] We call this method ***Binary Cellular Allometric Growth Analysis***.

The ***Binary Cellular Allometric Growth Equation*** Describes ***How the Body Makes its Parts***

In this communication, we have described our studies that have shown that ***Binary Cellular Allometric Growth Analysis*** of many ***tissues***, ***organs***, and ***anatomical structures*** from many animals (nematodes, tunicates, chicks, rats, zebrafish, mice, goldfish, and humans) has found that almost all fit an equation of the same form: the **Binary Cellular Allometric Growth Equation**.

The parameters of this equation, $B_{N1}$, the ***Binary Cellular Allometric Birth***, and $S_N$, the ***Binary Cellular Allometric Slope***, capture the basic biology of ***Body Part*** formation. The ***Binary Cellular Allometric Birth***, $B_{N1}$, lies where the ***Binary Cellular Allometric Growth Equation*** crosses the ***x***-axis, thus where log($N_p$)=**0**, thus where $N_p$=**1**, that is, when the ***Body Part*** could have been a ***Single Founder Cell***. The ***Binary Cellular Allometric Slope***, $S_N$, appears on log-log graphs as a measure of whether a ***Body Part*** is growing faster ($S_N$>**1**), slower ($S_N$<**1**), or at the same speed ($S_N$=**1**), as the body as a whole. $S_N$, is Determined by ***Cell-Heritable*** change in ***Cell Cycle Time***, $c_p$, of each ***Founder Cell***.

***Binary Cellular Allometric Growth Analysis*** Describes ***Body Parts*** Arising from ***Single Founder Cells***

The ***Binary Cellular Allometric Birth***, $B_{N1}$ for each ***Body Part***, $S_N$, tells us how big the embryo was, in numbers of cells, $N_w$, when the ***Body Part*** would have been but a single cell. In our examination of the growth of many ***tissues***, ***organs***, and ***anatomical structures***, in many animals, we have not found a single case where the creation of a ***Body Part*** appears to arise from more than one ***Founder Cell***. The species (and body parts) that we have examined by ***Binary Cellular Allometric Growth Analysis*** include: nematodes, tunicates, chicks (gizzards, livers, hearts, kidneys), rats (livers, brains, kidneys, forelegs, ears, stomachs, spinal cords), zebrafish (eye lens), mice (livers, brains, kidneys, forelegs), goldfish, and humans (brains, livers, kidneys, lungs, pancreases, adrenals, thymuses, spleens, lower extremities, upper extremities, stomach, heart, intestines).

From our ***Binary Cellular Allometric Growth Analysis*** of the growth ***Body Parts*** of those animals for which we have full cell lineage data (**Supplement-Figures-I.24-27**), nematode worms and tunicates, the data have always shown that the origin of each ***Body Part*** arises from a ***Single Founder Cell*** at its ***Binary Cellular Allometric Birth***, $B_{N1}$. Tunicates are of special importance here, as they are members of the chordate phylum to which we belong

From our ***Binary Cellular Allometric Growth Analysis*** of the growth ***Body Parts*** of those animals for which we don't have full cell lineage data, we have always found that the ***Binary Cellular Allometric Growth Equation*** points down to a ***Binary Cellular Allometric Birth***, $B_{N1}$, where we would expect to find a ***Single Founder Cell*** early in development, and never when embryos are smaller than **2** cells (**Supplement-Figures-I.27-33**). Regrettably, the animals for which we have such incomplete data are principally vertebrates, including humans, in whom ***Body Part*** growth has almost always been measured late in development. Fortunately, ***Binary Cellular Allometric Growth Analysis*** provides us with a guide for identifying when we should carry out the imaging studies that will make the informative counting of cells possible.

***Body Parts*** Arising from ***Few*** or ***Single Founder Cells*** Have Been Described in Many Published Studies

While few of the large structures from late in development have been measured to the $B_{N1}$ intersection point, the zebrafish eye lens comes close, as Greiling and Clark documented its growth from just 8 cells, when the embryo is 16 hours post fertilization.[90-92] ***Binary Cellular Allometric Growth Analysis*** of these data reveal that the lens's ***Binary Cellular Allometric Growth Equation*** points down to a ***Binary Cellular Allometric Birth*** when the embryo was about 2000 cell in size, and thus $B_{N1}\approx 2000$ when $N_p\approx 1$ (**Supplement-Figure-I.33**).

Other studies that have also found that many large anatomical structures arise from small numbers of cells.[186,187] These studies include recent CRISPR/Cas9 ***cell lineage*** labeling studies.[188,189]

Many parts of the ***Drosophila*** body have also been observed from small numbers of cells.[190-192] The spiracular anlage[193], the pole cells (which form the germ cells)[194-196], and genital imaginal discs[197,198] have been seen from 2 cells onward. Imaging before these points in time was not carried out, so these structures might well have arisen from single cells. Verma and Cohen[194] have created videos that show the growth of histoblast nests, which form the mass of the body, back to as few as 4 cells at the beginning of their videos; one wonders what would have been seen if one turned on the camera before then. Such a simple data collection exercise cries out for experimental examination. Furthermore, these videos make clear that histoblast nest cells increase in number by cell division rather than by recruiting unrelated cells.[194] For a deeper review of cellular origins of Drosophila ***Body Parts***, see the supplement to reference 199.

#### ***Binary Cellular Analysis*** Describes How the Body Controls the Growth of its ***Parts***

##### ***Body Part*** Growth, $S_N$, Arises from ***Cell-Heritable*** Change of the ***Cell Cycle Time***, $c_p$, in ***Founder Cells***

By examining ***Growth*** of our ***tissues***, ***organs***, and ***anatomical structures*** in units of numbers of cells, ***N***, we have been able to see that the ***Speed of Growth*** of each ***Body Part***, captured by its ***Binary Cellular Allometric Slope***, $S_N$, is determined by a ***Cell-Heritable*** change in the ***Cell Cycle Time***, $c_p$, of its ***Founder Cell***. That the change in the ***Cell Cycle Time***, $c_p$, of each ***Founder Cell*** is ***Cell-Heritable*** is made visible by the log-linearity of ***Body Part*** growth captured by ***Binary Cellular Allometric Growth Equation*** on log-log graphs, revealing the new value of $c_p$ arising in each ***Founder Cell*** continues in the ***Founder Cell's*** progeny.

##### ***Body Part*** Change in the ***Cell Cycle Time***, $c_p$, and DNA Methylation

There are a number of ***Cell-Heritable*** biological processes that influence the time it takes for a cell to divide, of which DNA methylation has been the most widely studied.[200,201] Methylated DNA takes longer to copy than unmethylated DNA, and thus methylation causes cells to take more time to divide.

##### ***Body Part*** Change in the ***Cell Cycle Time***, $c_p$, is But One Example of ***Cell-Heritable*** Change.

Of course, body parts display many ***Cell-Heritable*** changes in their phenotypes. Perhaps the most familiar examples are the ***Cell-Heritable*** changes in the expression of genes that mark the cell types that characterize the specific features of each of the body's ***tissues***, ***organs***, and ***anatomical structures***.

**_Binary Cellular Analysis_** Describes How **_Cellular Selection_** Drives the **_Formation_** and **_Growth_** of the Body

What is **_Cellular Selection_**?

The Merriam-Webster Dictionary notes that "Selection" is "the act or process of selecting", while "selecting" is "to choose (as by fitness or excellence) from a number or group". Etymologically, the word is derived from the Latin "se" (apart) and "legere" (choose), and is, perhaps most familiar in "***Natural Selection***", as coined by Darwin. ***Natural Selection*** is the occurrence of differential reproduction, or death, among organisms, which ***Ecological Analysis*** has revealed determines the ***Size*** and ***Evolution*** of Natural Populations. ***Cellular Selection*** is the occurrence of differential cellular proliferation,[112,114] or apoptosis[115] , among our cells, which ***Binary Cellular Analysis*** has revealed determines the ***Size*** and ***Development*** of the animal body.

In the world of our single cell ancestors, ***Natural Selection*** and ***Cellular Selection*** were the same thing. Perhaps our single cell forebears were able to bring their progeny together to become multicellular animals by domesticating ***Natural Selection*** into ***Cellular Selection***, bringing the Darwinian world inside to create the first animal, more than a half a billion years ago.[4,5]

1. **_Binary Cellular Analysis_** Describes **_Cellular Selection's_** Roles in Regulating the Growth of the Body
   a. **_Progeny Cell Selection_** in the body as a whole:

***Binary Cellular Analysis*** of data on the ***Size*** of the body of developing animals has revealed that embryos create growth to adult size by ***Cellular Selection***, as captured by the ***Binary Cellular Universal Mitotic Fraction Equation***. This has been seen with growth data from a great range of animals, including nematodes, frogs, chickens, cows, geese, quail, turkeys, mice, rats, fish, mollusks, and humans. For each of these creatures, ***Binary Cellular Analysis*** revealed that embryos start out with all cells dividing, soon smoothly transitioning away from ***Exponential Growth***, by having progressively fewer and fewer cells dividing. The ***Mitotic Fraction***, the fraction of cells dividing, whose value can be measured from ***Growth*** data with the ***Mitotic Fraction Method***, provides the measure of such ***Cellular Selection's*** action in the control of growth.

2. **_Binary Cellular Analysis_** Describes **_Cellular Selection's_** Role in **_Formation_** and **_Growth_** of Body Parts

***Binary Cellular Analysis*** has revealed that embryos create ***Body Parts*** by two varieties of ***Cellular Selection***, captured by the two parameters of the ***Binary Cellular Allometric Growth Equation***:

   a. **_Founder Cell Selection_** in the body as a whole:

***Binary Cellular Analysis*** of ***tissues***, ***organs***, and ***anatomical structures*** of the embryo reveals that ***Body Parts*** are created by the ***Selection*** of individual ***Founder Cells*** among the many other cells in the early embryo. The $B_{N1}$ parameter of the ***Binary Cellular Allometric Growth Equation***, the ***Binary Cellular Allometric Birth***, tells us the size of the embryo when each ***Founder Cell*** is ***Selected*** to give rise to its ***Body Part***.

   b. **_Progeny Cell Selection_** in Each **_Body Part_**:

***Binary Cellular Analysis*** of ***tissues***, ***organs***, and ***anatomical structures*** of the embryo also tells us that ***Body Part Size*** is determined by ***Differential Cellular Proliferation***, the most common form of ***Cellular Selection***. The $S_N$ parameter of the ***Binary Cellular Allometric Growth Equation***, the ***Binary Cellular Allometric Slope***, tells us the magnitude of this manifestation of ***Cellular Selection***. This ***Differential Cellular Proliferation*** is ascribable to ***Founder Cells*** undergoing a cell-heritable changes in their ***Cell Cycle Times***, $c_p$, by a mechanism such as DNA methylation, which continue in their progeny, determining whether a ***Body Part*** will grow faster or slower than the body as a whole.

**_Cellular Selection_**: in the Immune System and Liver

The creation of ***Body Parts*** by selecting ***Single Founder Cells***, and ***Cellular Selection*** of each of these clonal body parts to become dominant by growing more rapidly, are familiar features of the immune system, where the production of clones of cells producing antibodies and T-cell receptors arise from single lymphocytes (i.e. ***Founder Cell Selection***), and are expanded, by ***Clonal Selection*** (i.e. ***Progeny Cell Selection***).[202] Imaging of the cells in the liver has also revealed a similar process of clones arising from single ***Founder Cells***, with ***Cellular Selection*** driving the synthesis of many of the plasma proteins in our blood.[105,114]

***Binary Cellular Analysis*** Has Revealed Practical Methods for ***Managing Human Fetal Growth***.

As we have seen here, and in the next two papers in this series, these new ***Binary Cellular Equations*** have at least six practical applications in health:

1. As we shall see in the next paper in this series,[6] the ***Binary Cellular Estimated Fetal Weight Equations*** described above provides better, more accurate and more precise equations and methods for measuring of the ***Size*** of human fetuses from fetal ultrasound measurements.
2. As we shall see in the third paper in this series,[7] the ***Binary Cellular Equations*** described above provides better, more accurate, and more precise, equations and methods for detecting the abnormal ***Growth*** of human fetuses that leads to birthweights that are ***Too Small*** or ***Too Large***.
3. As we shall see also in the third paper in this series,[7] ***Binary Cellular Mathematics***, and specifically the ***Binary Cellular Universal Growth Equation***, provides the basis for deriving biologically-based, data-driven, ***Fetal Growth Curves***.
4. As we shall also see in the third paper in this series,[7] the ***Binary Cellular Equations*** described above reveals previously hidden forces by which human fetuses grow to birthweights that are ***Too Small*** or ***Too Large***.
5. As we shall also see in the third paper in this series,[7] the ***Binary Cellular Equations*** described above reveals previously hidden opportunities for managing human fetuses that grow ***Too Slow*** or ***Too Fast***, so as to lower the chance of each fetus being born ***Too Small*** or ***Too Large***.
6. ***Binary Cellular Analysis*** also has the potential to examine childhood growth.

These practical applications will be reviewed in greater detail in the DISCUSSION sections of the second and third paper in this series.[6,7]

Web-based Calculators Provide ***Binary Cellular*** Calculations That Aid Obstetric Care.

A first-generation Web-based calculator for calculating ***Fetal Weight*** from the ***Abdominal Circumference Binary Cellular Estimated Fetal Weight Equation***, is now available at “https://kidzgrowth.com”. Encoding the additional ***Binary Cellular*** calculations described here is straightforward,[203] and underway, including calculation of the speed of fetal growth from multiple, sequential, ***Ultrasound Measurements***, in units of ***Weight-Gain Rate*** (grams/day), and in units of the value of the ***“a” Parameter*** of the ***Binary Cellular Universal Growth Equation***, as well as projected values ***Fetal Weight*** in the days and weeks ahead. displayed on a ***Binary Cellular Fetal Growth Chart***, with the statistical limits of reliability also shown. Also to be provided by these calculators are tools for performance feedback, comparing each ***Ultrasound Measurement*** to the ultimate ***Birthweight***, so that sonographers and their supervisors can review performance over the long term, as well as the performance of each individual case. These web-based calculators make the results of these ***Binary Cellular Equations*** widely available for aiding in obstetric care of individual pregnancies.

### *The Binary Cellular Biology of Human Growth*:
#### II. Calculating the Size of the Human Fetus from the Size of its Parts
**204**


**ABSTRACT**

Assessing and understanding the ***Size*** of the human fetus provides essential information for the management of the health of the newborn and its mother, a quality that is commonly measured by ***Fetal Ultrasound***. Many equations for making such calculations have been considered, none of which have been entirely satisfactory. As we have shown in the previous paper in this series, a consideration of the formation of the body, in units of numbers of cells, ***N***, an approach we call ***Binary Cellular Analysis***, provides new quantitative tools for estimating the ***Size*** and ***Age*** of the fetus. These tools include a new mathematical basis for creating useful expressions, ***Binary Cellular Estimated Fetal Weight Equations***, for calculating human ***Fetal Weight*** from ***Ultrasound Measurements***. As we show here, the most promising of these expressions, the ***Abdominal Circumference Binary Cellular Estimated Fetal Weight Equation***, performs better than other currently used ***Fetal Weight Equations***, while also yielding up new methods for improving ultrasound size assessment. Web-based calculators (https://kidzgrowth.com) make these mathematical manipulations available for obstetric care.

#### INTRODUCTION

While many equations have been considered for estimating ***Fetal Weight*** from ***Ultrasound Measurements***, none have been entirely satisfactory.[205-218] To address this challenge, in this, the second of three articles in this series,[219,220] we again return to a consideration the ***Size*** the whole body, ***w***, and its parts, ***p***, in units of numbers of cells, $N_w$ and $N_p$, especially those parts measured by ***Fetal Ultrasound***. We call this method ***Binary Cellular Analysis***, as it examines the ***Binary*** choices made by our cells to be ***Mitotic*** or ***Quiescent***, determining the number of cells in our ***Bodies*** and our ***Parts***.[221] Thus, ***Binary Cellular Analysis*** provides a way to examine the body's process of ***Cellular Selection***, resulting from ***Differential Cellular Proliferation***, which molds the body from a single fertilized ovum into a multicellular animal. In this way, ***Binary Cellular Analysis*** gives us a method to decipher how we ***Grow*** and ***Develop***, from which we could construct biologically-based equations that relate ***Ultrasound Measurement*** of ***Body Parts*** to ***Fetal Weight***.[222,223]

As we have seen in the first article in this series, such a ***Binary Cellular*** approach provides new expressions for calculating the ***Size*** and ***Growth*** of human fetuses from ***Fetal Ultrasound Measurements***: ***Binary Cellular Estimated Fetal Weight Equations***. Here we shall see that the most promising of these expressions, the ***Abdominal Circumference Binary Cellular Estimated Fetal Weight Equation***, performs better than other widely used ***Estimated Fetal Weight Equations***, while also yielding up new methods for improving ultrasound size assessment.

#### METHODS

The ***Combined Dataset***

For this, and the following manuscript in this series,[220] we utilized a deidentified dataset of 821 pregnant women seen at the Brigham and Women's Hospital. We call this dataset the "***Combined Dataset***". 23 of these patients had newborns that were less than 1,800 grams, 27 of which were less than 2,000 grams, and 123 of which were 2,500 or fewer grams at birth. Of the 23 newborns for which were less than 1,800 grams newborns, we had full data, including ***Crown to Rump Length*** (***CRL***) measurements early in development, for 14. In addition to ***Birthweight***, these datasets contained information on ultrasound measurement of: ***Biparietal Diameter*** (BPD), ***Occipitofrontal Diameter*** (OFD), ***Abdominal Diameter*** (AD), ***Femur Length*** (FL), ***Head Circumference*** (HC), ***Abdominal Circumference*** (AC), and ***Crown Rump Length*** (CRL). The ***Combined Dataset*** was comprised of two subsets: a ***194" dataset*** of 194 women, and a ***627" dataset***, which contains data on 627 pregnant women. The two datasets were analyzed separately, as only the "***627" dataset*** contained full data on each newborn and its mother. Data were analyzed in Microsoft Excel.

The ***Combined Dataset*** was divided randomly into to two subsets: a ***Training Dataset***, used to derive the parameters for the new ***Binary Cellular Estimated Fetal Weight Equations***, and a ***Testing Dataset***, used to test the accuracy of these expressions and parameters. Both datasets contained 117 ultrasound examinations carried out within 7 days of birth.

The "***194" Dataset***

The 194 women of the "***194" dataset*** was comprised of 1022 ultrasound examinations. 302 of these examinations measured ***Crown to Rump Length*** (***CRL***) early in development. 720 of these examinations had other measurements made later in development.

The "***627" Dataset***: Overview

The 627 women "***627" dataset*** contains data on 215 ultrasound measurements made within 7 days of birth, which were used to test the accuracy of the ***Estimated Fetal Weight*** (***EFW***) ***Equations***, and especially the ***Binary Cellular Estimated Fetal Weight Equations*** and their parameters, whose values were extracted from the data for women in the "***194" dataset***.

Among the 627 patients in the "***627" dataset***, the ***Fetal Age*** at birth (the estimated time from conception to birth, calculated from the ***CRL Fetal Age Equation*** (#7a)) had a mean value of 277 days; a ***Standard Deviation*** of 15 days; a median of 278 days; the longest being 306 days; and the shortest being 175 days (**Supplement-Figure-III.1**). The birthweights had a mean value of 3178 grams; a ***Standard Deviation*** of 743 grams; a median of 3100 grams; the largest being 5180 grams; and the smallest being 700 grams (**Supplement-Figure-III.1**).

Newborns suitable for ***Binary Cellular Fetal Growth Analysis*** had at least one CRL measurement early in pregnancy, for ***Fetal Age*** calculation, with the ***CRL Fetal Age Equation*** (#7a), and at least two ultrasound measurements later in pregnancy, for ***Fetal Weight Estimation*** with the ***Abdominal Circumference Binary Cellular Estimated Fetal Weight Equation*** (#14)). The "***627" dataset*** has been utilized in references 224, 225, 226, and 227.

The "***627" Dataset***: ***Small***, ***Average***, and ***Big Baby Study Groups***

Detailed analysis was carried out of 19 ***Average Size Babies*** (birthweight 3355-3459 grams), 53 ***Big Babies*** (birthweight 4000-4725 grams), and 26 ***Small Babies*** (birthweight 1125-2065 grams). Of these, full information for all ultrasounds was available for 15 ***Average Size Babies*** (birthweight 3355-3459 grams), 30 ***Big Babies*** (birthweight 4000-4725 grams), and 23 ***Small Babies*** (birthweight 1125-2065 grams) (TABLE I).

Equations not derived or specified in the main text of this communication

As developed in the previous paper in this series,[262] the ***Binary Cellular Universal Growth Equation*** (#2), for capturing the relationship between ***size***, ***N***, and ***age***, ***t***:

$$N_w = \int ln2/c * N_w * a \hat{} \bar{N}_w \hat{} b \qquad (2c)$$

Within this expression lies the ***Binary Cellular Universal Mitotic Fraction Equation***:

$$m = a^{(N_w^b)} \qquad (4)$$

**RESULTS**

The Data: ***Ultrasound Measurements*** vs ***Birthweight***.

Let us begin by taking a general look at the relationships between various ***Ultrasound Measurements*** and ***Birthweights*** in our datasets (METHODS section). This can be seen simply by graphing each ***Ultrasound Measurement*** against the corresponding ***Birthweight*** of that newborn and fitting the data to a simple linear regression. Such graphs for ***Ultrasound Measurements*** carried out within 7 days of birth can be seen in **Supplement-Figure-II.1**. These graphs reveal that for each kilogram of ***Fetal Weight***, ***Abdominal Circumference*** (AC) increases by ~447 millimeters, while ***Abdominal Diameter*** (AD) increases by ~142 millimeters, ***Head Circumference*** (HC) by ~19.6 millimeters, ***Occipitofrontal Diameter*** (OFD) by ~7.8 millimeters, ***Biparietal Diameter*** (BPD) by ~6.2 millimeters, and ***Femur Length*** (FL) by ~5.9 millimeters (**Supplement-Figures II.1&2**). The ***Abdominal Circumference*** (AC) measurements also had a much lower ***Standard Deviation*** than the other ***Ultrasound Measurements***. (**Supplement-Figure-II.2**).

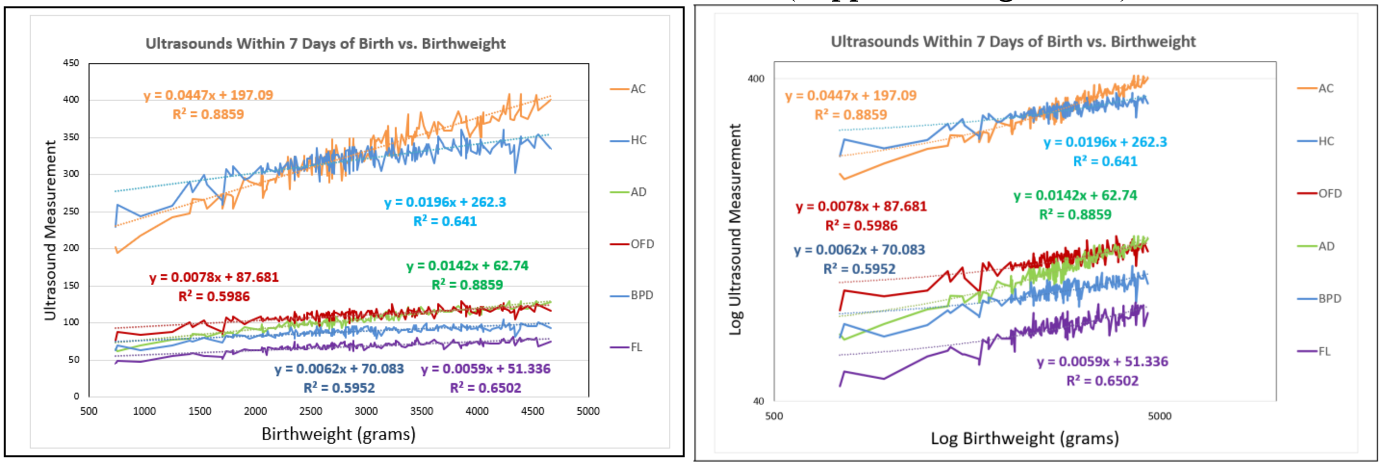


**Supplement-Figure-II.1** Comparison of ***Birthweights***, and ***Ultrasound Measurements*** made within 7 days of birth, and fit of the data to linear regressions. Linear scales (left), and logarithmic scales (right).

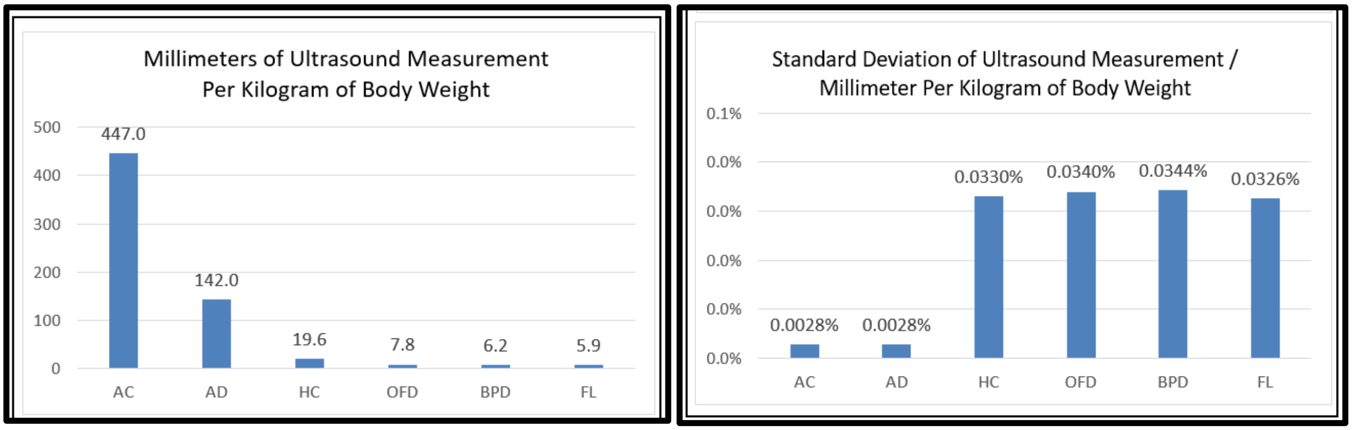


**Supplement-Figure-II.2** Comparison of ***Birthweights***, and ***Ultrasound Measurements*** made within 7 days of birth, in Millimeters/Kilogram (left), and Standard Deviation (right).

This first look at the relationship between ***Ultrasound Measurement*** and ***Birthweight*** reveals that ***Abdominal Circumference*** (AC) gives much greater leverage in capturing ***Fetal Weight*** than the other ***Ultrasound Measurements***. For example, ***Abdominal Circumference*** (AC) had 75-fold greater potential to capture change in ***Fetal Weight*** than ***Femur Length*** (FL), with 10-fold less error. Thus, ***Abdominal Circumference*** (AC) gives a much stronger, and much less noisy, signal for estimating ***Fetal Weight*** than the other ***Ultrasound Measurements***. These features make clear why ***Abdominal Circumference*** (AC) has repeatedly been proven to be the most effective ***Ultrasound Measurement*** for many ***Estimated Fetal Weight Equations***.[31-27]

We can visualize the noise of each ***Ultrasound Measurement*** simply by constructing scatter plots of ***Birthweight*** vs. ***Ultrasound Measurements***, when examined within a few days of birth. Note in **Supplement-Figure-II.3** how babies with the same ***Birthweight*** display a great variety of ***Ultrasound Measurements*** for all of the anatomical sites examined. Note also how the $r^2$ value for ***Abdominal Measurement*** is much smaller than for other anatomical ***Measurements***.

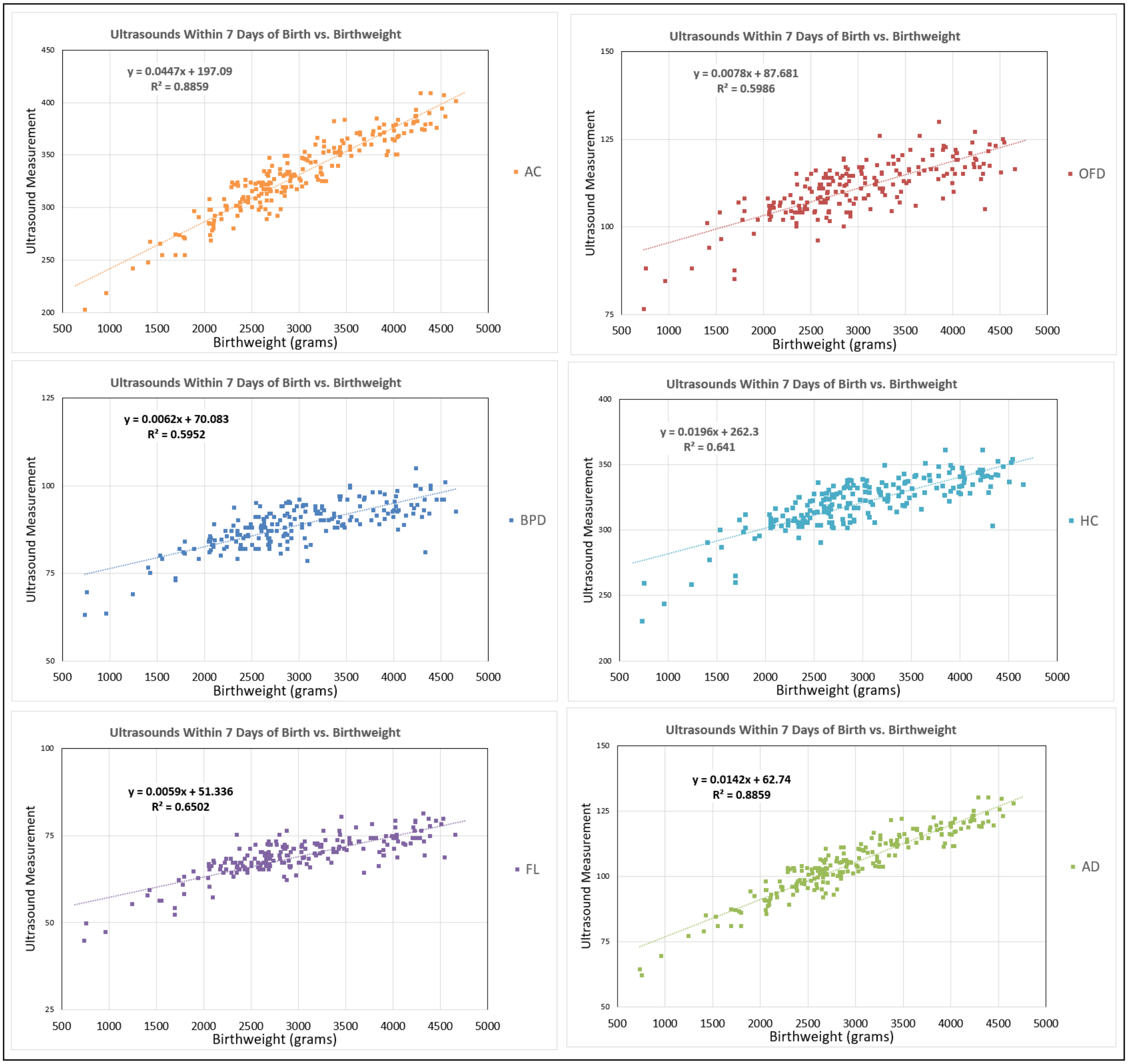


**<u>Supplement-Figure-II.3</u>** ***Ultrasound Measurement*** vs ***Birthweight***

We can also visualize the noise of ***Ultrasound Measurement*** by constructing scatter plots of ***Ultrasound Measurement*** vs. ***Ultrasound Measurement***, from individual sonographic sessions, of one anatomical site against another (**Supplement-Figure-II.4)**.

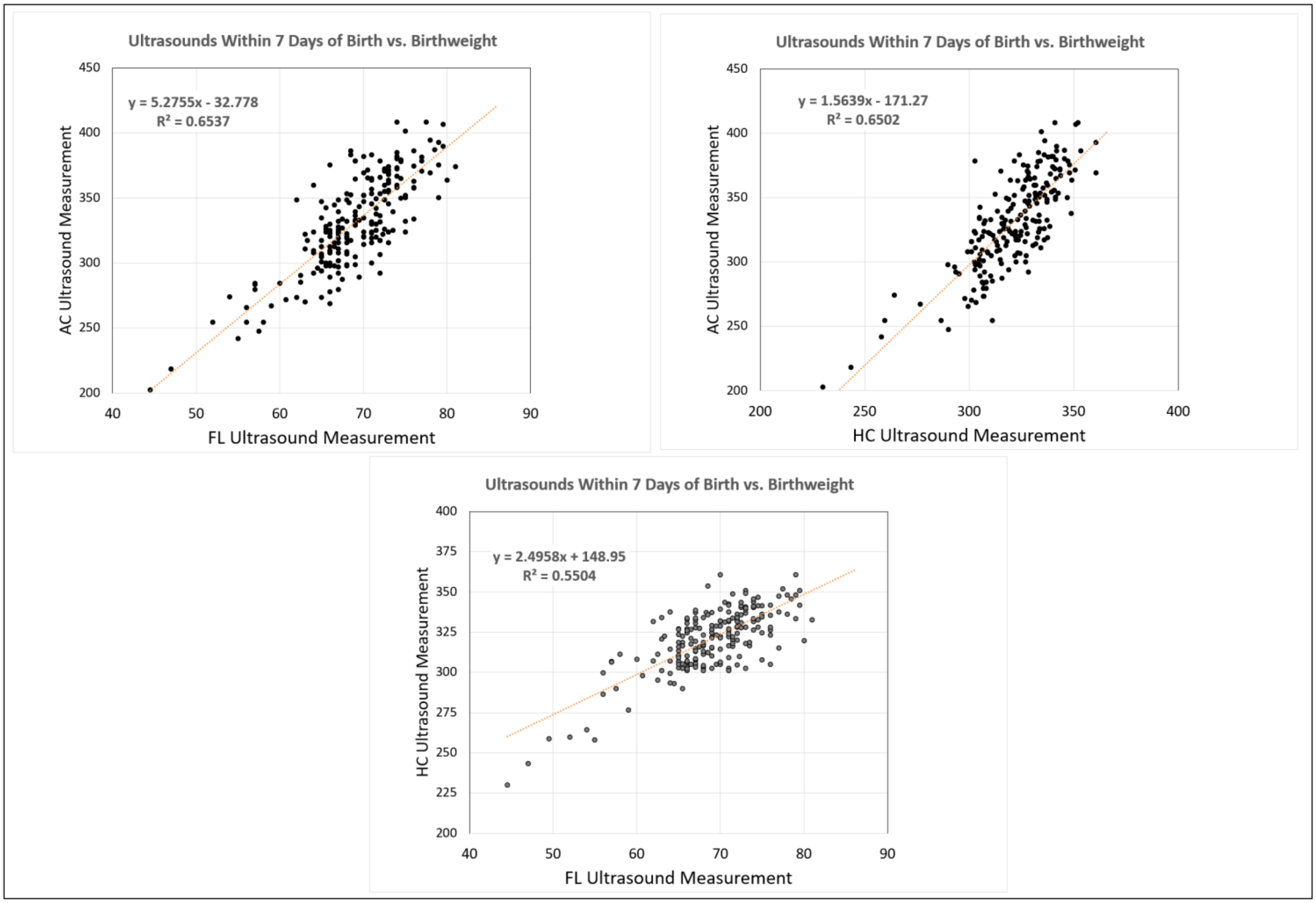


**Supplement-Figure-II.4** ***Ultrasound Measurement*** vs ***Ultrasound Measurement***

As we shall see below, and as others have long appreciated,[212] the noise in ***Ultrasound Measurement-Birthweight*** relationship is principally due to the ultrasound procedures not being carried out as satisfactorily as they could have been. Below we shall see that ***Binary Cellular Analysis*** of ***Fetal Ultrasound-Fetal Weight*** relationships can give us tools for reducing this noise.

Current ***Estimated Fetal Weight Equations*** for Determining ***Fetal Weight*** from ***Ultrasound Measurements***

Many ***Estimated Fetal Weight Equations*** have been constructed for estimating the size of the fetus from ultrasound measurements. Among the most common are:

***Hadlock Estimated Fetal Weight Equation-1***:

**EFW=10^((1.3596+0.0064*(HC/10)+0.0424*( AC /10)+0.174*(FL/10)+0.00061*(BDP/10)*(AC/10)-0.00386*(AC/10)*(FL/10)))** (900)

***Hadlock Estimated Fetal Weight Equation-2***:

**EFW=10^(1.304+0.05281(AC)+0.1938(FL)-0.004(AC)(FL)))** (901)

***Hadlock Estimated Fetal Weight Equation-3***:

**EFW=10^(1.335-0.0034(AC)(FL)+0.0316(BPD)+0.0457(AC)+0.1623(FL))** (902)

***Hadlock Estimated Fetal Weight Equation-4***:

**EFW=10^(0.3596+0.00061(BPD)(AC)+0.0424(AC)+0.174(FL)+0.0064(HC)-0.00386(AC)(FL))** (903)

***Hadlock Estimated Fetal Weight Equation-5***:

**EFW=10^(1.3596+0.006(HC)+0.0424(AC)(FL))** (904)

***INTERGROWTH Estimated Fetal Weight Equation***:

$$EFW=10^{(5.084820-54.06633\times(AC/100)^3-95.80076\times(AC/100)^3\times\log(AC/100)+3.136370\times(HC/100))} \quad (905)$$

***Hadlock Estimated Fetal Weight Equation-4*** is the most commonly used ***Estimated Fetal Weight Equation***, and many studies have found it to be superior to the others.[205-218]

New ***Binary Cellular Estimated Fetal Weight Equations*** for Determining ***Fetal Weight*** from ***Ultrasound***

As noted in the previous paper in this series, for more than a century, biologists have observed that the ***Size***, $\boldsymbol{p}$, of many parts of the body (***tissues***, ***organs***, and ***anatomical structures***), when compared with the ***Size***, $\boldsymbol{w}$, of the body as a whole, frequently appear on log-log graphs as straight rows of ***p-w***-pairs, that is:

$$\log(\boldsymbol{w}) = \boldsymbol{S} \cdot \log(\boldsymbol{p}) + \log(\boldsymbol{B}) \quad (10a)$$

which is equivalent to:

$$\boldsymbol{w} = \boldsymbol{B} \cdot \boldsymbol{p}^{\boldsymbol{s}} \quad (10b)$$

This observation is called ***Allometric Growth***, and Equation #10 is called the ***Allometric Growth Equation***. Reformatting such ***Allometric Growth*** into units of numbers of cells, ***N***:

$$\log(\boldsymbol{N_w}) = \boldsymbol{S_N} \cdot \log(\boldsymbol{N_p}) + \log(\boldsymbol{B_{N1}}) \quad \text{where} \quad \boldsymbol{S_N} = \frac{\boldsymbol{c_p}}{\boldsymbol{c_w}} \quad (11a)$$

which is equivalent to:

$$\boldsymbol{N_w} = \boldsymbol{B} \cdot \boldsymbol{N_p}^{\boldsymbol{s}} \quad (11b)$$

We call this expression the ***Binary Cellular Allometric Growth Equation*** (#11) (**Supplement-Figure-I.23**). With this expression, we could put ***Binary Cellular Biology*** to work against the ***Training Dataset*** (see METHODS section) to derive a new, biologically based, method for calculating ***Fetal Weight*** from ***Ultrasound Measurements***, ***Binary Cellular Estimated Fetal Weight Equations***.

Let us begin with the ***Binary Cellular Allometric Growth*** and ***Allometric Growth Equations*** (#10 and #11) to examine the ***Allometric*** relationship between individual ***Fetal Ultrasound Measurements*** (***U***) and ***Birthweights*** (***$B_w$***):

$$\log(\boldsymbol{Estimated\ Fetal\ Weight}) = \boldsymbol{\sigma} \cdot \log(\boldsymbol{Ultrasound\ Measurement}) + \log(\boldsymbol{\beta}) \quad (12a)$$

which is equivalent to:

$$\boldsymbol{Estimated\ Fetal\ Weight} = \boldsymbol{\beta} * (\boldsymbol{Ultrasound\ Measurement}^{\boldsymbol{\sigma}}) \quad (12b)$$

We call this expression the ***1st-Steps-Binary Cellular Estimated Fetal Weight Equation*** (#12). Values for the parameters were derived from ultrasound measurements that were carried out within 7 days of birth, from the ***Training Dataset*** (METHODS).

To get to the value of the **β** parameter for each ***Ultrasound Measurement***, several steps are required. With each step, we arrive at more finely tuned values (**$\beta_1$, $\beta_2$, $\beta_3$**), until the final value, **β,** is reached:

$$\boldsymbol{Estimated\ Fetal\ Weight_{step}} = \boldsymbol{\beta_{step}} * (\boldsymbol{Ultrasound\ Measurement}^{\boldsymbol{\sigma}}) \quad (13)$$

or

$$\boldsymbol{EFW_{step}} = \boldsymbol{\beta_{step}} * (\boldsymbol{U}^{\boldsymbol{\sigma}}) \quad (13b)$$

Where the ***Estimated Fetal Weight $_{step}$*** is the ***Estimated Fetal Weight*** for each step in our refinement of the parameters of this expression, as we shall describe next.

As described in the previous paper in this series, and confirmed in the next paper, over the last months of pregnancy, the average increase in fetal weight is ~26 grams/day, a feature of the ***Binary Cellular Universal Growth Equation*** (#2) we call the ***~Linear Phase of Human Fetal Growth.***[228] Therefore, we examined values for both ***Birthweight*** and ***Birthweight-(days-until-birth*26 grams)***, which captures the average change in ***Fetal Weight*** that would be expected to occur between ultrasound and birth against the ***Training Dataset***. The 26 grams/day adjusted values gave slightly better performance. The INTERGROWTH-21 study also generated their ***Estimated Fetal Weight Equation*** with such a ***days-until-birth*** correction.[211]

Of course, ***Estimated Fetal Weight Equations*** based on ***Birthweight-(days-until-birth*26 grams)***, provide estimates of ***Fetal Weight*** from data on ***Ultrasound Measurement*** and ***Birthweight***. ***Estimated Fetal Weight Equations*** based on ***Birthweight*** alone are subject to distortion by the amount of time between the ultrasound and birth.[229] In the next paper in this series, we shall examine how it is possible to combine ***Binary Cellular*** measurements of ***Fetal Size***, as we are examining here, with ***Binary Cellular*** measurements of ***Fetal Growth***, as we examined in the previous paper, so as to project values backwards and forward in time, to comprehend and calculate how much larger fetuses will become in the days ahead, and how much smaller they were in the days past. Such information goes right to the heart of understanding, from the past, and managing, for the future, the growth and health of fetuses and the newborns they will become.

The values of the $\sigma$ and $\beta_1$ parameter, for each ***Ultrasound Measurement*** were extracted from linear regressions of the LOG of each ***Ultrasound Measurement*** vs. the LOG of its corresponding ***Birthweight***, or ***Birthweight-(days-until-birth*26 grams)*** in the ***Training Dataset***. The linear regressions of each of these LOG-LOG comparisons also provided, in their correlation coefficients ($r^2$), a measure of the potential of each ***Ultrasound Measurement*** to capture ***Fetal Weight*** (**Supplement-Figure-II.5**). Note, for example, how the $r^2$ value for ***Abdominal Circumference*** (AC) reflects a superior performance in comparison to other ***Ultrasound Measurements*** (**Supplement-Figure-II.5**).

The error (EFW-BW/BW) for each $EFW_{step}$ of ***Ultrasound Measurement*** was calculated for both ***Birthweight*** and ***Birthweight-(days-until-birth*26 grams)***, and the average value for $EFW_1$ for BPD, OFD, AD, FL, HC, and AC, were calculated in excel. A slight improvement of each $\beta_1$ parameter value was made by manually refining this average value to yield zero error in the ***Training Dataset***, yielding $\beta_2$ values for BPD, OFD, AD, FL, HC, and AC (**Supplement-Figure-II.6**).

A problem that has arisen for many ***Estimated Fetal Weight Equations*** has been the inability to accurately capture the sizes of fetuses that will become ***Large*** and ***Small Babies***.[236,240,245] Note in **Supplement-Figure-II.7** how the linear regressions of $EFW_2$ versus ***Birthweight-(days-until-birth*26 grams)***, yields a line for ***Abdominal Circumference*** (AC) that has a slope that is close to 1, and the intercept close to 0,0, indicating roughly similar estimations of fetal weight for ***Large*** and ***Small Babies***, while this is not the case for the other ***Ultrasound Measurements*** (BPD, OFD, AD, FL, HC). This imbalance is easily solved by adding one more parameter, $\gamma$:

$$\textit{Fetal Weight} = \beta * (\textit{Ultrasound Measurement}^{\sigma}) - \gamma \tag{14a}$$

or

$$EFW = \beta * (U^{\sigma}) - \gamma \tag{14b}$$

We call Equation #14 the ***Binary Cellular Estimated Fetal Weight Equation***. The value of $\gamma$ could be determined by graphing $EFW_3$ versus ***Birthweight-(days-until-birth*26 grams)***, against the ***Training Dataset***, carrying out linear regression (**Supplement-Figure-II.8**), and using the slope and intercept to calculate the values of the $\gamma$ and $\beta_3$ parameter. A final slight adjustment of manually refining the $\beta_3$ parameter to yield zero error in the ***Training Dataset***, yields the final $\beta$ parameter value. The resulting ***EFW*** vs ***Birthweight***, and ***Birthweight-(days-until-birth*26 grams)*** graphs reveal the capacity of the ***Binary Cellular Estimated Fetal Weight Equations*** to generate values without any distortion for ***Large*** and ***Small Babies*** (**Supplement-Figure-II.8**). For ***Abdominal Circumference***: $\beta=0.001371156$; $\sigma=2.5187$; $\gamma=244.25$ and thus:

$$EFW = (0.001371156 * (AC\text{^}2.5187)) - 244.25 \tag{14c}$$

We call Equation #14c the ***Abdominal Circumference Binary Cellular Estimated Fetal Weight Equation***.

C:_1\BBCDG V7 DATA______Baby Math Data 10 20 23_____New EFW Calcs Nov 4 23__medium to medium 2 23 2024\[Deid POPs and LIFECODES Combined. Slope & Beta Adjustable. V20d 4 12 2024 Train & Test. 26 gr-day in Train & Test.xlsx]Test.7d 4th (2.5) (2)'!$E$124

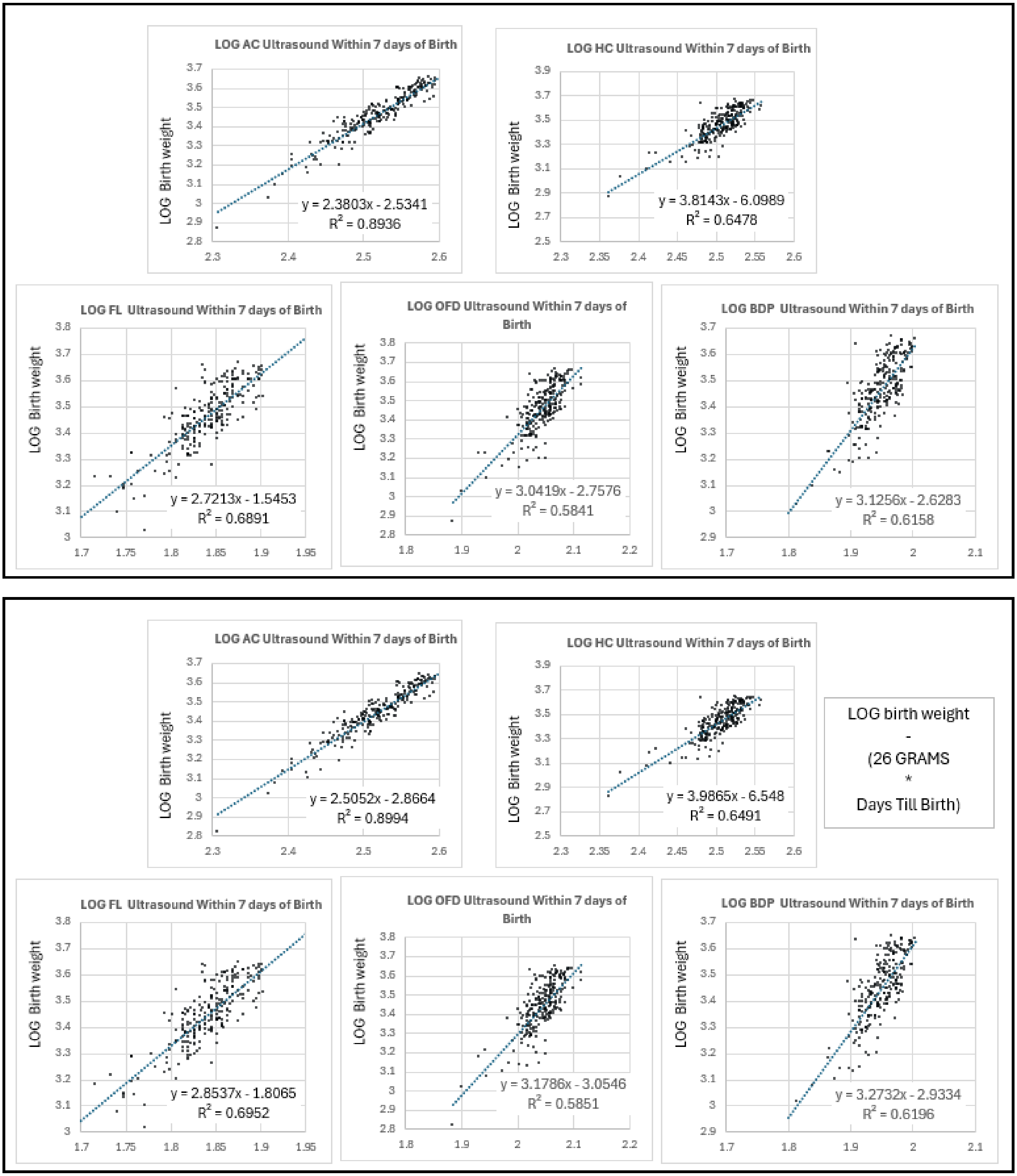


**Supplement-Figure-II.5**

Fit of the LOG of each ***Ultrasound Measurement*** carried out within 7 days of birth vs. the LOG of its corresponding ***Birthweight*** (top), or ***Birthweight-(days-until-birth*26 grams)*** (bottom), in the ***Training Dataset***.

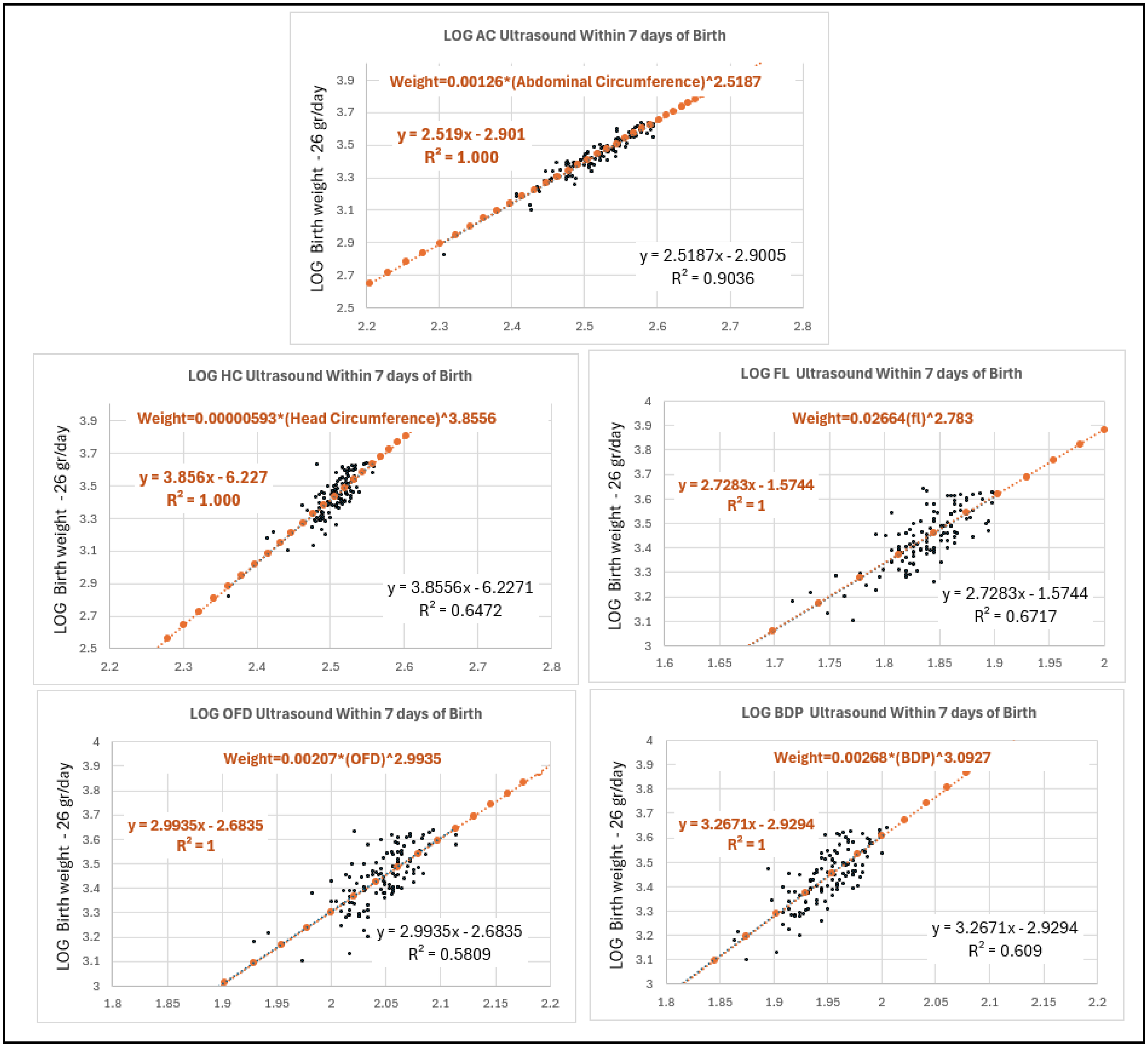


<u>**Supplement-Figure-II.6**</u>

Fit of ***Fetal Ultrasound*** measurements carried out within 7 days of birth vs ***Birthweight***, to the ***Binary Cellular Estimated Fetal Weight$_{step}$*** Equation #13 in the ***Training Dataset***.

C:_1\BBCDG V7 DATA_______Baby Math Data 10 20 23_____New EFW Calcs Nov 4 23__medium to medium 2 23 2024\[Deid POPs and LIFECODES Combined. Slope & Beta Adjustable. V20b 4 9 2024 Train & Test. 26 gr-day in Train & Test.xlsx]Train.7d'!$AM$218

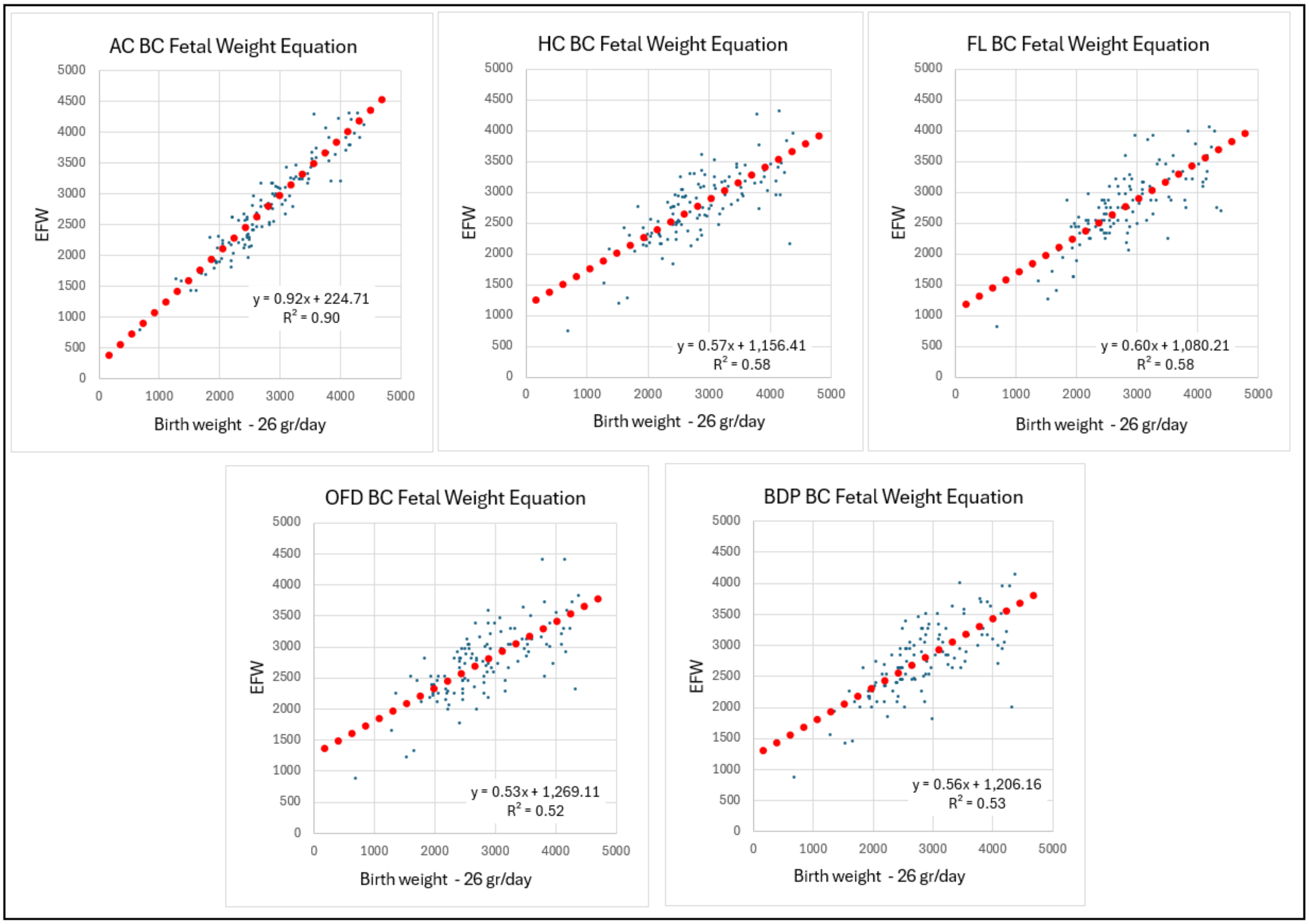


C:_1\BBCDG V7 DATA_______Baby Math Data 10 20 23_____New EFW Calcs Nov 4 23__medium to medium 2 23 2024\[Deid POPs and LIFECODES Combined. Slope & Beta Adjustable. V20b 4 9 2024 Train & Test. 26 gr-day in Train & Test.xlsx]Train.7d'!$AM$218

**Supplement-Figure-II.7**

Fit of ***Fetal Ultrasound*** measurements carried out within 7 days of birth vs ***Birthweight***-(***days-until-birth*26 grams***), to the ***Binary Cellular Estimated Fetal Weight***$_{step}$ Equation #13, in the ***Training Dataset***.

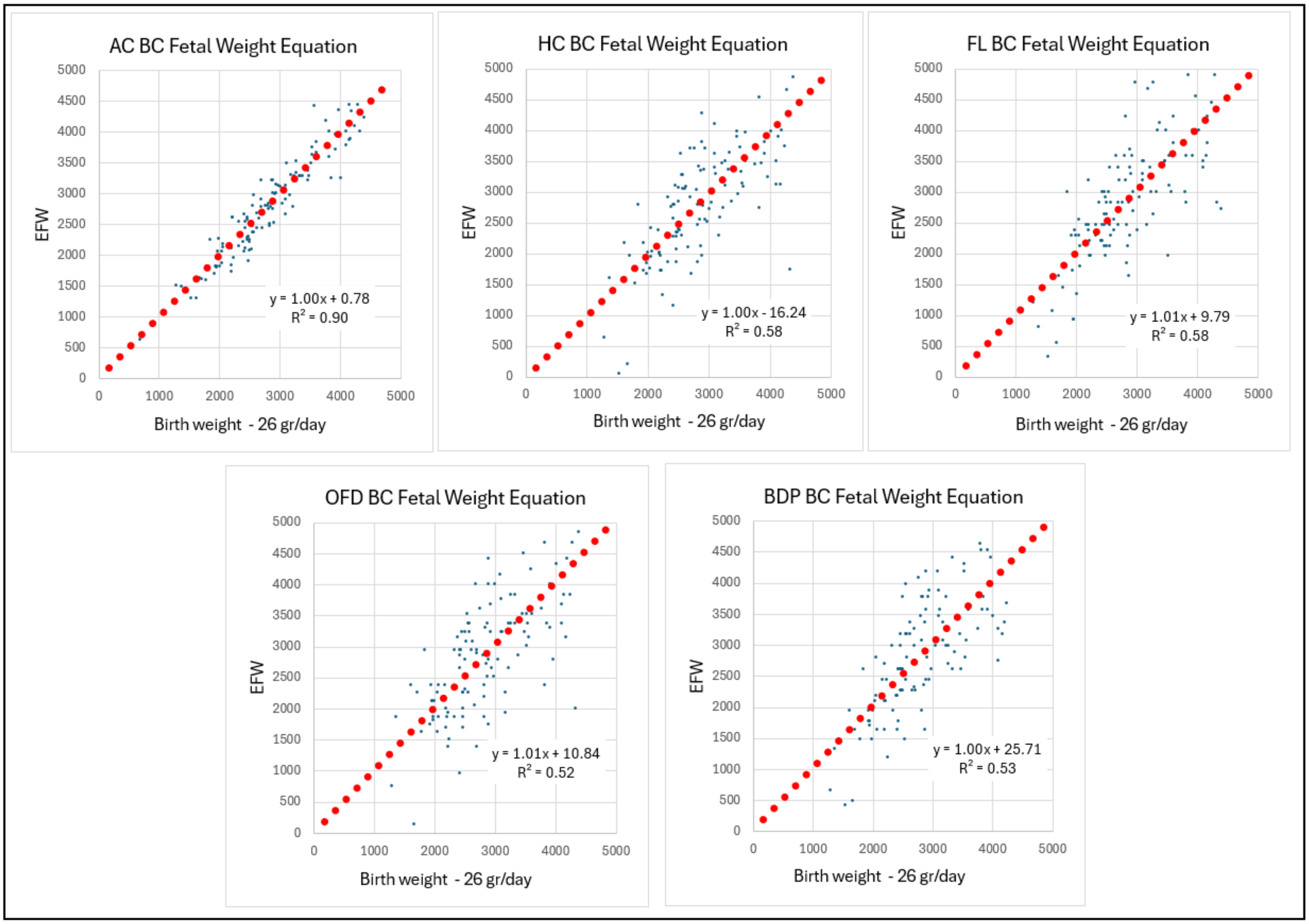


C:_1\BBCDG V7 DATA_______Baby Math Data 10 20 23_____New EFW Calcs Nov 4 23__medium to medium 2 23 2024\[Deid POPs and LIFECODES Combined. Slope & Beta Adjustable. V20b 4 9 2024 Train & Test. 26 gr-day in Train & Test.xlsx]Train.7d'!$AM$218

**Supplement-Figure-II.8.** Fit of the of ***Fetal Ultrasound*** measurements carried out within 7 days of birth vs ***Birthweight***-(***days-until-birth*26 grams***), to the ***Binary Cellular Estimated Fetal Weight*** Equation #14, in the ***Training Dataset***. Similar results were found when examined against ***Birthweight*** alone (not shown).

Testing ***Binary Cellular Estimated Fetal Weight Equations***: Comparing Equations

Overview.

Having determined the values for the parameters of the ***Binary Cellular Estimated Fetal Weight Equations***, derived from the ***Training Dataset*** (TABLE), we then tested their performance against the ***Testing Dataset***, along with other widely used ***Estimated Fetal Weight Equations***. Below, we shall examine the performance of each of these equations, when tested against the ***Testing Dataset***, focusing first on ***Birthweight***-(***days-until-birth****26 ***grams***). We shall also first examine the performance of the ***Abdominal Circumference Binary Cellular Estimated Fetal Weight Equation*** (#15) against these other expression: the ***INTERGROWTH*** and ***Hadlock-1***,***-2***,***-3***,***-4***, and ***-5 Estimated Fetal Weight Equations***, for a total of seven ***Estimated Fetal Weight Equations***. In the sections that follow, we shall examine other ***Binary Cellular Estimated Fetal Weight Equations*** (for ***Biparietal Diameter*** (BPD); ***Occipitofrontal Diameter*** (OFD); ***Femur Length*** (FL) and ***Head Circumference*** (HC)), as well as other aspects of these ***Ultrasound Measures***. The impact of sonographer performance will also be examined, as well as how ***Binary Cellular Estimated Fetal Weight Equations*** can be brought to bear to assist improvement. Performance against ***Birthweight*** will also be examined in comparison to performance against ***Birthweight***-(***days-until-birth****26 ***grams***), to gain insight into the relationship of ***Ultrasound Measurement*** to ***Birthweight*** and ***Fetal Weight***.

Methods.

For each of seven ***Estimated Fetal Weight Equations*** noted above (the ***Abdominal Circumference Binary Cellular Estimated Fetal Weight Equation*** (#15), and the ***INTERGROWTH*** and ***Hadlock-1***,***-2***,***-3***,***-4***, and ***-5 Estimated Fetal Weight Equations***) estimates of fetal weight were made for the 117 ultrasound examinations in the ***Testing Dataset*** that were carried out within 7 days of birth. Error values (EFW-***Birthweight***-(***days-until-birth****26 ***grams***)/***Birthweight***-(***days-until-birth****26 ***grams***)) were calculated for each of the ***Estimated Fetal Weight Equations***, as well as ***Absolute Error*** calculations. The mean and median values, in percentages, and grams, were determined for seven ***Estimated Fetal Weight Equations*** for each visit of the 117 ultrasound examinations in the ***Testing Dataset***. ***Standard Deviation*** of the ***Error*** was also calculated, and from these were calculated the minimum error, in grams, expected for ~1-in-300 ultrasound measurements, which is +/- (1 ***Standard Deviation*** x average birthweight), as wells as the minimum error, in grams, expected for ~1-in-20 ultrasound measurements, which is +/-(2 ***Standard Deviations*** x average birthweight), and the minimum error, in grams, expected for ~1-in-300 ultrasound measurements, which is +/- (3 ***Standard Deviations*** x average birthweight). The ***Mean*** and ***Median*** values for the ***Error*** and ***Absolute Error*** in grams are shown at the same scale as these measures of ***Standard Deviation*** variation, so that their relative impacts can be seen.

***Mean*** and ***Median Error***.

For these seven ***Estimated Fetal Weight Equations***, the ***Mean*** and ***Median Errors*** (**Supplement-Figure-II.9**) and ***Absolute Errors*** (**Supplement-Figure-II.10**) for the ***Abdominal Circumference Binary Cellular Estimated Fetal Weight Equation*** (#14c), proved to be superior to these generated by the ***INTERGROWTH*** and ***Hadlock-1***,***-2***,***-3***,***-4***, and ***-5 Estimated Fetal Weight Equations***. For the ***Abdominal Circumference Binary Cellular Estimated Fetal Weight Equation*** (#15), the ***Mean Error*** was 0.5%, while the next, the ***Hadlock-4 Estimated Fetal Weight Equation*** had a ***Mean Error*** of 5.4%. In terms of ***Birthweight***, for the ***Abdominal Circumference Binary Cellular Estimated Fetal Weight Equation*** (#15), the ***Mean Error*** was 18 grams, while for the ***Hadlock-4 Estimated Fetal Weight Equation***, the ***Mean Error*** was 186 grams.

***Standard Deviation***.

The ***Standard Deviations*** of all 7 ***Estimated Fetal Weight Equations*** were quite similar, but all quite large, displaying a much greater problem than the ***Mean*** and ***Median Errors*** (**Supplement-Figures-II.9&10**). The PEAK of the Bell Curve is roughly where it should be, but the FLANKS stretch WAY out.

The Problem for ***Ultrasounds*** isn't that, on Average, All are a BIT Off: It's that a Few are a LOT Off.

Among all 7 ***Estimated Fetal Weight Equations*** shown in **Supplement-Figures-II.9&10**, the average ***Error*** for predicting ***Birthweight*** among all newborns varies from as little as 18 grams for the ***Abdominal Circumference Binary Cellular Estimated Fetal Weight Equation*** (#15) to as much 194 grams for the ***Hadlock-5 Estimated Fetal Weight Equation.*** In contrast, the values for the ***Standard Deviation*** of the ***Error*** show us that the prediction of ***Birthweight*** for ~1-in-300 fetuses is off by 892 grams (when estimated by the ***Hadlock-4 Estimated Fetal Weight Equation***), and off by 1099 grams (when estimated by the ***Hadlock-5 Estimated Fetal Weight Equation***), and off by 1029 grams (when estimated by the ***Abdominal Circumference Binary Cellular Estimated Fetal Weight Equation***). The prediction of ***Birthweight*** for ~1-in-20 fetuses is off by 595 grams (when estimated by the ***Hadlock-4 Estimated Fetal Weight Equation***), and off by 733 grams (when estimated by the ***Hadlock-5 Estimated Fetal Weight Equation***), and off by 686 grams (when estimated by the ***Abdominal Circumference Binary Cellular Estimated Fetal Weight Equation***). The prediction of ***Birthweight*** for ~1-in-3 fetuses is off by 297 grams (when estimated by the ***Hadlock-4 Estimated Fetal Weight Equation***), and off by 366 grams (when estimated by the ***Hadlock-5 Estimated Fetal Weight Equation***), and off by 343 grams (when estimated by the ***Abdominal Circumference Binary Cellular Estimated Fetal Weight Equation***).

The Problem isn't the ***Error*** but the ***Standard Deviation*** of the ***Error***.

These calculations show that it's not the ***Mean Error*** of ***Fetal Weight*** estimation by ultrasound that causes the most trouble, but the ***Standard Deviation*** of the ***Error***, a problem we see regardless of which of the 7 ***Estimated Fetal Weight Equations*** are used (**Supplement-Figures-II.9&10**). The distinction is a bit like the difference between how much over the limit is the ***Average Speed*** of all the cars on the highway, vs how much over the limit is the 1-in-300$^{th}$ car that is going at the ***Fastest Speed***. And when even the 1-in-10$^{th}$ ***Fastest*** car is going much fater than the ***Average*** car, we have a problem. Below, and in the DISCUSSION, we shall expand upon these features of the ***Mean Error*** and ***Standard Deviation*** of the ***Error*** of the ***Estimated Fetal Weight***, and how their implications can be understood and minimized with the help of ***Binary Cellular Analysis***.

The Problem Isn't Which ***Estimated Fetal Weight Equation*** is Used.

The ***Standard Deviations*** of all seven ***Estimated Fetal Weight Equations***, were quite similar, with ***Hadlock-4***, at 8.7%, being very slightly better than the ***Abdominal Circumference Binary Cellular Estimated Fetal Weight Equation*** (#15) at 10.0% (**Supplement-Figure-II.9**). However, the practical difference is negligible; the greatest impact, which could be expected for ~1-in-300 ultrasound measurements, displays a difference between ***Abdominal Circumference Binary Cellular Estimated Fetal Weight Equation*** (#15), and the ***Hadlock-4 Estimated Fetal Weight Equation***, was a modest 137 grams, roughly 4% of the weight of an average newborn. For 2 out of 3 ultrasound measurements, the difference between was an even more modest, at 28 grams, roughly 1% of the weight of an average newborn.

Looked at from another standpoint, for every 2-out-of 3 ultrasound measurements, the difference between the two equations is less than 28 grams, ~1% of the weight of an average newborn. For every 299-out-of-300 ultrasound measurement, the difference between to the equations is less than 137 grams, ~4% of the weight of an average newborn.

As we shall see below, the source of these disappointing ***Standard Deviations*** seen for all 7 ***Estimated Fetal Weight Equations*** lies not in their calculations of ***Fetal Weight***, but in the measurement of ***Fetal Size*** by ultrasound, when the patient was examined by the sonographer. Fortunately, as we shall see, this is a problem for which ***Binary Cellular Analysis*** can offer assistance.

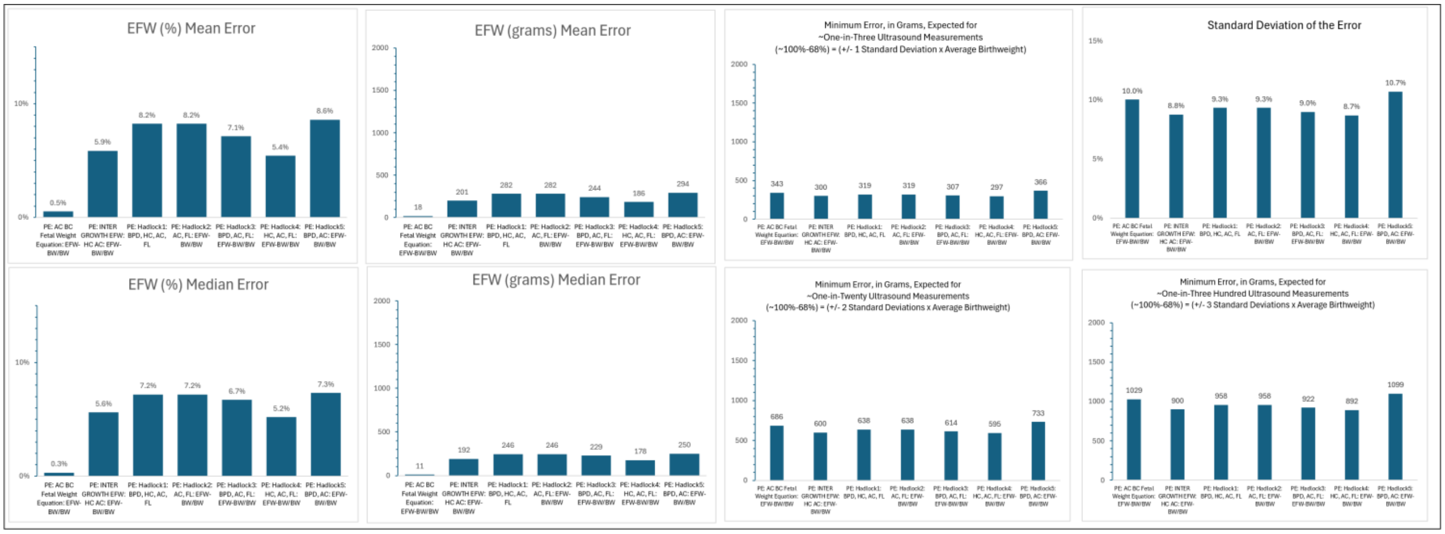


C:_1\BBCDG V7 DATA_______Baby Math Data 10 20 23_____New EFW Calcs Nov 4 23__medium to medium 2 23 2024 [Deid POPs and LIFECODES Combined. Slope & Beta Adjustable. V20c 4 11 2024 Train & Test. 26 gr-day in Train & Test.xlsx]Test.7d 4th (2.5)'!$DS$10
Checked 11/11/2024: Deid POPs and LIFECODES Combined. Slope & Beta Adjustable. V20d 4 12 2024 Train & Test. 26 gr-day in Train & Test.xlsx ::: updated to: Deid POPs and LIFECODES Combined. Slope & Beta Adjustable. V20e 11 11 2024 Train & Test. 26 gr-day in Train & Test.xlsx

**Supplement-Figure-II.9** ***Mean*** and ***Median Errors*** and ***Standard Deviation*** of the ***Error***, or seven ***Estimated Fetal Weight Equations***: ***Abdominal Circumference Binary Cellular Estimated Fetal Weight Equation*** (#14c), ***INTERGROWTH Estimated Fetal Weight Equation***, ***Hadlock-1***,***-2***,***-3***,***-4***, and ***-5 Estimated Fetal Weight Equations***, in the ***Testing Dataset***, based on ultrasound carried out within 7 days of birth.

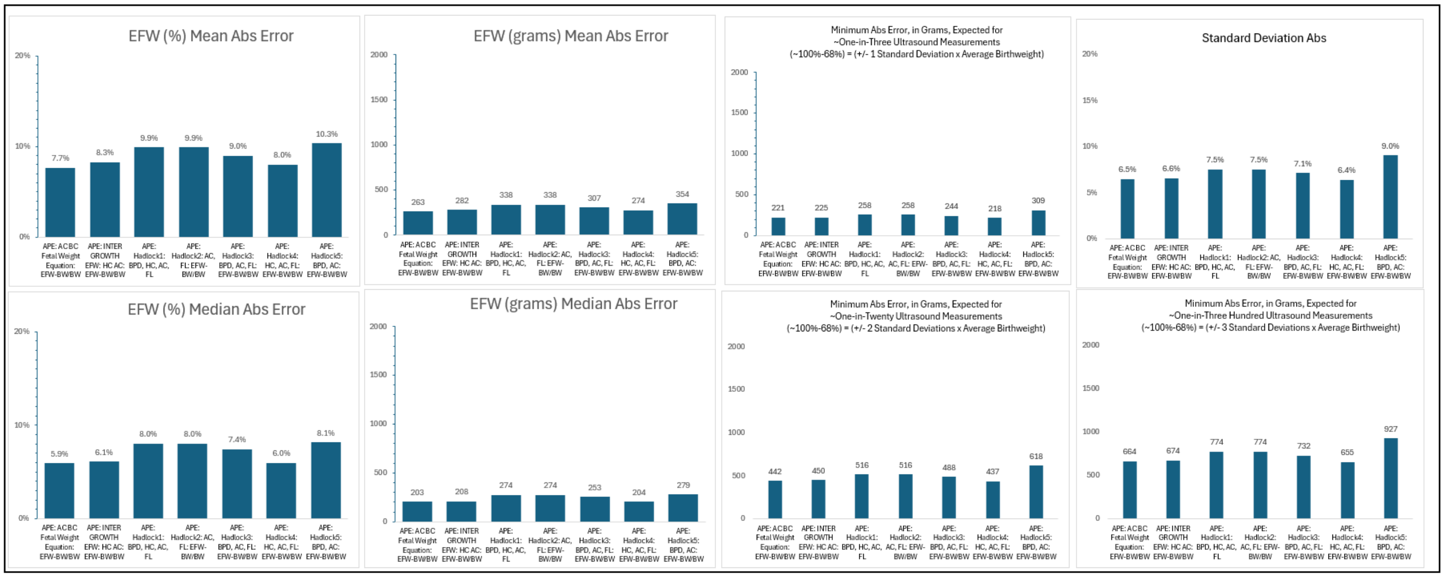


C:_1\BBCDG V7 DATA_______Baby Math Data 10 20 23_____New EFW Calcs Nov 4 23__medium to medium 2 23 2024\ [Deid POPs and LIFECODES Combined. Slope & Beta Adjustable. V20c 4 11 2024 Train & Test. 26 gr-day in Train & Test.xlsx]Test.7d 4th (2.5)'!$DS$10

**Supplement-Figure-II.10.** ***Absolute Errors*** and ***Standard Deviation*** of the ***Error*** for seven ***Estimated Fetal Weight Equations***: ***Abdominal Circumference Binary Cellular Estimated Fetal Weight Equation*** (#14c), ***INTERGROWTH Estimated Fetal Weight Equation***, ***Hadlock-1***,***-2***,***-3***,***-4***, and ***-5 Estimated Fetal Weight Equations***, in the ***Testing Dataset***, based on ultrasound carried out within 7 days of birth.

Testing ***Binary Cellular Estimated Fetal Weight Equations***: Comparing Equations (cont.)

Assessment of Small and Large Newborns

As noted above, many ***Estimated Fetal Weight Equations*** have been degraded by an inability to accurately capture the sizes of fetuses that will become ***Large*** and ***Small Babies*** (see METHODS section). This can be seen by linear regression of the comparison of ***Estimated Fetal Weight*** vs ***Birthweight***-(***days-until-birth*26 grams***) (**Supplement-Figure-II.11**) and ***Birthweight*** (**Supplement-Figure-II.12**) and by examining mean percentage error as a function of ***Birthweight*** (**Supplement-Figure-II.13**). For ***INTERGROWTH*** and ***Hadlock-1***,***-2***,***-3***,***-4***, and ***-5 Estimated Fetal Weight Equations***, these expressions overestimate the ***Weight*** for fetuses that become newborns of ***Small Size***, but this is not the case for the ***Abdominal Circumference Binary Cellular Estimated Fetal Weight Equation*** (#15), which was equally precise for fetuses that become newborns of all sizes.

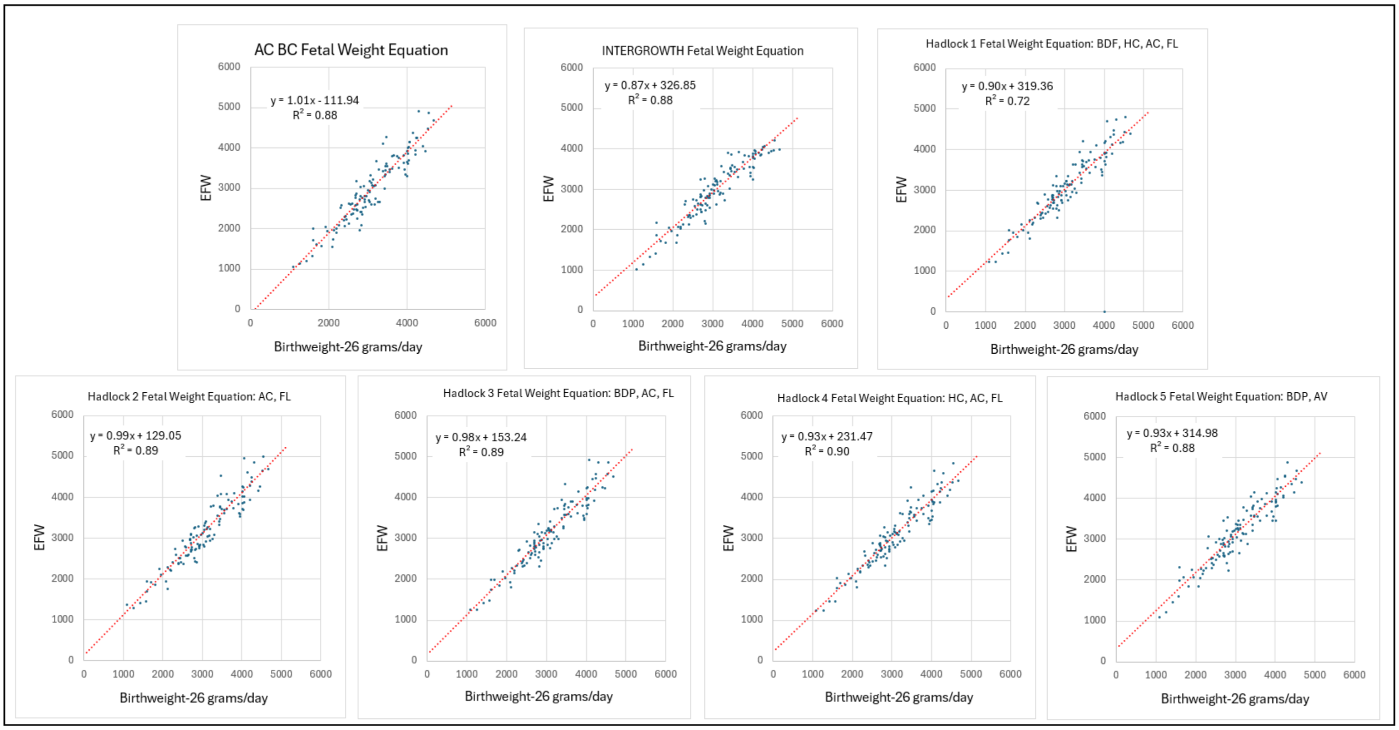


**Supplement-Figure-II.11.** Comparison of ***Estimated Fetal Weight*** vs ***Birthweight***-(***days-until-birth*26 grams***) for seven ***Estimated Fetal Weight Equations***: ***Abdominal Circumference Binary Cellular Estimated Fetal Weight Equation*** (#14c), ***INTERGROWTH Estimated Fetal Weight Equation***, ***Hadlock-1***,***-2***,***-3***,***-4***, and ***-5 Estimated Fetal Weight Equations***, in the ***Testing Dataset***, based on ultrasound carried out within 7 days of birth.

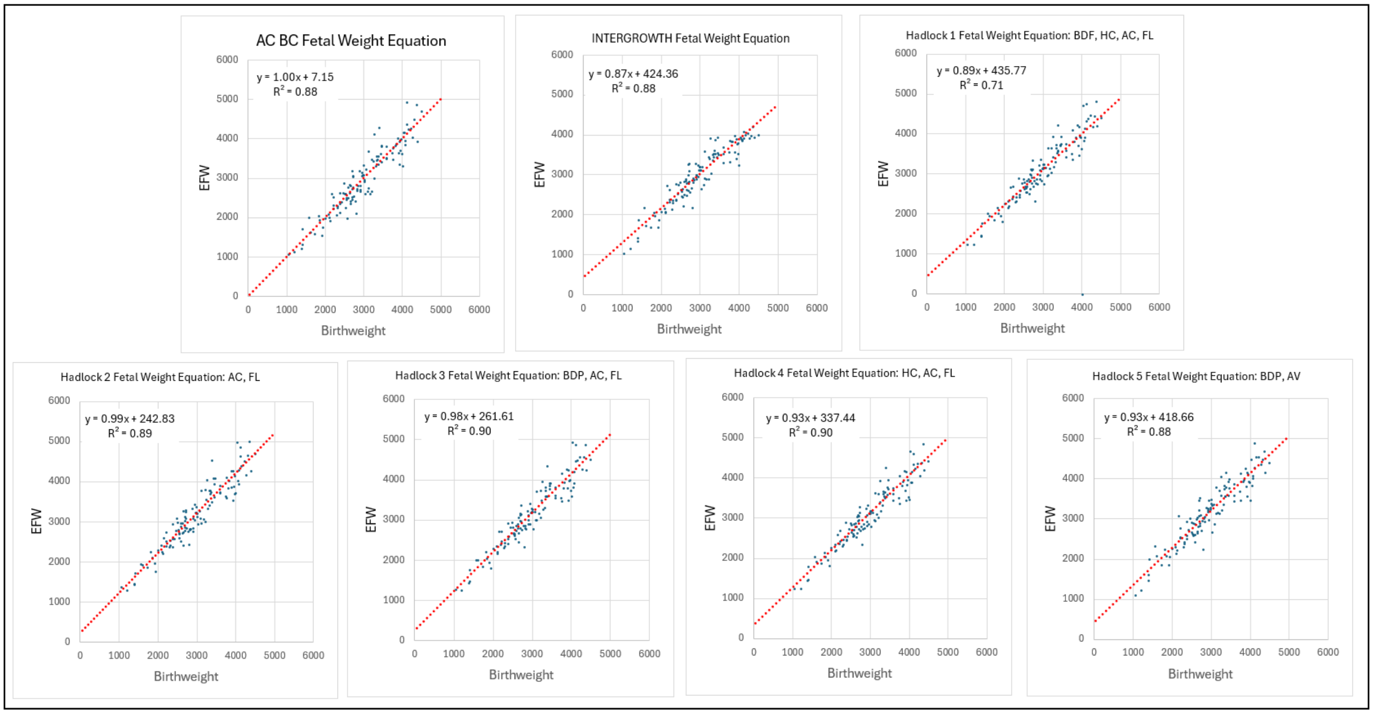


**<u>Supplement-Figure-II.12</u>** Comparison of ***Estimated Fetal Weight*** vs ***Birthweight*** for seven ***Estimated Fetal Weight Equations***: ***Abdominal Circumference Binary Cellular Estimated Fetal Weight Equation*** (#14c), ***INTERGROWTH Estimated Fetal Weight Equation***, ***Hadlock-1,-2,-3,-4***, and ***-5 Estimated Fetal Weight Equations***, in the ***Testing Dataset***, based on ultrasound carried out within 7 days of birth.

C:\ 1\BBCDG V7 DATA\ Baby Math Data 10 20 23\ New EFW Calcs Nov 4 23\ medium to medium 2 23 2024\ [Deid POPs and LIFECODES Combined. Slope & Beta Adjustable. V20c 4 11 2024 Train & Test. 26 gr-day in Train & Test.xlsx]Test.7d 4th (1)'!$EV$9

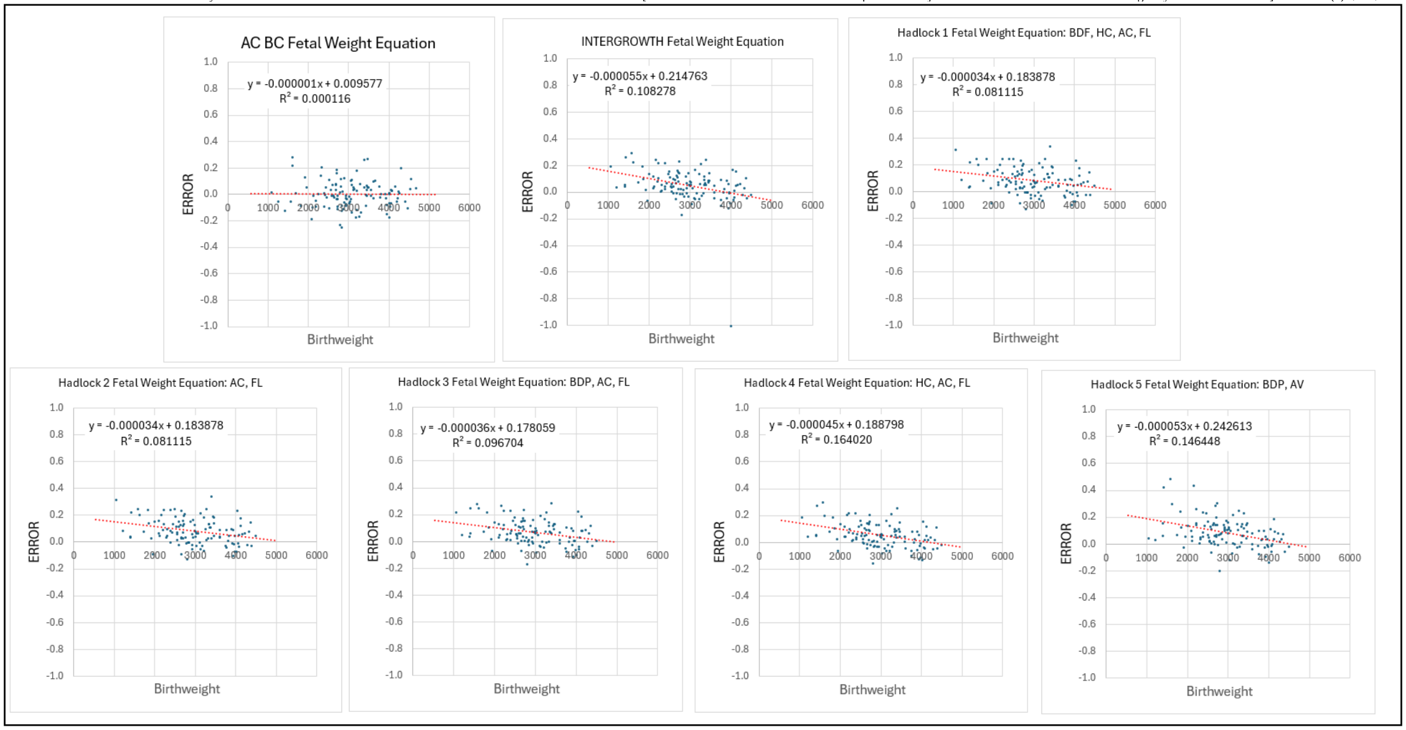


C:_1\BBCDG V7 DATA_______Baby Math Data 10 20 23_____New EFW Calcs Nov 4 23__medium to medium 2 23 2024\ [Deid POPs and LIFECODES Combined. Slope & Beta Adjustable. V20c 4 11 2024 Train & Test. 26 gr-day in Train & Test.xlsx]Test.7d 4th (1)'!$EV$9

**<u>Supplement-Figure-II.13</u>**. Mean percentage error as a function of ***Birthweight*** for seven ***Estimated Fetal Weight Equations***: ***Abdominal Circumference Binary Cellular Estimated Fetal Weight Equation*** (#14c), ***INTERGROWTH Estimated Fetal Weight Equation***, ***Hadlock-1,-2,-3,-4***, and ***-5 Estimated Fetal Weight Equations***, in the ***Testing Dataset***, based on ultrasound carried out within 7 days of birth.

Testing ***Binary Cellular Estimated Fetal Weight Equations***: Comparison of ***Body Parts***

As its name indicates, the ***Abdominal Circumference Binary Cellular Estimated Fetal Weight Equation*** (#15) is based on a single ***Ultrasound Measurement***, while the other ***Estimated Fetal Weight Equations*** (***INTERGROWTH*** and ***Hadlock-1***,***-2***,***-3***,***-4***, and***-5***), utilize multiple ***Ultrasound Measurements***. In addition to ***Abdominal Circumference*** (AC), we also have measurements of ***Biparietal Diameter*** (BPD); ***Occipitofrontal Diameter*** (OFD); ***Femur Length*** (FL); ***Head Circumference*** (HC). ***Binary Cellular Biology*** gives us insight into the potential of these measurements to estimate ***Fetal Weight***, by generating ***Binary Cellular Estimated Fetal Weight Equations*** for each of these ***Ultrasound Measurements***, and examining their performances.

In agreement with the crude comparisons of each of these ***Ultrasound Measurements*** to ***Birthweight*** among the 117 ultrasound examinations in the ***Training Dataset*** recorded within 7 days of birth (**Supplement-Figure-II.9-13**), comparison of the performance of these five ***Binary Cellular Estimated Fetal Weight Equations***, reveals the striking superiority of ***Abdominal Circumference Binary Cellular Estimated Fetal Weight Equation*** (#15) to the ***Biparietal Diameter*** (BPD)***, Occipitofrontal Diameter*** (OFD), ***Femur Length*** (FL)**,** and ***Head Circumference*** (HC) ***Binary Cellular Estimated Fetal Weight Equations***. This could be seen by examining data on the 117 ultrasound examinations in the ***Testing Dataset*** recorded within 7 days of birth (**Supplement-Figure-II.14-18**).

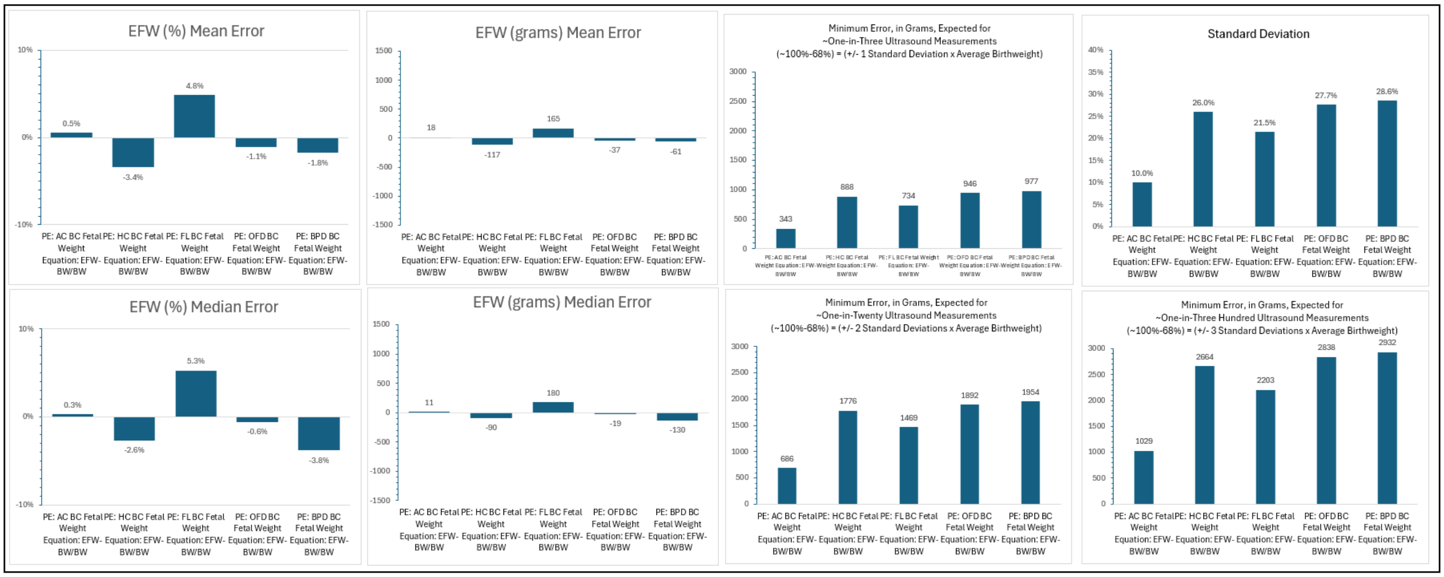

C:_1\BBCDG V7 DATA_______Baby Math Data 10 20 23_____New EFW Calcs Nov 4 23__medium to medium 2 23 2024\ [Deid POPs and LIFECODES Combined. Slope & Beta Adjustable. V20c 4 11 2024 Train & Test. 26 gr-day in Train & Test.xlsx]Test.7d 4th (5.2)'!$EV$9

**Supplement-Figure-II.14**. ***Mean*** and ***Median Errors*** and ***Standard Deviation*** of the ***Error*** for ***Abdominal Circumference*** (#14c), ***Biparietal Diameter*** (BPD)***, Occipitofrontal Diameter*** (OFD), ***Femur Length*** (FL)**,** and ***Head Circumference*** (HC) ***Binary Cellular Estimated Fetal Weight Equations***, in the ***Testing Dataset***, based on ultrasound carried out within 7 days of birth.

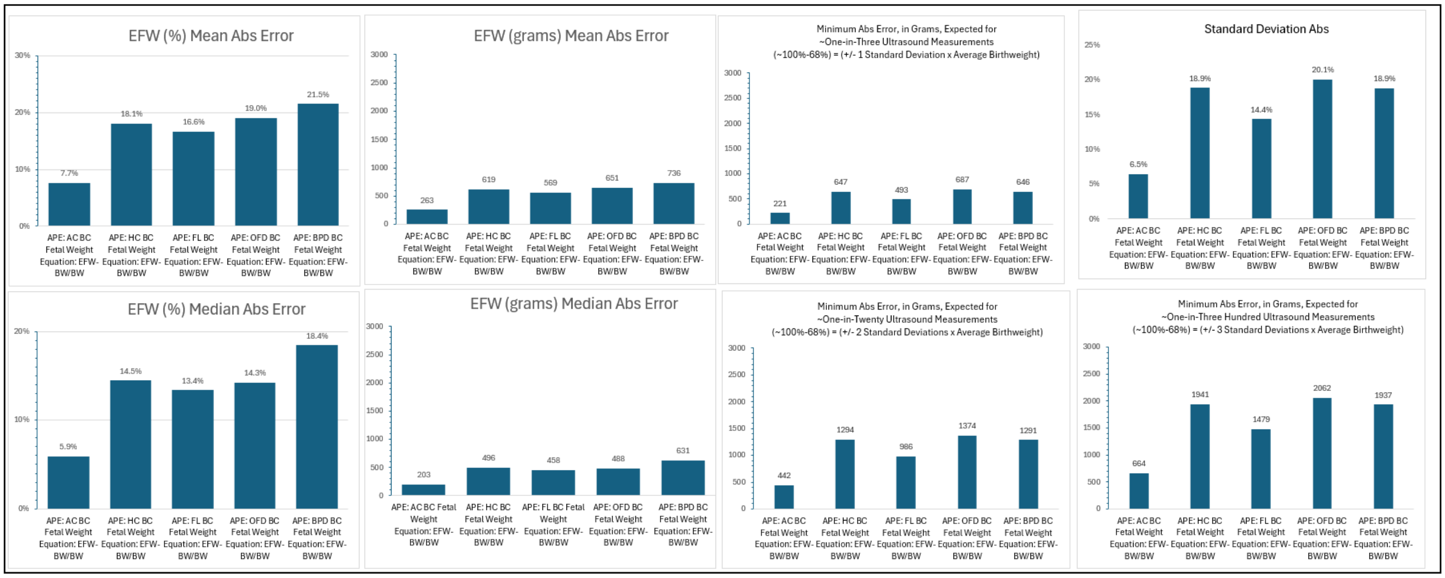


C:_1\BBCDG V7 DATA______Baby Math Data 10 20 23_____New EFW Calcs Nov 4 23__medium to medium 2 23 2024\ [Deid POPs and LIFECODES Combined. Slope & Beta Adjustable. V20c 4 11 2024 Train & Test. 26 gr-day in Train & Test.xlsx]Test.7d 4th (5.2)'!$EV$9

**Supplement-Figure-II.15**, ***Mean*** and ***Median Absolute Errors*** and ***Standard Deviation*** of the ***Error*** for ***Abdominal Circumference*** (#14c), ***Biparietal Diameter*** (BPD)**,** ***Occipitofrontal Diameter*** (OFD), ***Femur Length*** (FL)**,** and ***Head Circumference*** (HC) ***Binary Cellular Estimated Fetal Weight Equations***, in the ***Testing Dataset***, based on ultrasound carried out within 7 days of birth.

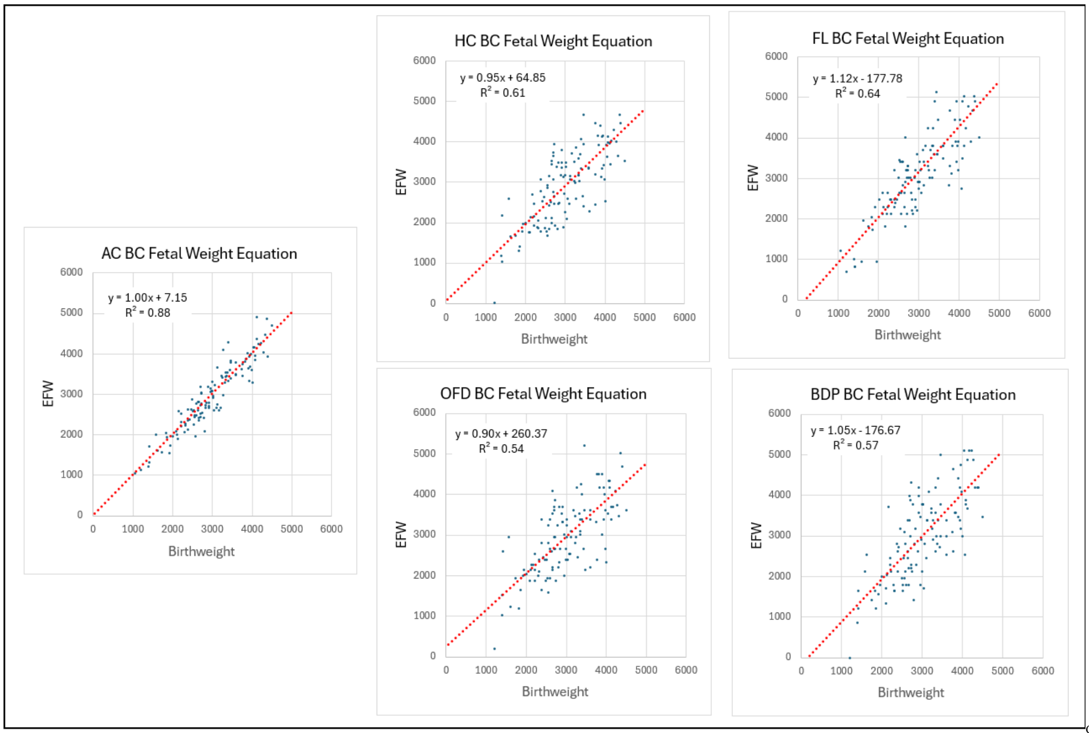


C:_1\BBCDG V7 DATA______Baby Math Data 10 20 23_____New EFW Calcs Nov 4 23__medium to medium 2 23 2024\ [Deid POPs and LIFECODES Combined. Slope & Beta Adjustable. V20c 4 11 2024 Train & Test. 26 gr-day in Train & Test.xlsx]Test.7d 4th (5)'!$EV$9

**Supplement-Figure-II.16**. Comparison of ***Birthweight*** with ***Abdominal Circumference*** (#14c), ***Biparietal Diameter*** (BPD)**,** ***Occipitofrontal Diameter*** (OFD), ***Femur Length*** (FL)**,** and ***Head Circumference*** (HC) ***Binary Cellular Estimated Fetal Weight Equations***, in the ***Testing Dataset***, based on ultrasound carried out within 7 days of birth.

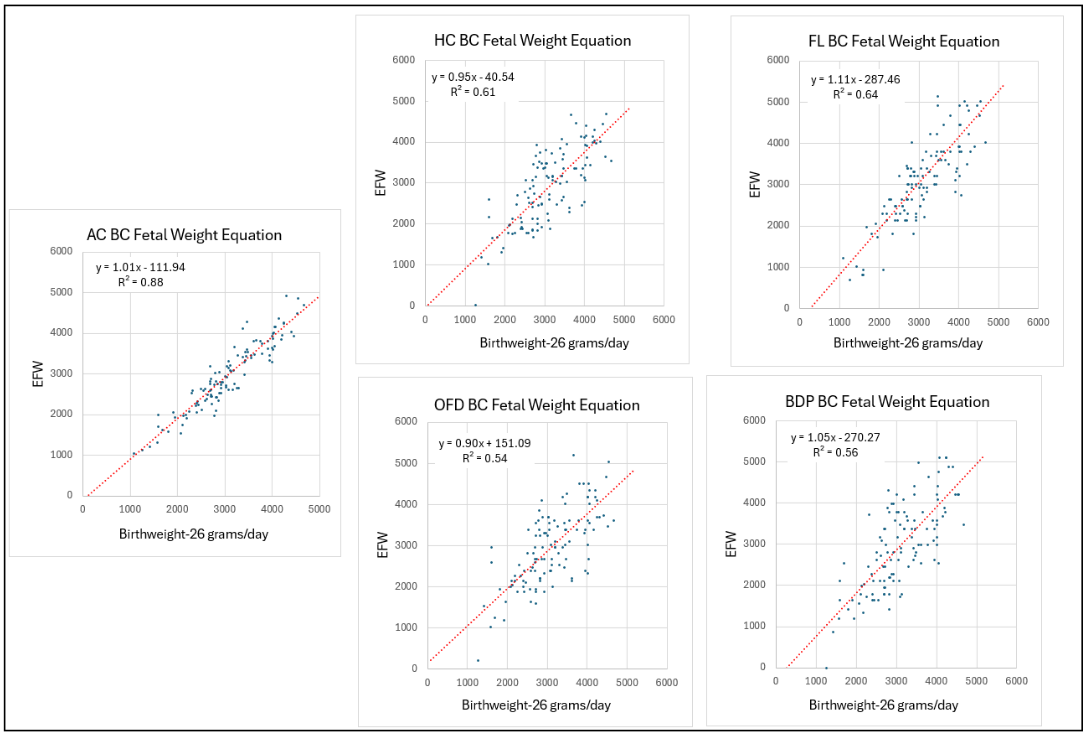


C:_1\BBCDG V7 DATA_______Baby Math Data 10 20 23_____New EFW Calcs Nov 4 23__medium to medium 2 23 2024\ [Deid POPs and LIFECODES Combined. Slope & Beta Adjustable. V20c 4 11 2024 Train & Test. 26 gr-day in Train & Test.xlsx]Test.7d 4th (5)'!$EV$9

**Supplement-Figure-II.17**, Comparison of ***Birthweight-(days-until-birth*26 grams)*** with ***Abdominal Circumference*** (#14c), ***Biparietal Diameter*** (BPD)**,** ***Occipitofrontal Diameter*** (OFD), ***Femur Length*** (FL)**,** and ***Head Circumference*** (HC) ***Binary Cellular Estimated Fetal Weight Equations***, based on ultrasound carried out within 7 days of birth.

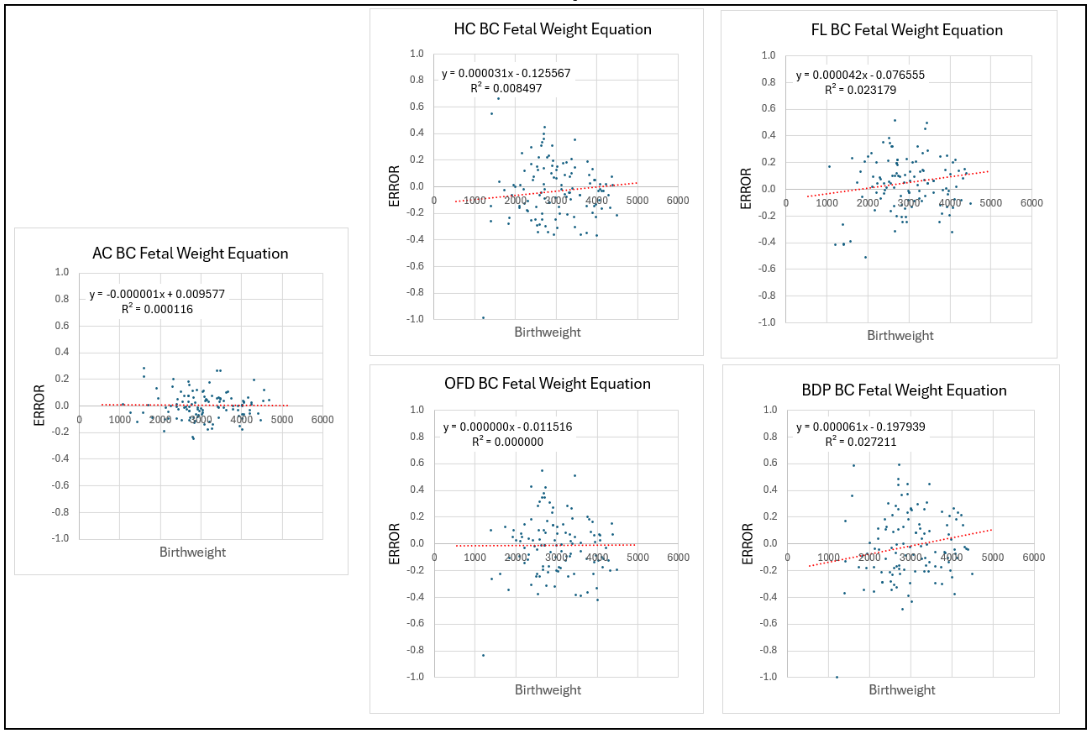


C:_1\BBCDG V7 DATA_______Baby Math Data 10 20 23_____New EFW Calcs Nov 4 23__medium to medium 2 23 2024\ [Deid POPs and LIFECODES Combined. Slope & Beta Adjustable. V20c 4 11 2024 Train & Test. 26 gr-day in Train & Test.xlsx]Test.7d 4th (5.2)'!$EV$9

**Supplement-Figure-II.18**, Mean percentage error as a function of ***Birthweight*** for ***Abdominal Circumference*** (#14c), ***Biparietal Diameter*** (BPD)**,** ***Occipitofrontal Diameter*** (OFD), ***Femur Length*** (FL)**,** and ***Head Circumference*** (HC) ***Binary Cellular Estimated Fetal Weight Equations***, in the ***Testing Dataset***, based on ultrasound carried out within 7 days of birth.

Testing ***Binary Cellular Estimated Fetal Weight Equations***: Averaging of Multiple ***Body Parts***

Despite the striking superiority of ***Abdominal Circumference Binary Cellular Estimated Fetal Weight Equation*** (#15) to the ***Femur Length Binary Cellular Estimated Fetal Weight Equation,*** and ***Head Circumference Binary Cellular Estimated Fetal Weight Equation***, averaging them yielded almost the same outcome as the ***Abdominal Circumference Binary Cellular Estimated Fetal Weight Equation*** (#15) alone (**Supplement-Figure-II.19-23**). While the Average of the ***Abdominal Circumference / Femur Length / Head Circumference Binary Cellular Estimated Fetal Weight Equations***, does not get quite as good as the ***Abdominal Circumference Binary Cellular Estimated Fetal Weight Equation*** (#15) alone, it does get quite close. This suggests practical improvements of these measurements may make averaging a way to smooth out and improve the performance of any single measurement.

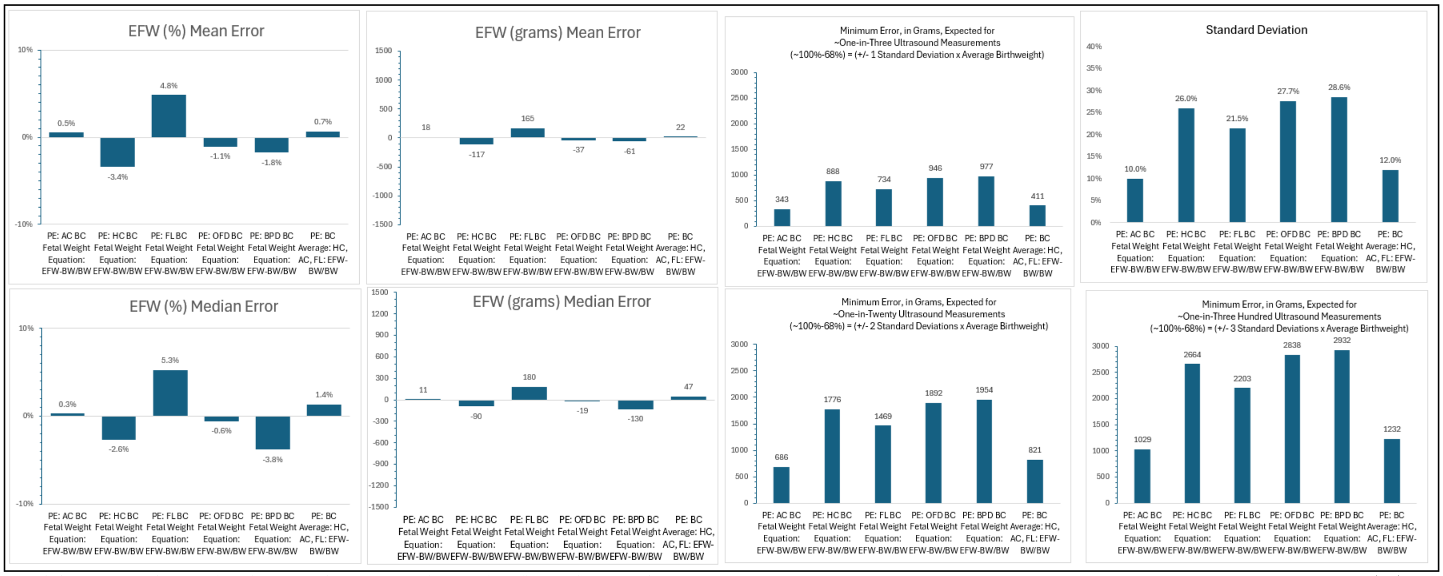


C:_1\BBCDG V7 DATA_______Baby Math Data 10 20 23_____New EFW Calcs Nov 4 23__medium to medium 2 23 2024\ [Deid POPs and LIFECODES Combined. Slope & Beta Adjustable. V20c 4 11 2024 Train & Test. 26 gr-day in Train & Test.xlsx]Test.7d 4th (3.5)'!$EV$9

**Supplement-Figure-II.19**, ***Mean*** and ***Median Errors*** and ***Standard Deviation*** of the ***Error*** of Averaged of ***Binary Cellular Estimated Fetal Weight Equations*** for various ultrasound measurements, of babies in the ***Testing Dataset***, based on ultrasounds carried out within 7 days of birth.

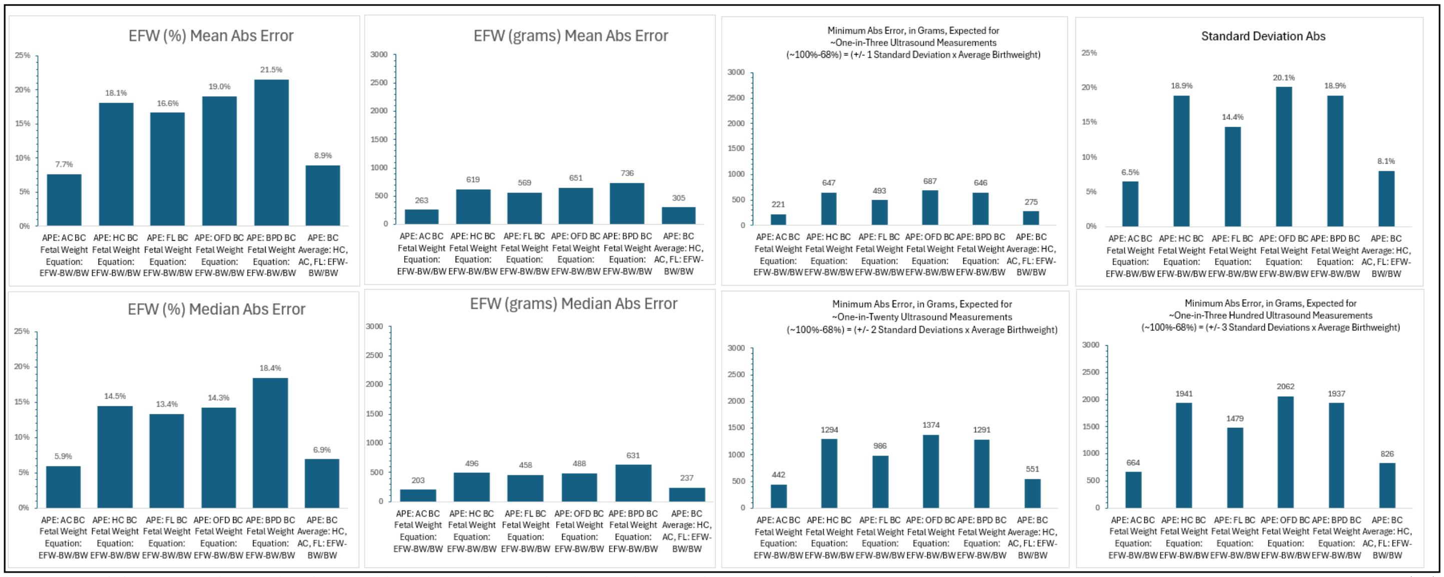


C:_1\BBCDG V7 DATA_______Baby Math Data 10 20 23_____New EFW Calcs Nov 4 23__medium to medium 2 23 2024\ [Deid POPs and LIFECODES Combined. Slope & Beta Adjustable. V20c 4 11 2024 Train & Test. 26 gr-day in Train & Test.xlsx]Test.7d 4th (4)'!$EV$9

**Supplement-Figure-II.20**, ***Mean*** and ***Median Absolute Errors*** and ***Standard Deviation*** of the ***Error*** of Averaged of ***Binary Cellular Estimated Fetal Weight Equations*** for various ultrasound measurements, of babies in the ***Testing Dataset***, based on ultrasounds carried out within 7 days of birth.

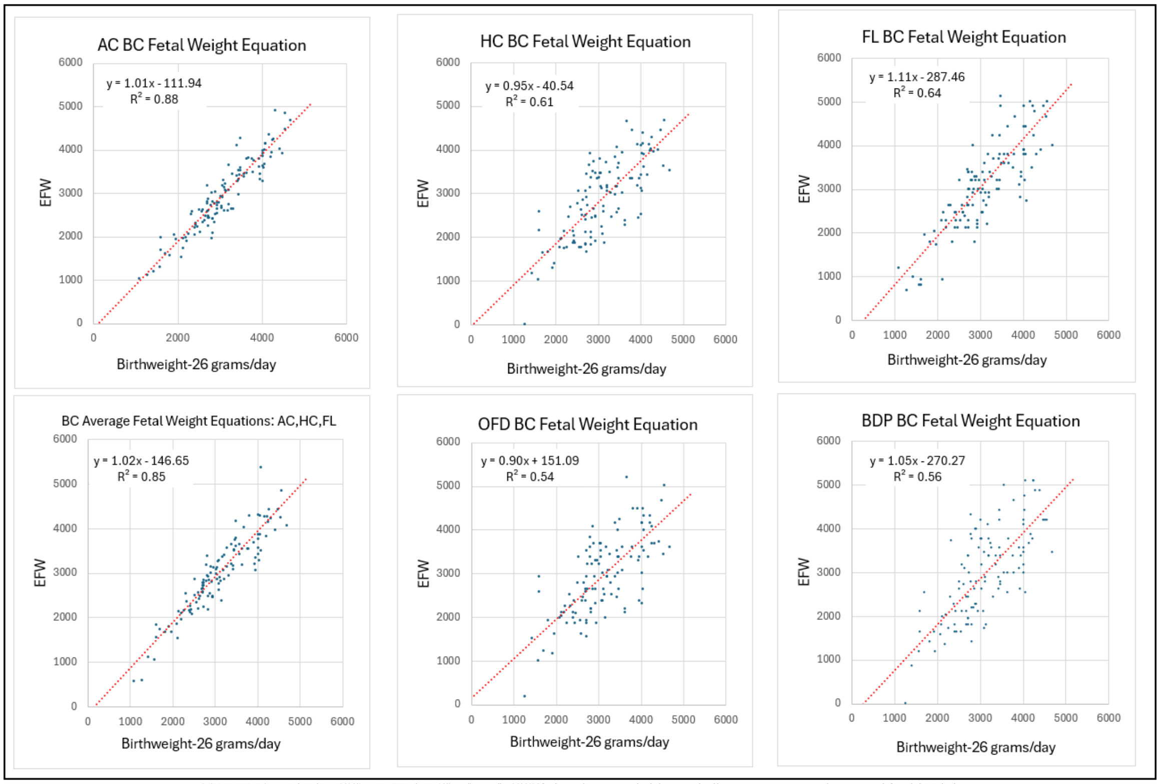


C:_1\BBCDG V7 DATA_____Baby Math Data 10 20 23____New EFW Calcs Nov 4 23__medium to medium 2 23 2024\ [Deid POPs and LIFECODES Combined. Slope & Beta Adjustable. V20c 4 11 2024 Train & Test. 26 gr-day in Train & Test.xlsx]Test.7d 4th (3.5)'!$EV$9

**Supplement-Figure-II.21**. Comparisons of Averaged of ***Binary Cellular Estimated Fetal Weight Equations*** with ***Birthweight*-(*days-until-birth**26 grams**) for various ultrasound measurements of babies in the ***Testing Dataset***, based on ultrasounds carried out within 7 days of birth.

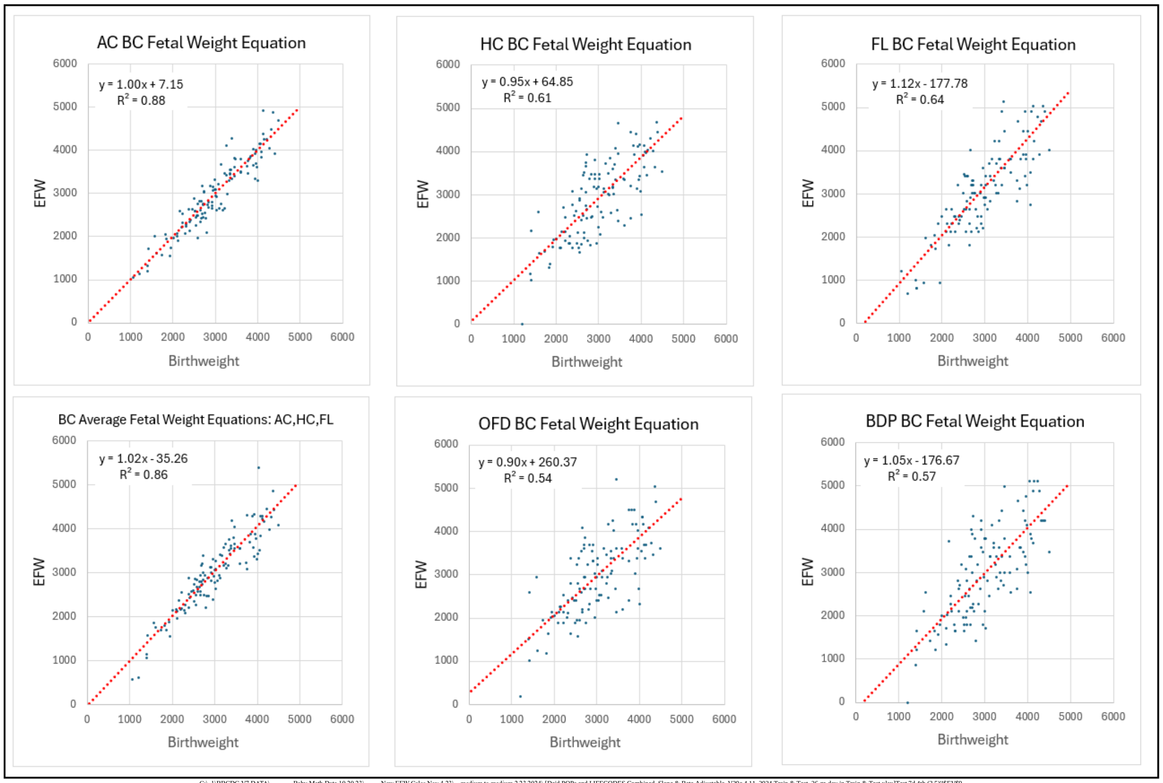


C:_1\BBCDG V7 DATA_____Baby Math Data 10 20 23____New EFW Calcs Nov 4 23__medium to medium 2 23 2024\ [Deid POPs and LIFECODES Combined. Slope & Beta Adjustable. V20c 4 11 2024 Train & Test. 26 gr-day in Train & Test.xlsx]Test.7d 4th (3.5)'!$EV$9

**Supplement-Figure-II.22**, Comparisons of Averaged of ***Binary Cellular Estimated Fetal Weight Equations*** with ***Birthweight*** for various ultrasound measurements of babies in the ***Testing Dataset***, based on ultrasounds carried out within 7 days of birth.

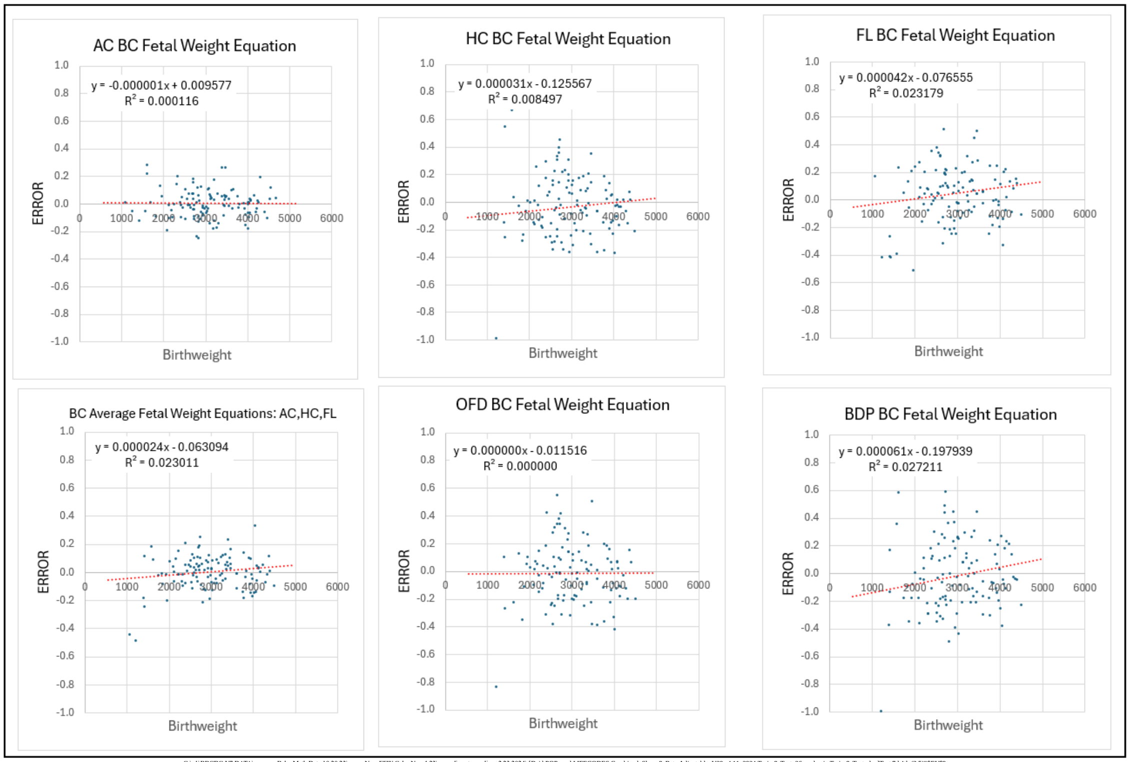


C:_1\BBCDG V7 DATA_______Baby Math Data 10 20 23_____New EFW Calcs Nov 4 23__medium to medium 2 23 2024\ [Deid POPs and LIFECODES Combined. Slope & Beta Adjustable. V20c 4 11 2024 Train & Test. 26 gr-day in Train & Test.xlsx]Test.7d 4th (3.5)'!$EV$9

**Supplement-Figure-II.23**, Mean percentage error as a function of ***Birthweight*** for Averaged of ***Binary Cellular Estimated Fetal Weight Equations*** with ***Birthweight*** for various ultrasound measurements of babies in the ***Testing Dataset***, based on ultrasounds carried out within 7 days of birth.

Testing ***Binary Cellular Estimated Fetal Weight Equations***: The Biology of ***Body Part*** Choice.

Why does the ***Abdominal Circumference*** provide a better estimate of ***Fetal Weight*** than ***Femur Length*** and ***Head Circumference***? As we saw from our first look at crude relationship between ***Ultrasound Measurement*** and ***Birthweight***, seen on ordinary x-y graphs, the ***Abdominal Circumference*** simply makes a steeper line in relationship to ***Birthweight*** than the other ***Ultrasound Measurements***: ***Biparietal Diameter*** (BPD); ***Occipitofrontal Diameter*** (OFD); ***Femur Length*** (FL); and ***Head Circumference*** (HC) (**Supplement-Figure-II.9**). This results in better performance of the ***Abdominal Circumference Binary Cellular Estimated Fetal Weight Equation*** than the ***Head Circumference Binary Cellular Estimated Fetal Weight Equation*** and the ***Femur Length Binary Cellular Estimated Fetal Weight Equation*** (**Supplement-Figures II.24 and 25**). We shall return to the mathematical and biological basis for this finding in the DISCUSSION.

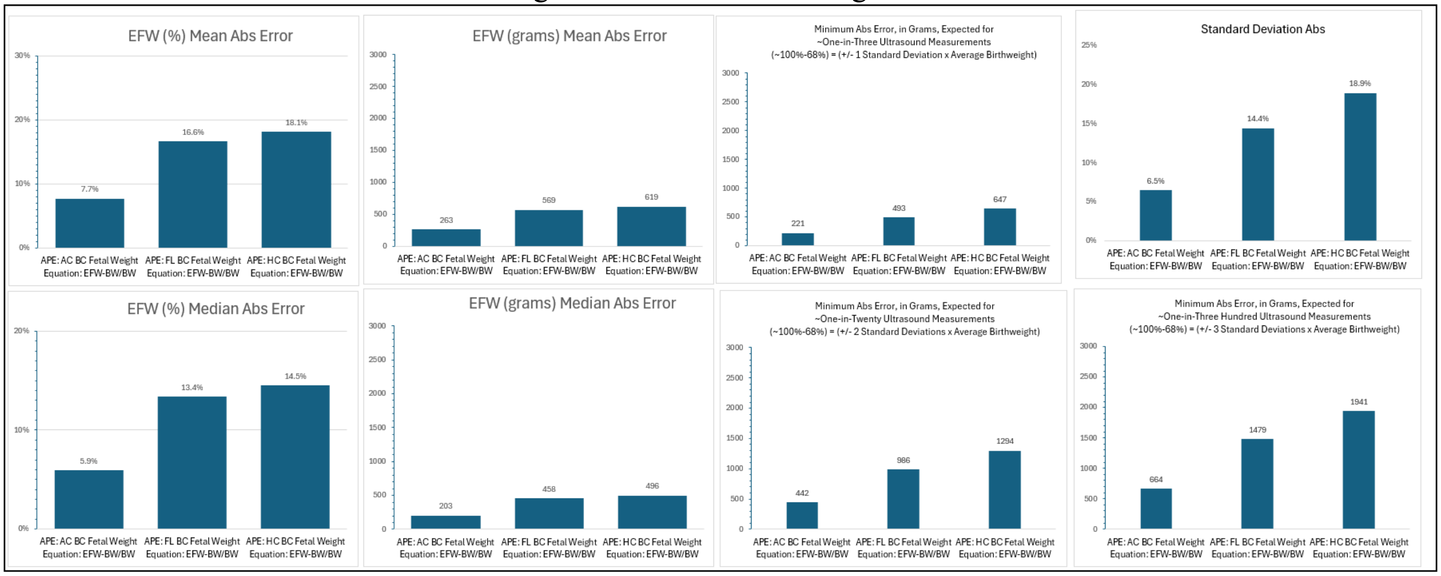


C:_1\BBCDG V7 DATA_______Baby Math Data 10 20 23_____New EFW Calcs Nov 4 23__medium to medium 2 23 2024\ [Deid POPs and LIFECODES Combined. Slope & Beta Adjustable. V20d 4 12 2024 Train & Test. 26 gr-day in Train & Test.xlsx]Test.7d 4th (5.2) (3)'!$DK$50

**Supplement-Figure-II.24**, ***Mean*** and ***Median Errors*** and ***Standard Deviation*** of the ***Error*** for ***Binary Cellular Estimated Fetal Weight Equations***, for abdomen, head, and femur, in the ***Testing Dataset***, based on ultrasound carried out within 7 days of birth.

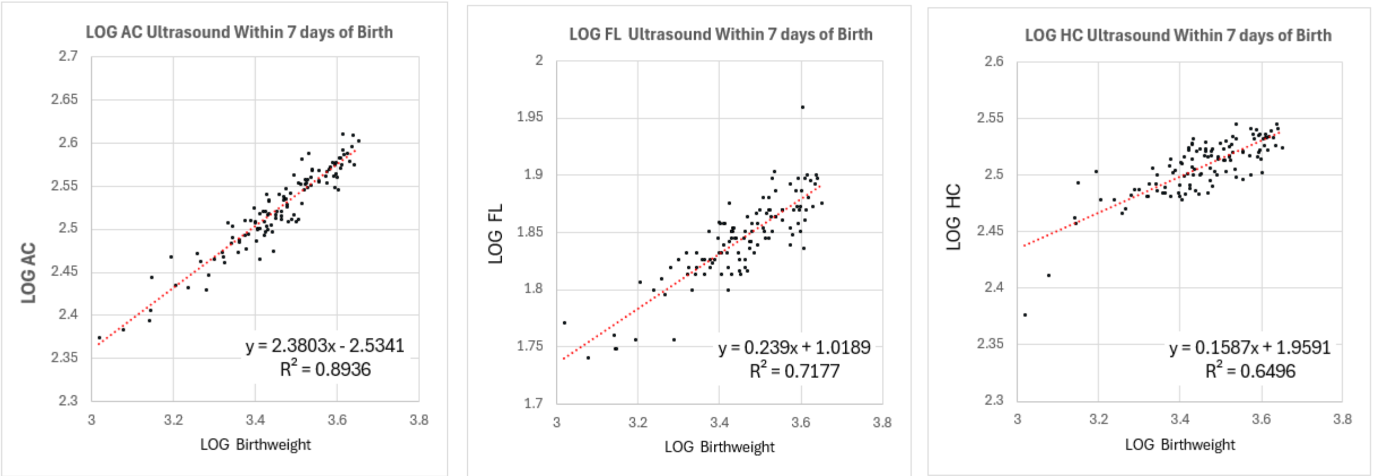


C:_1\BBCDG V7 DATA_______Baby Math Data 10 20 23_____New EFW Calcs Nov 4 23__medium to medium 2 23 2024\ [Deid POPs and LIFECODES Combined. Slope & Beta Adjustable. V20d 4 12 2024 Train & Test. 26 gr-day in Train & Test.xlsx]Test.7d 4th (5.2) (3)'!$DK$50

**Supplement-Figure-II.25**, Comparison of ***Binary Cellular Estimated Fetal Weight Equations***, for abdomen, head, and femur, in the ***Testing Dataset***, based on ultrasound carried out within 7 days of birth.

Testing ***Binary Cellular Estimated Fetal Weight Equations***: Sonographer Performance

The Problem isn't the ***Error*** but the ***Standard Deviation*** of the ***Error***, from ***Sonographer Performance***

Variability in the outcome value for a ***Fetal Ultrasound Measurements*** can be expected to come from a variety of sources, among which are: 1) Which equation is employed to interpret the ***Fetal Ultrasound Measurement***? 2) Which person is engaged in carrying out the ***Fetal Ultrasound Measurement***? An examination of results reported for 13 studies reveals an enormous amount of variation in both the ***Mean Error***, and the ***Standard Deviation*** of the ***Error*** (**Supplement-Figures-II.26 &27**), when examining the performance of a single equation, the ***Hadlock-4***, ***Estimated Fetal Weight Equation*** (#20)[230-242]. There is no evident correlation between ***Mean Error*** and ***Standard Deviation*** of the ***Error*** (**Supplement-Figure-II.26**). These studies have been carried out over four decades, but there was no indication of improvement in performance over time (**Supplement Figure-II.27**)

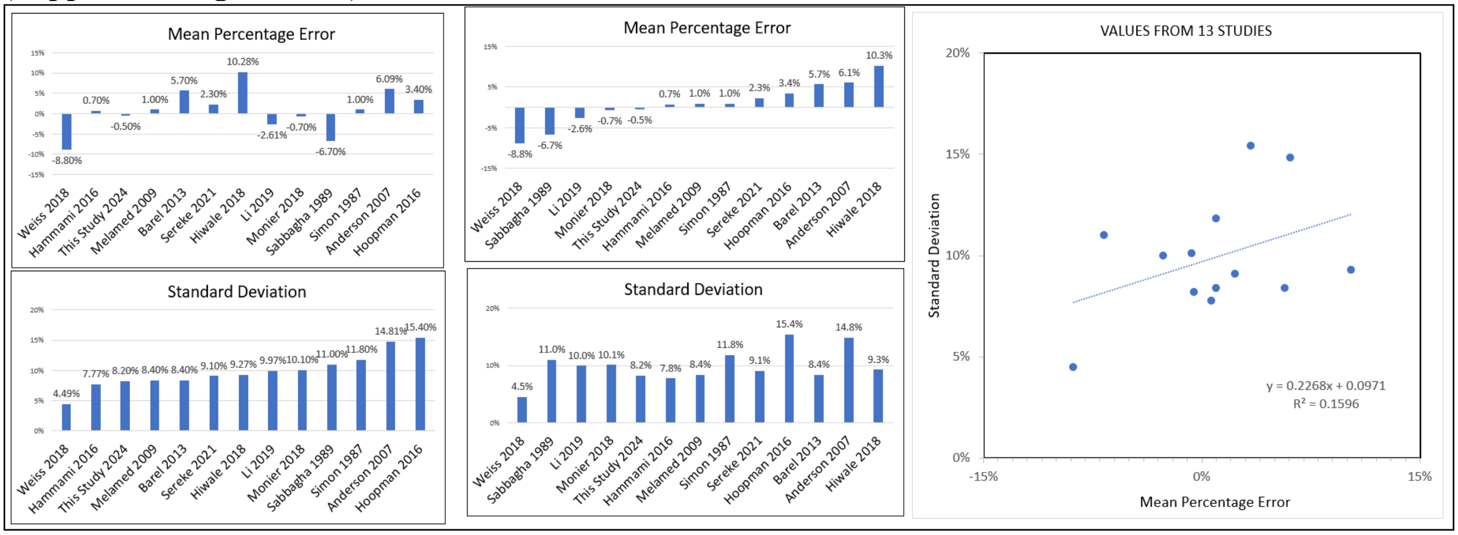


**Supplement-Figure-II.26**. Comparison of the performance of the ***Hadlock-4***, ***Estimated Fetal Weight Equation*** (#20) from 13 different studies.[230-242]

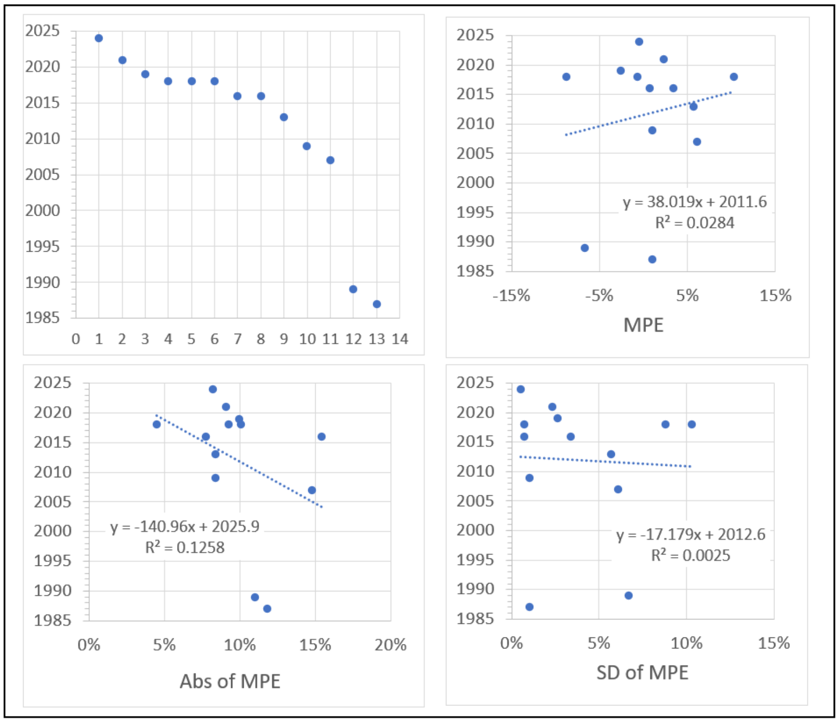


**Supplement-Figure-II.27** Comparison of the performance of the ***Hadlock-4***, ***Estimated Fetal Weight Equation*** (#20) from 13 different studies, by date of study.[230-242]

A number of studies have compared the performance of individuals in carrying out Fetal Ultrasound. One study examined the performance of 11 sonographers, reported in terms of ***Mean Error*** and the ***Standard Deviation*** of the ***Error***.[243] The variation from highest to lowest values was striking; the ***Mean Error*** of the ***Hadlock-4***, ***Estimated Fetal Weight Equation*** value varied from -0.9% to -6.3%, while the ***Standard Deviation*** of the ***Error*** varied from 8.4% to 12.0% (**Supplement-Figure-II.28**). There was no evident correlation among the results of the 11 sonographers between the ***Mean Error*** and the ***Standard Deviation*** of the ***Error***.

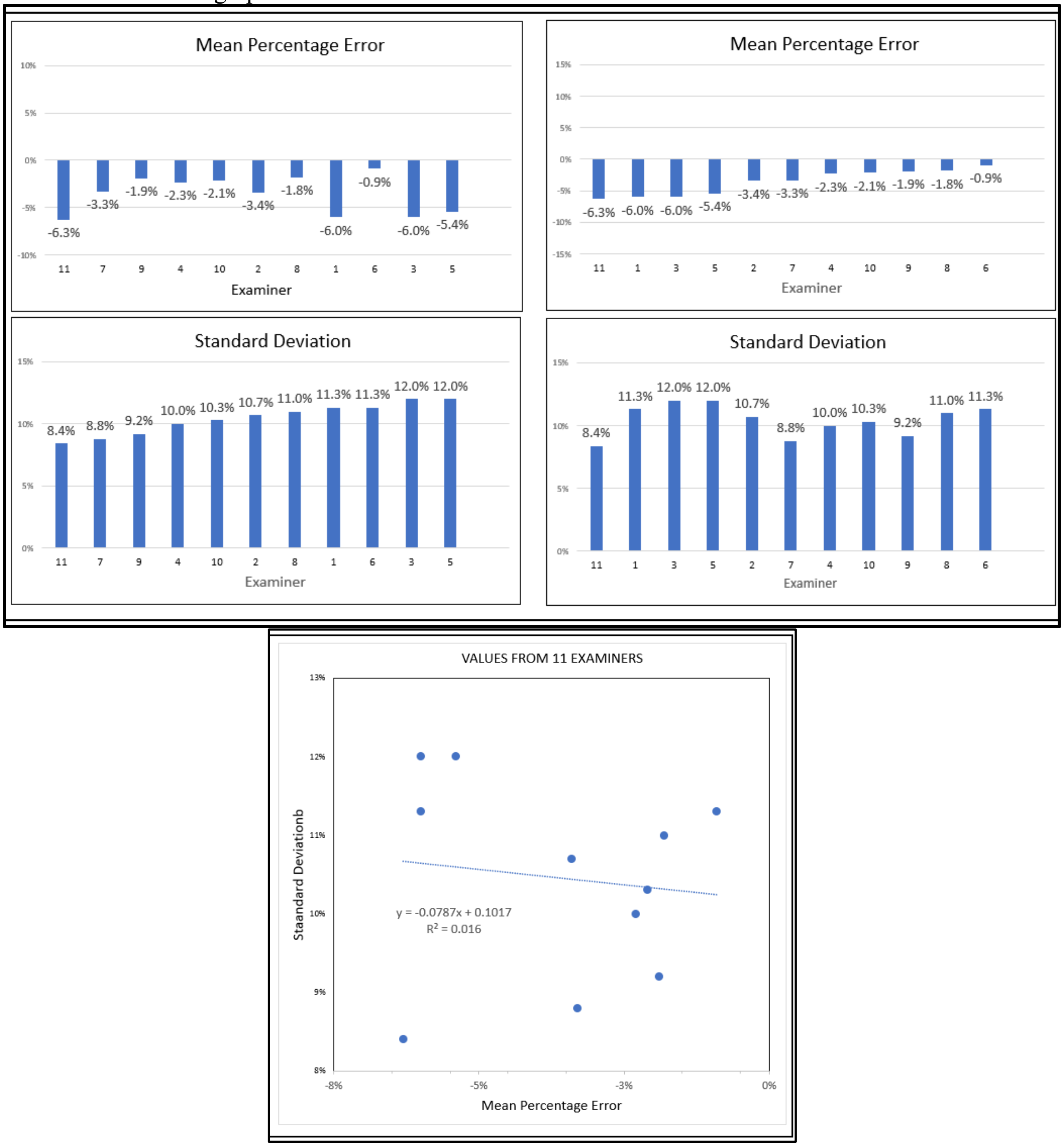


**Supplement-Figure-II.28**. Comparison of the performance of the ***Hadlock-4***, ***Estimated Fetal Weight Equation*** (#20) by 11 different examiners, as reported by Siemer et al[243]

Testing ***Binary Cellular Estimated Fetal Weight Equations***: Sonographer Performance (cont.)

Another study compared the performance of 259 ***Experienced Examiners*** with 1682 ***Inexperienced Examiners***, whose ***Ultrasound Measurements*** were applied to 11 ***Estimated Fetal Weight Equations***. The variation in performance betwewen the ***Experienced Examiners*** and ***Inexperienced Examiners*** was much greater than the performace of the various ***Estimated Fetal Weight Equations***. The correlation coeffficient ($r^2$) varied by 16.7% between the ***Experienced Examiners*** and ***Inexperienced Examiners***. The ***Estimated Fetal Weight Equations*** varied much less than the ***Examiners***, ranging highest to lowest varied by 3.5% among the data collected by the the ***Experienced Examiners***, and 9.3% among the deata collected by ***Inexperienced Examiners*** (**Supplement-Figure-II.29**).

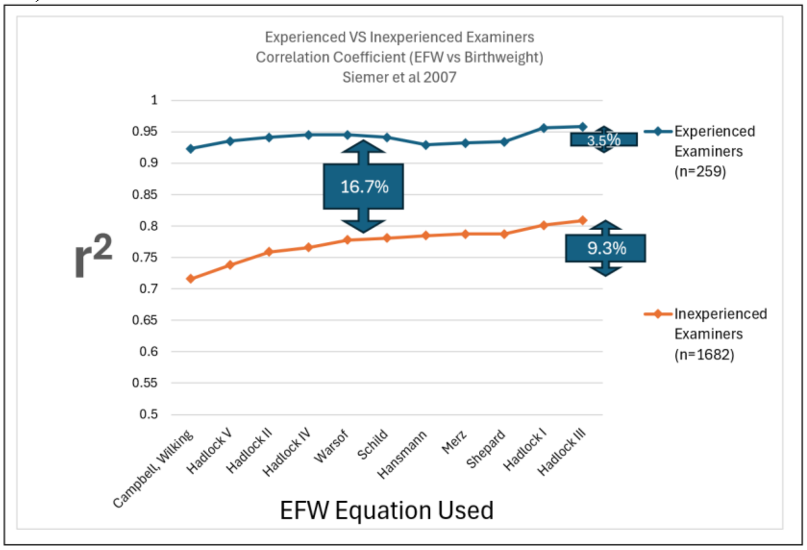


**Supplement-Figure-II.29** Comparison of the performance of a variety of ***Estimated Fetal Weight Equations*** by 259 ***Experienced Examiners*** versus 1682 ***Inexperienced Examiners***, as reported by Siemer et al 2007.[243]

Another example of variation from site to site in the performance of fetal ultrasound could be seen by the correlation coefficient ($r^2$) when comparing the results derived by the ***Hadlock-4 Estimated Fetal Weight Equation*** (#20). In the study results reported here, $r^2$=0.90, while in the study reported by Sharma et al[244] , $r^2$=0.3468 (**Supplement-Figure-II.30**).

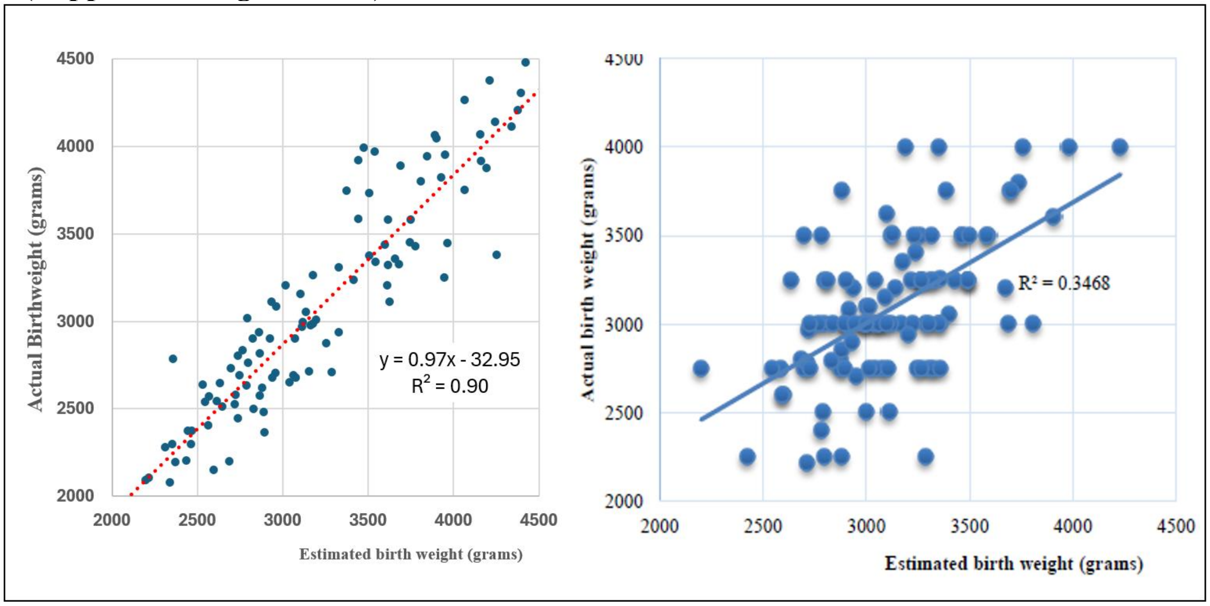


**Supplement-Figure-II.30**. Comparison of the performance of the ***Hadlock-4***, ***Estimated Fetal Weight Equation*** (#20) in this study, and in the study reported by Sharma et al.[244]

Testing ***Binary Cellular Estimated Fetal Weight Equations***: Sonographer Performance (cont.)

That performance can be traced to image quality was shown by Townsend et al,[245] who separated ***Ultrasound Measurements*** whose images they judge to be "Good" from those they judged to be "Poor". The ***Mean Error*** of the ***Hadlock-4***, ***Estimated Fetal Weight Equation*** (#20) value was 2.9% for studies with "Poor Images", and twice as high, at 6%, for studies with "Good Images". The ***Standard Deviation*** of the ***Error*** of the ***Hadlock-4***, ***Estimated Fetal Weight Equation*** (#20) value was 8.9% for studies with "Poor Images", and 15%, for studies with "Good Images" (**Supplement-Figure-II.31**).

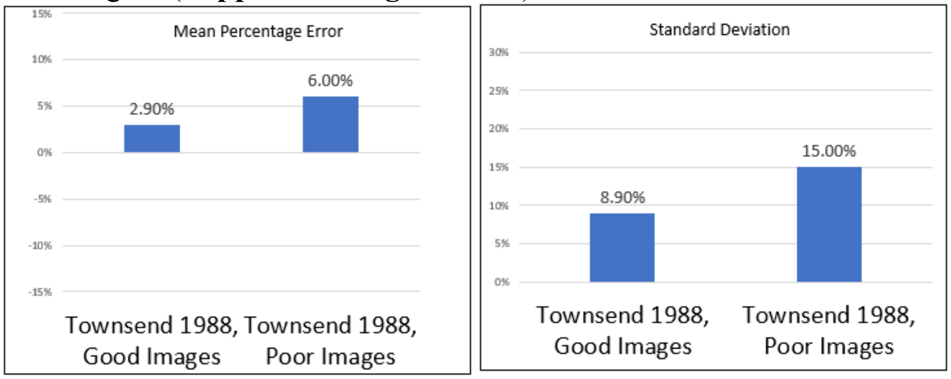


**Supplement-Figure-II.31** Comparison of the performance of the ***Hadlock-4***, ***Estimated Fetal Weight Equation*** (#20), sorted by the quality of the image, as reported by Townsend 1988.[245]

***Fetal Weight*** Over Time.

We can gain further insight into these three methods of estimating ***Fetal Weight*** from ***Ultrasound Measurement*** by examining how these values are distributed as pregnancy proceeds. Again, we shall examine the data from the 720 ***Ultrasound Measurements*** made late in pregnancy among 194 women seen at the Brigham and Women's Hospital, utilizing data only from ultrasound visits where all ***Ultrasound Measurements*** were recorded. With these values, we could compare estimations of ***Fetal Weight*** (red dots) made by the ***Abdominal Circumference Binary Cellular Estimated Fetal Weight Equation*** (#13), the ***Hadlock EFW Equation-4*** (#16), and ***INTERGROWTH EFW Equation*** (#15), examining how they lie in relation to the ***Binary Cellular Universal Growth Equation*** (#2), with its ***Empirical Ripples*** adjustments (blue line) (**Supplement-Figure-II.32**). As a crude first look, we can see that the ***Abdominal Circumference Binary Cellular Estimated Fetal Weight Equation*** (#13) and ***Hadlock EFW Equation-4*** (#16) yielded values roughly in accord with the ***Binary Cellular Universal Growth Equation*** (#2) out to the earliest measurement at 180 days before birth, while values generated by the ***INTERGROWTH EFW Equation*** (#15) departed strikingly for values measured earlier than 150 days before birth (**Supplement-Figure-II.32**).

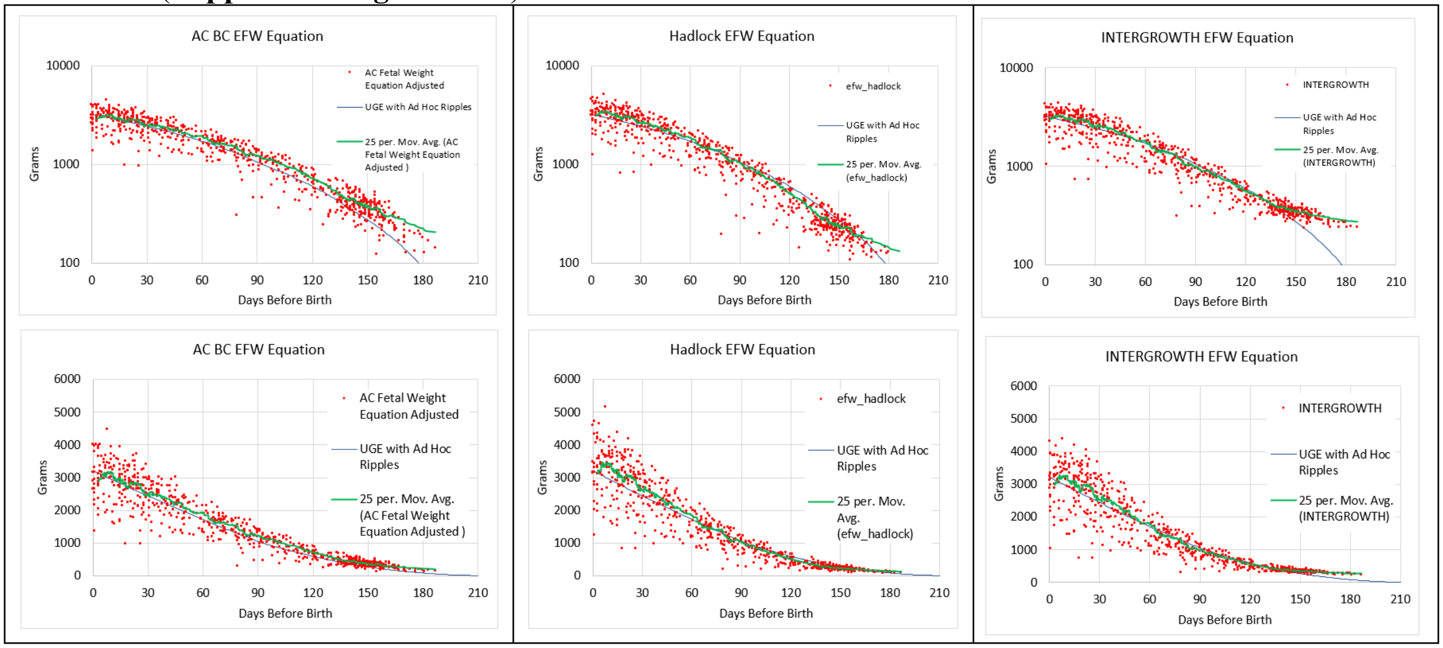


**Supplement-Figure-II.32**. Comparison of estimated ***Fetal Weight*** made by the ***Abdominal Circumference Binary Cellular Fetal Weight***) (#13), ***Hadlock EFW*** (#15) and ***INTERGROWTH EFW*** (#16) ***Equations***, for measurement up to 180 days before birth in comparison to values generated by the ***Binary Cellular Universal Growth Equation*** (#2), with ***Empirical Ripples*** adjustment.

C:_1\BBCDG V7 DATA_______Baby Math Data 10 20 23\4 21 23 Ivans Data AC FW Equation\[UGA & AC Equations 6 20 2023.xlsx]deid_POPS_with_Hadlock_4_21 (6)

The ***Binary Cellular Biology*** of Morphological Defect: ***Microcephaly Detection***.

Another use of ***Fetal Ultrasound*** is to determine whether the various parts of the body are in the correct relative size to each other, exemplified by the problem of microcephaly. As we have shown previously,[228] if each of two parts of the body, relate in an ***Allometric*** fashion to the weight of the body as a whole, they will relate in an ***Allometric*** fashion to the each other. This is borne out by the data shown in **Supplement-Figure-II.33** for the women in our dataset and provides more ***Binary Cellular Mathematics*** for yet another fetal analysis task: ***Microcephaly Detection***.

There is a small degree of curviness at the top and the bottom of the dataset. However, practical adjustment for this small departure can be accommodated by a polynomial approximation:

$$\textit{Microcephaly Ratio} = \mathrm{LOG(HC)}/[(-1.6233(\mathrm{LOG(AC)})^3+10.675(\mathrm{LOG(AC)})^2-22.451(\mathrm{LOG(AC)})+17.272] \quad (14)$$

We call this expression the ***Binary Cellular Microcephaly Detection Equation*** (#14).

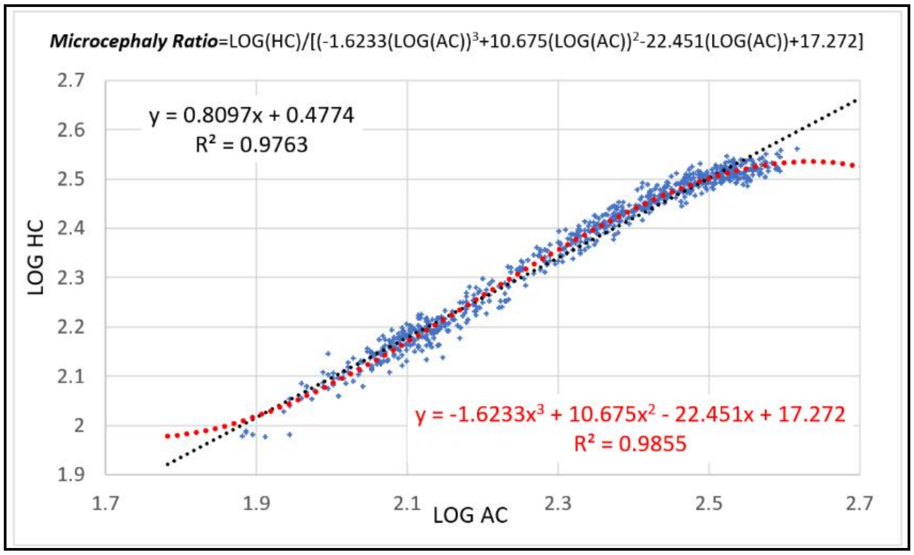


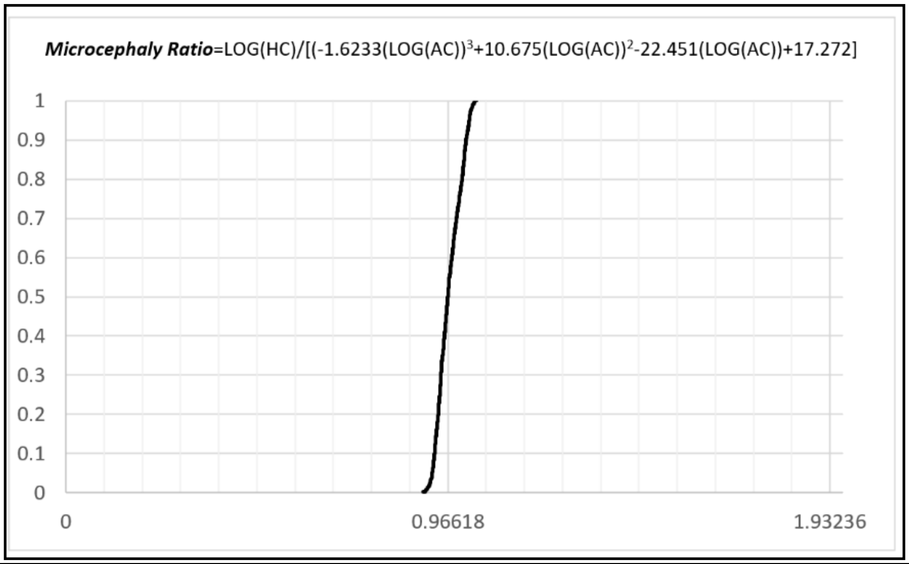


**Supplement-Figure-II.33**: The ***Binary Cellular Microcephaly Detection Equation*** (#14), with values from 720 ***Fetal Ultrasound*** measurements, among 194 women seen at the Brigham and Women's Hospital. TOP: log-log values of ***Head Circumference*** (HC) vs ***Abdominal Circumference*** (AC), with linear and polynomial regressions. BOTTOM: CDF of the equation's values from this dataset.

**DISCUSSION:**

***Binary Cellular Mathematics*** Provides ***Estimated Fetal Weight Equations*** That Are Better

Here we have seen that the ***Binary Cellular Estimated Fetal Weight Equation***, and specifically the ***Abdominal Circumference Binary Cellular Estimated Fetal Weight Equation***, performs better than six other ***Estimated Fetal Weight Equations***: the ***INTERGROWTH*** and ***Hadlock-1,-2,-3,-4***, and ***-5 Estimated Fetal Weight Equations***.[205-218] The ***Hadlock-4 Estimated Fetal Weight Equation*** is of special note, as it is among the most widely used, and has often been found to be the most accurate and precise of the many ***Estimated Fetal Weight Equations*** developed over the past five decades.[211-218]

The ***Abdominal Circumference Binary Cellular Estimated Fetal Weight Equation*** yielded a lower ***Mean Error*** than the six other ***Estimated Fetal Weight Equations***. Furthermore, unlike these six current ***Estimated Fetal Weight Equations***, the ***Abdominal Circumference Binary Cellular Estimated Fetal Weight Equation*** showed an equal performance among fetuses that become newborns of all sizes.

The ***Hadlock-4 Estimated Fetal Weight Equation*** has a marginally lower ***Standard Deviation*** of the ***Error*** than the ***Abdominal Circumference Binary Cellular Estimated Fetal Weight Equation***, but the difference was negligible. For 2-out-of 3 ultrasound measurements, the difference between the two equations is less than 28 grams, ~1% of the weight of an average newborn, while for 299-out-of-300 ultrasound measurements, the difference between to the equations is less than 137 grams, ~4% of the weight of an average newborn.

***Binary Cellular Mathematics*** Makes Possible Better ***Sonographer Performance*** of ***Fetal Weigh Estimation***

Here we have seen that the main source of error in estimating ***Fetal Weight*** lies not in the choice of the equation, but in the quality of the ***Ultrasound*** measurement carried out by the sonographer.[246,247,248] ***Mean Errors*** and ***Standard Deviation*** of the ***Error*** seen from different institutions, and from different sonographers, were greater than the ***Mean Errors*** and ***Standard Deviation*** of the ***Error*** from different equations.[230-245]

Why don't all sonographers perform as well as the best sonographers? One likely source of this shortcoming may well be the inability of ***Estimated Fetal Weight Equations*** to provide sonographers and their supervisors with feedback on performance, since they rely on multiple ***Fetal Ultrasound Measurements***. Even if the relationship between an ***Ultrasound Measurement*** and the subsequent ***Birthweight*** does not match up, the sonographer, and the sonographer's supervisor, are unaware of this, because multiple ***Ultrasound Measurement*** are combined into the estimate of ***Fetal Weight***. ***Binary Cellular Mathematics*** now makes such feedback possible, as each ***Binary Cellular Estimated Fetal Weight Equation*** relies on a single ***Fetal Ultrasound Measurement***, related to a single ***Birthweight***. Furthermore, it is impossible to "fudge" such a test, since the ***Birthweight*** is not known at the time of the ***Ultrasound Measurement***. Thus, performance feedback is now made possible with each ***Binary Cellular Estimated Fetal Weight Equation***, as each relates individual ***Ultrasound Measurement*** to individual ***Birthweights***.

<u>***Binary Cellular Biology*** Tells Us Why Some ***Ultrasound Measures*** Are Better Than Others</u>

Why does ***Abdominal Circumference*** provide a better estimate of ***Fetal Weight*** than ***Femur Length*** and ***Head Circumference***? As we outlined in the previous paper in this series, the underlying biology for why some parts of the body increase in size more rapidly than other parts of the body emerges from ***Binary Cellular Analysis*** of data on the size of many ***Body Parts*** (***p***), relative the size of the body as a whole (***w***), that is, the examination of the growth of the body and its parts in discrete units of numbers of cells ($N_p$ and $N_w$).

This ***Binary Cellular Analysis*** revealed that ***Body Part*** creation and growth is captured by a simple expression, the ***Binary Cellular Allometric Growth Equation*** (#11), which is governed by two parameters: the ***Binary Cellular Allometric Birth***, $B_{N1}$ and the ***Binary Cellular Allometric Slope***, $S_N$. The ***Binary Cellular Allometric Slope***, $S_N$, is determined by the average ***Cell Cycle Time***, $c_p$, of the cells in a ***Body Part***, which is specified in ***Cell-Heritable*** fashion, by a mechanisms such as DNA methylation, in the part's ***Founder Cell*** at its ***Binary Cellular Allometric Birth***, $B_{N1}$. Therefore, ***Binary Cellular Analysis*** shows us that it is ***Differential Cellular Proliferation***, that is, ***Cellular Selection***,[259,260] that make ***Ultrasound Measurements*** of some fetal ***Body Parts*** more effective objects for estimating ***Fetal Weight*** than other fetal ***Body Parts***.

Thus, ***Founder Cells*** that undergo ***Cell-Heritable*** changes in the shortening of their ***Cell Cycle Time***, $c_p$, by processes such as demethylation of their DNA, leading to more rapid growth of the resulting ***Body Part***, give rise to ***Body Parts*** with ***Ultrasound Measurement*** that do the best possible job in calculating ***Fetal Weight***. It is from this reasoning that we can understand why ***Abdominal Circumference*** has given better estimates of ***Fetal Weight*** than ***Biparietal Diameter***, ***Occipitofrontal Diameter***, ***Femur Length***, and ***Head Circumference***. It is also from this reasoning that we can revisit ***Body Part*** characteristics, biochemically, anatomically, and morphometrically, so as to identify which ***Ultrasound Measurements*** can give the most reliable measures of ***Fetal Weight***, as we shall examine below.

<u>***Binary Cellular Mathematics*** Can Combine ***Ultrasound Measurements*** of Multiple Anatomical Sites</u>

***Binary Cellular Analysis*** has revealed has ultrasound measurements of multiple anatomical sites can be combined by utilizing the ***Binary Cellular Estimated Fetal Weight Equation*** for each anatomical site, and then averaging the estimates into a single value for ***Fetal Weight***. Measurement of performance by the sonographer is thus retained. Furthermore, as we shall see in the next paper in this series,[219] ***Binary Cellular Analysis*** provides a basis by which multiple measurements made at different points in time can be combined.

<u>***Binary Cellular Biology*** Makes it Possible to Find Better Things to Measure.</u>

Note how our ***Cellular Allometric Analysis*** of human ***Body Parts*** shown in **Supplement-Figure-I.30** and **31** shows components of the abdomen (kidney and spleen) having steeper ***Binary Cellular Allometric Slopes***, $S_N$, than the brain. This is why the ***Abdomen*** gives a better result in predicting ***Birthweight*** than the ***Head.*** It follows that revisiting ***Cellular Allometric Analysis*** of ***Body Parts***, such as that shown in **Supplement-Figure-I.30** and **31** can give insight into which ***Ultrasound Measurements*** can be expected to give the most accurate and precise measurements of ***Fetal Weight***. This also means that we can go back to the basic anatomy of human fetuses, available to us from fetal MRI data,[249] 3D Sonography,[250,251] or fetal autopsy imaging, by methods such as high-resolution X-ray based Micro CT,[252,253] to search systematically for which fetal measurement should be expected to give us the most accurate and precise ***Estimated Fetal Weights***. ***Binary Cellular Mathematics*** thus gives us the power to re-invent ***Fetal Weight Estimation*** by returning to the 3D anatomy of the human fetus with tools not available to the pioneers of this field.

<u>Web-based Calculators Will Provide Clinicians with ***Binary Cellular*** Calculations That Aid Obstetric Care</u>

A first-generation Web-based calculator for calculating ***Fetal Weight*** from the ***Abdominal Circumference Binary Cellular Estimated Fetal Weight Equation***, is now available at "https://kidzgrowth.com". Encoding the additional ***Binary Cellular*** calculations described here is straightforward,[254] and underway. Also to be provided by these calculators will be tools for performance feedback, comparing each ***Ultrasound Measurement*** to the ultimate ***Birthweight,*** so that sonographers and their supervisors can review performance over the long term, as well as the performance of each individual case. These web-based calculators will make the results of these ***Binary Cellular*** mathematical manipulations widely available for aiding in obstetric care of individual pregnancies.

### *The Binary Cellular Biology of Human Growth*: III. Identifying, Comprehending, and Ameliorating Normal and Abnormal Fetal Size and Growth [255]


#### ABSTRACT

Assessing and understanding the ***Size*** and ***Growth*** and ***Age*** of each fetus provides essential information for managing the health of the newborn and its mother. As we have shown in the previous papers in this series, a consideration of the formation of the body, in units of numbers of cells, *N*, an approach we call ***Binary Cellular Analysis***, provides new equations for relating ***Fetal Ultrasound Measurement*** to ***Fetal Weight***, ***Binary Cellular Estimated Fetal Weight Equations***, and for capturing growth from conception onward, the ***Binary Cellular Universal Growth Equation***, which provides a basis for creating biologically-based, data-driven, ***Growth Curves***. These equations make it possible to compare ***Discrete Fetal Age/Fetal Weigh*** values determined by ultrasound with ***Continuous Fetal Age/Fetal Weigh*** values generated by the ***Binary Cellular Estimated Fetal Weight Equation***, to isolate and quantify the ***Speed of Growth*** and ***Pregnancy Length***. We call this new method ***Binary Cellular Fetal Growth Analysis***, which has found that ***Slow***, ***Average*** and ***Fast*** growing fetuses become ***Small***, ***Average***, and ***Large*** newborns by differing by the ***"a" Parameter*** of the ***Binary Cellular Universal Growth Equation***, the parameter that modeling suggests may reflect endocrinological control of growth, and that ***Short Pregnancies*** are associated with, and likely caused by, ***Slow Growth***. While ***Speed of Growth*** and ***Pregnancy Length*** are continuous metrics, ***Binary Cellular Fetal Growth Analysis*** captures at least 4 categories of growth: ***GROWTH TO AVERAGE SIZE***, normal growth along the ***Binary Cellular Universal Growth Equation***, arriving at a ***Healthy Square*** bounded by a healthy ***Birthweight*** at a healthy ***Fetal Age***; ***FAST GROWTH TO LARGE SIZE***, faster growth, to large size; ***SLOW GROWTH TO SMALL SIZE***, growth that is both too slow and too short; ***SLOW GROWTH TO AVERAGE SIZE***, growth that is slow but long enough. These findings reveal possibilities for the detection and pharmacological treatment of fetuses that grow ***Too Fast*** or ***Too Slowly***, as well as new ways to help obstetricians guide each fetus to a ***Birthweight*** and ***Fetal Age*** that has the lowest possible chance of complication. Web-based calculators (https://kidzgrowth.com) make these mathematical manipulations available for obstetric care.

#### INTRODUCTION

Assessing the ***Size*** and ***Growth*** of each fetus is a key task for understanding, and managing, obstetric care. In the previous two papers in this series, we presented a consideration of the formation of the body, in units of numbers of cells, ***N***, and the change in these numbers by the binary either/or decisions of cells to divide, or remain quiescent.[256-258] We call this approach ***Binary Cellular Analysis***, and it provides a new method for comprehending embryology, ***Binary Cellular Biology***, a link of ***Obstetrics*** and ***Biology*** to the common mitotic mechanism of ***Cellular Selection***, resulting from ***Differential Cellular Proliferation***, [259-261] and a variety of new equations for following the ***Creation***, ***Growth***, and ***Size*** and of the body and its parts.[262,] These equations include:

1. The ***Abdominal Circumference Binary Cellular Estimated Fetal Weight Equation***, for estimating the weight of a fetus from ***Abdominal Circumference Ultrasound Measurements***.
2. The ***CRL Fetal Age Equation***, for ascertaining the date of conception from ***Crown to Rump Length Ultrasound Measurements***.
3. The ***Binary Cellular Universal Growth Equation***, which captures the growth of the fetus from conception onward, and provides a method for creating biologically-based, data-driven, ***Growth Curves***.[276]

Here, we report the ***Binary Cellular Analysis*** of the ***Size*** of human fetuses, determined by the ***Abdominal Circumference Binary Cellular Estimated Fetal Weight Equation***, and their growth, captured by the ***Binary Cellular Universal Growth Equation***. We call this method ***Binary Cellular Fetal Growth Analysis***, and the images that capture this information ***Binary Cellular Fetal Growth Charts***.

The ***Binary Cellular Fetal Growth Analysis*** of a dataset of newborns described here reveals new insight into why some newborns are born ***Too Small***, and others, ***Too Large***, and how we can best foresee and ameliorate the arrival of these abnormal instances of growth.

#### METHODS

Data were analyzed in Microsoft Excel.

##### The ***Combined Dataset***

For this, and the previous manuscript in this series,[263] we utilized a deidentified dataset of 821 pregnant women seen at the Brigham and Women's Hospital. We call this dataset the "***Combined Dataset***". The ***Combined Dataset*** was comprised of two subsets: a ***194" dataset*** of 194 women, and a ***627" dataset***, which contains data on 627 pregnant women. The "***627" dataset*** had also been utilized in references 264, 265, 266, and 267. The two datasets were analyzed separately, as only the "***627" dataset*** contained full data on each newborn and its mother.

##### The "***627" Dataset***

Among the 627 patients in the "***627" dataset***, the ***Fetal Age*** at birth (the estimated time from conception to birth, calculated from the ***CRL Fetal Age Equation*** (#7a)) had a mean value of 277 days; a ***Standard Deviation*** of 15 days; a median of 278 days; the longest being 306 days; and the shortest being 175 days (**Supplement-Figure-III.1**). The birthweights had a mean value of 3178 grams; a ***Standard Deviation*** of 743 grams; a median of 3100 grams; the largest being 5180 grams; and the smallest being 700 grams (**Supplement-Figure-III.1**).

Newborns suitable for ***Binary Cellular Fetal Growth Analysis*** had at least one CRL measurement early in pregnancy, for ***Fetal Age*** calculation, with the ***CRL Fetal Age Equation*** (#7a), and at least two ultrasound measurements later in pregnancy, for ***Fetal Weight Estimation*** with the ***Abdominal Circumference Binary Cellular Estimated Fetal Weight Equation*** (#14)). Detailed analysis was carried out of 19 ***Average Size Babies*** (birthweight 3355-3459 grams), 53 ***Big Babies*** (birthweight 4000-4725 grams), and 26 ***Small Babies*** (birthweight 1125-2065 grams). Of these, full information for all necessary ultrasounds was available for 15 ***Average Size Babies*** (birthweight 3355-3459 grams) (***Average Size Baby Study Group***), 30 ***Big Babies*** (birthweight 4000-4725 grams) (***Big Baby Study Group***), and 23 ***Small Babies*** (birthweight 1125-2065 grams) (***Small Baby Study Group***) (**TABLE III.1**).

C:_1\BBCDG V7 DATA_______Baby Math Data 10 20 23______New Luke Lin Studies 1 2 2024\Slope & Size of Small Average & ***Big Babies***. jsm. V34d 3 28 2024.xlsx

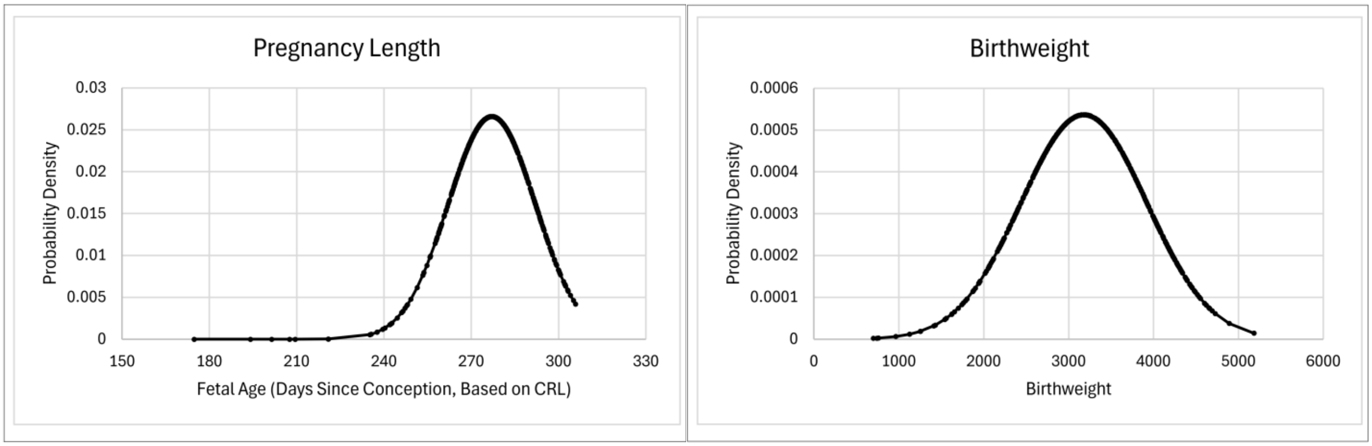


**Supplement-Figure-III.1**

Pregnancy Length, and Birthweights, in patients in the "***627" dataset***

##### The "***194" Dataset***

The ""***194" dataset***" of 194 women, who had comprised of 1022 ultrasound examinations, 302 of which measured ***Crown to Rump Length*** (***CRL***) early in development, and 720 of which measured other features later in development, most relevantly, ***Abdominal Circumference*** (***AC***).

Equations not derived or specified in the main text of this communication

As developed in the previous paper in this series, the ***Binary Cellular Universal Growth Equation*** (#2), for capturing the relationship between ***size***, ***N***, and ***age***, ***t***:

$$N_w = \int ln2/c * N_w * a\hat{}\bar{N}_w\hat{}b \quad (2c)$$

Within this expression lies the ***Binary Cellular Universal Mitotic Fraction Equation***:

$$m = a^{(N_w^b)} \quad (1)$$

The ***Abdominal Circumference Binary Cellular Estimated Fetal Weight Equation*** (#13) for estimating the weight of a fetus from ***Abdominal Circumference Ultrasound Measurements***, with the parameters used for calculation in this paper, is:

$$EFW = 0.0123(AC)\hat{}2.125 \quad (13c)$$

The reader will note a slight difference between this expression and the final ***Abdominal Circumference Binary Cellular Estimated Fetal Weight Equation*** arrived at in the previous paper in this series:

$$EFW = 0.001371156 * (AC\hat{}2.5187) - 244.25 \quad (14c)$$

This difference resulted from the fact that the calculations carried out in the previous paper, and specifically the working through of the various steps to arrive at Equation #14c, and the calculations reported in this paper, were carried out simultaneously, and when we have only reached the step for Equations #13 (**Supplement-Figure-II.5&6**). Fortunately, the difference between the two expressions is trivial from a practical standpoint. Future work, though, will no doubt use Equations #14c, with the updates of the parameters in that expression, as larger datasets become available.

The ***CRL Fetal Age Equation*** (#7a) for ascertaining of the date of conception from ***Crown to Rump Length Ultrasound Measurements***:

$$Fetal\ Age = (40.9041+(3.21585*(((CRL)\hat{}(0.5)))+(0.348956*(CRL)))) - 14 \quad (7a)$$

Derivations and testing can be found in the first[262] and second[263] papers in this series.

Terminology for ***Fetal Growth***: ***Age***, ***Gestational Age***, ***Conceptional Age, Fetal Age***

***Gestational Age***, the time since the last menstrual period, has long been the most common way for thinking about the age of the human fetus. Another commonly used term is ***Conceptional Age***, but the date of conception is not always known. As we shall be examining fetal, childhood and adult growth data, as well as growth data of animals such as mollusks, tunicates, and nematodes, which don't have a fetal period, we shall be using the simplest term, ***Age***, with modifiers to indicate the period examined, which, for our work here, will be ***Fetal Age***. Here we shall calculate ***Fetal Age*** from ***Crown to Rump Length*** (***CRL***) ***Ultrasound*** measurements with the ***CRL Fetal Age Equation*** (#7a).

Terminology for ***Fetal Growth***: ***Cells***, ***Grams***, ***Mitotic Fraction***, ***Weight-Gain Rate, Growth Velocity***

As we have noted in the first paper in this series,[262] while ***Binary Cellular Analysis,*** and specifically, the ***Binary Cellular Universal Growth Equation*** (#2), captures human growth over time, in units of number of cells, ***N*** (**Supplement-Figure-I.1**0, conversion to grams, ***g***, is also easily accomplished, as there are approximately $10^8$ cells in every gram of tissue.[273] Such a conversion to grams, ***g***, still retains an easily accessible, if perhaps temporarily hidden, biologically rooted mathematics for comprehending the biochemical mechanisms of growth, as we shall see in all three papers in this series**.** Further, when cell number values, ***N***, are derived from weight data, conversion back to grams, ***g***, cancels out any error in the estimation of cells per gram.

***Human Fetal Size*** and ***Growth*** can thus be examined in units of ***Cells***, ***Mitotic Fraction***, ***Grams***, or ***Weight-Gain Rate*** in units of ***Cells***/***Day*** or ***Grams***/***Day***, allowing comparison to other commonly used standards for assessing fetal developments, while retaining the analytical power of ***Binary Cellular Analysis***, as we shall see here.

The term ***Fetal Growth Velocity*** is sometime used, but we shall not be using it for several reasons. ***First***, although it has a long history in the study of growth, it is, in fact, a misnomer, as velocity technically is used to describe motion with a direction (a vector), such as a satellite, car, or bullet, while the growth of a fetus has no such directionality. ***Second***, in practice, there is no standardized definition of ***Fetal Growth Velocity***, and it has been used in various ways. ***Fetal Growth Velocity*** is sometime used to describe grams/time, based on ***Estimated Fetal Weight***,[268,269] and sometimes in millimeters/time, for ultrasound measurements.[270] Partarp Sovio and Smith capture ***Fetal Growth Velocity*** by measuring the change in the Z score of various ultrasound measurements between 28 and 37 weeks of gestational age.[271]

Of course, fetal growth is not linear, so ***Weight-Gain Rate***, in units of ***Cells***/***Day*** or ***Grams***/***Day***, even for the same fetus at various time points, will vary as to when the measurements are made. However, as Mongelli and colleagues[272] have noted, fetal growth is relatively constant at the end of pregnancy, a quality they call "quasi-linearity". As well shall see below, we also see this linearity, which we refer to as the ***~Linear Phase of Human Fetal Growth***. This linearity allows a first approximation of growth to be assessed in terms that we shall refer to as the ***Weight-Gain Rate***.

| Birthweigh Group | Number of Babies | Biggest Birthweight | Smallest Birthweight | Average Birthweight | Median Birthweight | Biggest Weight Grain (grams/day), by Linear Regression from 180 Days after Conception, from Ultrasound Measurement. | Smallest Weight Grain (grams/day), by Linear Regression from 180 Days after Conception, from Ultrasound Measurement. | Average Weight Grain (grams/day), from Ultrasound Measurement, by Linear Regression from 180 Days after Conception. | Median Weight Grain (grams/day), by Linear Regression from 180 Days after Conception, from Ultrasound Measurement. | Standard Deviation of Weight Grain (grams/day), by Linear Regression from 180 Days after Conception, from Ultrasound Measurement. | 0.95% | p | Biggest Weight Grain (grams/day), from the Last 2 Ultrasounds | Smallest Weight Grain (grams/day), from the Last 2 Ultrasounds | Average Weight Grain (grams/day), from the Last 2 Ultrasounds | Median Weight Grain (grams/day), from the Last 2 Ultrasounds | Standard Deviation of Weight Grain (grams/day), from the Last 2 Ultrasounds | 0.95% | p |
|---|---|---|---|---|---|---|---|---|---|---|---|---|---|---|---|---|---|---|---|
| ***Average Size Baby Study Group*** (15) | 15 | 3459 | 3355 | 3414 | 3420 | 31.5 | 20 | 23.81 | 23.20 | 2.89 | 5.66 | - | 39.1 | 6.5 | 23.29 | 22.5 | 9.56 | 18.73025 | - |
| ***Big Baby Study Group*** (30) | 30 | 4725 | 4210 | 4406 | 4385 | 46.4 | 12.1 | 30.97 | 30.80 | 6.74 | 13.21 | 0.000195 | 83.5 | 12.1 | 33.38 | 30.5 | 15.23 | 29.8476 | 2.85E-10 |
| Big Babies 4300-4725 grams (19) | 19 | 4725 | 4300 | 4499 | 4490 | 46.4 | 12.1 | 31.61 | 32.40 | 7.95 | 15.59 | 0.000689 | 60.4 | 12.1 | 33.71 | 32.4 | 13.35 | 26.16817 | 4.06E-08 |
| Big Babies 4500-4725 grams (8) | 8 | 4725 | 4515 | 4616 | 4625 | 46.4 | 12.1 | 32.45 | 32.95 | 9.26 | 18.16 | 0.002287 | 56.8 | 12.1 | 33.68 | 31.1 | 15.69 | 30.76135 | 0.001567 |
| Big Babies 4600-4725 grams (5) | 5 | 4725 | 4600 | 4665 | 4660 | 51.2 | 28.3 | 37.62 | 32.90 | 9.38 | 18.38 | 1.7E-05 | 51.2 | 28.3 | 37.62 | 32.9 | 9.38 | 18.38367 | 0.009708 |
| Big Babies 4660-4725 grams (3) | 3 | 4725 | 4660 | 4692 | 4690 | 57.4 | 32.9 | 38.07 | 38.20 | 5.95 | 11.66 | 1.06E-05 | 51.2 | 32.9 | 43.50 | 46.4 | 7.75 | 15.18462 | 0.055152 |
| ***Small Baby Study Group*** (23) | 23 | 2065 | 1125 | 1779 | 1870 | 34.4 | -28 | 14.08 | 15.40 | 10.37 | 20.32 | 0.000146 | 34.4 | -28 | 12.20 | 14.65 | 11.88 | 23.28665 | 0.002218 |

**Supplement -TABLE III.1.**

**Newborns from the "*627 Dataset*" Selected for Detailed *Binary Cellular Fetal Growth Analysis*.**

| Birthweight Range | Average Birthweight | Number | Average Pregnancy Length (Days) | Shortest Pregnancy (Days) | Longest Pregnancy (Days) | SD | -95% | +95% |
|---|---|---|---|---|---|---|---|---|
| 700-999 | 791 | 4 | 194 | 175 | 208 | 12 | 170 | 219 |
| 1000-1499 | 1117 | 8 | 224 | 175 | 266 | 29 | 167 | 282 |
| 1500-1999 | 1803 | 17 | 252 | 209 | 283 | 18 | 216 | 288 |
| 2000-2499 | 2299 | 68 | 269 | 242 | 300 | 11 | 249 | 290 |
| 2500-2999 | 2776 | 174 | 278 | 240 | 303 | 12 | 255 | 300 |
| 3000-3499 | 3239 | 124 | 281 | 253 | 303 | 10 | 261 | 301 |
| 3500-3999 | 3728 | 89 | 282 | 256 | 304 | 10 | 263 | 301 |
| 4000-4999 | 4201 | 98 | 283 | 261 | 306 | 10 | 263 | 304 |
| 4500-4599 | 4683 | 13 | 279 | 259 | 292 | 11 | 258 | 300 |

**Supplement -TABLE III.2**

Pregnancy Length, and Birthweights, by Birthweight Range, in Patients in the "***627*** ***dataset***"

C:_1\BBCDG V7 DATA_______Baby Math Data 10 20 23___Lifecodes 1 Line Per 10 20 2023\[LIFECODES_all_us_measure_wide_10_19_23 jsm REEXAMINED 4 28 2024.xlsx]LIFECODES_all_us_measure_wi (4)'!$AF$

#### RESULTS

***Binary Cellular Fetal Growth Analysis***

The growth of individual fetuses, and groups of fetuses, can be displayed with the ***Birthweight***, and ***Estimated Fetal Weight*** values calculated with the ***Abdominal Circumference Binary Cellular Estimated Fetal Weight Equation*** (#13), or the ***Crown to Rump Length Binary Cellular Fetal Weight*** (***CRL-BC EFW***) ***Equation*** (#14d), with each measure shown with the number of days since conception, as calculated by the ***CRL Fetal Age Equation*** (#7a). These ***Discrete Fetal Age/Fetal Weigh*** values determined by ultrasound can then be examined relative to the ***Continuous Fetal Age/Fetal Weigh*** values generated by the ***Binary Cellular Estimated Fetal Weight Equation***, to isolate and quantify the ***Speed of Growth*** and ***Pregnancy Length***. (**Supplement-Figure-III.2** and others below). We call this method ***Binary Cellular Fetal Growth Analysis***.

***Binary Cellular Fetal Growth Charts***

The graphs created by ***Binary Cellular Fetal Growth Analysis*** are called ***Binary Cellular Fetal Growth Charts***. These ***Binary Cellular Fetal Growth Charts*** can be displayed in grams, or units of numbers of cells, ***N***, since there are about $10^8$ cells in every gram of tissue[273] (**Supplement-Figure-III.2**).

***Binary Cellular Fetal Growth Charts*** in units of grams are frequently more practical, while displays in units of numbers of cells provide the starting point for deciphering the fundamental mechanism of growth, cellular proliferation, as we shall see below. Thus, even when we are examining growth in units of weight, behind the ***Binary Cellular Fetal Growth Charts*** shown in grams lie the biological agents of growth, cells, each marked by its essential ***Binary*** decision of whether, or not, to enter into mitosis.

The ***~Linear Phase of Human Fetal Growth***

As described in the first paper in this series,[262,261] the ***Binary Cellular Universal Growth Equation*** (#2) reveals that the growth of human fetuses flattens into an approximately constant growth over the last months of pregnancy. We call this feature the ***~Linear Phase of Human Fetal Growth***, as shown in the lower right ***Binary Cellular Fetal Growth Chart*** of **Supplement-Figure-III.2**.

The ***Healthy Square***, the ***Healthy Island***

We have placed on ***Binary Cellular Fetal Growth Charts*** lines that identify the conventional limits of a healthy birth: being born below 4,000 grams and above 2,500 grams, and more than 240 days after conception, the time that the ***Binary Cellular Universal Growth Equation*** (#2) of an ***Average Fetus*** could be expected to reach 2,500 grams. Later in this manuscript, we shall see how ultrasound data lets us refine that limit to 230 days. We shall also see that the rectangular boundaries of the ***Healthy Square*** can be refined by replacing them with the meandering concentric topological boundaries of a ***Healthy Island***, based on newborn morbidity and mortality data.

***<u>Binary Cellular Fetal Growth Analysis</u>*** <u>of</u> ***<u>Average Size Babies</u>***<u>, examined individually.</u>

***Binary Cellular Fetal Growth Charts*** of the 15 ***Average Size Babies*** ("***Average Size Baby Study Group***") were analyzed in depth (TABLE III.1). One typical example can be seen in **Supplement-Figure-III.2**, where growth occurs right along the ***Binary Cellular Universal Growth Equation*** (#2),[263,263] with a length of pregnancy that lands the fetus into the ***Healthy Square*** of ***Average Birthweight***.

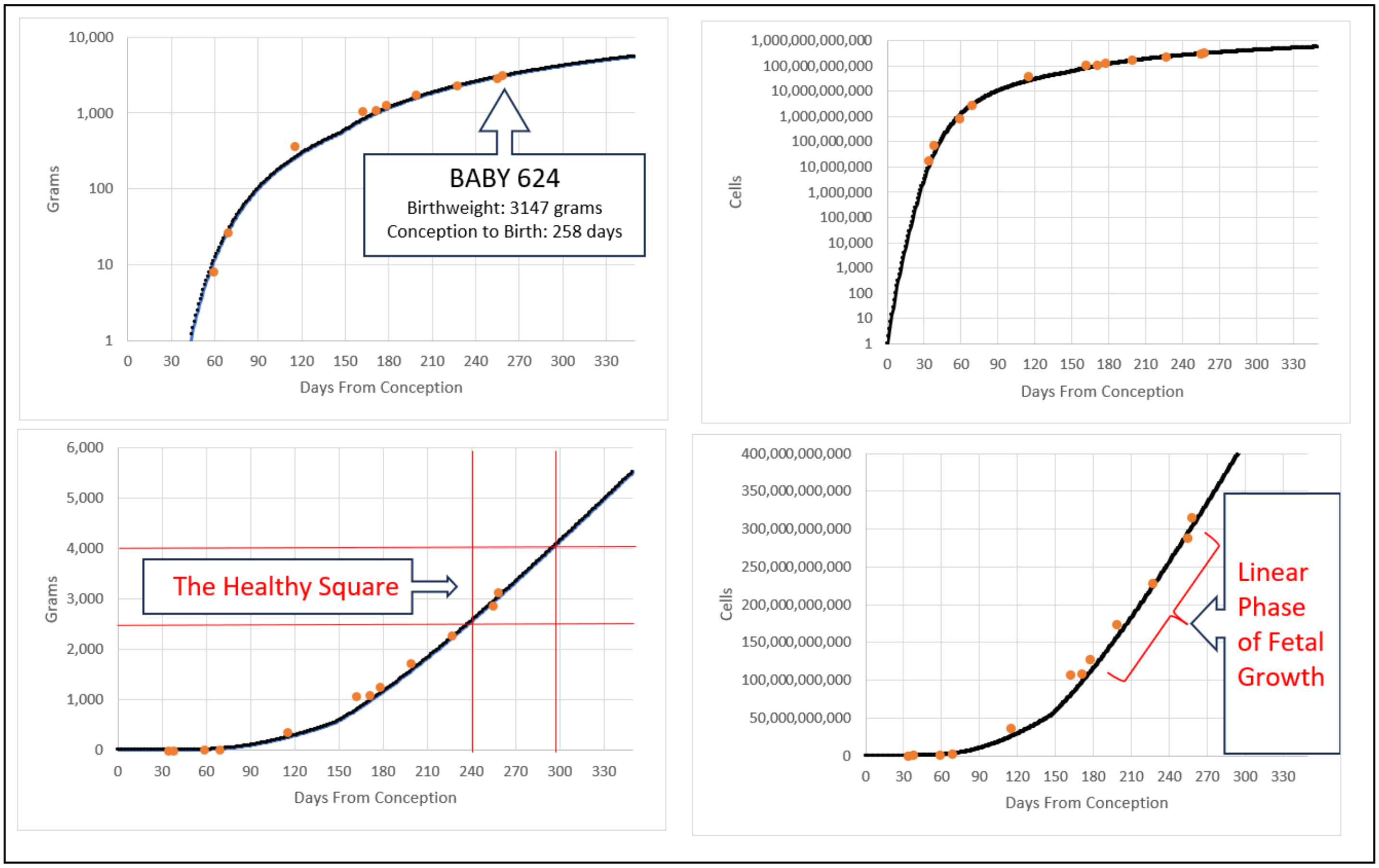


**<u>Supplement-Figure-III.2</u>** ***Binary Cellular Fetal Growth Charts*** for an ***Average Size Baby*** (newborn born between 2,500 and 4,000-grams). The last datapoint identifies birthweight; the other values are from ultrasound by the ***Abdominal Circumference Binary Cellular Estimated Fetal Weight Equation*** (#13), or the ***Crown to Rump Length Binary Cellular Fetal Weight*** (***CRL-BC EFW***) ***Equation*** (#14d), rooted in the time of conception, as determined by the ***CRL Fetal Age Equation*** (#7a). The dark line is the ***Binary Cellular Universal Growth Equation*** (#2).

***Binary Cellular Fetal Growth Analysis*** of ***Big Babies***, Examined Individually.

***Binary Cellular Fetal Growth Charts*** of the 30 ***Big Babies*** ("***Big Baby Study Group***") were analyzed in depth (TABLE III.1). One typical example can be seen in **Supplement-Figure-III.3**, where growth occurs above the ***Binary Cellular Universal Growth Equation*** (#2). The length of the pregnancy of the ***Big Baby*** shown in **Supplement-Figure-III.3**, and the other 29, are quite similar to that of ***Average Size Babies*** (**TABLE III.2**), but the faster growth of fetuses that become the ***Big Babies*** land them above the ***Healthy Square*** at a ***Large Birthweight*** (***Macrosomia***).

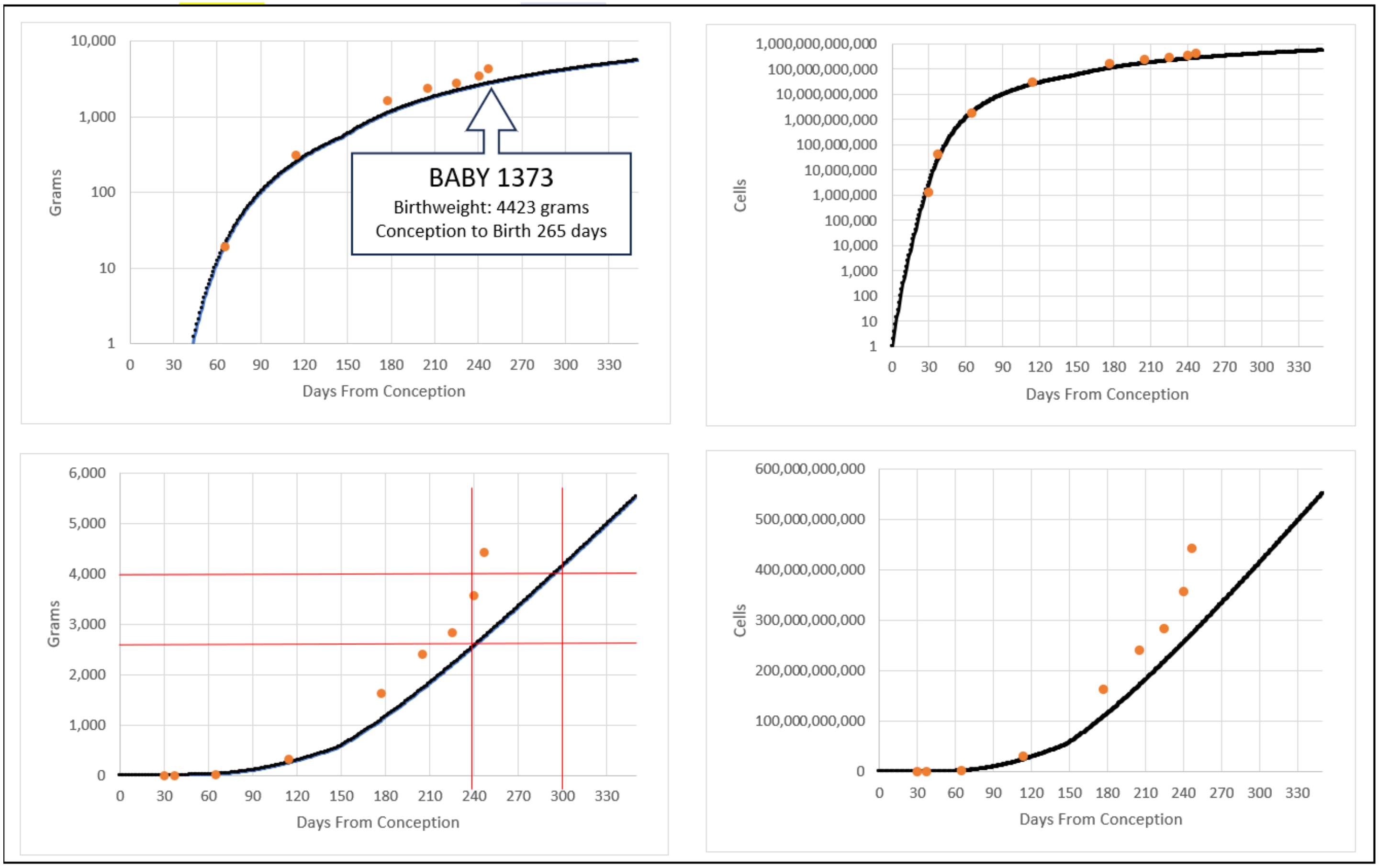


**Supplement-Figure-III.3** ***Binary Cellular Fetal Growth Charts*** of a ***Big Baby*** (macrosomia: newborn born above 4,000-grams). The last datapoint identifies birthweight; the other values are from ultrasound by the ***Abdominal Circumference Binary Cellular Estimated Fetal Weight Equation*** (#13), or the ***Crown to Rump Length Binary Cellular Fetal Weight*** (***CRL-BC EFW***) ***Equation*** (#14d), rooted in the time of conception, as determined by the ***CRL Fetal Age Equation*** (#7a). The dark line is the ***Binary Cellular Universal Growth Equation*** (#2).

**_Binary Cellular Fetal Growth Analysis_ of _Small Babies_, Examined Individually.**

***Binary Cellular Fetal Growth Charts*** of the 23 ***Small Babies*** ("***Small Baby Study Group***") were analyzed in depth (**TABLE III.1**). One typical example can be seen in **Supplement-Figure-III.4**. The length of the pregnancy of the ***Small Baby*** shown in **Supplement-Figure-III.4**, is shorter than that of the ***Average Size*** and ***Big Babies*** (**TABLE III.2**), and its rate of growth was slower, with growth occurring below the ***Binary Cellular Universal Growth Equation*** (#2). As was generally the case for all 23 ***Small Babies*** in the ***Small Baby Study Group***, the shortness of their pregnancies was the principal cause of their low ***Birthweights***, which when combined with their slow growth rates, land them below the ***Healthy Square*** of normal growth.

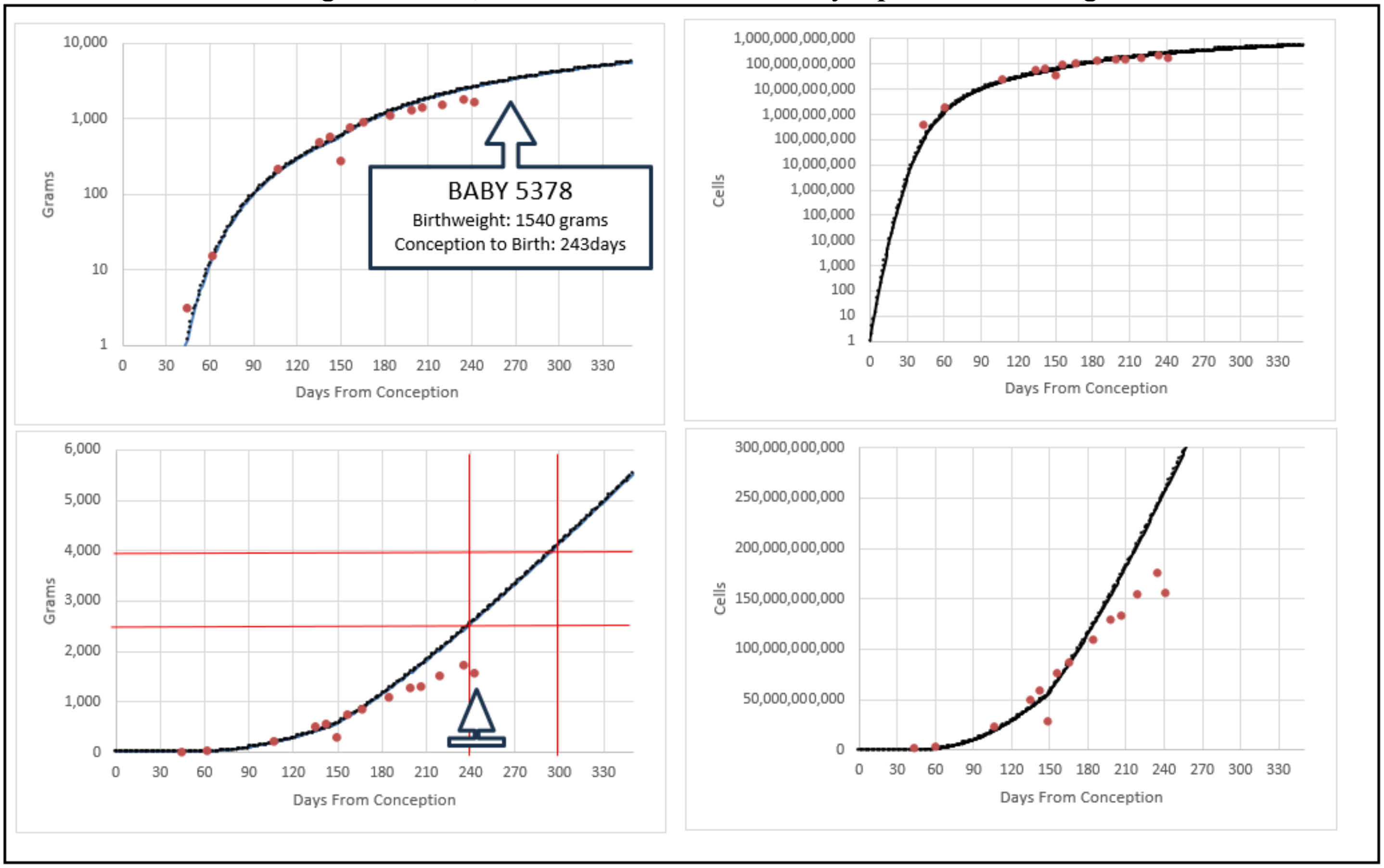


**Supplement-Figure-III.4.** ***Binary Cellular Fetal Growth Charts*** of a ***Small Baby*** (newborn born below 2.500-grams). The small arrow identifies birthweight. The last datapoint identifies birthweight; the other values are from ultrasound by the ***Abdominal Circumference Binary Cellular Estimated Fetal Weight Equation*** (#13), or the ***Crown to Rump Length Binary Cellular Fetal Weight*** (***CRL-BC EFW***) ***Equation*** (#14d), rooted in the time of conception, as determined by the ***CRL Fetal Age Equation*** (#7a). The dark line is the ***Binary Cellular Universal Growth Equation*** (#2).

***Binary Cellular Fetal Growth Analysis*** of ***Small, Average***, and ***Big Babies***, Examined in Small Groups

Let us now move on from examining the growth of fetuses one-at-a time, to examining the growth of each of the groups fetuses, together (**TABLE III.1**): the ***Small Baby Study Group*** made up of 23 ***Small Babies*** (birthweight 3355-3459 grams); the ***Average Size Baby Study Group*** made up of 15 ***Average Size Babies*** (birthweight 3355-3459 grams); and, the ***Big Baby Study Group*** made up of 30 ***Big Babies*** (birthweight 3355-3459 grams).

Using the ***~Linear Phase of Human Fetal Growth*** to Capture Fetal Growth by ***Weight-Gain Rate*** (***Grams/Day***)

As described below and above, the measurement of the growth of fetuses can be made by using ultrasound measurements from multiple dates, by several methods. The most straightforward method takes advantage ***~Linear Phase of Human Fetal Growth,*** which is especially pronounced for ultrasound measurement made at ***Fetal Ages*** from ~210 days (≈7 months) from conception to birth, but which can also be accomplished, with a slight degradation in quality, with ultrasound data starting from ***Fetal Ages*** as young as ~180 days (≈6 months) from conception. The ***~Linear Phase of Human Fetal Growth*** allows such simple measurements of fetal growth to be made by regression with 2 or more ultrasound measurements, yielding measurements of growth in units of ***Weight-Gain Rate*** (***Grams/Day***) (**Supplement-Figure-III.5**).

Using the ***Binary Cellular Universal Growth Equation*** to Capture Fetal Growth by ***"a" Parameter***

More complex, and rigorous, although arithmetically more demanding, measurements of fetal growth can also be made by fitting multiple ultrasound measurements to the ***Binary Cellular Universal Growth Equation*** (#2). As we shall see below, such calculations yield measurements of growth in units of the ***"a" Parameter*** the ***Binary Cellular Universal Growth Equation*** (#2). These ***"a" Parameter*** estimates can rely on ultrasound measurements much earlier than the 180-day ***Fetal Age*** limit that restricts ***Weight-Gain Rate*** (***Grams/Day***) measurements noted in the previous section.

Fetuses that Become ***Big Babies*** Grow Rapidly; Fetuses that Become ***Small Babies*** Grow Slowly

The fetal growth measurements of the 15 ***Average Size Babies*** ("***Average Size Baby Study Group***"), birthweight 3355-3459 grams), the 30 ***Big Babies*** ("***Big Baby Study Group***", birthweight 4000-4725 grams), and the 23 ***Small Babies*** ("***Small Baby Study Group***", birthweight 1125-2065 grams) are summarized in **TABLE III.1** and **Supplement-Figures III.5&6.** Shown are ***Weight-Gain Rate*** values (***Grams/Day***), measured by linear regression with the ***Abdominal Circumference*** measurements made from 180 days after conception, and ***Weight-Gain Rate*** values (***Grams/Day***), measured from just the last two ***Abdominal Circumference*** measurements carried out before birth. Calculations of ***Weight-Gain Rate*** (***Grams/Day***), derived from the Estimated Fetal Weight calculated by the ***Abdominal Circumference Binary Cellular Estimated Fetal Weight Equation*** (#2) show us that fetuses that become ***Big Babies*** grow rapidly while fetuses that become ***Small Babies*** grow slowly.

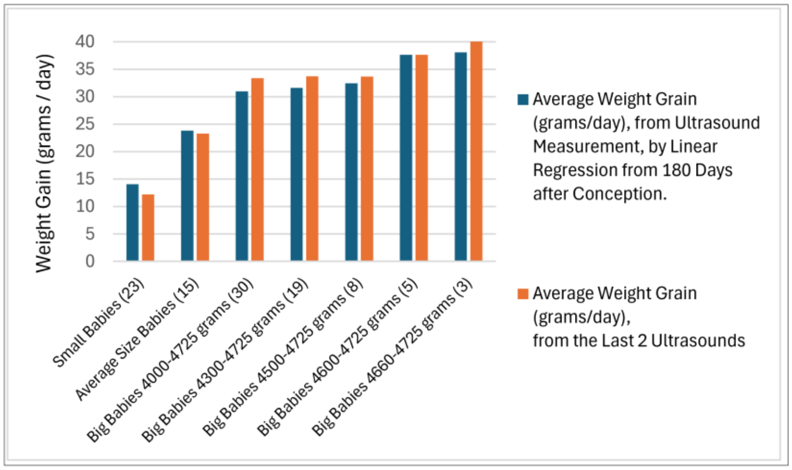


**Supplement-Figure-III.5** ***Weight-Gain Rate Rate*** (***grams/day***) for fetuses that become babies in the "***Average Size Baby Study Group***", "***Small Baby Study Group***", "***Big Baby Study Group***", and various subgroups of the "***Big Baby Study Group***"

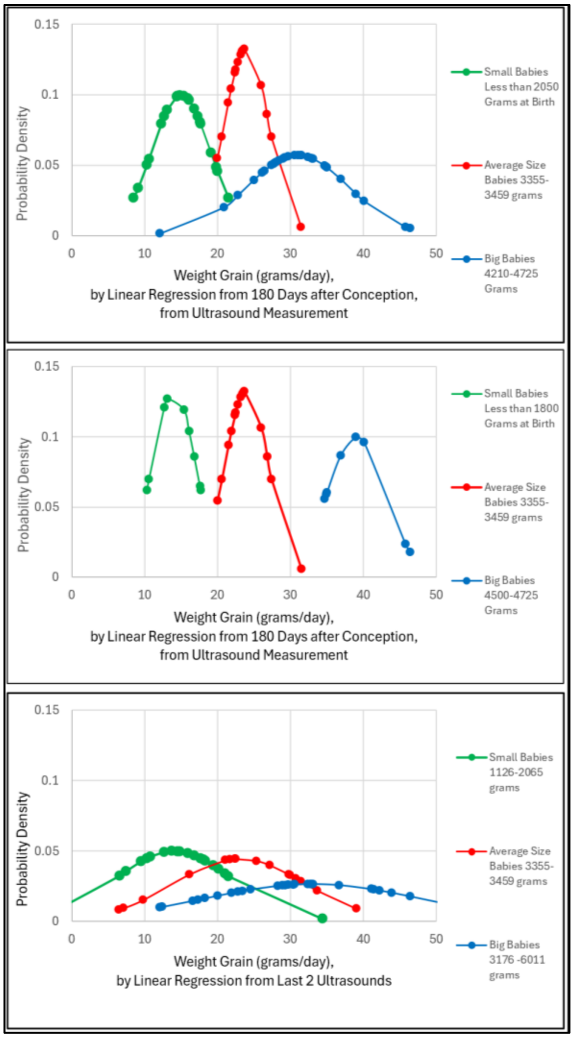


**Supplement-Figure-III.6**. Distribution of **Weight-Gain Rate Rate** (**grams/day**) for fetuses that become babies in the "***Average Size Baby Study Group***", "***Small Baby Study Group***", "***Big Baby Study Group***", and various subgroups of the "***Big Baby Study Group***"

***Binary Cellular Fetal Growth Analysis*** of Fetal Ultrasound Can Aid Prediction of ***Big*** and ***Small Birthweight.***

Fetal ***Weight-Gain Rate*** (***Grams/Day***), measured by linear regression with the ***Abdominal Circumference*** measurements made from 180 days after conception, was greatest for the ***Big Babies*** ("***Big Baby Study Group***"), less so for the ***Average Size Babies*** ("***Average Size Baby Study Group***"), and lowest for the ***Small Babies*** ("***Small Baby Study Group***") (**TABLE III.1**). This was seen for the 15 ***Average Size Babies*** of birthweight 3355-3459 grams, the 30 ***Big Babies*** of birthweight 4000-4725 grams, and the 23 ***Small Babies*** of birthweight 1125-2065 grams), where the three ***Study Groups*** had a small number of overlaps (**Supplement-Figure-III.6**). When these calculations are narrowed a bit, to ***Big Babies*** of birthweight 4500-4725 grams, and ***Small Babies*** of birthweight 1125-1800 grams, the three groups show no overlap whatsoever (**Supplement-Figure-III.6, middle**).

When ***Weight-Gain Rate*** (***Grams/Day***), is measured from just the last two ***Abdominal Circumference*** measurements carried out before birth, the three ***Study Groups*** also were significantly distinguishable (**TABLE III.1**). However, with only these two measurements, there was considerable overlap between the three groups (**Supplement-Figure-III.6, bottom**).

These observations suggest that the detection of fetuses that will become ***Big***, ***Small***, and ***Average Size Babies*** is possible. However, doing so with the accuracy and precision required for taking medical action will require more than two measurements, at optimal intervals. Approaches for accomplishing this goal are outlined in the DISCUSSION.

__***Binary Cellular Fetal Growth Analysis*** of ***Average Size Babies***, Examined as a Group, in Grams.__

Images of the change in size of fetuses over time were created by graphing ***Binary Cellular Fetal Growth Charts*** capturing the ***Fetal Weight*** of each ultrasound measurement, calculated by the ***Abdominal Circumference Binary Cellular Estimated Fetal Weight Equation*** (#13), against the ***Fetal Age*** when each ultrasound measurement was made, calculated by the ***CRL Fetal Age Equation*** (#7a). In **Supplement-Figure-III.7**, this is shown for the 15 ***Average Size Babies*** ("***Average Size Baby Study Group***", **TABLE III.1**), together with data from the first week of life, calculated from cell counts made of IVF embryos.[300] Note the "cloud" of the individual datapoints, and the "curve" of the ***Moving Average*** that smooths these values.

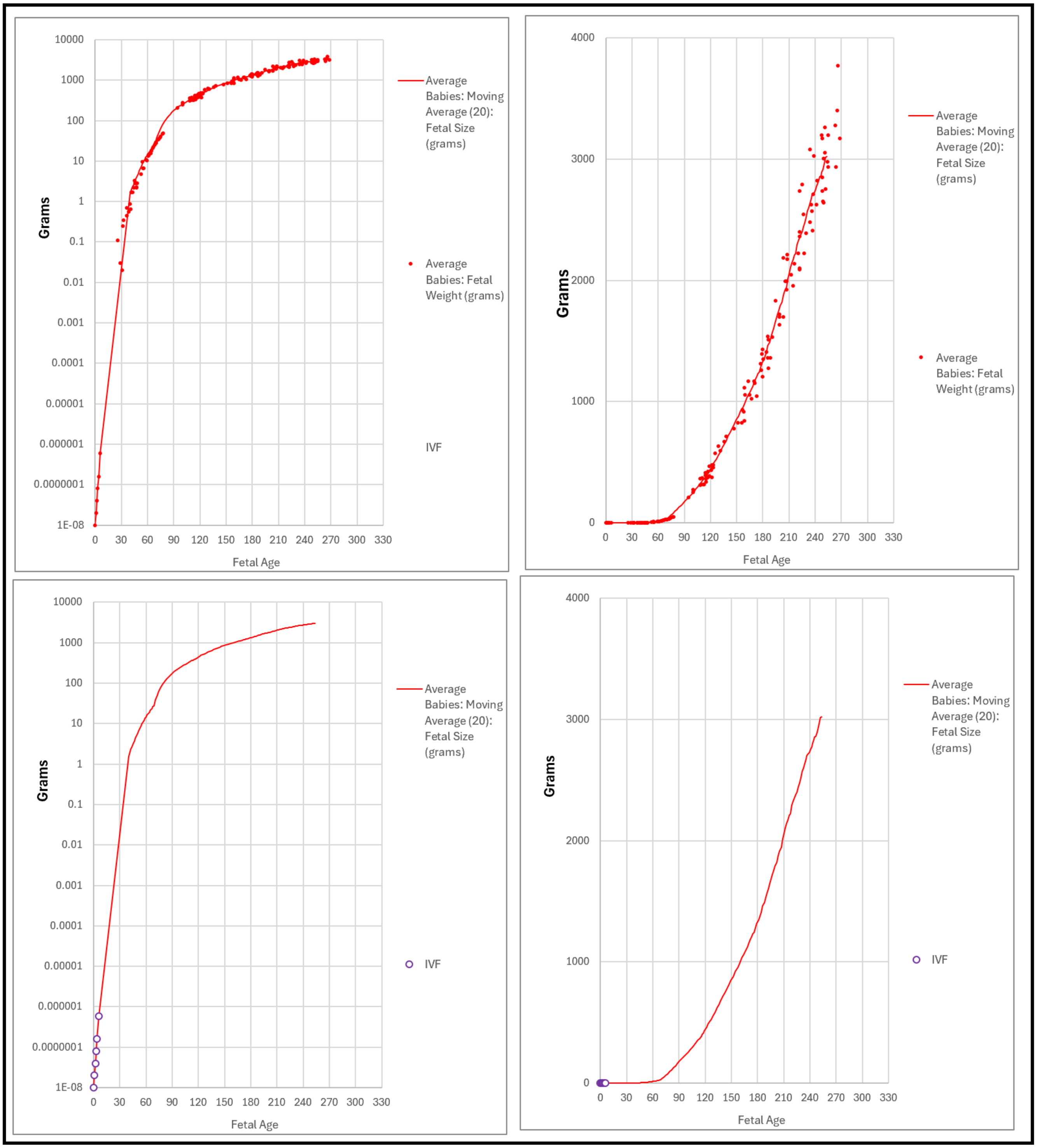

**_Binary Cellular Fetal Growth Analysis_ of _Average Size Babies_, Examined as a Group, in Cells.**

Again, it is straightforward to convert these weight-based data into units of cells, ***N***, since there are about $10^8$ cells in every gram of tissue (**Supplement-Figure-III.8**).[273] By this simple step, one can get the first view of the underlying principal mechanism of growth, mitosis.

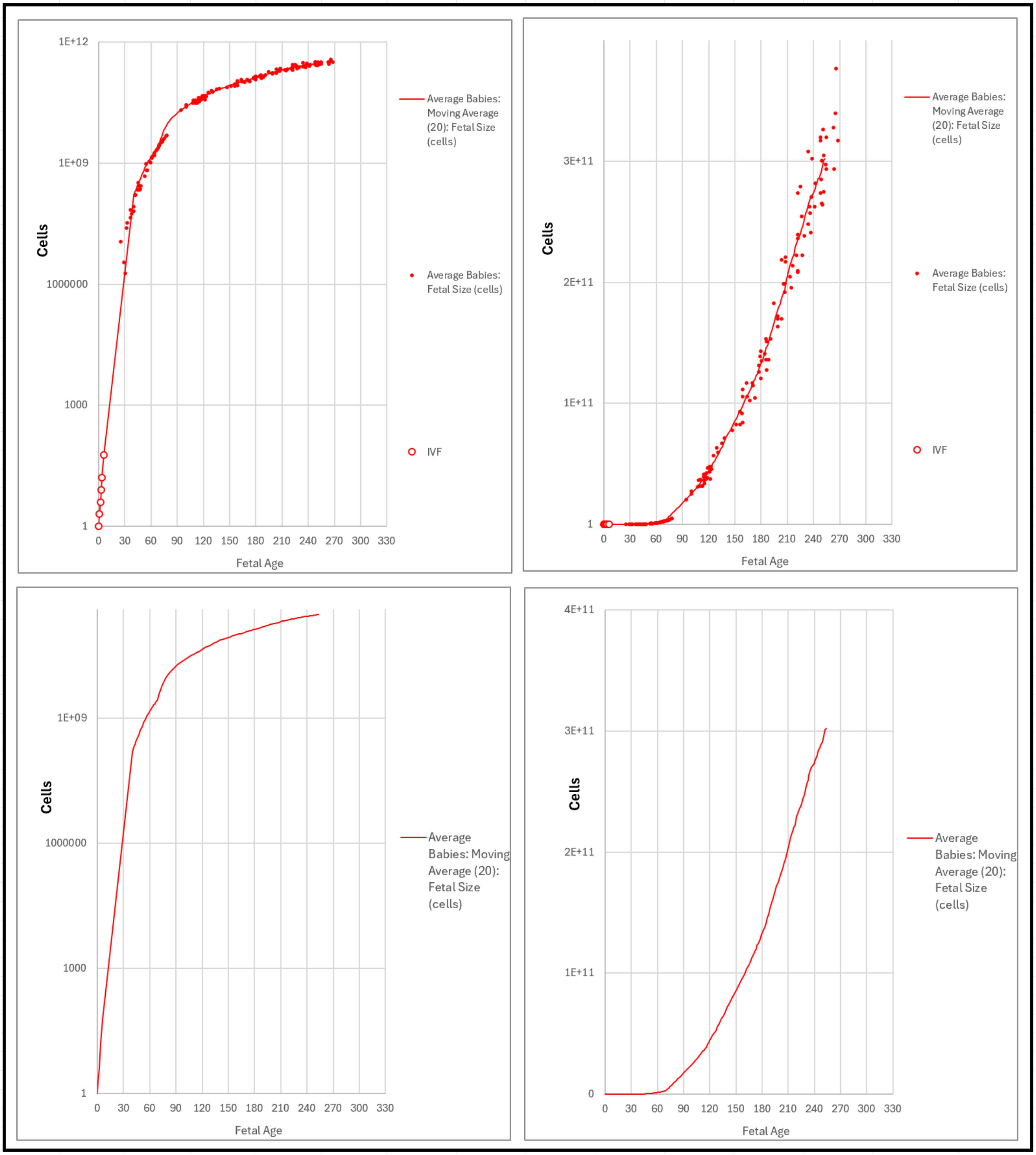


**Supplement-Figure-III.8** ***Binary Cellular Fetal Growth Charts*** capturing the ***Fetal Size*** (cells) of each ultrasound measurement, calculated by the ***Abdominal Circumference Binary Cellular Estimated Fetal Weight Equation*** (#13), against the ***Fetal Age*** when each ultrasound measurement was made, calculated by the ***CRL Fetal Age Equation*** (#7a), for the 15 babies in the "***Average Size Baby Study Group***".

Fit of the ***Binary Cellular Universal Growth Equation*** to the Fetuses that Become ***Average Size Babies***

In the first paper in this series, we executed a fit of fetal size data to the ***Binary Cellular Universal Growth Equation*** (#2), by determining the optimal values for the equations three parameters: ***c***, the ***Cell Cycle Time***, which measures how long it takes cells to divide; and ***a*** and ***b***, which determine how many cells are engaged in mitosis, the ***Mitotic Fraction***, as captured by the ***Binary Cellular Universal Mitotic Fraction Equation*** (#1); and ***R***, the ***Ripples in Growth***, which adjusts the fine scale details of growth over the full expanse of life.[263] In that first paper, we used fetal autopsy data to measure these ***Ripples***. With the ultrasound data described here, we refined the values of these ***Ripples*** for the data from the "***Average Size Baby Study Group***" (**Supplement-Figure-III.9**). While the results from these two sources are quite similar, the new ultrasound-based ***Binary Cellular Fetal Growth Charts*** are likely to provide more accurate representations of fetal growth, and, most importantly, provide more internally consistent assessments of growth based on ultrasound (**Supplement-Figure-III.10-12**).

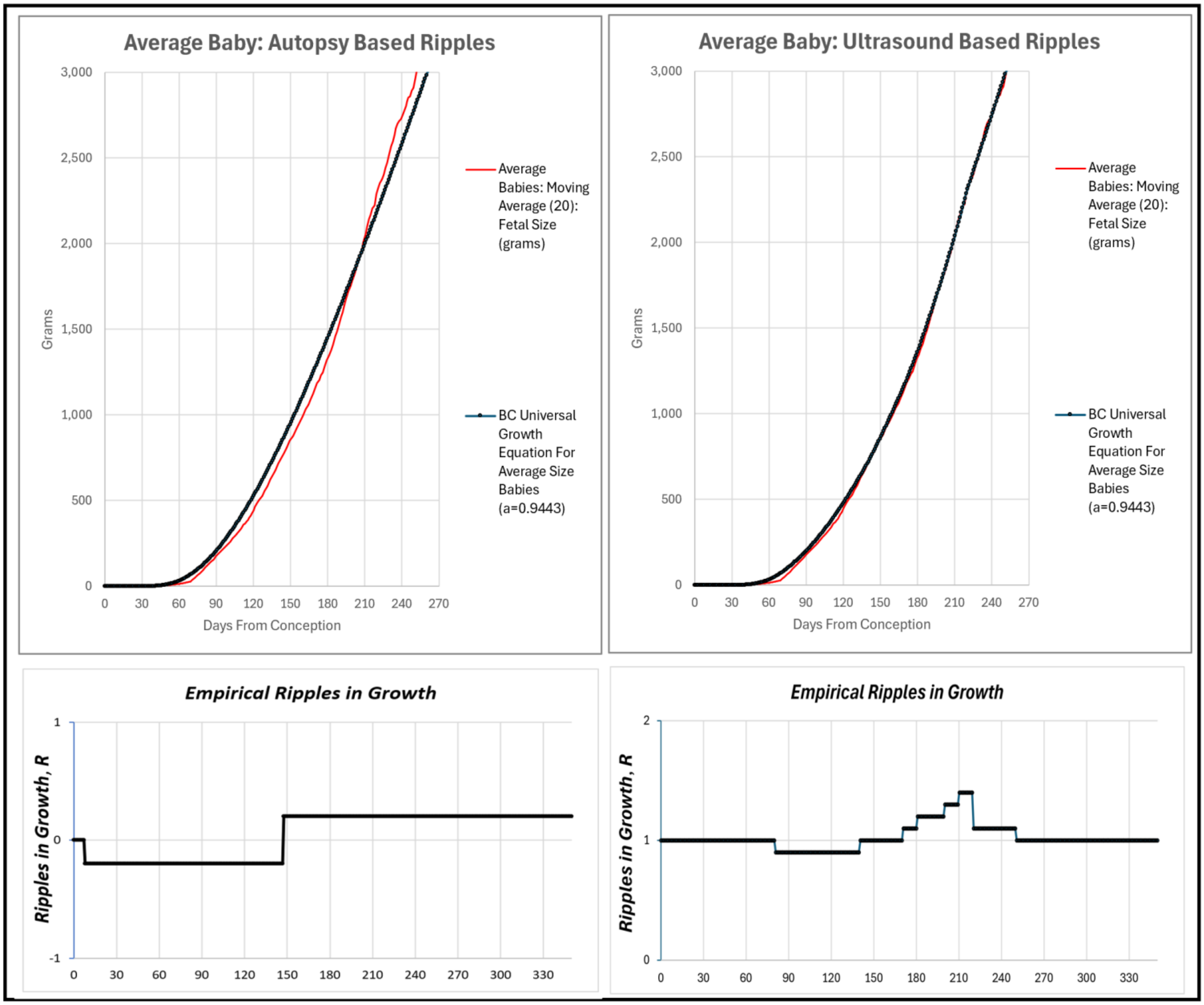


**Supplement-Figure-III.9** **Empirical *Ripples*** for data from the "***Average Size Baby Study Group***"

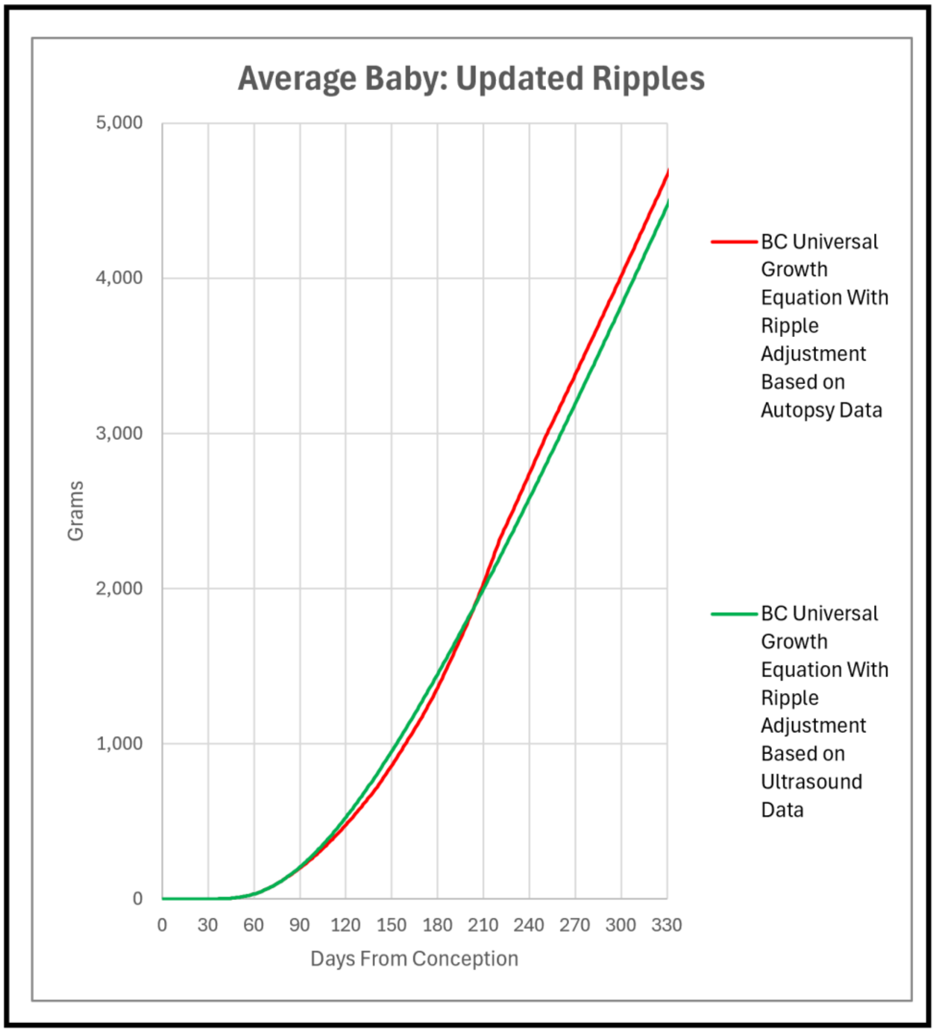


**<u>Supplement-Figure-III.10</u>** . ***Ripple*** corrected ***Binary Cellular Universal Growth Equation*** (#2), in grams, from data from the "***Average Size Baby Study Group***

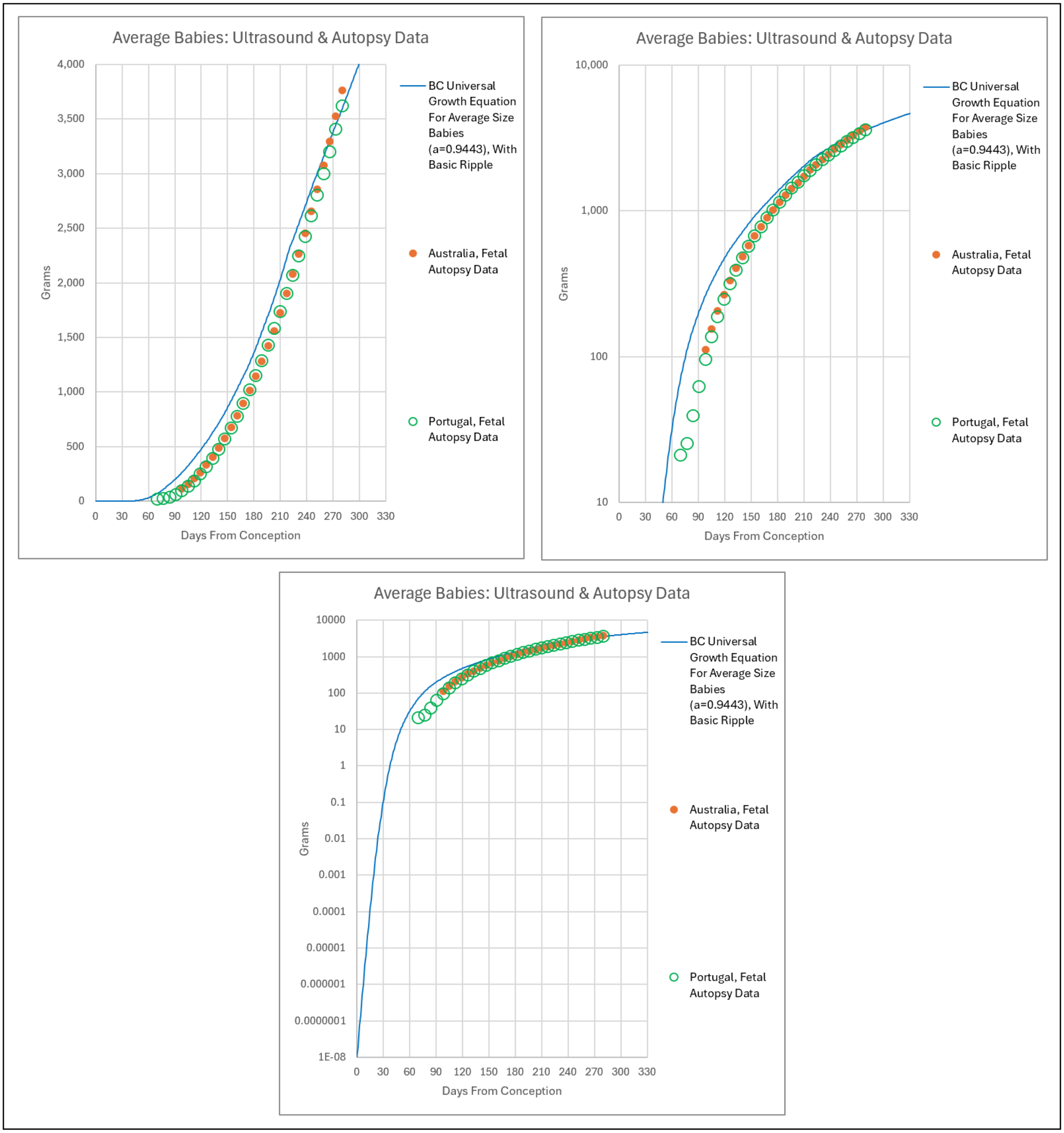


**<u>Supplement-Figure-III.11</u>** ***Ripple*** corrected ***Binary Cellular Universal Growth Equation*** (#2), in grams, from data from the "***Average Size Baby Study Group***, compared to fetal autopsy data.

Again, it is straightforward to convert such weight-based data into units of cells, ***N***.[273]

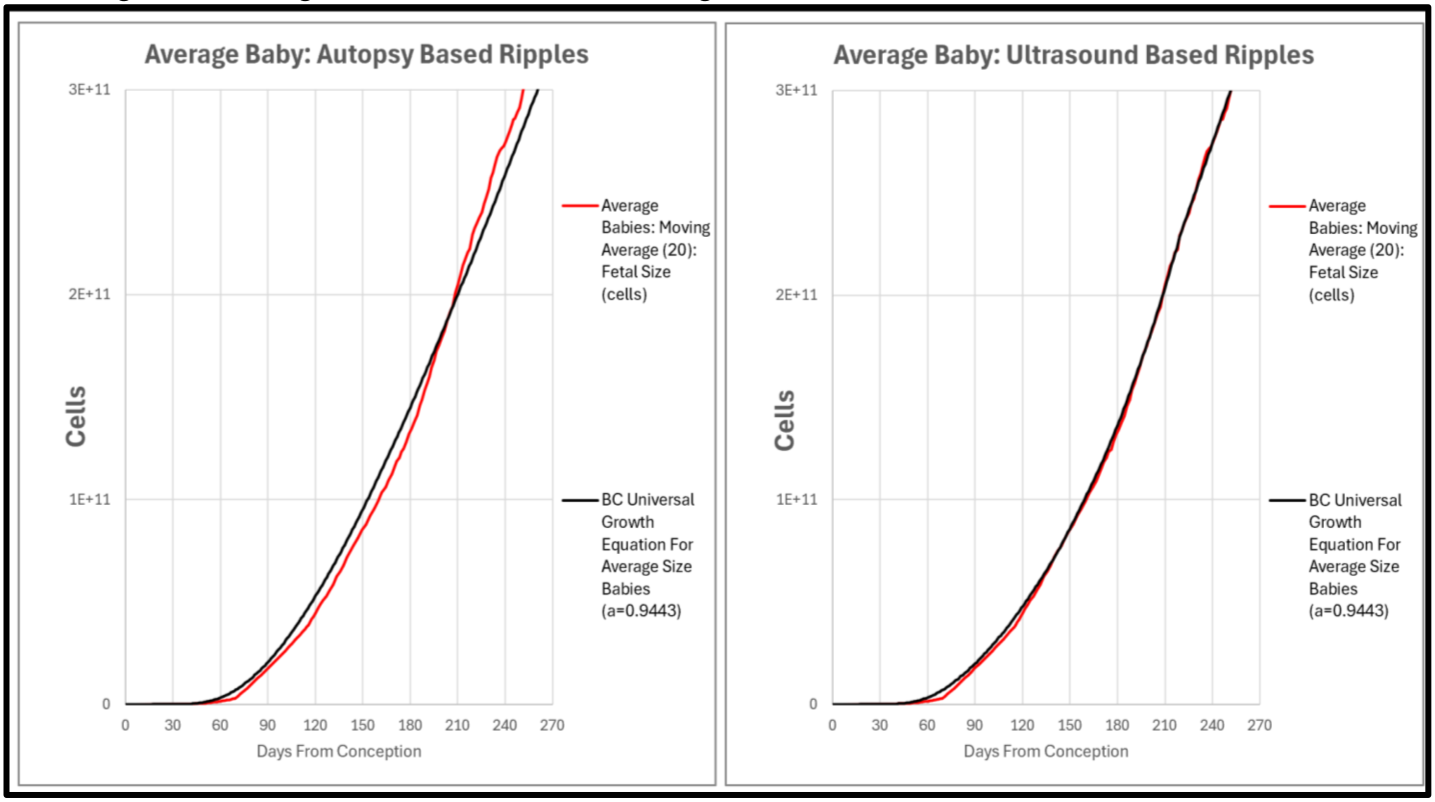


**<u>Supplement-Figure-III.12</u>**. Updated ***Ripple*** corrected ***Binary Cellular Universal Growth Equation*** (#2), in units of numbers of cells, ***N***, from data from the "***Average Size Baby Study Group***"

***Binary Cellular Fetal Growth Analysis*** of ***Small***, ***Average***, and ***Big Babies***, Examined as Groups, in Grams.

Extending the ***Binary Cellular Growth*** method described above makes it possible to create ***Binary Cellular Fetal Growth Charts*** of the change in size of fetuses over time for fetuses that grow to different ***Birthweights***: "***Average Size Baby Study Group***", "***Small Baby Study Group***", and "***Big Baby Study Group***", (**TABLE III.1**, **Supplement-Figures III.5&6**). To recap, we graph the ***Fetal Weight*** of each ultrasound measurement, calculated by the ***Abdominal Circumference Binary Cellular Estimated Fetal Weight Equation*** (#2), against the ***Fetal Age*** when each ultrasound measurement was made, calculated by the ***CRL Fetal Age Equation*** (#7a). The resulting cloud of datapoints, and curves for the ***Moving Averages*** that smooth these values, can be seen in **Supplement-Figure-III.13**&14.

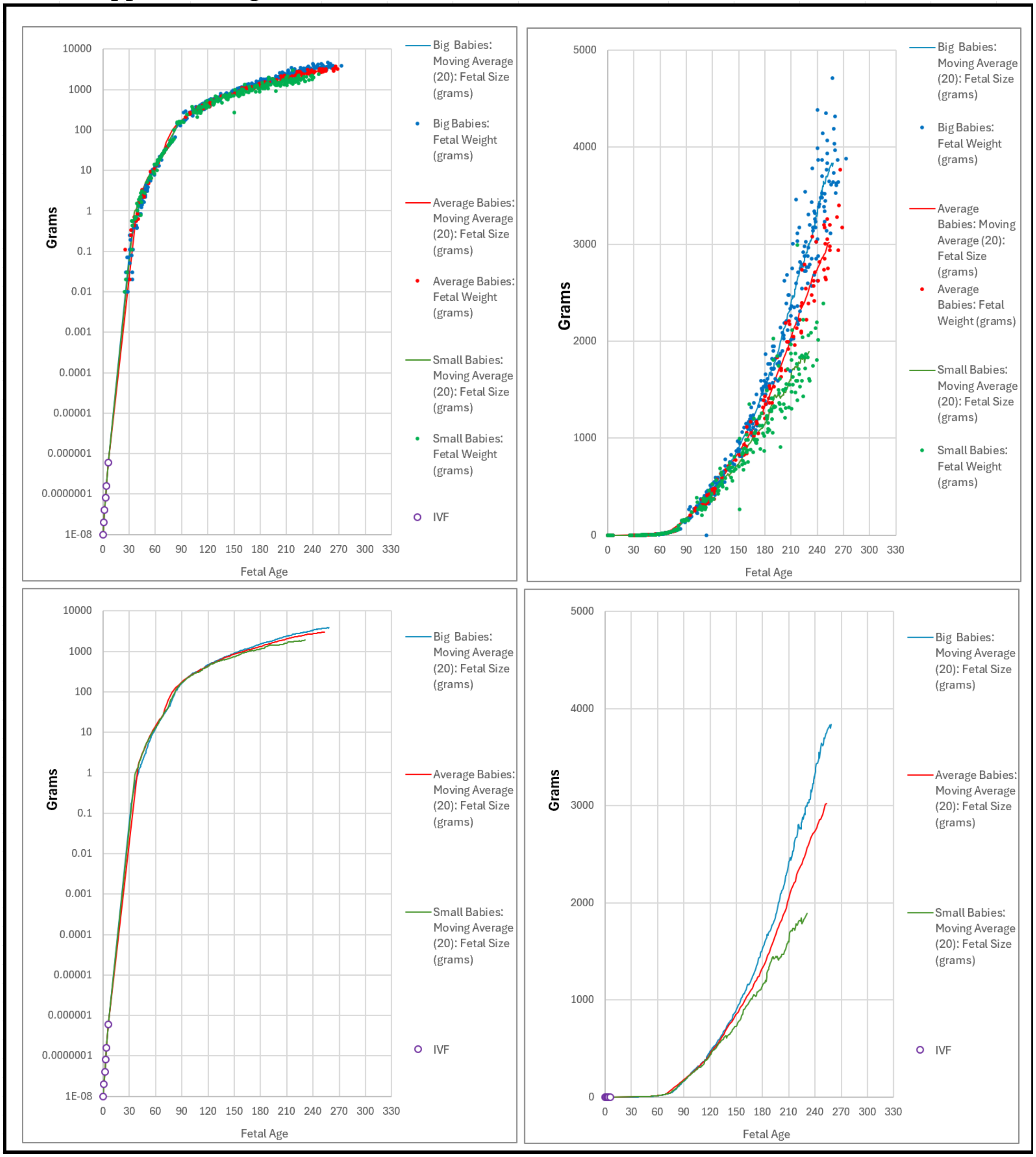


**Supplement-Figure-III.13** ***Binary Cellular Fetal Growth Charts*** capturing the ***Fetal Weight*** (grams) of each ultrasound measurement, calculated by the ***Abdominal Circumference Binary Cellular Estimated Fetal Weight Equation*** (#13), against the ***Fetal Age*** when each ultrasound measurement was made, calculated by the ***CRL Fetal Age Equation*** (#7a), ***for the babies in the*** "***Average Size Baby Study Group***, "***Small Baby Study Group***", **and** "***Big Baby Study Group***"

***Binary Cellular Fetal Growth Analysis*** of ***Small***, ***Average***, and ***Big Babies***, Examined as Groups, in Cells

Yet once again, it is straightforward to convert these weight-based data into units of cells, ***N***.[273]

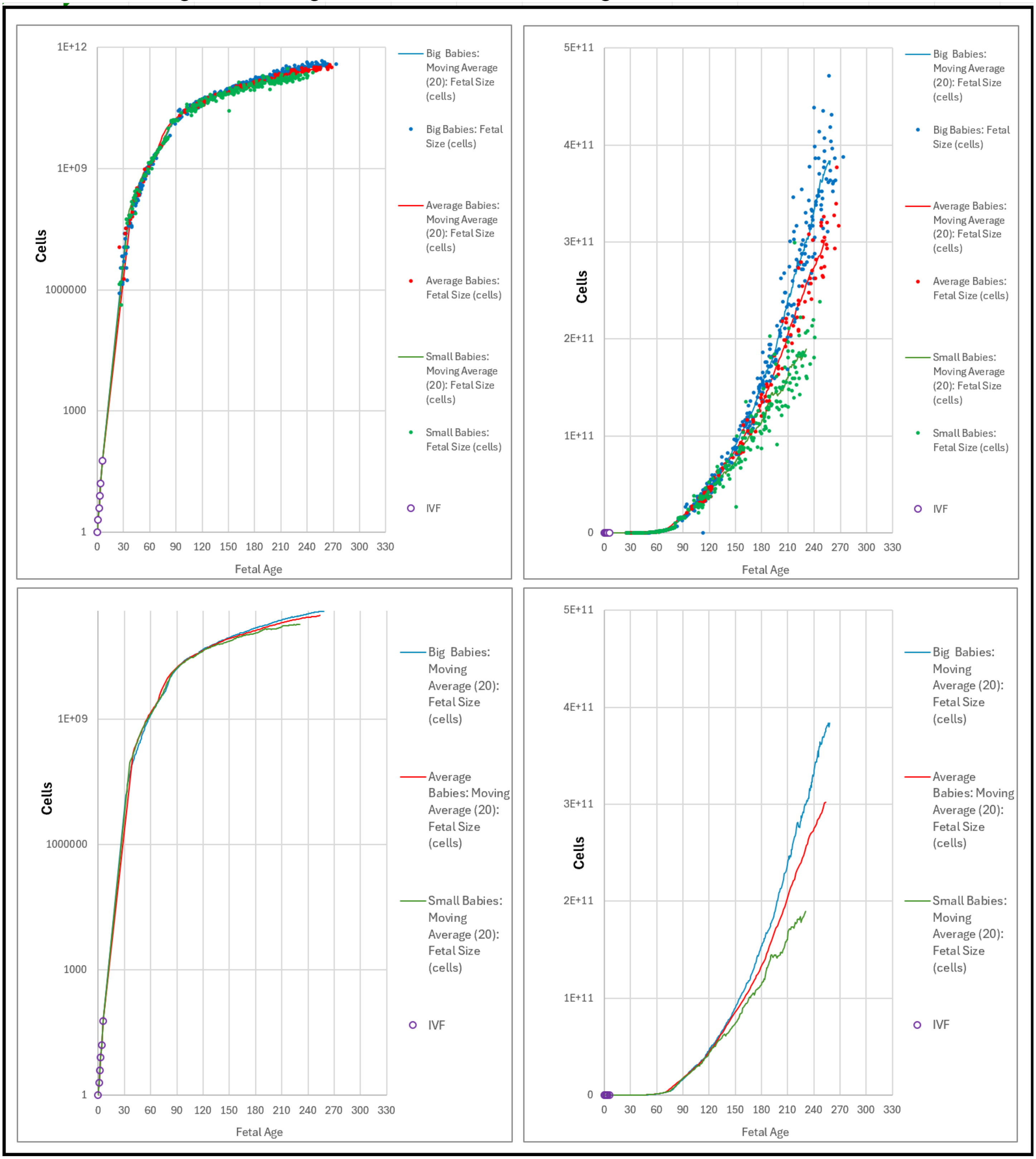


**Supplement-Figure-III.14** ***Binary Cellular Fetal Growth Charts*** capturing the ***Fetal Size*** (Cells) of each ultrasound measurement, calculated by the ***Abdominal Circumference Binary Cellular Estimated Fetal Weight Equation*** (#13), against the ***Fetal Age*** when each ultrasound measurement was made, calculated by the ***CRL Fetal Age Equation*** (***#7a***), ***for the babies in the*** "***Average Size Baby Study Group***, "***Small Baby Study Group***", **and** "***Big Baby Study Group***"

***Very Big Babies*** Arise from More Rapid Growth than ***Marginally Big Babies***.

***Large Babies*** (***Macrosomia***) are classified into three categories: ***Grade 1*** (4000 to 4499 grams), ***Grade 2*** (4500 to 4999 grams), and ***Grade 3*** (over 5000 grams). We didn't have enough ***Grade 3 Babies*** to create ***Binary Cellular Fetal Growth Charts***, but it was possible to do so for ***Grade 1*** and ***Grade II Macrosomia***. As one might suspect, the group of fetuses that become ***Grade 2 Macrosomia Babies*** show a faster growth curve than the group of fetuses that become ***Grade 1 Macrosomia Babies***. The groups of both ***Grade 1*** and ***Grade 2 Macrosomia Babies*** show faster growth curves than the group of 15 fetuses that become ***Average Size Babies*** **Supplement-Figure-III. 15**.

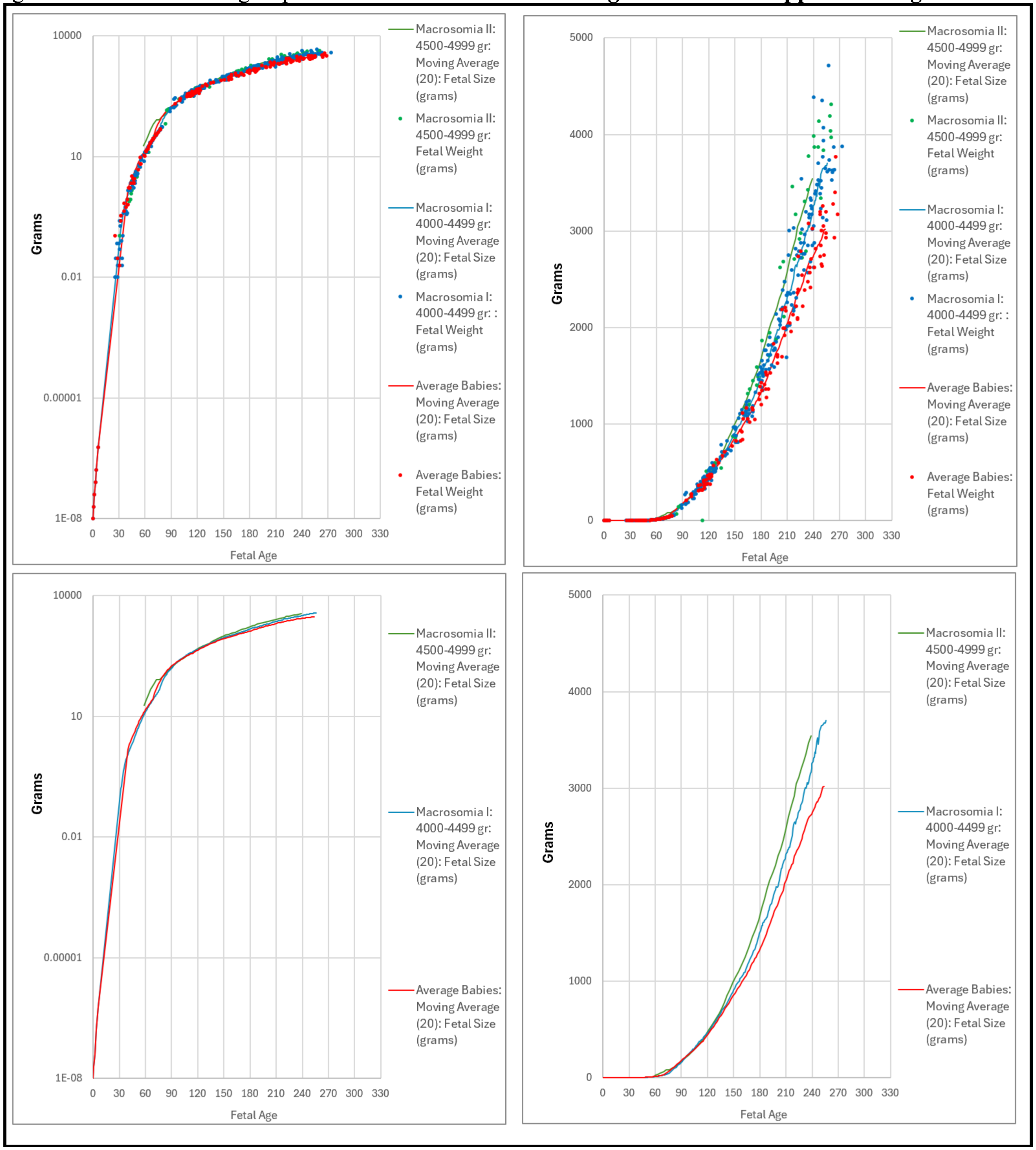


**Supplement-Figure-III. 15** ***Binary Cellular Fetal Growth Charts*** capturing the ***Fetal Weight*** (grams) of each ultrasound measurement, calculated by the ***Abdominal Circumference Binary Cellular Estimated Fetal Weight Equation*** (#13), against the ***Fetal Age*** when each ultrasound measurement was made, calculated by the ***CRL Fetal Age Equation*** **(*#7a*),** for ***Grade 1*** and ***Grade II Macrosomic Babies***, and ***Average Size Babies***

***Binary Cellular Fetal Growth Analysis*** Reveals How ***Size***, ***Weight-Gain*** and ***Mitotic Fraction*** Occur Over Time.

Underneath the change in the ***Size*** of a fetus over time lies ***Cellular Selection,*** that is ***Differential Cellular Proliferation***, the main mechanism for the creation and control of ***Growth***. The ***Binary Cellular Universal Mitotic Fraction Equation***, which lies within the ***Binary Cellular Universal Growth Equation***, captures this mechanism of ***Growth*** to adult size, control of cell division, through the metric of the ***Mitotic Fraction***, ***m***, the fraction of cells that must be dividing to account for the growth of the body. The ***Mitotic Fraction***, ***m***, is calculated by examining ***Growth*** in units of numbers of cells, ***N***, that is to say, the ***Mitotic Fraction Method*** of ***Binary Cellular Analysis***. The ***Weight-Gain Rate*** is calculated by new grams per day, which can then be transformed into new cells per day, by using the approximation of $10^8$ cells/gram[82].

This information is captured statistically, by the ***Moving Average*** of the fetal weight values derived from ultrasound measurements (Black Lines, **Supplement-Figure-III.16**), and analytically, by the ***Binary Cellular Universal Growth Equation*** (#2) (Red Lines). These two methods yield subtle differences, especially early in pregnancy, when we have few ultrasound measurements to work with. However, these differences can be narrowed as more data becomes available, but for now, these minor gaps do not impede gaining useful new insights, as we shall see below.

Roughly speaking, the ***Weight-Gain Rate***, most easily pictured when we consider it in numbers of new cells per day, is the ***Mitotic Fraction***, ***m***, ***TIMES*** the ***Fetal Size***, also considered in numbers of cells, ***N***. Note that while the ***Mitotic Fraction***, ***m***, declines progressively with time, the ***Weight-Gain Rate*** increases, until about 230 days after conception, and then declines.

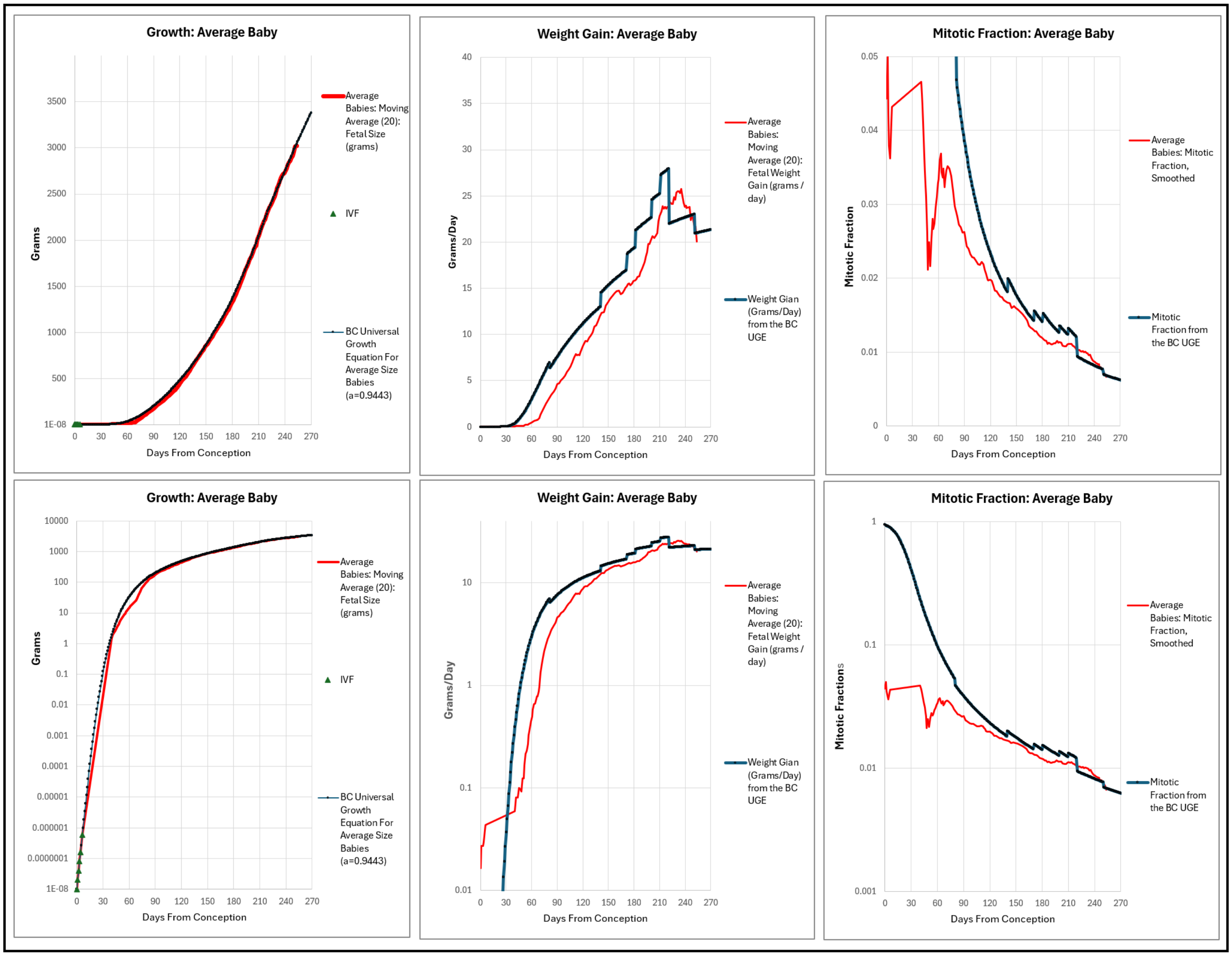


**Supplement-Figure-III.16** ***Growth***, ***Mitotic Fraction***, and ***Weight-Gain Rate***, for "***Average Size Baby Study Group***"

#### The Mitotic Basis of ***Growth*** to ***Small***, ***Average,*** and ***Big Babies*** Revealed by ***Weight-Gain*** and ***Mitotic Fraction***

***Binary Cellular*** calculations of ***Mitotic Fraction*** (the fraction of cells that must be dividing to account for the growth of the body), and ***Weight-Gain Rate*** (growth, in ***Grams***/***Day***), are higher for fetuses that will become ***Big Babies***, and lower for fetuses that will become ***Small Babies*** ("***Average Size Baby Study Group***", "***Small Baby Study Group***", and "***Big Baby Study Group***", **Supplement-Figure-III.?**).  Graphing these data show that these differences occur over much of the ***Fetal Age*** range for which we have data, that is, from about 120 days after conception until birth (**Supplement-Figure-III.17**).

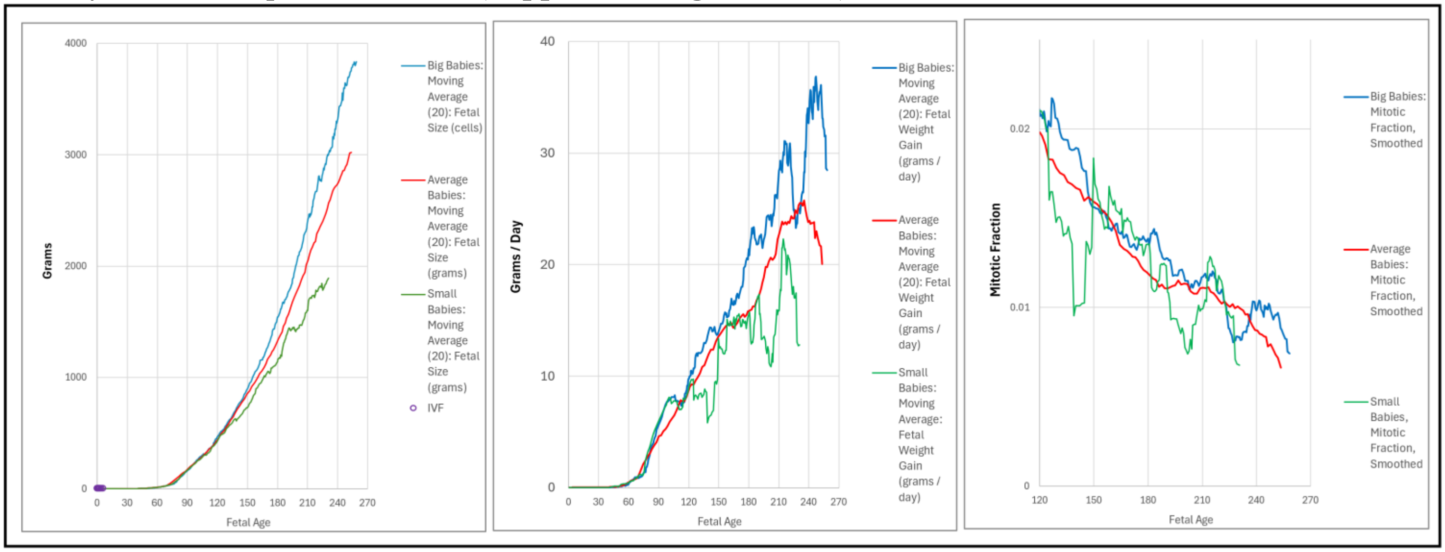


**Supplement-Figure-III.17**   ***Growth***, ***Mitotic Fraction***, and ***Weight-Gain Rate***, for "***Average Size Baby Study Group***", "***Small Baby Study Group***" and "***Big Baby Study Group***"

Fetuses that Become ***Average*** and ***Big Babies*** Have ***Mitotic Fractions*** that Differ Over Long Periods of Time.

***Binary Cellular Fetal Growth Analysis*** reveals that the ***Mitotic Fraction*** of fetuses that become ***Big Babies*** runs roughly parallel and higher than the ***Mitotic Fraction*** of the fetuses that will become ***Average Size Babies***, over much of the span for which we have data, that is, from ~120 days after conception until birth ("***Average Size Baby Study Group***" and "***Small Baby Study Group***", **Supplement-Figure-III.18**). Thus, the data suggest that it is the ***Mitotic Fraction***, the fraction of cells that must be dividing to account for the growth of the body, persisting at a higher level over much, perhaps all, of the length of pregnancy, that drives fetuses to become ***Big Babies***.

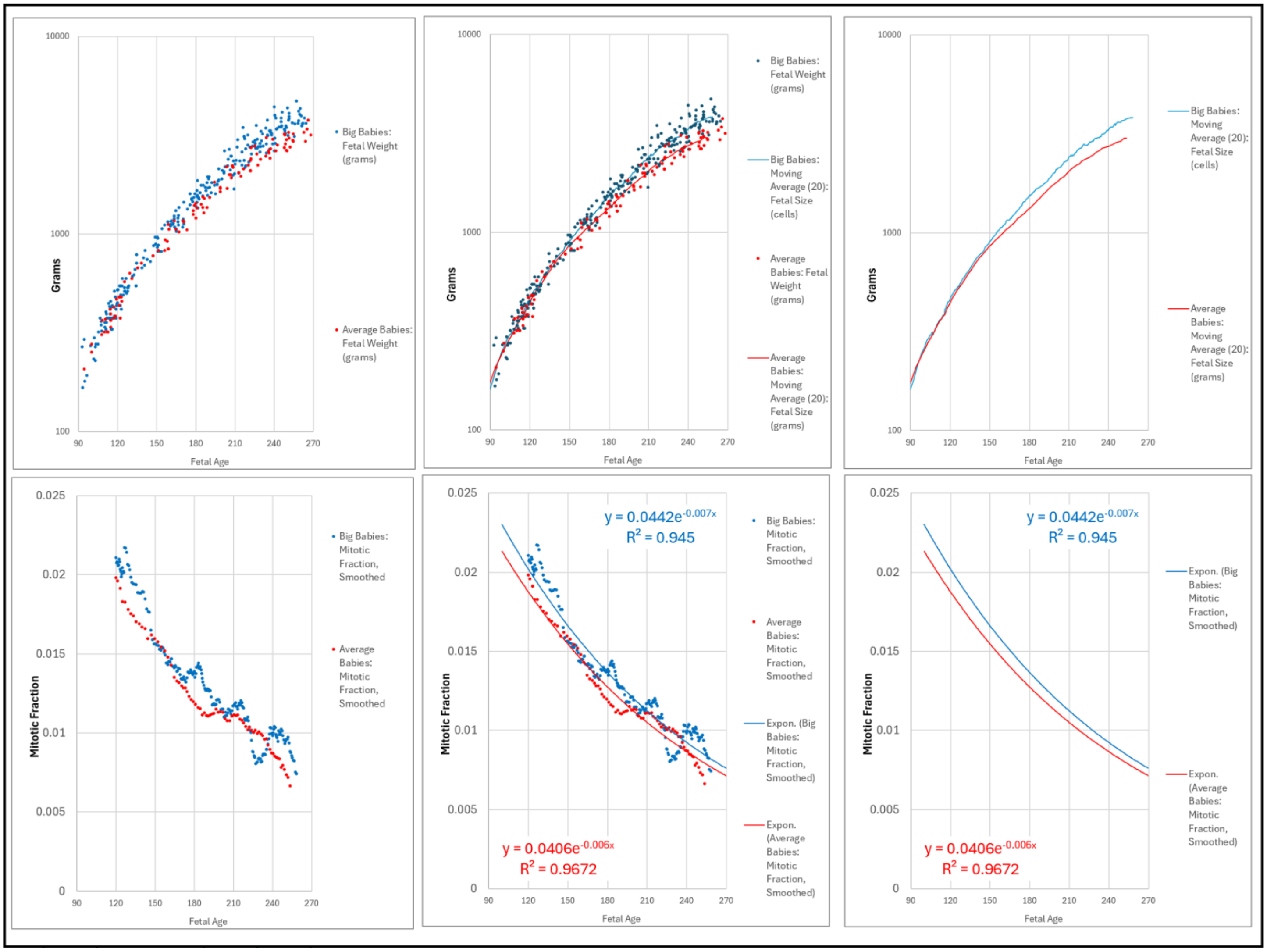


**Supplement-Figure-III.18**. ***Growth***, and ***Mitotic Fraction***, for "***Average Size Baby Study Group***" and "***Big Baby Study Group***", fit to an exponential function and moving average.

***Very Big Babies*** Arise from Higher ***Mitotic Fractions*** than ***Marginally Big Babies***.

Recall that, as we saw above (**Supplement-Figure-III.18**), ***Binary Cellular Fetal Growth Analysis*** of ultrasound data told us that fetuses that become ***Grade 2 Macrosomia Babies*** (4500 to 4999 grams) show faster growth curves than fetuses that become ***Grade 1 Macrosomia Babies*** (4000 to 4499 grams), and both are faster than ***Average Size Babies***. As can be seen in **Supplement-Figure-III.19**), this difference in the two ***Grades*** of ***Macrosomia Birthweight*** is also reflected in the ***Mitotic Fraction***, the fraction of cells that must be dividing to account for the growth of the body.

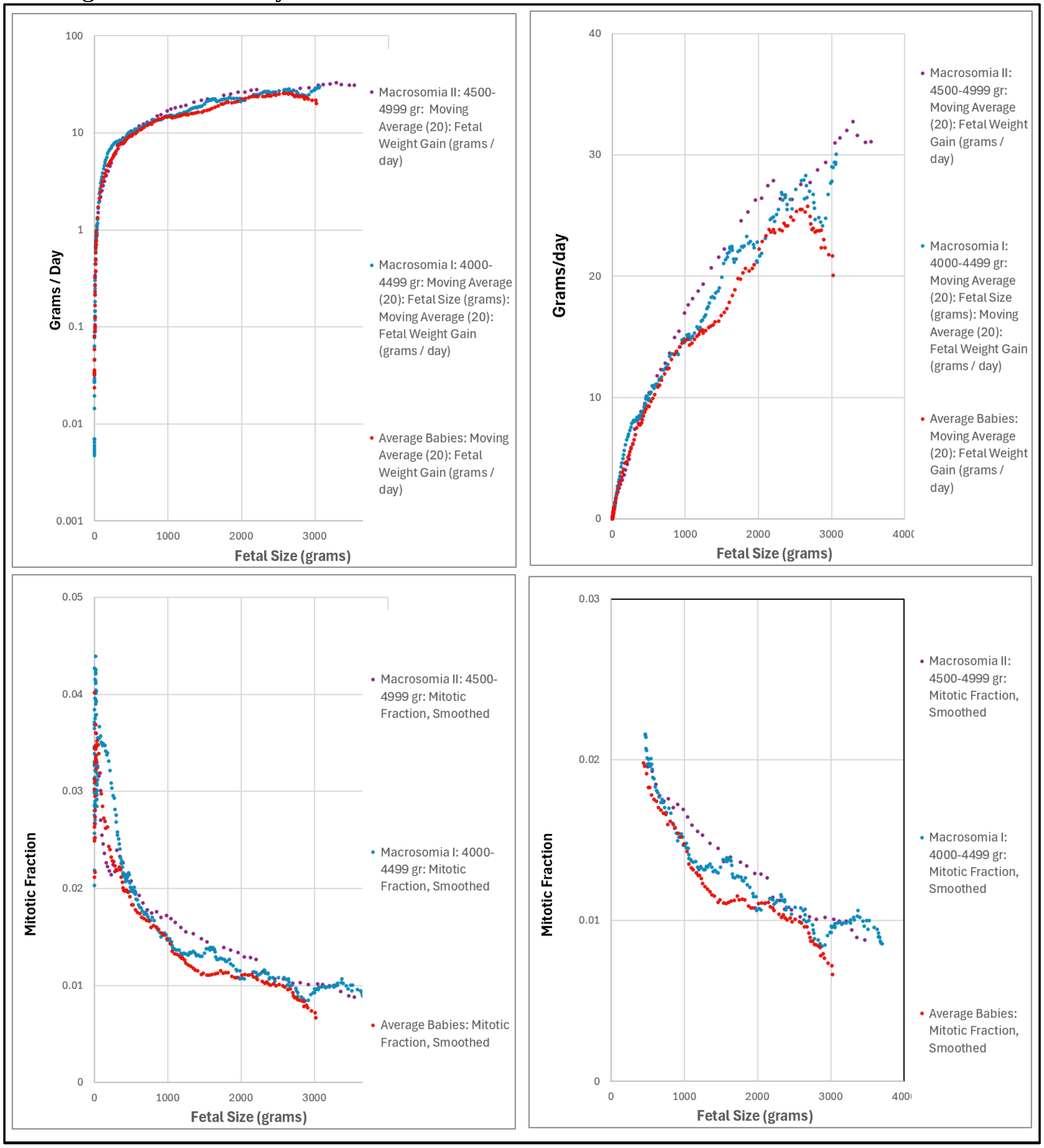


**Supplement-Figure-III.19**. ***Growth***, and ***Mitotic Fraction***, for "***Average Size Baby Study Group***" and the ***Grade 2 Macrosomia Babies*** (4500 to 4999 grams) and ***Grade 1 Macrosomia Babies*** (4000 to 4499 grams) of the"***Big Baby Study Group***".

Fetuses of ***Big Babies*** Grow Bigger by Having Bigger ***"a" Parameters***, Giving Bigger ***Mitotic Fractions***

At the core of the ***Binary Cellular Universal Growth Equation*** (#2), and the growth of human fetuses which it captures, lies the ***Mitotic Fraction***, ***m***, the fraction of cells that must be dividing to account for the growth of the body, which declines as we get bigger. As outlined in the first paper in this series, for a great variety of species, including humans, the ***Mitotic Fraction*** declines by the ***Binary Cellular Mitotic Fraction Equation*** (#4):

$$\boldsymbol{mitotic\ fraction = a^{(N_w^b)}} \quad (2)$$

where ***a*** and ***b*** characterize the growth of each species, and ***N*** is the size of the animal, in units of numbers of cells.

That the ***Mitotic Fraction***, ***m***, declines by the ***Binary Cellular Mitotic Fraction Equation*** (#4) was seen by graphing the log(-log) of ***m***, the ***Mitotic Fraction***, against the log of ***N***, revealing a straight line. We call this the ***Mitotic Fraction Log(Log)/Log Test***, and the line that confirms the ***Binary Cellular Mitotic Fraction Equation*** (#4), the ***Mitotic Fraction Log(Log)/Log Line***. To recap the arithmetic:

***m = a^(N^b)*** (90)

***-Log(m) = -Log (a^(N^b))*** (91)

***-Log(m) = (N^b) * (-Log (a)*** (92)

***Log (-Log(m)) = Log ( (N^b) * (-Log (a) )*** (93)

***Log (-Log(m)) = Log( (N^b) ) + Log (-Log (a) )*** (94)

***Log (-Log(m)) = b * Log( (N) + Log (-Log (a) )*** (95)

***Log (-Log(m)) = b * Log( (N) + Z*** (96)

where

***Z = Log (-Log (a)*** (97)

Since the calculations we carried out used ***Log*** to the base 10: (98)

***-Log(a) = 10^Z*** (99)

***a = 10^(-10^Z)*** (999)

Thus the ***Mitotic Fraction Log(Log)/Log Line*** has a slope that reflects the ***b*** parameter of the ***Binary Cellular Mitotic Fraction Equation*** (#4), and a height that reflects the ***"a" Parameter***, which we shall see below, provides the key to predicting the size at which a human fetus is born.

This ***Log(Log)/Log*** linearity over the full lifespan can be seen in **Supplement-Figure-I.2**, and for ***Average Size Babies*** and ***Big Babies*** in **Supplement-Figure-III.20** below ("***Average Size Baby Study Group***" vs "***Big Baby Study Group***"). Note in **Supplement-Figure-III.20** that that ***Mitotic Fraction Log(Log)/Log Test*** reveals the ***Mitotic Fraction Log(Log)/Log Lines*** of ***Average Size Babies*** and ***Big Babies*** are roughly parallel in slope, but of different in height. That is to say, the ***Mitotic Fraction Log(Log)/Log Test*** reveals that ***Big Babies*** grow bigger than ***Average Size Babies***, by a difference in the ***"a" Parameter*** of the ***Binary Cellular Mitotic Fraction Equation*** (#2).

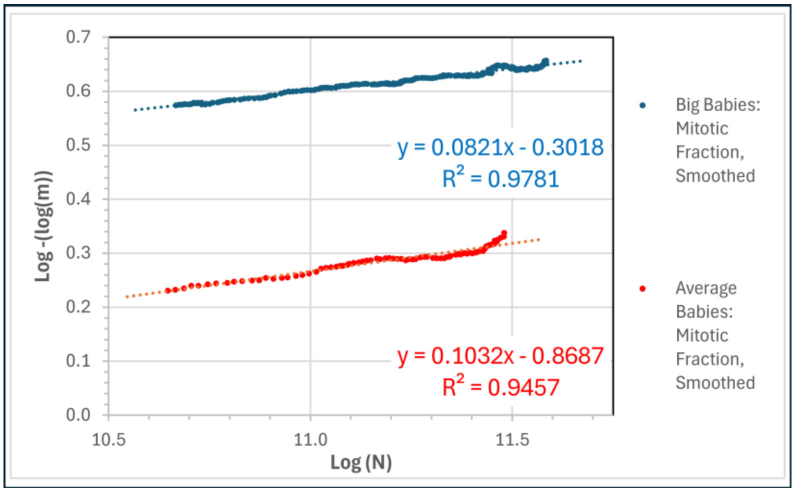


**Supplement-Figure-III.20** ***Mitotic fraction Log(Log)/Log Test*** for "***Average Size Baby Study Group***" and "***Big Baby Study Group***".

***Big Babies*** are Bigger by***Binary Cellular Universal Growth Equations*** with Bigger ***"a" Parameters***.

Replacing the ***Average Size Baby "a" Parameter*** value of 0.9443 with the ***Big Baby "a" Parameter*** value of 0.94475 in the ***Binary Cellular Universal Growth Equation*** (#2) shows how well it now fits the growth data of fetuses that will become ***Big Babies*** ("***Average Size Baby Study Group***" vs "***Big Baby Study Group***", **Supplement-Figure-III.21**).

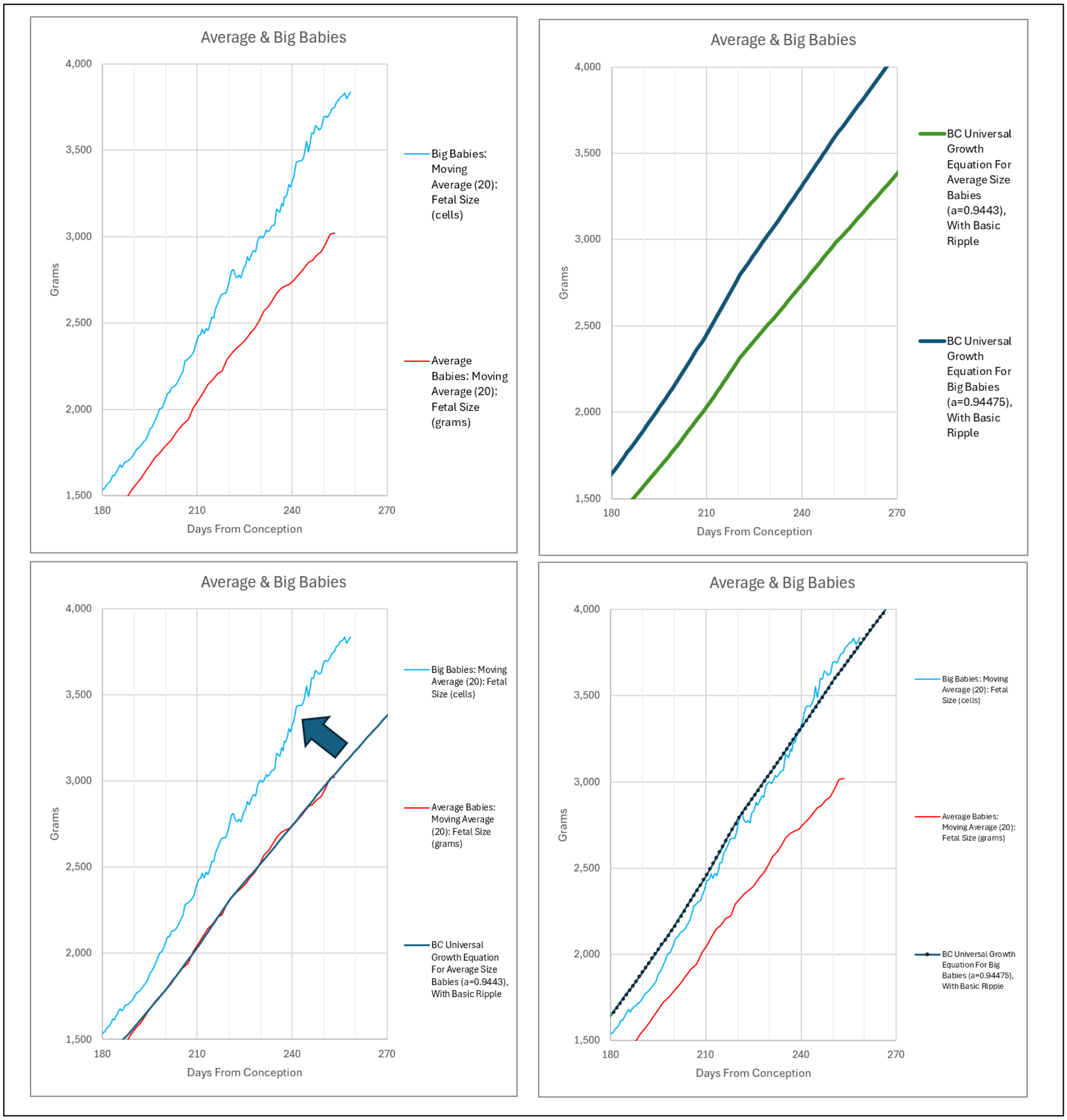


**Supplement-Figure-III.21**. Replacing the ***Average Size Baby "a" Parameter*** value of 0.9443 with the ***Big Baby "a" Parameter*** value of 0.94475 in the ***Binary Cellular Universal Growth Equation*** (#2), in comparison to growth data, from fetal ultrasound moving averages of fetuses that will become ***Average Size Babies*** ("***Average Size Baby Study Group***") vs ***Big Babies*** ("***Big Baby Study Group***").

***Small Babies*** are Small by ***Binary Cellular Universal Growth Equations*** at Small ***“a” Parameters***.

Replacing the ***Average Size Baby “a” Parameter*** value of 0.9443 with the ***Small Baby “a” Parameter*** value of 0.9441 in the ***Binary Cellular Universal Growth Equation*** (#2) shows how well it now fits the growth data of fetuses that will become ***Small Babies*** ((“***Average Size Baby Study Group***” vs “***Small Baby Study Group***”, **Supplement-Figure-III.22**,) Unfortunately, the numbers of ***Small Babies*** for which we have data are too few to carry out a ***Mitotic Fraction Log***(***Log***)***/Log Test***, and this certainly would be worthwhile when more data become available.

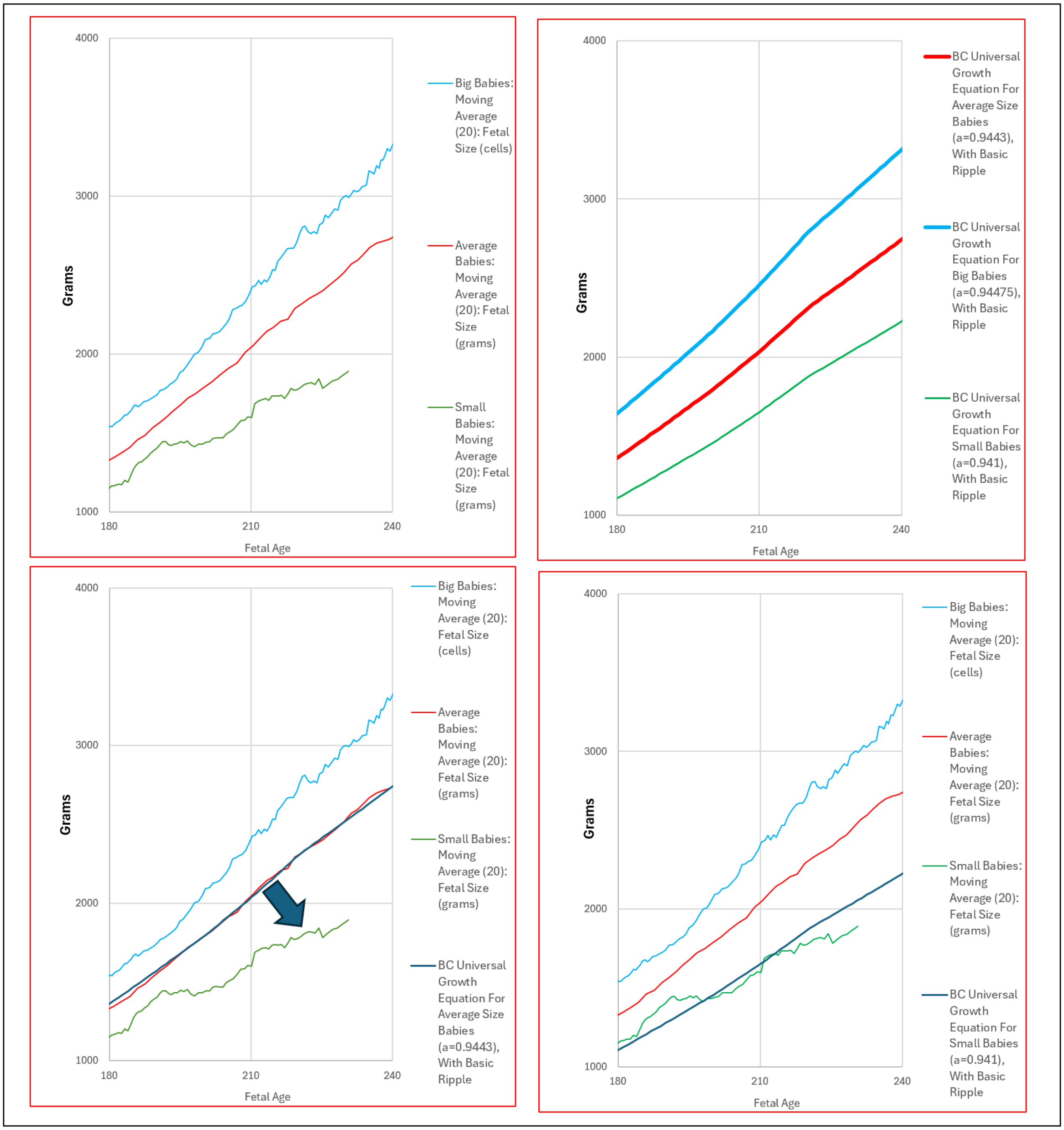


**Supplement-Figure-III.22** Replacing the ***Average Size Baby “a” Parameter*** value of 0.9443 with the ***Small Baby “a” Parameter*** value of 0.941 in the ***Binary Cellular Universal Growth Equation*** (#2), in comparison to growth data, from fetal ultrasound moving averages of fetuses that will become ***Average Size Babies*** (“***Average Size Baby Study Group***”) vs ***Small Babies*** (“***Small Baby Study Group***”).

***Binary Cellular Universal Growth Equation*** with ***"a" Parameters*** of ***Small, Average, Big Babies***

***Binary Cellular Fetal Growth Charts*** showing the changes in the ***"a" Parameter*** of the ***Binary Cellular Universal Growth Equation*** (#2) that fit the growth of fetuses that become ***Small***, ***Average,*** and ***Big Babies*** can be seen in **Supplement-Figure-III.23** ("***Average Size Baby Study Group***", "***Small Baby Study Group***", and "***Big Baby Study Group***"). ***Binary Cellular Fetal Growth Charts*** of these three curves over a variety of ranges can be seen in **Supplement-Figure-III.24**.

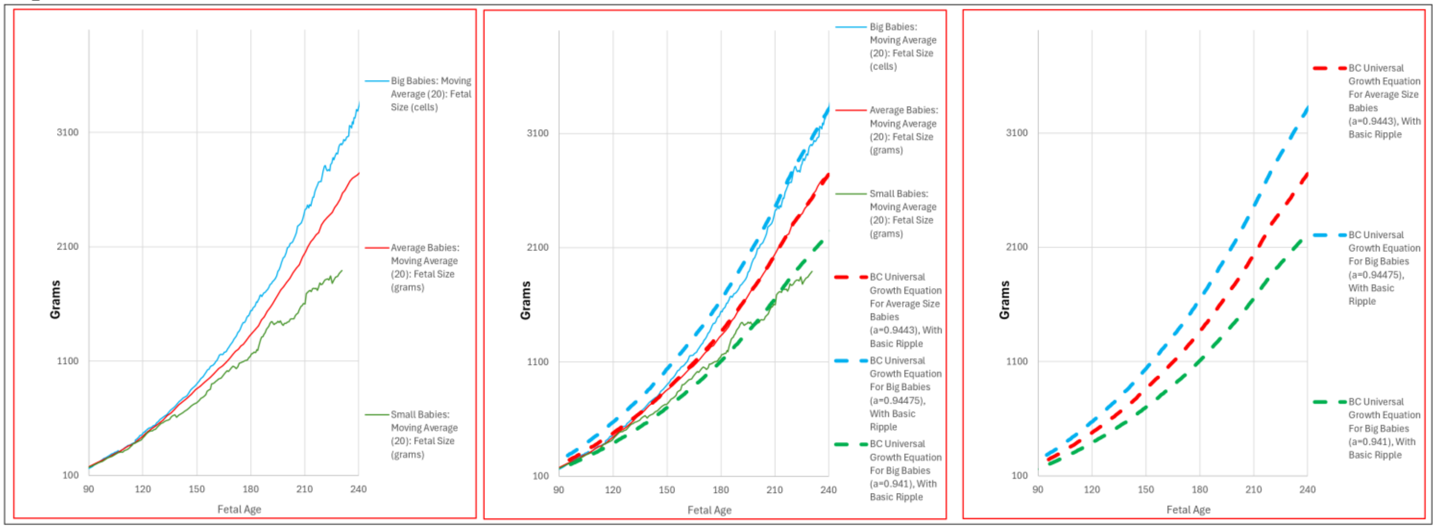


**Supplement-Figure-III.23** "***Average Size Baby Study Group***", "***Small Baby Study Group***", **and** "***Big Baby Study Group***", captured by the ***"a" Parameter*** value in the ***Binary Cellular Universal Growth Equation***.

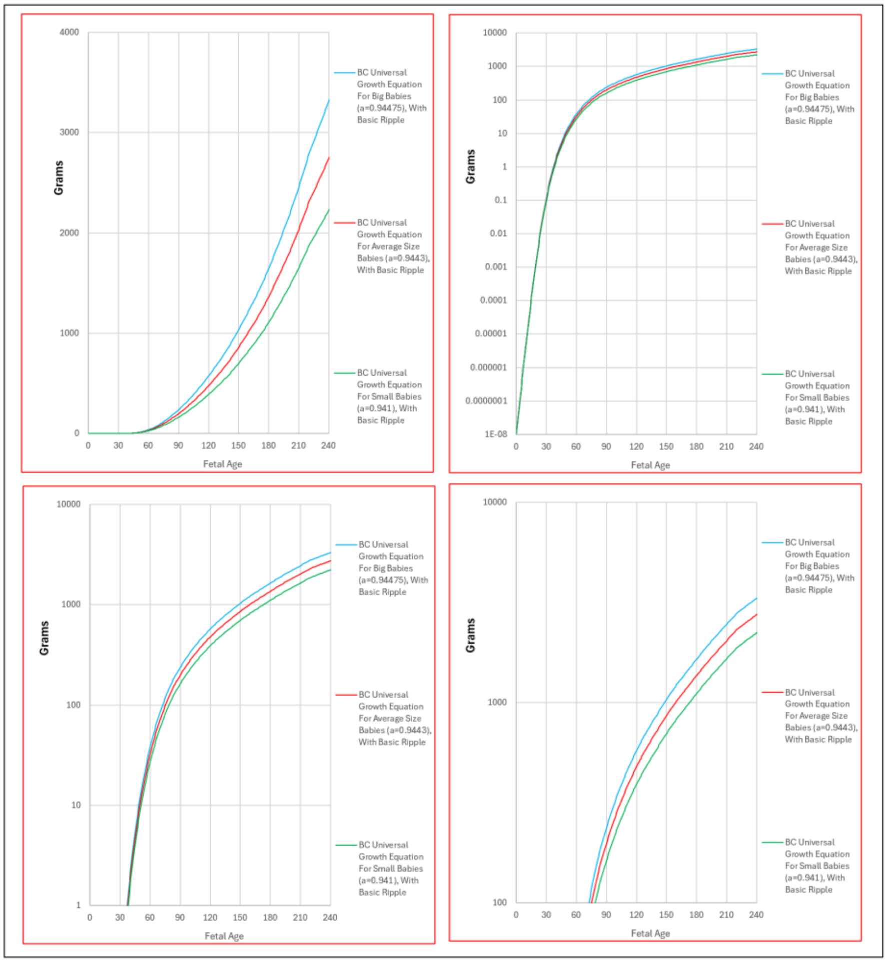


**Supplement-Figure-III.24** "***Average Size Baby Study Group***", "***Small Baby Study Group***", **and** "***Big Baby Study Group***", captured by the ***"a" Parameter*** value in the ***Binary Cellular Universal Growth Equation***.

Fetuses That Become ***Small***, ***Average,*** and ***Big Babies*** Display ~Linear Growth, at ***Slopes*** Reflecting ***Birthweight***.

Recall, as we showed in the first paper in this series, that the ***Binary Cellular Universal Growth Equation*** (#2) displays a high degree of curvature over the first months of pregnancy, then settling into a relatively constant rate of growth in the last months of pregnancy. We call this the ***~Linear Phase of Human Fetal Growth***.

The ***Moving Average*** value for the fetal weights in our datasets show this ***~Linearity*** to be the case for fetuses that become ***Big***, ***Average***, and ***Small Babies*** ("***Average Size Baby Study Group***", "***Small Baby Study Group***", and "***Big Baby Study Group***", **TABLE III.1**) with each group displaying different ***Weight-Gain Rates***. Fetuses that become ***Big Babies*** ("***Big Baby Study Group***") have a ***Weight-Gain Rate*** of ~31 grams/day. Fetuses that became ***Average Size Babies*** ("***Average Size Baby Study Group***") have a ***Weight-Gain Rate*** of ~24 grams/day. Fetuses that become ***Small Babies*** ("***Small Baby Study Group***") have a ***Weight-Gain Rate*** of ~14 grams/day (**Supplement-Figure-III.25**).

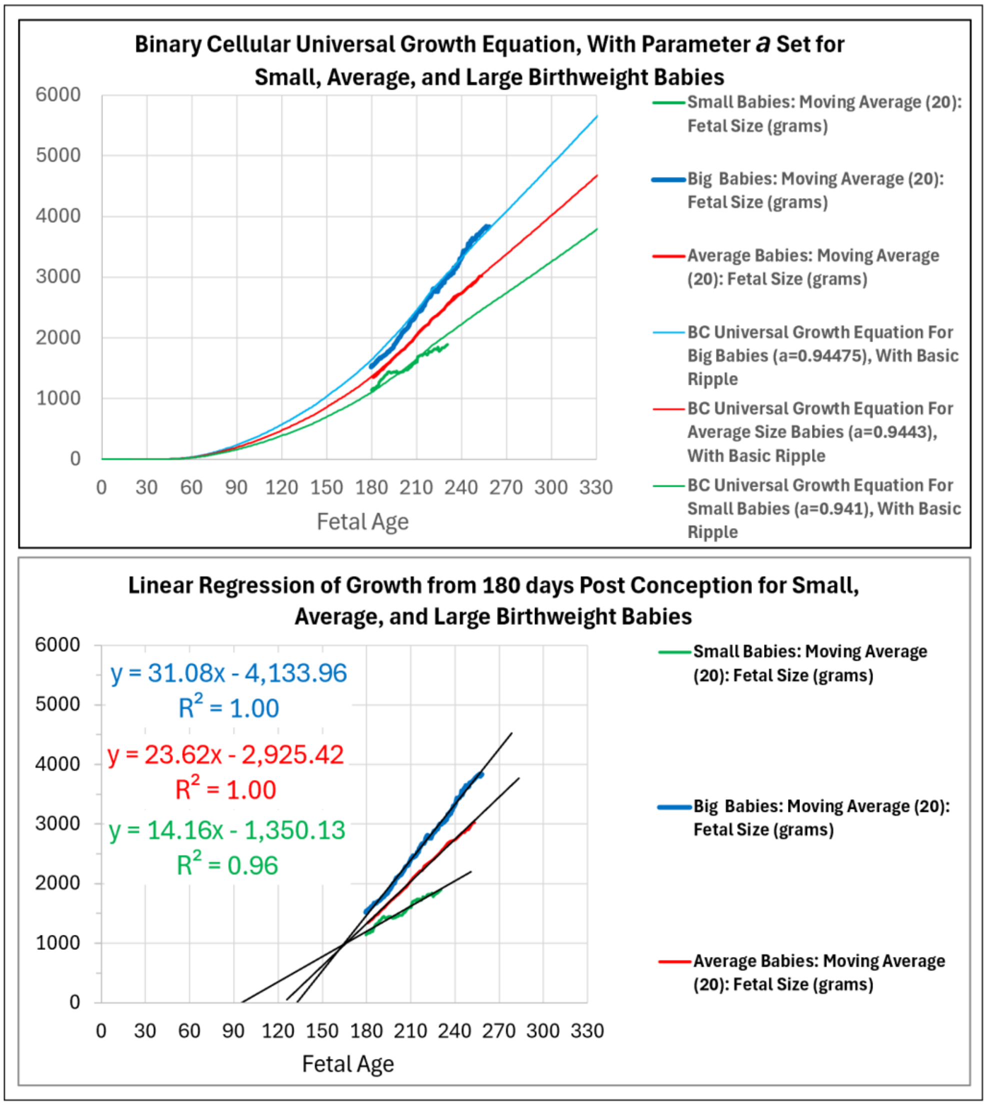


<u>**Supplement-Figure-III.25**</u> ***Weight-Gain Rate***, and fit by variation of the ***"a" Parameter*** value in the ***Binary Cellular Universal Growth Equation***, to babies of the "***Average Size Baby Study Group***", "***Small Baby Study Group***", and "***Big Baby Study Group***".

**_Binary Cellular Fetal Growth Analysis_: _Fetal Weight, Fetal Growth_, _Pregnancy Length_ and _Birthweight_**

We can begin to visualize the relationship between ***Fetal Weight***, the ***Rate of Growth***, ***Pregnancy Length***, and the ultimate ***Birthweight*** at the end of pregnancy, by displaying all of these values on a ***Binary Cellular Fetal Growth Chart*** (**Supplement-Figure-III.26**).

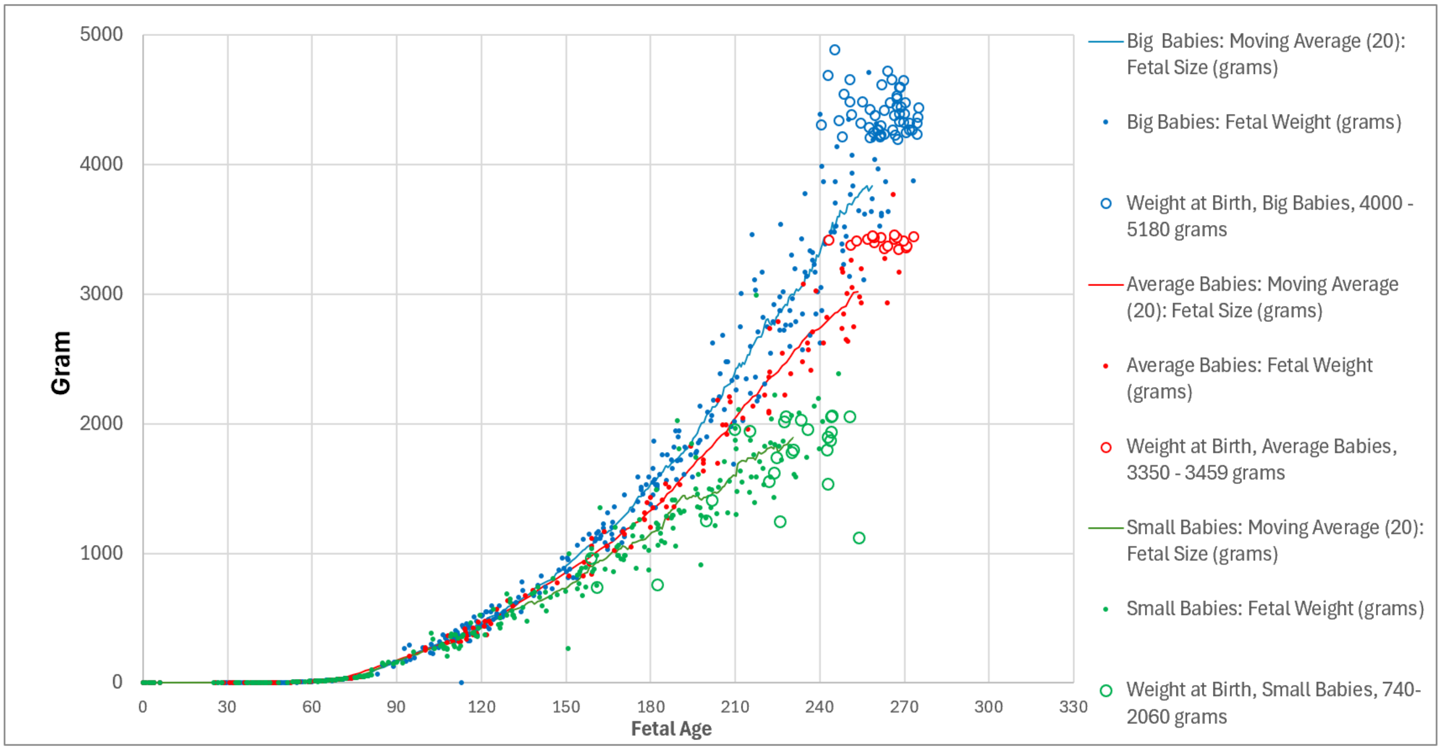


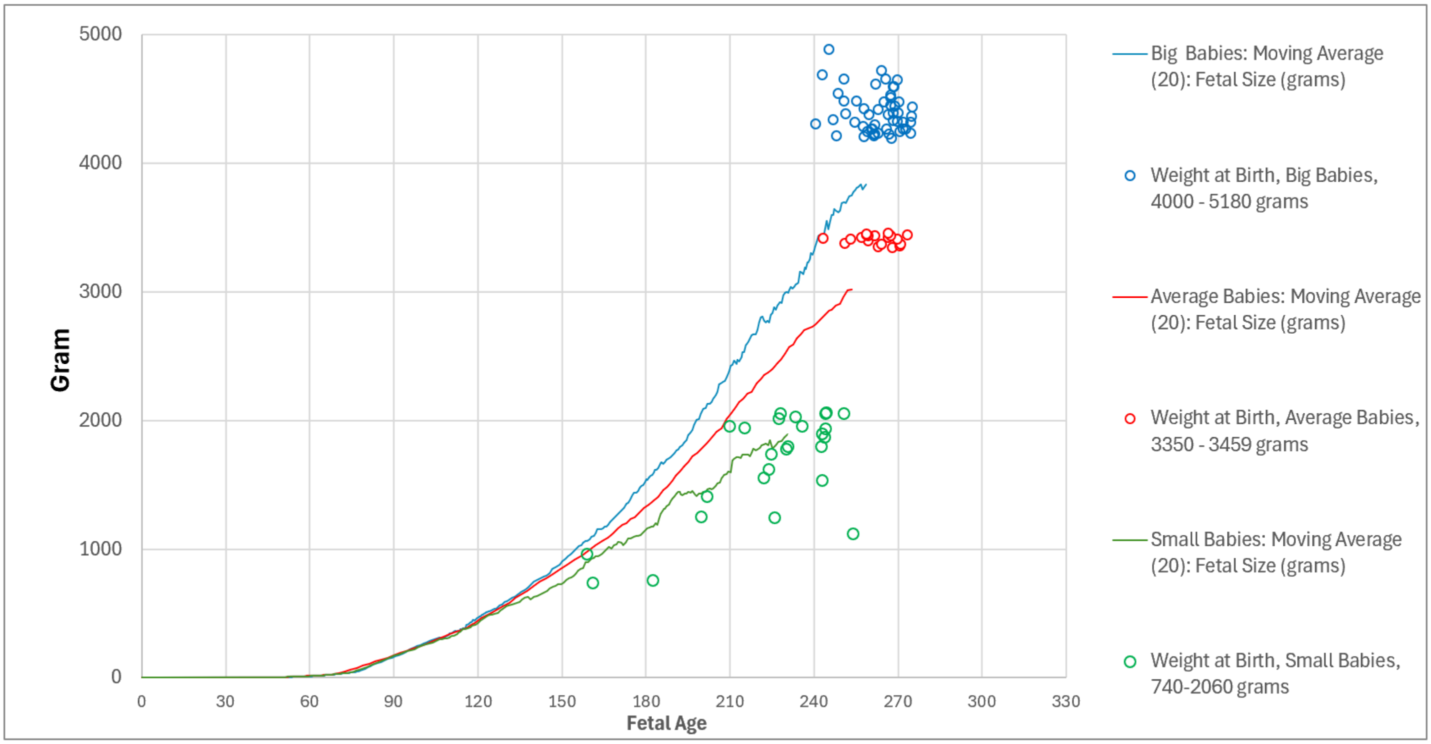


**Supplement-Figure-III.26**. Relationship between ***Fetal Weights*** (grams) of each ultrasound measurement (closed circles above, and running averages, above and below), calculated by the ***Abdominal Circumference Binary Cellular Estimated Fetal Weight Equation*** (#13), and ***Birthweights*** (open circles) by displaying all of these values on a ***Binary Cellular Fetal Growth Chart***.

**_Binary Cellular Fetal Growth Analysis_** Reveals Why Fetuses Become Babies with Bad **_Birthweights_**

By removing the individual ultrasound datapoints and showing just the ***Moving Average*** lines that capture this ultrasound data, together with the ***Birthweights*** of each of the babies in the three ***Birthweight*** groups, we can begin to get a picture of the combined effect of the ***Fetal Weight***, the ***Rate of Growth***, and ***Pregnancy Length***, on ***Birthweight*** (**Supplement-Figure-III.27**). The "***Big Baby Study Group***" had pregnancies of almost the same length as the "***Average Size Baby Study Group***". (**TABLE III.2**). The ***Big Babies*** were born big because they grew rapidly. In contrast, the babies in the "***Small Baby Study Group***" both grew more slowly than the ***Average Size Babies***, but also had pregnancies that were shorter (**TABLE III.2**). ***Small Babies*** were born ***Small*** partially because they grew too slowly, but mainly because they were born too soon (**Supplement-Figure-III.27**).

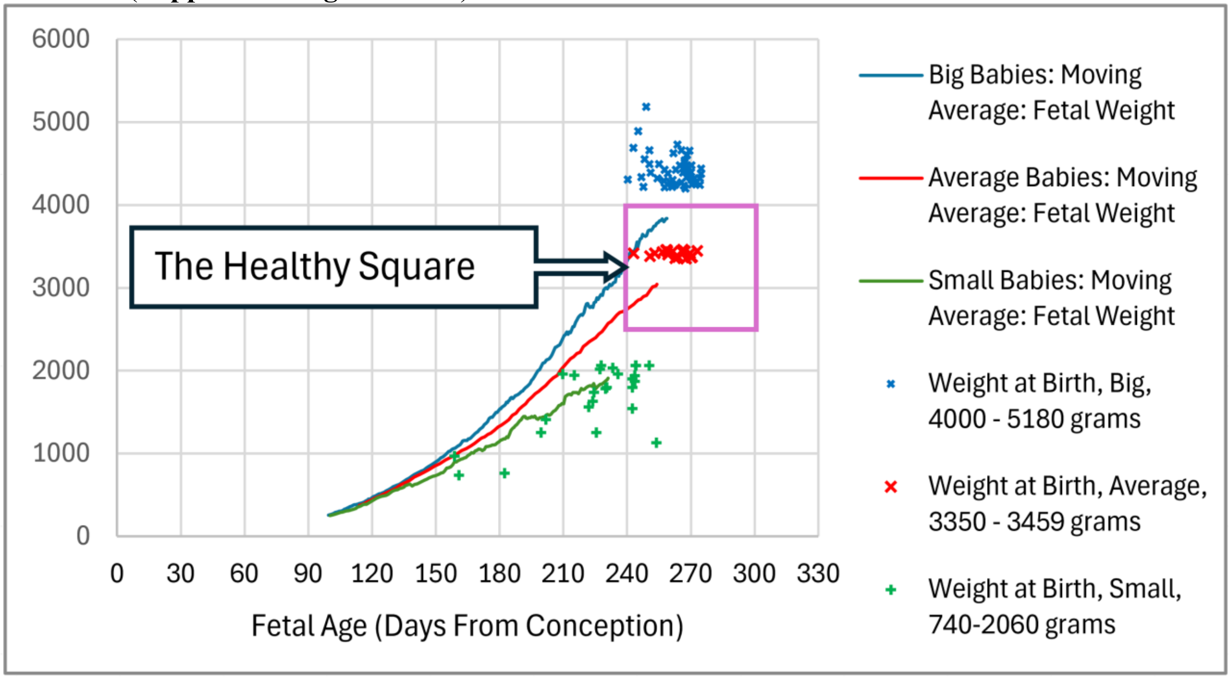


**Supplement-Figure-III.27**. Combined effect of the ***Speed of Growth***, and the ***Pregnancy Length***, on ***Birthweight*** "***Average Size Baby Study Group***", "***Small Baby Study Group***", and "***Big Baby Study Group***"

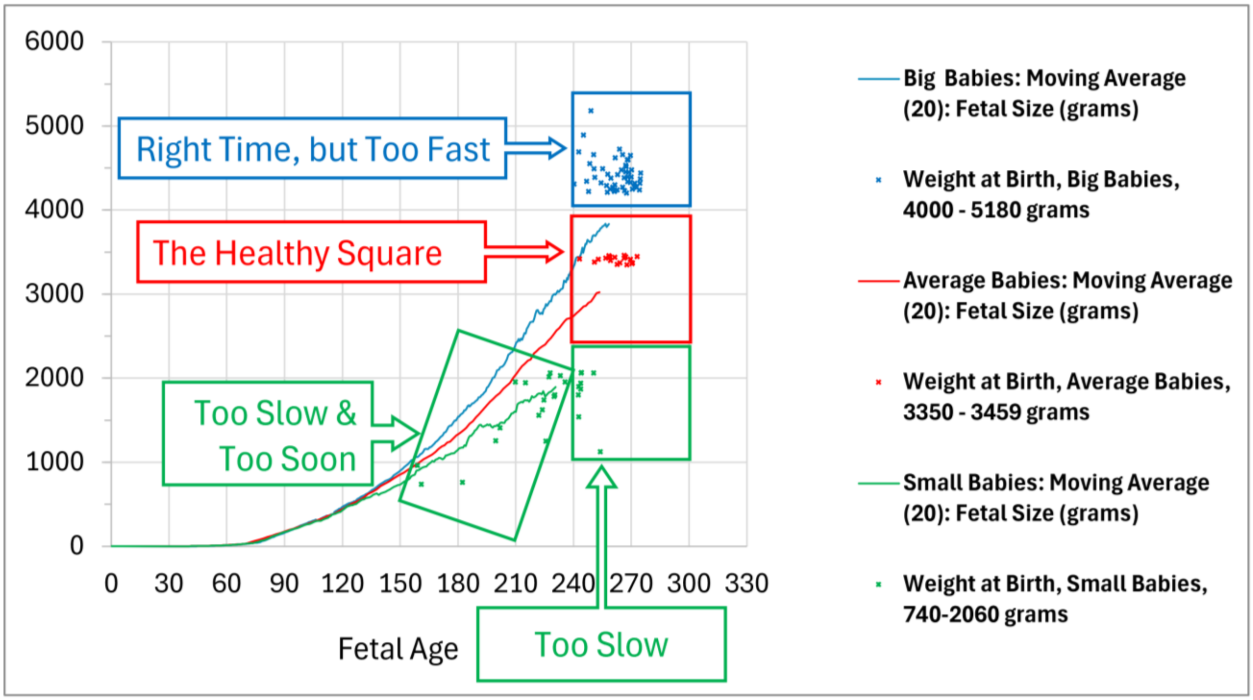


**Supplement-Figure-III.28**. Isolating the effects of the ***Speed of Growth***, and the ***Pregnancy Length***, on ***Birthweight*** "***Average Size Baby Study Group***", "***Small Baby Study Group***", and "***Big Baby Study Group***"

***Binary Cellular Fetal Growth Analysis*** Shows What Would Have Been If All Pregnancies Had Same Length.

We can begin to tease out the impact of the ***Rate of Growth***, and ***Pregnancy Length***, by calculating what would have been the ***Birthweights*** if they had all been born after the same period of time. We chose 268 days, as that was the average Pregnancy Length in the ***"627" dataset*** (**Supplement-Figure-III.1**).

We can easily make such calculations because our data show that in the last months of pregnancy, the fetuses that became the babies in our "***Big Baby Study Group"*** grew at ~31 grams/day (**Supplement-Figure-III.5**, **TABLE III.1**). The fetuses that became the babies in our "***Average Size Baby Study Group"*** grew at ~24 grams/day. The fetuses that became the babies in our "***Small Baby Study Group"*** grew at ~14 grams/day.

When we use these ***Weight-Gain Rate*** values, we can see that if all of the 30 ***Big Babies*** in our dataset ("***Big Baby Study Group***") were born after 268 days, there would have been even more babies over 5000 grams, and generally many would have been born even higher above the ***Healthy Square*** (**Supplement-Figures III.29 and 30**). We can also see that if all of the 15 ***Average Size Babies*** ("***Average Size Baby Study Group"***) in our dataset were born after 268 days, they all would still have been born comfortably within the ***Healthy Square***. Finally, we can see that if all of the 23 ***Small Babies*** ("***Small Baby Study Group"***) in our dataset were born at 268 days, many would have been born comfortably within the ***Healthy Square***, and most would have been born close to it.

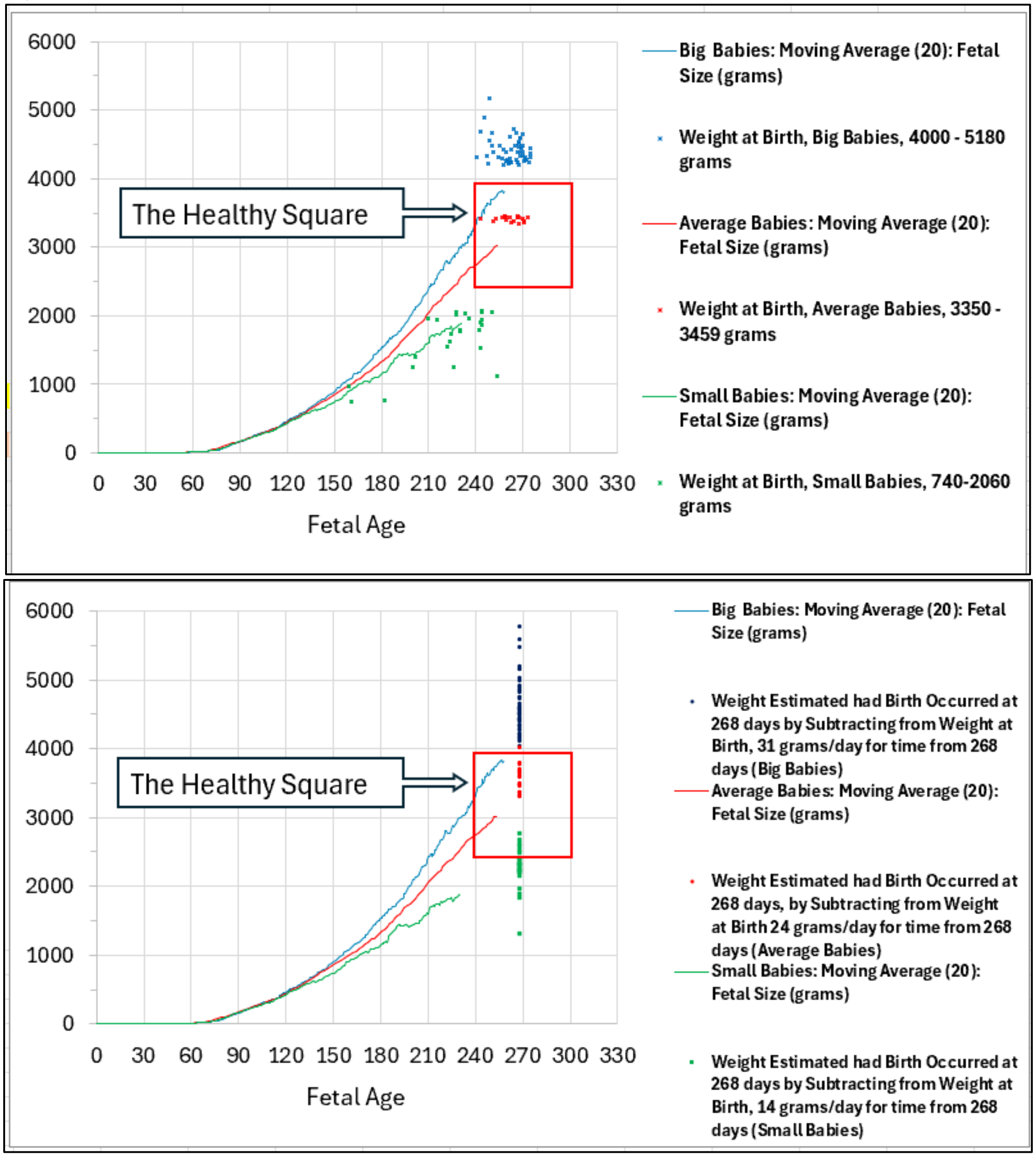


**Supplement-Figure-III.29**. Visualization of what would have been the ***Birthweights*** if all pregnancies had the same lengths: see text for description of calculations.

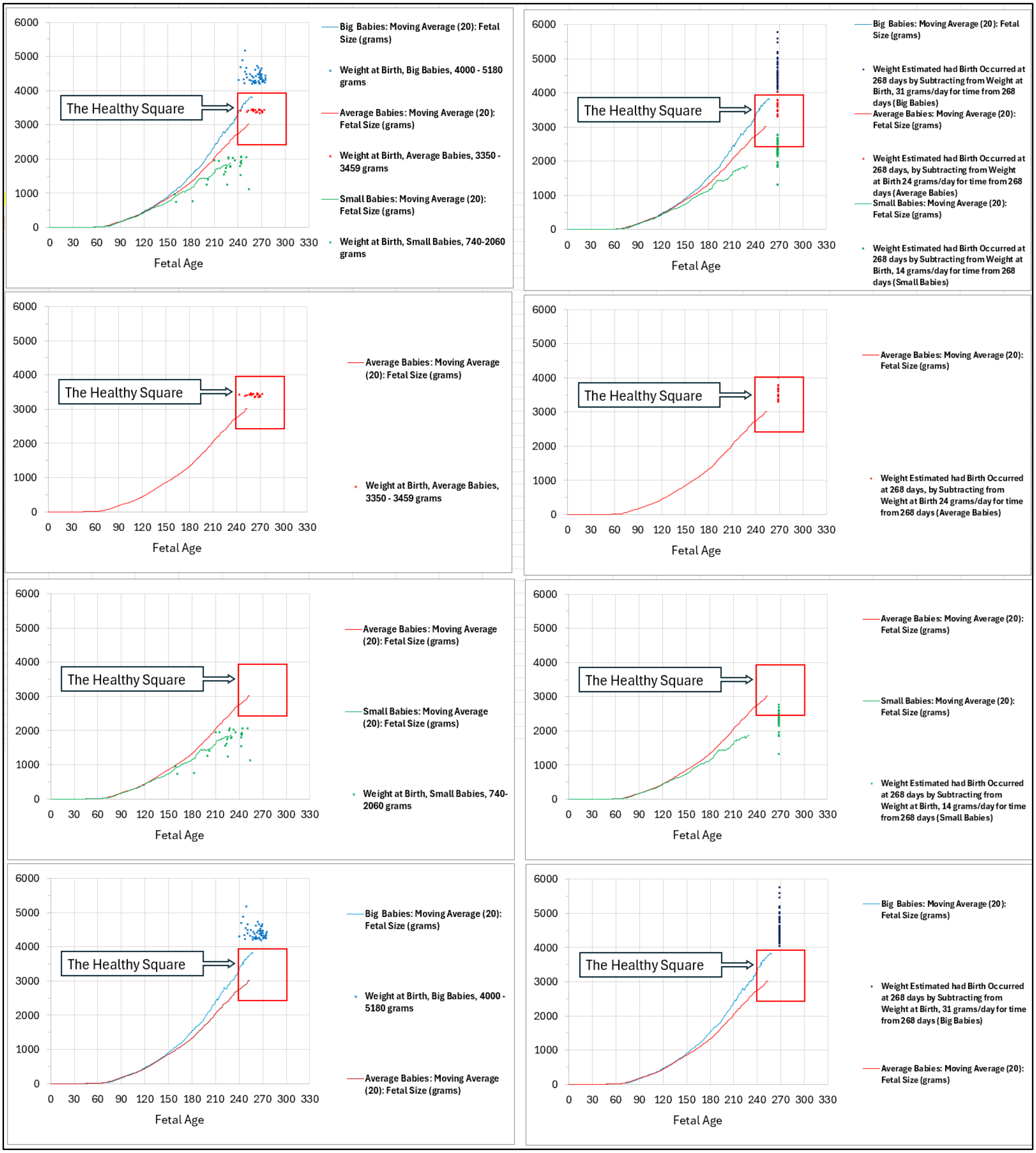


**Supplement-Figure-III.30** Visualization of what would have been the ***Birthweights*** if all pregnancies had the same lengths: see text for description of calculations.

***Binary Cellular Fetal Growth Analysis*** of ***All Babies*** in our Dataset

Above, we have examined in detail the rather small set of the 68 babies in the "***Big***", "***Average***", and "***Small*** ***Baby Study Groups***. Let us now expand that view, by considering all 663 babies in our ***Combined Dataset***, for which we have CRL measurements that allow us to measure the length of each pregnancy by the ***CRL Fetal Age Equation*** (#7a) (**Supplement-Figure-III.31**). With this information, we can display the ***Birthweights*** for each of the 663 babies on ***Binary Cellular Fetal Growth Charts***, to tease out the ***Fetal Weight***, the ***Rate of Growth***, and ***Pregnancy Length***, for each baby.

***Binary Cellular Fetal Growth Analysis*** Gives Biologically, Data-Driven, ***Fetal Growth Curves***.

As we have seen above, ***Fast***, ***Average***, and ***Slow*** growing fetuses, that become ***Large***, ***Average***, and ***Small Newborns***, differ in the value of the ***"a" Parameter*** of the ***Binary Cellular Universal Growth Equation***. This affords us the opportunity to create biologically, data-driven, ***Fetal Growth Curves*** (**Supplement-Figure-III.31**). The ***"a" Parameter*** values shown in **Supplement-Figure-III.31** were taken from ***Average***, ***Small***, and ***Big Baby Study Groups*** (**Supplement-Figures III.21-25**), plus values above, below, and in between, but, of course, one can generate curves with whichever ***"a" Parameter*** value would be informative.

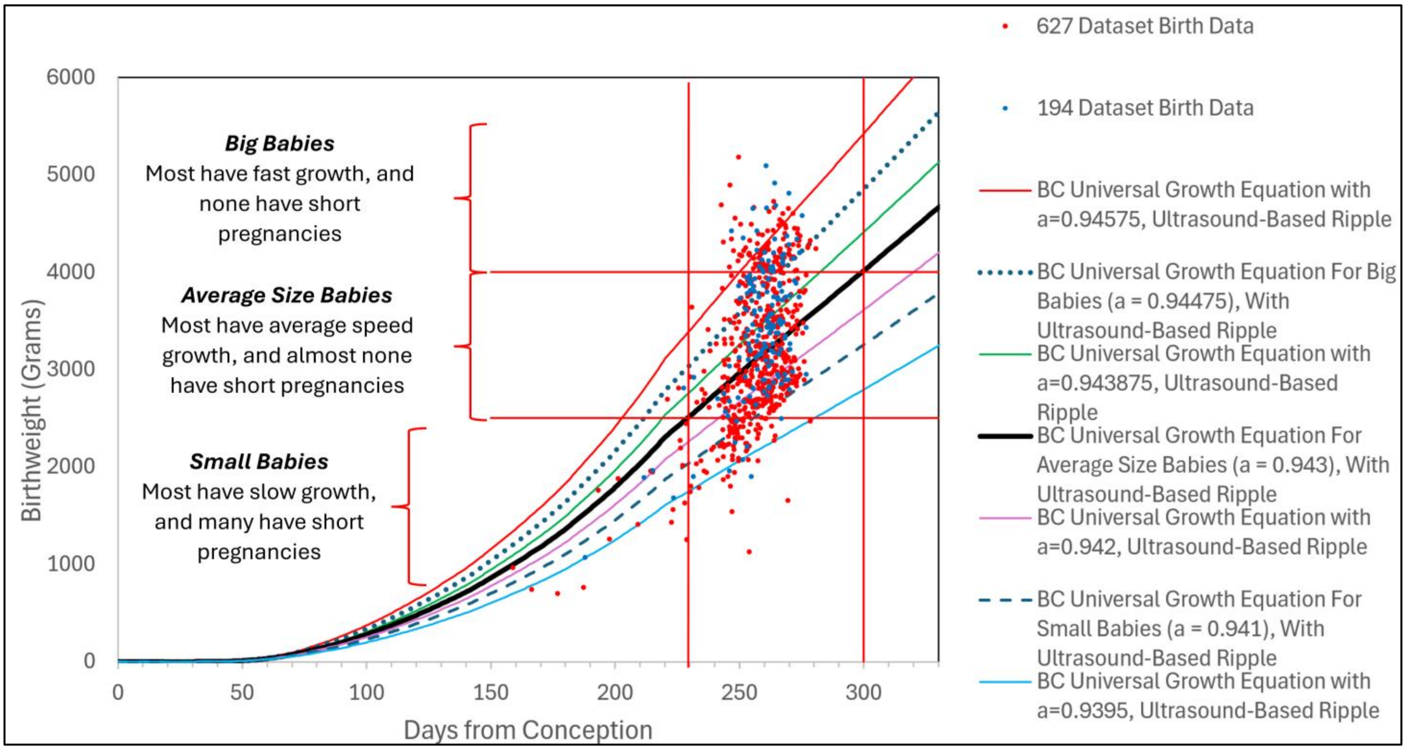


**Supplement-Figure- III.31**. ***Fetal Growth Curves***, differing in the ***"a" Parameter*** of the ***Binary Cellular Universal Growth Equation***. The ***"a" Parameter*** values shown in this figure were taken from ***Average***, ***Small***, and ***Big Baby Groups***, plus values above, below, and in between those values

***Fetal Growth Curves*** formulated with various values of the ***"a" Parameter*** of the ***Binary Cellular Universal Growth Equation***, such as those shown in **Supplement-Figure-III.31**, can tell us a great deal about ***Fetal Growth***. For example, such ***Fetal Growth Curves*** can make it possible to examine the relationship between the ***"a" Parameter*** values for individual fetuses, determined from ultrasound measurements with ***Binary Cellular Estimated Fetal Weight Equations***, and the ***"a" Parameter*** values determined from ***Birthweight***. Similarly, such ***Fetal Growth Curves*** can tell us how constant the ***"a" Parameter*** value is for individual fetuses, as can be determined from multiple ultrasound measurements. It will also be possible to identify curves with ***"a" Parameter*** values that map to various centiles (3rd, 50th, and 97th percentiles are common examples); the reverse calculation is also possible, that is to say, to back calculate the percentile to which each fetus belongs. These studies are underway and will be reported later. However, let us first examine what these ***Fetal Growth Curves*** can tell us about the relationship between ***Rate of Growth***, ***Pregnancy Length***, and ***Birthweight***.

***Binary Cellular Fetal Growth Analysis*** Gives Independent Measures of ***Rate of Growth*** & ***Pregnancy Length***.

In **Supplement-Figure-III.32a**, we have shown the ***Birthweights*** of each of these 663 babies in our ***Combined Dataset***, together with ***Binary Cellular Universal Growth Equations*** with various ***"a" Parameter*** values. Note that such a ***Binary Cellular Fetal Growth Analysis*** lets us see, for each of the 663 babies, what its ***Pregnancy Length*** was, and what its ***Rate of Growth*** was. The ***Pregnancy Length*** can be known by the location of each ***Birthweight*** on the X-axis. The ***Rate of Growth*** can be known by which of the various ***Binary Cellular Universal Growth Equations***, each with different ***"a" Parameter*** value, the ***Birthweight*** value sits on. Note that this way to measure the ***Rate of Growth*** is independent of when the baby was born.

A useful, if somewhat arbitrary, way to talk about the ***Pregnancy Length*** is to consider whether a ***Pregnancy*** is shorter, or longer, than the left-hand boundary of ***Healthy Square***, which is 230 days after conception. With these measures, we can begin to use ***Binary Cellular Fetal Growth Analysis*** to decipher what makes some babies ***Big*** at birth, other babies ***Average***, and others ***Small***.

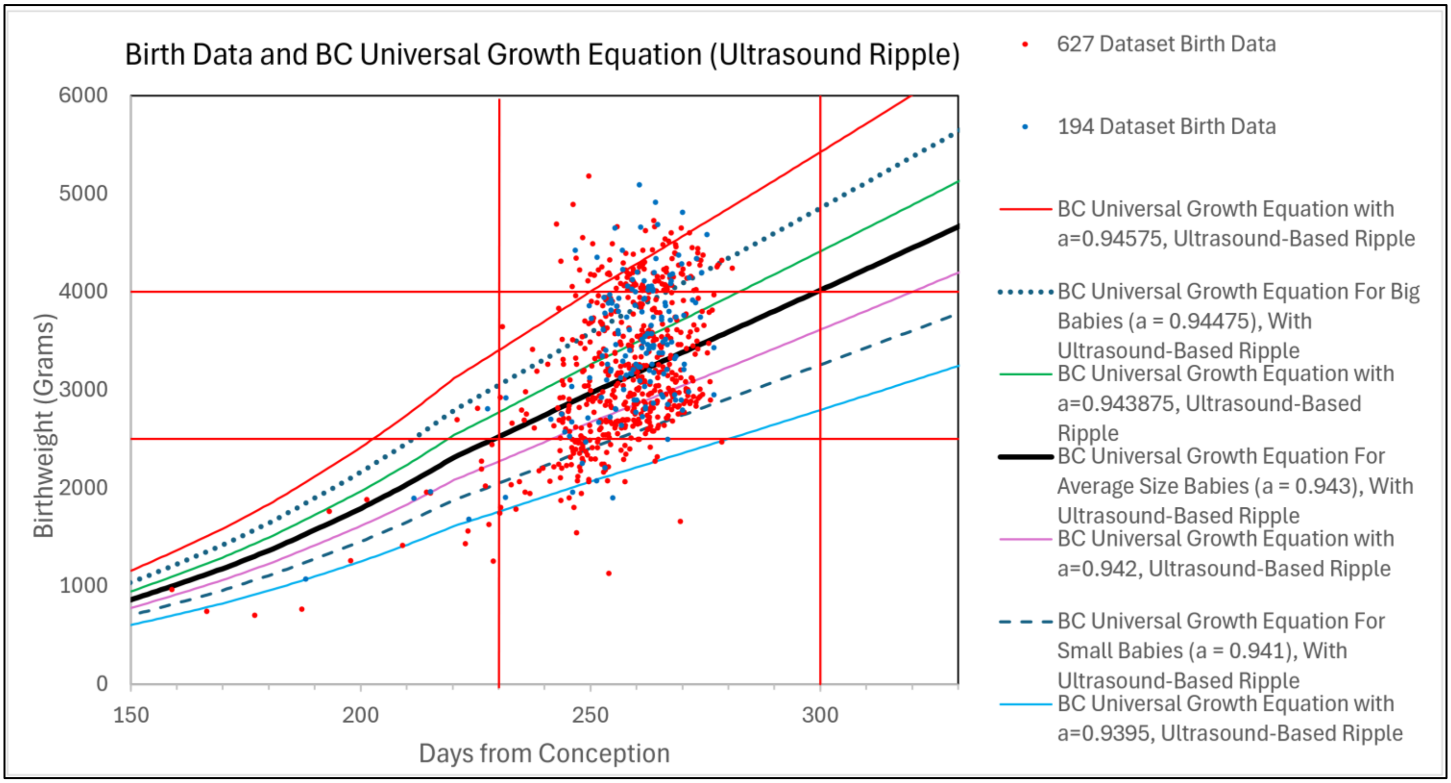


**Supplement-Figure-III.32a**. The ***"a" Parameter*** values shown in this figure were taken from "***Average***, ***Small***, and ***Big Baby Study Groups***", and values above, below, and in between, but we can, of course, generate curves with any ***"a" Parameter*** value we wish.

***Binary Cellular Fetal Growth Analysis*** of ***Categories of Fetal Growth*** of ***Big***, ***Average***, and ***Small Newborns***

While the results of the ***Binary Cellular Fetal Growth Analysis*** of the patient dataset we have examined here reveals that the ***Speed of Growth*** and the ***Pregnancy Length*** are continuous qualities, one can capture some of the general principles by lumping babies into various categories of fetal growth. Let us next use ***Binary Cellular Fetal Growth Analysis*** to examine the growth of fetuses that become ***Big***, ***Average***, and ***Small Newborns***.

***Big Babies*** Become ***Big*** by ***Fast Growth*** with ***Normal Length Pregnancies***.

Let us begin by examining the ***Big Babies***, Macroscopic Babies, conventionally defined as those newborns more than 4000 grams (**Supplement-Figure-III.32b**). Note that all of these ***Big Babies*** had resided alongside ***Binary Cellular Universal Growth Equations*** with values of the ***"a" Parameter*** indicative of ***Fast Growth***. Note also that all of these ***Big Babies*** had ***Pregnancy Lengths*** of 230 days or more after conception, that is to say, had ***Normal Length Pregnancies***. Thus, the data show that ***Big Babies*** become ***Big*** by ***Fast Growth*** with ***Normal Length Pregnancies***.

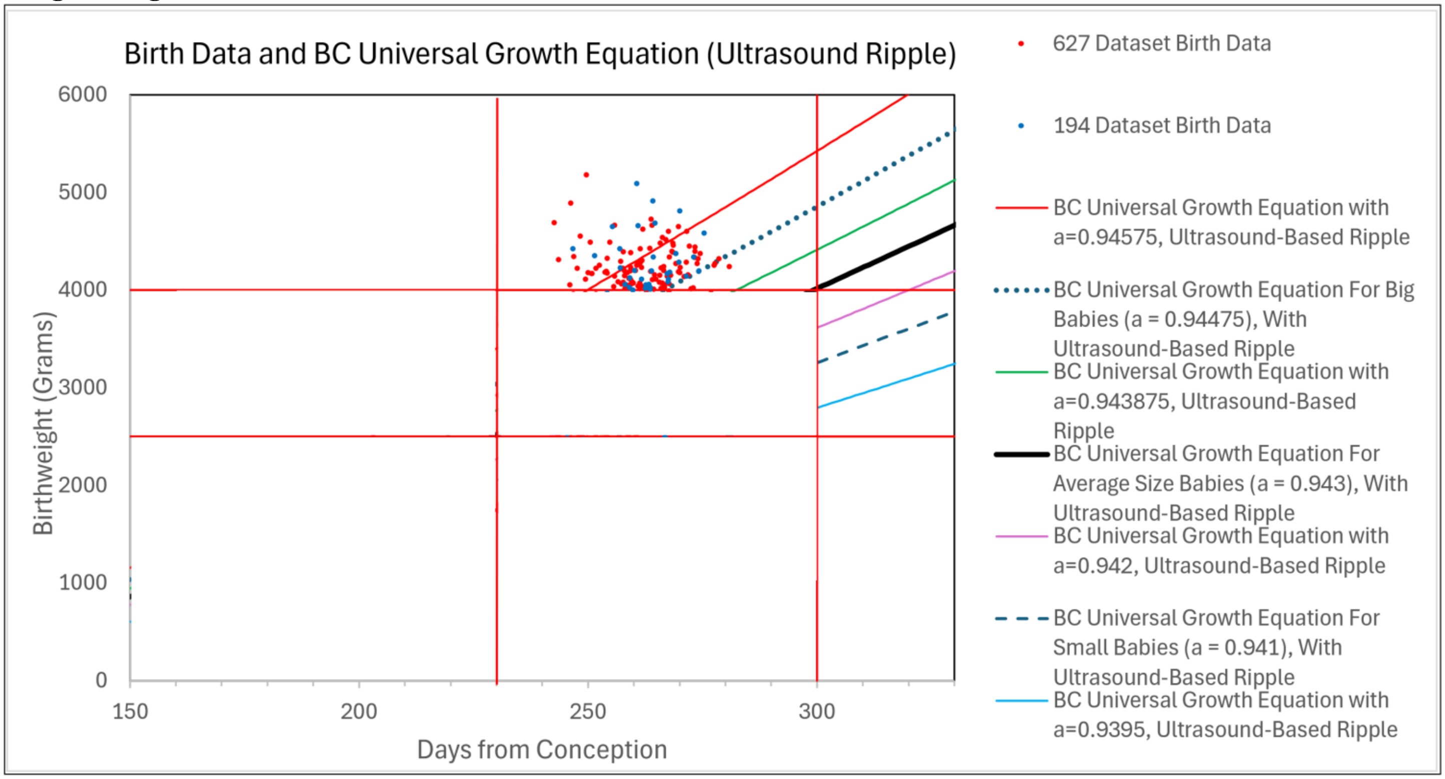


**Supplement-Figure-III.32b**. ***Big Babies***, Macroscopic Babies, conventionally defined as those newborns more than 4000 grams. The ***"a" Parameter*** values shown in this figure were taken from "***Average***, ***Small***, and ***Big Baby Study Groups***", and values above, below, and in between, but we can, of course, generate curves with any ***"a" Parameter*** value we wish.

<u>Most ***Average Size Babies*** Become ***Average Size*** by ***Average Growth*** with ***Normal Length Pregnancies***.</u>

Let us next examine the ***Average Size Babies***, conventionally defined as those newborns weighing between 2500 and 4000 grams (**Supplement-Figure-III.32c**). Note that almost all of these ***Average Size Babies*** had resided alongside ***Binary Cellular Universal Growth Equations*** with values of the ***"a" Parameter*** indicative of ***Average Growth***. Note also that all but three of these ***Big Babies*** had ***Pregnancy Lengths*** of 230 days or more after conception, that is to say, had ***Normal Length Pregnancies***. Thus, the data show that almost all ***Average Size Babies*** become ***Average Size*** by ***Average Growth*** with ***Normal Length Pregnancies***.

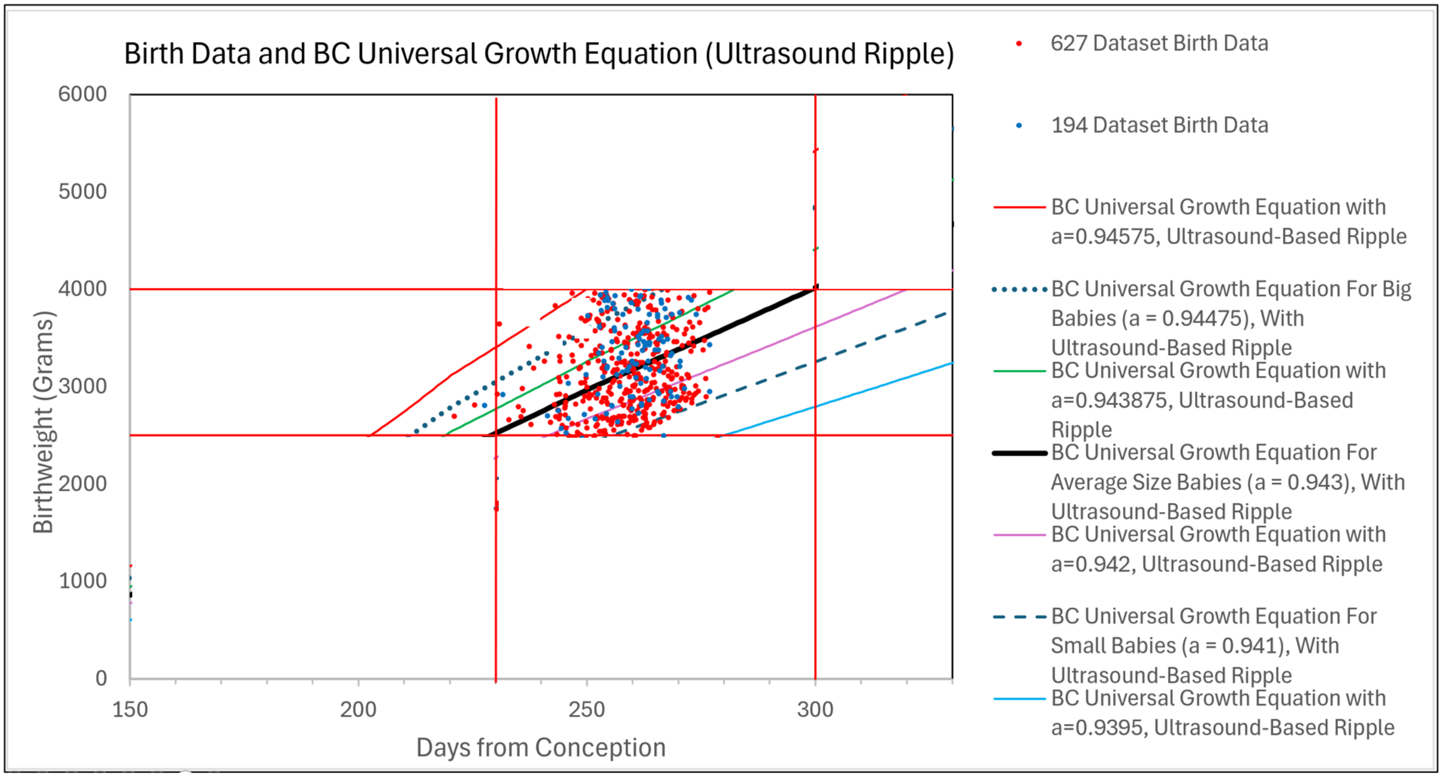


**Supplement-Figure-III.32c**. The ***"a" Parameter*** values shown in this figure were taken from "***Average***, ***Small***, and ***Big Baby Study Groups***", and values above, below, and in between, but we can, of course, generate curves with any ***"a" Parameter*** value we wish.

***Small Babies*** Become ***Small*** by ***Slow Growth*** and ***Short Pregnancies***.

Let us next examine the ***Small Babies***, conventionally defined as those newborns weighing less than 2500 grams (**Supplement-Figure-III.32d**). Note that almost all of these ***Small Babies*** had resided alongside ***Binary Cellular Universal Growth Equations*** with values of the ***"a" Parameter*** indicative of ***Slow Growth***. Not also that quite a few of these ***Small Babies*** had ***Pregnancy Lengths*** of less than 230 days after conception, that is to say, had ***Short Pregnancies***. Thus, the data show that ***Small Babies*** become ***Small*** by having either ***Slow Growth***, or ***Short Pregnancies***, or both.

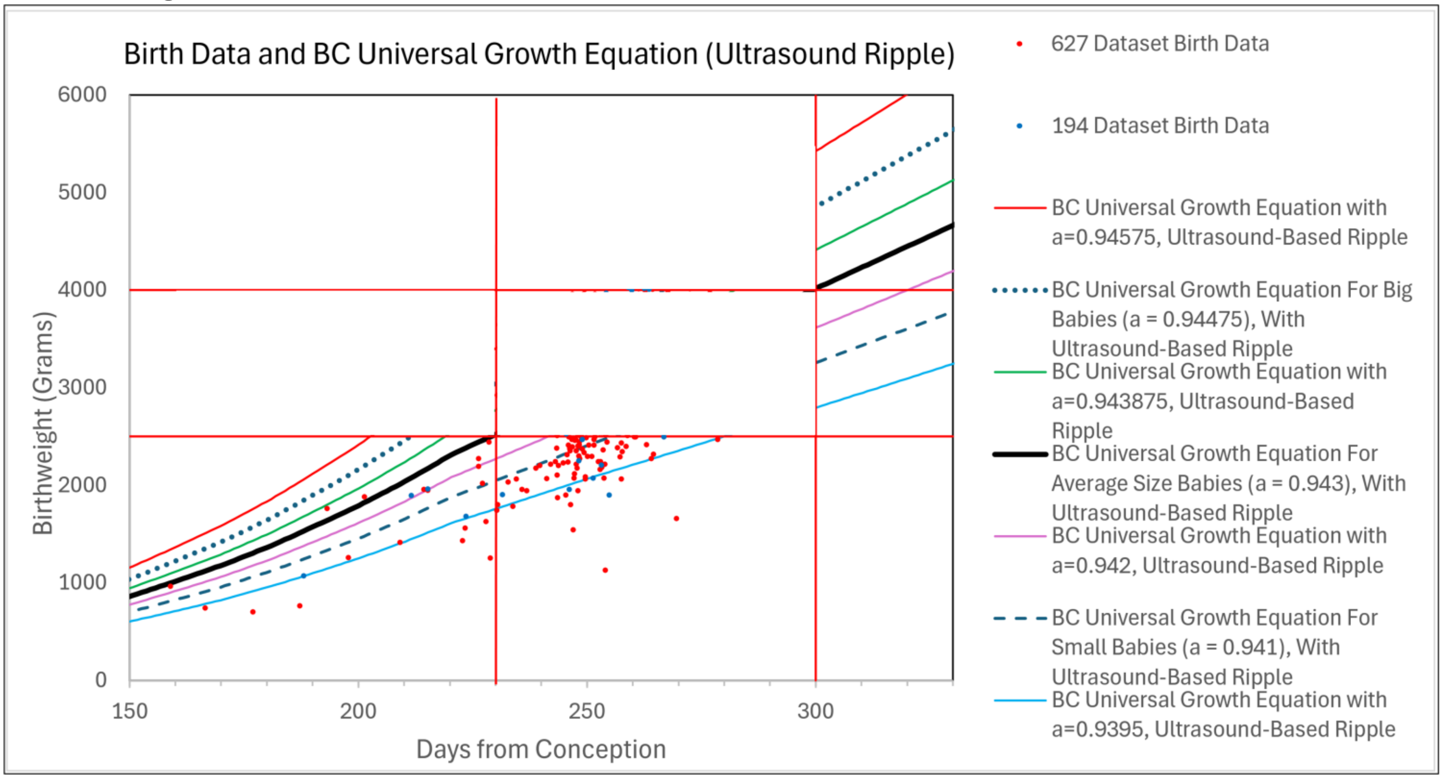


**Supplement-Figure-III.32d**. The ***"a" Parameter*** values shown in this figure were taken from "***Average***, ***Small***, and ***Big Baby Study Groups***", and values above, below, and in between, but we can, of course, generate curves with any ***"a" Parameter*** value we wish.

***<u>Small Birthweights</u>*** <u>Have</u> ***<u>Short Pregnancies</u>***<u>;</u> ***<u>Average & Large Birthweights</u>*** <u>Have</u> ***<u>Same Length Pregnancies</u>***

In **SUPPLEMENT-TABLE III.3** and **Supplement-Figures III.33** are shown data from the "***627" dataset*** on babies sorted into groups by ***Birthweight***, together with the ***Pregnancy Length*** of each group. Babies in the 700-999 grams ***Birthweight*** group were born after an average ***Pregnancy Length*** of 172 days. Babies in the 1000-1499 grams ***Birthweight*** group were born after an average ***Pregnancy Length*** of 223 days. Babies in the 1500-1999 grams ***Birthweight*** group were born after an average ***Pregnancy Length*** of 232 days. Babies in the 2000-2499 grams ***Birthweight*** group were born after an average ***Pregnancy Length*** of 249 days. Babies in all of the ***Birthweight*** groups above 2500 grams (~85% of the babies in this dataset) were born after similar average ***Pregnancy Lengths*** of 257-263 days.

Thus, for the most part, babies of ***Small Birthweight*** tended to have ***Pregnancies*** of ***Short Length***. In contrast, babies of ***Average*** or ***Big Birthweights*** tended to have ***Pregnancies*** of roughly ***Similar***, ***Average***, ***Length***.

Babies of the ***Smallest Birthweights*** tended to have ***Pregnancy Lengths*** that increased with ***Birthweight*** until about 2500 grams (700-999, 1000-1499, 1500-1999, 2000-2499 grams: 172, 223, 232, 249 days). However, babies with ***Average*** or ***Big Birthweights*** of 2500 grams and more tended to have ***Pregnancies*** of about the ***Same Normal Length*** (2500-2999, 3000-3499, 3500-3999, 4000-4999, 4500-4599 grams; 257, 260, 260, 263, 260 days)

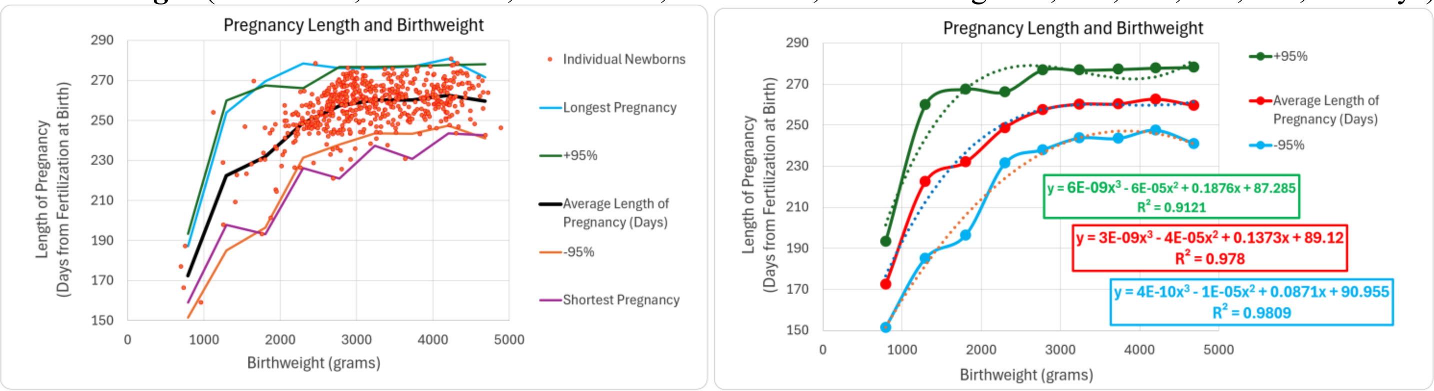


**Supplement-Figure-III.33**

| Birthweight Range | Average Birthweight | Number | Average Pregnancy Length (Days) | Shortest Pregnancy | Longest Pregnancy | SD | -95% | +95% |
|---|---|---|---|---|---|---|---|---|
| 700-999 | 791 | 4 | 172 | 159 | 187 | 11 | 152 | 193 |
| 1000-1499 | 1294 | 5 | 223 | 198 | 254 | 19 | 185 | 260 |
| 1500-1999 | 1803 | 17 | 232 | 193 | 270 | 18 | 196 | 268 |
| 2000-2499 | 2299 | 68 | 249 | 226 | 279 | 9 | 231 | 266 |
| 2500-2999 | 2776 | 174 | 257 | 221 | 276 | 10 | 238 | 277 |
| 3000-3499 | 3239 | 124 | 260 | 237 | 276 | 8 | 244 | 277 |
| 3500-3999 | 3728 | 89 | 260 | 231 | 277 | 9 | 243 | 277 |
| 4000-4999 | 4201 | 98 | 263 | 244 | 281 | 8 | 247 | 278 |
| 4500-4599 | 4683 | 13 | 260 | 243 | 272 | 9 | 241 | 278 |

**SUPPLEMENT-TABLE III.4**

These babies were counted by examining the dots on the figures shown above. Because of overlap of these dots, values shown above undercount these babies. Computationally-driven counts will refine these values, but the general impressions from such data are unlikely to change greatly.

***Short Pregnancy*** is Correlated with ***Slow Growth***; ***Average Length*** is Correlated with ***Average& Fast Growth***.

In **SUPPLEMENT-TABLE III.4** and **Supplement-Figure-III.34** are shown data from the "*627" **dataset*** on babies sorted into groups by ***Pregnancy Length***, together with the ***Birthweight*** of each group. To do so, we rely on World Health Organization and ***Healthy Square*** categories. The WHO divides preterm babies into three groups: ***Extremely Preterm*** (<28 weeks of gestation=180 days since conception); ***Very Preterm*** (28–32 weeks of gestation =180->210 days since conception); and ***Moderate or Late Preterm*** (>32 weeks of gestation=210 days since conception). **Supplement-Figures III.876** and **999** also shows the latest ***Binary Cellular Universal Growth Equation*** with the refined Ripple defined by ultrasound measurements, rather than autopsy data, which slightly revises the left size of the ***Healthy Square*** as beginning at 230 days rather than 240. Thus, we sorted babies into five groups: ***Extremely Preterm*** Babies Born at less than 180 days after conception; ***Very Preterm*** Babies Born at 180-210 days after conception; ***Moderate Preterm***, Babies Born to the ***Left of the Healthy Square*** at 210-230 days after conception; ***Moderate Preterm***, Babies Born to the ***Right of the Healthy Square*** at 210-230 days after conception; ***Normal*** Babies Born at 245+ days after conception. We then sorted these five ***Pregnancy Length*** groups to into three ***Speed of Growth*** groups, differing by the ***"a" Parameter*** of the ***Binary Cellular Universal Growth Equation*** for fetuses that become the newborns in the ***Big***, ***Average*** and ***Small Baby*** groups: Babies that arose from fast-growing fetuses, ***a***≈0.945; Babies that arose from average-growing fetuses, ***a***≈0.943; Babies that arose from slow-growing fetuses, ***a***≈0.941.

For ***Extremely*** and ***Very Preterm*** Babies Born less than 210 days after conception, none arose from ***Fast-Growing*** Fetuses (***a***≈0.945), 33% (3 babies) arose from ***Average-Growing Fetuses*** (***a***≈0.943), and 66% (6 babies) arose from ***Slow-Growing Fetuses*** (***a***≈0.941). For ***Moderate Preterm*** babies born before the ***Healthy Square*** at 210-230 days after conception, 20% (3) arose from ***Fast-Growing*** Fetuses (***a***≈0.945), 40% (6 babies) arose from ***Average-Growing Fetuses*** (***a***≈0.943), and 40% (6 babies) arose from ***Slow-Growing Fetuses*** (***a***≈0.941). For ***Moderate Preterm*** babies born within the ***Healthy Square*** at 230-244 days after conception, 14% (9) arose from ***Fast-Growing*** Fetuses (***a***≈0.945), 49% (32 babies) arose from ***Average-Growing Fetuses*** (***a***≈0.943), and 37% (24 babies) arose from ***Slow-Growing Fetuses*** (***a***≈0.941). For ***Normal Term*** babies born at 245 or more days after conception, 20% (3) arose from ***Fast-Growing*** Fetuses (***a***≈0.945), 49% (32 babies) arose from ***Average-Growing Fetuses*** (***a***≈0.943), and 37% (24 babies) arose from ***Slow-Growing Fetuses*** (***a***≈0.941).

| ***Pregnancy Length Group*** ➡<br>***Speed of Growth Group*** by the ⬇ ***Binary Cellular Universal Growth Equation*** | Babies Born at less than 180 days after conception: ***Extremely Preterm*** | Babies Born at 180-210 days after conception: ***Very Preterm*** | Babies Born at 210-230 days after conception: ***Moderate Preterm***, before the ***Healthy Square*** | Babies Born at 230-244 days after conception: ***Moderate Preterm***, within the ***Healthy Square*** | Babies Born at 245+ days after conception |
|---|---|---|---|---|---|
| Babies That Arose from Fast-Growing Fetuses, ***a***≈0.945 | 0% (0) | 0% (0) | 20% (3) | 14% (9) | 48% (228) |
| Babies That Arose from Average-Growing Fetuses, ***a***≈0.943 | 33% (1) | 33% (2) | 40% (6) | 49% (32) | 33% (156) |
| Babies That Arose from Slow-Growing Fetuses, ***a***≈0.941 | 67% (2) | 67% (4) | 40% (6) | 37% (24) | 19% (93) |

**SUPPLEMENT-TABLE III.4**

Thus, babies with the shortest ***Extreme*** or ***Very Preterm Pregnancy Lengths*** (fewer than 210 days) tended to arise from fetuses that were ***Growing*** at the ***Slowest Rate***. In contrast, babies with the longest, normal ***Pregnancy Lengths*** (more than 245 days) tended to arise from fetuses that were ***Growing*** at the ***Average Rate*** or ***Fast Rate***. In between, babies with the ***Moderate Preterm***, ***Pregnancy Lengths*** (210-245 days) tended to arise from fetuses that were ***Growing*** at the ***Average Rate***.

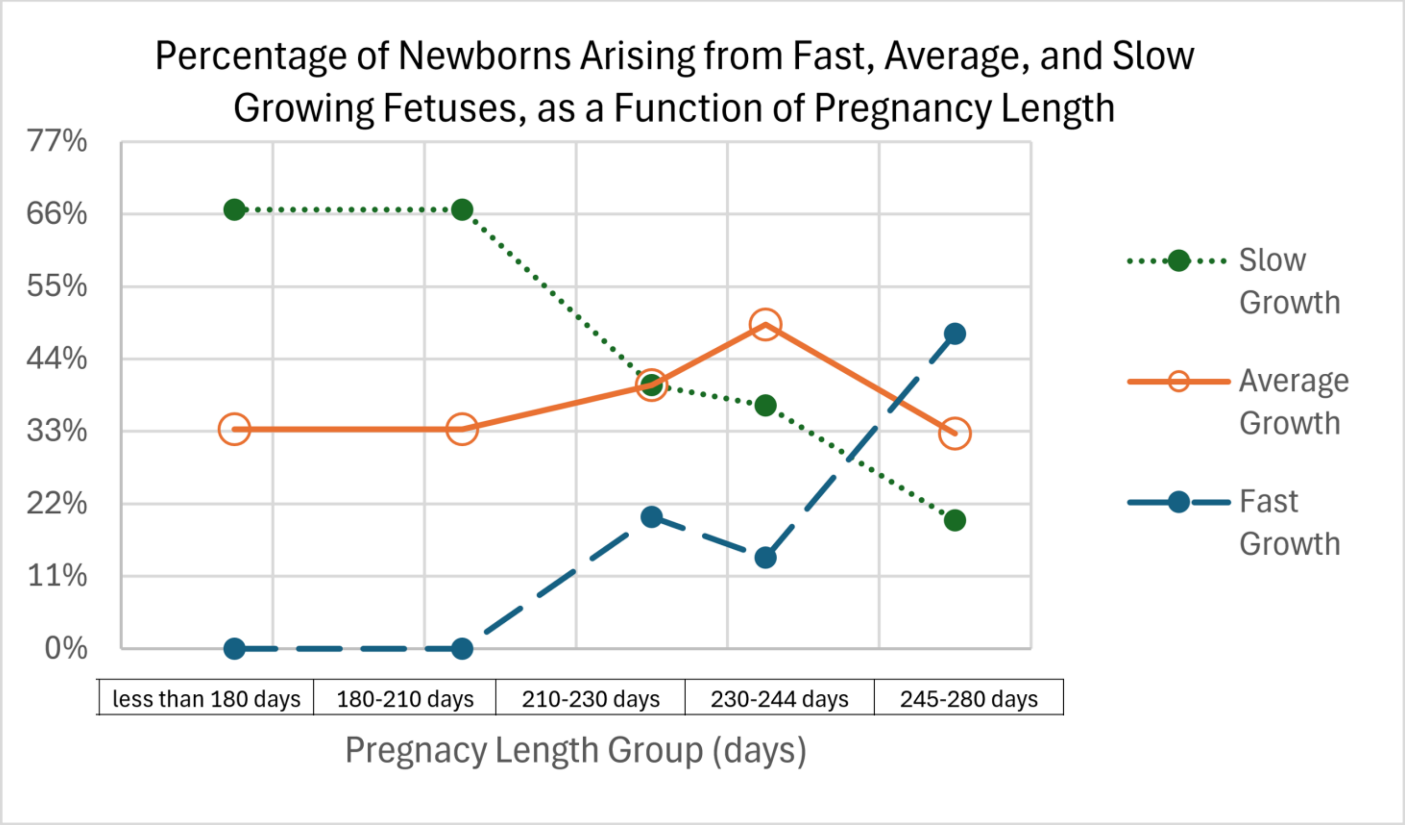


**Supplement-Figure-III.34**

<u>***Short Pregnancy*** Appears to Be ***Caused*** by ***Slow Growth***</u>

In **Figure 10**, we can see that about 50% of the babies in our dataset grew with a "***Fast***" ***"a" Parameter*** typical of the ***Big Baby Group*** (~0.9447), about 30% of the babies grew with an "***Average Speed***" ***"a" Parameter*** typical of the ***Average Size Baby Group*** (~0.943), ), and about 20% of the babies grew with a "***Slow***" ***"a" Parameter*** typical of the ***Small Baby Group*** (~0.943) (For the counts behind the numbers in this section, see the SUPPLEMENT). On the other hand, none (0%) of the babies with a ***Pregnancy Length*** of less than 220 days grew with a "***Fast***" ***"a" Parameter***, about one-third (33%) of the babies with a ***Pregnancy Length*** of less than 220 days grew with an "***Average***" ***"a" Parameter***, and about two-thirds (~66%) of the babies with a ***Pregnancy Length*** of less than 220 days grew with a "***Slow***" ***"a" Parameter***. Clearly, ***Slow Growth*** predisposes for ***Short Pregnancy***. That is, the numbers seem to be telling us that ***Short Pregnancy*** is ***Caused*** by ***Slow Growth***. In the DISCUSSION, we will consider some of the biological reasons for why this causal relationship between ***Slow Growth*** and ***Short Pregnancy*** may occur, and how this information offers some possibilities for preventing this from occurring.

The ***Healthy Square***, the ***Healthy Island***, the ***Healthiest Possible Birth***.

The ***Healthy Square*** defined by the ***Binary Cellular Universal Growth Equation*** (#2) captures in its borders the conventional measure of a fetus that is born ***Too Small*** (less than 2,500 grams) or born ***Too Large*** (more than 4000 grams). However, there is no reason why the ***Healthy Square*** couldn't be transformed into the ***Healthy Island***, whose meandering concentric topological borders could be used to define the risks for fetal and maternal morbidity and mortality.

Shown in **Supplement-Figure-III.35**, we can see the overlap of the ***Binary Cellular Fetal Growth Charts*** showing ***Birthweights*** from the "***627" and*** "***197" datasets*** with the ***Binary Cellular Universal Growth Equation*** (#2), rooted in the time of conception, as determined by the ***CRL Fetal Age Equation*** (#7a) (dark line) together with the ***Infant Mortality Rates*** from Solis et al[274] Note how the ***Infant Mortality Rate*** below 6 deaths per 1,000 births roughly overlaps with the ***Healthy Square***, and how the ***Infant Mortality Rate*** below 1.5 deaths per 1,000 births sits at the top of the ***Healthy Square***. One can easily envisage the practical application of this to help guide obstetricians in making the best decisions allowing pregnancies to end at the point where the smallest chance of ***Mortality*** might be.

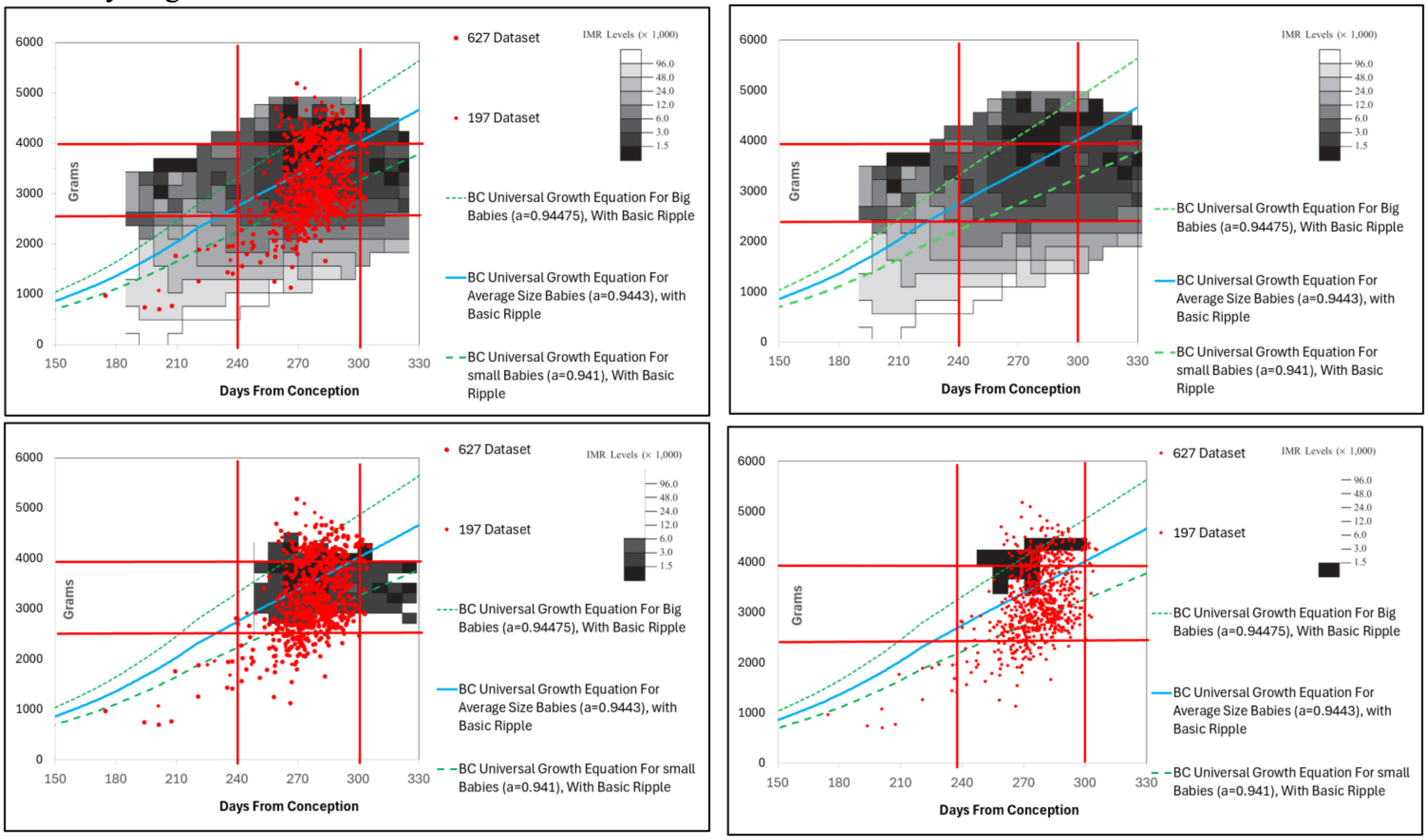


**Supplement-Figure-III.35** ***Binary Cellular Fetal Growth Charts*** showing only ***Birthweights*** from the "***194" and*** "***197" datasets***. Dark line is the ***Binary Cellular Universal Growth Equation*** (#2), rooted in the time of conception, as determined by the ***CRL Fetal Age Equation*** (#7a). Infant Mortality Rates (IMR) from Solis et al[274]

Not only does the concept of the ***Healthy Island*** offer the opportunity to provide guidance to obstetricians in their work of achieving the best outcome for babies and their mothers, the underlying data is transparent, and thus capable of continuous improvement and application to different mothers, and different medical professionals, in different populations, in different socioeconomic groups, in different parts of the world. The general calculations made possible by this approach should also be an aid for policy makers in their efforts to identify the best medical guidelines and organizational choices for achieving the best outcomes at the lowest possible costs.

In **Supplement-Figure-III.36** is shown a graph ***Binary Cellular Universal Growth Equations for Small, Average***, and ***Big Babies***, together with the ***Large, Appropriate***, and ***Small for Gestational Age*** categories created by Battaglia & Lubchenco (1967)[275]. Thus, we can see how ***Binary Cellular Fetal Growth Analysis*** can aid in refining the methods and goals of this admirable approach to improving the outcome of pregnancy.

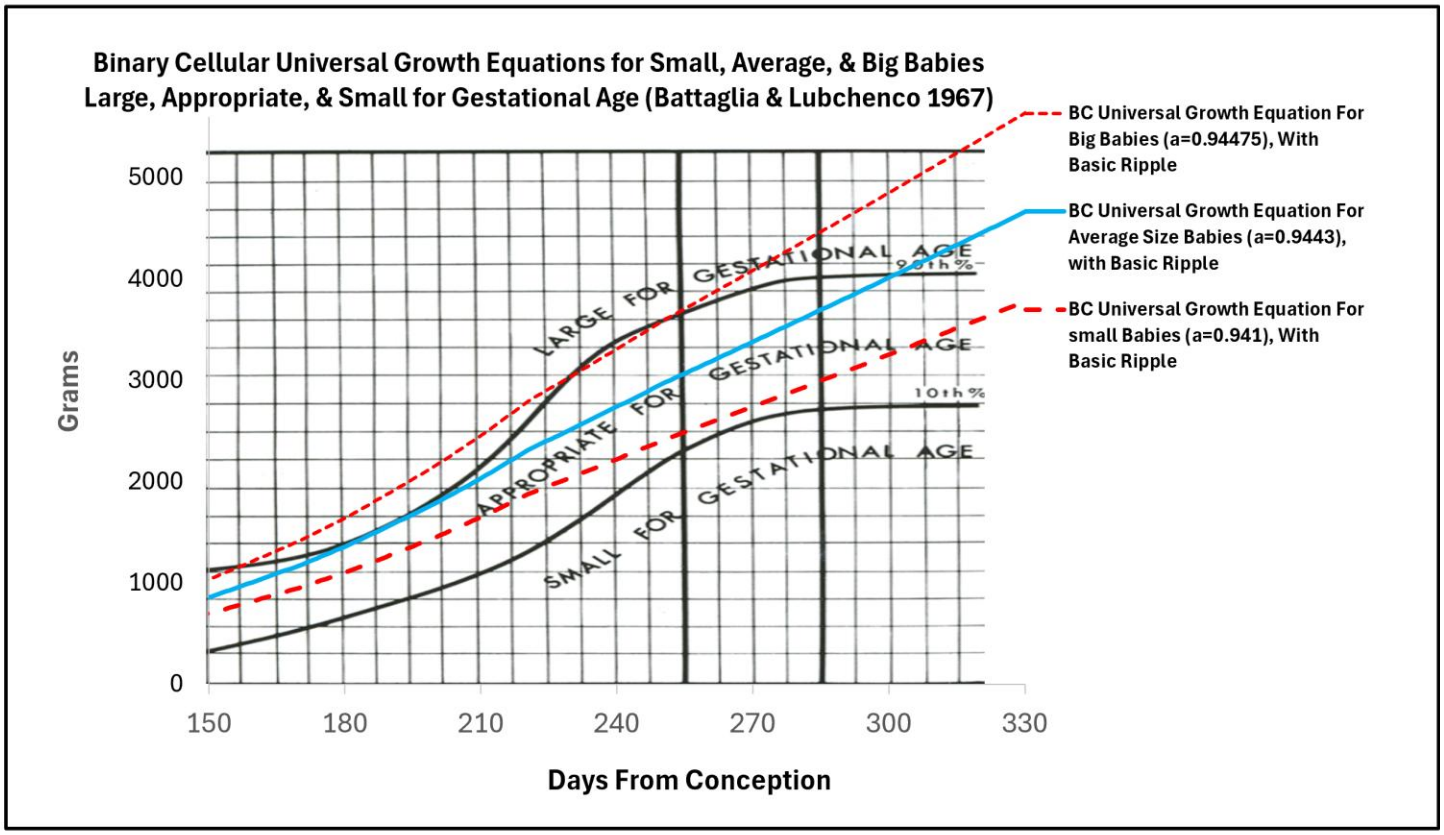


**<u>Supplement-Figure-III.36</u>.** ***Binary Cellular Universal Growth Equations for Small, Average, & Big Babies***, and ***Large, Appropriate, & Small for Gestational Age*** (Battaglia & Lubchenco 1967)[275275].

The Curve of the ***Binary Cellular Universal Growth Equation*** is Quite Similar to Other Fetal Growth Curves.

Returning to the ***Binary Cellular Universal Growth Equation*** (#2), loaded with the parameters/modifiers ***a***, ***b***, ***c*** and ***R***, derived from the babies in our dataset as a whole, we can compare them with other standards, specifically, the NICHD, and INTERGROWTH-21, fetal growth curves for the 50$^{th}$ percentile (**Supplement-Figure-III.37**).

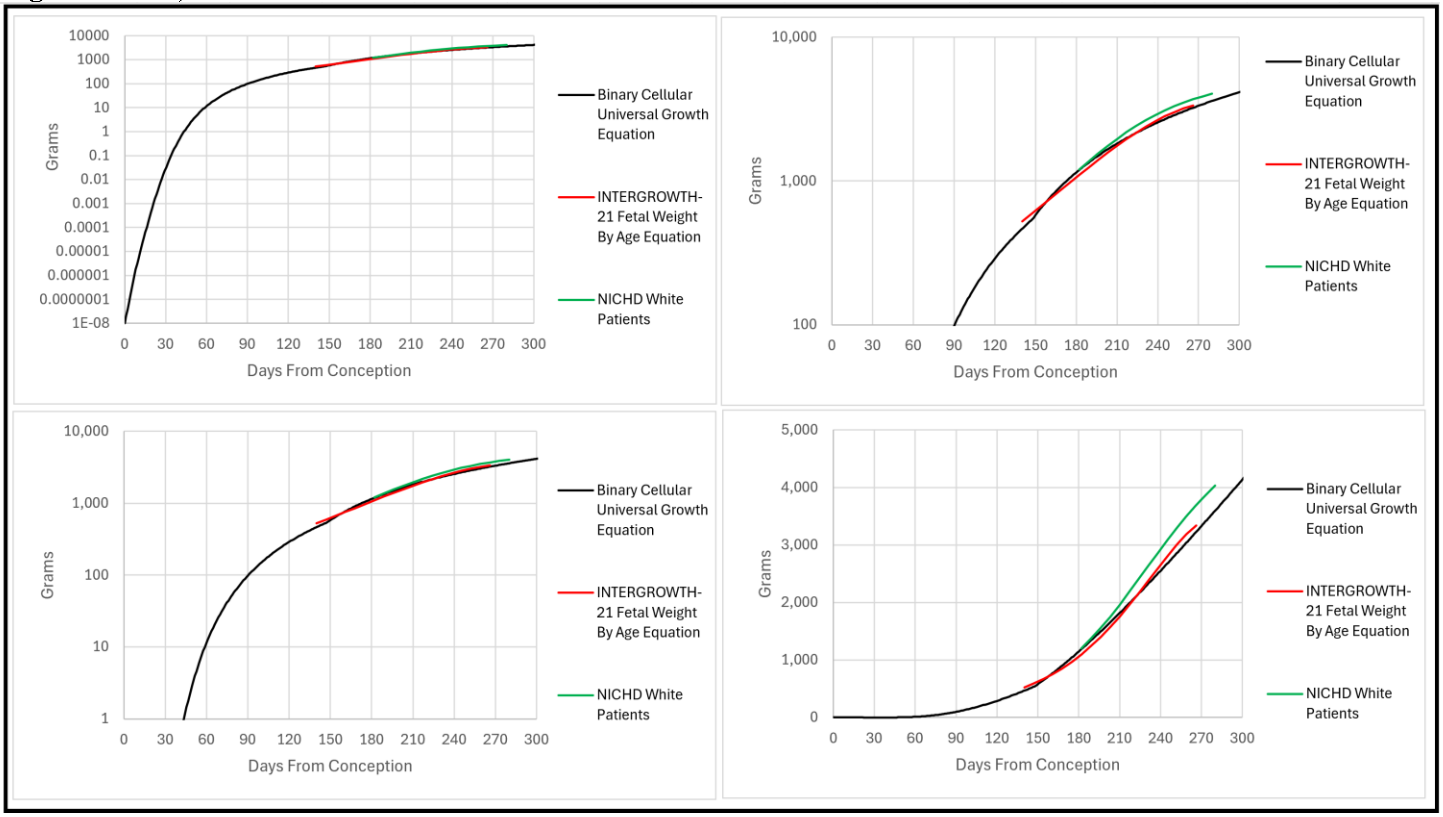


**Supplement-Figure-III.37**

***Weight-Gain Rate*** and ***Fetal Growth Velocity***.

Comparison of ***Weight-Gain Rate***, based on the ***Binary Cellular Universal Growth Equation*** (#2), (to use the terminology we have adopted), or 50th Percentile ***Fetal Growth Velocity*** (with timing to conception adjusted from gestational age by subtracting 14 days) from the NICHD[276], and INTERGROWTH-21[277] fetal growth curves for the 50th percentile, can be seen in **Supplement-Figure-III.38**

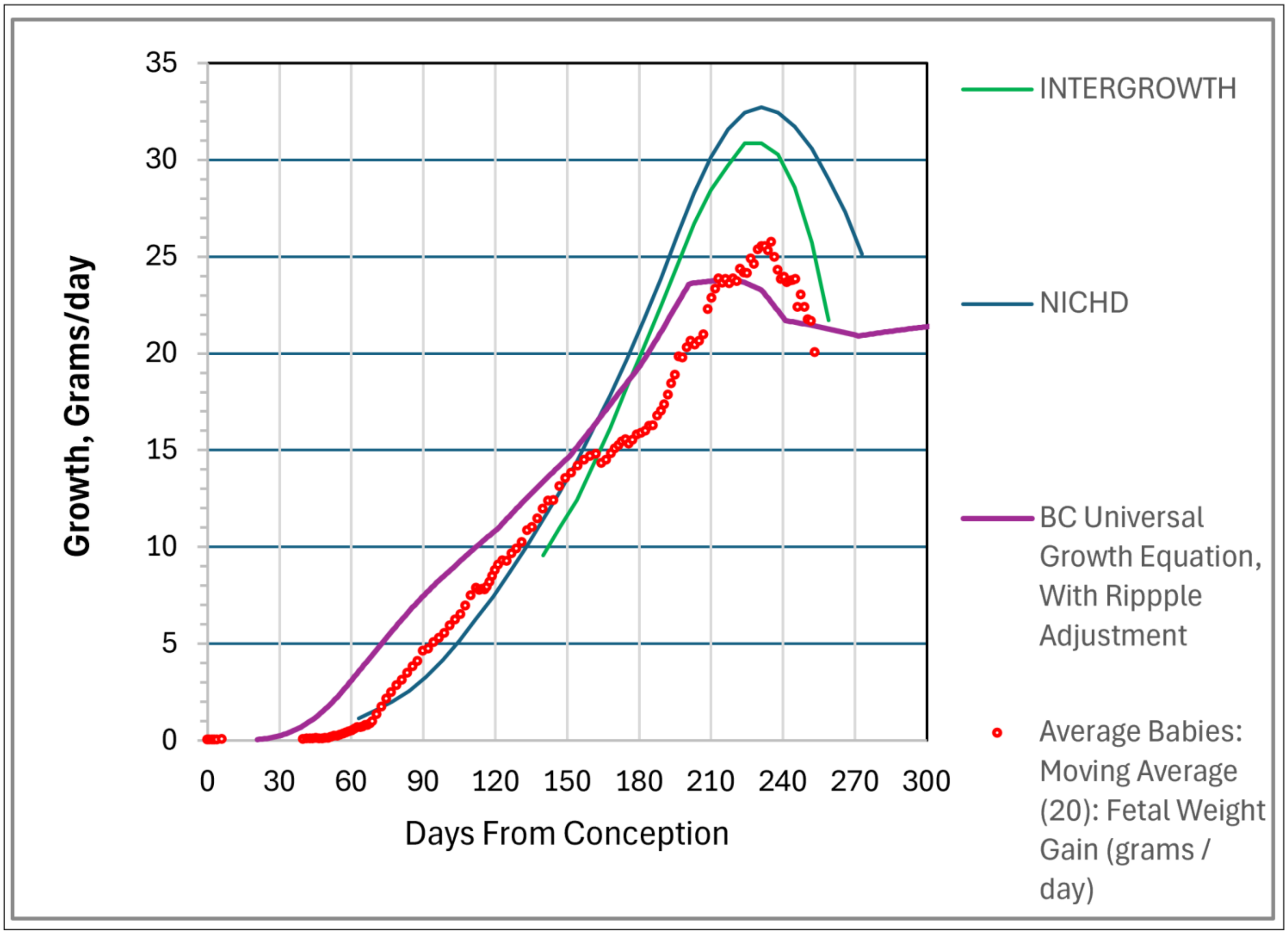


**Supplement-Figure-III.38** ***Weight-Gain Rate***, based on the ***Binary Cellular Universal Growth Equation*** (#2), and the 50th Percentile ***Fetal Growth Velocity*** (with timing to conception adjusted from Gestational age by subtracting 14 days) from the NICHD[276], and INTERGROWTH-21[277].

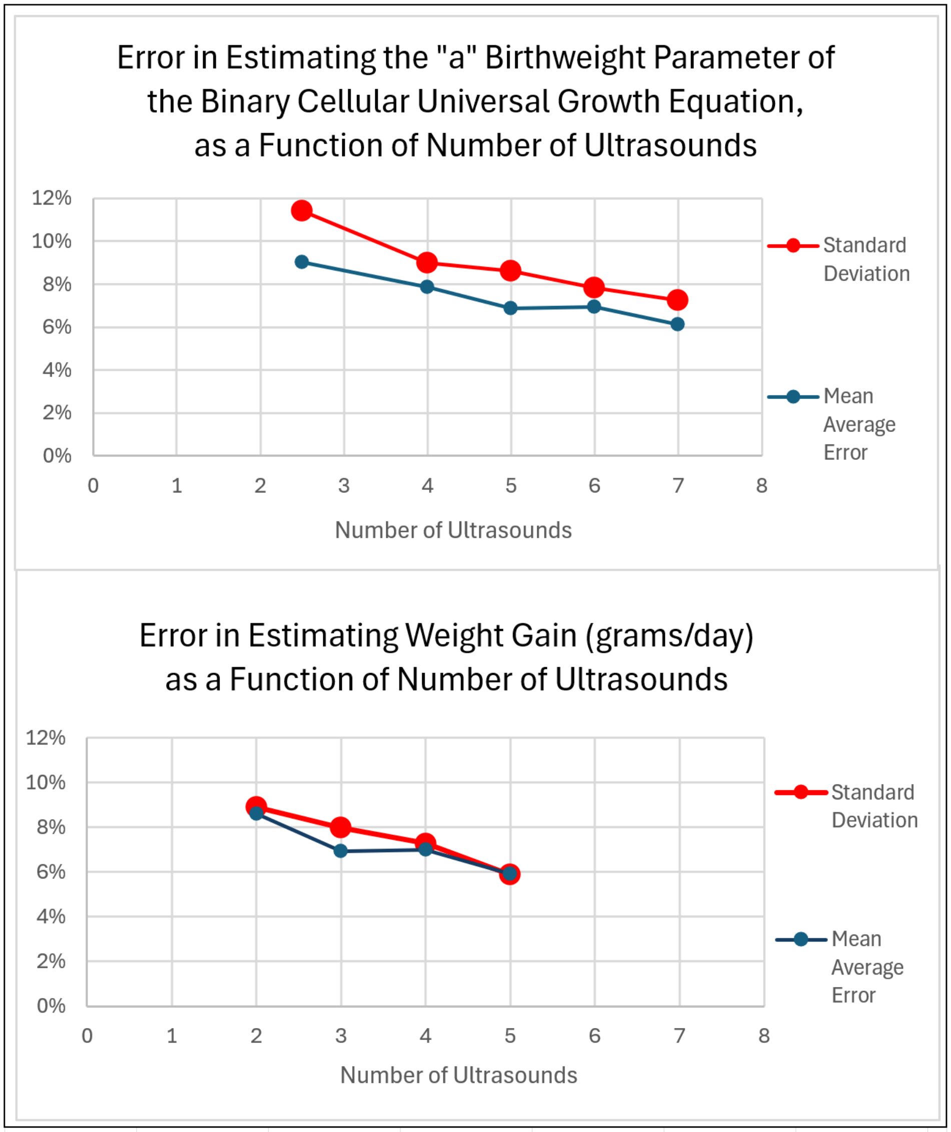


**Supplement-Figure-III.39**

***Error in Estimating the "a" Parameter of the Binary Cellular Universal Growth Equation, as a Function of Number of Ultrasounds***

***Error in Estimating Weight Gain (grams/day) as a Function of Number of Ultrasounds***

***Weight-Gain Rate***, based on the ***Binary Cellular Universal Growth Equation*** (#2), and the 50$^{th}$ Percentile ***Fetal Growth Velocity***

**_Binary Cellular Mathematics_** Provides New Methods for Predicting and Managing **_Fetal Growth_** and **_Birth_**

Birthweight = -171.98 * (EF $weight_1$ - EF $weight_2$) / time between ultrasounds) + 1185.7

Fetal Weight = EF $weight_2$ * (EF $weight_1$ - EF $weight_2$) / time between ultrasounds) * Days From Now

Days Until conception =

Days Until Birth =

C:_1\BBCDG V7 DATA_______Baby Math Data 10 20 23______New Luke Lin Studies 1 2 2024\what to predict 5 22 2024.xlsx

**DISCUSSION**

In this, and the previous two papers in this series, we have seen that a consideration of animal ***Size***, in units of numbers of cells, ***N***, a method we call ***Binary Cellular Analysis***, provides new insights into understanding, and measuring, ***Growth*** and ***Development***, an approach we call ***Binary Cellular Biology***. The ***Binary Cellular Analysis*** of growth data examined in the three papers in this series has revealed new equations for estimating and following ***Size*** and ***Growth*** from conception to maturity, deeper insights in the role of ***Cellular Selection*** in molding ***Growth*** and ***Development***, and new findings for the causes of ***Growth*** to ***Sizes*** that are ***Too Small***, or ***Too Large***, especially for human fetuses.

In the ***First Paper*** in this series,[262] we have seen that ***Binary Cellular Analysis*** could generate ***Binary Cellular Equations***, including:

1. The ***Binary Cellular Universal Growth Equation***, which captures growth from conception to maturity, and provides a method for creating biologically-based, data-driven, ***Growth Curves***.
2. The ***Binary Cellular Universal Mitotic Fraction Equation***, which lies within the ***Binary Cellular Universal Growth Equation***, and captures the mechanism of growth to adult size, ***Cellular Selection***, the result of the regulation of the fraction of cells dividing, the ***Mitotic Fraction***.
3. The ***Binary Cellular Allometric Growth Equation***, which captures the creation and growth of the ***tissues***, ***organs***, and ***anatomical structures*** of the body relative to the body as a whole, and its mechanism, again ***Cellular Selection***, by the regulation of cell division.
4. The ***Binary Cellular Estimated Fetal Weight Equation***, derived from the ***Binary Cellular Allometric Growth Equation***, which captures the relationship between human fetal ***Ultrasound Measurement*** and the ***Size*** of the body as a whole.

In the ***Second Paper*** in this series[263], we have seen that the most promising of these ***Binary Cellular Estimated Fetal Weight Equations***, the ***Abdominal Circumference Binary Cellular Estimated Fetal Weight Equation***, is superior to currently used ***Fetal Weight Equations***, while also yielding up new methods for improving ultrasound size assessment.

In this, the ***Third Paper*** in this series, we have seen that the ***Fetal Weight*** and subsequent ***Birthweight***, of individual fetuses, and groups of fetuses, can be examined against the ***Binary Cellular Universal Growth Equation***. ***Estimated Fetal Weight*** was calculated with the ***Abdominal Circumference Binary Cellular Estimated Fetal Weight Equation***, while the number of days since conception was calculated by the ***CRL Fetal Age Equation***. These equations make it possible to compare ***Discrete Fetal Age/Fetal Weigh*** values determined by ultrasound with ***Continuous Fetal Age/Fetal Weigh*** values generated by the ***Binary Cellular Estimated Fetal Weight Equation***, to isolate and quantify the ***Speed of Growth*** and ***Pregnancy Length***. We call this new method ***Binary Cellular Fetal Growth Analysis***. We call these graphs ***Binary Cellular Fetal Growth Charts***, which display whether fetuses arrive at, or outside of, the ***Healthy Square***, bounded by a healthy ***Birthweight***, at a healthy ***Fetal Age***.

***Binary Cellular Fetal Growth Analysis*** Isolates, and Measures, ***Speed of Growth*** and ***Pregnancy Length***

***Binary Cellular Fetal Growth Analysis*** makes it possible to isolate and measure the ***Speed of Growth*** and the ***Pregnancy Length***.[285] ***Binary Cellular Fetal Growth Analysis*** makes it possible to identify the mechanisms, and features, of human fetal growth to ***Birthweights*** that are ***Too Small***, or ***Too Large***, as well as ***Pregnancy Lengths*** that are ***Too Short***.

***Binary Cellular Fetal Growth Analysis*** Has Identified ***Four Categories of Fetal Growth***

While the ***Binary Cellular Fetal Growth Analysis*** of the patient dataset we have examined here reveals that the ***Speed of Growth*** and the ***Pregnancy Length*** are continuous qualities, with ***Slow Growth*** and ***Short Pregnancy*** statistically linked, one can capture some of the general principles by lumping babies into at least four categories of fetal growth:

1. ***AVERAGE GROWTH TO AVERAGE SIZE***, normal growth along the ***Binary Cellular Universal Growth Equation***, thus landing the newborn into the ***Healthy Square*** on the last day of pregnancy.
2. ***FAST GROWTH TO LARGE SIZE***, faster growth, captured by the ***"a" Parameter*** of the ***Binary Cellular Universal Growth Equation***, thus landing the newborn above the ***Healthy Square*** on the last day of pregnancy.
3. ***SLOW GROWTH TO SMALL SIZE***, growth that is both too slow, captured by the ***"a" Parameter*** of the ***Binary Cellular Universal Growth Equation***, and too short, thus landing the newborn below the ***Healthy Square*** on the last day of pregnancy.
4. ***SLOW GROWTH TO AVERAGE SIZE***, growth that is slow, captured by the ***"a" Parameter*** of the ***Binary Cellular Universal Growth Equation***, but long enough, thus landing the newborn into, or close to, the ***Healthy Square*** on the last day of pregnancy.

***Small*** and ***Large*** Babies Differ by ***"a" Parameter***.

Here we have seen that fetuses that become ***Small***, ***Average*** and ***Large*** babies differ in the ***Speed of Growth***, which can be traced to the value of the ***"a" Parameter*** of the ***Binary Cellular Universal Growth Equation***. These differences in growth leading to ***Small***, ***Average*** and ***Large*** babies can also be captured empirically by a measure of the ***Weight-Gain Rate*** (Grams/Day), made by two or more ultrasound measurements in the last months of pregnancy.

The ***"a" Parameter*** Links ***Small*** and ***Large Birth*** to the Endocrine System.

Modeling described in the first paper in this series[262] suggests that variation in the value of the ***"a" Parameter*** may well reflect the endocrinological control of fetal growth. The possibility that the ***Fast*** growing fetuses that become ***Large*** babies may be the result of endocrinology is not a surprise, as ***Diabetes*** is a well-known cause of such growth.[278-281,] However, the possibility that ***Slow*** growing fetuses that become ***Small*** babies could also be the result a hormonal effect, but in the opposite direction, is striking, and worthy of deeper analysis, a point we shall return to below.

***Short Pregnancy*** is Caused by ***Slow Growth***.

***Binary Cellular Fetal Growth Analysis*** reveals that almost all babies with ***Short Pregnancy Lengths*** were ***Growing Slowly***. Indeed, babies with the ***Slowest Growth*** and ***Smallest Birthweight*** tended to have the ***Shortest Pregnancy Lengths***. This correlation was especially pronounced in the babies with ***Smallest Birthweight***, starting at 700 grams, with ***Pregnancy Length*** increasing with ***Birthweight*** until about 2500 grams. After that 2500-gram ***Birthweight***, size, all groups of babies in the 2500-to-4599-gram range appeared to settle into roughly the same ***Pregnancy Length***. Apparently, ***Slow Growth*** and ***Pregnancy Length*** are not independent qualities, or one would expect that ***Pregnancy Lengths*** would increase for babies with ***Birthweights*** increasing from 2500 to 4599 grams.

The Darwinian Logic of ***Slow Growth*** Causing ***Short Pregnancy***.

These observations raise the possibility that ***Slow Growing*** fetuses being ***Born Early*** may be the natural consequences of our Darwinian origins, rather than medical accidents. In the pre-modern era, ***Natural Selection*** may well have favored mothers that expel fetuses that were growing ***Slowly***, simply because they were unlikely to survive. Such a biologically ruthless process of expelling ***Slow Growing*** fetuses could have arisen evolutionarily, by its potential to increase the reproductive fitness of the mother, who would save biological resources for a subsequent birth, thus increasing the chance of the survival of the subsequent birth, as well as causing the subsequent birth to occur sooner.[282-285] Of course, a major goal of medicine is to avoid such lamentable biology. Fortunately, ***Binary Cellular Fetal Growth Analysis*** points to some ways to avert such tragedies, as we shall examine below.

The Darwinian Logic of ***Gestational Diabetes***.

Another biological consideration of the findings of the ***Binary Cellular Fetal Growth Analysis*** of ***Fetal Ultrasound*** and ***Birthweight*** data reported here offers a hypothesis concerning ***Gestational Diabetes***, which affects roughly 8% of all pregnancies.[278-281] We perceive the consequences of ***Gestational Diabetes*** in ***Large Babies***, that is ***Macrosomia***, no doubt due to the conversion of ***Average Growing Fetuses*** into ***Fast Growing Fetuses***, likely traceable to the increase in the value of the ***"a" Parameter*** of the ***Binary Cellular Universal Growth Equation***. What we can't perceive is the conversion of ***Slow Growing Fetuses*** into ***Average Growing Fetuses***, again likely traceable to the increase in the value of the ***"a" Parameter***. Thus, perhaps ***Gestational Diabetes*** is not a disease at all, but an adaptive biological mechanism, which acts to make ***Slow Growing Fetuses*** transition into ***Average Growing Fetuses***, thus avoiding ***Early Birth*** at ***Small Size***. The adaptive loss due to some babies being born ***Too Large*** may be made up for by the adaptive benefit of preventing other babies from being born ***Too Small***.

***Binary Cellular Analysis*** Tells Us That ***Multiple*** Ultrasounds are Better than ***Individual*** Ultrasounds.

As we can see in **Supplement-Figure-III.6**, measurement of ***Fetal Growth***, and the ultimate distinction between fetuses that will become ***Small***, ***Average***, and ***Big Babies***, is most effectively carried out with ***Multiple*** ultrasound measurements, at sequential points in time, spaced as far apart as is feasible.

***Binary Cellular Analysis*** Tells Us How Multiple Ultrasounds Can be Combined for Best Measure.

Note that fit of ***Multiple*** ultrasound measurements can be made to the ***~Linear Phase of Human Fetal Growth***, in units of ***Weight-Gain Rate*** (grams/day). Even more accurate and precise measurements, based on the ***Binary Cellular Universal Growth Equation***, in terms of the ***"a" Parameter***, can be made by taking advantage of ultrasound measurements made earlier in life.

***Binary Cellular Math*** Can Identify Optimal Ultrasounds Schedules.

We now have the tools at hand to determine what would be the optimal schedule for making ultrasound measurements[286-292], so as to achieve the most accurate, precise, and useful calculation results. Such an analysis is mathematically comparable to identifying optimal scheduling for cancer screening tests.[293,294]

***Binary Cellular Analysis*** Tells Us That ***Fetal Growth Speed*** Is More Informative Than ***Fetal Size***.

While we tend to think of abnormal fetuses as ***Small for Gestational Age*** and ***Large for Gestational Age***, values that are identified by ***Single*** ultrasound measurements, it may be more productive to consider fetuses as growing ***Fast*** or ***Slow***, qualities that the findings reported here show can be more accurately measured with ***Multiple*** ultrasound measurements. Furthermore, unlike identifying ***Small for Gestational Age*** and ***Large for Gestational Age***, identifying ***Fast*** or ***Slow*** growth requires no information on ***Fetal Age***. Fit of ***Multiple*** ultrasound measurements to the ***Binary Cellular Universal Growth Equation*** is sufficient for such assessments. Such measurements can be made in terms of ***Weight-Gain Rate*** (grams/day) or in terms of the ***"a" Parameter***.

***Binary Cellular Math*** Can Identify and Quantify Other Types of Fetal Growth.

While the four categories of fetal growth outlined above provide a first view of fetal growth, other varieties of growth remain to be discovered as more data becomes available. For example, one might expect that abrupt changes in the ***Mitotic Fraction*** could also occur, resulting in ***Large*** and ***Small Birthweights***. There may also be much benefit in examining the ***Speed of Growth***, and ***Pregnancy Length***, as continuous qualities, rather than categorical entities.

It has been suggested that slow growth is different when occurring early in pregnancy, or late.[295,296] Whether the differences in the value of the ***"a" Parameter*** that we have found to distinguish ***Slow***, ***Average*** and ***Fast*** growing fetuses become ***Small***, ***Average***, and ***Large*** babies occur throughout pregnancy, or arise at some period of time after conception, has yet to be determined. The dataset examined here was too small to examine such a question, but with larger datasets, ***Binary Cellular Fetal Growth Analysis*** has the potential to settle this point rigorously.

Also, yet to be determined is whether ***Small***, ***Average*** and ***Large*** babies differ in the values of the other two parameters of the ***Binary Cellular Universal Growth Equation***, ***b***, and ***c***, or in the modifier, ***R***. Again, as more data become available, is should be possible to identify how the ***Binary Cellular Growth Parameters***, of the ***Binary Cellular Universal Growth Equation***, vary among fetuses with various outcomes.

***Binary Cellular Math*** Can Identify Patient Characteristics Matching to ***Speed of Growth***.

The capacity of ***Binary Cellular Fetal Growth Analysis*** to isolate out the ***Speed of Growth***, as measured by the ***"a" Parameter*** of the ***Binary Cellular Universal Growth Equation***, and the ***Pregnancy Length***, as well as by this method's ability to sort babies into four categories of fetal growth, allows these features of fetal growth to be examined for their associations with the genotype[297-299] and phenotype of the mother and the child. Abundant evidence already suggests that diabetes would be expected to be associated with ***FAST GROWTH TO LARGE SIZE***, and its higher value of the ***"a" Parameter*** of the ***Binary Cellular Universal Growth Equation***.[278-281] The finding reported here that ***SLOW GROWTH TO SMALL SIZE*** is also associated with a change in the value of the ***"a" Parameter***, but in the opposite direction, also makes one wonder whether ***Slow Growth*** may also be associated with a hormonal cause, also in the opposite direction. This possibility provides a motivation for searching for endocrinological correlates of both ***Fast*** and ***Slow*** growth, leading to ***Large*** and ***Small Birthweights***.

***Binary Cellular Math*** Can Identify Patient Characteristics Matching to ***Pregnancy Length.***

***Binary Cellular Fetal Growth Analysis*** reveals that not all ***Slow Growing*** fetuses are ***Born Early***. Indeed, as we have seen here, ***Binary Cellular Fetal Growth Analysis*** provides a way to independently measure the ***Speed of Growth*** and ***Fetal Age*** at birth, that is, ***Pregnancy Length***[285], which we can treat as either continuous or categorical measures. In this way, ***Binary Cellular Fetal Growth Analysis*** provides a way to explore what genetic, physiologic, and other characteristics distinguish between the two fates of ***SLOW GROWTH TO SMALL SIZE***, and ***SLOW GROWTH TO AVERAGE SIZE***, so as to see how we might help the yet to be born land in the ***Healthy Square***.

***Binary Cellular Math*** Can Characterize Populations by Their ***Binary Cellular Growth Parameters***

Here we have seen that ***Small***, ***Average*** and ***Large*** babies differ by ***"a" Parameter*** of the ***Binary Cellular Universal Growth Equation***. Whether babies also differ in the other two parameters, ***b***, and ***c***, or in the modifier, ***R***, remains to be determined. The values of the ***"a" Parameter*** in our population represents a continuous distribution, and this is what one might expect for the ***b*** and ***c*** parameters, and the modifier, ***R***, as well. It follows that that each fetus can be expected to have a characteristic set of such values, determined by its genetic, environmental, medical, and stochastic experience, while populations of fetuses can be expected to have distributions of these values. It also follows that populations differing by genetics,[297-299] such as male vs female,[300,301] or ethnicity/race,[317,319,] or altitude,[302-324] or hormonal status[278-281], or any of another of the plethora of possibilities, may well differ in the values of these ***Growth Parameters***. Because each of these ***Growth Parameters*** is linked to a specific feature of the biology of cell division, such information allows one to gain insight into the mechanism of ***Growth*** from the data on ***Growth***.

***Binary Cellular Math*** Can Calculate What the ***Size*** of Each Fetus Will Be in the Days Ahead.

Once ***Binary Cellular Fetal Growth Analysis*** has measured the growth of a fetus, either in terms of ***Weight-Gain Rate*** (grams/day), or in terms of the value of the ***"a" Parameter*** of the ***Binary Cellular Universal Growth Equation***, it is then possible to predict growth in the future, together with statistical boundaries for such predictions. In that way, obstetricians can not only assess the ***Speed of Growth*** of the fetus, but also what ***Size*** it will be in the days and weeks ahead. This information can readily be displayed for each fetus on a ***Binary Cellular Fetal Growth Chart***.

***Binary Cellular Math*** Offers Definite Possibilities for Preventing Macrosomia.

Perhaps the most promising use of ***Binary Cellular Fetal Growth Analysis*** is the ability to identify the optimal timepoint by which ***Fast Growing*** fetuses should be born before they become ***Too Large*** to have a healthy birth.[264-310] For each fetus, such information may readily be seen on a ***Binary Cellular Fetal Growth Chart***.

***Binary Cellular Math*** Offers Testable Possibilities for Preventing Prematurity.

Should ***Gestational Diabetes*** be a natural biological mechanism for reversing ***Slow Growth***, and thus preventing ***Early Birth***, (SEE ABOVE), it may well be worth considering whether inducing a pharmacological equivalent in mothers of ***Slow Growing Fetuses*** may have a therapeutic benefit.[311-313] As these ***Slow Growing Fetuses*** can now be detected early by ***Binary Cellular Fetal Growth Analysis***, such a therapy could well offer a way to reverse ***Slow Growth*** and prevent ***Early Birth*** at ***Small Size***.

***Binary Cellular Math*** Can Aid Obstetricians in Guiding Fetuses into the ***Healthy Square*** at Birth.

As a starting point, obstetricians would like to see babies born within the confines of the conventional boundaries for a healthy birth, which is to be heavier than 2,500 grams, the conventional measure of being ***Too Small***, and smaller than 4,000 grams, the conventional measure of being ***Too Large***, and by at least 230 days post conception, that is, the ***Healthy Square*** (**Supplement-Figure-III.31**). ***Binary Cellular Fetal Growth Analysis*** of each fetus makes this possible, as the ***Abdominal Circumference Binary Cellular Estimated Fetal Weight Equation*** can help the obstetrician know how big the fetus is at the time of ultrasound. In addition, the ***Binary Cellular Universal Growth Equation***, loaded with the ***"a" Parameter***, derived from 2 or more ultrasound measurements, allows the obstetrician see what the fetus's ***Size*** is going to be in the days and weeks ahead, as visualized on its ***Binary Cellular Fetal Growth Chart***. The obstetrician can then make the appropriate clinical judgments, based on when the fetus will arrive at, and remain in, the ***Healthy Square***.

***Binary Cellular Math*** Can Transform ***Healthy Squares*** into More Informative ***Healthy Islands***.

Refining the point noted in the previous section, newborn outcome data can be used to further refine the borders of where a fetus is heading on its ***Binary Cellular Fetal Growth Chart***, by replacing the rectangular boundaries of the ***Healthy Square*** with the meandering concentric topological boundaries of a ***Healthy Island***, based on newborn morbidity and mortality data (**Supplement-Figure-III.38**). The obstetrician will then be provided with the essential information for making the appropriate clinical judgments, based on when the fetus will arrive at, and remain in, the irregularly bounded central locations of the ***Healthy Island***, where each pregnancy has its highest chance of a successful outcome.

***Binary Cellular Math*** Can Provide a ***Universal Reference*** for ***Fetal Growth Curves***.

There has been much concern as to how to construct ***Fetal Growth Curves*** that are applicable to all populations of pregnant women.[314-326,] As Grantz has articulated[327] "The findings from three modern, multicenter fetal growth reference studies, INTERGROWTH, NICHD and WHO, demonstrate wide variation in fetal growth among different countries and racial and ethnic groups. … Given the natural variation in fetal growth and differences in fetal growth percentiles among references, selecting a ***Universal Reference*** [ital. added] would make comparison of fetal growth simpler for clinical use and for comparison across populations." The ***Binary Cellular Universal Growth Equation*** offers the ability to provide such a ***Universal Reference*** for ***Fetal Growth Curves***, with the specific characteristics of each population captured by the values of the ***a***, ***b***, and ***c*** parameter, and the ***R*** modifier, found for each population. Grantz continues, by noting that "it is critical to test how a fetal growth reference performs in relation to clinically meaningful outcomes including neonatal morbidity and mortality in the local population to which it is being applied." Again, such information can be organized with the topologically concentric maps of the ***Healthy Islands***, which can characterize the risk of mortality and morbidity in each population.

The ***Binary Cellular Universal Growth Equation*** offers the possibility of identifying ***Fetal Growth Curves*** with ***"a" Parameter*** values that map to various centiles (3rd, 50th, and 97th percentiles are common examples). The reverse calculation is also possible, that is to say, to back calculate the percentile to which each fetus belongs. The distribution of these ***"a" Parameter*** values, and the ***Fetal Growth Curves*** generated by the ***Binary Cellular Universal Growth Equation*** offers the possibility of continuous improvement and application of making ***Fetal Growth Curves*** specific to various mothers, in different populations, in different racial and socioeconomic groups, in different parts of the world.

Web-based Calculators Will Provide Clinicians with ***Binary Cellular*** Calculations That Aid in Care

A first-generation Web-based calculator for calculating ***Fetal Weight*** from the ***Abdominal Circumference Binary Cellular Estimated Fetal Weight Equation***, is now available at "https://kidzgrowth.com". Encoding the additional ***Binary Cellular*** calculations described here is straightforward,[328] and underway, including calculation of the speed of fetal growth from multiple, sequential, ***Ultrasound Measurements***, in units of ***Weight-Gain Rate*** (grams/day), and in units of the value of the ***"a" Parameter*** of the ***Binary Cellular Universal Growth Equation***, as well as projected values ***Fetal Weight*** in the days and weeks ahead. displayed on a ***Binary Cellular Fetal Growth Chart***, with the statistical limits of reliability also shown. Also to be provided by these calculators are tools for performance feedback, comparing each ***Ultrasound Measurement*** to the ultimate ***Birthweight***, so that sonographers and their supervisors can review performance over the long term, as well as the performance of each individual case. These web-based calculators make the results of these ***Binary Cellular*** Mathematical manipulations widely available for aiding in obstetric care of individual pregnancies.

### REFERENCES

#### PAPER I


1 Michaelson J. Cell selection in development. Biological Reviews 1987; 62:115-139.
2 Michaelson, J. Cell Selection in the Genesis of Multicellular Organization Lab Invest. 69: 136-150 1993
3 Michaelson J. The role of molecular discreteness in normal and cancerous growth. Anticancer Res. 1999 Nov-Dec;19(6A):4853-67. PMID: 10697599.
4 dos Reis M, Thawornwattana Y, Angelis K, Telford MJ, Donoghue PC, Yang Z. Uncertainty in the Timing of Origin of Animals and the Limits of Precision in Molecular Timescales. Curr Biol. 2015 Nov 16;25(22):2939-50. doi: 10.1016/j.cub.2015.09.066. Epub 2015 Oct 22. PMID: 26603774; PMCID: PMC4651906.
5 Dunn FS, Kenchington CG, Parry LA, Clark JW, Kendall RS, Wilby PR. A crown-group cnidarian from the Ediacaran of Charnwood Forest, UK. Nat Ecol Evol. 2022 Aug;6(8):1095-1104. doi: 10.1038/s41559-022-01807-x. Epub 2022 Jul 25. PMID: 35879540; PMCID: PMC9349040.
6 The Binary Cellular Biology of Human Growth: II. Calculating the Size of the Human Fetus from the Size of its Parts
7 The Binary Cellular Biology of Human Growth: III. Identifying, Comprehending, and Ameliorating Normal and Abnormal Fetal Size and Growth
8 Tsoularis, A, Wallace J. Analysis of logistic growth models. Mathematical biosciences 179, 21–55 (2001)
9 Verhulst, P.-F. "Deuxième mémoire sur la loi d'accroissement de la population." Mém. de l'Academie Royale des Sci., des Lettres et des Beaux-Arts de Belgique 2 0, 1-32, 1847
10 Wright S. Book review. Jour AmStat Assoc. 1926;21:494
11 Bertalanffy, L. von, (1934). Untersuchungen über die Gesetzlichkeit des Wachstums. I. Allgemeine Grundlagen der Theorie; mathematische und physiologische Gesetzlichkeiten des Wachstums bei Wassertieren. Arch. Entwicklungsmech., 131:613-652
12 Richards, F. J. (1959). A Flexible Growth Function for Empirical Use. Journal of Experimental Botany 10 (2): 290–300
13 West GB, Brown JH, Enquist BJ. A general model for ontogenetic growth. Nature. 2001 Oct 11;413(6856):628-31.
14 Warsof, Steven L., "Ultrasonic estimation of fetal weight for the detection of intrauterine growth retardation by computer-assisted analysis" (1977). Yale Medicine Thesis Digital Library. 3285. https://elischolar.library.yale.edu/ymtdl/3285
15 Shepard MJ, Richards VA, Berkowitz RL, Warsof SL, Hobbins JC. An evaluation of two equations for predicting fetal weight by ultrasound. Am J Obstet Gynecol. 1982 Jan 1;142(1):47-54. doi: 10.1016/s0002-9378(16)32283-9. PMID: 7055171.
16 Hadlock FP, Harrist RB, Carpenter RJ, Deter RL, Park SK. Sonographic estimation of fetal weight. The value of femur length in addition to head and abdomen measurements. Radiology. 1984 Feb;150(2):535-40. doi: 10.1148/radiology.150.2.6691115. PMID: 6691115.
17 Hadlock FP, Harrist RB, Sharman RS, Deter RL, Park SK. Estimation of fetal weight with the use of head, body, and femur measurements--a prospective study. Am J Obstet Gynecol. 1985 Feb 1;151(3):333-7. doi: 10.1016/0002-9378(85)90298-4. PMID: 3881966.
18 Combs CA, Jaekle RK, Rosenn B, Pope M, Miodovnik M, Siddiqi TA. Sonographic estimation of fetal weight based on a model of fetal volume. Obstet Gynecol. 1993 Sep;82(3):365-70. PMID: 8355935.
19 Hiwale S, Firtion C. Analysis of factors influencing accuracy of ultrasound-based fetal weight estimation. Indian J Radiol Imaging. 2020 Apr-Jun;30(2):156-162. doi: 10.4103/ijri.IJRI_167_19. Epub 2020 Jul 13. PMID: 33100682; PMCID: PMC7546305.
20 Stirnemann J, Villar J, Salomon LJ, Ohuma E, Ruyan P, Altman DG, Nosten F, Craik R, Munim S, Cheikh Ismail L, Barros FC, Lambert A, Norris S, Carvalho M, Jaffer YA, Noble JA, Bertino E, Gravett MG, Purwar M, Victora CG, Uauy R, Bhutta Z, Kennedy S, Papageorghiou AT; International Fetal and Newborn Growth Consortium for the 21st Century (INTERGROWTH-21st). International estimated fetal weight standards of the INTERGROWTH-21st Project. Ultrasound Obstet Gynecol. 2017 Apr;49(4):478-486. doi: 10.1002/uog.17347. Epub 2017 Mar 5. PMID: 27804212; PMCID: PMC5516164.
21 Dudley NJ. A systematic review of the ultrasound estimation of fetal weight. Ultrasound Obstet Gynecol. 2005 Jan;25(1):80-9. doi: 10.1002/uog.1751. PMID: 15505877.
22 Hammami A, Mazer Zumaeta A, Syngelaki A, Akolekar R, Nicolaides KH. Ultrasonographic estimation of fetal weight: development of new model and assessment of performance of previous models. Ultrasound Obstet Gynecol. 2018 Jul;52(1):35-43. doi: 10.1002/uog.19066. Epub 2018 Jun 3. PMID: 29611251.
23 Barel O, Vaknin Z, Tovbin J, Herman A, Maymon R. Assessment of the accuracy of multiple sonographic fetal weight estimation formulas: a 10-year experience from a single center. J Ultrasound Med. 2013 May;32(5):815-23. doi: 10.7863/ultra.32.5.815. PMID: 23620324.
24 Melamed N, Yogev Y, Meizner I, Mashiach R, Bardin R, Ben-Haroush A. Sonographic fetal weight estimation: which model should be used? J Ultrasound Med. 2009 May;28(5):617-29. doi: 10.7863/jum.2009.28.5.617. PMID: 19389901.
25 Hoopmann M, Kagan KO, Sauter A, Abele H, Wagner P. Comparison of Errors of 35 Weight Estimation Formulae in a Standard Collective. Geburtshilfe Frauenheilkd. 2016 Nov;76(11):1172-1179. doi: 10.1055/s-0042-118598. PMID: 27904167; PMCID: PMC5123881.

26 Ma J, Cheng D, Zhang Z, Cai B, Xu X. Evaluating the accuracy of sonographic fetal weight estimations using the Hadlock IV formula in a Chinese population. Quant Imaging Med Surg. 2023 Jun 1;13(6):3726-3734. doi: 10.21037/qims-22-778. Epub 2023 Apr 3. PMID: 37284125; PMCID: PMC10240036.
27 Milner J, Arezina J. The accuracy of ultrasound estimation of fetal weight in comparison to birth weight: A systematic review. Ultrasound. 2018 Feb;26(1):32-41. doi: 10.1177/1742271X17732807. Epub 2018 Feb 7. PMID: 29456580; PMCID: PMC5810856.
28 Knight W. Asymptotic Growth: An Example of Nonsense Disguised as Mathematics Journal of the Fisheries Research Board of Canada, 1968, 25(6): 1303-1307, 10.1139/f68-114
29 Pardo, S. A., Cooper, A. B., Dulvy, N. K. (2013), Avoiding fishy growth curves. Methods in Ecology and Evolution. Volume 4, Issue 4, pages 353–360, 2013
30 Hirst AG, Forster J. When growth models are not universal: evidence from marine invertebrates. Proc Biol Sci. 2013 14;280(1768):20131546.
31 Bricker L, Medley N, Pratt JJ. Routine ultrasound in late pregnancy (after 24 weeks' gestation). Cochrane Database Syst Rev 2015;(6):CD001451.
32 Lees CC, Stampalija T, Baschat A, et al. ISUOG Practice Guidelines: diagnosis and management of small-for-gestational-age fetus and fetal growth restriction. Ultrasound Obstet Gynecol 2020;56(2):298–312.
33 Melamed N, Baschat A, Yinon Y, et al. FIGO (International Federation of Gynecology and Obstetrics) initiative on fetal growth: best practice advice for screening, diagnosis, and management of fetal growth restriction. Int J Gynaecol Obstet 2021;152(Suppl. 1):3–57.
34 Kleinrouweler CE, Cheong-See FM, Collins GS, et al. Prognostic models in obstetrics: available, but far from applicable. Am J Obstet Gynecol 2016 Jan;214(1):79–90.e36. https://doi.org/10.1016/j.ajog.2015.06.013. Epub 2015 Jun 10. PMID: 26070707.
35 Hadlock FP, Harrist RB, Martinez-Poyer J. In utero analysis of fetal growth: a sonographic weight standard. Radiology 1991;181:129–33.
36 Nwabuobi C, Odibo L, Camisasca- Lopina H, Leavitt K, Tuuli M, Odibo AO. Comparing INTERGROWTH-21st Century and Hadlock growth standards to predict small for gestational age and short-term neonatal outcomes. J Matern Fetal Neonatal Med 2020;33: 1906–12.
37 Duncan JR, Odibo L, Hoover EA, Odibo AO. Prediction of large-for-gestational-age neonates by different growth standards. J Ultrasound Med 2021;40:963–70.
38 Blue NR, Beddow ME, Savabi M, Katukuri VR, Chao CR. Comparing the Hadlock fetal growth standard to the Eunice Kennedy Shriver National Institute of Child Health and Human Development racial/ethnic standard for the prediction of neonatal morbidity and small for gestational age. Am J Obstet Gynecol 2018;219:474.e1–12.
39 Kiserud T, Piaggio G, Carroli G, et al. The World Health Organization fetal growth charts: A multinational longitudinal study of ultrasound biometric measurements and estimated fetal weight. PLoS Med 2017;14:e1002220.
40 Buck Louis GM, Grewal J, Albert PS, et al. Racial/ethnic standards for fetal growth: the NICHD Fetal Growth Studies. Am J Obstet Gynecol 2015;213:449.e1–41.
41 Papageorghiou AT, Ohuma EO, Altman DG, et al. International standards for fetal growth based on serial ultrasound measurements: the Fetal Growth Longitudinal Study of the INTERGROWTH-21st Project. Lancet 2014;384:869–79.
42 Anderson NH, Sadler LC, McKinlay CJD, McCowan LME. INTERGROWTH-21st vs customized birthweight standards for identification of perinatal mortality and morbidity. Am J Obstet Gynecol 2016;214:509.e1–7
43 Fay E, Hugh O, Francis A, Katz R, Sitcov K, Souter V, Gardosi J. Customized GROW vs INTERGROWTH-21st birthweight standards to identify small for gestational age associated perinatal outcomes at term. Am J Obstet Gynecol MFM. 2022 Mar;4(2):100545. doi: 10.1016/j.ajogmf.2021.100545. Epub 2021 Dec 4. PMID: 34875415
44 Odibo AO, Nwabuobi C, Odibo L, Leavitt K, Obican S, Tuuli MG. Customized fetal growth standard compared with the INTERGROWTH-21st century standard at predicting small-for gestational-age neonates. Acta Obstet Gynecol Scand 2018;97:1381-7.
45 Cheng Y, Leung TY, Lao T, Chan YM, Sahota DS. Impact of replacing Chinese ethnicity-specific fetal biometry charts with the INTERGROWTH-21(st) standard. BJOG 2016;123(Suppl3):48–55
46 Hua X, Shen M, Reddy UM, et al. Comparison of the INTERGROWTH-21st, National Institute of Child Health and Human Development, and WHO fetal growth standards. Int J Gynaecol Obstet 2018;143:156–63.
47 Kabiri D, Romero R, Gudicha DW, et al. Prediction of adverse perinatal outcome by fetal biometry: comparison of customized and population-based standards. Ultrasound Obstet Gynecol 2020;55:177–88.
48 Grantz KL. Fetal growth curves: is there a universal reference? Obstet Gynecol Clin North Am 2021;48:281–96.
49 Rokas A, Mesiano S, Tamam O, LaBella A, Zhang G, Muglia L. Developing a theoretical evolutionary framework to solve the mystery of parturition initiation. Elife. 2020 Dec 31;9:e58343. doi: 10.7554/eLife.58343. PMID: 33380346; PMCID: PMC7775106
50 Manuck TA, Gyamfi-Bannerman C, Saade G. What now? A critical evaluation of over 20 years of clinical and research experience with 17-alpha hydroxyprogesterone caproate for recurrent preterm birth prevention. Am J Obstet Gynecol MFM. 2023 Oct;5(10):101108. doi: 10.1016/j.ajogmf.2023.101108. Epub 2023 Jul 30. PMID: 37527737; PMCID: PMC10591827.
51 Manuck TA. Refining Pharmacologic Research to Prevent and Treat Spontaneous Preterm Birth. Front Pharmacol. 2017 Mar 15;8:118. doi: 10.3389/fphar.2017.00118. PMID: 28360854; PMCID: PMC5350137.
52 Spiliopoulos D, Economides DL. Early fetal growth in progesterone-treated IVF pregnancies. Arch Gynecol Obstet. 2016 Jul;294(1):63-9. doi: 10.1007/s00404-015-3951-3. Epub 2015 Nov 13. PMID: 26563314.

53 P. Go´mez-Requeni, L. E. C. Conceic¸a˜o, A.-E. Olderbakk Jordal, I. Rønnestad A reference growth curve for nutritional experiments in zebrafish (Danio rerio) and changes in whole body proteome during development Fish Physiol Biochem (2010) 36:1199–1215
54 P. N. Claridge and I. C. Potter Movements, abundance, age composition and growth of bass, Dicentrarchus labrax, in the Severn Estuary and inner Bristol Channel. Journal of the Marine Biological Association of the United Kingdom Volume 63, Issue 4 November 1983, pp. 871-879
55 H. de Verdal, M. Vandeputte, E. Pepey, M.O. Vidal, C. Ouedraogo,M. Canonne, H. D'Cotta, J.F. Baroiller, E. Baras, B. Chatain Estimation of body weight of European sea bass (Dicentrarchus labrax) and Nile tilapia (Oreochromis niloticus) larvae by image analysis Proceedings, 10th World Congress of Genetics Applied to Livestock Production
56 Macdowell, E. C., Allen, E., MacDowell, C. G. (1927). The Prenatal Growth of the Mouse. Journal of General Physiologyy, 11(1), 57-70.
57 Poiley SM. Growth tables for 66 strains and stocks of laboratory animals. Lab Anim Sci. 1972 Oct;22(5):758-79.
58 Donaldson, H. H. The Rat: Reference Tables and Data for the Albino (Mus norvegicus albinus) and the Norway rat (Mus norvegicus). 278 (The Wistar Institute: 1915).
59 Mao, WH, Albrecht E, Teuscher F,Yang Q, Zhao RQ, Wegner J, Growth- and Breed-related Changes of Fetal Development in Cattle. Asian-Aust. J. Anim. Sci. Vol. 21, No. 5 : p 640 – 647 2008
60 Kratochvílová M, Hyánková L, Knížetová H,Fiedler J, URBAN F, Growth curve analysis in cattle from early maturity and mature body size viewpoints Czech J. Anim. Sci., 47, (4): 125–132 2002
61 Romanoff, A. L.. Romanoff, A. J. Biochemistry of the Avian Embryo; a quantitative analysis of prenatal development. Interscience Publishers (Wiley). 1967 New York
62 Blem, C. R., Zara, J. The Energetics of Young Bobwhite. Comparative Biochemistry and Physiology Part A: Physiology 61, 611–615 (1980)
63 Byerly, B. Y. T. C. Growth of the chick embryo in relation to its food supply. J. Exp. Biol., 9: 15-44. J. Exp. Biol. 9:15−44. 1932
64 Aggrey S. E. Comparison of Three Nonlinear and Spline Regression Models for Describing Chicken Growth Curves Poult Sci. 81(12):1782-8. 2002.
65 Škrtić Z, Kralik G, Gajčević Z, Growth evaluation of turkey heavy hybrid by means of asymmetric s-function agriculture, Vol.13 No.1 June 2007.
66 Adolph EF The Size of The Body And The Size of The Environment In The Growth of Tadpoles Biol. Bull. vol. 61 no. 3 350-375 1931
67 Loosanoff VL. Davis HC Conditioning V. Mercenaria for Spawning in Winter and Breeding Its Larvae in the Laboratory Biological Bulletin Vol. 98, No. 1 (Feb., 1950), pp. 60-65
68 Carriker. Embryogenesis and Organogenesis of Veligers and Early Juveniles in: Biology of the Hard Clam, Volume 31 (Developments in Aquaculture and Fisheries Science) J.N. Kraeuter (Editor), M. Castagna (Editor)
69 Harding JM. Relationships and growth patterns in the York River, Virginia 1954 TO 1970 Journal of Shellfish Research, Vol. 26, No. 1, 101–107, 2007.
70 Blaas H.-G. K., Taipale P, Torp H, Eik-Nes SH, Three-dimensional ultrasound volume calculations of human embryos and young fetuses: a study on the volumetry of compound structures and its reproducibility Ultrasound Obstet Gynecol 2006; 27: 640–646
71 Fenton TR, New growth chart for preterm babies: Babson and Benda's chart updated with recent data and a new format BMC Pediatrics 3 p13 2003
72 Flegal KM, Cole TJ. Construction of LMS parameters for the Centers for Disease Control and Prevention 2000 growth chart. National health statistics reports; no 63. Hyattsville, MD: National Center for Health Statistics. 2013. Percentile Data Files with LMS Values 0 - 5 years WHO Child Growth Standards WFA_boys_0_5_percentiles
http://www.cdc.gov/growthcharts/percentile_data_files.htm
73 Harlow GM Quinn P. Development of Preimplantation Mouse Embryos in vivo and in vitro. Aust J Biol Sci. 1982;35(2):187-93
74 Handyside, A. H., Hunter, S. Cell division and death in the mouse blastocyst before implantation. Roux 's Archives of Developmental Biology 519–526 (1986).
75 Bailey, F.R. and Miller, A.M. Text-Book of Embryology. New York: William Wood and Co (1921).
76 Leidenfrost S, Boelhauve M, Reichenbach, Tuna Gu¨ngo¨ M, Reichenbach HD, Sinowatz F, Wolf E, Habermann FA. Cell Arrest and Cell Death in Mammalian Preimplantation Development: Lessons from the Bovine Model PLoS ONE Volume 6, Issue 7, e22121 1 July 2011
77 Bakst MR, Gupta SK Akuffo V, Comparative Development of the Turkey and Chicken Embryo from Cleavage Through Hypoblast Formation Poult. Sci. vol. 76 no. 1 83-90 1997
78 Gupta SK Bakst MR, Turkey Embryo Staging From Cleavage Through Hypoblast Formation Journal of Morphology 217:313-325 (1993)
79 A. W. Marrable Variations in Early Cleavage of the Zebra Fish Nature 184, 1160 - 1161 (10 October 1959).
80 Patricia Cucchi, Elliott Sucre, Raphae¨l Santos, Jeremy Lecle`re, Guy Charmantier, Rene´ Castille Embryonic development of the sea bass Dicentrarchus labrax Helgol Mar Res (2012) 66:199–209
81 Ray PF, Conaghan, WinstonM, Handyside AH. Increased number of cells and metabolic activity in male human preimplantation embryos following in vitro fertilization J Reprod Fertil 104(1):165-71. 1995
82 Del Monte U Does the cell number 10^9 still really fit one gram of tumor tissue? Cell Cycle 8:3, 505-506; 1 February 2009

83 Sulston JE, Schierenberg E, White JG, Thomson JN. The embryonic cell lineage of the nematode Caenorhabditis elegans. Dev Biol. 1983 Nov;100(1):64-119.
84 Calderón-Urrea A, Vanholme B, Vangestel S, Kane SM, Bahaji A, Pha K, Garcia M, Snider A, Gheysen G. Early development of the root-knot nematode Meloidogyne incognita. BMC Dev Biol. 2016 Apr 28;16:10
85 Stach T, Anselmi C. High-precision morphology: bifocal 4D-microscopy enables the comparison of detailed cell lineages of two chordate species separated for more than 525 million years. BMC Biol. 2015 Dec 23;13:113.
86 Del Monte U. Does the cell number 10(9) still really fit one gram of tumor tissue? Cell Cycle. 2009 Feb 1;8(3):505-6. doi: 10.4161/cc.8.3.7608. Epub 2009 Feb 11. PMID: 19176997.
87 Romanoff AL The Avian Embryo Structural and Functional Development. The Macmillan Co 1960
88 Goedbloed JF.Embryonic and postnatal growth of rat and mouse. IV. Prenatal growth of organs and tissues: age determination, and general growth pattern. Acta Anat (Basel). 1976;95(1):8-33.
89 Collery RF, Veth KN, Dubis AM, Carroll J, Link BA. Rapid, accurate, and non-invasive measurement of zebrafish axial length and other eye dimensions using SD-OCT allows longitudinal analysis of myopia and emmetropization. Send to PLoS One. 2014 Oct 21;9(10):e110699.
90 Greiling TM, Clark JI.Early lens development in the zebrafish: a three-dimensional time-lapse analysis.Dev Dyn. 2009 Sep;238(9):2254-65.
91 A. W. Marrable Variations in Early Cleavage of the Zebra Fish Nature 184, 1160 - 1161 (10 October 1959).
92 Gomez-Requeni P, Conceicao L. E. C. Olderbakk AE, Rønnestad I. A reference growth curve for nutritional experiments in zebrafish (Danio rerio) and changes in whole body proteome during development Fish Physiol Biochem (2010) 36:1199–1215
93 de Boer BA, van den Berg G, de Boer PA, Moorman AF, Ruijter JM. Growth of the developing mouse heart: an interactive qualitative and quantitative 3D atlas.Dev Biol. 2012 Aug 15;368(2):203-13.
94 Jackson CM On the prenatal growth of the human body and the relative growth of the various organs and parts Am. J. Anat. 9:119–165 1909
95 Champman RL The Quantitative Analysis of Environmental Factors Ecology Vol. 9, No. 2 (Apr., 1928), pp. 111-122
96 Chang M, A ParkerEA, Muller TJM, Haenen C, Mistry M, Finkielstain GP, Murphy-Ryan M, Barnes KM, Sundaram R and Baron J. Changes in Cell-Cycle Kinetics Responsible for Limiting Somatic Growth in Mice FREE Pediatr Res 64: 240-245;
97 O'Farrell PH. Quiescence: early evolutionary origins and universality do not imply uniformity. Philos Trans R Soc Lond B Biol Sci. 2011 Dec 27;366 (1584):3498-507.
98 Hartwell LH- Nobel Lecture: Yeast and Cancer. Nobelprize.org. Nobel Media AB 2014. Web. 13 Sep 2015. <http://www.nobelprize.org/nobel_prizes/medicine/laureates/2001/hartwell-lecture.html>
99 Bertoli C, Skotheim JM, de Bruin RA. Control of cell cycle transcription during G1 and S phases. Nat Rev Mol Cell Biol. 2013 Aug;14(8):518-28. doi: 10.1038/nrm3629.
100 Chodrow, P, Su, J, Lee, D, Ahmed, T, He, N, De Man, R, Tiwari, A, Mannherz, W, Citi, L, DiCorpo, D, Michaelson, J S. How Our Cells Become Our Selves: The Cellular Phylodynamic Biology of Growth and Development. Preprint at: https://biorxiv.org/cgi/content/short/2021.09.22.461268v1 (2021)
101 Del Monte U Does the cell number 10^9 still really fit one gram of tumor tissue? Cell Cycle 8:3, 505-506; 1 February 2009
102 https://en.wikipedia.org/wiki/Exponential_integral
103 Edward O. Wilson and William H. Bossert, 1971, A Primer of Population Biology
104 Neal D, Introduction to Population Biology, 2nd Edition Cambridge University Press; 2 edition (January 17, 2019) ISBN-10: 1107605121 ISBN-13: 978-1107605121
105 Michaelson, J. The Role of Molecular Discreteness in Normal and Cancerous Growth. Anticancer Research 19: 4853-4868 (1999)
106 http://stattrek.com/probability-distributions/poisson.aspx
107 Grantz KL, Kim S, Grobman WA, Newman R, Owen J, Skupski D, Grewal J, Chien EK, Wing DA, Wapner RJ, Ranzini AC, Nageotte MP, Hinkle SN, Pugh S, Li H, Fuchs K, Hediger M, Buck Louis GM, Albert PS. Fetal growth velocity: the NICHD fetal growth studies. Am J Obstet Gynecol. 2018 Sep;219(3):285.e1-285.e36. doi: 10.1016/j.ajog.2018.05.016. Epub 2018 May 24. PMID: 29803819; PMCID: PMC7035912.
108 Wu T, Gong X, Zhao Y, Zhang L, You Y, Wei H, Zuo X, Zhou Y, Xing X, Meng Z, Lv Q, Liu Z, Zhang J, Hu L, Li J, Li L, Chen C, Liu C, Sun G, Liu A, Chen J, Lv Y, Wang X, Wei Y. Fetal growth velocity references from a Chinese population-based fetal growth study. BMC Pregnancy Childbirth. 2021 Oct 9;21(1):688. doi: 10.1186/s12884-021-04149-x. PMID: 34627184; PMCID: PMC8501691.
109 Ohuma EO, Villar J, Feng Y, Xiao L, Salomon L, Barros FC, Cheikh Ismail L, Stones W, Jaffer Y, Oberto M, Noble JA, Gravett MG, Wu Q, Victora CG, Lambert A, Di Nicola P, Purwar M, Bhutta ZA, Kennedy SH, Papageorghiou AT; International Fetal and Newborn Growth Consortium for the 21st Century (INTERGROWTH-21st Project). Fetal growth velocity standards from the Fetal Growth Longitudinal Study of the INTERGROWTH-21st Project. Am J Obstet Gynecol. 2021 Feb;224(2):208.e1-208.e18. doi: 10.1016/j.ajog.2020.07.054. Epub 2020 Aug 5. PMID: 32768431; PMCID: PMC7858163.
110 Partap U, Sovio U, Smith GC. Fetal Growth and the Risk of Spontaneous Preterm Birth in a Prospective Cohort Study of Nulliparous Women. Am J Epidemiol. 2016 Jul 15;184(2):110-9. doi: 10.1093/aje/kwv345. Epub 2016 Jul 1. PMID: 27370790; PMCID: PMC5860261.
111 Mongelli M, Benzie R, Condous G. Average fetal weekly weight gain: a novel measure of fetal growth velocity. J Matern Fetal Neonatal Med. 2016;29(4):676-9. doi: 10.3109/14767058.2015.1015981. Epub 2015 Feb 25. PMID: 25714480..

112 Michaelson J. Cell selection in development. Biological Reviews 1987; 62:115-139.
113 Luca Citi, Jessica Su, Luke Huang, James S Michaelson. Counting Cells by Age Tells Us About How, and Why, and When, We Grow, and Become Old and Ill. medRxiv 2023.01.05.23284244; doi: https://doi.org/10.1101/2023.01.05.23284244
114 Michaelson, J. Cell Selection in the Genesis of Multicellular Organization Lab Invest. 69: 136-150 1993
115 Michaelson J The significance of Cell Death. In: Apoptosis: The molecular basis of Cell Death. Cold Springs Harbor Laboratory Press 1991
116 Bartosch C, Vilar I, Rodrigues M, Costa L, Botelho N, Brandão O. Fetal autopsy parameters standards: biometry, organ weights, and long bone lengths. Virchows Arch. 2019 Oct;475(4):499-511. doi: 10.1007/s00428-019-02639-0. Epub 2019 Aug 16. PMID: 31420733.
117 Phillips JB, Billson VR, Forbes AB. Autopsy standards for fetal lengths and organ weights of an Australian perinatal population. Pathology. 2009;41(6):515-26. doi: 10.1080/00313020903041093. PMID: 19588281.
118 Mongelli M, Benzie R, Condous G. Average fetal weekly weight gain: a novel measure of fetal growth velocity. J Matern Fetal Neonatal Med. 2016;29(4):676-9. doi: 10.3109/14767058.2015.1015981. Epub 2015 Feb 25. PMID: 25714480.
119 Grantz KL, Kim S, Grobman WA, Newman R, Owen J, Skupski D, Grewal J, Chien EK, Wing DA, Wapner RJ, Ranzini AC, Nageotte MP, Hinkle SN, Pugh S, Li H, Fuchs K, Hediger M, Buck Louis GM, Albert PS. Fetal growth velocity: the NICHD fetal growth studies. Am J Obstet Gynecol. 2018 Sep;219(3):285.e1-285.e36. doi: 10.1016/j.ajog.2018.05.016. Epub 2018 May 24. PMID: 29803819; PMCID: PMC7035912
120 Ohuma EO, Villar J, Feng Y, Xiao L, Salomon L, Barros FC, Cheikh Ismail L, Stones W, Jaffer Y, Oberto M, Noble JA, Gravett MG, Wu Q, Victora CG, Lambert A, Di Nicola P, Purwar M, Bhutta ZA, Kennedy SH, Papageorghiou AT; International Fetal and Newborn Growth Consortium for the 21st Century (INTERGROWTH-21st Project). Fetal growth velocity standards from the Fetal Growth Longitudinal Study of the INTERGROWTH-21st Project. Am J Obstet Gynecol. 2021 Feb;224(2):208.e1-208.e18. doi: 10.1016/j.ajog.2020.07.054. Epub 2020 Aug 5. PMID: 32768431; PMCID: PMC7858163
121 Huxley JS (1932) Problems of relative growth. Methuen, London
122 Gould SJ Geometric Similarity in Allometric Growth: A Contribution to the Problem of Scaling in the Evolution of Size The American Naturalist, Vol. 105, No. 942 pp. 113-136 Mar - Apr. 1971
123 White JF Gould SJ Interpretation of the Coefficient in the Allometric Equation. The American Naturalist Vol. 99, No. 904 pp. 5-18 Jan. - Feb., 1965
124 Sober E. Reconstructing the Past: Parsimony, Evolution, and Inference. MIT Press. ISBN-10: 026219273X 1989
125 Fitch, W. Toward defining the course of evolution: minimum change for a specific tree topology. Syst. Zool., 20:406–416
126 Legant WR, Shao L, Grimm JB, Brown TA, Milkie DE, Avants BB, Lavis LD, Betzig E. High-density three-dimensional localization microscopy across large volumes. Nat Methods. 2016 Apr;13(4):359-65.
127 Khan S. Proof: d/dx(ln x) = 1/x. https://www.khanacademy.org/math/in-in-grade-12-ncert/in-in-advanced-differentiation-two/copy-of-proofs-for-derivatives-of-ex-and-lnx-ab/v/proof-d-dx-ln-x-1-x-old ; https://www.khanacademy.org/math/ap-calculus-ab/ab-antiderivatives-ftc/ab-common-indefinite-int/v/antiderivative-of-x-1
128 Phillips JB, Billson VR, Forbes AB. Autopsy standards for fetal lengths and organ weights of an Australian perinatal population. Pathology. 2009;41(6):515-26
129 Phillips JB, Billson VR, Forbes AB. Autopsy standards for fetal lengths and organ weights of an Australian perinatal population. Pathology. 2009;41(6):515-26
130 Papageorghiou AT, Kennedy SH, Salomon LJ, Ohuma EO, Cheikh Ismail L, Barros FC, Lambert A, Carvalho M, Jaffer YA, Bertino E, Gravett MG, Altman DG, Purwar M, Noble JA, Pang R, Victora CG, Bhutta ZA, Villar J; International Fetal and Newborn Growth Consortium for the 21st Century (INTERGROWTH-21st). International standards for early fetal size and pregnancy dating based on ultrasound measurement of crown-rump length in the first trimester of pregnancy. Ultrasound Obstet Gynecol. 2014 Dec;44(6):641-8. doi: 10.1002/uog.13448. Epub 2014 Nov 2. PMID: 25044000; PMCID: PMC4286014.
131 Zalokar, M.; Erk, I.; Division and migration of nuclei during early embryogenesis of Drosophila melanogaster Journal de microscopie et de biologie cellulaire 26 97-108 [1976]
132 Sawala A, Gould AP The sex of specific neurons controls female body growth in Drosophila PLoS Biol . 2017 Oct 4;15(10):e2002252. doi: 10.1371/journal.pbio.2002252. 2017
133 Church RB Robertson FW A biochemical study of the growth of Drosophila melanogaster. Journal of Experimental Zoology. Volume 162, 337-351, 1966.
134 Crick FH, Lawrence PA. Compartments and polyclones in insect development. Science. 1975 Aug 1;189(4200):340-7
135 Laugé G. Recherches expérimentales sur la détermination et la différenciation des caractères morphologiques et histologiques des intersexués Ann Embryol. Morph 2 245-270 1969
136 Nöthiger R. The Larval Development of Imaginal Disks. The Biology of Imaginal Disks pp 1-34 1972
137 Madhavan MM, Madhavan K. Morphogenesis of the epidermis of adult abdomen of Drosophila. J Embryol Exp Morphol 60:1-31. 1980
138 Madhavan MM Schneiderman HA. Histological Analysis of the Dynamics of Growth of Imaginal Discs and Histoblast Nests During the Larval Development of Drosophila melanogaster. Wilehm Roux Arch Dev Biol 183(4):269-305. 1977
139 Graves BJ, Schubiger G Cell cycle changes during growth and differentiation of imaginal leg discs in Drosophila melanogaster Dev Biol 93(1):104-10 1982 Sep;
140 Bate M, Arias AM The embryonic origin of imaginal discs in Drosophila Development 112(3):755-61 1991

141 P J Bryant PJ, Levinson P Intrinsic growth control in the imaginal primordia of Drosophila, and the autonomous action of a lethal mutation causing overgrowth Dev Biol 107(2):355-63. 1985
142 Verma P, Cohen SM miR-965 controls cell proliferation and migration during tissue morphogenesis in the Drosophila abdomen. Elife 30;4:e07389. 2015
143 Rabinowitz M. Studies on the cytology and early embryology of the egg of Drosophila melanogaster. Journal of Morphology 69(1):1 - 49 1941
144 Parker NF, Shingleton AW., The coordination of growth among Drosophila organs in response to localized growth-perturbation. Dev Biol 2011 Sep 15;357(2):318-25.
145 Ulrike N. On growth and form and patterning of drosophila wing imaginal discs October 2013 Dissertation. https://www.zora.uzh.ch/id/eprint/93491/1/diss_ulrike2.pdf
146 B Mintz B Clonal basis of mammalian differentiation Symp Soc Exp Biol. 1971;25:345-70
147 Jerne NK The Natural-Selection Theory of Antibody Formation. Proc Natl Acad Sci U S A. 1955 Nov 15;41(11):849-57
148 Humphries A Wright NA. Colonic crypt organization and tumorigenesis. Nature Reviews Cancer volume 8, pages415–424 (2008)
149 Klein AM, Nakagawa T, Ichikawa R, Yoshida S, Simons BD. Mouse germ line stem cells undergo rapid and stochastic turnover. Cell Stem Cell. 2010 Aug 6;7(2):214-24.
150 Wieschaus E, Gehring W. Clonal analysis of primordial disc cells in the early embryo of Drosophila melanogaster. Dev Biol. 1976 Jun;50(2):249-63
151 Postlethwait JH, Schneiderman HA. A clonal analysis of development in Drosophila melanogaster: Morphogenesis, determination, and growth in the wild-type antenna. Developmental Biology Volume 24, Issue 4, April 1971, Pages 477-519
152 Bryant PJ. Cell lineage relationships in the imaginal wing disc of Drosophila melanogaster Dev Biol. 1970 Jul;22(3):389-411
153 Larsen-Rapport E W. Imaginal Disc Determination: Molecular and Cellular Correlates. Annual Review of Entomology Vol. 31:145-175 January 1986
154 Bryant PJ, Schneiderman HA. Cell lineage, growth, and determination in the imaginal leg discs of Drosophila melanogaster. Dev Biol. 1969 Sep;20(3):263-90
155 Ripoll P. The embryonic organization of the imaginal wing disc of Drosophila melanogaster. Wilhelm Roux Arch Entwickl Mech Org. 1972 Sep;169(3):200-215.
156 Morata G, Garcia-Bellido A. Developmental analysis of some mutants of the bithorax system of Drosophila Wilehm Roux Arch Dev Biol. 1976 Jun;179(2):125-143.
157 Garcia-Bellido A, Merriam JR. Cell lineage of the imaginal discs in Drosophila gynandromorphs. J Exp Zool. 1969 Jan;170(1):61-75.
158 Hotta Y, Benzer S. Mapping of behaviour in Drosophila mosaics. Nature. 1972 Dec 29;240(5383):527-35.
159 Zalokar M, Erk I, Santamaría P. Distribution of ring-X chromosomes in the blastoderm of gynandromorphic D. melanogaster. Cell. 1980 Jan;19(1):133-41
160 Portin P. Analysis of the negative complementation of abruptex alleles in gynandromorphs of Drosophila melanogaster. Genetics.1977 Jun;86(2 Pt. 1):309-19.
161 Newman SA. Animal egg as evolutionary innovation: a solution to the "embryonic hourglass" puzzle. J Exp Zool B Mol Dev Evol. 2011 Nov 15;316(7):467-83. doi: 10.1002/jez.b.21417. Epub 2011 May 9. PMID: 21557469.
162 Warrington NM, Beaumont RN, Horikoshi M, Day FR, Helgeland Ø, Laurin C, Bacelis J, Peng S, Hao K, Feenstra B, Wood AR, Mahajan A, Tyrrell J, Robertson NR, Rayner NW, Qiao Z, Moen GH, Vaudel M, Marsit CJ, Chen J, Nodzenski M, Schnurr TM, Zafarmand MH, Bradfield JP, Grarup N, Kooijman MN, Li-Gao R, Geller F, Ahluwalia TS, Paternoster L, Rueedi R, Huikari V, Hottenga JJ, Lyytikäinen LP, Cavadino A, Metrustry S, Cousminer DL, Wu Y, Thiering E, Wang CA, Have CT, Vilor-Tejedor N, Joshi PK, Painter JN, Ntalla I, Myhre R, Pitkänen N, van Leeuwen EM, Joro R, Lagou V, Richmond RC, Espinosa A, Barton SJ, Inskip HM, Holloway JW, Santa-Marina L, Estivill X, Ang W, Marsh JA, Reichetzeder C, Marullo L, Hocher B, Lunetta KL, Murabito JM, Relton CL, Kogevinas M, Chatzi L, Allard C, Bouchard L, Hivert MF, Zhang G, Muglia LJ, Heikkinen J; EGG Consortium; Morgen CS, van Kampen AHC, van Schaik BDC, Mentch FD, Langenberg C, Luan J, Scott RA, Zhao JH, Hemani G, Ring SM, Bennett AJ, Gaulton KJ, Fernandez-Tajes J, van Zuydam NR, Medina-Gomez C, de Haan HG, Rosendaal FR, Kutalik Z, Marques-Vidal P, Das S, Willemsen G, Mbarek H, Müller-Nurasyid M, Standl M, Appel EVR, Fonvig CE, Trier C, van Beijsterveldt CEM, Murcia M, Bustamante M, Bonas-Guarch S, Hougaard DM, Mercader JM, Linneberg A, Schraut KE, Lind PA, Medland SE, Shields BM, Knight BA, Chai JF, Panoutsopoulou K, Bartels M, Sánchez F, Stokholm J, Torrents D, Vinding RK, Willems SM, Atalay M, Chawes BL, Kovacs P, Prokopenko I, Tuke MA, Yaghootkar H, Ruth KS, Jones SE, Loh PR, Murray A, Weedon MN, Tönjes A, Stumvoll M, Michaelsen KF, Eloranta AM, Lakka TA, van Duijn CM, Kiess W, Körner A, Niinikoski H, Pahkala K, Raitakari OT, Jacobsson B, Zeggini E, Dedoussis GV, Teo YY, Saw SM, Montgomery GW, Campbell H, Wilson JF, Vrijkotte TGM, Vrijheid M, de Geus EJCN, Hayes MG, Kadarmideen HN, Holm JC, Beilin LJ, Pennell CE, Heinrich J, Adair LS, Borja JB, Mohlke KL, Eriksson JG, Widén EE, Hattersley AT, Spector TD, Kähönen M, Viikari JS, Lehtimäki T, Boomsma DI, Sebert S, Vollenweider P, Sørensen TIA, Bisgaard H, Bønnelykke K, Murray JC, Melbye M, Nohr EA, Mook-Kanamori DO, Rivadeneira F, Hofman A, Felix JF, Jaddoe VWV, Hansen T, Pisinger C, Vaag AA, Pedersen O, Uitterlinden AG, Järvelin MR, Power C, Hyppönen E, Scholtens DM, Lowe WL Jr, Davey Smith G, Timpson NJ, Morris AP, Wareham NJ, Hakonarson H, Grant SFA, Frayling TM, Lawlor DA, Njølstad PR, Johansson S, Ong KK, McCarthy MI, Perry JRB, Evans DM, Freathy RM. Maternal and fetal genetic effects on birth weight and their relevance to cardio-metabolic risk factors. Nat Genet. 2019 May;51(5):804-814. doi: 10.1038/s41588-019-0403-1. Epub 2019 May 1. PMID: 31043758; PMCID: PMC6522365.

163 Juliusdottir T, Steinthorsdottir V, Stefansdottir L, Sveinbjornsson G, Ivarsdottir EV, Thorolfsdottir RB, Sigurdsson JK, Tragante V, Hjorleifsson KE, Helgadottir A, Frigge ML, Thorgeirsson G, Benediktsson R, Sigurdsson EL, Arnar DO, Steingrimsdottir T, Jonsdottir I, Holm H, Gudbjartsson DF, Thorleifsson G, Thorsteinsdottir U, Stefansson K. Distinction between the effects of parental and fetal genomes on fetal growth. Nat Genet. 2021 Aug;53(8):1135-1142. doi: 10.1038/s41588-021-00896-x. Epub 2021 Jul 19. PMID: 34282336
164 Solé-Navais P, Flatley C, Steinthorsdottir V, Vaudel M, Juodakis J, Chen J, Laisk T, LaBella AL, Westergaard D, Bacelis J, Brumpton B, Skotte L, Borges MC, Helgeland Ø, Mahajan A, Wielscher M, Lin F, Briggs C, Wang CA, Moen GH, Beaumont RN, Bradfield JP, Abraham A, Thorleifsson G, Gabrielsen ME, Ostrowski SR, Modzelewska D, Nohr EA, Hypponen E, Srivastava A, Talbot O, Allard C, Williams SM, Menon R, Shields BM, Sveinbjornsson G, Xu H, Melbye M, Lowe W Jr, Bouchard L, Oken E, Pedersen OB, Gudbjartsson DF, Erikstrup C, Sørensen E; Early Growth Genetics Consortium; Estonian Biobank Research Team; Danish Blood Donor Study Genomic Consortium; Lie RT, Teramo K, Hallman M, Juliusdottir T, Hakonarson H, Ullum H, Hattersley AT, Sletner L, Merialdi M, Rifas-Shiman SL, Steingrimsdottir T, Scholtens D, Power C, West J, Nyegaard M, Capra JA, Skogholt AH, Magnus P, Andreassen OA, Thorsteinsdottir U, Grant SFA, Qvigstad E, Pennell CE, Hivert MF, Hayes GM, Jarvelin MR, McCarthy MI, Lawlor DA, Nielsen HS, Mägi R, Rokas A, Hveem K, Stefansson K, Feenstra B, Njolstad P, Muglia LJ, Freathy RM, Johansson S, Zhang G, Jacobsson B. Genetic effects on the timing of parturition and links to fetal birth weight. Nat Genet. 2023 Apr;55(4):559-567. doi: 10.1038/s41588-023-01343-9. Epub 2023 Apr 3. Erratum in: Nat Genet. 2023 Jul;55(7):1250. doi: 10.1038/s41588-023-01412-z. PMID: 37012456; PMCID: PMC10101852
165 McIntyre HD, Catalano P, Zhang C, Desoye G, Mathiesen ER, Damm P. Gestational diabetes mellitus. Nat Rev Dis Primers. 2019 Jul 11;5(1):47. doi: 10.1038/s41572-019-0098-8. PMID: 31296866
166 Lovrić B, Šijanović S, Zmijanović I, Jurić G, Juras J. ULTRASOUND DIAGNOSIS OF MACROSOMIA AMONG WOMEN WITH GESTATIONAL DIABETES - REVIEW OF THE LITERATURE. Acta Clin Croat. 2022 Mar;61(1):95-106. doi: 10.20471/acc.2022.61.01.12. PMID: 36398082; PMCID: PMC9616028
167 J. Van't Hof and A. H. Sparrow A relationship between DNA content, nuclear volume, and minimum mitotic cycle time Proc Natl Acad Sci U S A. 1963 Jun; 49(6): 897–902
168 Cavalier-Smith T Nuclear volume control by nucleoskeletal DNA, selection for cell volume and cell growth rate, and the solution of the DNA C-value paradox. J Cell Sci. 1978 Dec;34:247-78
169 Francis D, Davies MS, Barlow PW. A strong nucleotypic effect on the cell cycle regardless of ploidy level. Ann Bot. 2008 Apr;101(6):747-57. 2008
170 Eddy SR. The ENCODE project: missteps overshadowing a success. Curr Biol. 2013 Apr 8;23(7)
171 Orgel LE, Crick F. Selfish DNA: the ultimate parasite. Nature. 1980 Apr 17;284(5757):604-7.
172 Alexander F. Palazzo, T. Ryan Gregory. The Case for Junk DNA. PLoS Genet. 2014 May; 10(5)
173 E. L. Jockusch An evolutionary correlate of genome size change in plethodontid salamanders. Proc Biol Sci. 1997 Apr 22; 264(1381): 597–604.
174 Chipman AD, Khaner O, Haas A, Tchernov E. The evolution of genome size: what can be learned from anuran development? J Exp Zool. 2001 Dec 15;291(4):365-74
175 Horner HA, Macgregor HC. C value and cell volume: their significance in the evolution and development of amphibians. J Cell Sci. 1983 Sep;63:135-46
176 Ferrari JA, Rai KS. 1989. Phenotypic correlates of genome size variation in Aedes albopictus. Evolution 43: 895–899.
177 I. A. McLaren, J.-M. Sevigny, C. J. Corkett Body sizes, development rates, and genome sizes among Calanus species Volume 47 of the series Developments in Hydrobiology pp 275-284 1988 Hydrobiologia pp 275-284 1988
178 Fabian, D., Flatt, T. Life History Evolution. Nature Education Knowledge 3(10):24. (2012)
179 Fox, C.W. & F.J. Messina. 2013. Life History. In Oxford Bibliographies in Ecology (Online Series). http://www.oxfordbibliographies.com/view/document/obo-9780199830060/obo-9780199830060-0016.xml?rskey=EYOgWp&result=3&q=Life+History#firstMatch.
180 Beta J, Khan N, Khalil A, Fiolna M, Ramadan G, Akolekar R, Maternal and neonatal complications of fetal macrosomia: systematic review and meta-analysis. Ultrasound Obstet Gynecol. 2019 Apr 2.
181 Moffat AS. Genetics. Transposons help sculpt a dynamic genome. Science. 2000 Sep 1;289(5484):1455-7.
182 Petrov DA, Sangster TA, Johnston JS, Hartl DL, Shaw KL. Evidence for DNA loss as a determinant of genome size. Science. 2000 Feb 11;287(5455):1060-2.
183 Nam K, Ellegren H. Recombination drives vertebrate genome contraction. PLoS Genet. 2012;8(5):e1002680. doi: 10.1371/journal.pgen.1002680. Epub 2012 May 3.
184 Gardosi J, Hugh O. Outcome-based comparative analysis of five fetal growth velocity models to define slow growth. Ultrasound Obstet Gynecol. 2023 May 16. doi: 10.1002/uog.26248. Epub ahead of print. PMID: 37191400.
185 J. Van't Hof and A. H. Sparrow A relationship between DNA content, nuclear volume, and minimum mitotic cycle time Proc Natl Acad Sci U S A. 1963 Jun; 49(6): 897–902
186 Mintz B. Gene control of mammalian pigmentary differentiation. I. Clonal origin of melanocytes. Proc Natl Acad Sci U S A. 1967 Jul;58(1):344-51
187 Gupta V, Poss KD Clonally dominant cardiomyocytes direct heart morphogenesis. Nature. 2012 Apr 25;484(7395):479-84
188 McKenna A, Findlay GM, Gagnon JA, Horwitz MS, Schier AF, Shendure J.Whole-organism lineage tracing by combinatorial and cumulative genome editing. Science.;353(6298) 2016

189 Raj B, Wagner DE, McKenna A, Pandey S, Klein AM, Shendure J, Gagnon JA, Schier AF Simultaneous single-cell profiling of lineages and cell types in the vertebrate brain. Nat Biotechnol. Mar 28. 2018
190 Sawala A, Gould AP The sex of specific neurons controls female body growth in Drosophila PLoS Biol . 2017 Oct 4;15(10):e2002252. doi: 10.1371/journal.pbio.2002252. 2017
191 Church RB Robertson FW A biochemical study of the growth of Drosophila melanogaster. Journal of Experimental Zoology. Volume 162, 337-351, 1966.
192 Crick FH, Lawrence PA. Compartments and polyclones in insect development. Science. 1975 Aug 1;189(4200):340-7
193 Madhavan MM, Madhavan K. Morphogenesis of the epidermis of adult abdomen of Drosophila J Embryol Exp Morphol 60:1-31. 1980
194 Verma P, Cohen SM miR-965 controls cell proliferation and migration during tissue morphogenesis in the Drosophila abdomen. Elife 30;4:e07389. 2015
195 Zalokar, M.; Erk, I.; Division and migration of nuclei during early embryogenesis of Drosophila melanogaster Journal de microscopie et de biologie cellulaire 26 97-108 [1976]
196 Rabinowitz M. Studies on the cytology and early embryology of the egg of Drosophila melanogaster. Journal of Morphology 69(1):1 - 49 1941
197 Laugé G. Recherches expérimentales sur la détermination et la différenciation des caractères morphologiques et histologiques des intersexués Ann Embryol. Morph 2 245-270 1969
198 Nöthiger R. The Larval Development of Imaginal Disks. The Biology of Imaginal Disks pp 1-34 1972
199 Luca Citi, Jessica Su, Luke Huang, James S Michaelson Counting Cells by Age Tells Us About How, and Why, and When, We Grow, and Become Old and Ill medRxiv 2023.01.05.23284244; doi: https://doi.org/10.1101/2023.01.05.23284244
200 Desjobert C, El Maï M, Gérard-Hirne T, Guianvarc'h D, Carrier A, Pottier C, Arimondo PB, Riond J. Combined analysis of DNA methylation and cell cycle in cancer cells. Epigenetics. 2015;10(1):82-91. doi: 10.1080/15592294.2014.995542. Epub 2015 Jan 23.
201 Reisenauer A, Kahng LS, McCollum S, Shapiro L. Bacterial DNA methylation: a cell cycle regulator? J Bacteriol. 1999 Sep;181(17):5135-9.
202 Burnet, F. M. (1957). "A modification of Jerne's theory of antibody production using the concept of clonal selection". Australian Journal Science. 20 (2): 67–69.
203 Michaelson JS, Chen LL, Bush D, Fong A, Smith B, Younger J. Improved web-based calculators for predicting breast carcinoma outcomes. Breast Cancer Res Treat. 2011 Aug;128(3):827-35
[204] **PAPER II**
205 Warsof, Steven L., "Ultrasonic estimation of fetal weight for the detection of intrauterine growth retardation by computer-assisted analysis" (1977). Yale Medicine Thesis Digital Library. 3285. https://elischolar.library.yale.edu/ymtdl/3285
206 Shepard MJ, Richards VA, Berkowitz RL, Warsof SL, Hobbins JC. An evaluation of two equations for predicting fetal weight by ultrasound. Am J Obstet Gynecol. 1982 Jan 1;142(1):47-54. doi: 10.1016/s0002-9378(16)32283-9. PMID: 7055171.
207 Hadlock FP, Harrist RB, Carpenter RJ, Deter RL, Park SK. Sonographic estimation of fetal weight. The value of femur length in addition to head and abdomen measurements. Radiology. 1984 Feb;150(2):535-40. doi: 10.1148/radiology.150.2.6691115. PMID: 6691115.
208 Hadlock FP, Harrist RB, Sharman RS, Deter RL, Park SK. Estimation of fetal weight with the use of head, body, and femur measurements--a prospective study. Am J Obstet Gynecol. 1985 Feb 1;151(3):333-7. doi: 10.1016/0002-9378(85)90298-4. PMID: 3881966.
209 Combs CA, Jaekle RK, Rosenn B, Pope M, Miodovnik M, Siddiqi TA. Sonographic estimation of fetal weight based on a model of fetal volume. Obstet Gynecol. 1993 Sep;82(3):365-70. PMID: 8355935.
210 Hiwale S, Firtion C. Analysis of factors influencing accuracy of ultrasound-based fetal weight estimation. Indian J Radiol Imaging. 2020 Apr-Jun;30(2):156-162. doi: 10.4103/ijri.IJRI_167_19. Epub 2020 Jul 13. PMID: 33100682; PMCID: PMC7546305.
211 Stirnemann J, Villar J, Salomon LJ, Ohuma E, Ruyan P, Altman DG, Nosten F, Craik R, Munim S, Cheikh Ismail L, Barros FC, Lambert A, Norris S, Carvalho M, Jaffer YA, Noble JA, Bertino E, Gravett MG, Purwar M, Victora CG, Uauy R, Bhutta Z, Kennedy S, Papageorghiou AT; International Fetal and Newborn Growth Consortium for the 21st Century (INTERGROWTH-21st). International estimated fetal weight standards of the INTERGROWTH-21st Project. Ultrasound Obstet Gynecol. 2017 Apr;49(4):478-486. doi: 10.1002/uog.17347. Epub 2017 Mar 5. PMID: 27804212; PMCID: PMC5516164.
212 Dudley NJ. A systematic review of the ultrasound estimation of fetal weight. Ultrasound Obstet Gynecol. 2005 Jan;25(1):80-9. doi: 10.1002/uog.1751. PMID: 15505877.
213 Hammami A, Mazer Zumaeta A, Syngelaki A, Akolekar R, Nicolaides KH. Ultrasonographic estimation of fetal weight: development of new model and assessment of performance of previous models. Ultrasound Obstet Gynecol. 2018 Jul;52(1):35-43. doi: 10.1002/uog.19066. Epub 2018 Jun 3. PMID: 29611251.
214 Barel O, Vaknin Z, Tovbin J, Herman A, Maymon R. Assessment of the accuracy of multiple sonographic fetal weight estimation formulas: a 10-year experience from a single center. J Ultrasound Med. 2013 May;32(5):815-23. doi: 10.7863/ultra.32.5.815. PMID: 23620324.
215 Melamed N, Yogev Y, Meizner I, Mashiach R, Bardin R, Ben-Haroush A. Sonographic fetal weight estimation: which model should be used? J Ultrasound Med. 2009 May;28(5):617-29. doi: 10.7863/jum.2009.28.5.617. PMID: 19389901.

216 Hoopmann M, Kagan KO, Sauter A, Abele H, Wagner P. Comparison of Errors of 35 Weight Estimation Formulae in a Standard Collective. Geburtshilfe Frauenheilkd. 2016 Nov;76(11):1172-1179. doi: 10.1055/s-0042-118598. PMID: 27904167; PMCID: PMC5123881.
217 Ma J, Cheng D, Zhang Z, Cai B, Xu X. Evaluating the accuracy of sonographic fetal weight estimations using the Hadlock IV formula in a Chinese population. Quant Imaging Med Surg. 2023 Jun 1;13(6):3726-3734. doi: 10.21037/qims-22-778. Epub 2023 Apr 3. PMID: 37284125; PMCID: PMC10240036.
218 Milner J, Arezina J. The accuracy of ultrasound estimation of fetal weight in comparison to birth weight: A systematic review. Ultrasound. 2018 Feb;26(1):32-41. doi: 10.1177/1742271X17732807. Epub 2018 Feb 7. PMID: 29456580; PMCID: PMC5810856.
219 The Binary Cellular Biology of Human Growth: III. Identifying, Comprehending, and Ameliorating Normal and Abnormal Fetal Size and Growth
220 The Binary Cellular Biology of Human Growth: I. Calculating the Growth of the Human Fetus from Conception Onward
221 Michaelson J. The role of molecular discreteness in normal and cancerous growth. Anticancer Res. 1999 Nov-Dec;19(6A):4853-67. PMID: 10697599
222 Michaelson, J. Cell Selection in the Genesis of Multicellular Organization Lab Invest. 69: 136-150 1993
223 Michaelson J. Cell selection in development. Biological Reviews 1987; 62:115-139.
224 Bommarito PA, Cantonwine DE, Stevens DR, Welch BM, Davalos AD, Zhao S, McElrath TF, Ferguson KK. Fetal growth trajectories of babies born large-for-gestational age in the LIFECODES Fetal Growth Study. Am J Obstet Gynecol. 2023 Mar;228(3):340.e1-340.e20. doi: 10.1016/j.ajog.2022.10.006. Epub 2022 Oct 12. PMID: 36241081; PMCID: PMC9974610.
225 Boyer TM, Bommarito PA, Welch BM, Meeker JD, James-Todd T, Cantonwine DE, McElrath TF, Ferguson KK. Maternal exposure to phthalates and total gestational Weight-Gain Rate Rate in the LIFECODES birth cohort. Reprod Toxicol. 2023 Apr;117:108354. doi: 10.1016/j.reprotox.2023.108354. Epub 2023 Feb 24. PMID: 36841368; PMCID: PMC10073336.
226 Bommarito PA, Friedman A, Welch BM, Cantonwine DE, Ospina M, Calafat AM, Meeker JD, McElrath TF, Ferguson KK. Temporal trends and predictors of gestational exposure to organophosphate ester flame retardants and plasticizers. Environ Int. 2023 Oct;180:108194. doi: 10.1016/j.envint.2023.108194. Epub 2023 Sep 7. PMID: 37708814; PMCID: PMC10591987.
227 Bommarito PA, Cantonwine DE, Stevens DR, Welch BM, Davalos AD, Zhao S, McElrath TF, Ferguson KK. An application of group-based trajectory modeling to define fetal growth phenotypes among small-for-gestational-age births in the LIFECODES Fetal Growth Study. Am J Obstet Gynecol. 2023 Mar;228(3):334.e1-334.e21. doi: 10.1016/j.ajog.2022.08.041. Epub 2022 Aug 24. PMID: 36027952; PMCID: PMC9950285.
228 Binary Cellular Biology: I. Calculating the Growth of the Human Fetus from Conception Onward
229 Cohen JM, Hutcheon JA, Kramer MS, Joseph KS, Abenhaim H, Platt RW. Influence of ultrasound-to-delivery interval and maternal-fetal characteristics on validity of estimated fetal weight. Ultrasound Obstet Gynecol. 2010 Apr;35(4):434-41. doi: 10.1002/uog.7506. PMID: 20069655.
230 Monier I, Ego A, Benachi A, Ancel PY, Goffinet F, Zeitlin J. Comparison of the Hadlock and INTERGROWTH formulas for calculating estimated fetal weight in a preterm population in France. Am J Obstet Gynecol. 2018 Nov;219(5):476.e1-476.e12. doi: 10.1016/j.ajog.2018.08.012. Epub 2018 Aug 14. PMID: 30118693.
231 Anderson NG, Jolley IJ, Wells JE. Sonographic estimation of fetal weight: comparison of bias, precision and consistency using 12 different formulae. Ultrasound Obstet Gynecol. 2007 Aug;30(2):173-9. doi: 10.1002/uog.4037. PMID: 17557378.
232 Barel O, Vaknin Z, Tovbin J, Herman A, Maymon R. Assessment of the accuracy of multiple sonographic fetal weight estimation formulas: a 10-year experience from a single center. J Ultrasound Med. 2013 May;32(5):815-23. doi: 10.7863/ultra.32.5.815. PMID: 23620324.
233 Hammami A, Mazer Zumaeta A, Syngelaki A, Akolekar R, Nicolaides KH. Ultrasonographic estimation of fetal weight: development of new model and assessment of performance of previous models. Ultrasound Obstet Gynecol. 2018 Jul;52(1):35-43. doi: 10.1002/uog.19066. Epub 2018 Jun 3. PMID: 29611251.
234 Hiwale S, Misra H, Ulman S. Fetal weight estimation by ultrasound: development of Indian population-based models. Ultrasonography. 2019 Jan;38(1):50-57. doi: 10.14366/usg.18004. Epub 2018 Apr 14. PMID: 29976038; PMCID: PMC6323311.
235 Hoopmann M, Kagan KO, Sauter A, Abele H, Wagner P. Comparison of Errors of 35 Weight Estimation Formulae in a Standard Collective. Geburtshilfe Frauenheilkd. 2016 Nov;76(11):1172-1179. doi: 10.1055/s-0042-118598. PMID: 27904167; PMCID: PMC5123881
236 Weiss C, Oppelt P, Mayer RB. Disadvantages of a weight estimation formula for macrosomic fetuses: the Hart formula from a clinical perspective. Arch Gynecol Obstet. 2018 Dec;298(6):1101-1106. doi: 10.1007/s00404-018-4917-z. Epub 2018 Oct 4. PMID: 30284620; PMCID: PMC6244680.
237 Melamed N, Yogev Y, Meizner I, Mashiach R, Bardin R, Ben-Haroush A. Sonographic fetal weight estimation: which model should be used? J Ultrasound Med. 2009 May;28(5):617-29. doi: 10.7863/jum.2009.28.5.617. PMID: 19389901.
238 Sereke SG, Omara RO, Bongomin F, Nakubulwa S, Kisembo HN. Prospective verification of sonographic fetal weight estimators among term parturients in Uganda. BMC Pregnancy Childbirth. 2021 Mar 4;21(1):175. doi: 10.1186/s12884-021-03645-4. PMID: 33663407; PMCID: PMC7934251.
239 Simon NV, Levisky JS, Shearer DM, O'Lear MS, Flood JT. Influence of fetal growth patterns on sonographic estimation of fetal weight. J Clin Ultrasound. 1987 Jul-Aug;15(6):376-83. doi: 10.1002/jcu.1870150604. PMID: 3134435.

240 Sabbagha RE, Minogue J, Tamura RK, Hungerford SA. Estimation of birth weight by use of ultrasonographic formulas targeted to large-, appropriate-, and small-for-gestational-age fetuses. Am J Obstet Gynecol. 1989 Apr;160(4):854-60; discussion 860-2. doi: 10.1016/0002-9378(89)90301-3. PMID: 2653039.
241 Weiss C, Oppelt P, Mayer RB. Disadvantages of a weight estimation formula for macrosomic fetuses: the Hart formula from a clinical perspective. Arch Gynecol Obstet. 2018 Dec;298(6):1101-1106. doi: 10.1007/s00404-018-4917-z. Epub 2018 Oct 4. PMID: 30284620; PMCID: PMC6244680.
242 Li C, Peng Y, Zhang B, Ji W, Li L, Gong J, Xia W, Li Y, Jin S, Song R, Wang Y, Xu S. Birth weight prediction models for the different gestational age stages in a Chinese population. Sci Rep. 2019 Jul 25;9(1):10834. doi: 10.1038/s41598-019-47056-0. PMID: 31346206; PMCID: PMC6658529.
243 Siemer J, Egger N, Hart N, Meurer B, Müller A, Dathe O, Goecke T, Schild RL. Fetal weight estimation by ultrasound: comparison of 11 different formulae and examiners with differing skill levels. Ultraschall Med. 2008 Apr;29(2):159-64. doi: 10.1055/s-2007-963165. Epub 2007 Jun 29. PMID: 17602369.
244 Sharma R, Bhoil R, Dogra P, Kaushal Sharma A. Accuracy and reliability of ultrasound estimation of fetal weight in women with a singleton term pregnancy. International Journal of Reproduction, Contraception, Obstetrics and Gynecology Vol. 9 No. 1 (2020): January 2020 https://doi.org/10.18203/2320-1770.ijrcog20196042
245 Townsend RR, Filly RA, Callen PW, Laros RK. Factors affecting prenatal sonographic estimation of weight in extremely low birthweight infants. J Ultrasound Med. 1988 Apr;7(4):183-7. doi: 10.7863/jum.1988.7.4.183. PMID: 3285021.
246 Dudley NJ. The management of error in ultrasound fetal growth monitoring. Ultrasound. 2021 Feb;29(1):4-9. doi: 10.1177/1742271X20945749. Epub 2020 Aug 3. PMID: 33552222; PMCID: PMC7844476.
247 Dudley NJ, Jindal S, Varley H. An evaluation of systematic and random errors in ultrasound estimated fetal weight during serial ultrasound. Ultrasound. 2023 Nov;31(4):259-265. doi: 10.1177/1742271X221139796. Epub 2023 Jan 4. PMID: 37929245; PMCID: PMC10621486.
248 Yaqub M, Kelly B, Stobart H, Napolitano R, Noble JA, Papageorghiou AT. Quality-improvement program for ultrasound-based fetal anatomy screening using large-scale clinical audit. Ultrasound Obstet Gynecol. 2019 Aug;54(2):239-245. doi: 10.1002/uog.20144. PMID: 30302849; PMCID: PMC6771606.
249 Badr DA, Cannie MM, Kadji C, Kang X, Carlin A, Jani JC. Performance of fetal ultrasound and magnetic resonance imaging in predicting birthweight according to the test-to-delivery interval: A cohort study. Eur J Obstet Gynecol Reprod Biol. 2024 Jan;292:138-146. doi: 10.1016/j.ejogrb.2023.11.025. Epub 2023 Nov 20. PMID: 38006818.
250 Grantz KL, Lee W, Mack LM, Sanz Cortes M, Goncalves LF, Espinoza J, Newman RB, Grobman WA, Wapner RJ, Fuchs K, D'Alton ME, Skupski DW, Owen J, Sciscione A, Wing DA, Nageotte MP, Ranzini AC, Chien EK, Craigo S, Sherman S, Gore-Langton RE, He D, Tekola-Ayele F, Zhang C, Grewal J, Chen Z. Multiethnic growth standards for fetal body composition and organ volumes derived from 3D ultrasonography. Am J Obstet Gynecol. 2024 Jun 3:S0002-9378(24)00658-6. doi: 10.1016/j.ajog.2024.05.049. Epub ahead of print. PMID: 38838912.
251 Grantz KL, Lee W, Chen Z, Hinkle S, Mack L, Cortes MS, Goncalves LF, Espinoza J, Gore-Langton RE, Sherman S, He D, Zhang C, Grewal J. The NICHD Fetal 3D Study: A Pregnancy Cohort Study of Fetal Body Composition and Volumes. Am J Epidemiol. 2024 Apr 8;193(4):580-595. doi: 10.1093/aje/kwad210. PMID: 37946325.
252 Simcock IC, Arthurs OJ, Hutchinson JC, Sebire NJ, Jacques TS, Sekar T, Shelmerdine SC; GOSH Pathology. Impact of non-invasive post-mortem micro-CT imaging on a fetal autopsy service: a single centre retrospective study. Clin Radiol. 2024 Jul 6:S0009-9260(24)00338-6. doi: 10.1016/j.crad.2024.06.015. Epub ahead of print. PMID: 39068113.
253 Shelmerdine SC, Hutchinson JC, Arthurs OJ, Sebire NJ. Latest developments in post-mortem foetal imaging. Prenat Diagn. 2020 Jan;40(1):28-37. doi: 10.1002/pd.5562. Epub 2019 Dec 3. PMID: 31525275.
254 Michaelson JS, Chen LL, Bush D, Fong A, Smith B, Younger J. Improved web-based calculators for predicting breast carcinoma outcomes. Breast Cancer Res Treat. 2011 Aug;128(3):827-35
255 **PAPER III**
256 O'Farrell PH. Quiescence: early evolutionary origins and universality do not imply uniformity. Philos Trans R Soc Lond B Biol Sci. 2011 Dec 27;366 (1584):3498-507.
257 Hartwell LH- Nobel Lecture: Yeast and Cancer. Nobelprize.org. Nobel Media AB 2014. Web. 13 Sep 2015. <http://www.nobelprize.org/nobel_prizes/medicine/laureates/2001/hartwell-lecture.html>
258 Bertoli C, Skotheim JM, de Bruin RA. Control of cell cycle transcription during G1 and S phases. Nat Rev Mol Cell Biol. 2013 Aug;14(8):518-28. doi: 10.1038/nrm3629.
259 Michaelson J. Cell selection in development. Biological Reviews 1987; 62:115-139.
260 Michaelson, J. Cell Selection in the Genesis of Multicellular Organization Lab Invest. 69: 136-150 1993
261 Michaelson J. The role of molecular discreteness in normal and cancerous growth. Anticancer Res. 1999 Nov-Dec;19(6A):4853-67. PMID: 10697599.
262 The Binary Cellular Biology of Human Growth: I. Calculating the Growth of the Human Fetus from Conception Onward
263 The Binary Cellular Biology of Human Growth: II. Calculating the Size and Age of the Human Fetus from the Size of its Parts
264 Bommarito PA, Cantonwine DE, Stevens DR, Welch BM, Davalos AD, Zhao S, McElrath TF, Ferguson KK. Fetal growth trajectories of babies born large-for-gestational age in the LIFECODES Fetal Growth Study. Am J Obstet Gynecol. 2023 Mar;228(3):340.e1-340.e20. doi: 10.1016/j.ajog.2022.10.006. Epub 2022 Oct 12. PMID: 36241081; PMCID: PMC9974610.

265 Boyer TM, Bommarito PA, Welch BM, Meeker JD, James-Todd T, Cantonwine DE, McElrath TF, Ferguson KK. Maternal exposure to phthalates and total gestational Weight-Gain Rate Rate in the LIFECODES birth cohort. Reprod Toxicol. 2023 Apr;117:108354. doi: 10.1016/j.reprotox.2023.108354. Epub 2023 Feb 24. PMID: 36841368; PMCID: PMC10073336.
266 Bommarito PA, Friedman A, Welch BM, Cantonwine DE, Ospina M, Calafat AM, Meeker JD, McElrath TF, Ferguson KK. Temporal trends and predictors of gestational exposure to organophosphate ester flame retardants and plasticizers. Environ Int. 2023 Oct;180:108194. doi: 10.1016/j.envint.2023.108194. Epub 2023 Sep 7. PMID: 37708814; PMCID: PMC10591987.
267 Bommarito PA, Cantonwine DE, Stevens DR, Welch BM, Davalos AD, Zhao S, McElrath TF, Ferguson KK. An application of group-based trajectory modeling to define fetal growth phenotypes among small-for-gestational-age births in the LIFECODES Fetal Growth Study. Am J Obstet Gynecol. 2023 Mar;228(3):334.e1-334.e21. doi: 10.1016/j.ajog.2022.08.041. Epub 2022 Aug 24. PMID: 36027952; PMCID: PMC9950285.
268 Grantz KL, Kim S, Grobman WA, Newman R, Owen J, Skupski D, Grewal J, Chien EK, Wing DA, Wapner RJ, Ranzini AC, Nageotte MP, Hinkle SN, Pugh S, Li H, Fuchs K, Hediger M, Buck Louis GM, Albert PS. Fetal growth velocity: the NICHD fetal growth studies. Am J Obstet Gynecol. 2018 Sep;219(3):285.e1-285.e36. doi: 10.1016/j.ajog.2018.05.016. Epub 2018 May 24. PMID: 29803819; PMCID: PMC7035912.
269 Wu T, Gong X, Zhao Y, Zhang L, You Y, Wei H, Zuo X, Zhou Y, Xing X, Meng Z, Lv Q, Liu Z, Zhang J, Hu L, Li J, Li L, Chen C, Liu C, Sun G, Liu A, Chen J, Lv Y, Wang X, Wei Y. Fetal growth velocity references from a Chinese population-based fetal growth study. BMC Pregnancy Childbirth. 2021 Oct 9;21(1):688. doi: 10.1186/s12884-021-04149-x. PMID: 34627184; PMCID: PMC8501691.
270 Ohuma EO, Villar J, Feng Y, Xiao L, Salomon L, Barros FC, Cheikh Ismail L, Stones W, Jaffer Y, Oberto M, Noble JA, Gravett MG, Wu Q, Victora CG, Lambert A, Di Nicola P, Purwar M, Bhutta ZA, Kennedy SH, Papageorghiou AT; International Fetal and Newborn Growth Consortium for the 21st Century (INTERGROWTH-21st Project). Fetal growth velocity standards from the Fetal Growth Longitudinal Study of the INTERGROWTH-21st Project. Am J Obstet Gynecol. 2021 Feb;224(2):208.e1-208.e18. doi: 10.1016/j.ajog.2020.07.054. Epub 2020 Aug 5. PMID: 32768431; PMCID: PMC7858163.
271 Partap U, Sovio U, Smith GC. Fetal Growth and the Risk of Spontaneous Preterm Birth in a Prospective Cohort Study of Nulliparous Women. Am J Epidemiol. 2016 Jul 15;184(2):110-9. doi: 10.1093/aje/kwv345. Epub 2016 Jul 1. PMID: 27370790; PMCID: PMC5860261.
272 Mongelli M, Benzie R, Condous G. Average fetal weekly weight gain: a novel measure of fetal growth velocity. J Matern Fetal Neonatal Med. 2016;29(4):676-9. doi: 10.3109/14767058.2015.1015981. Epub 2015 Feb 25. PMID: 25714480..
273 Del Monte U Does the cell number 10^9 still really fit one gram of tumor tissue? Cell Cycle 8:3, 505-506; 1 February 2009
274 Hsu ST, Hsieh CJ, Chen HW, Jeng SF, Wu HC, Chen MH, Chen CY, Chou HC, Tsao PN, Chen PC, Hsieh WS. Nationwide Birth Weight and Gestational Age-specific Neonatal Mortality Rate in Taiwan. Pediatr Neonatol. 2015 Jun;56(3):149-58. doi: 10.1016/j.pedneo.2014.07.006. Epub 2014 Oct 30. PMID: 25440779.
275 Battaglia FC, Lubchenco LO. A practical classification of newborn infants by weight and gestational age. J Pediatr. 1967 Aug;71(2):159-63. doi: 10.1016/s0022-3476(67)80066-0. PMID: 6029463.
276 Grantz KL, Kim S, Grobman WA, Newman R, Owen J, Skupski D, Grewal J, Chien EK, Wing DA, Wapner RJ, Ranzini AC, Nageotte MP, Hinkle SN, Pugh S, Li H, Fuchs K, Hediger M, Buck Louis GM, Albert PS. Fetal growth velocity: the NICHD fetal growth studies. Am J Obstet Gynecol. 2018 Sep;219(3):285.e1-285.e36. doi: 10.1016/j.ajog.2018.05.016. Epub 2018 May 24. PMID: 29803819; PMCID: PMC7035912
277 Ohuma EO, Villar J, Feng Y, Xiao L, Salomon L, Barros FC, Cheikh Ismail L, Stones W, Jaffer Y, Oberto M, Noble JA, Gravett MG, Wu Q, Victora CG, Lambert A, Di Nicola P, Purwar M, Bhutta ZA, Kennedy SH, Papageorghiou AT; International Fetal and Newborn Growth Consortium for the 21st Century (INTERGROWTH-21st Project). Fetal growth velocity standards from the Fetal Growth Longitudinal Study of the INTERGROWTH-21st Project. Am J Obstet Gynecol. 2021 Feb;224(2):208.e1-208.e18. doi: 10.1016/j.ajog.2020.07.054. Epub 2020 Aug 5. PMID: 32768431; PMCID: PMC7858163
278 Yin XG, Wang P, Zhou MT, Li DQ, Tao RX, Tao FB, Wang Y, Zhu P. Timing of gestational diabetes diagnosis, gestational weight gains and offspring growth trajectory: a prospective birth cohort study. BMC Pregnancy Childbirth. 2023 Sep 7;23(1):642. doi: 10.1186/s12884-023-05954-2. PMID: 37679668; PMCID: PMC10483803.
279 Andersen AS, Linneberg Rathcke S, Tang Christensen T, Sørensen A. The association between second trimester ultrasound fetal biometrics and gestational diabetes. Eur J Obstet Gynecol Reprod Biol. 2022 Sep;276:139-143. doi: 10.1016/j.ejogrb.2022.07.015. Epub 2022 Jul 21. PMID: 35905541.
280 McIntyre HD, Catalano P, Zhang C, Desoye G, Mathiesen ER, Damm P. Gestational diabetes mellitus. Nat Rev Dis Primers. 2019 Jul 11;5(1):47. doi: 10.1038/s41572-019-0098-8. PMID: 31296866.
281 Lovrić B, Šijanović S, Zmijanović I, Jurić G, Juras J. ULTRASOUND DIAGNOSIS OF MACROSOMIA AMONG WOMEN WITH GESTATIONAL DIABETES - REVIEW OF THE LITERATURE. Acta Clin Croat. 2022 Mar;61(1):95-106. doi: 10.20471/acc.2022.61.01.12. PMID: 36398082; PMCID: PMC9616028
282 Bruckner TA, Catalano R. Selection in utero and population health: Theory and typology of research. SSM Popul Health. 2018 Jun 2;5:101-113. doi: 10.1016/j.ssmph.2018.05.010. Erratum in: SSM Popul Health. 2020 Dec 10;12:100714. doi: 10.1016/j.ssmph.2020.100714. PMID: 29928686; PMCID: PMC6008283.
283 Baird DD. The gestational timing of pregnancy loss: adaptive strategy? Am J Hum Biol. 2009 Nov-Dec;21(6):725-7. doi: 10.1002/ajhb.20935. PMID: 19408286.

284 Magnus MC, Wilcox AJ, Morken NH, Weinberg CR, Håberg SE. Role of maternal age and pregnancy history in risk of miscarriage: prospective register based study. BMJ. 2019 Mar 20;364:l869. doi: 10.1136/bmj.l869. PMID: 30894356; PMCID: PMC6425455.
285 Rokas A, Mesiano S, Tamam O, LaBella A, Zhang G, Muglia L. Developing a theoretical evolutionary framework to solve the mystery of parturition initiation. Elife. 2020 Dec 31;9:e58343. doi: 10.7554/eLife.58343. PMID: 33380346; PMCID: PMC7775106.
286 Committee on Practice Bulletins—Obstetrics and the American Institute of Ultrasound in Medicine. Practice Bulletin No. 175: Ultrasound in Pregnancy. Obstet Gynecol. 2016 Dec;128(6):e241-e256. doi: 10.1097/AOG.0000000000001815. PMID: 27875472.
287 International Society of Ultrasound in Obstetrics and Gynecology; Bilardo CM, Chaoui R, Hyett JA, Kagan KO, Karim JN, Papageorghiou AT, Poon LC, Salomon LJ, Syngelaki A, Nicolaides KH. ISUOG Practice Guidelines (updated): performance of 11-14-week ultrasound scan. Ultrasound Obstet Gynecol. 2023 Jan;61(1):127-143. doi: 10.1002/uog.26106. PMID: 36594739.
288 Salomon LJ, Alfirevic Z, Berghella V, Bilardo CM, Chalouhi GE, Da Silva Costa F, Hernandez-Andrade E, Malinger G, Munoz H, Paladini D, Prefumo F, Sotiriadis A, Toi A, Lee W. ISUOG Practice Guidelines (updated): performance of the routine mid-trimester fetal ultrasound scan. Ultrasound Obstet Gynecol. 2022 Jun;59(6):840-856. doi: 10.1002/uog.24888. Epub 2022 May 20. Erratum in: Ultrasound Obstet Gynecol. 2022 Oct;60(4):591. doi: 10.1002/uog.26067. PMID: 35592929.
289 Khalil A, Sotiriadis A, D'Antonio F, Da Silva Costa F, Odibo A, Prefumo F, Papageorghiou AT, Salomon LJ. ISUOG Practice Guidelines: performance of third-trimester obstetric ultrasound scan. Ultrasound Obstet Gynecol. 2024 Jan;63(1):131-147. doi: 10.1002/uog.27538. PMID: 38166001.
290 Peahl AF, Howell JD. The evolution of prenatal care delivery guidelines in the United States. Am J Obstet Gynecol. 2021 Apr;224(4):339-347. doi: 10.1016/j.ajog.2020.12.016. Epub 2020 Dec 13. PMID: 33316276; PMCID: PMC9745905.
291 WHO recommendations on antenatal care for a positive pregnancy experience 28 November 2016 https://www.who.int/publications/i/item/9789241549912
292 Dr. Sarah J. Kilpatrick MD PhD FACOG, LuAnn Papile MD FAAP, George A. Macones MD FACOG , Kristi L. Watterberg MD FAAP , AAP Committee on Fetus and Newborn , ACOG Committee on Obstetric Practice , Guidelines for Perinatal Care Eighth Edition ISBN-10 : 1610020871
293 Michaelson, J, Halpern, E, Kopans, D. A Computer Simulation Method For Estimating The Optimal Intervals For Breast Cancer Screening. Radiology. 212:551-560 1999
294 Oeffinger KC, Fontham ET, Etzioni R, Herzig A, Michaelson JS, Shih YC, Walter LC, Church TR, Flowers CR, LaMonte SJ, Wolf AM, DeSantis C, Lortet-Tieulent J, Andrews K, Manassaram-Baptiste D, Saslow D, Smith RA, Brawley OW, Wender R; American Cancer Society. Breast Cancer Screening for Women at Average Risk: 2015 Guideline Update From the American Cancer Society. JAMA. 2015 Oct 20;314(15):1599-614. doi: 10.1001/jama.2015.12783. Erratum in: JAMA. 2016 Apr 5;315(13):1406. doi: 0.1001/jama.2016.3404. PMID: 26501536; PMCID: PMC4831582.
295 Figueras F, Gratacós E. Update on the diagnosis and classification of fetal growth restriction and proposal of a stage-based management protocol. Fetal Diagn Ther 2014;36: 86–98.
296 Lees CC, Romero R, Stampalija T, Dall'Asta A, DeVore GA, Prefumo F, Frusca T, Visser GHA, Hobbins JC, Baschat AA, Bilardo CM, Galan HL, Campbell S, Maulik D, Figueras F, Lee W, Unterscheider J, Valensise H, Da Silva Costa F, Salomon LJ, Poon LC, Ferrazzi E, Mari G, Rizzo G, Kingdom JC, Kiserud T, Hecher K. Clinical Opinion: The diagnosis and management of suspected fetal growth restriction: an evidence-based approach. Am J Obstet Gynecol. 2022 Mar;226(3):366-378. doi: 10.1016/j.ajog.2021.11.1357. Epub 2022 Jan 10. PMID: 35026129; PMCID: PMC9125563.
297 Warrington NM, Beaumont RN, Horikoshi M, Day FR, Helgeland Ø, Laurin C, Bacelis J, Peng S, Hao K, Feenstra B, Wood AR, Mahajan A, Tyrrell J, Robertson NR, Rayner NW, Qiao Z, Moen GH, Vaudel M, Marsit CJ, Chen J, Nodzenski M, Schnurr TM, Zafarmand MH, Bradfield JP, Grarup N, Kooijman MN, Li-Gao R, Geller F, Ahluwalia TS, Paternoster L, Rueedi R, Huikari V, Hottenga JJ, Lyytikäinen LP, Cavadino A, Metrustry S, Cousminer DL, Wu Y, Thiering E, Wang CA, Have CT, Vilor-Tejedor N, Joshi PK, Painter JN, Ntalla I, Myhre R, Pitkänen N, van Leeuwen EM, Joro R, Lagou V, Richmond RC, Espinosa A, Barton SJ, Inskip HM, Holloway JW, Santa-Marina L, Estivill X, Ang W, Marsh JA, Reichetzeder C, Marullo L, Hocher B, Lunetta KL, Murabito JM, Relton CL, Kogevinas M, Chatzi L, Allard C, Bouchard L, Hivert MF, Zhang G, Muglia LJ, Heikkinen J; EGG Consortium; Morgen CS, van Kampen AHC, van Schaik BDC, Mentch FD, Langenberg C, Luan J, Scott RA, Zhao JH, Hemani G, Ring SM, Bennett AJ, Gaulton KJ, Fernandez-Tajes J, van Zuydam NR, Medina-Gomez C, de Haan HG, Rosendaal FR, Kutalik Z, Marques-Vidal P, Das S, Willemsen G, Mbarek H, Müller-Nurasyid M, Standl M, Appel EVR, Fonvig CE, Trier C, van Beijsterveldt CEM, Murcia M, Bustamante M, Bonas-Guarch S, Hougaard DM, Mercader JM, Linneberg A, Schraut KE, Lind PA, Medland SE, Shields BM, Knight BA, Chai JF, Panoutsopoulou K, Bartels M, Sánchez F, Stokholm J, Torrents D, Vinding RK, Willems SM, Atalay M, Chawes BL, Kovacs P, Prokopenko I, Tuke MA, Yaghootkar H, Ruth KS, Jones SE, Loh PR, Murray A, Weedon MN, Tönjes A, Stumvoll M, Michaelsen KF, Eloranta AM, Lakka TA, van Duijn CM, Kiess W, Körner A, Niinikoski H, Pahkala K, Raitakari OT, Jacobsson B, Zeggini E, Dedoussis GV, Teo YY, Saw SM, Montgomery GW, Campbell H, Wilson JF, Vrijkotte TGM, Vrijheid M, de Geus EJCN, Hayes MG, Kadarmideen HN, Holm JC, Beilin LJ, Pennell CE, Heinrich J, Adair LS, Borja JB, Mohlke KL, Eriksson JG, Widén EE, Hattersley AT, Spector TD, Kähönen M, Viikari JS, Lehtimäki T, Boomsma DI, Sebert S, Vollenweider P, Sørensen TIA, Bisgaard H, Bønnelykke K, Murray JC, Melbye M, Nohr EA, Mook-Kanamori DO, Rivadeneira F, Hofman A, Felix JF, Jaddoe VWV, Hansen T, Pisinger C, Vaag AA, Pedersen O, Uitterlinden AG, Järvelin MR, Power C, Hyppönen E, Scholtens DM, Lowe WL Jr, Davey Smith G, Timpson NJ, Morris AP, Wareham NJ, Hakonarson H, Grant SFA, Frayling TM, Lawlor DA, Njølstad PR, Johansson S, Ong KK, McCarthy MI, Perry JRB, Evans DM, Freathy RM. Maternal and fetal genetic effects on birth weight and their relevance to cardio-

metabolic risk factors. Nat Genet. 2019 May;51(5):804-814. doi: 10.1038/s41588-019-0403-1. Epub 2019 May 1. PMID: 31043758; PMCID: PMC6522365.
298 Juliusdottir T, Steinthorsdottir V, Stefansdottir L, Sveinbjornsson G, Ivarsdottir EV, Thorolfsdottir RB, Sigurdsson JK, Tragante V, Hjorleifsson KE, Helgadottir A, Frigge ML, Thorgeirsson G, Benediktsson R, Sigurdsson EL, Arnar DO, Steingrimsdottir T, Jonsdottir I, Holm H, Gudbjartsson DF, Thorleifsson G, Thorsteinsdottir U, Stefansson K. Distinction between the effects of parental and fetal genomes on fetal growth. Nat Genet. 2021 Aug;53(8):1135-1142. doi: 10.1038/s41588-021-00896-x. Epub 2021 Jul 19. PMID: 34282336.
299 Solé-Navais P, Flatley C, Steinthorsdottir V, Vaudel M, Juodakis J, Chen J, Laisk T, LaBella AL, Westergaard D, Bacelis J, Brumpton B, Skotte L, Borges MC, Helgeland Ø, Mahajan A, Wielscher M, Lin F, Briggs C, Wang CA, Moen GH, Beaumont RN, Bradfield JP, Abraham A, Thorleifsson G, Gabrielsen ME, Ostrowski SR, Modzelewska D, Nohr EA, Hypponen E, Srivastava A, Talbot O, Allard C, Williams SM, Menon R, Shields BM, Sveinbjornsson G, Xu H, Melbye M, Lowe W Jr, Bouchard L, Oken E, Pedersen OB, Gudbjartsson DF, Erikstrup C, Sørensen E; Early Growth Genetics Consortium; Estonian Biobank Research Team; Danish Blood Donor Study Genomic Consortium; Lie RT, Teramo K, Hallman M, Juliusdottir T, Hakonarson H, Ullum H, Hattersley AT, Sletner L, Merialdi M, Rifas-Shiman SL, Steingrimsdottir T, Scholtens D, Power C, West J, Nyegaard M, Capra JA, Skogholt AH, Magnus P, Andreassen OA, Thorsteinsdottir U, Grant SFA, Qvigstad E, Pennell CE, Hivert MF, Hayes GM, Jarvelin MR, McCarthy MI, Lawlor DA, Nielsen HS, Mägi R, Rokas A, Hveem K, Stefansson K, Feenstra B, Njolstad P, Muglia LJ, Freathy RM, Johansson S, Zhang G, Jacobsson B. Genetic effects on the timing of parturition and links to fetal birth weight. Nat Genet. 2023 Apr;55(4):559-567. doi: 10.1038/s41588-023-01343-9. Epub 2023 Apr 3. Erratum in: Nat Genet. 2023 Jul;55(7):1250. doi: 10.1038/s41588-023-01412-z. PMID: 37012456; PMCID: PMC10101852.
300 Ray PF, Conaghan J, Winston RM, Handysd metabolic activity in male human preimplantation embryos following in vitro fertilization. J Reprod Fertil. 1995 May;104(1):165-71. doi: 10.1530/jrf.0.1040165. PMID: 7636798.
301 Kiserud T, Piaggio G, Carroli G, et al. The World Health Organization fetal growth charts: A multinational longitudinal study of ultrasound biometric measurements and estimated fetal weight. PLoS Med 2017;14:e1002220.
302 Brown ER, Giussani DA. Cause of fetal growth restriction during high-altitude pregnancy. iScience. 2024 Apr 8;27(5):109702. doi: 10.1016/j.isci.2024.109702. PMID: 38694168; PMCID: PMC11061758.
303 Moore LG. How hypoxia slows fetal growth: insights from high altitude. Pediatr Res. 2022 Jan;91(1):17-18. doi: 10.1038/s41390-021-01784-0. Epub 2021 Oct 14. PMID: 34650197; PMCID: PMC8776573.
304 Boulvain M, Thornton JG. Induction of labour at or near term for suspected fetal macrosomia. Cochrane Database Syst Rev. 2023 Mar 8;3(3):CD000938. doi: 10.1002/14651858.CD000938.pub3. PMID: 36884238; PMCID: PMC9995561.
305 Macrosomia: ACOG Practice Bulletin, Number 216. Obstet Gynecol. 2020 Jan;135(1):e18-e35. doi: 10.1097/AOG.0000000000003606. PMID: 31856124.
306 Ewington LJ, Gardosi J, Lall R, Underwood M, Fisher JD, Wood S, Griffin R, Harris K, Bick D, Booth K, Brown J, Butler E, Fowler K, Williams M, Deshpande S, Gornall A, Dewdney J, Hillyer K, Gates S, Jones C, Mistry H, Petrou S, Slowther AM, Willis A, Quenby S. Induction of labour for predicted macrosomia: study protocol for the 'Big Baby' randomised controlled trial. BMJ Open. 2022 Nov 11;12(11):e058176. doi: 10.1136/bmjopen-2021-058176. PMID: 36368760; PMCID: PMC9660609.
307 Smith GC, Moraitis AA, Wastlund D, Thornton JG, Papageorghiou A, Sanders J, Heazell AE, Robson SC, Sovio U, Brocklehurst P, Wilson EC. Universal late pregnancy ultrasound screening to predict adverse outcomes in nulliparous women: a systematic review and cost-effectiveness analysis. Health Technol Assess. 2021 Feb;25(15):1-190. doi: 10.3310/hta25150. PMID: 33656977; PMCID: PMC7958245.
308 Sovio U, White IR, Dacey A, Pasupathy D, Smith GCS. Screening for fetal growth restriction with universal third trimester ultrasonography in nulliparous women in the Pregnancy Outcome Prediction (POP) study: a prospective cohort study. Lancet 2015;386:2089–97.
309 Inducing labour. London: National Institute for Health and Care Excellence (NICE); 2021 Nov 4. PMID: 35438865.
310 Moraitis AA, Shreeve N, Sovio U, Brocklehurst P, Heazell AEP, Thornton JG, Robson SC, Papageorghiou A, Smith GC. Universal third-trimester ultrasonic screening using fetal macrosomia in the prediction of adverse perinatal outcome: A systematic review and meta-analysis of diagnostic test accuracy. PLoS Med. 2020 Oct 13;17(10):e1003190. doi: 10.1371/journal.pmed.1003190. PMID: 33048935; PMCID: PMC7553291.
311 Manuck TA, Gyamfi-Bannerman C, Saade G. What now? A critical evaluation of over 20 years of clinical and research experience with 17-alpha hydroxyprogesterone caproate for recurrent preterm birth prevention. Am J Obstet Gynecol MFM. 2023 Oct;5(10):101108. doi: 10.1016/j.ajogmf.2023.101108. Epub 2023 Jul 30. PMID: 37527737; PMCID: PMC10591827.
312 Manuck TA. Refining Pharmacologic Research to Prevent and Treat Spontaneous Preterm Birth. Front Pharmacol. 2017 Mar 15;8:118. doi: 10.3389/fphar.2017.00118. PMID: 28360854; PMCID: PMC5350137.
313 Spiliopoulos D, Economides DL. Early fetal growth in progesterone-treated IVF pregnancies. Arch Gynecol Obstet. 2016 Jul;294(1):63-9. doi: 10.1007/s00404-015-3951-3. Epub 2015 Nov 13. PMID: 26563314.
314 Hadlock FP, Harrist RB, Martinez-Poyer J. In utero analysis of fetal growth: a sonographic weight standard. Radiology 1991;181:129–33.
315 Nwabuobi C, Odibo L, Camisasca- Lopina H, Leavitt K, Tuuli M, Odibo AO. Comparing INTERGROWTH-21st Century and Hadlock growth standards to predict small for gestational age and short-term neonatal outcomes. J Matern Fetal Neonatal Med 2020;33: 1906–12.

316 Duncan JR, Odibo L, Hoover EA, Odibo AO. Prediction of large-for-gestational-age neonates by different growth standards. J Ultrasound Med 2021;40:963–70.
317 Blue NR, Beddow ME, Savabi M, Katukuri VR, Chao CR. Comparing the Hadlock fetal growth standard to the Eunice Kennedy Shriver National Institute of Child Health and Human Development racial/ethnic standard for the prediction of neonatal morbidity and small for gestational age. Am J Obstet Gynecol 2018;219:474.e1–12.
318 Kiserud T, Piaggio G, Carroli G, et al. The World Health Organization fetal growth charts: A multinational longitudinal study of ultrasound biometric measurements and estimated fetal weight. PLoS Med 2017;14:e1002220.
319 Buck Louis GM, Grewal J, Albert PS, et al. Racial/ethnic standards for fetal growth: the NICHD Fetal Growth Studies. Am J Obstet Gynecol 2015;213:449.e1–41.
320 Papageorghiou AT, Ohuma EO, Altman DG, et al. International standards for fetal growth based on serial ultrasound measurements: the Fetal Growth Longitudinal Study of the INTERGROWTH-21st Project. Lancet 2014;384:869–79.
321 Anderson NH, Sadler LC, McKinlay CJD, McCowan LME. INTERGROWTH-21st vs customized birthweight standards for identification of perinatal mortality and morbidity. Am J Obstet Gynecol 2016;214:509.e1–7
322 Fay E, Hugh O, Francis A, Katz R, Sitcov K, Souter V, Gardosi J. Customized GROW vs INTERGROWTH-21st birthweight standards to identify small for gestational age associated perinatal outcomes at term. Am J Obstet Gynecol MFM. 2022 Mar;4(2):100545. doi: 10.1016/j.ajogmf.2021.100545. Epub 2021 Dec 4. PMID: 34875415
323 Odibo AO, Nwabuobi C, Odibo L, Leavitt K, Obican S, Tuuli MG. Customized fetal growth standard compared with the INTERGROWTH- 21st century standard at predicting small-for gestational- age neonates. Acta Obstet Gynecol Scand 2018;97:1381–7.
324 Cheng Y, Leung TY, Lao T, Chan YM, Sahota DS. Impact of replacing Chinese ethnicity-specific fetal biometry charts with the INTERGROWTH-21(st) standard. BJOG 2016;123(Suppl3):48–55
325 Hua X, Shen M, Reddy UM, et al. Comparison of the INTERGROWTH-21st, National Institute of Child Health and Human Development, and WHO fetal growth standards. Int J Gynaecol Obstet 2018;143:156–63.
326 Kabiri D, Romero R, Gudicha DW, et al. Prediction of adverse perinatal outcome by fetal biometry: comparison of customized and population-based standards. Ultrasound Obstet Gynecol 2020;55:177–88.
327 Grantz KL. Fetal growth curves: is there a universal reference? Obstet Gynecol Clin North Am 2021;48:281–96.
328 Michaelson JS, Chen LL, Bush D, Fong A, Smith B, Younger J. Improved web-based calculators for predicting breast carcinoma outcomes. Breast Cancer Res Treat. 2011 Aug;128(3):827-35
